# The transcription factor ATML1 maintains giant cell identity by inducing synthesis of its own long-chain fatty acid-containing ligands

**DOI:** 10.1101/2024.03.14.584694

**Authors:** Batthula Vijaya Lakshmi Vadde, Nicholas J. Russell, Saket Rahul Bagde, Bryce Askey, Michael Saint-Antoine, Bryce Brownfield, Salaiha Mughal, Lauren E. Apprill, Aashima Khosla, Frances K. Clark, Erich M. Schwarz, Saleh Alseekh, Alisdair R. Fernie, Abhyudai Singh, Kathrin Schrick, J. Christopher Fromme, Aleksandra Skirycz, Pau Formosa-Jordan, Adrienne H. K. Roeder

## Abstract

During development, cells not only adopt specialized identities but also maintain those identities. Endoreduplication is thought to maintain cell identity. High concentrations of ARABIDOPSIS THALIANA MERISTEM LAYER1 (ATML1) specify giant cell identity and induce endoreduplication in sepals. How different concentrations of ATML1 can specify different identities remains unclear. Here, we show that high concentrations of ATML1 induce the biosynthesis of both long-chain and very long-chain fatty acids (LCFAs/VLCFAs), and these fatty acids are required for the maintenance of giant cell identity. Inhibition of VLCFA biosynthesis causes endoreduplicated giant cells to resume division and lose their identity, indicating that endoreduplication is not sufficient to maintain cell identity. Structural predictions suggest that LCFA-containing lipids bind to the START domain 2 of ATML1, causing ATML1 dimerization and its auto-activation. Our data and modeling imply that ATML1 induces biosynthesis of its own lipid ligands in a positive feedback loop, shedding light on the intricate network dynamics that specify and maintain giant cell identity.

**Teaser:** Endoreduplicated cells in *Arabidopsis thaliana* sepals divide and de-differentiate in the absence of VLCFA biosynthesis.

## Introduction

Maintaining differentiated cell status is important for the growth and physiology of the plant and animal cells, tissues, organs, and organisms. Perturbations in the maintenance of differentiated cell identity result in diseases, including cancers in animals (*1*). The mechanisms or molecular pathways for forming differentiated cells from stem cells have been extensively investigated in both plants and animals (*2–4*), whereas the mechanisms for maintaining the differentiated status of cells are less understood. Specifically, the maintenance of epidermal cell fate is crucial for the survival of the organism because the epidermis acts as a protective barrier between the internal cells and the harsh external environment. Although little is known about the maintenance of epidermal identity in plants, the specification of epidermal cell identity in *Arabidopsis thaliana* (henceforth Arabidopsis) requires two redundant transcription factors: ARABIDOPSIS THALIANA MERISTEM LAYER1 (ATML1) and PROTODERMAL FACTOR2 (PDF2) (*5–7*). Strong *atml1 pdf2* double mutants arrest as embryos without the specification of an epidermis; likewise, seedlings of *atml1 pdf2* double mutants with weaker alleles lack an epidermis, have exposed mesophyll cells, and are seedling lethal, emphasizing the importance of an epidermis in survival (*7*, *8*).

In addition to specifying the epidermis, the ATML1 transcription factor has a secondary role in specifying giant cell identity within the Arabidopsis sepal epidermis. The Arabidopsis sepal epidermis is an accessible system to study development at the cellular and tissue scale (*9*). Sepals are the outermost organs of the flower that protect the inner reproductive organs. The outer sepal epidermis consists of pavement cells, stomata (guard cells surrounding pores for gas exchange), and trichomes (hairs). Pavement cells can be further classified into small and giant cells on the basis of their size. Giant cells in sepals are large, highly endoreduplicated (16C to 32C), elongated cells, and the size of a giant cell is proportional to its endoreduplication level (*10*, *11*). Endoreduplication is associated with terminal differentiation, and has been thought to maintain giant cell identity and size. Studies to identify the genes involved in giant cell development identified ATML1 as a key protein (*12*). Giant cells are nearly absent from *atml1* mutant sepals, whereas giant cells nearly cover the entire sepal in *ATML1* over expression (*12*, *13*). Although *atml1* single mutants are specifically affected in giant cell development, ATML1 is expressed in both small and giant cells, consistent with its overarching role in specifying epidermal identity.

Meticulous tracking of ATML1 protein concentrations in sepal epidermal cells throughout early sepal development revealed that nuclear ATML1 protein levels fluctuate over time (*13*). On the basis of these observations, we previously developed a model wherein giant cells are specified when the ATML1 concentration surpasses a soft threshold specifically during the G2 stage of the cell cycle. Once the ATML1 concentration crosses this threshold, the cell differentiates by entering into a specialized endoreduplication cycle and elongates to form giant cells. Cells in which the concentration of ATML1 does not surpass this soft threshold during the G2 phase divide and remain as small cells. The concept of thresholds in developmental biology is common. Recently, it was shown that Woolly (Wo) governs digitate and peltate trichome differentiation in a dose dependent manner in tomato (*14*). Wo encodes a homolog of ATML1 and PDF2. However, how ATML1 can specify different cell fates at different concentrations remains elusive.

ATML1 belongs to the class IV homeodomain leucine-zipper (HD-ZIP IV) transcription factor family (*5*). The HD-ZIP family is grouped into four classes (I – IV) (*15*). Classes III and IV are characterized by the presence of a homeodomain (HD), a leucine zipper (ZIP), a steroidogenic acute regulatory protein (StAR)-related lipid transfer (START) domain, and a START-adjacent domain (SAD) (*16*). Class III HD-ZIP proteins contain an additional MEKHLA domain (*17*). In Arabidopsis, the HD-ZIP IV class is the largest group and contains 16 proteins with HD and START domains (*6*, *18*). The ZIP domain in HD-ZIP IV proteins is unique compared with that of other HD-ZIP members because it contains a loop called zipper loop zipper (ZLZ) (*16*). Many HD-ZIP IV transcription factors are involved in specifying the identity of different epidermal cell types (*6*, *12*, *19*, *20*). What remains unknown is whether these transcription factors also have a role in maintaining cell identity after cell fate is specified.

Although HD-ZIP START-containing proteins were identified in the mid-1990s in Arabidopsis, the role of the START domain in these proteins remains elusive (*20*, *21*). The related mammalian StAR/STARD1 protein was first identified in the mouse cell line MA-10 and transports cholesterol into the mitochondria for steroid synthesis (*22*). The structure of the START domain was initially determined for the MLN64/STARD3 protein, which is a close homolog to StAR and is involved in steroidogenesis (*23*, *24*). START domains were first characterized in mammals for binding lipids such as cholesterol and phosphatidylcholine (PC) (*22*, *25*). Clinically, mutations in the StAR protein result in congenital adrenal hyperplasia (CAH) (*26*). Plants are unique in possessing transcription factors containing a START domain. Several studies have implicated the START domain in the dimerization of HD-ZIP proteins (*18*). Although the dimerization is thought to occur via the ZLZ/ZIP domain, deletion of the START domain blocks dimerization (*27–29*). Phospholipids and sphingolipids, such as PC, and ceramide (Cer), are among the primary ligands for the START domain that are thought to be common to plant and animal systems (*25*, *29–31*). Recent studies in plants showed that the START domain of ATML1 can bind to ceramides and restrict its expression to the epidermis (*32*, *33*). The START domains in PDF2 and PHABULOSA (PHB, a class III HD-ZIP) bind to the phospholipids lysophosphatidylcholine (LysoPC) and PC, respectively, to regulate target gene transcription (*29*, *30*). However, how the START domain, and in particular its ligand binding, regulate the activity of the transcription factor remains unclear.

Very long-chain fatty acids (VLCFAs) are defined as fatty acids that contain more than 20 carbon atoms (*34*), and these are incorporated into major plant lipids including ceramides and other sphingolipids, waxes, and phospholipids. Fatty acids (FAs) that contain carbon chains from 11–20 are called long-chain fatty acids (LCFAs) which can be further extended into VLCFAs in endoplasmic reticulum (ER) (*34*). In plants, VLCFA synthesis occurs in ER by the sequential action of four different enzymes: (1) 3-keto-acyl-CoA synthase (KCS), (2) 3-keto-acyl-CoA reductase (KCR), (3) 3-hydroxy-acyl-CoA dehydratase (HCD), and (4) trans-2, 3-enoyl-CoA reductase (ECR). These four enzymes sequentially elongate the fatty acid chain by two carbons. Multiple rounds of KCS, KCR, HCD and ECR activity result in carbon chain lengths of C20 to C38 (*28*, *29*). Among these enzymes, KCSs are rate-limiting (*37*). The Arabidopsis genome encodes 21 KCS genes (*36*), whereas the other enzymes in the pathway are encoded by either one or two genes (*38–41*). Loss-of-function of one of the 21 KCS-encoding genes, *CER2*, results in loss of waxes greater than C28, suggesting that CER2 is required for the synthesis of VLCFAs greater than C28 (*42*). *cer2* mutants have glossy stems compared with wild type (WT) (*42*). Complete loss of *KCR1,* the single KCR-encoding gene in Arabidopsis, is embryo lethal; however, knockdown of *KCR1* using RNA interference (RNAi) lines produces pleiotropic effects, including fused leaves, sensitivity to dehydration, reduced VLCFA content, and retarded growth (*38*). Loss of function of *CER10,* the single Arabidopsis ECR encoding gene, results in shorter plants with reduced cell expansion and reduced VLCFA content (*41*). To what extent these pleiotropic phenotypes relate to the role of VLCFA-containing lipids as ligands for HD-ZIP-START transcription factors versus the myriad of cellular roles of VLCFAs remains unknown.

In this study, we show that high concentrations of ATML1 activate VLCFA and LCFA synthesis. We predict VLCFA/LCFA containing ligands bind to the newly identified ATML1 START2 domain to promote dimerization and activation of ATML1 in a feedback loop. We have further shown that this feedback loop leads not only to the specification of giant cell identity, but also to its maintenance. Our work suggests that even terminally differentiated cell identity must be maintained.

## Results

### High concentrations of ATML1 activate the expression of very long-chain fatty acid (VLCFA) biosynthesis and metabolism genes

We have previously shown that high concentrations of ATML1 specify giant cell identity whereas low concentrations do not (*13*); therefore, we asked whether high concentrations of ATML1 induce different downstream genes than low ATML1 concentrations. To test this, we induced *ATML1* to different levels in Arabidopsis inflorescence tissue and performed RNA-seq. Specifically, we treated the inducible *ATML1* line, *RPS5A*>>*ATML1* (*43*), with 0.1 µM, 1 µM, and 10 µM of the inducer estradiol, and harvested tissue at 8, 16, 24, and 32 h after induction (Fig. 1A). We confirmed by RT-qPCR that *ATML1* expression in the harvested tissues was induced to different levels before sending the samples for sequencing. RT-qPCR results showed that increasing the estradiol concentration increased the *ATML1* transcript level at all time points, except in the 16 h 10 µM sample, which did not induce well and was excluded from further analysis (Fig. 1B).

**Fig. 1.**
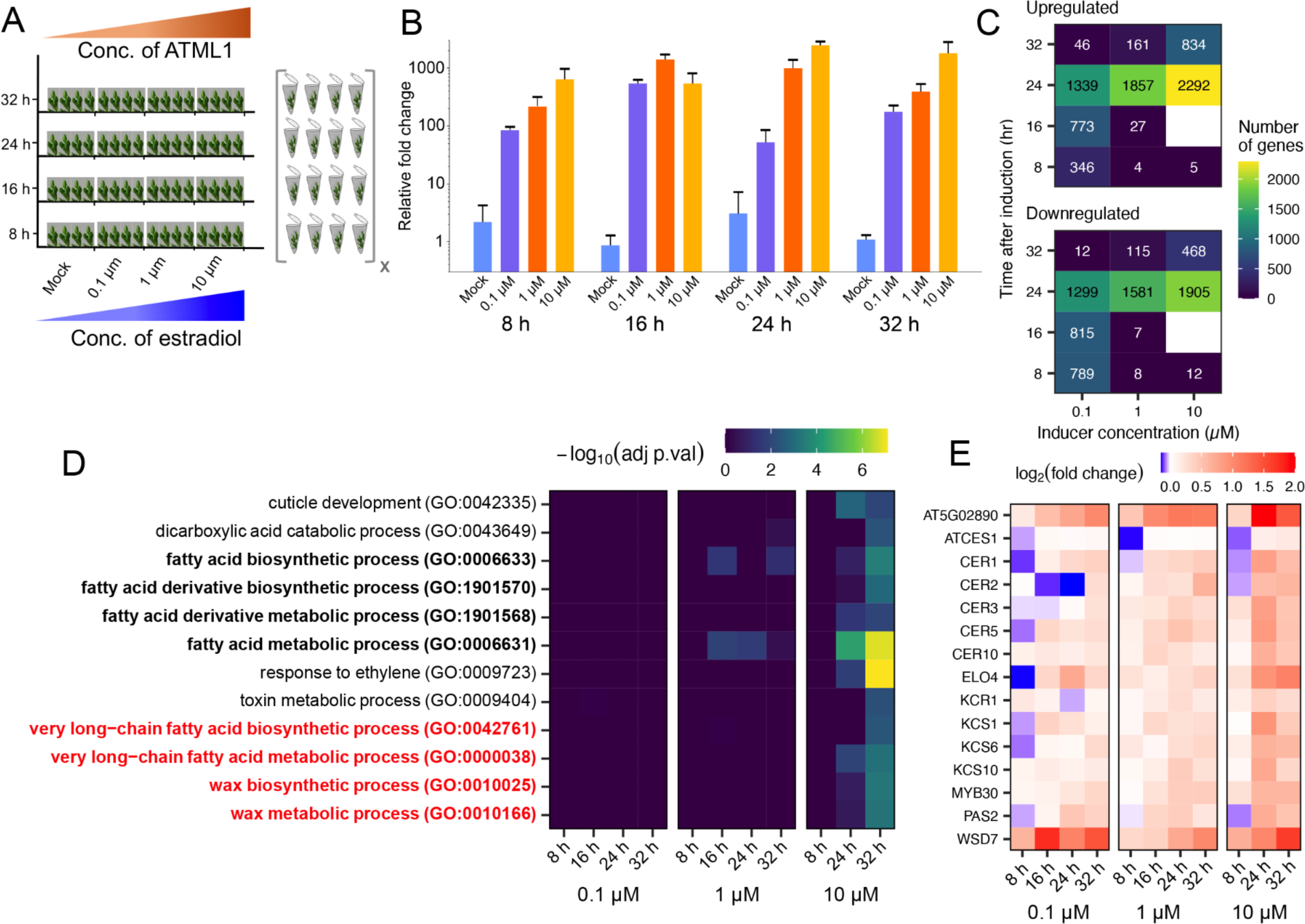
ATML1 activates genes involved in the VLCFA pathway. (**A**) Schematic of ATML1 induction with estradiol concentrations of 0 (mock), 0.1 µM, 1 µM, and 10 µM (x-direction), and sampling times (y-direction) at 8 h, 16 h, 24 h, and 32 h of induction. Tissues at each concentration and time were collected in biological triplicates. Gradient colors of orange and blue indicate ATML1 and estradiol levels respectively. (**B**) Validation of *ATML1* induction using RT-qPCR from the samples collected for RNA-seq (*n* = 3 biological replicates, same as for RNA-seq). Note that the 16 h 10 µM sample did not induce well and was excluded from further analysis. Errorbars indicate 1 SD. (**C**) The number of significantly (adjusted *P* < 0.05) up- and down-regulated genes in each inducible RNA-seq sample. Heat map blue to yellow indicates the number of genes significantly regulated by ATML1. (**D**) Significance of biological process GO terms that were highly enriched (adjusted *P* < 0.05, fold enrichment > 4) in the upregulated genes of the 32 h 10 µM sample. VLCFA biosynthesis, wax biosynthesis, and metabolism are highlighted in red, and all fatty acid-related terms are in bold. (**E**) Heat map of log_2_ (fold change) of gene expression associated with the bolded and highlighted VLCFA and wax GO terms that were upregulated in the 32 h 10 µM sample.

The highest level of *ATML1* induction occurred at the 24 h time point, and RNA-seq data also demonstrated that the greatest number of genes were differentially expressed at the 24 h time point across all inducer concentrations (Fig. 1C). To understand the pathways that ATML1 activates at higher concentrations, we performed gene ontology (GO) analysis on the up- and downregulated genes of each sample to identify overrepresented terms (Data S1). Although ATML1 expression and the number of differentially expressed genes decreased at 32 h, we found that many intriguing biological process terms were enriched in the upregulated genes of the 10 µM 32 h sample (Fig. 1D). These terms included those associated with cuticle development, fatty acid biosynthetic and metabolic processes, VLCFA biosynthetic and metabolic processes, and wax biosynthetic and metabolic processes. Many genes associated with the wax GO terms also participate in VLCFA biosynthetic and metabolic processes, because VLCFAs are core components of waxes (*44*). Genes associated with the VLCFA and wax GO terms were generally strongly upregulated by the high 10 µM concentration at 24 h and 32 h compared with lower 0.1 µM and 1 µM concentrations (Fig. 1E). Additionally, most of these genes were most highly expressed at 24 h but remained significantly upregulated at 32 h. These observations suggest that high concentrations of ATML1 induce the expression of these genes, which then remain upregulated. Thus, our expression analysis highlights VLCFAs as candidate factors involved in the specification of giant cell identity.

We next asked how the differentially expressed genes responded to an increase in ATML1 concentration and if they were induced/repressed only when ATML1 reached a threshold concentration. To this end, we performed gene correlation analysis to identify genes that were co-expressed with *ATML1* (Fig. S1A). We calculated the Spearman correlation and identified 141 genes that were significantly correlated with *ATML1* (Bonferroni-corrected *p*-values less than or equal to 0.05; Fig. S1A; Data S2). Ten of the VLCFA and wax-associated genes were included in this list of highly correlated genes. The expression of these genes ranged from a graded response (Hill coefficient ∼1) to a switch-like response (Hill coefficients ∼5 to 20) to ATML1 concentration (Data S3). The VLCFA genes *CER1*, *PAS2*, and *FDH* showed a graded response to ATML1, whereas *CER3* and *CER5* responded in a switch-like manner to the ATML1 concentration (Fig. S1B and Data S3). These mixed modes of response to ATML1 concentration suggest that although concentration dependance is important, a strictly switch-like response to a threshold concentration of ATML1 is not necessary for giant cell specification. These results indicate that ATML1 activates VLCFA biosynthetic process genes in a concentration-dependent manner, suggesting that VLCFA might contribute to the specification of giant cell identity.

### ATML1 induces (V)LCFA and (V)LCFA-containing lipid production

To identify the lipids that change with increasing concentrations of ATML1, we again induced ATML1 for 24 and 32 h with mock, 0.1 µM, 1 µM, or 10 µM estradiol, and analyzed lipid composition by mass spectrometry (Data S4). Induction of ATML1 increased the concentrations of several families of lipids with 18–26 chain lengths (LCFAs and VLCFAs), ceramides, glycolipids including glucosylceramides, and LysoPCs at 24 and/or 32 h (Fig. 2A-C). Other lipids such as phosphatidylserine (PS), monogalactosyldiacylglycerol (MGDG), and digalactosyldiacylglycerol (DGDG) were significantly downregulated at one or both time points (Fig. S2A–C). These results confirm that ATML1 induces the biogenesis of free LCFAs/VLCFAs (henceforth (V)LCFA) and specific families of (V)LCFA-containing lipids.

**Fig. 2.**
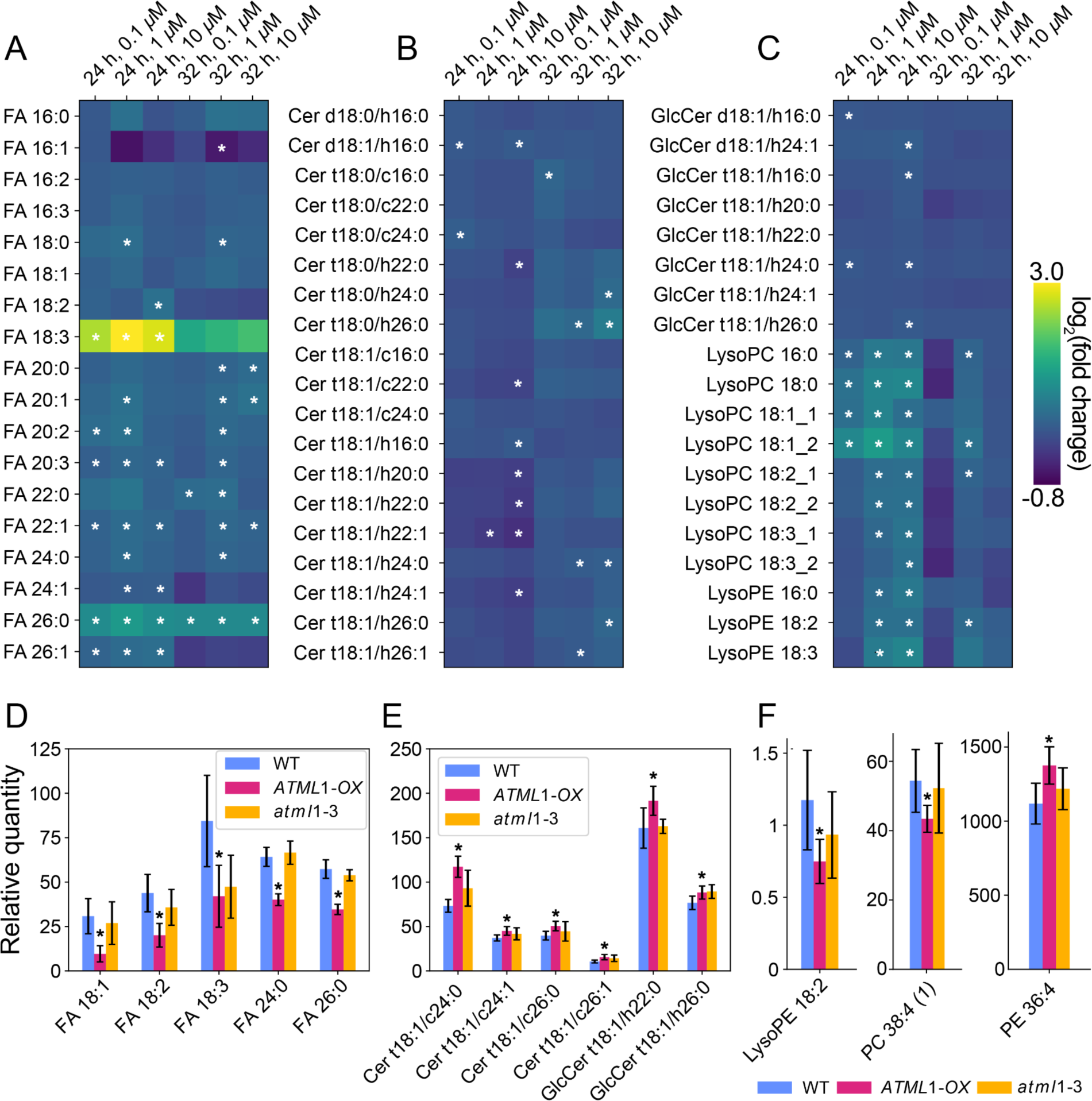
ATML1 induces the synthesis of FA and LCFA-containing lipids. (**A–C**) Induction of ATML1 using estradiol (0.1 µM, 1 µM, 10 µM) for 24 h and 32 h and measurement of (**A**) free fatty acids including LCFAs (FA), (**B**) ceramides, and (**C**) lysophosphatidylcholines, lysophosphatidylethanolamines (LysoPE), and glycosylceramides (GlcCer) using mass spectrometry. Bold and bordered values are statistically significant (Unpaired student’s *t*-test, adjusted *P* < 0.05 with *n* = 5 biological replicates). **(D–F**) Steady-state levels of (**D**) FA including LCFAs, (**E**) ceramides (Cer), (**F**) phosphatidylcholines (PC), phosphatidylethanolamines (PE), and LysoPE in wild type (Col-0), *ATML1-OX* and *atml1-3*. Biological replicates: *n* = 6 for Col-0 and *atml1-3*, whereas *n* = 5 for *ATML1-OX*. * indicates statistically significant values (*P <* 0.05) from wild type and significance was determined using an unpaired student’s *t*-test.

To gain further insight into the relationship between ATML1 and lipid content, we compared the steady-state levels of lipids in WT, *atml1-3*, and *ATML1-OX* (overexpression of *ATML1* in the epidermis; *PDF1::Flag-ATML1*; Data S5). Similar to the results of the induction experiment, specific ceramides and glucosylceramides were more abundant in *ATML1-OX* than in WT, although their concentration was unchanged in *atml1-3* (Fig. 2E). By contrast, the concentration of free fatty acids (FAs) as well as of the phospholipids LysoPE 18:2 and PC 38:4 was significantly reduced in *ATML1-OX* (Fig. 2D, F), whereas the amount of MGDG and DGDG were significantly increased (Fig. S2D, S2F). We suspect this opposite behavior between the inducible system and stable ATML1 overexpression might be due to feedback loops. However, these results again suggest that ceramides, which are sphingolipids that incorporate (V)LCFAs, are associated with high levels of ATML1, which is notable because ceramides were previously shown to bind to ATML1 *in vitro* (*32*).

### Mutation of VLCFA biosynthesis genes decreases giant cell development

To test the role of VLCFA biosynthesis genes in giant cell development, we examined the phenotype of available mutants for these genes. Among the mutants that we screened, the largest giant cells were absent in *cer2-1* (one of the 21 KCSs) and only medium- and smaller-sized giant cells remained (Fig. 3A–E, Fig. S3). The *cer2-1* mutant phenotype showed a smaller reduction in the size of giant cells than the *atml1* mutant (Fig. 3C–E). As mentioned previously, Arabidopsis has only one *KCR* gene and complete loss of *KCR1* is embryo lethal; hence, we examined the *KCR1RNAi* line (*38*). Only the smallest of the giant cells remained in *KCR1RNAi* sepals, a phenotype comparable to the *atml1* mutant (Fig. 3B–C). Thus, VLCFAs are required for the normal development of giant cells in the sepal epidermis.

**Fig. 3.**
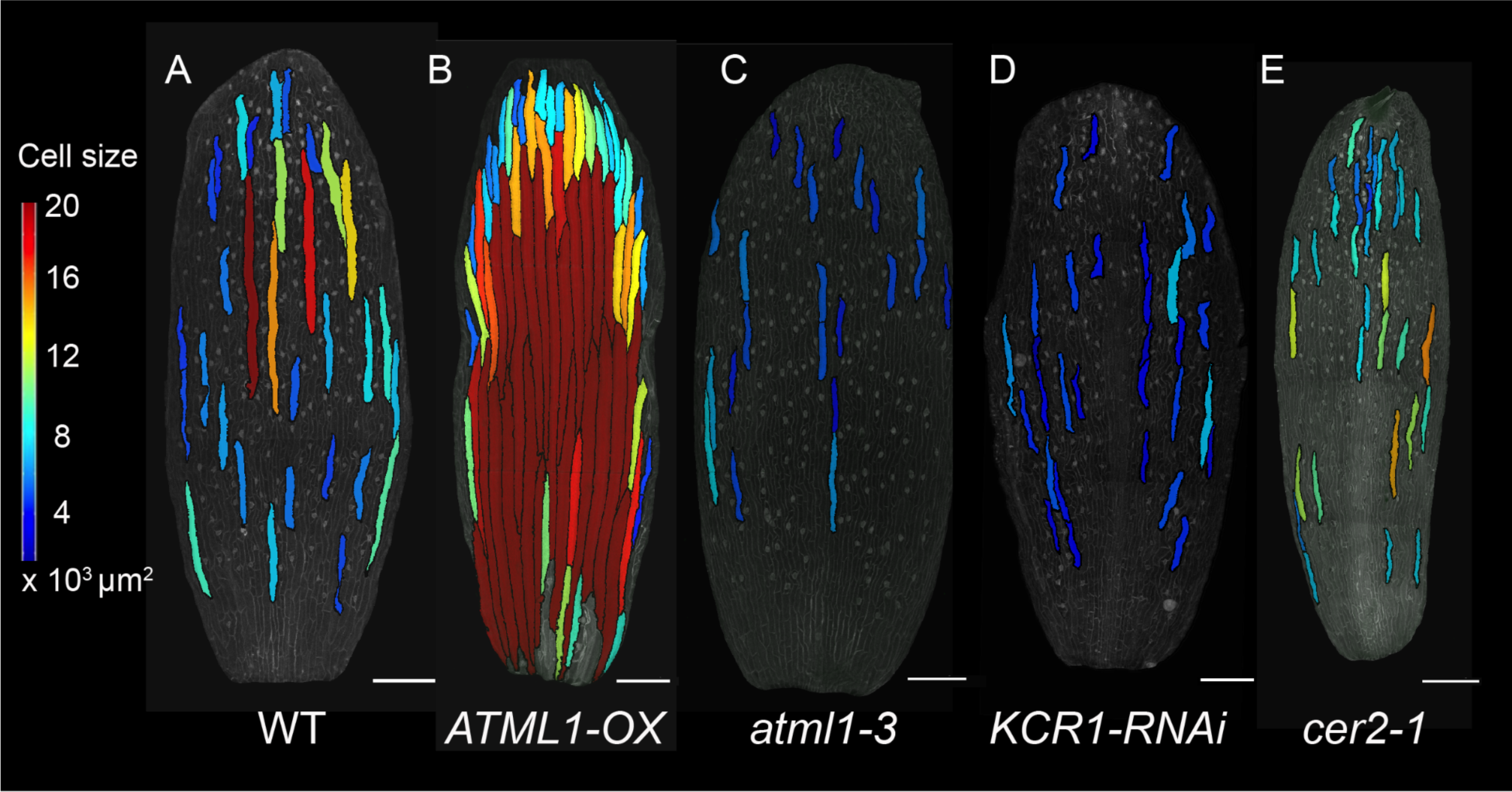
Giant cell development is reduced in VLCFA synthesis mutants. Giant cell segmentation using MorphoGraphX in (**A**) wild type (Col-0), (**B**) *ATML1-OX*, (**C**) *atml1-3*, (**D**) *KCR1RNAi*, and (**E**) *cer2-1*. Heatmap scale bar indicates cell size from blue (medium) to red (extremely large). Images are representative of three biological replicates. Scale bars = 200 µm.

### ATML1-mediated VLCFA synthesis is required to maintain giant cell differentiation

To further test the role of VLCFA synthesis in giant cell development and to probe its relationship with ATML1, we used two well-established chemical inhibitors of VLCFA synthesis: cafenstrole and fentrazamide (*45*). We cultured *ATML1-OX* inflorescences with mock, 30 nM, or 300 nM cafenstrole for 7 days. With 300 nM cafenstrole treatment, the number of giant cells on the sepal epidermis was reduced in all 4 biological samples tested compared with mock and 30 nM cafensterole treatments (Fig. 4A). Similarly, fentrazamide treatment also reduced the number of giant cells in all 3 biological replicates of *ATML1-OX* sepals (Fig. S4A), indicating that the giant cell phenotype is specific to the inhibition of VLCFA synthesis but not to the chemical inhibitor used. We conclude that VLCFA synthesis is required for ATML1 to produce ectopic giant cells in sepals. We further tested whether inhibition of VLCFA synthesis could block the development of giant cells in WT. Treatment of WT with 300 nM cafenstrole reduced sepal giant cell number in 2 out of 4 biological replicates (Fig. S4B). Notably, ATML1 levels did not decrease significantly after cafenstrole application in WT (Fig. S4C–D). These experiments suggest that ATML1 requires VLCFA synthesis to induce giant cell development.

**Fig. 4.**
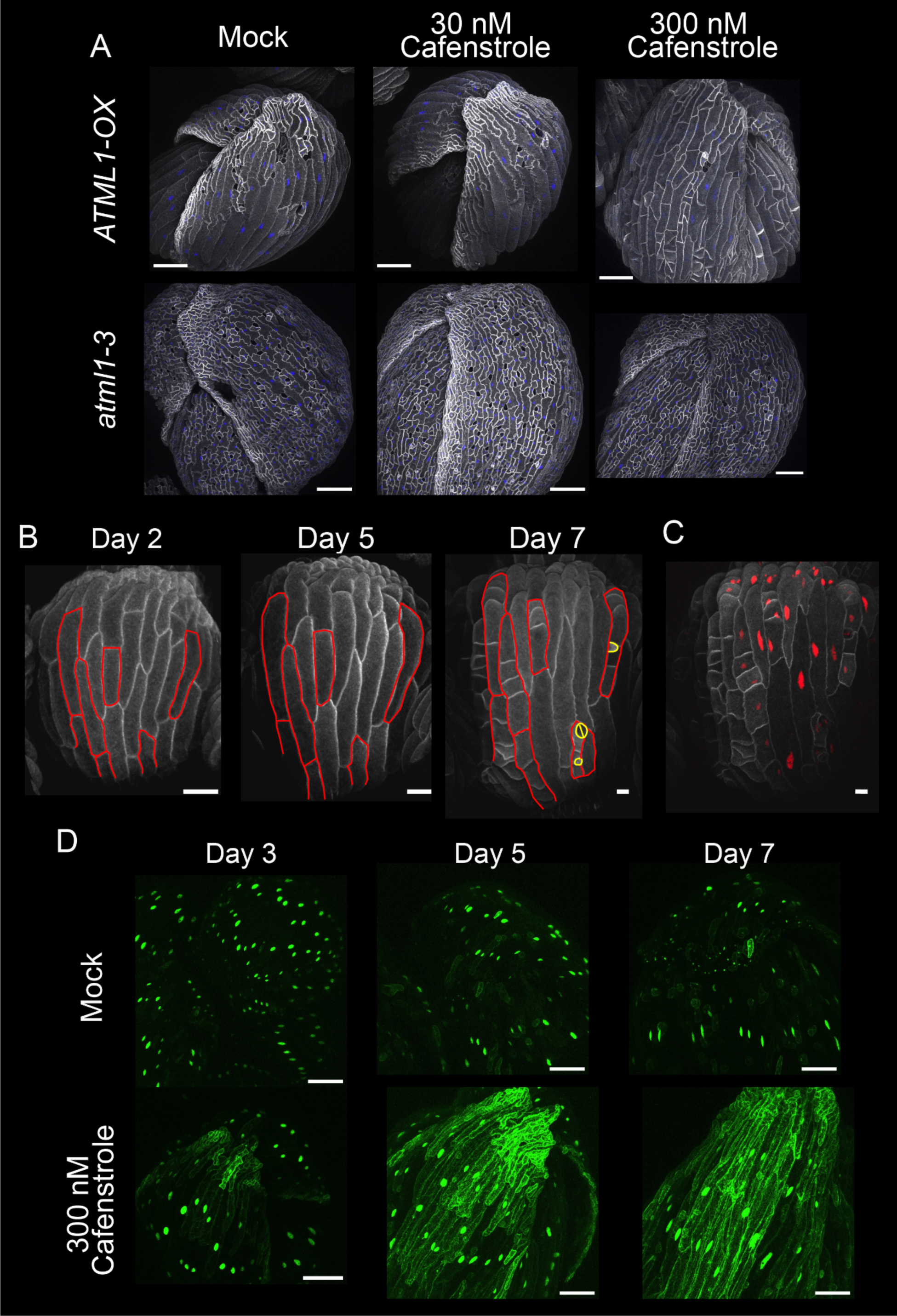
VLCFA synthesis is required for giant cell formation and maintenance. (**A**) Pharmacological inhibition of VLCFA synthesis using cafenstrole in *ATML1-OX* and *atml1-3* inflorescence tissue. Cell outlines are visualized by the presence of *35S::mCitrine-RCI2A;* nuclei are visualized in blue with *pUBQ10::H2B-mTFP*. Images are representative of three biological replicates. **(B**) Live imaging of *ATML1-OX* plants with *35S::mCitrine-RCI2A* treated with 300 nM cafenstrole for 7 days. On day 7, many differentiated giant cells divided, which could lead to stomata formation. For clarity, the giant cells that divided on day 7 are outlined in red color manually on days 2, 5, and 7. Stomata and meristemoid mother cells are highlighted in yellow. Scale bars = 50 µm. (**C**) Day 7 *ATML1-OX* plant from (**B**), imaged with *35S::mCherry-H2B* (red) localized to the nucleus. Cell outlines were visualized by the presence of *35S::mCitrine-RCI2A*. Scale bar = 50 µm. (**D**) Live imaging of giant (nuclei shown in green) and small cell (ER shown in green) markers expression in *ATML1-OX* flowers from day 3 to day 7 at intervals of 48 h. Scale bars = 100 µm.

We next tested whether VLCFA synthesis was required for maintenance of giant cell identity by live imaging *ATML1-OX* treated with 300 nM cafenstrole for 7 days. Until day 5, giant cells developed normally, including elongation and endoreduplication. Between days 5 to 7, many of the giant cells underwent multiple rounds of division, which subdivided the giant cell into several small cells. The VLCFA synthesis inhibitor experiments showed that giant cells initially formed, but then subsequently divided into several small cells, suggesting that the maintenance of giant cell identity was disrupted. In some cases, the smaller cells resulting from division of the giant cell entered the stomatal patterning pathway and differentiated as guard cells, indicating that the giant cell had de-differentiated (Fig. 4B). Analysis of a nuclear marker revealed that the highly endoreduplicated giant cell nuclei were larger than the nuclei of the daughter cells after cell division (Fig. 4C), suggesting the daughter cells had a reduced ploidy. Analysis of epidermal ploidy by flow cytometry showed that 300 nM cafenstrole treatment reduced the proportion of highly endoreduplicated 32C cells compared with mock-treated inflorescences (Fig. S4E). These data show that inhibition of VLCFA synthesis induces cell division in giant cells, and sometimes reprograms the fate of the small daughter cell.

We further tested whether, and to what extent, giant cells lose their identity and de-differentiate by examining the expression of molecular markers specific to giant and small cells (*12*, *13*, *46*). Live imaging of *ATML1-OX* inflorescences containing the giant and small cell markers treated with mock and 300 nM cafenstrole, revealed that in 300 nM cafenstrole, cells started to express the small cell marker basipetally from the sepal tip, compared with mock-treated sepals (Fig. 4D). Expression of the small cell marker was initiated in cells before division occurred and concomitant with continued expression of the giant cell marker, suggesting that cells start to lose their identity and progress through a mixed identity state rapidly after VLCFA synthesis is inhibited. Overall, these results show that ATML1-mediated VLCFA synthesis is required for the maintenance of giant cell identity.

### ATML1 has two START domains that are predicted to interact with a LCFA-containing lipid/ceramide, resulting in ATML1 dimerization

Because the genetic and pharmacological experiments suggested that ATML1-mediated VLCFA synthesis is required for giant cell formation and maintenance, we next asked how these small molecules regulate giant cell development. Previous work showed that LCFA-containing lipids, including ceramides, can bind to the START domain of ATML1 and its closely related paralog PDF2 (*30*, *32*, *33*), suggesting the potential existence of a feedback loop. To consider this possibility, we tested whether cafensterole treatment requires ATML1 to promote cell division or alternatively, whether cafensterole treatment promotes cell division generally. Treatment with 30 nM and 300 nM cafenstrole did not affect the *atml1-3* sepal cells compared with those in the mock treatment, suggesting that ATML1 is necessary for the effect of cafenstrole on cellular division (Fig. 4A). Thus, inhibition of VLCFA biosynthesis does not promote cell division in general, but instead acts specifically through ATML1 in the giant cell development pathway.

To gain insights into the mechanism, we analyzed predicted structures of ATML1. Previous studies have shown that the ATML1 protein has a DNA-binding homeodomain (HD), a dimerization zipper-loop-zipper (ZLZ) domain, a START domain and a START-adjacent domain (SAD) (*11*, *41*). We obtained a predicted structural model of Arabidopsis ATML1 from the AlphaFold 2 database (Fig. 5A–B) (*48*, *49*). In this model, the HD forms a structure consisting of three α-helices characteristic of homeodomain proteins (*50*). The third helix of the HD domain extends to form the leucine zipper, which is divided into two zippers due to the presence of a loop (loop I) and forms a ZLZ motif, characteristic of HD-ZIP IV proteins. The HD-ZLZ domains are connected to the START domain via a 46-residue loop (loop II). The model for the N-terminal residues preceding the HD domain and loops II and III have low predicted local distance difference test (pLDDT) scores, suggesting these regions are likely unstructured and flexible. The START domain forms an interface with the SAD domain to give a START/SAD didomain structure (Fig. 5B). Notably, superposition of the predicted model of the SAD domain on the crystal structure of the START domain from human ceramide transfer protein (CERT/STARD11) in complex with C18-ceramide (PDB 2E3Q) (PMID: 18184806) showed that the SAD domain is predicted to adopt a START domain fold (root-mean-square deviation (RMSD), 4.02Å) (Fig. 5C). This domain was not recognized previously as a START domain likely due to low amino acid sequence identity (18% between START1 and SAD/START2 domains of ATML1) and due to a large (16 residue) insertion into loop III of SAD/START2 (Fig. 5C). Similar superposition analysis of AlphaFold 2 predicted models of HD-ZIP III and IV proteins from Arabidopsis revealed that the SAD domain of all class III and class IV HD-ZIP members is predicted to adopt a START domain fold (Table S1). This prediction is supported by circular dichroism performed by another group, which found that the HD-ZIP III SAD and classical START domains adopt virtually identical secondary structures (Aman Husbands, personal communication, November 18, 2023). Hence, we term the first START domain of HD-ZIP III and IV proteins START1 and the SAD domain START2. Together, we refer to these domains as the START1/2 didomain.

**Fig. 5.**
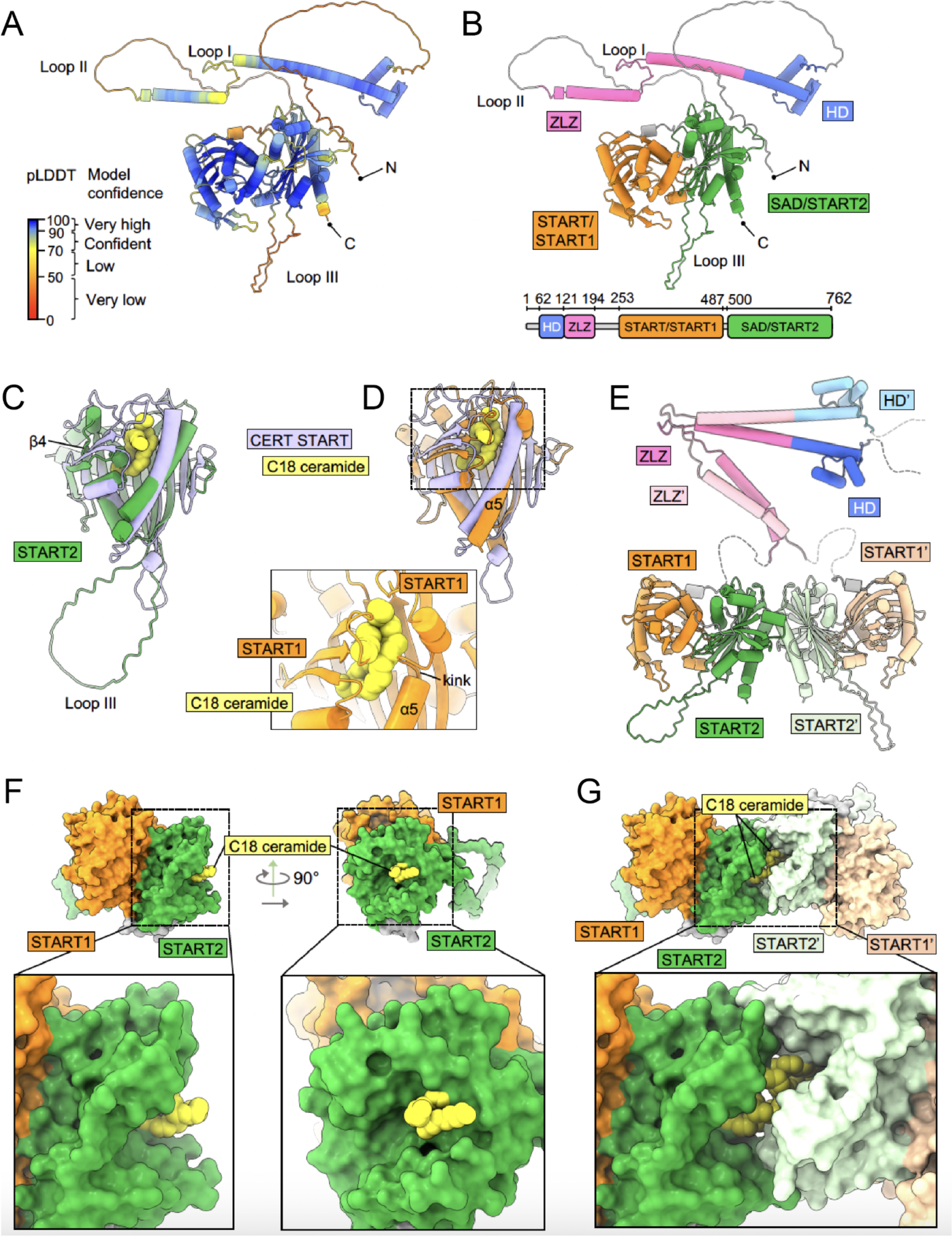
AlphaFold2 predicts ATML1 has two START domains. AlphFold-Multimer predicts ATML1 homodimerizes at the START2 interface, completing the VLCFA-binding pockets. (**A**) AlphaFold2 predicted structure of ATML1 colored by pLDDT score to show model confidence. (**B**) Alphafold2 predicted structure of ATML1 colored by domains: homeodomain (HD), zipper loop zipper (ZLZ), START domain 1 (previously identified), START domain 2 (newly predicted). Schematic showing the linear N-to C-terminal domain arrangement of ATML1. Models of ATML1 (**C**) START1 and (**D**) START2 domains, aligned to the crystal structure of the START domain from Human ceramide transfer protein (CERT) in complex with C18-ceramide (PDB 2E3Q). The magnified view in (**D**) shows a model of C18-ceramide docked on ATML1 START1 based on the crystal structure of CERT in complex with C18-ceramide. A kink in helix 5 (α5) in the predicted ATML1 START1 model constricts the predicted lipid-binding site and introduces steric clash with docked C18-ceramide. ATML1 START2 has a longer Loop III than that in CERT START. (**E**) Model of dimeric ATML1 predicted using AlphaFold-Multimer. Note the predicted homodimerization via the START2 domains in addition to the ZLZ domain. (**F**) Predicted model of ATML1 START1–START2 didomain with C18-ceramide modeled in the START2 domain (shown in surface representation). Note the binding pocket is not complete in the monomer and the ceramide is exposed. (**G**) Dimer model of the ATML1 START1–START2 didomain (in surface representation) with two C18-ceramide molecules modeled in the two START2 domain-binding pockets. Note that dimerization completes both binding pockets and the ceramides are enclosed.

Next, we analyzed the putative lipid-binding pockets of the START1 and START2 domains in the predicted model of ATML1. Comparison of the START1 model with the crystal structure of the START domain from CERT in complex with C18-ceramide revealed that a kink in helix 5 (α5) in the predicted ATML1 START1 model constricts the predicted lipid-binding site and introduces steric clash with the docked lipid (Fig. 5D). On this basis, we predict that the START1 domain is not able to bind a lipid molecule in the predicted conformation. On the other hand, beta strands β4 and β5 in the START2 model are bent away from the putative lipid binding site, leading to an incomplete pocket that may expose the bound lipid to the surrounding solution. In this case, stable binding of lipids to START2 will likely require completion of this pocket. One possibility is that a conformational change may lead to completion of the predicted lipid-binding pocket of START2. Another possibility is that binding of another protein to the START2 domain is needed to complete the pocket.

We then used AlphaFold-Multimer to predict homodimeric structures of ATML1 (*51*). As expected, the ZLZ domain dimerizes via the leucine zipper helices in the predicted dimer model (Fig. 5F). Surprisingly, the START1/2 didomain is also predicted to form a dimer (Figs. 5F, S6A). Importantly, the START2 domains from each copy of ATML1 assemble at the predicted dimer interface such that the lipid binding pocket of each START2 domain is completed by the START2 domain of the other monomer (Fig. 5G). This suggests the possibility that binding of a LCFA-containing lipid, such as ceramide, would be stabilized by homodimerization of the START1/2 didomain, and that this lipid binding to the START2 domain might induce homodimerization of the START1/2 didomain.

In support of these predictions, BiFC experiments using split GFP tagged to ATML1 suggested that ATML1 forms a homodimer (Figs. 6A–B, S7A). We also tested dimer formation using quantitative yeast two-hybrid assays. Full-length ATML1 exhibited homodimerization in the yeast system and we additionally detected that when ATML1 START1/2 was used as a bait, the didomain interacted with the full-length ATML1 (Fig. 6C–D). Yeast two-hybrid assays also corroborated that START1 and START2 form intramolecular interactions (Fig. S7B), consistent with the AlphaFold model. We assume that in both yeast and in the BiFC experiments that sufficient LCFA-containing lipid ligands are available in the system. Finally, co-immunoprecipitation confirms that ATML1 proteins dimerize *in vivo* in Arabidopsis inflorescences (Fig. 6E). Our results support dimerization of ATML1 through the START1/2 didomain in addition to the ZLZ domain.

**Fig. 6.**
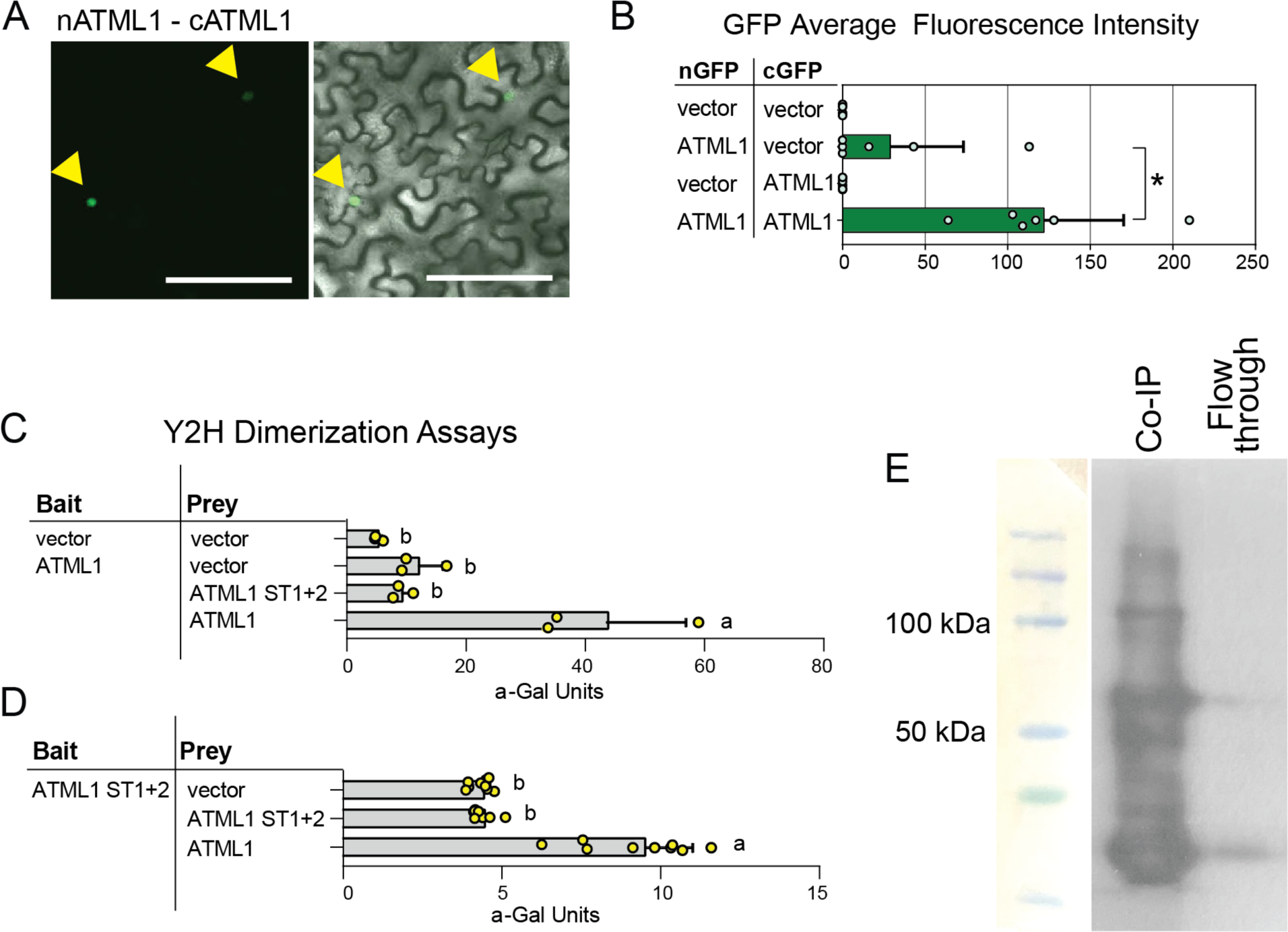
ATML1 homodimerizes. (**A**) Bimolecular fluorescence complementation (BiFC) assays indicate homodimerization of ATML1. *Nicotiana benthamiana* leaf abaxial epidermal cells were transiently transformed with constructs expressing split GFP segments fused to ATML1. Interaction was detected between ATML1 tagged with N-terminal GFP (nGFP:ATML1) and ATML1 tagged with C-terminal GFP (cGFP:ATML1). Nuclear expression of GFP is indicated by arrowheads. Scale bar = 100 µm. See also Fig. S7. (**B**) Quantitative analysis of BiFC images. Mean pixel intensities are from *n* = 6 images taken from two independent transformants in two trials. Error bars indicate standard deviations (SD). *, *P* = 0.0060 by unpaired *t-*test. (**C–D**) Dimerization of ATML1 in yeast two-hybrid (Y2H) assays. Diploid yeast containing the bait and prey constructs were assayed for *MEL1* reporter gene activity in quantitative α-galactosidase assays. Empty bait and prey vectors served as negative controls. (**C**) Homodimerization of ATML1 in the Y2H assay. Weaker reporter gene activity is observed for the combination of ATML1 as a bait and the START1+2 didomain as the prey. Data represent means of *n* = 3 independent yeast transformants. Error bars indicate SD. Significant differences between constructs are marked by letters (one-way ANOVA, Tukey’s test, *p* < 0.0001). (**D**) Dimerization of the ATML1 START1+2 didomain, when expressed as a bait, with full-length ATML1 as the prey. Data represent means of *n* = 3 independent yeast transformants with *n* = 3 technical replicates each. Error bars indicate SD. Significant differences between constructs are marked by letters (one-way ANOVA, Tukey’s test, *P* < 0.0001). (**E**) Co-IP of FLAG-ATML1 from the protein extracts of *pPDF1::FLAG-ATML1×pATML1::mCitrine-ATML1* to check ATML1 dimerization. ATML1 protein was pulled down with FLAG beads and a western with GFP antibody detected mCitrine-ATML1. mCitrine-ATML1 fusion protein is ∼110 KDa (degradation products are also observed).

To explore the potential conservation of the ATML1 structural predictions, we also predicted dimeric models of all HD-ZIP IV family members using AlphaFold-Multimer (*51*). To specifically assess dimerization of the START1/2 didomain, we focused the predictions on the isolated START1/2 didomain (Fig. S6 and Table S1). Among HD-ZIP IV members, only ATML1, PDF2, HDG2 and GLABRA2/ATHB10 (GL2) are predicted to dimerize via their START1/2 didomains (Fig. S6A–D). The START1/2 didomains in PDF2 and HDG2 are predicted to dimerize in the same fashion as described for ATML1 above, involving an interface formed by the START2 domains (Figs. 5F, S6B–C). However, GL2 is predicted to have a different mode of dimerization that involves interfaces formed by both START1 and START2 domains (Fig. 6A, D). This pattern closely follows phylogenetic relationships of HD-ZIP IV proteins, because PDF2 and HDG2 are most closely related to ATML1 among all HD-ZIP IV family members in Arabidopsis (*52*).

The predicted dimeric architecture of HD-ZIP III family proteins is different from that predicted for the HD-ZIP IV family proteins (Figs. S5, S6E–I; Table S1). In the HD-ZIP III predictions, the START1 domains interface with the leucine zipper dimer to form an overall cyclic architecture. Comparison of the AlphaFold2 predicted START1 and START2 models of HD-ZIP III protein REVOLUTA/IFL1 with the crystal structure of the START domain from CERT in complex with C18-ceramide revealed that the predicted lipid binding pocket of REVOLUTA/IFL1 START1 is much smaller than that of the CERT START domain (Fig. S5D–F) (*53*). Furthermore, the START2 domains of HD-ZIP III family proteins have a helical insertion (α5) between helix 4 (α4) and the beta strand (β4). In the REVOLUTA/IFL1 model, helix 5 is predicted to interfere with lipid binding (Fig. S5G, H). Therefore, we expect neither the START1 nor START2 domain of REVOLUTA/IFL1 is able to bind a lipid molecule in the predicted conformation. Conformational changes may enable lipid binding to the pockets of the START domains of REVOLUTA/IFL1 or it is possible that other smaller secondary metabolites may bind the START domains in the predicted conformation.

Although our protein structure model predicts that binding of ceramide to ATML1 may induce or stabilize dimerization, which we hypothesize could enhance the activation of transcription by ATML1, there are other possible roles for this lipid-binding activity. Previously, Nagata et al. showed that VLCFAs promote the stability of ATML1 protein in root meristems and that treatment with cafenstrole destabilized the ATML1 protein (*32*). To test the effect of cafenstrole on ATML1 protein concentration in developing sepals, we treated inflorescences expressing *mcitrine-ATML1* (*pATML1::mCitrine-ATML1;atml1-3*) with mock and 300 nM cafenstrole and imaged them every day for 8 days. In developing sepals, mcitrine-ATML1 fluorescence levels were similar between the mock- and 300 nM cafenstrole-treated samples (Fig. S4C–D), suggesting that ATML1 protein stability was not dramatically affected in sepals. Thus, it is likely cafenstrole treatment has tissue-specific effects. For sepals, our structural models suggest that LCFA/VLCFA-containing lipids binding to ATML1 regulates its activity as a transcription factor.

### Modeling of the ATML1–(V)LCFA regulatory network reproduces the giant cell pattern and identity maintenance *in silico*

We created an analytical and computational model to explore the dynamic behavior of our hypothesized ATML1–(V)LCFA feedback network. We had previously modeled fluctuations in ATML1 protein levels and showed that they were sufficient to generate the pattern of giant cells and small cells in the sepal (*13*). However, that model did not include (V)LCFAs and (V)LCFA-containing lipids nor maintenance of giant cell identity; therefore, on the basis of our experimental data, we hypothesized an updated molecular regulatory network for giant cell specification and maintenance (Fig. 7A), building on the previous network (see Methods).

**Fig. 7.**
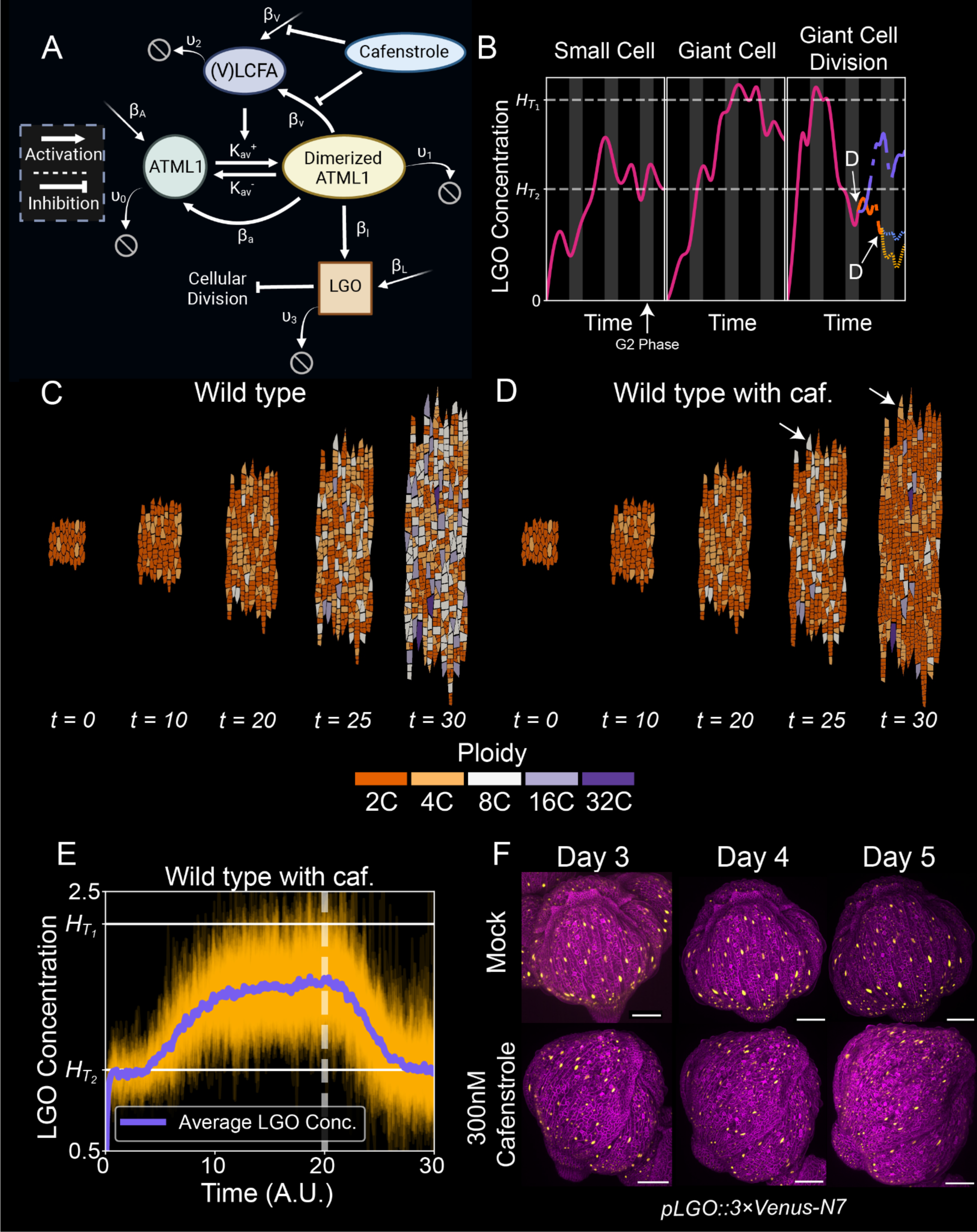
*In silico* simulation of giant cell development recapitulates loss of giant cell maintenance upon inhibition of (V)LCFA biosynthesis. (**A**) Scheme of the gene regulatory pathway derived from the genetic and cell biology observations (see Methods). (**B**) Illustration showing examples of LGO concentration trajectories in a small cell, a giant cell, and a giant cell undergoing division of the proposed model. Black and grey areas represent the G1 and G2 phases, respectively. *H*_T1_ and *H*_T2_ represent the higher and lower LGO thresholds, respectively. The LGO concentration of small cells never reaches the higher threshold, *H*_T1_. The LGO concentration of a giant cell exceeds *H*_T1_ during the G2 phase and never drops below the lower threshold *H*_T2_ In the rightmost panel, because the LGO concentration of an endoreduplicating giant cell eventually goes below the lower threshold *H*_T_*_2_*, the cell divides into two equal ploidy cells (denoted by D). At the start of the next G2 phase, the purple trajectory endoreduplicated again since the concentration is above the threshold *H*_T_*_2_*, but the orange trajectory is below *H*_T_*_2_*, causing it to divide again (see Methods). (**C–D**) Simulation of a wild type (WT) sepal epidermis (**C**) without and (**D**) with cafenstrole. Arrows denote an example of giant cell division (see Movies S1–S2). (**E**) LGO concentration for the WT simulation with cafenstrole that are shown in (**D**). Orange lines represent individual trajectories of cells, and the purple line represents the average LGO concentration in all cells. At *t* = 20, cafenstrole is added. (**F**) Live imaging of *pLGO::3×Venus-N7* (yellow); *35S::mCherry-RCI2A* (magenta) in mock and 300 nM cafenstrole. Scale bars = 100 µm. Quantification is presented in Fig. S10.

In brief, we assumed that (V)LCFA-containing lipids bind to ATML1, enabling ATML1 to dimerize. We further assumed that only ATML1 dimers transcriptionally activate downstream gene expression, including activating the biosynthesis of (V)LCFAs (Fig. 7A). Note that we attempt to understand the dynamics of the (V)LCFA that are available to bind to ATML1, not all of the (V)LCFA in the tissue. We have previously shown that ATML1 operates in a weak positive feedback loop and activates its own expression (*13*), and this is now included in the model as activation of ATML1 expression by the ATML1 dimer. The cyclin-dependent kinase (CDK) inhibitor, LOSS OF GIANT CELLS FROM ORGANS (LGO), acts downstream of ATML1 to induce endoreduplication of giant cells (*10*, *13*). In our model, the ATML1 dimer activates *LGO* expression, and if the *LGO* concentration exceeds a predetermined threshold when the cell is at the G2 phase, the cell will endoreduplicate and become a giant cell (Fig. 7B, threshold *H*_T_*_1_*). We included the effect of the cafenstrole drug into our model. Specifically, cafenstrole impairs the basal production of (V)LCFAs as well as the activation of (V)LCFA synthesis by the ATML1 dimer.

To simulate the observed behavior of giant cells losing their fate and dividing in cafenstrole treated plants, we postulated a second threshold in LGO expression, allowing a giant cell to de-differentiate and resume division. Giant cell fate will only be affected if the cell’s expression dips below a second threshold *H*_T_*_2_*< *H*_T_*_1_*(Fig. 7B). If this occurs at any time during the cell cycle, then the giant cell undergoes mitosis and divides by the end of the next G2 phase. These daughter cells divide yet again in the next cell cycle if, by the time they reach the G2 phase, their LGO levels do not exceed the second threshold *H*_T_*_2_*. If these levels are above the second threshold, the cell resumes endoreduplication (Fig. 7B).

To understand the effects of cafenstrole, we first created a deterministic model of our network for a single cell (without growth, division, or stochastic expression, but incorporating dilution effects to emulate the deterministic dynamics of a growing tissue; see Methods). We plotted the protein and (V)LCFA concentration trajectories before and after the addition of varying levels of cafenstrole in three settings: WT, *ATML1-OX*, and *LGO-OX* (Fig. S8A–C). For both WT and *ATML1-OX*, once cafenstrole is added into the system, the cell’s LGO concentration declines towards the second, lower threshold (*H*_T_*_2_*). This brings the LGO concentration level low enough that giant cell fate could be reversed if stochastic fluctuations cause the LGO concentration to dip below the threshold (Fig. S8A-B). However, cafenstrole does not have a prominent impact on *LGO-OX*, showing that giant cells in *LGO-OX* may be able to overcome the reduction in (V)LCFAs and maintain their giant cell identity (Fig. S8C). Therefore, the deterministic model predicts that cafenstrole should allow giant cell de-differentiation in WT and *ATML1-OX*, but not in *LGO-OX*.

### Cafenstrole application in our model can de-differentate giant cells, similarly to experimental data

Next, we implemented the model in an *in silico* multicellular tissue of growing and dividing cells. We added stochastic fluctuations in the gene regulatory dynamics, growth, and cell division to determine whether this molecular network could reproduce the pattern of giant cells observed in experiments (see Methods section and Table S2). We first constructed a WT simulation, in which we recapitulated an interspersed pattern of giant cells (Fig. 7C, Movie S1) with a similar ploidy distribution to WT and where no giant cells (8C and above) divide (Fig. S4E and Fig. S9). To assess whether we could induce giant cell division, we simulated the addition of cafenstrole at time *t* = 20 after some giant cells had formed (Fig. 7D, Movie S2). At *t* = 25, the sepal appeared largely similar to the simulation of WT sepals without cafenstrole. However, fewer giant cells were present at this stage of the simulation because many cells could not reach the higher *H*_T_*_2_*to induce endoreduplication. By *t* = 30, most of the giant cells divided and several cells that would have become giant remained small (see arrows in Fig. 7D). These sepals have fewer giant cells, similar to the experimental results (Fig. 4B, S4E and S9). Similar to the deterministic model, the addition of cafenstrole lowered the average LGO concentration to the *H*_T_*_2_*threshold and, due to the stochastic fluctuations, some of the giant cells dipped below the threshold *H*_T_*_2_*, inducing divisions (Fig. 7D–E).

Our model predicted that application of cafenstrole reduced LGO concentration before giant cell divisions occurred (Figs. 7E, S8). To validate our simulation data of reduced LGO expression, we grew inflorescences of *pLGO::3×Venus-N7* (LGO transcriptional reporter) *35S::mCherry-RCI2A* (fluorescent membrane marker) doubly transgenic plants on mock and 300 nM cafenstrole media. Tracking this *LGO* reporter showed that *LGO* transcription declined on the fifth day of cafenstrole treatment compared with its mock control (Figs. 7F, S10), preceding the initiation of giant cell division in our live-imaging experiments (Fig. 4B). This delayed reduction of *LGO* from the addition of cafenstrole is captured in our model (Fig. 7E). These data along with the data from (*13*), indicating *LGO* acts downstream of *ATML1*, are consistent with our model that (V)LCFA-mediated ATML1 dimerization is required for the maintenance of *LGO* transcription and giant cell fate maintenance.

### The model captures the LGO dynamics observed in the experiments with ATML1 and LGO mutants

To further test the ATML1–(V)LCFA network model, we predicted the expression pattern of *LGO* in *ATML1-OX* and *atml1-3* mutants and tested these predictions experimentally (Fig. 8A). In order to simulate the *atml1-3* mutant, we assume a very low basal production rate to account for redundancies in the regulatory network such as that due to the *ATML1* paralog *PDF2*. Our simulation data showed that compared with WT, in *ATML1-OX*, high levels of LGO concentration were reached more rapidly (*t ≈* 10 in WT whereas *t ≈* 3 in *ATML1-OX*) (Fig. 8A, B). However, in *atml1-3*, LGO expression was slightly reduced and delayed (*t ≈* 15) compared with WT (Fig. 8A–B). To validate our simulation data, we expressed the *pLGO::3*×*Venus-N7* reporter in *ATML1-OX* and *atml1-3* mutant backgrounds. Confocal imaging showed LGO reporter was strongly expressed in *ATML1-OX* at stage 4, strongly expressed in WT at stage 5, and only very weakly expressed in *atml1-3* mutants at stage 6, similar to the time order predicted by the model (Fig. 8C– E). In the *atml1-3* mutant, *LGO* reporter was not detected at the critical stage 5 of flower development, when *LGO* reporter expression initiates in WT, suggesting that lack of *LGO* expression at this time point prevents giant cell development (Fig. 8D). However, in *atml1-3*, a low level of *LGO* reporter expression started from stage 6 of flower development (Fig. 8E, see arrow). At later stages of sepal development (stage 9), *LGO* reporter expression in *atml1-3* was roughly equivalent to that in WT (Fig. 8F).

**Fig. 8.**
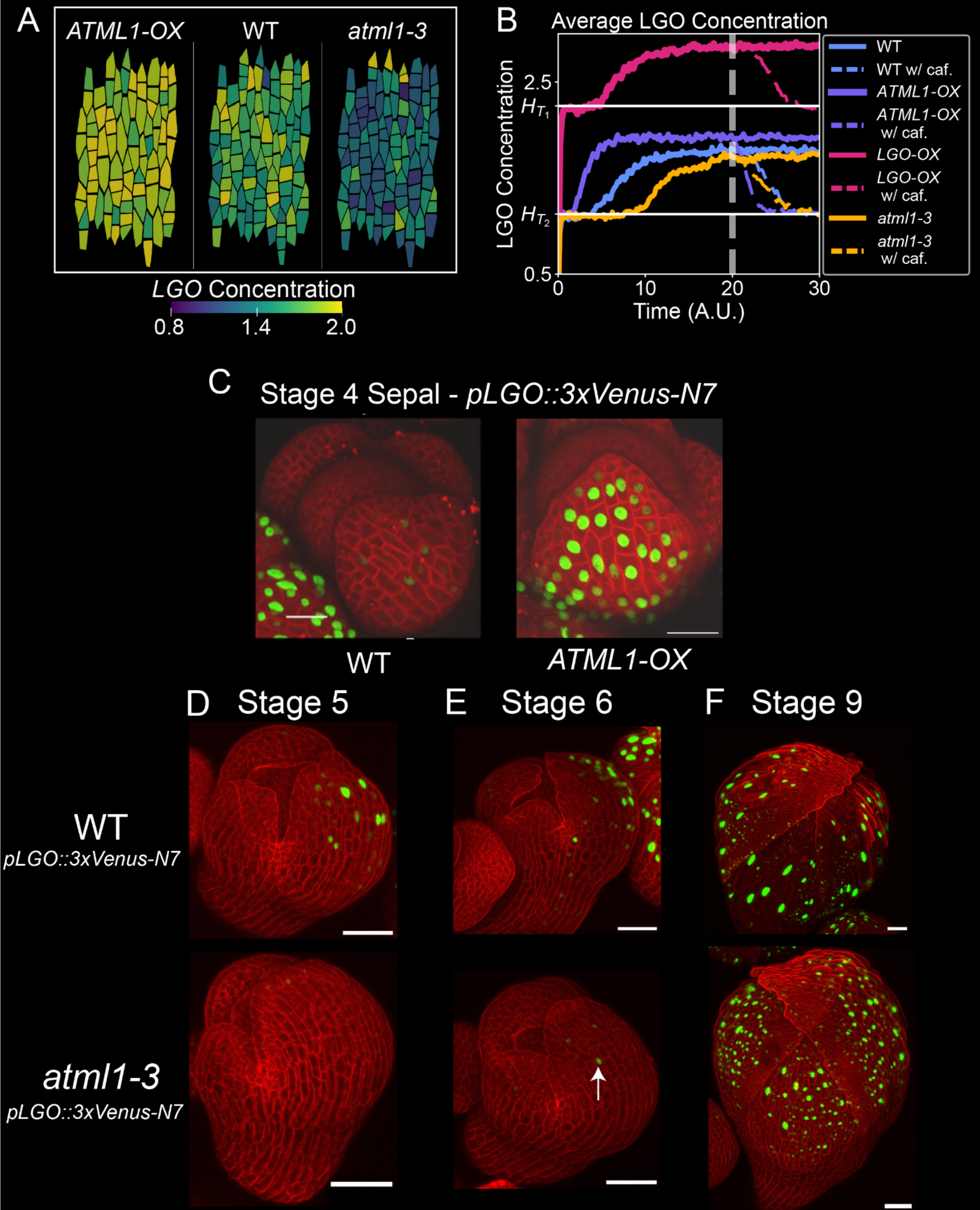
*LGO* expression levels are regulated by ATML1. (**A**) Simulation results showing the cellular concentrations of LGO at a middle time point *t* = 10 in simulated wild type (WT), *ATML1-OX*, and *atml1-3* developing sepals. (**B**) Simulated trajectories of average LGO concentrations in WT, *ATML1-OX*, *LGO-OX*, and *atml1-3* with cafenstrole added at *t* = 20. Upon cafenstrole treatment, LGO concentration drops significantly in WT, *ATML1-OX*, and *atml1-3*. With stochastic fluctuations that take LGO concentration below the *H*_T2_ threshold (e.g. see Fig. 7E), giant cells can lose their giant cell fate, consistent with experimental data (Fig. 4A). For *LGO-OX*, LGO expression is high enough that cafenstrole cannot reduce LGO concentrations down to *H*_T2_, predicting that *LGO-OX* will maintain giant cell fate even in the presence of cafenstrole. (**C**) *LGO* expression in F1 population of WT × *pLGO::3×Venus-N7 35S::mCherry-RCI2A* and *ATML1-OX* × *pLGO::3×Venus 35S::mCherry-RCI2A* backgrounds. Cell outlines are marked by *35S::mCherrry-RCI2A* (red) and LGO expression is marked by *pLGO::3×Venus-N7* (green). Scale bars = 20 µm. (**D–F**) Developmental time series of *LGO* expression levels in WT and *atml1-3* sepals at (**D**) stage 5, (**E**) stage 6, and (**F**) stage 9. Cell outlines are marked by the *35S::mCherrry-RCI2A* (red) and *LGO* expression is marked by *pLGO::3×Venus-N7* (green). Note the faint *LGO* expression in (**E**) for *atml1-3*. Scale bars = 40 µm.

Our model can recapitulate what we observed *in vivo*. This is because in all simulated genotypes, there is a transient in which LGO levels reached a first plateau, mainly driven by the LGO constitutive turnover, and, subsequently, there is a second increase of LGO concentration levels (Fig. 8B), driven by the ATML1-(V)LCFA feedback loops. The first transient arises because there is not enough dimerized ATML1 to induce *LGO* expression. Once sufficient dimerized ATML1 has been created, monomeric ATML1 can be induced, which eventually increases the level of dimerized AMTL1 sufficiently such that *LGO* can be induced downstream (Figs. 8B and S11). However, in the *atml1-3* simulated mutant, this increase in *LGO* is not high enough to promote giant cell development and it also occurs at a later stage, which may impact giant cell fate. Overall, the experimentally determined influence of ATML1 on the timing of *LGO* expression is consistent with the outcomes of the model during giant cell specification.

We then tested whether our model could reproduce the *ATML1-OX* phenotype upon cafenstrole treatment. In *ATML1-OX*, almost every cell becomes a giant cell due to the increase in the basal production of ATML1 in the simulation (Fig. 9A, Movie S3). A higher ATML1 basal production leads to a more rapid increase in dimerized ATML1, which in turn, more rapidly activates the production of LGO (Figs. 8B, S8B, S11A). This early increase in the level of LGO allows all cells to start endoreduplicating within the first two complete cell cycles. When cafenstrole was added at *t* = 20, some of the giant cells divided and became smaller ploidy cells by *t* = 30 (Fig. 9B, Movie S4). Our model suggests that with cafenstrole treatment, LGO expression will decrease more rapidly in *ATML1-OX* than WT (Fig. 8B). This is due to the lack of free (V)LCFA ligands, as the (V)LCFA ligands are being used to create the dimeric ATML1. Once cafenstrole is added, synthesis of more (V)LCFAs is strongly inhibited and thus, the amount of dimerized ATML1 goes down rapidly, causing LGO concentrations to go down with it (Fig. S11B). As well, our model postulates symmetrical giant cell division events in which both daughter cells acquire the same ploidy. However, in the model, some of the de-differentiated giant cells lead to progenies of cells with heterogeneous cell ploidies because some daughter cells continue to go through mitosis and others go through the endoreduplication process. This reflects what we observe *in vivo* (Fig. 4B).

**Fig. 9.**
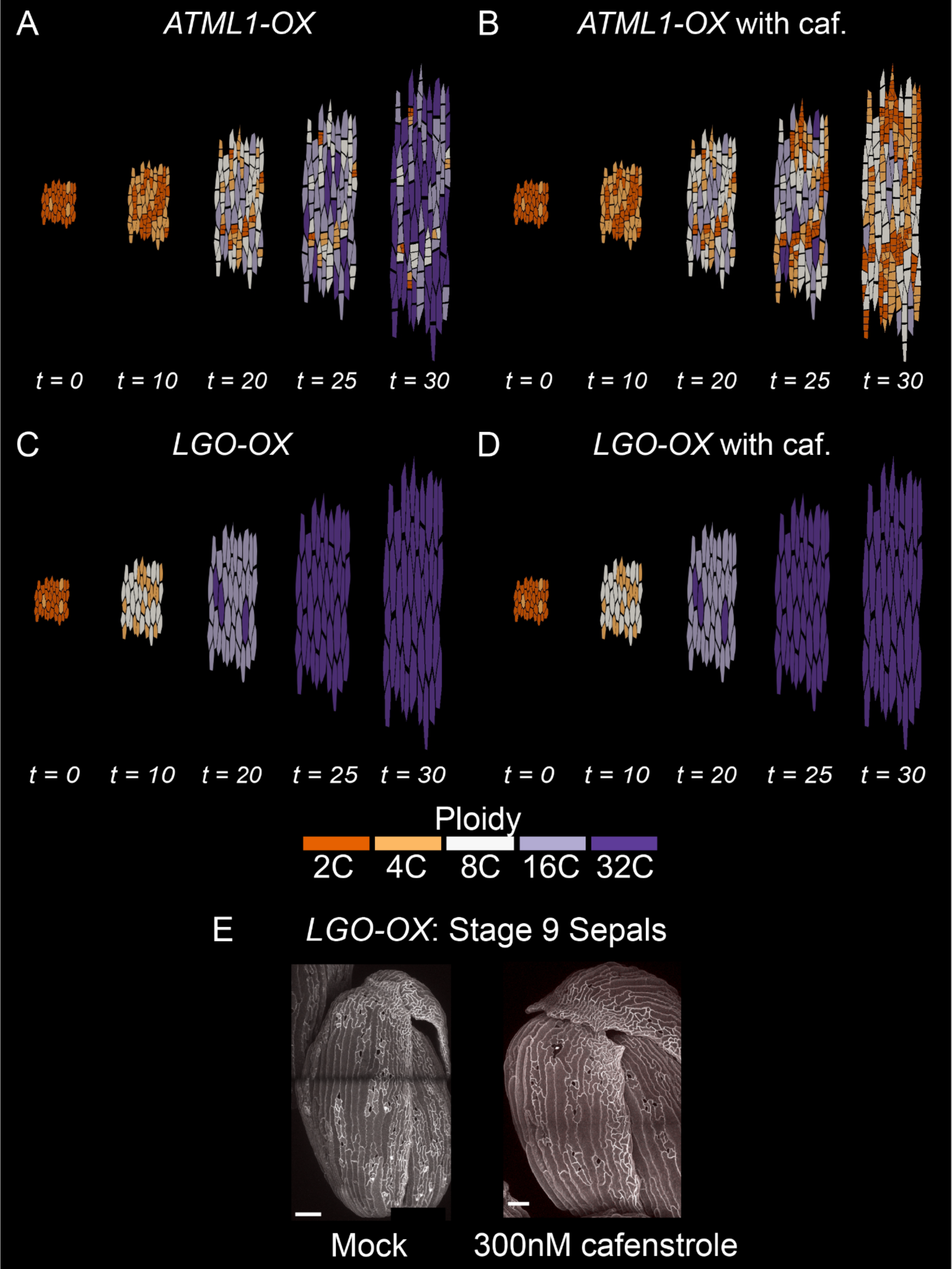
The model accurately predicts the behavior of *LGO* downstream of ATML1 dimerization. (**A–B**) *In silico* simulation of *ATML1* overexpression (**A**) without and (**B**) with cafenstrole predicts that giant cells in *ATML1-OX* can divide upon cafenstrole treatment, emulating the experimental data (see Fig. 4A). (**C-D**) *In silico* simulation of *LGO-OX* without (**C**) and with cafenstrole (**D**) predicts that giant cells generated by LGO overexpression will not resume divisions upon cafenstrole treatment. (**E**) *LGO-OX* flower buds with mock and 300 nM cafenstrole, in which giant cells do not divide. Cells outlines are visualized with the fluorescent membrane marker *35S::mCitrine-RCI2A*. Scale bars = 100 µm.

Lastly, we simulated cafenstrole addition to *LGO-OX* sepals (Fig. 9C–D, Movies S5–S6). We increased the basal production rate of *LGO*, and this resulted in sepals with large numbers of giant cells (Fig. 9C), mimicking those of *LGO-OX* plants (Fig. 9E). After adding cafenstrole at time point *t* = 20 to *LGO-OX,* none of the giant cells divided by *t* = 30. This is consistent with the results in the deterministic model of *LGO-OX*, where we find the *LGO* levels are so high that cafenstrole does not cause much decrease in *LGO* levels (Fig. S8C). This contrasts with the *ATML1-OX* line, where the simulated addition of cafenstrole in the deterministic model brings the *LGO* levels low enough that the giant cell fate could be reversed (Fig. S8B). We tested these outcomes *in vivo* by treating *LGO-OX* sepals with 300 nM cafenstrole. We did not observe giant cell divisions after 7 days of 300 nM cafenstrole treatment (Fig. 9E), a time point by which *ATML1-OX* cafenstrole-treated giant cells divide (Fig. 4B), which matches our simulation results. Thus, ectopic expression of *LGO* overcomes the lack of (V)LCFAs, confirming that *LGO* acts downstream of the ATML1– (V)LCFA feedback loop in the network as predicted. Thus, our experimental tests support our ATML1–(V)LCFA network model of giant cell specification and maintenance.

## Discussion

Through a combination of modeling and experimentation, we identified the central positive autoregulatory loop of a network that produces two cell fates: giant cells and small cells. Our findings suggest that the transcription factor ATML1 plays a key role in this feedback loop by activating the biosynthesis of (V)LCFAs, which are incorporated into lipid ligands of ATML1. Previous research has suggested that START domains in ATML1 and related HD-ZIP transcription factors bind to (V)LCFA-containing lipids (*32*, *33*). Using AlphaFold protein structure prediction, we found that ATML1 contains two adjacent START domains, not just one as previously thought. Our structural prediction suggests that binding of ceramide C18 to the START2 domain of ATML1 triggers homodimerization of ATML1. In our model, these homodimers activate transcription of downstream genes including *ATML1* itself, genes involved in (V)LCFA biosynthesis, and the cell-cycle regulator *LGO*. LGO expression leads to endoreduplication and differentiation of giant cells. Our proposed model is that if ATML1 reaches high concentration during its fate decision phase, this autoregulatory loop triggers specification of giant cell identity and differentiation via endoreduplication by promoting LGO expression. Alternatively, if ATML1 concentration is low during this phase, the cell divides and remains small because its LGO concentration is not promoted by the ATML1–(V)LCFA pathway. The accumulation of these individual cell decisions gives rise to the pattern of scattered giant cells and small cells in the sepal.

To our surprise, not only is this putative feedback loop required for the specification of giant cell identity, but it is also required for the maintenance of giant cell differentiation. When we inhibited VLCFA biosynthesis in this pathway after giant cells had already differentiated, the giant cells lost their identity and started to divide again, presumably due to the decrease of LGO. A few of the daughter cells even adopted stomatal lineage identities and entered a completely different cell fate pathway. The re-entry into division was not anticipated because endoreduplication has been thought to be a terminally differentiated status, since many intertwined copies of the chromosomes would be difficult to segregate during the division process. This behavior is reminiscent of what has been found in the leaf epidermis of *lgo* mutants, whereby differentiated pavement cells start dividing by acting like cells in the stomatal lineage (*54*). As well, previous studies showed that endoreduplication is necessary for the maintenance of trichome (hair) cell identity (*55*). Thus, we reveal that the same regulatory network is not only involved in specification and patterning, but also in the maintenance of cell identity, which is much less understood.

### HD-ZIP transcription factors have (V)LCFA containing ligands

Evidence is growing that the HD-ZIP III and IV transcription factors bind (V)LCFA-containing lipid ligands. Using a protein lipid overlay, it was detected that START1 domain of ATML1 binds to C24 ceramide (*32*). Recent studies on PDF2, a close paralog of ATML1, showed that LysoPC 18:2 (2), LysoPC 18:1 (2), MGDG 34:3, and ceramide t18:1/c24:0 copurified with PDF2 from cell culture and LysoPC 18:1 bound to PDF2 in microscale thermophoresis with liposomes (*30*). Another recent study showed that the START1 domain of the HD-ZIP III protein PHABULOSA (PHB) binds PC with various lengths of fatty acid chains (*56*). It is possible that these proteins bind to different lipids to execute their function and/or that the START domains bind a range of (V)LCFA-containing lipids or (V)LCFAs directly. In the future, it will be relevant to test whether these START domain-containing proteins modulate cellular functions by binding different lipids in different conditions. These studies are all focused on the previously identified START domain, which we call START1. In our analysis, we discovered that the C-terminal START-adjacent domain (SAD) of ATML1 is predicted to form another START domain, START2, which provides a second mechanism through which different lipids may bind and modulate the activity of ATML1. Ceramides are plausible ligands for ATML1 to promote giant cell development because ceramides are strongly reduced in *KCR1RNAi* plants, which also have the strongest giant cell phenotype (*35*). We also observed an increase in ceramide level when we induced ATML1 and in *ATML1-OX* (Fig. 2B, E), corroborating the possibility that ceramides might be involved in a positive feedback loop governing ATML1-mediated giant cell development.

One of the remaining open questions is where and how ATML1 and other members of HD-ZIP III and IV proteins acquire their lipid ligands. It is possible that they acquire ceramides or other lipids from either the nuclear envelope, protein–protein interaction with lipid transfer proteins (LTPs), or during translation on rough endoplasmic reticulum (ER) because the ER is where VLCFAs are synthesized. The nuclear envelope is contiguous with the ER, possibly providing a way for lipids to move between compartments. In our transcriptomic data, several LTPs were upregulated by ATML1 induction, suggesting they are a potential transport candidate. Future experimentation will be necessary to distinguish which of these mechanisms provide (V)LCFAs for START-domain binding.

### START2 dimerization upon lipid binding

Using AlphaFold-Multimer, we predicted that ceramide C18 binding would be stabilized by homodimerization of the START2 domain, and also that LCFA binding to the START2 domain could induce homodimerization of the START1/2 didomain (*51*). A few studies in addition to ours have indicated that ATML1 can form homodimers (*27*, *57*, *58*). Our AlphaFold analysis of HD-ZIP IV proteins suggested that only ATML1, PDF2, HDG2, and GL2 are able to form homodimers using the START2 domain (Fig. S6). GL2 is predicted to dimerize in a different orientation from the other three proteins, which does not involve lipid binding at the interface of the dimer. Traditionally, these HD-ZIP proteins are thought to dimerize through the leucine zipper (*59*, *60*); however, the extra loop in the ZLZ domain of class IV proteins might weaken ZLZ domain dimer formation. Dimerization through the START1/2 didomain might provide enhanced dimer stability. By contrast, AlphaFold-Multimer predicts HD-ZIP III proteins dimerize exclusively through their leucine zipper (LZ) domains and that this does not involve dimerization of the START2 domain. Not having a loop in the LZ domain in HD-ZIP III proteins might give enough strength to form stable dimers, thus not requiring START2 to be involved in dimer formation. Additionally, the C-terminal MEKLA domain in these proteins interfere with START2 dimerization. These contrasting predictions for the dimerization of these proteins might explain why the predictions for the roles of the START domains in the activity of these proteins are also thought to differ.

### A double feedback loop in our mathematical and computational model largely recapitulates experimental observations of giant cell differentiation and de-differentiation

Based on our experimental findings, we are able to propose a new analytical and computational model for giant cell differentiation in a multicellular tissue with growing and dividing cells, as well as the de-differentiation of giant cells when cafenstrole is applied to the tissue. We are also able to model the *atml1-3*, *ATML1-OX* and *LGO-OX* mutants and confirm the model with experimental observations. Interestingly, in all simulated genotypes, the average LGO concentration shows a continuous increase, predicting that LGO concentration has an integrator-like dynamical behavior due to its upstream regulators. Our new model consists of two feedback loops, in which the dimerized ATML1 promotes the production of both (V)LCFAs and monomeric ATML1. We have found that both feedback loops are essential in both differentiating and de-differentiating giant cells, because without these two feedback loops, we are unable to capture all of the observations from experiments. Mathematically, this double feedback loop structure will be particularly interesting for further studies.

Experiments show that cafenstrole application does not induce an immediate drop in LGO concentration (Figs. 7F, S10). This delayed downregulation of LGO is captured in the model (Fig. 7E) by inducing a very low degradation rate for the (V)LCFAs (Fig. S8D). Our model predicts that LGO concentration decreases on day 5 due to the availability of a large pool of (V)LCFAs that can facilitate ATML1 dimerization and which take a long time to fully degrade (Fig. S11B). Once the (V)LCFAs degrade enough, LGO concentration will eventually decrease due to the lack of dimerized ATML1 (Figs. 8B, S11B).

Although the model can recapitulate several key findings from the experiments, there are some limitations. The model suggests that the total ATML1 concentration should exhibit a very large decrease upon cafenstrole treatment by about 30%-40% (Fig. S8A). This differs with our experimental observations, which show an insignificant reduction in total ATML1 concentration (Fig. S4C, D). Our model can tolerate such a small reduction in total ATML1 concentration if we reduce the feedback strength between dimerized ATML1 and monomeric ATML1. However, this has other downstream consequences due to the stochastic nature of the simulations, resulting in either too many or insufficient dividing cells in *ATML1-OX*. As well, we note that previous work on the root has shown a substantial decline in ATML1 expression after cafenstrole treatment (*32*). Therefore, more work is needed to explore the consequences of the stability of ATML1 expression and whether this is tissue dependent. Overall, our study sheds light on the intricate regulatory network underlying the generation and maintenance of specialized cell identities.

## Methods

### Plant growth conditions and mutants used in this study

Seeds were planted directly on Lambert general purpose mix LM111 soil and were stratified for 2 days before placing them in the Percival chamber. Plants were grown at 22°C in continuous light to avoid gene expression changes associated with the circadian clock. Mutant seeds of *atml1-3* (SALK_033408), *cer2-1* (CS32), *cer1-1* (CS31), *cer3-1* (CS33), *cer6-2* (CS6242), *cer6-2R* (CS3946), *cer1-10* (CS92), and *kcs2 kcs20* (CS71729) were obtained from Arabidopsis Biological Resource Center (ABRC). *KCR1-RNAi*, estradiol inducible *RPS5A>>ATML1* (proRPS5A-ATML1/pER8 and proATML1-nls-3×GFP line #7), *pPDF1::FLAG-ATML1* (*ATML1-OX*), *pATML1::mCitrine-ATML1* were previously described (*13*, *38*, *43*). Accession Col-0 was used as the wild type accession throughout the experiments, except for in comparison to the *cer1-1*, *cer3-1*, *cer6-2*, and *cer6-2R* mutants which are in the Landsberg *erecta* (L*er*) background. The small cell (CS70134) and giant cell markers (pAR111) were previously described (*12*, *46*).

### Tissue collection for RNA-seq

Inflorescences including flower buds through stage 12 were cut from estradiol inducible RPS5A>>ATML1 (proRPS5A-ATML1/pER8 and proATML1-nls-3×GFP line #7; (*43*)) plants and cultured on MS medium 1% sucrose 0.5 g/L MES pH5.7 plates supplemented with estradiol (0.1 μM, 1 μM, 10 μM) or mock (0.1% ethanol; estradiol stocks were dissolved in ethanol). Inflorescence tissues were incubated on the treatment plates for 0 h, 8 h, 16 h, 24 h, and 32h hours and three to four inflorescences were collected in an Eppendorf tube per biological replicate. The tissue was snap-frozen in liquid nitrogen and stored at −80°C.

### RNA extraction and RT-qPCR

Total RNA was isolated from frozen tissue (see Tissue Collection for RNA-seq above) using the RNeasy Plant Mini Kit (QIAGEN Cat #74904) as per the manufacturer’s instructions. One microgram of isolated RNA was treated with Invitrogen DNase I (Cat #18068015) and incubated at room temperature for 25 min. DNase was inactivated by adding EDTA and samples were incubated at 65°C for 10 min. First-strand cDNA synthesis was carried out using oligo (dT) and Invitrogen Superscript II (Cat # 18064014) according to the manufacturer’s protocol. RT-qPCR was performed using Roche SYBR Green Master Mix (Cat #4707516001) according to the manufacturer’s instructions on a Roche LightCycler 440 system. For each sample, three biological replicates and three technical replicates were used. ATML1 was quantified using primers oHM58 (GAGCTAGAGTCGTTCTTCAAGG) and oHM62 (GTTCTCGTGCCTCTCATGTTGTG). Gene expression was quantified using the ΔΔCt method.

### RNA-sequencing

Libraries for sequencing were prepared according to the protocol described in (*61*). In brief, 5 µg RNA was treated with DNase and incubated at RT for 15 min. DNase was inactivated by adding EDTA, and RNA was purified using Invitrogen Dynabeads oligo (dT)25 (Cat# 61002). Purified RNA was fragmented at 94°C and first-strand cDNA synthesis was performed using Invitrogen random hexamer (Cat# 48190011) and Invitrogen Superscript II (Cat# 18064014) followed by second-strand synthesis using Fermentas DNA Pol I (Cat# EP0041). The resultant cDNA was end-repaired using NEBNext end pair enzyme (Cat# E6050S) and Klenow DNA polymerase (Cat# M0210S). cDNA was A-tailed with Fermentas Klenow 3′ to 5′ exonuclease (Cat# EP0421) and ligated with adaptor oligonucleotides (NEB NEXT adaptor oligos) using Mighty Mix Ligase (Cat# TAK6023). Adaptor-ligated cDNA was purified using XP bead purification according to the manufacturer’s protocol. PCR enrichment was performed with PE 1.0 (5′-AAT GAT ACG GCG ACC ACC GAG ATC TAC ACT CTT TCC CTA CAC GAC GCT CTT CCG ATC* T-3′) and PE 2.0 (5′-CAA GCA GAA GAC GGC ATA CGA GAT CGG TCT CGG CAT TCC TGC TGA ACC GCT CTT CCG ATC* T-3′) for 15 cycles using Phusion polymerase (NEB M0530S). The library was purified by electrophoresis on a 1.2% agarose gel to get rid of adapter dimers. Libraries were sequenced at the Genomics facility of Cornell University Biological Resource Center using an Illumina NextSeq 500 instrument to generate 75-nt reads.

### RNA-seq data analysis

We obtained the sequencing data as gzip-compressed FastQ files (fastq.gz) from an Illumina NextSeq 500 sequencer. We performed the RNA-seq analysis essentially as in (*62*). Briefly, reads were first quality filtered with *quality_trim_fastq.pl* (https://github.com/SchwarzEM/ems_perl) to remove reads that failed CHASTITY and read sequences over 84 nt. We then further quality-filtered the reads with fastp 0.20.1 (*63*) using the arguments ‘*--dont_overwrite--length_required 75--max_len1 84*’ to remove adapter and low-quality sequences. The resulting trimmed and quality-filtered reads are available in the Sequence Read Archive (SRA; https://www.ncbi.nlm.nih.gov/sra) in BioProject PRJNA1074358. We mapped RNA-seq reads to *Arabidopsis thaliana* TAIR10 cDNA sequences (coding sequences plus 5’ and 3’ UTRs) with Salmon 1.4.0 using the arguments ‘*quant--validateMappings--seqBias--gcBias--libType A -- geneMap [transcript-to-gene index]*’; for each RNA-seq replicate, this determined the level of gene expression in transcripts per million (TPM) and number of mapped reads per gene.

### Differential gene expression and GO term analysis

DESeq2 (version 1.18.1) was used to conduct the differential expression analysis (*62*, *64*). Fold change in gene expression was calculated for each sample relative to mock treated tissue collected at the same time point. Shrinkage was applied to fold change values by DESeq2. Accompanying *p*-values were calculated with the Wald test, and adjusted for multiple testing with the Benjamini-Hochberg procedure by DESeq2.

An adjusted *P*-value cutoff of 0.05 was applied to identify differentially expressed genes in each sample. After separating up- and downregulated genes, gene lists were supplied to the PANTHER web server (release 17.0) for GO overrepresentation testing (*65*). The reference list used was the complete set of protein-coding genes for *A thaliana*, which PANTHER obtained from the Reference Proteomes project (*66*). Complete biological process, cellular component, and molecular function annotation datasets were tested for overrepresentation using Fisher’s exact test, and *P*-values were adjusted for multiple testing via Bonferroni correction. Data were collected for any term with at least one associated gene, and adjusted *P*-value and fold enrichment cutoffs were applied *post hoc* to identify significant and highly enriched terms.

### Transcriptomic analysis to predict ATML1 targets

We created a normalized expression dataset containing Transcripts per Kilobase Million (TPM) measurements for 27,655 genes under four different conditions: a control condition, a 0.1 µM induction of ATML1, a 1 µM induction of ATML1, and a 10 µM induction of ATML1. For each ATML1 induction condition, measurements were taken at four time points: 8 h, 16 h, 24 h and 32 h (except for the 10 µM condition, which did not have a 16 h measurement). For the control condition, measurements were taken at only the 0 h time point. For each condition at each time point, measurements were taken for three replicates.

To predict potential targets of ATML1, the Spearman correlation was calculated between ATML1 and every expressed gene across the 24 h and 32 h time points, and the genes were selected that were significantly correlated with ATML1 concentration with a Bonferroni-corrected *P*-value less than or equal to 0.05. The code for this correlation analysis can be found in the “get_spearman_correlations.py” file in the data repository (DOI: 10.17605/OSF.IO/QEXHR).

This selection process yielded 141 potential target genes, including 110 that were positively correlated with *ATML1* and 31 that were negatively correlated with *ATML1*.

From time series plots of TPM, we visually inspected the expression of these genes relative to *ATML1* to check whether they followed the same pattern (which would suggest a possible regulatory relationship). The plots for these correlated genes can be found in the folder called “timeseries_plots” in the code repository, and the code for generating the plots can be found in the “timeseries_plots.py” file.

In addition to the time series plots, scatter plots of potential target gene expression against ATML1 expression were generated using data from the 24 h and 32 h timepoints. Hill functions were then fitted to these plots to gain insight into possible regulatory dynamics.

We used the following Hill function to model positive regulation between ATML1 and a target:

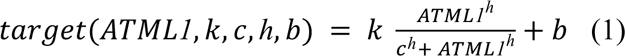

Here, the target expression is represented as a function of the ATML1 expression and four constant parameters, *k*, *c*, *h*, and *b*, which can be described as follows: *k* defines the maximum level of target expression that can be activated by ATML1, *c* is the macroscopic dissociation constant, *h* is the Hill coefficient, and *b* represents a basal expression level of the target that occurs when there is no ATML1 expression.

To fit our model to the data, we defined an objective function as follows:

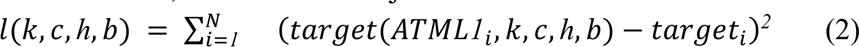

where *target_i_* and *ATML1_i_* represent the experimental data points for target and ATML1 expression and *target*(*ATML1_i_*, *k*, *c*, *h*, *b*) represents the model prediction for target expression given a parameter set. We applied a numerical optimization in Python to find the values of *k*, *c*, *h*, and *b* which minimized the objective function in Eq. 2.

In addition to modeling positive regulation, we defined a Hill function to model negative regulation between ATML1 and a target:

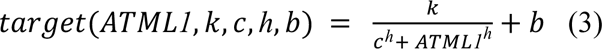

This function is similar to the positive Hill function, except that the target expression decreases with increasing ATML1 expression. We then used Eq. 2 along with the same minimization pipeline to find the best-fit parameters.

### Lipidomics

ATML1 inducible lipidomics was performed using RPS5A>>ATML1 inflorescence tissues. In brief, inflorescence tissues were dissected by removing old and opened flowers and were placed on plates containing MS media supplemented with 0.1 µM, 1 µM, or 10 µM estradiol or ethanol (Mock). Approximately 25–28 mg tissue was collected in Eppendorf tubes after 24 and 32 h of treatment. Biological pentuplicates were collected for each treatment.

Total lipids from inflorescence tissues were isolated using the protocol from (*67*). Briefly, inflorescence tissues were ground into a fine powder using liquid nitrogen and 1 mL methyl-tert-butyl-ester:methanol (3:1), followed by a phase separation adding H2O:MeOH (3:1 v/v). For lipid analysis, 800 µL from the lipid-containing organic phase was dried under vacuum. The dried residue was resuspended in 200 µL UPLC-grade acetonitrile:isopropanol (70:30) mixture. From this mixture, 2 µL was injected individually onto an Acquity UPLC system using an RP C8 column and analyzed by MS (*67*). The samples were measured in positive and negative ionization mode. The mass spectra were acquired using a Q-exactive Orbitrap high-resolution mass spectrometer: Fourier-transform mass spectrometer (FT-MS) (ThermoFisher Scientific, Waltham, MA, USA, https://www.thermofisher.com). Processing of chromatograms, peak detection, and integration were performed using RefinerMS (version 5.3; GeneData). Mass features were annotated using an in-house lipid database (*68*) allowing for 10 ppm mass error, and a dynamic retention-time shift window of 0.1.

### VLCFA inhibitor treatments

A stock solution of 0.1 M cafenstrole (FUJIFILM Wako Pure Chemical Corporation) in acetone was made and added to MS plates to make final concentrations of 30 nM and 300 nM. Equal volumes of acetone were used as mock controls. Similarly, fentrazamide (Sigma Cat #37903-100MG) stock of 10 mM was made in isopropanol appropriate volumes and were added to MS medium immediately before pouring the plates to make final concentrations of 3 µM. Equal volumes of isopropanol were used as mock controls.

### Confocal imaging

All reporter lines were imaged using a Zeiss 710 confocal microscope with a 20x water dipping objective NA 1.0 and or a 10x air objective for imaging the epidermis of PI stained sepals. For sepal epidermis staining, the mature sepals were dissected from flowers and placed on a glass slide with a coverslip and a drop of 0.01% Triton X-100, and were stained with propidium iodide (PI) solution for 15 min. After staining, sepals were mounted on a glass slide and covered with a coverslip. PI-stained sepals were imaged with a 514 nm laser for excitation, and emission was collected at 566 to 650 nM. VLCFA inhibitor-treated inflorescence tissues were imaged on the plates on which they were treated.

### Flow cytometry

Inflorescence tissues of transgenic plants of *pPDF1::FLAG-ATML1 pML1::H2B-mGFP* were kept on 300 nM cafenstrole and mock MS medium plates for 10 days of growth. Nuclei were isolated from the entire inflorescence tissue and stained with PI according to the protocol in (*69*). Flow cytometry was performed with a BD FACS machine. Nuclei were gated for GFP-positive nuclei to select the epidermal nuclei. GFP-positive nuclei were analyzed for ploidy on the basis of PI staining.

### Structure prediction using AlphaFold-Multimer

Predictions were performed using AlphaFold-Multimer via ColabFold 1.3.0 (*51*, *70*). Amino-acid sequences for Arabidopsis HD-ZIP III and IV proteins obtained from UniProt and were used as input to predict dimer structures with default settings. Ceramide C18 was d18:1/18:0. The predicted interface TM-score (ipTM) scores for the top three models obtained from AlphaFold-Multimer are shown in Table S1.

### Yeast two-hybrid (Y2H)

The pENTR/D-TOPO plasmid containing the wild-type *ATML1* cDNA was previously described (*28*). Sequence fragments of *ATML1* (START1, START2, and didomain) were generated using the Q5 Site-Directed Mutagenesis Kit (New England Biolabs) with the following primers: ATML1START1F (CACCATACCTTCTGAG GCTGATAAG), ATML1START1R (CTACATGGAACTGGCGAGCCG), ATML1START2F (CACCGCCAGCAACATTCCGG) and ATML1CR (ATCGATTAGGCTCCGTCGCAG). *ATML1* fragments were transferred to pDEST32 bait and pDEST22 prey vectors (Proquest Two-Hybrid System, Invitrogen) using Gateway LR Clonase II Enzyme mix (ThermoFisher Scientific). Bait and prey constructs were transformed into the haploid yeast strains Y2HGold and Y187 (Matchmaker Gold Yeast Two-Hybrid System, Clontech), respectively, using the LiAc method (*71*). Matings to form diploids were performed as described previously (*28*). To assay for interactions of bait and prey in diploids, cells were grown to a density of OD_600_=0.5–1.0. Four serial 4-fold dilutions were prepared from normalized cultures. Cells were spotted onto permissive and selective media with a 48-pin multiplex Frogger (Dankar, Inc.), followed by incubation at 30°C for 3–5 days, and imaging using a GelDoc XR+ System (BioRad).

### Y2H quantitative α-galactosidase assay

Three independent colonies from each yeast diploid were selected to perform quantitative α-galactosidase liquid assays, modified from a β-galactosidase assay (*31*). Cells were cultured in permissive (-Leu-Trp) medium in a 24-well plate to a density of OD_600_=0.5–0.8, followed by pelleting 1.2 mL cells and resuspension in 100 µL Z-buffer (60 mM Na_2_HPO_4_, 40 mM NaH_2_PO_4_, 10 mM KCl, 1 mM MgSO_4_•7H_2_O, 50 mM β-mercaptoethanol, pH 7.0). Cells were subjected to three freeze–thaw cycles followed by pelleting. The assays were performed in 96-well microplates. In each well, 30 µL cell supernatant was incubated with 80 µL substrate (10 mg/ml *p*-Nitrophenyl α-D-galactopyranoside (ThermoScientific L13376.ME) in 340 mM sodium acetate, pH 4.5) for 1–3 h at 30°C. Reactions were stopped with 190 µL of 1 M Na_2_CO_3_. Yellow color formation (A_405_) was measured in a BioTek Epoch2 microplate reader (Agilent). The following formula was used for quantification: Units α-galactosidase = V(30 µL) × A**_405_**/time × V(200 µL) × OD_600_.

### Bimolecular fluorescence complementation (BiFC) in *N. benthamiana*

To generate constructs for split GFP, Gateway-compatible BiFC vectors 125-NXGW and 127-CXGW for N-terminal fusions of split GFP were used in LR recombination reactions with pENTR/D-TOPO plasmids containing *ATML1*. Inoculation of *N. benthamiana* via *Agrobacterium tumefaciens* was performed as described in (*72*). Propagation of *N. benthaniana* plants, *A. tumefaciens* GV3101 transformation, infiltration of leaves, and incubation prior to microscopy was conducted as previously described (*28*). GFP fluorescence was imaged with a Zeiss LSM-5 Pascal microscope having a 488 nm Argon laser for excitation, and 510-540 nm band pass filter for emission. Mean pixel intensities were quantified using ImageJ software from six images for each construct combination.

### Co-immunoprecipitation (Co-IP)

100 mg of dissected inflorescence tissues of *pPDF1::FLAG-ATML1* X *pATML1::mCitrine-ATML1* (F1 generation) plants including flower buds up to stage 8-9 of flower development were used for protein extraction and Co-IP as described in (*73*) with modifications in antibodies used. In short, we used EZview red ANTI-FLAG M2 affinity gel (Sigma Cat #F2426 lot #SLBN6224V) according to the manufacturer’s protocol to bind the FLAG-ATML1 protein and washed it with stringent washing buffer conditions to remove the non-specific binding proteins. We then ran a western blot using the GFP antibody (Takara living colors EGFP monoclonal antibody JL-8, Cat# 632569) at 1:4000 dilution followed by secondary antibody (EMD Millipore goat anti-mouse antibody Cat# AP308P Lot # 3436981) conjugated with HRP. HRP signal was detected using chemiluminiscence (Thermo Scientific Pierce ECL Western Blotting Substrate Cat# PI32109).

### Quantification of ATML1 and LGO reporters

Segmentation of the nuclear-expressed *pATML1::mCitrine-ATML1* and *pLGO::3×Venus-N7* constructs were completed using ilastik (*74*). The Pixel Classification module was used to construct a probability map, which was then transferred to the Object Classification module to perform segmentation. The hysteresis thresholding scheme was used with a smoothing parameter of 1.0 for all three dimensions, 0.85 and 0.55 for the high and low thresholds, respectively, and 10 pixels as the minimum size for a nucleus. A CSV file with key characteristics was then generated for each segmented nucleus, including total signal intensity and total volume of the nucleus.

For the LGO quantification, segmentation of the membrane marker (*35S::mCherry-RCI2A*) was completed using ilastik’s boundary-based segmentation using multicut (*75*). For each time point and replicate, the mean nuclear concentration of *pLGO::3×Venus-N7* was calculated by dividing the total fluorescence signal per nucleus by the total nuclear volume, and then computing its mean across the cell population in the sepal, both epidermal and subepidermal. To account for the cells which did not express *pLGO::3×Venus-N7*, we counted the number of cells segmented from the cell membrane quantification. This was performed by importing the segmented numpy matrix file (.npy) from ilastik into a Python script as a matrix. Each element of the matrix corresponded to a voxel and the value of the element corresponded to the cell label it is a part of, thus allowing the number of unique cell labels to be used as a proxy for the number of cells in the original image. If the number of nuclei segmented is smaller than the number of cells segmented, there are cells that are not expressing any *pLGO::3×Venus-N7* and thus we prescribe that these cells have a nuclear concentration of 0 for that reporter. The mean concentration was then calculated using these values and was normalized by the mean concentration of the replicate at day 0 (Fig. S12). For the ATML1 quantification, the mean nuclear concentration of *pATML1::mCitrine-ATML1* was calculated by dividing the total fluorescence signal per nucleus by the total nuclear volume and then computing its mean across the segmented nuclei.

### Mathematical Modeling

A stochastic computational model was constructed and implemented for ATML1, LGO, and (V)LCFA-mediated giant cell fate decisions. The schematic of the model can be visualized in Fig. 7A. ATML1 cell concentration dynamics was modeled by treating ATML1 as a protein that is basally produced and linearly degraded. Similarly, (V)LCFAs were considered to be basally produced and linearly degraded. Note that we attempt to understand the dynamics of the (V)LCFA that are available to bind to ATML1, not all of the fatty acids in the tissue. Therefore, the term (V)LCFA in this model refers to the fatty acids that are available to bind to ATML1. ATML1 and (V)LCFAs must bind together and dimerize to activate the production of LGO. For simplicity, the resulting complex will be referred to as dimerized ATML1. The dimerized ATML1, which also linearly degrades (albeit at a slower rate than ATML1), activates ATML1 and (V)LCFAs production. The dimerized ATML1 also activates a downstream target, LGO. LGO is also basally produced and linearly degraded, and if LGO levels exceed a certain threshold at a certain time of a cell cycle, cell division is inhibited (see further details below on the cell cycle implementation through a timer). We also implemented an inhibitory drug to understand the effect of a (V)LCFA-synthesis inhibitor on the system, mimicking the cafenstrole treatments from our experiments. When applied to our system, this inhibitor blocks the basal production of (V)LCFAs and the activation of (V)LCFAs production by the dimerized ATML1. If the LGO concentration level in an endoreduplicating cell falls below a second threshold (lower than the first threshold), the cell will then be able to divide again, reversing its giant cell fate decision (Fig. 7B).

The deterministic expressions for the dynamics of ATML1, (V)LCFAs, dimerized ATML1, and LGO concentrations in cell *i* can be written as follows, where *A_i_* = [*ATML1*]*_i_*, *B_i_* = [Dimerized *ATML1*]*_i_, V_i_* = [(V)LCFA]*_i_*, *L_i_* = [*LGO*]*_i_*:

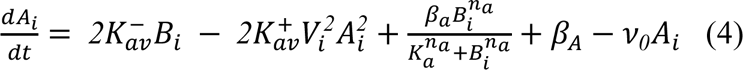

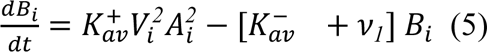

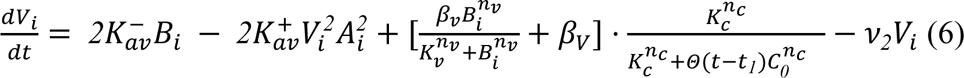

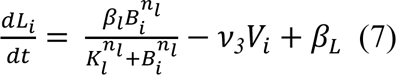

where the parameters are defined in Table S2 and Θ(*t*) is the Heaviside function. We assume that the ATML1 dimerization rate is much faster than the other production and degradation rates. For Fig. S8, the deterministic simulations were completed using values of 0 for all initial conditions. A fourth order Runga-Kutta method was used with *dt* = 0.0001 to avoid numerical instabilities.

For the computational simulations, similar to (*13*), a cell is defined by a set of vertices in 2D, and each cell grows exponentially and anisotropically by moving the vertices away from the center of mass of the tissue. We note that all cells grow anisotropically, and they divide or endoreduplicate according to a timer variable unique to each cell. This timer variable increases linearly with time and is reset once it reaches a threshold. Therefore, an equation for this timer can be written as

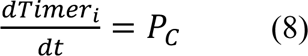

where *Timer_i_* is the timer variable for cell *i* and *P_C_* is the basal timer production rate. We update the timer every timestep using the following equation:

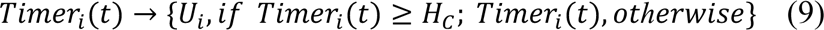

where *U*_i_ is a uniform randomly distributed number chosen from the interval [0,0.5), and *H_C_* is the threshold for the timer at which it resets to start a new cell cycle, coinciding with the timer value at which cells divide. To balance the effects of cellular growth, we implemented dilution terms for ATML1, (V)LCFA, the dimerized ATML1, and LGO.

Cell ploidy was modeled as a discrete variable that is dependent on the cell timer, cell division, and the ATML1 network. Cell ploidy increases from 2C to 4C when the timer reaches a threshold *H*_s_, which represents the S phase, and decreases to 2C if the cell divides. Cell division for cell *i* occurs at the 4C stage when the timer reaches a second threshold *H_C_* and [*LGO*]*_i_* is higher than a specific threshold *H*_!_*_1_*during this 4C stage (i.e., the G2 phase, which is the cell cycle stage while the timer is between *H_S_* and *H_C_*). If this happens, endoreduplication occurs and the cells reset their time according to Eq. 9. Cells which endoreduplicate may divide again if [*LGO*]*_i_* falls below another specified threshold *H*_!_*_2_* when the timer reaches *H_S_*. If a cell divides, each of the two daughter cells inherits half the ploidy of the mother cell (e.g. a single 4C cell becomes two 2C cells). The daughter cells also inherit the ATML1, (V)LCFA, Dimerized ATML1, and LGO concentration levels of the mother cell, as well as having their timers reset by Eq. 9. These daughter cells then can either divide again or can continue to endoreduplicate. This is determined when they enter S phase, i.e. once their timer has reached *H_S_*. If a daughter cell’s LGO levels are above *H*_!_*_2_*, the cell does not divide anymore and enters the endoreduplication cycle. If a daughter cell’s LGO levels are below *H*_!_*_2_*, it will divide yet again into two equal ploidy cells and those daughter cells will go through this exact same process, unless the daughter cells are at 2C.

Dynamic stochasticity was introduced into the ATML1, (V)LCFA, dimerized ATML1, LGO, and timer variables using a similar algorithm explained in (*13*). For every ATML1, dimerized ATML1, LCFA, LGO, and Timer variable in a cell *i*, the resulting stochastic equations read

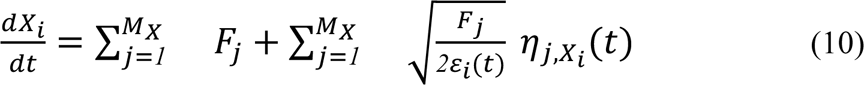

where *F*_j_ is a generic function consisting of any of the generic variables *X*, including the production and degradation terms and *M_x_* is the number of generic functions needed to simulate the system for a given variable. The normalized cell area, ε_i_(*t*), is assumed to be ε_i_(*t*) = *E_0_E_i_*(*t*), where *E_0_* is an effective cell area used to normalize the noise and *E_i_*(*t*) is the current area of cell *i* in arbitrary units. Lastly, *η*_j,xi_ are temporally and spatially uncorrelated, statistically independent Gaussian white noise random variables, which follow 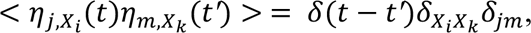 where *j* and *m* are cell indices, *X_i_* and *X_k_* are the modeled variables, δ*_xi,xk_* and δ_.1_ are Kronecker deltas and δ(*t* −*t′*) is the Dirac delta. For the terms in Eqs. 4-6, which correspond to the mass action law for the dimerization process, the random noises are correlated so that the mass conservation law is held.

Integration of the stochastic Langevin equations with the Îto interpretation was performed using a variation of the Heun algorithm (*76*) using an absorptive barrier at 0 to prevent negative values. Growth and dilution effects were considered deterministic; thus, these were integrated with an Euler algorithm. To avoid numerical instabilities, stochastic integration was performed with a timestep of *dt =* 0.001. Cell division was performed according to the shortest path rule in which the new wall passes through the center of mass of the mother cell (*77*). As explained above, the daughter cells inherit the same ATML1, (V)LCFA, Dimerized ATML1, and LGO concentrations when they are generated, but they may have different sizes. Additionally, these cells will have different initial timer variables due to Eq. 9.

For all cells, we set uniformly distributed random initial conditions for ATML1 and (V)LCFA variables in the interval of [0,1) and set the dimerized ATML1 and LGO variables to 0. Timer initial conditions were set to be correlated to the cell size of the initial template with the following expression:

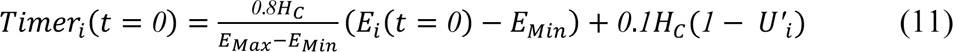

where *U_i_^′^* is a uniformly distributed random number between [0,0.1) and *E_Min_* and *E_Max_* are the areas of the smallest and largest cells, respectively. This allows larger cells to start at a more advanced cell-cycle stage and thus be more likely to divide sooner. Ploidies were initially set to either 2C or 4C depending on whether the initial timer set by Eq. 11 was above or below the S phase threshold *H_S_*.

Dilution effects in the deterministic simulations were accounted for by rescaling the degradation rates of each concentration variable, i.e., by adding the average cell area growth rate to the different degradation rates appearing in the Eqs. 4–7. The average cell area growth rate was computed by dividing the change in cell area from *t =* 29.5 to *t* = 30 by area of each cell at *t* = 30. In this case, the value was 2/25.

The computational implementation of our model was performed using the open source C++ Tissue package (*78*). We simulated eight different situations: wild type, wild type with cafenstrole, *ATML1-OX*, *ATML1-OX* with cafenstrole, *LGO-OX*, *LGO-OX* with cafenstrole, *atml1-3*, and *atml1-3* with cafenstrole. To ensure that our simulations were reproducible, we have simulated each of these situations using three different seeds. In the file myRandom.cc in Tissue, we changed the starting seed, MSEED, to 300, 400, and 500 for the three simulations. This ensures that the seed we selected was not the determining factor in our results. Data analysis and plots from the simulation outputs were performed with Python 3.10, the Matplotlib package (*79*), and MATLAB. The visualization of the simulated growing tissues was performed in Paraview (*80*). See Table S2 for parameter values used in all simulations.

### Statistics

Data are presented as means ± SD from at least three independent samples/experiments (*n ≥* 3). Error bars represent one SD. An *\** implies *P* < 0.05, which is considered significant. Analysis was computed using one-or two-way analysis of variance (ANOVA) and an unpaired student *t*-test.

## Acknowledgements

We thank ABRC for the mutant seeds of VLCFA biosynthesis genes, Frederic Beaudoin for kindly sharing the *KCR1 RNAi* lines with us, and John Chandler for his helpful comments and suggestions in the writing of this manuscript. We acknowledge Carly Rodriguez, Emily Phung, Hoaxing Harry Hou for taking care of plants, assisting with confocal imaging, and tissue collection.

## Funding

National Science Foundation IOS-1553030 (AHKR)

National Science Foundation MCB-1616818 (KS)

Max Planck Society (NR, PFJ)

The U.S. National Science Foundation grant 2226270 (AS)

Samuel and Nancy Fleming Research Fellowship (BVLV)

National Institute of General Medical Sciences of the National Institute of Health Award No. P20GM103418 (LEA)

The USDA National Institute of Food and Agriculture Hatch/Multi-State project 1013013 (KS)

Johnson Cancer Research Center at Kansas State University (AK, LEA)

European Union’s Horizon 2020 Research and Innovation Programme, Project PlantaSYST (SGA-CSA No. 739582 under FPA No. 664620) financed by the European Regional Development Fund through the Bulgarian “Science and Education for Smart Growth” Operational Programme (SA, ARF)

## Author Contributions

BVLV and AHKR conceived and designed most of the experiments. BVLV carried out the RNA-seq, confocal imaging, pharmacological treatments, and generation of transgenic lines for the confocal microscopy. RNA-seq analysis was done by BA, EMS and BVLV. Correlation analysis of ATML1 concentration with downstream gene expression was done by MS-A and AS. VLCFA mutants from ABRC were screened by BVLV and SM. SRB and JCF did the ATML1 structural predictions. BB, SRB, JCF performed ATML1 biochemistry. NJR and PFJ designed and constructed the mathematical model. NJR performed the deterministic and stochastic simulations. NJR performed quantitative image analysis. LEA conducted Y2H experiments. AK performed BiFC experiments. KS designed BiFC and Y2H experiments and participated in data analysis. Lipidomics was done by BVLV, ASkirycz, SA, ARF. BVLV, NJR, SRB, KS, and AHKR wrote the manuscript. PFJ, MSA, KS, and all authors edited and approved the manuscript.

## Competing interests

The authors declare that they have no competing interests.

## Data availability

All data are available in the main text, in the supplementary materials, or in publicly available data repositories. Image data, mathematical modeling code, and transcriptomic analysis code are available at the Open Science Framework (osf.io), DOI: 10.17605/OSF.IO/8YSQA. RNA-seq data is available at the Sequence Read Archive (SRA; BioProject PRJNA1074358).

## Notes

https://doi.org/10.17605/OSF.IO/8YSQA

https://www.ncbi.nlm.nih.gov/bioproject/PRJNA1074358

### Competing Interest Statement

The authors have declared no competing interest.

