## Supplementary Figures for "The transcription factor ATML1 maintains giant cell identity by inducing synthesis of its own long-chain fatty acid-containing ligands"

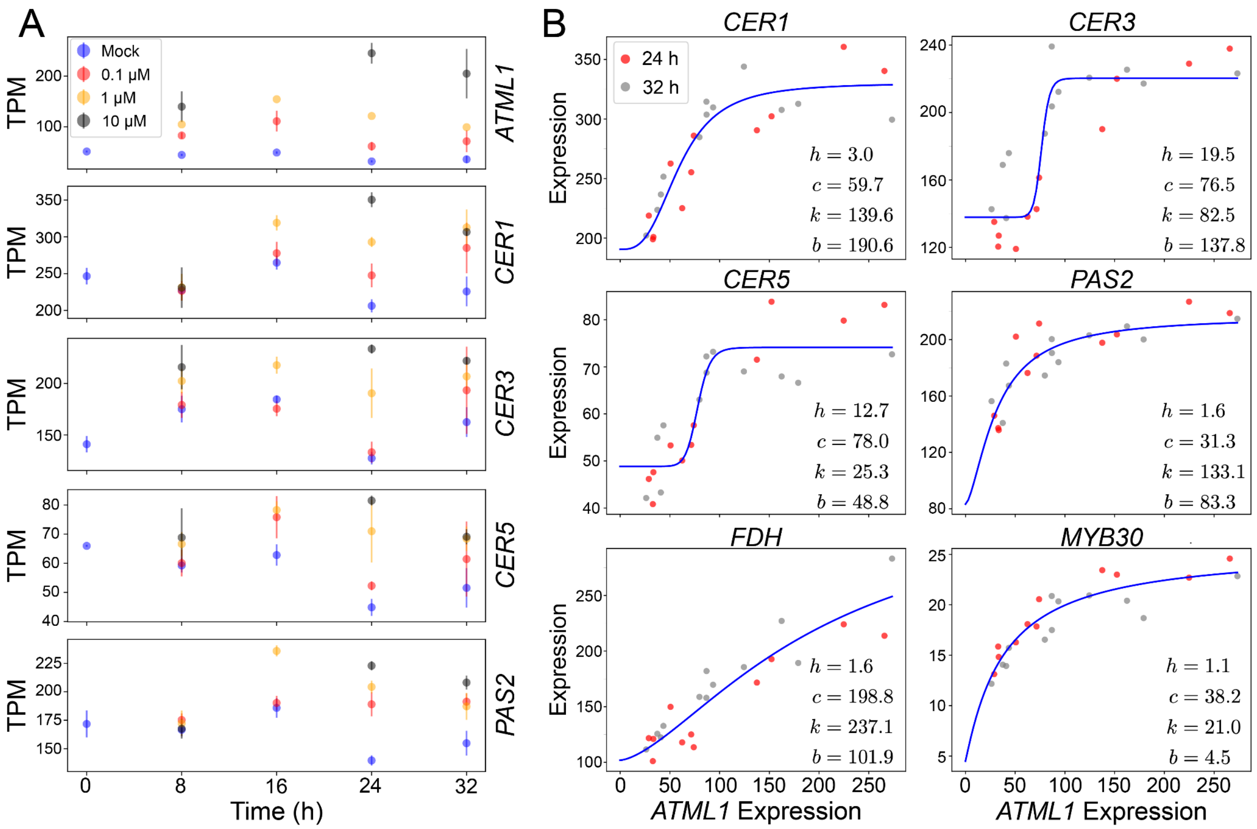

**Fig. S1. ATML1 activates (V)LCFA biosynthetic process genes in a concentration-dependent manner, associated with Fig. 1.**

(**A**) The top plot shows the transcript per million (TPM) expression of *ATML1* at different time points and induction levels (bars show one standard deviation). Higher expression at the higher induction levels confirms that the experiment worked as intended. The Spearman correlation was calculated between *ATML1* and every other gene in the dataset across the 24 h and 32 h time points and genes were selected that had a Bonferroni-corrected $p$-value less than or equal to 0.05. This analysis yielded 141 genes that were significantly correlated with *ATML1*. The plots for four of those genes (*CER1*, *CER3*, *CER5*, and *PAS2*) are shown in the figure. The purpose of these plots is to visually inspect the expression levels of the genes that were found to be significantly correlated with ATML1 at different time points and induction levels. If a gene is significantly correlated with ATML1 and differentially expressed at different induction levels of ATML1, we consider it to be a possible downstream target worthy of further investigation. See the Methods section for more details on this analysis and Data S2 for the plots of all 141 significantly correlated genes. (**B**) For each of the 141 genes that were significantly correlated with *ATML1* across the 24 h and 32 h timepoints, their expression was plotted against the expression of *ATML1*, and Hill function curves were fitted to these plots. Examples for six genes (*CER1*, *CER3*, *CER5*, *PAS2*, *FDH*, and *MYB30*) are shown on the right, as well as their fitted values for the Hill function parameters. The goal of this analysis was to assess whether the relationship between ATML1 and each of its potential downstream targets was graded or switch-like. The difference between these two types of potential regulatory relationships can be seen both visually and with the fit Hill parameters: fit values of $h$ close to 1 correspond to a graded relationship, and larger values of $h$ correspond to a switch-like relationship. See the Methods section for more details on this analysis, and Data S3 for the plots of all 141 significantly correlated genes.

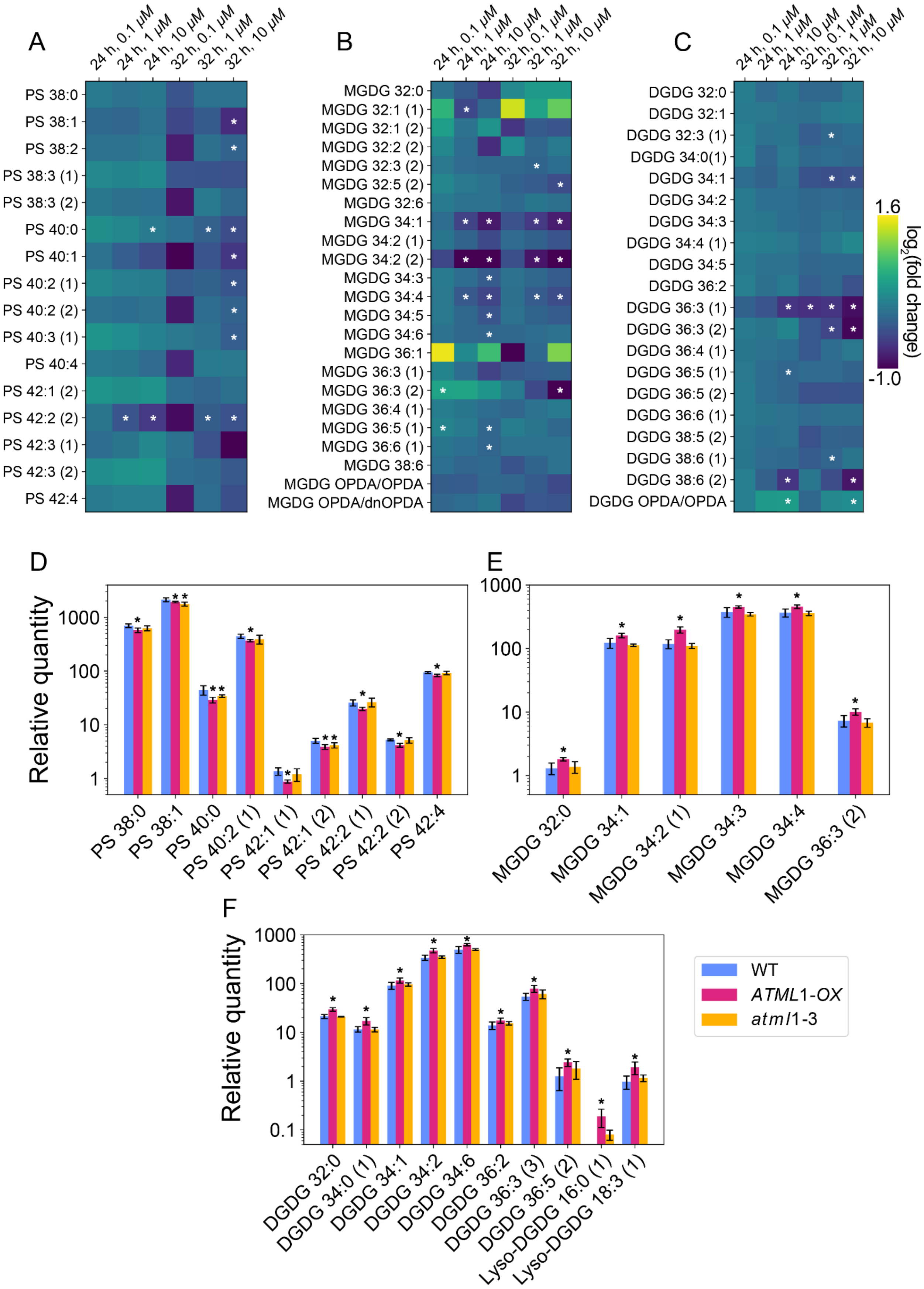

A
24 h, 0.1 μM
24 h, 1 μM
24 h, 10 μM
32 h, 0.1 μM
32 h, 1 μM
32 h, 10 μM
PS 38:0
PS 38:1
PS 38:2
PS 38:3 (1)
PS 38:3 (2)
PS 40:0
PS 40:1
PS 40:2 (1)
PS 40:2 (2)
PS 40:3 (1)
PS 40:4
PS 42:1 (2)
PS 42:2 (2)
PS 42:3 (1)
PS 42:3 (2)
PS 42:4
B
MGDG 32:0
MGDG 32:1 (1)
MGDG 32:1 (2)
MGDG 32:2 (2)
MGDG 32:3 (2)
MGDG 32:5 (2)
MGDG 32:6
MGDG 34:1
MGDG 34:2 (1)
MGDG 34:2 (2)
MGDG 34:3
MGDG 34:4
MGDG 34:5
MGDG 34:6
MGDG 36:1
MGDG 36:3 (1)
MGDG 36:3 (2)
MGDG 36:4 (1)
MGDG 36:5 (1)
MGDG 36:6 (1)
MGDG 38:6
MGDG OPDA/OPDA
MGDG OPDA/dnOPDA
C
DGDG 32:0
DGDG 32:1
DGDG 32:3 (1)
DGDG 34:0(1)
DGDG 34:1
DGDG 34:2
DGDG 34:3
DGDG 34:4 (1)
DGDG 34:5
DGDG 36:2
DGDG 36:3 (1)
DGDG 36:3 (2)
DGDG 36:4 (1)
DGDG 36:5 (1)
DGDG 36:5 (2)
DGDG 36:6 (1)
DGDG 38:5 (2)
DGDG 38:6 (1)
DGDG 38:6 (2)
DGDG OPDA/OPDA
1.6
-1.0
log2(fold change)
D
Relative quantity
1000
100
10
1
PS 38:0
PS 38:1
PS 40:0
PS 40:2 (1)
PS 42:1 (1)
PS 42:1 (2)
PS 42:2 (1)
PS 42:2 (2)
PS 42:4
E
1000
100
10
1
MGDG 32:0
MGDG 34:1
MGDG 34:2 (1)
MGDG 34:3
MGDG 34:4
MGDG 36:3 (2)
F
Relative quantity
1000
100
10
1
0.1
DGDG 32:0
DGDG 34:0 (1)
DGDG 34:1
DGDG 34:2
DGDG 34:6
DGDG 36:2
DGDG 36:3 (3)
DGDG 36:5 (2)
Lyso-DGDG 16:0 (1)
Lyso-DGDG 18:3 (1)
WT
ATML1-OX
atml1-3

**Fig. S2. Lipid content in ATML1 mutants, associated with Fig. 2.**

(**A–C**) Lipid levels including (**A**) phosphatidylserine (PS), (**B**) monogalactosyldiacylglycerol (MGDG), and (**C**) digalactosyldiacylglycerol (DGDG) after ATML1 induction in inflorescence tissue. Yellow and violet indicate the increased and decreased levels of lipids, respectively. * indicate statistically significant changes ($P < 0.05$, unpaired student $t$-test) from wild type. (**D–F**) Steady-state levels of lipids including (**D**) PS, (**E**) MGDG, and (**F**) DGDG in wild type (Col-0), *ATML1-OX,* and *atml1-3* inflorescences. An asterisk indicates the statistically significant difference of $P < 0.05$ determined using an unpaired student $t$-test.

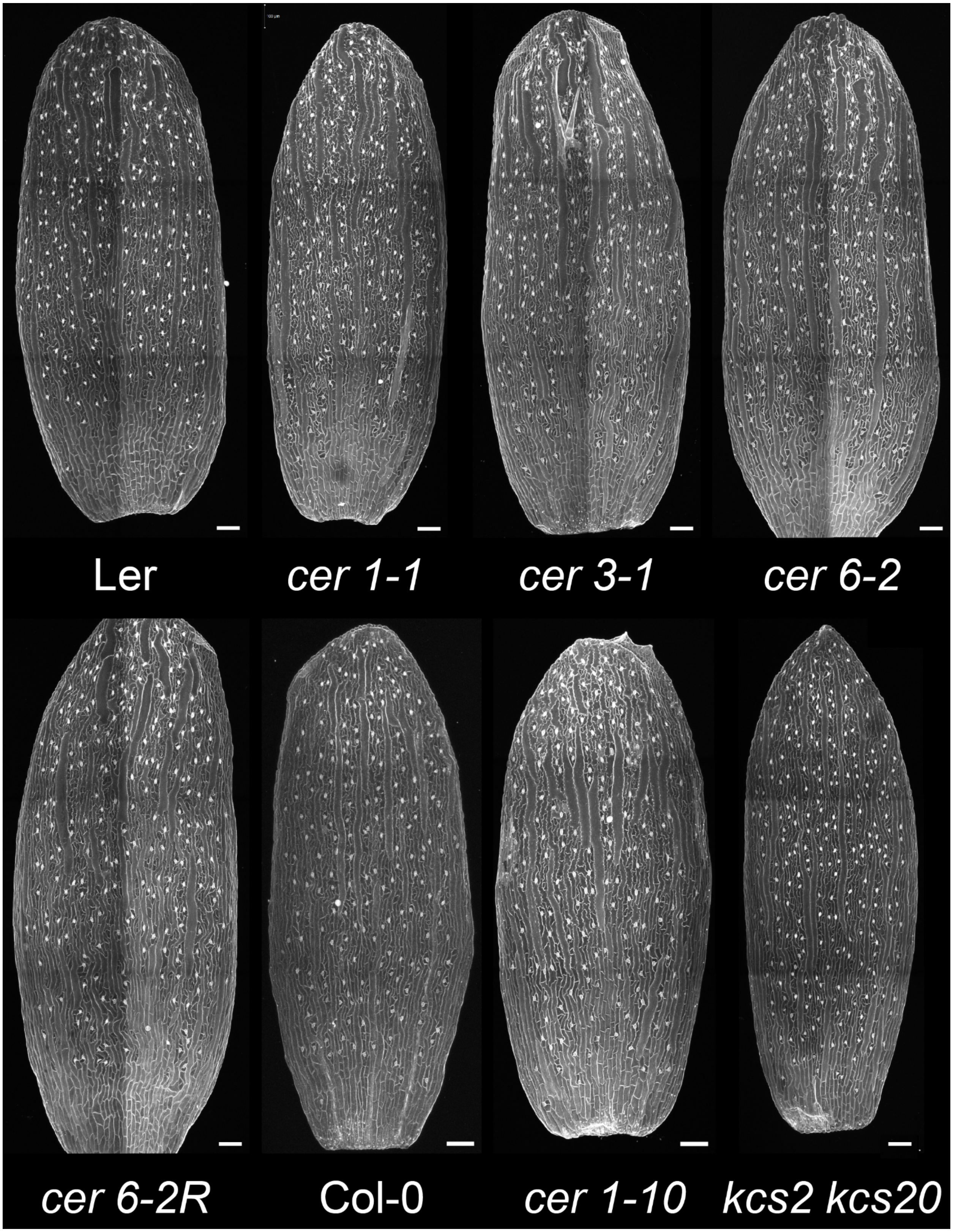

**Fig. S3. Giant cell phenotype in VLCFA mutants, associated with Fig. 3**. Confocal images of sepal epidermis of genotypes *cer 1-1*, *cer 3-1*, *cer 6-2*, *cer 6-2R* and their corresponding L*er* wild type stained with PI. Confocal images of sepal epidermis of genotypes *cer 1-10* and *kcs2 kcs20* and their corresponding Col-0 wild type stained with PI. Scale bars = 100 µm.

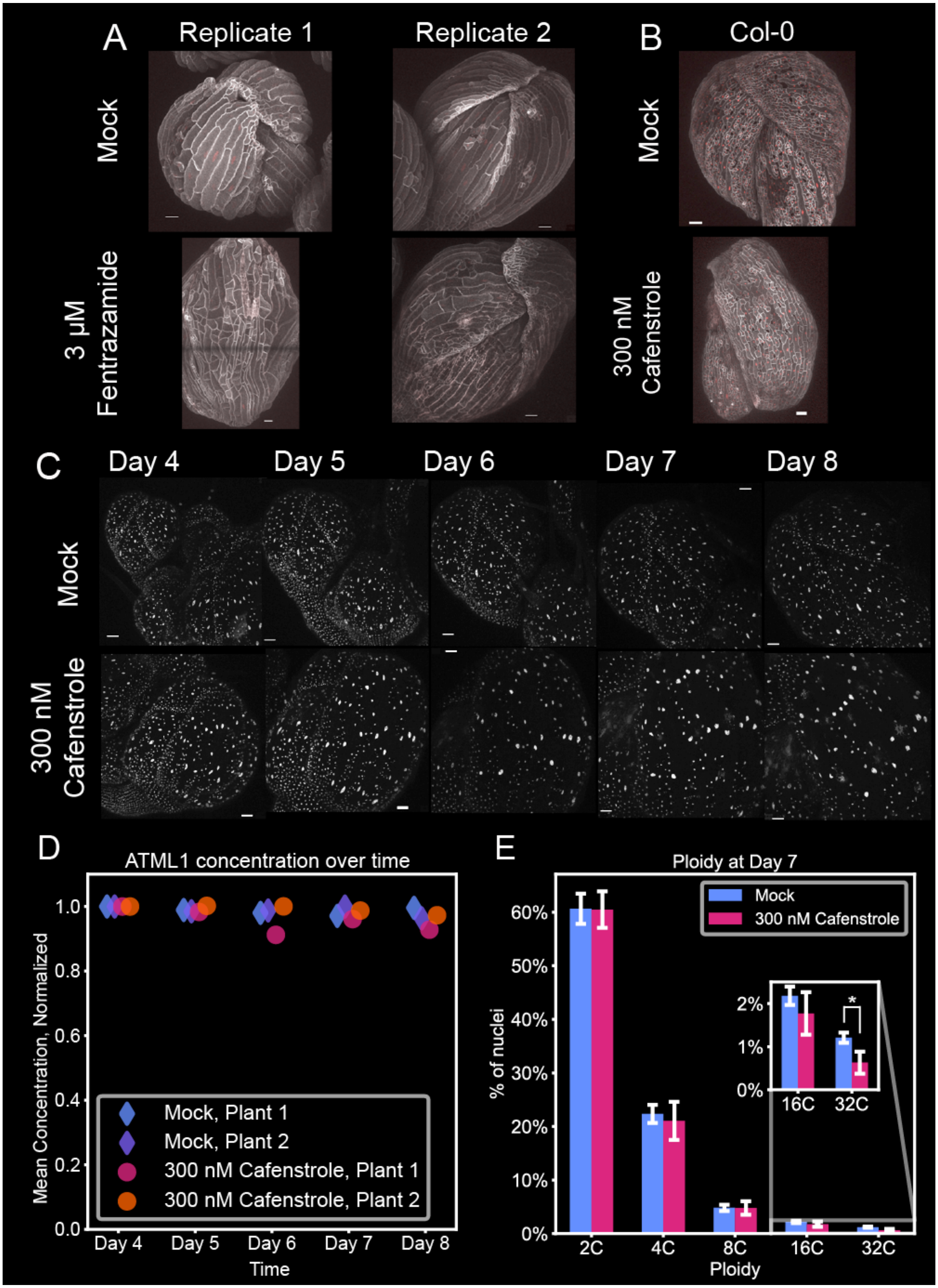
A
Replicate 1
Replicate 2
Mock
3 µM Fentrazamide
B
Col-0
Mock
300 nM Cafenstrole
C
Day 4
Day 5
Day 6
Day 7
Day 8
Mock
300 nM Cafenstrole
D
ATML1 concentration over time
Mean Concentration, Normalized
1.0
0.8
0.6
0.4
0.2
0.0
Mock, Plant 1
Mock, Plant 2
300 nM Cafenstrole, Plant 1
300 nM Cafenstrole, Plant 2
Day 4
Day 5
Day 6
Day 7
Day 8
Time
E
Ploidy at Day 7
% of nuclei
60%
50%
40%
30%
20%
10%
0%
Mock
300 nM Cafenstrole
2%
1%
0%
*
16C
32C
2C
4C
8C
16C
32C
Ploidy

**Fig. S4. VLCFA synthesis is required for giant cell formation, associated with Fig. 4.**
(**A**) *ATML1-OX* flower buds treated with Mock and the VLCFA inhibitor, fentrazamide. Scale bar = 50 µm. (**B**) Flower buds of wild type (Col-0) treated with Mock and 300 nM cafenstrole. Cell outlines are marked by *35S::mCitrine-RCI2A*. Red indicates the *35S:mCherry-H2B* nuclear marker. Scale bar = 50 µm. 2 out of 4 biological replicates shows the phenotype. (**C**) ATML1 protein levels in the inflorescence tissue of *pATML1::mCitrine-ATML1* tracked over 8 days after treatment with Mock and 300 nM cafenstrole. Scale bar = 40 µm. (**D**) Quantification of mean ATML1 nuclear concentration from (C). Nuclear quantification was completed in ilastik using the Pixel and Object modules, and two replicates were used. Values were normalized by the day 4 average intensity. (**E**) Epidermal cell ploidy measurement using flow cytometry in WT inflorescence tissue 7 days after treatment with Mock and 300 nM cafenstrole from Fig. 4B. The $P$-value for the 32C plots is $0.011 < 0.05$ determined using a one-tailed Student's $t$-test. Note that the bars do not add up to 100% because there are background cells that have been discarded.

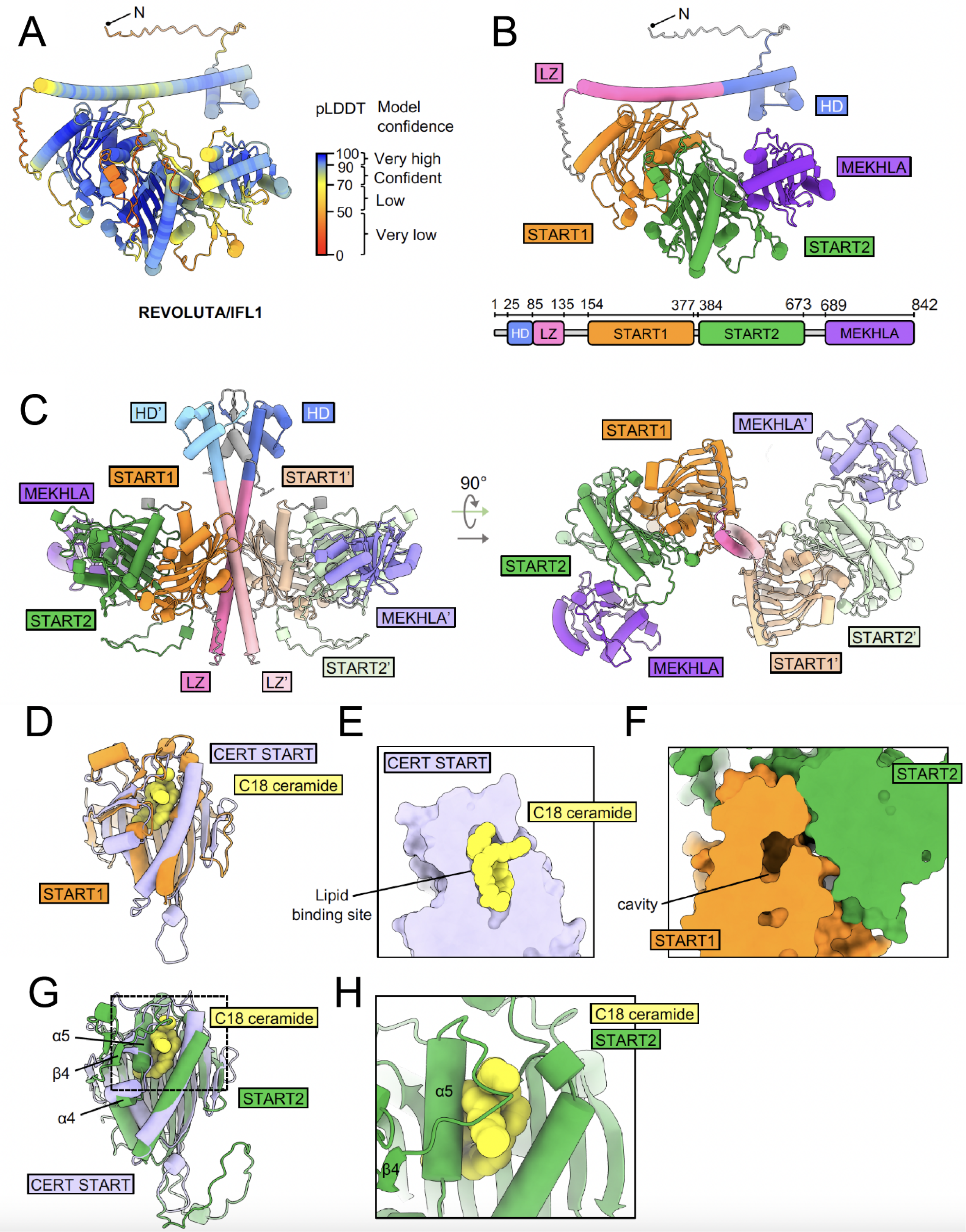
A
N
pLDDT Model confidence
100
90
70
50
0
Very high
Confident
Low
Very low
REVOLUTA/IFL1
B
N
LZ
HD
MEKHLA
START1
START2
1 25 85 135 154 377 384 673 689 842
HD
LZ
START1
START2
MEKHLA
C
HD'
HD
START1
START1'
MEKHLA
START2
MEKHLA'
LZ
LZ'
START2'
90°
START1
MEKHLA'
START2
MEKHLA
START1'
START2'
D
CERT START
C18 ceramide
START1
E
CERT START
C18 ceramide
Lipid binding site
F
START2
cavity
START1
G
C18 ceramide
α5
β4
α4
START2
CERT START
H
C18 ceramide
START2
α5
β4

**Fig. S5. Analysis of predicted structures of the HD-ZIP III protein REVOLUTA/IFL1 also predicts two START domains, but different dimerization, associated with Fig. 5.** (**A**) AlphaFold2 predicted the structure of REVOLUTA/IFL1 (REV) colored by pLDDT score to show model confidence. (**B**) AlphaFold2 predicted structure of REV colored according to domain: homeodomain (HD), leucine zipper (LZ), START domain 1 (previously identified), START domain 2 (newly predicted), MEKHLA domain. Schematic showing linear N- to C- terminal domain arrangement of REV. Note that two START domains are predicted in REV, similar to ATML1. Unlike HD-ZIP IV proteins, REV and other HD-ZIP III proteins also contain a C-terminal MEKHLA domain. (**C**) Model of dimeric REV predicted using AlphaFoldMultimer. Note that homodimerization is predicted only through the leucine zipper and not the START2 domains. Models of REV START1 and START2 domains, (**D**) and (**G**) respectively, aligned to the crystal structure of the START domain from Human ceramide transfer protein (CERT) in complex with C18-ceramide (PDB 2E3Q). (E) Cross-section of the surface model of the crystal structure of the START domain from CERT in complex with C18-ceramide. (**F**) A cavity in the model of the REV START1 domain that is expected to bind lipids is marked in the cross-section of the surface model. The close-up view in (**H**) shows helix 5 (α5) in the REV START1 predicted model clashes with C18-ceramide docked based on the crystal structure of CERT in complex with C18-ceramide.

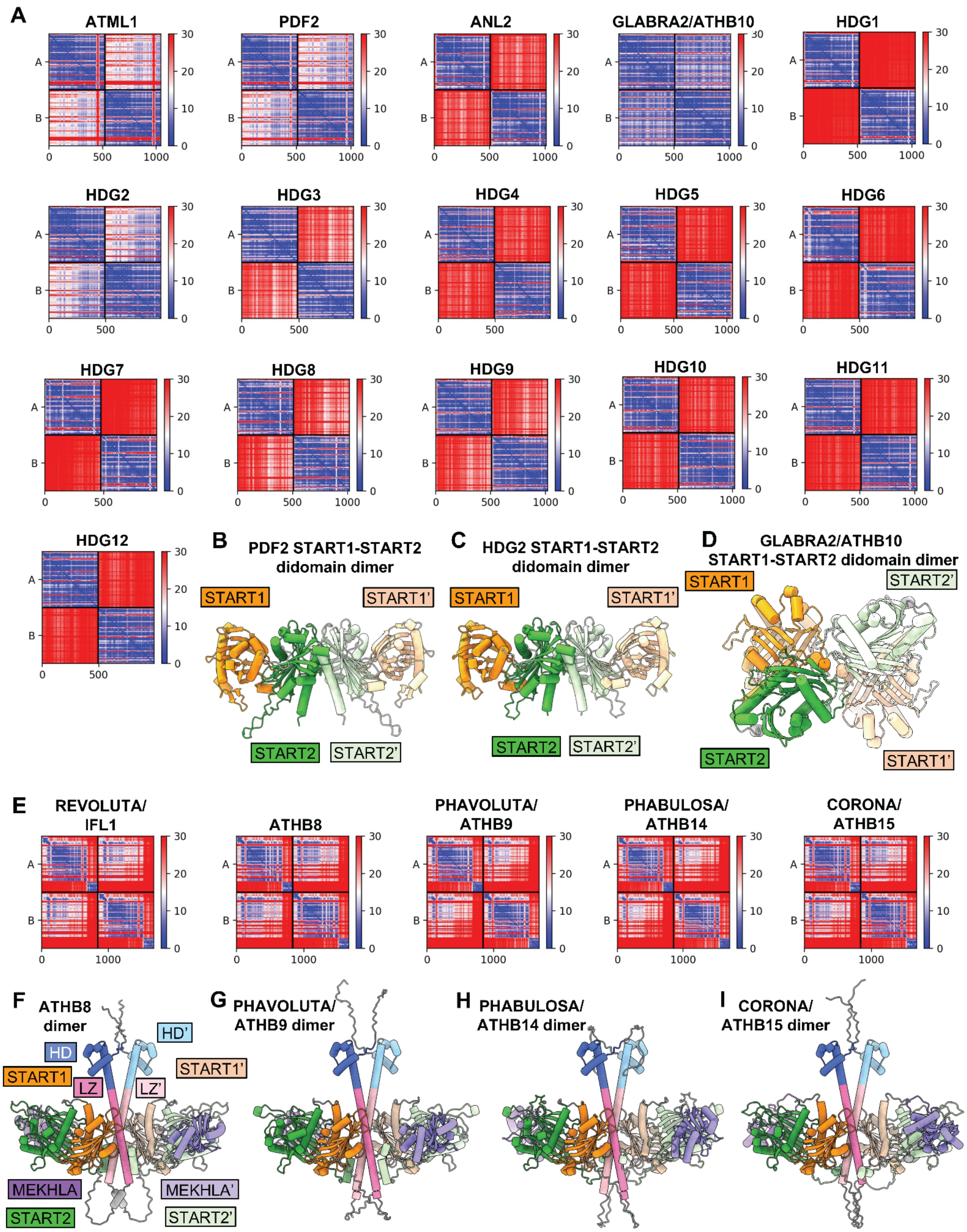

A
ATML1
PDF2
ANL2
GLABRA2/ATHB10
HDG1
HDG2
HDG3
HDG4
HDG5
HDG6
HDG7
HDG8
HDG9
HDG10
HDG11
HDG12
B
PDF2 START1-START2 didomain dimer
START1
START1'
START2
START2'
C
HDG2 START1-START2 didomain dimer
START1
START1'
START2
START2'
D
GLABRA2/ATHB10 START1-START2 didomain dimer
START1
START2'
START2
START1'
E
REVOLUTA/ IFL1
ATHB8
PHAVOLUTA/ ATHB9
PHABULOSA/ ATHB14
CORONA/ ATHB15
F
ATHB8 dimer
HD
HD'
START1
START1'
LZ
LZ'
MEKHLA
MEKHLA'
START2
START2'
G
PHAVOLUTA/ ATHB9 dimer
H
PHABULOSA/ ATHB14 dimer
I
CORONA/ ATHB15 dimer

**Fig. S6. HD-ZIP III and IV members are predicted to have different homodimerization structures, associated with Fig. 5.** (**A**) Predicted Aligned Error (PAE) plots for predicted dimeric structures of START1–START2 didomains of HD-ZIP IV members with the highest Interface predicted TM-score (ipTM). (**B**), (**C**), and (**D**) Predicted dimer models of START1–START2 didomains of PDF2, HDG2, and GLABRA2/ATHB10, respectively. (**E**) Predicted Aligned Error (PAE) plots for predicted dimeric structures of HD-ZIP Class III members with the highest Interface predicted TM-score (ipTM). PAE (in Å) between pairs of amino acids (x,y) is colored from blue (0Å) to red (30Å) in the plots in panels (**A**) and (**E**). (**F**), (**G**), (**H**), and (**I**) Predicted dimer models of ATHB8, PHAVOLUTA/ATHB9, PHABULOSA/ATHB14, and CORONA/ATHB15, respectively.

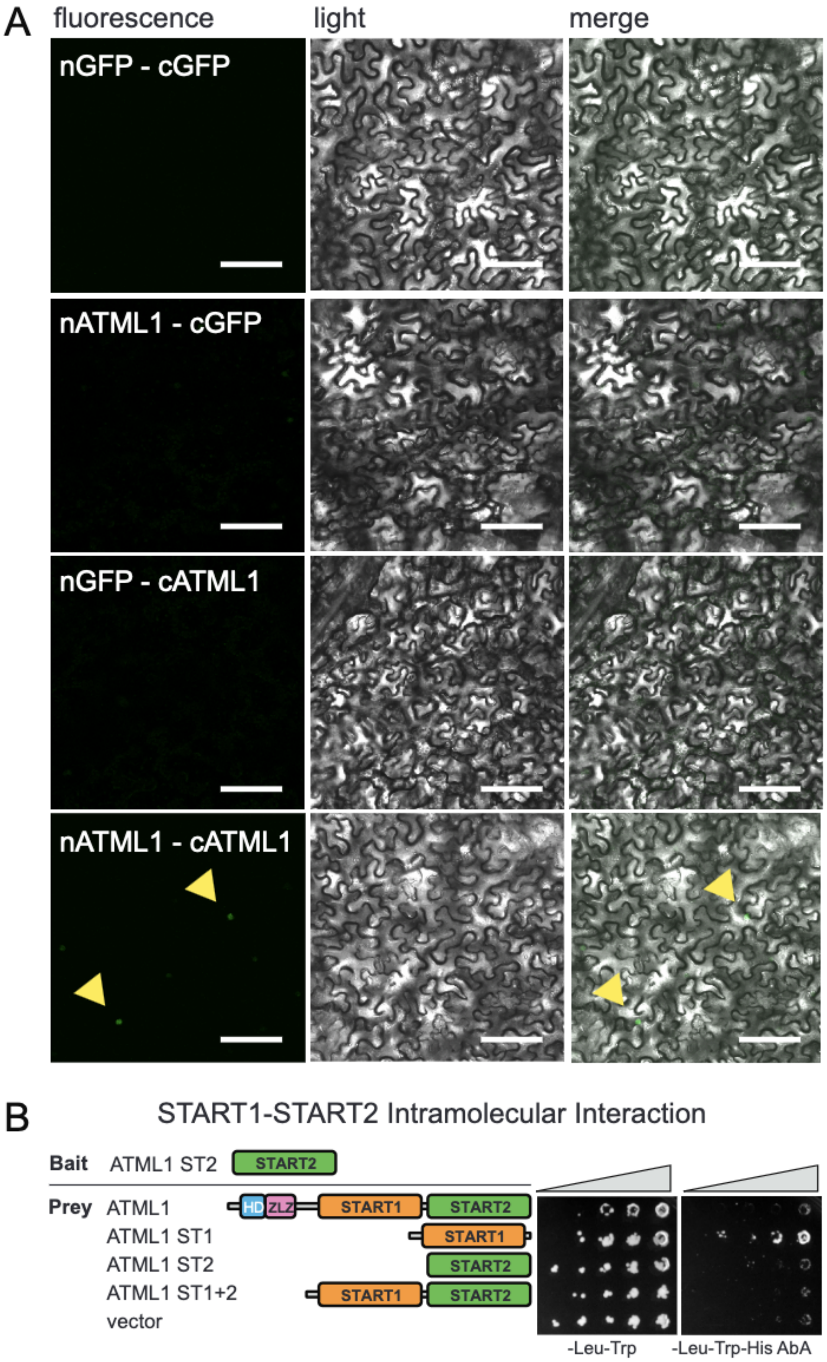

**Fig. S7. Y2H and BiFC assays reveal interaction properties of ATML1, associated with Fig. 6A–C.** (**A**) Bimolecular fluorescence complementation (BiFC) assays indicate homodimerization of ATML1. Leaf abaxial epidermal cells from *Nicotiana benthamiana* plants were transiently transformed with constructs expressing split GFP segments for BiFC assays. Representative matching GFP fluorescence (left), white light (middle), and merge (right) images are shown for vector–vector (nGFP–cGFP) control, nGFP:ATML1 with cGFP vector control, and cGFP:ATML1 with nGFP vector control. Nuclear expression, as indicated by arrowheads, is only observed when nGFP:ATML1 is expressed together with cGFP:ATML1. Scale bar = 50 µm. (**B**) START1 and START2 participate in intramolecular interaction. A schematic of full-length ATML1 protein in comparison with START1 (ST1) and START2 (ST2) segments is illustrated. . Diploid yeast was transformed with bait and prey constructs to test for interaction. ATML1 START2 (ST2) was expressed as bait in combination with full-length ATML1, START1, START2, START1+2 didomain, or vector alone. Growth was assayed on permissive medium (-Leu-Trp) and selective medium (-Leu-Trp-His) with Aureobasidin A (AbA). The interaction of START2 with START1 resulted in a signal greater than that from the vector alone.

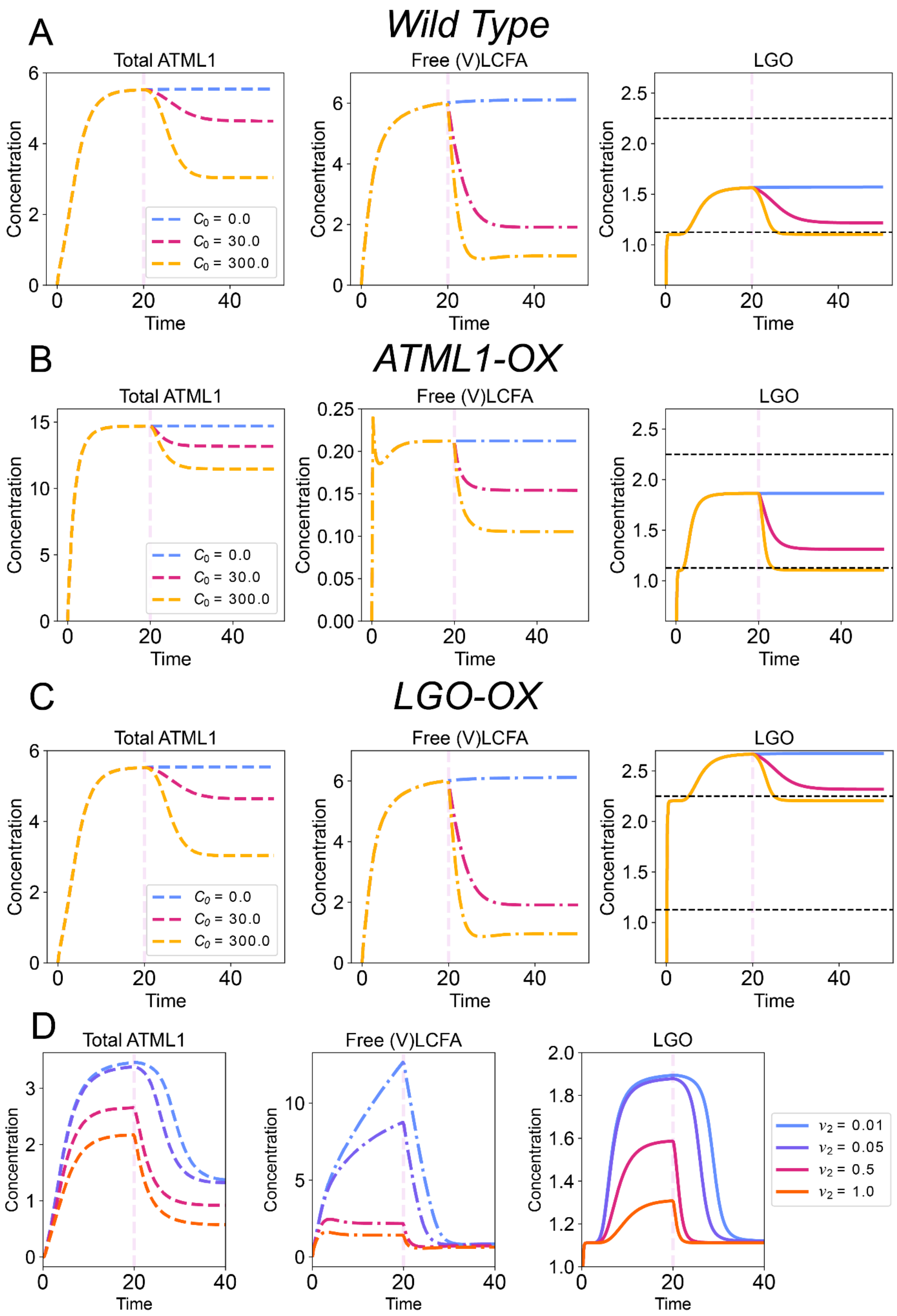

A
Wild Type
Total ATML1
Free (V)LCFA
LGO
Concentration
Time
$C_0$ = 0.0
$C_0$ = 30.0
$C_0$ = 300.0
B
ATML1-OX
Total ATML1
Free (V)LCFA
LGO
Concentration
Time
$C_0$ = 0.0
$C_0$ = 30.0
$C_0$ = 300.0
C
LGO-OX
Total ATML1
Free (V)LCFA
LGO
Concentration
Time
$C_0$ = 0.0
$C_0$ = 30.0
$C_0$ = 300.0
D
Total ATML1
Free (V)LCFA
LGO
Concentration
Time
$v_2$ = 0.01
$v_2$ = 0.05
$v_2$ = 0.5
$v_2$ = 1.0

**Fig. S8. Deterministic simulations of the ATML1–(V)LCFA model show *ATML1-OX* – but not *LGO-OX* – will be affected by cafenstrole, associated with Fig. 8.** Simulations of a single cell using the Eqs. 4–7 in the Mathematical Modeling section of the Methods. Because plants carried the fluorescent marker *mcitrine-AMTL1*, which is a proxy for experimentally measuring total ATML1 levels, the total ATML1 concentration level is plotted, i.e., $[ATML1]_i + 2\times[Dimerized\ ATML1]_i$. Pink dashed vertical lines indicate the time of cafenstrole application (blue, no cafenstrole, $C_0 = 0$; magenta, $C_0 = 30$; and yellow, $C_0 = 300$) in the simulation at time $t = 20$. In the LGO plot, the horizontal dashed lines represent $H_{T_1}$, i.e., the threshold for endoreduplication, and $H_{T_2}$, i.e., the threshold for division of an endoreduplicated cell (top and bottom lines, respectively). Dilution effects are accounted for by rescaling the degradation terms of each concentration variable by the average cell area growth rate, 2/25 (see Methods section). Trajectories of total ATML1, free (V)LCFA that are available to bind to ATML1, and LGO in a simulation of (**A**) wild type (WT), (**B**) *ATML1-OX*, and (**C**) *LGO-OX* with three levels of cafenstrole treatment. (**D**) Trajectories of total ATML1, (V)LCFA, and LGO concentrations in WT with various levels of (V)LCFA degradation rates ($\nu_2$) after $C_0 = 300$ are implemented at $t = 20$. Note that LGO declines much slower with lower degradation rates of (V)LCFA.

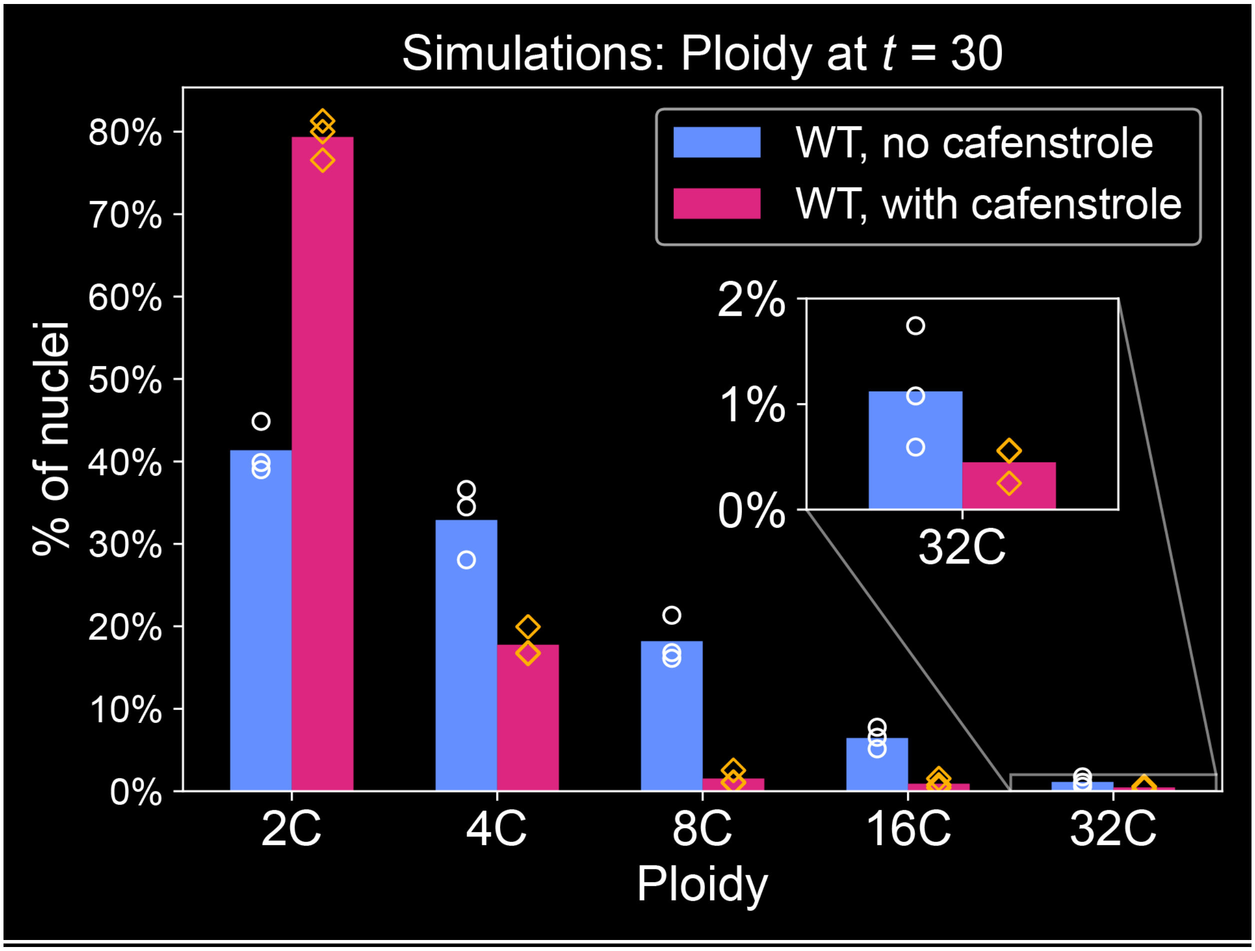

**Fig. S9. The model predicts a large reduction in the number of giant cells after cafenstrole treatment.** Ploidy levels at the end of simulations ($t$ = 30) for wild type with and without cafenstrole. Each bar represents the mean of three simulations. Each simulation is represented by a circle or diamond on the respective bars. The inset panel shows a decrease in the number of 32C cells; strikingly, it shows that not all giant cells divide once cafenstrole is added, which captures the dynamics observed in the experiments (see Figs. 4B and S4E).

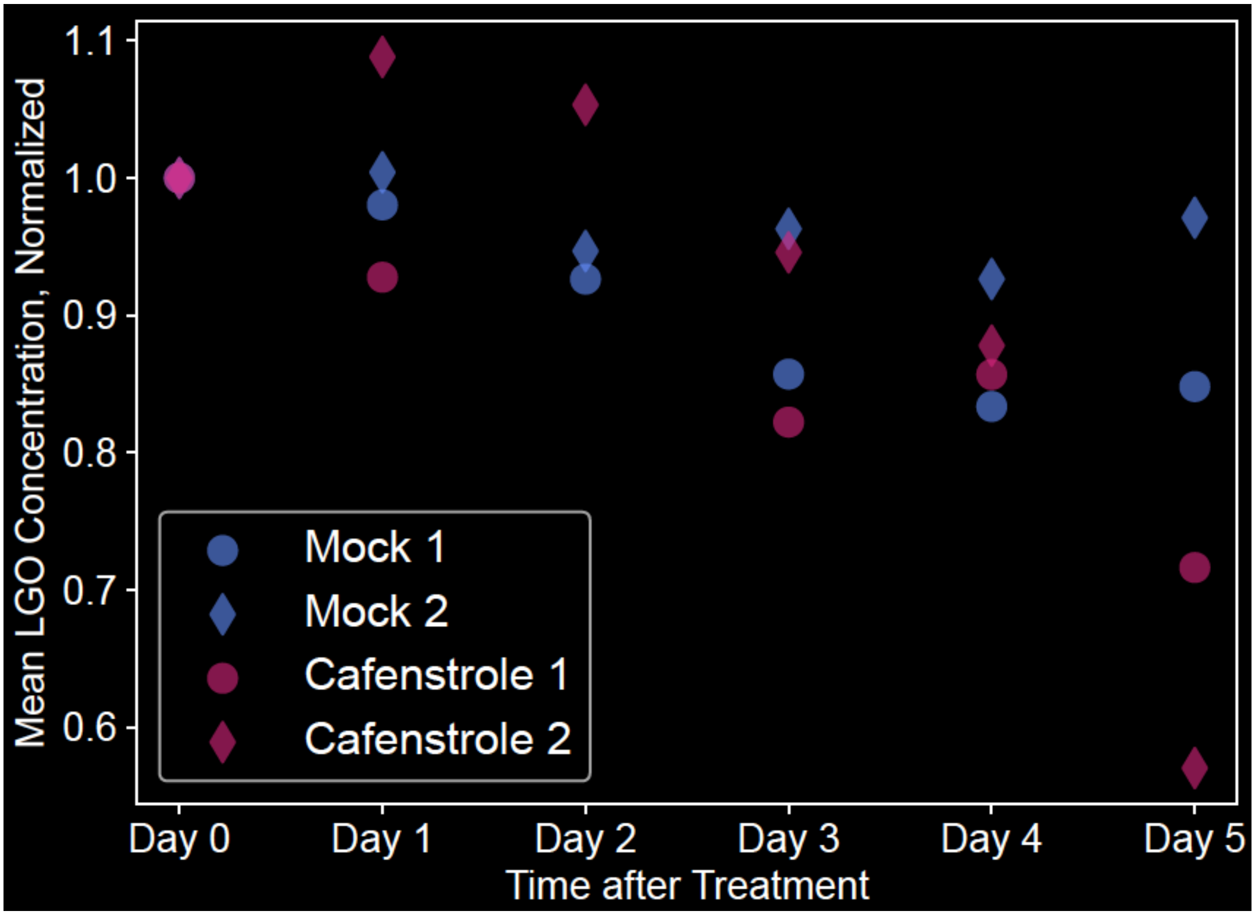

**Fig. S10. Levels of the cyclin-dependent kinase inhibitor, *LGO*, decrease with cafenstrole treatment, associated with Figure 7F.** Quantification of *LGO* levels using ilastik from the live imaging data shown in Figure 7F. All concentrations were normalized by the day 0 concentration to compare the decrease in *LGO* concentrations. Two experimental replicates were used for this quantification. See the Materials and Methods section (specifically the subsection "Quantification of ATML1 and LGO reporters") and Fig. S12 for the full image analysis protocol. Note that days 0, 1, and 2 are included in the quantification whereas they are excluded from Fig. 7F for the sake of space.

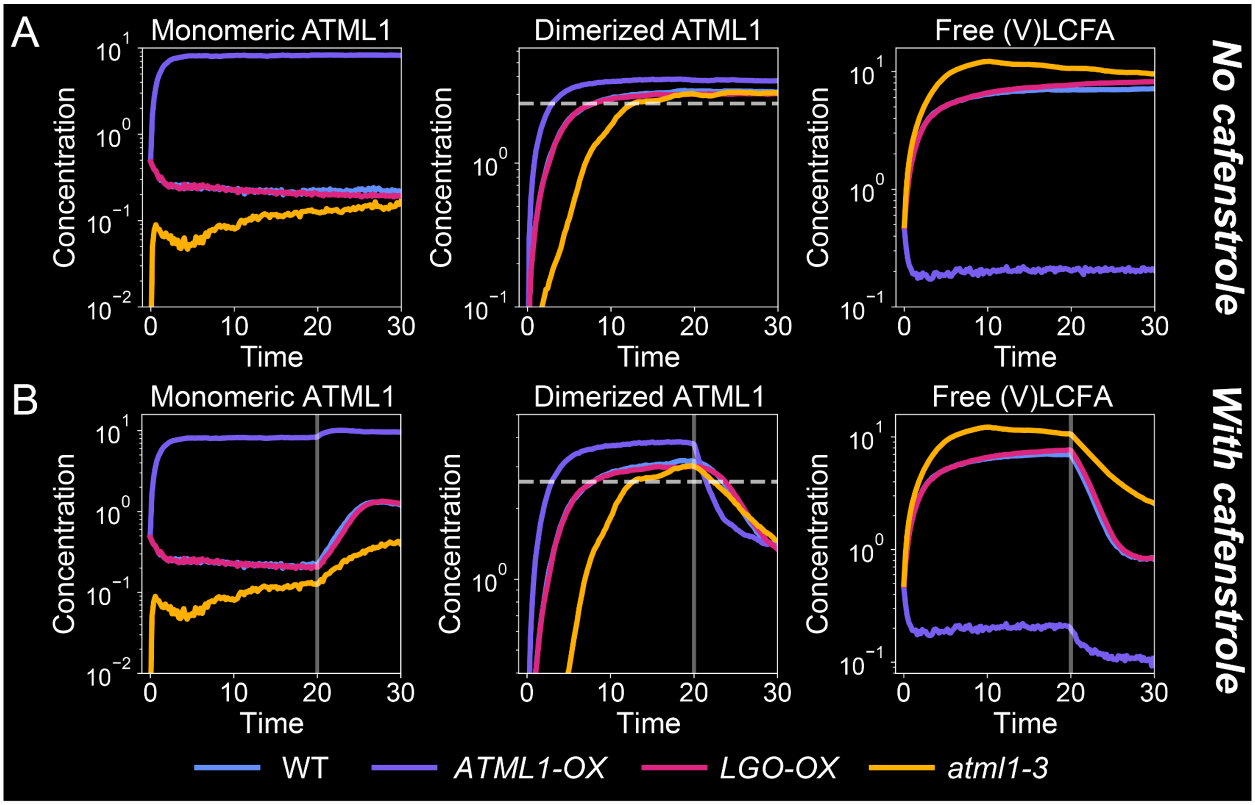

**Fig. S11.** ***LGO*** **expression levels are regulated by dimerized ATML1, associated with Fig. 8B.** Average concentrations of monomeric AMTL1, dimerized ATML1, and free (V)LCFAs for different simulated genotypes, including wild type (WT), *ATML1-OX*, *LGO-OX*, and *atml1-3* (**A**) without cafenstrole and (**B**) with cafenstrole added at $t = 20$ (see Materials and Methods for further details). In the *atml1-3* mutant, we assume some remaining "ATML1" activity due to the paralog *PDF2*. For WT, the basal production rate of ATML1 is $\beta_A = 1.1$, whereas for the *atml1-3* mutant, the basal production term is $\beta_A = 0.2$ (see Table S2). Note that in *ATML1-OX*, the average dimerized ATML1 concentration quickly reaches the $K_l$ threshold, i.e., the amount of dimerized ATML1 needed to induce a significant production of LGO. *ATML1-OX* is followed by WT and *LGO-OX* simultaneously in reaching this threshold, with *atml1-3* achieving this threshold last. This agrees with the timing at which the second rise of LGO occurs in these simulated genotypes (Fig. 8B). When cafenstrole is applied in (B), *ATML1-OX* has such a low level of free (V)LCFA that the decrease in Dimerized ATML1 is quicker than in WT, resulting in the more rapid decrease in LGO levels seen in Fig. 8B. The y-axis for the dimerized ATML1 plot in (B) is different from (A) to emphasize this point. Note that (V)LCFA in this model refers to the fatty acids that are available to bind to ATML1.

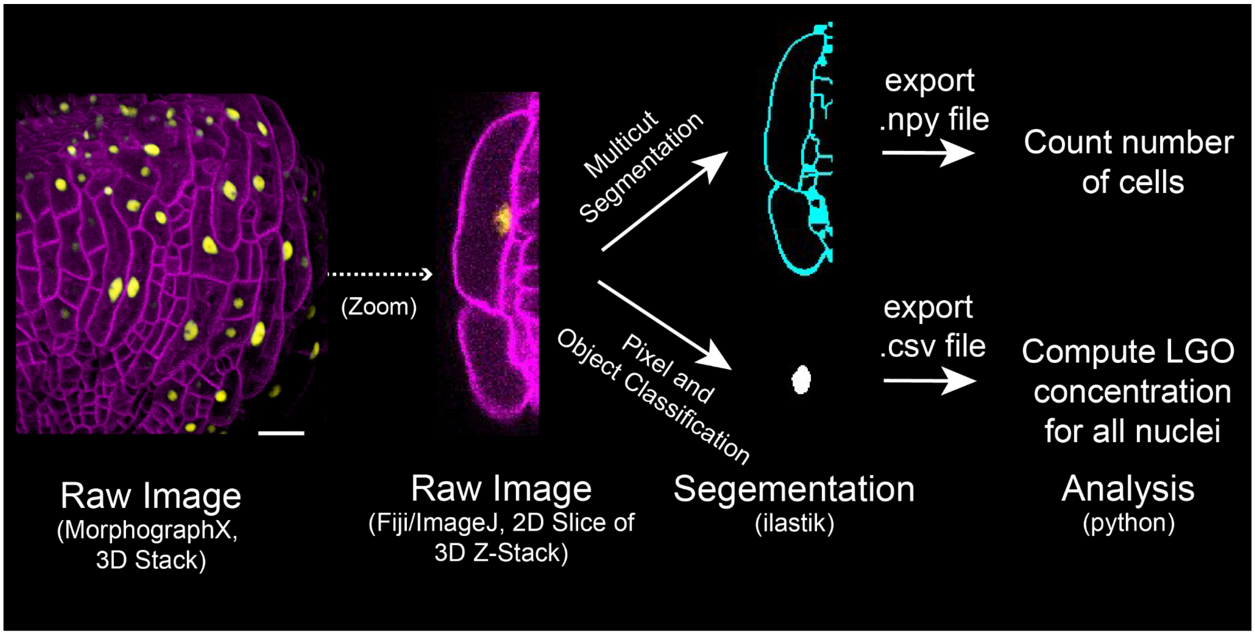

**Fig. S12: Image quantification schematic for Figs. 7F and S10.** A simple schematic of the image analysis pipeline to quantify the change in the average *pLGO::3×Venus-N7* fluorescence intensity per nuclei. We consider this as a proxy for inferring the average LGO mRNA concentration per nuclei. This method is explained in full in the Materials and Methods section (see the subsection "Quantification of ATML1 and LGO reporters"). In brief, we use ilastik to perform both the segmentation of the nuclei and the plasma membrane. Exporting the membrane segmentation as a .npy file allows us to count the number of cells segmented. Not all nuclei will express the *LGO* reporter. Therefore, the difference in the number of cells counted and the number of nuclei segmented gives us the number of cells that are not expressing the the *LGO* reporter. To quantify the average *LGO* reporter concentration, we consider these cells to have 0 as their concentration. Scale bar = 20 µm.

| HD ZIP Class III | | |
|---|---|---|
| | | iPTM scores for predicted dimeric models |
| 1 | REVOLUTA/IFL1 | 0.642, 0.538, 0.522 |
| 2 | ATHB8 | 0.661, 0.648, 0.642 |
| 3 | PHAVOLUTA/ATHB9 | 0.599, 0.59, 0.586 |
| 4 | PHABULOSA/ATHB1 4 | 0.579, 0.567, 0.529 |
| 5 | CORONA/ATHB15 | 0.635, 0.607, 0.553 |
| HD ZIP Class IV | | |

| | | iPTM scores for predicted START1-START2 didomain dimeric models |
|---|---|---|
| 1 | ATML1 | 0.593, 0.477, 0.155 |
| 2 | PDF2 | 0.648, 0.477, 0.243 |
| 3 | ANL2 | 0.241, 0.151, 0.135 |
| 4 | GLABRA2/ATHB10 | 0.807, 0.729, 0.702 |
| 5 | HDG1 | 0.149, 0.147, 0.127 |
| 6 | HDG2 | 0.675, 0.533, 0.135 |
| 7 | HDG3 | 0.312, 0.259, 0.119 |
| 8 | HDG4 | 0.235, 0.145, 0.118 |
| 9 | HDG5 | 0.204, 0.137, 0.116 |
| 10 | HDG6 | 0.188, 0.154, 0.134 |
| 11 | HDG7 | 0.171, 0.153, 0.131 |
| 12 | HDG8 | 0.285, 0.126, 0.191 |
| 13 | HDG9 | 0.278, 0.162, 0.126 |
| 14 | HDG10 | 0.215, 0.152, 0.15 |
| 15 | HDG11 | 0.191, 0.126, 0.107 |
| 15 | HDG12 | 0.19, 0.125, 0.11 |

**Table S1. ipTM scores for predicted dimeric models of HD-ZIP III and IV members.** Proteins for which the highest Interface predicted TM-score (ipTM) is greater than 0.5 are marked in green.

| Parameter | Description | Values |
|---|---|---|
| $C_0$ | (V)LCFA-synthesis inhibitor Concentration | 0.0-300.0 |
| $t_1$ | Time when LCFA-synthesis inhibitor is added | 20.0 |
| $\beta_A$ | ATML1 basal production rate | 1.1 |
| $\beta_L$ | LGO basal production rate | 10.0 |
| $\beta_V$ | (V)LCFA basal production rate | 1.1 |
| $\beta_a$ | Dimerized ATML1 maximal production rate for ATML1 | 2.0 |
| $\beta_v$ | Dimerized ATML1 maximal production rate for (V)LCFA | 1.4 |
| $\beta_l$ | Dimerized ATML1 maximal transcription rate for LGO | 8.0 |
| $v_0$ | ATML1 degradation rate | 1.0 |
| $v_1$ | Dimerized ATML1 degradation rate | 0.4 |
| $v_2$ | (V)LCFA degradation rate | 0.03 |

| Parameter | Description | Values |
|---|---|---|
| $v_3$ | LGO degradation rate | 9.0 |
| $K_a$ | Dissociation constant for dimerized ATML1 mediated ATML1 induction | 1.1 |
| $K_v$ | Dissociation constant for dimerized ATML1 mediated (V)LCFA induction | 1.0 |
| $K_l$ | Dissociation constant for dimerized ATML1 mediated LGO induction | 2.6 |
| $K_c$ | Dissociation constant for cafenstrole to inhibit (V)LCFA production | 30.0 |
| $K_{av}+$ | ATML1 dimerization rate | 15.0 |
| $K_{av}-$ | Dimerized ATML1 dissociation rate | 12.0 |
| $n_a$ | Hill coefficient for dimerized ATML1 mediated ATML1 induction | 2.0 |
| $n_v$ | Hill coefficient for dimerized ATML1 mediated (V)LCFA induction | 1.0 |
| $n_l$ | Hill coefficient for dimerized ATML1 mediated LGO induction | 7.0 |
| $n_c$ | Hill coefficient for (V)LCFA-Synthesis mediated (V)LCFA inhibition | 5.0 |
| $P_C$ | Timer basal production rate | 0.45 |
| $H_C$ | Timer threshold for resetting | 3.0 |
| $H_S$ | Timer threshold for synthesis | 2.5 |
| $H_{T_1}$ | LGO high threshold for endoreduplication | 2.25 |
| $H_{T_2}$ | LGO low threshold for giant-cell fate reversal | 1.125 |
| $E_0$ | Characteristic effective volume | 15 |
| | Exponential radial growth rate | 0.018 |
| | Exponential vertical growth rate | 0.041 |

**Table S2. Parameter values used for computational simulations.** These parameter values are used in Figs. 7–9, Figs. S8 and S11, and Movies S1–6. See Eqs. 4–11. Time and concentration units are arbitrary. The following are the exceptions: *ATML1-OX*: $\beta_A = 10$; *LGO-OX*: $\beta_L = 20$; *atml1-3*: $\beta_A = 0.2$.

**Movies S1–S6:** Simulations of a growing and dividing tissue using the gene regulatory network shown in Fig. 7A (see Methods section). These videos correspond to simulations shown in Fig. 7C–D and Fig. 9A–D. The movies show the development of a tissue in the following phenotypes: (**1**) wild type (WT), (**2**) WT with cafenstrole, (**3**) *ATML1-OX*, (**4**) *ATML1-OX* with cafenstrole, (**5**) *LGO-OX*, and (**6**) *LGO-OX* with cafenstrole.

**Data S1 (uploaded as Excel): Data showing the** Gene ontology (GO) analysis of the up- and down-regulated genes when overexpressing ATML1.

**Data S2 (uploaded as a zip file):** Gene correlation analysis, time-series data.

**Data S3 (uploaded as a zip file): Hill plots showing gene correlation analysis with respect to ATML1.**

**Data S4 (uploaded as Excel): Data showing the (V)LCFA and (V)LCFA-containing lipids levels in inducible ATML1 inflorescence tissue using mass spectrometry**.

**Data S5 (uploaded as Excel): Data showing the (V)LCFA and (V)LCFA-containing lipids levels in WT, *atml1-3*, and *ATML1-OX*.**
