## Supplementary figures and images for "The transcription factor ATML1 maintains giant cell identity by inducing synthesis of its own long-chain fatty acid-containing ligands"

### AAO3.pdf

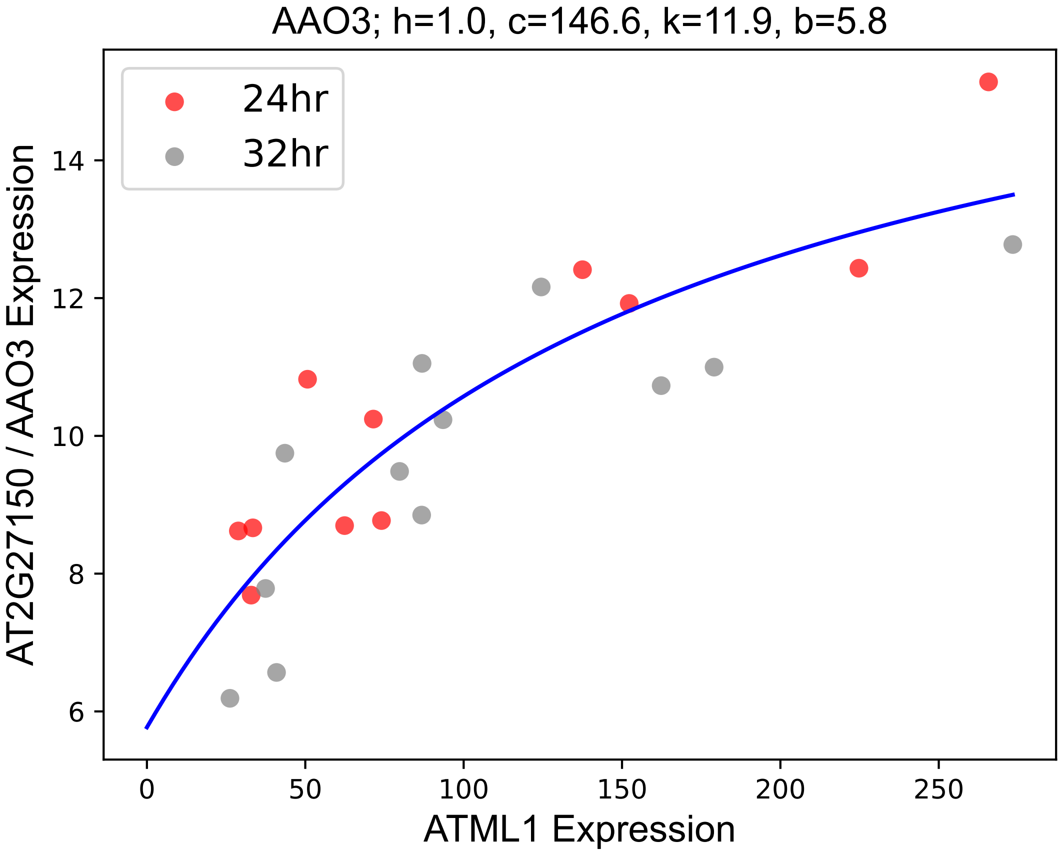

### AIM1.pdf

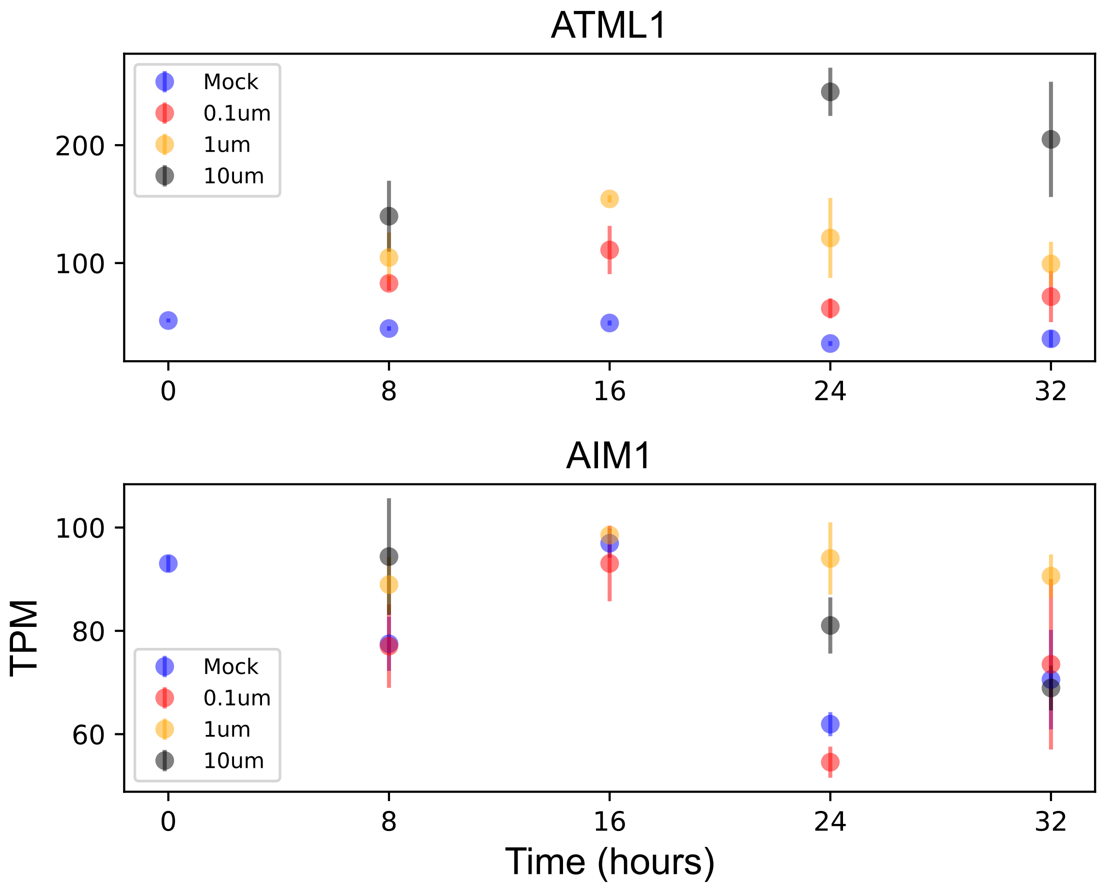

### ASG4.pdf

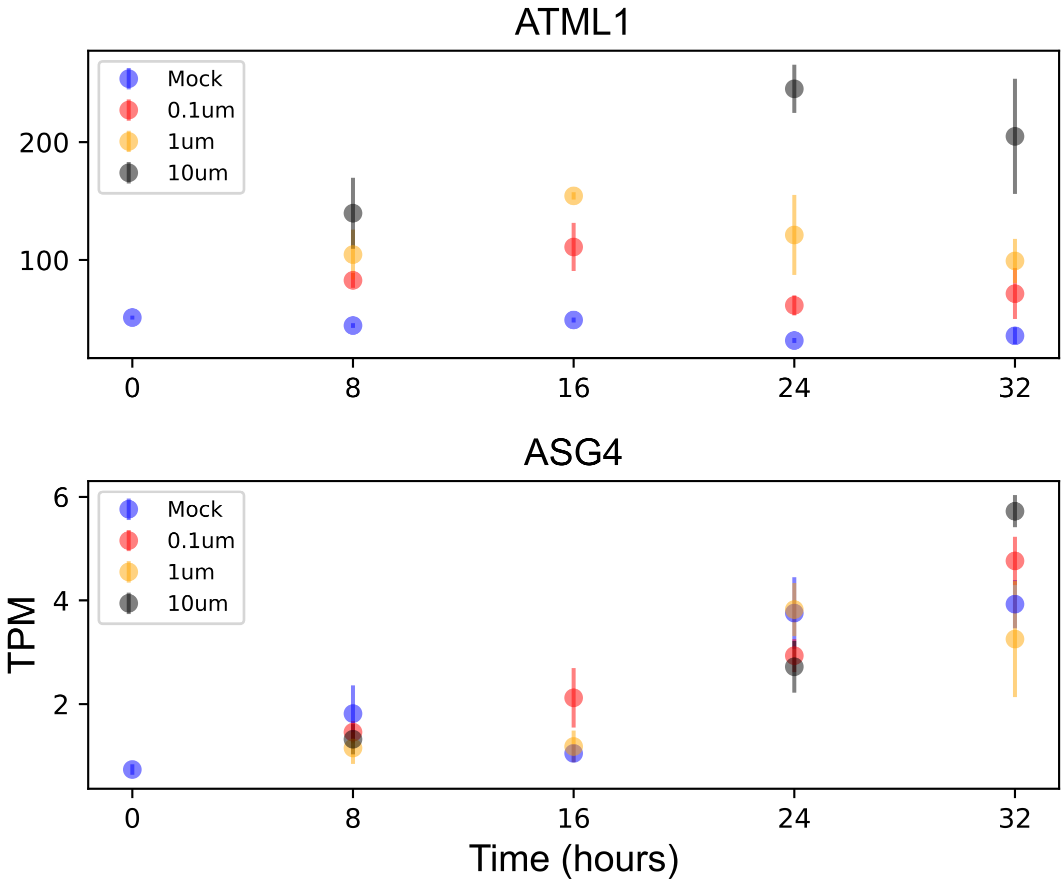

### AT1G10310.pdf

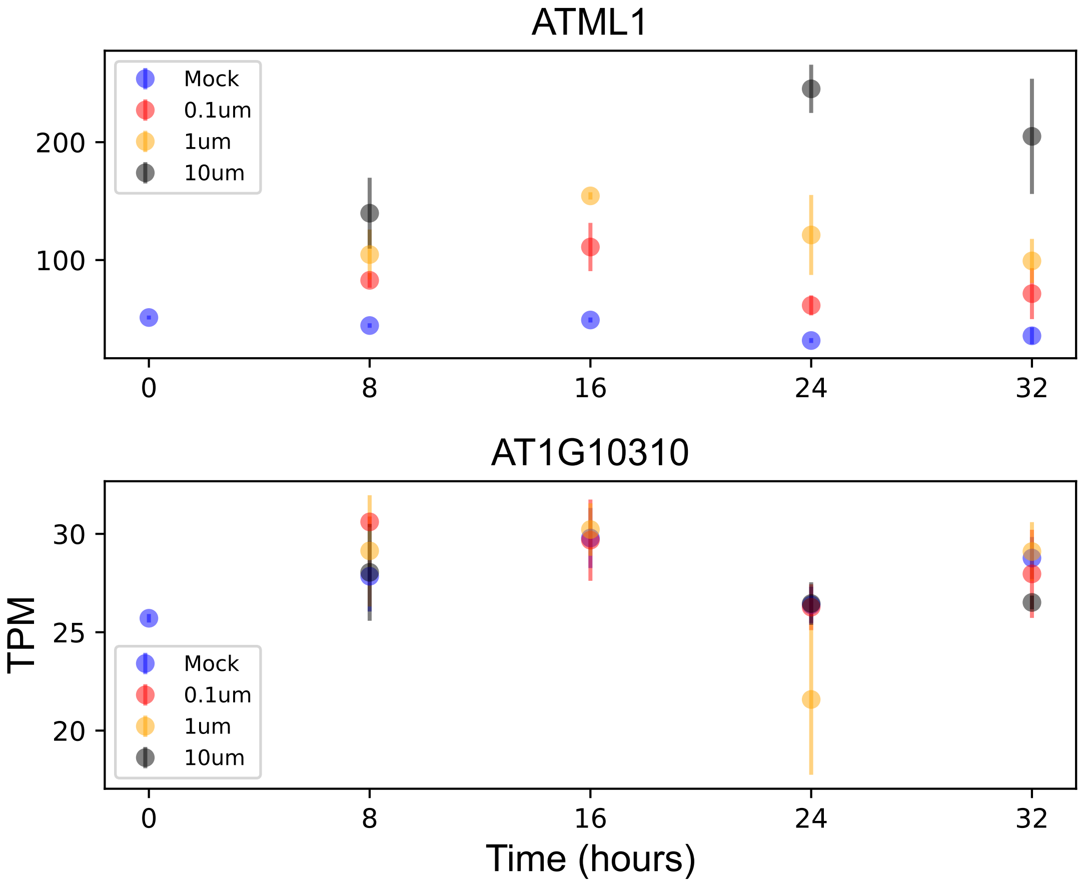

### AT1G19020.pdf

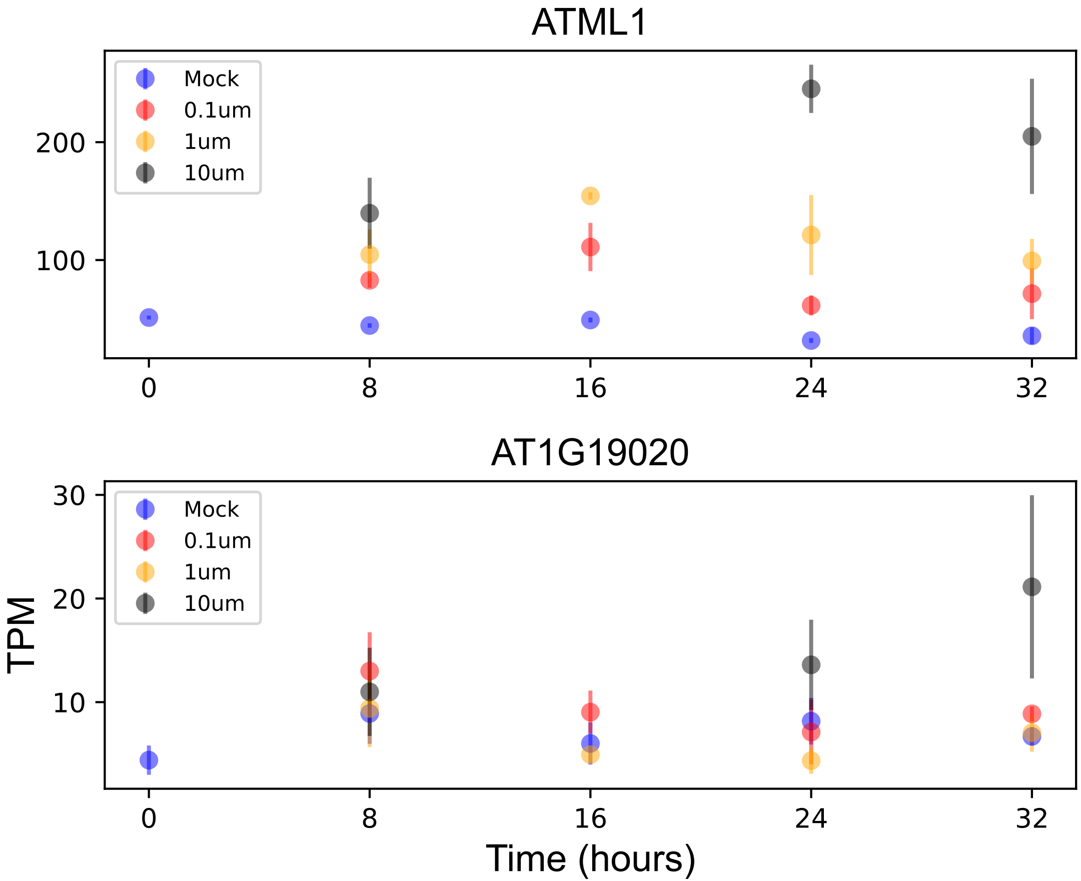

### AT1G24010.pdf

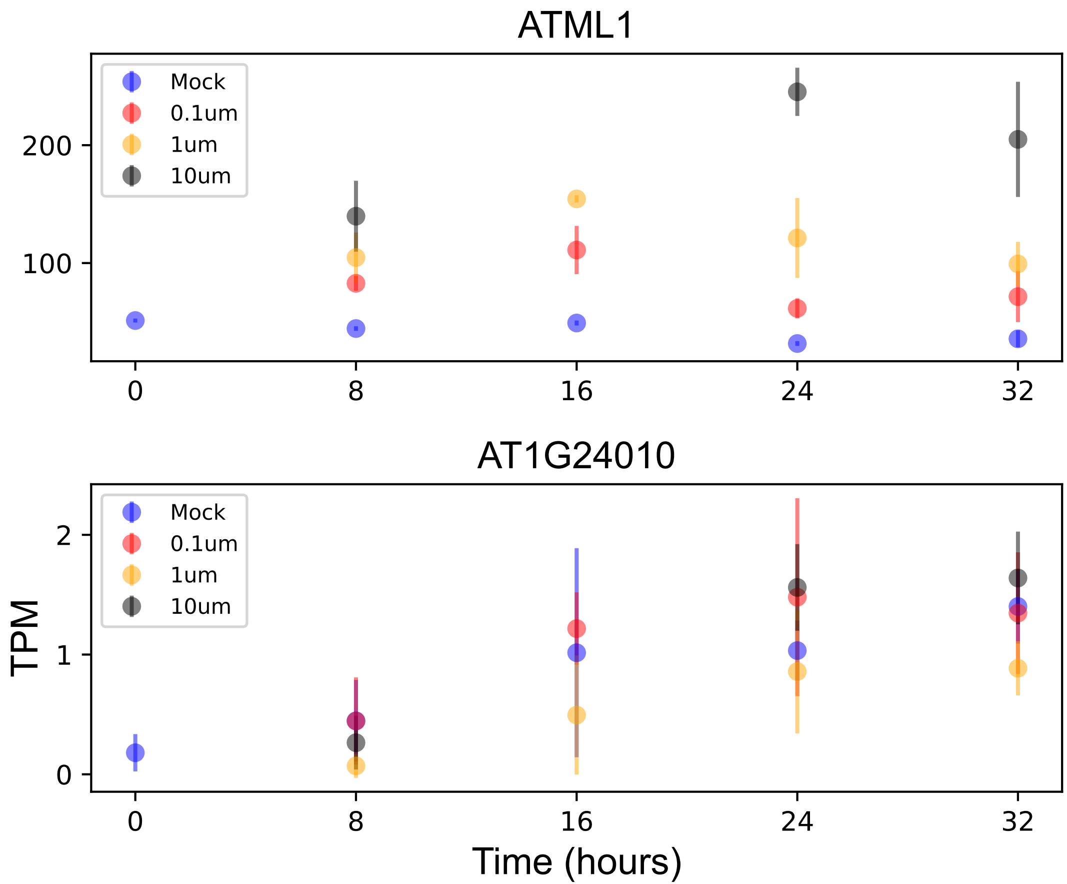

### AT1G27170.pdf

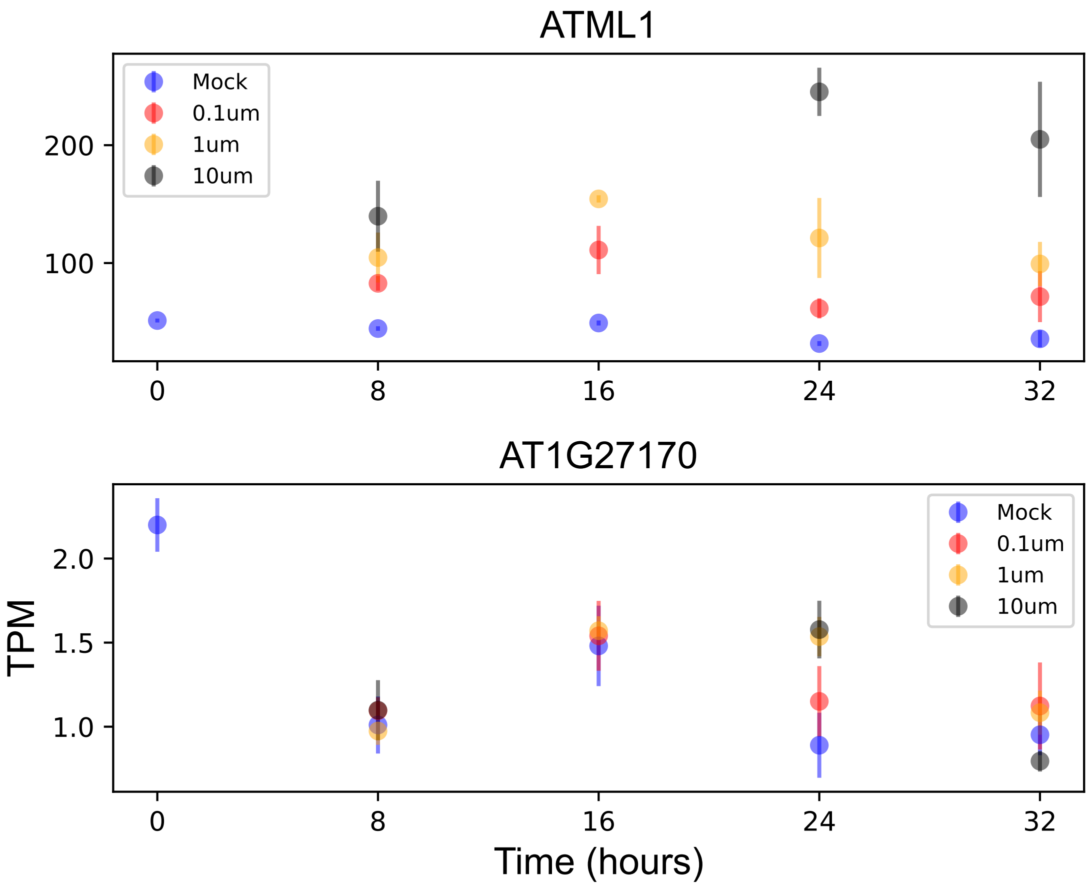

### AT1G27461.pdf

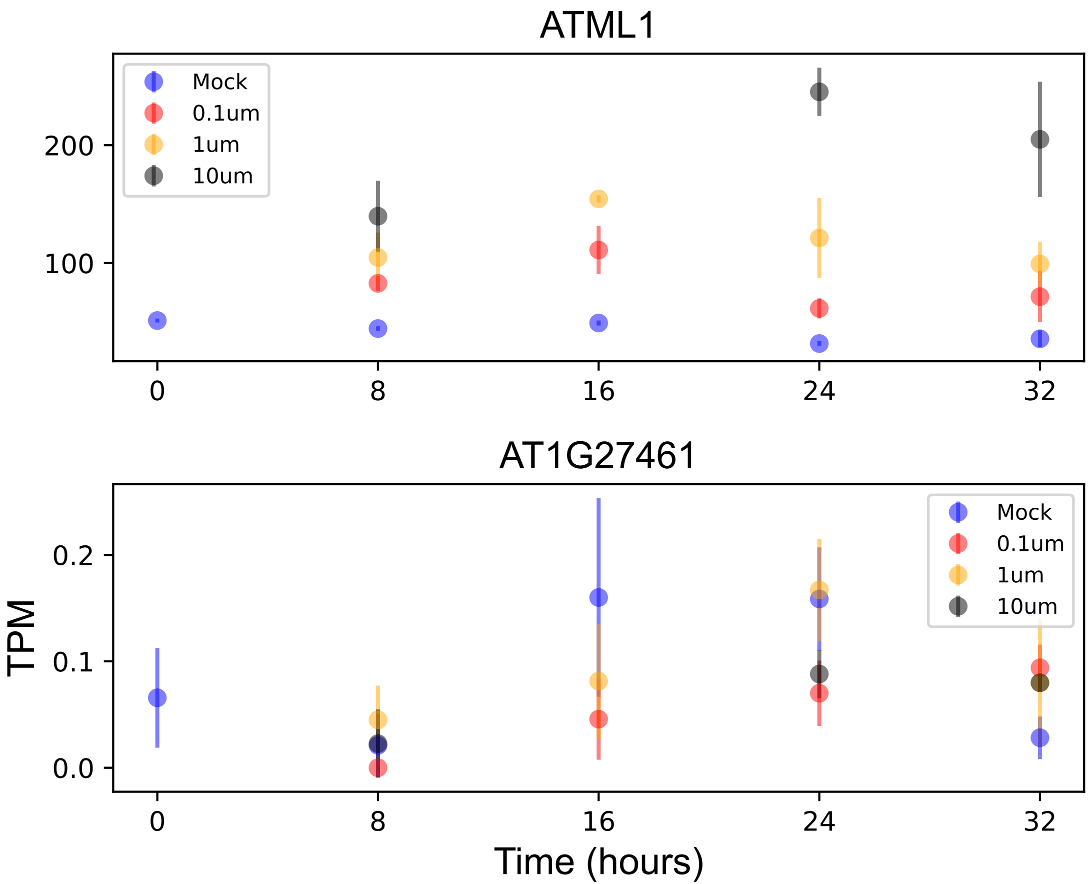

### AT1G28610.pdf

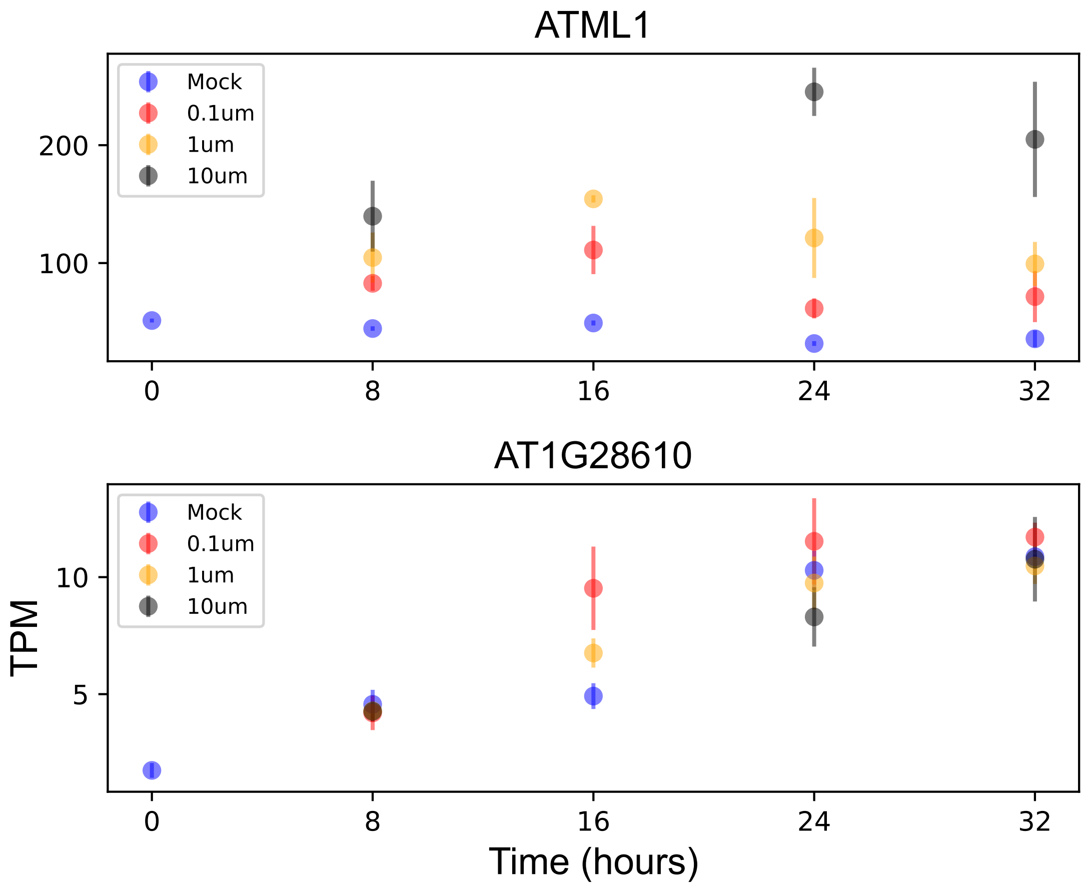

### AT1G29965.pdf

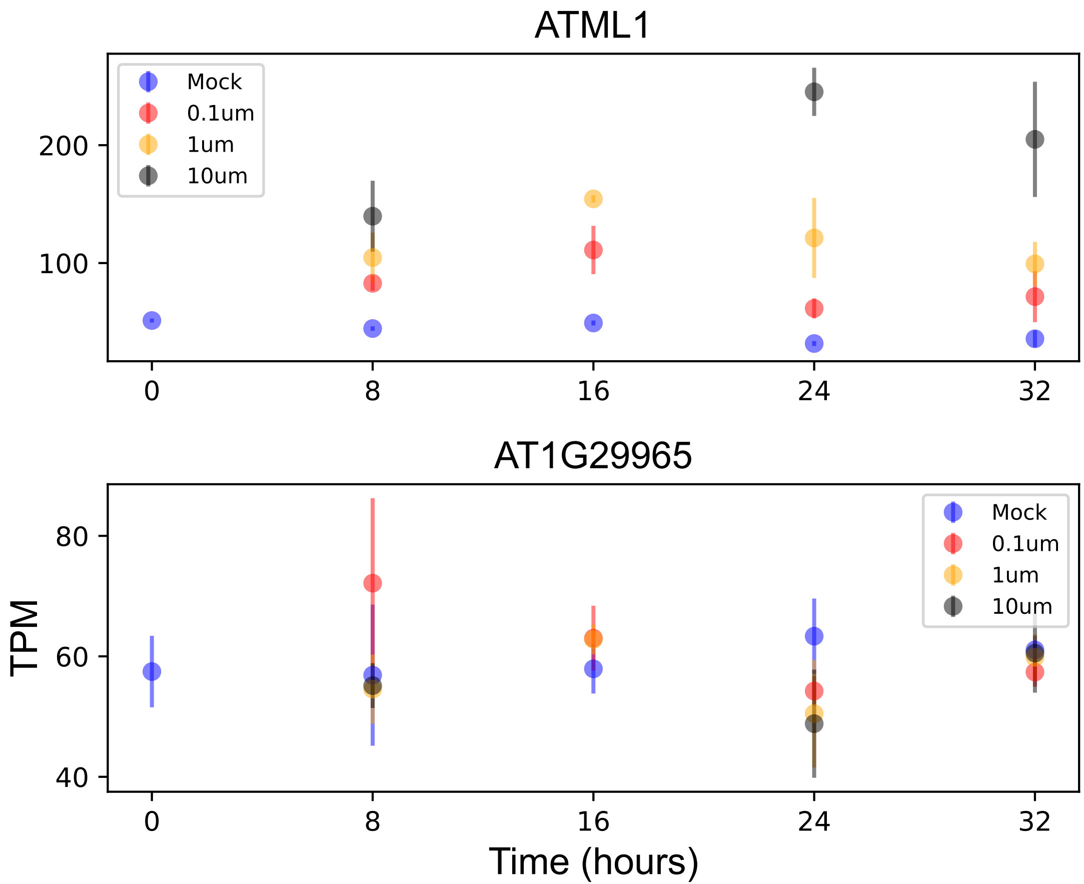

### AT1G45100.pdf

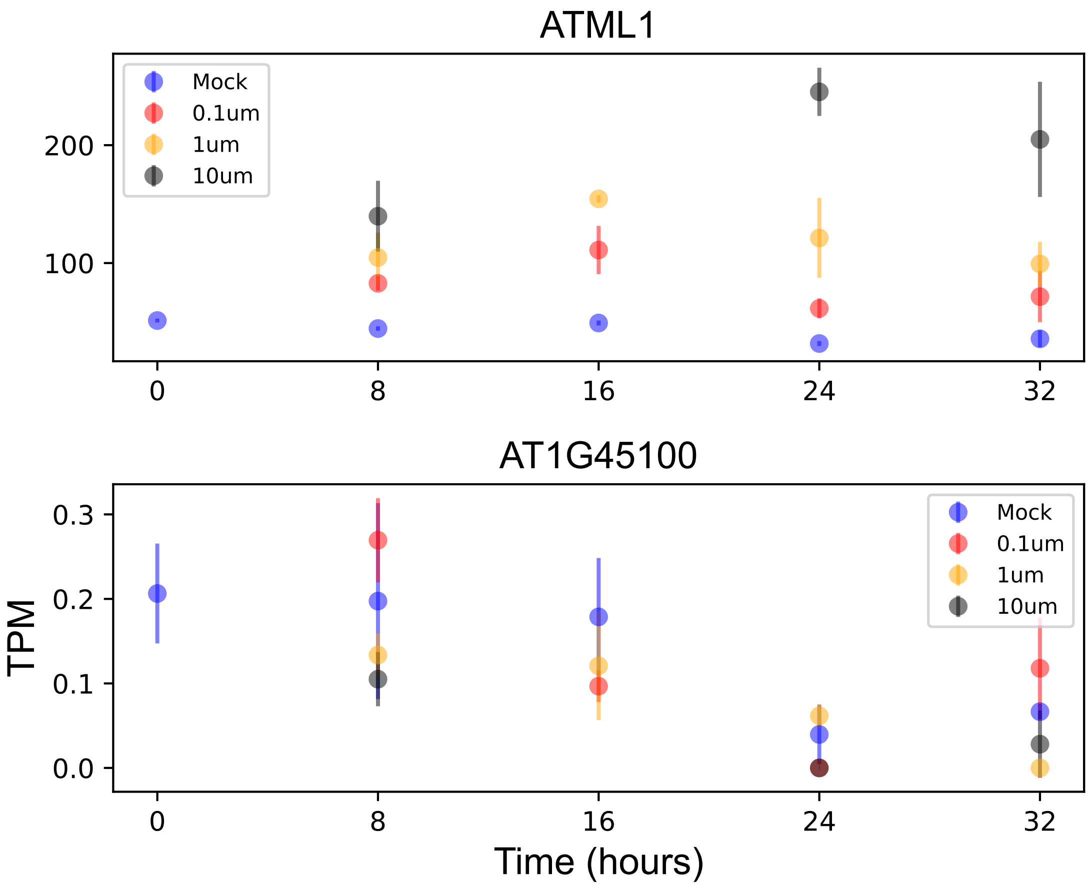

### AT1G55750.pdf

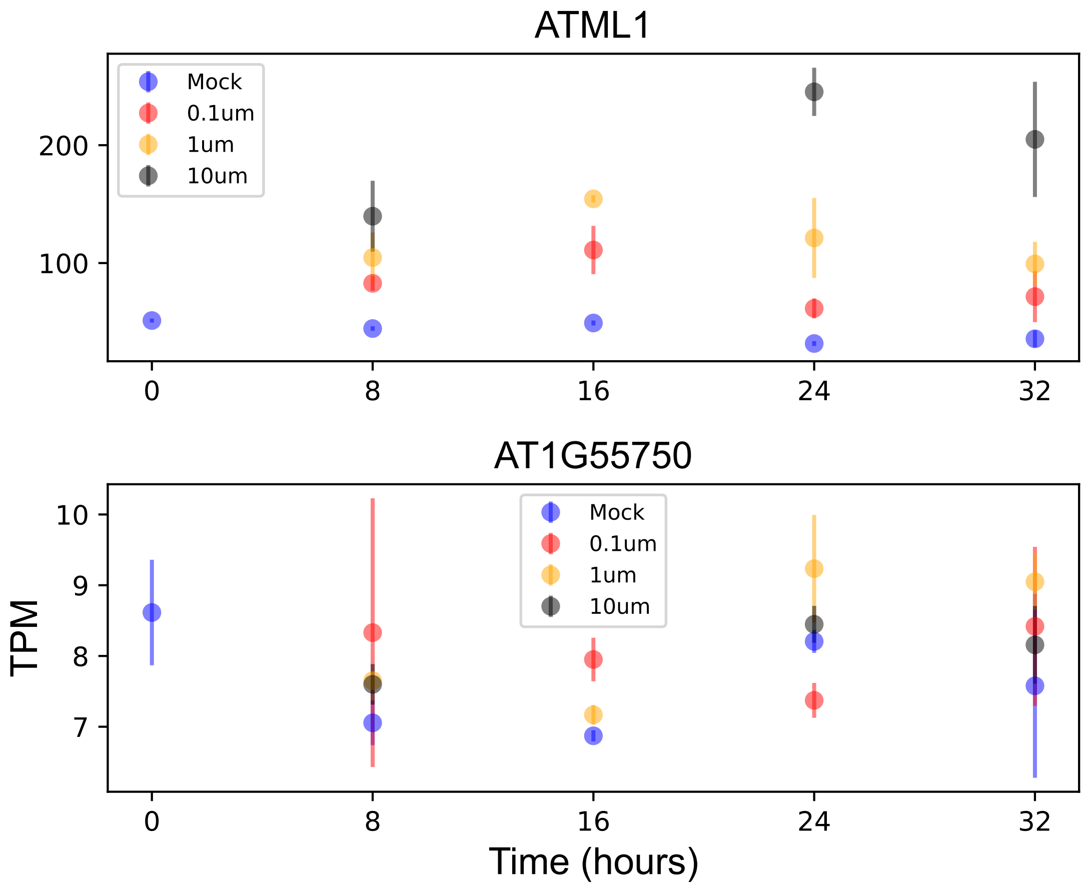

### AT1G56460.pdf

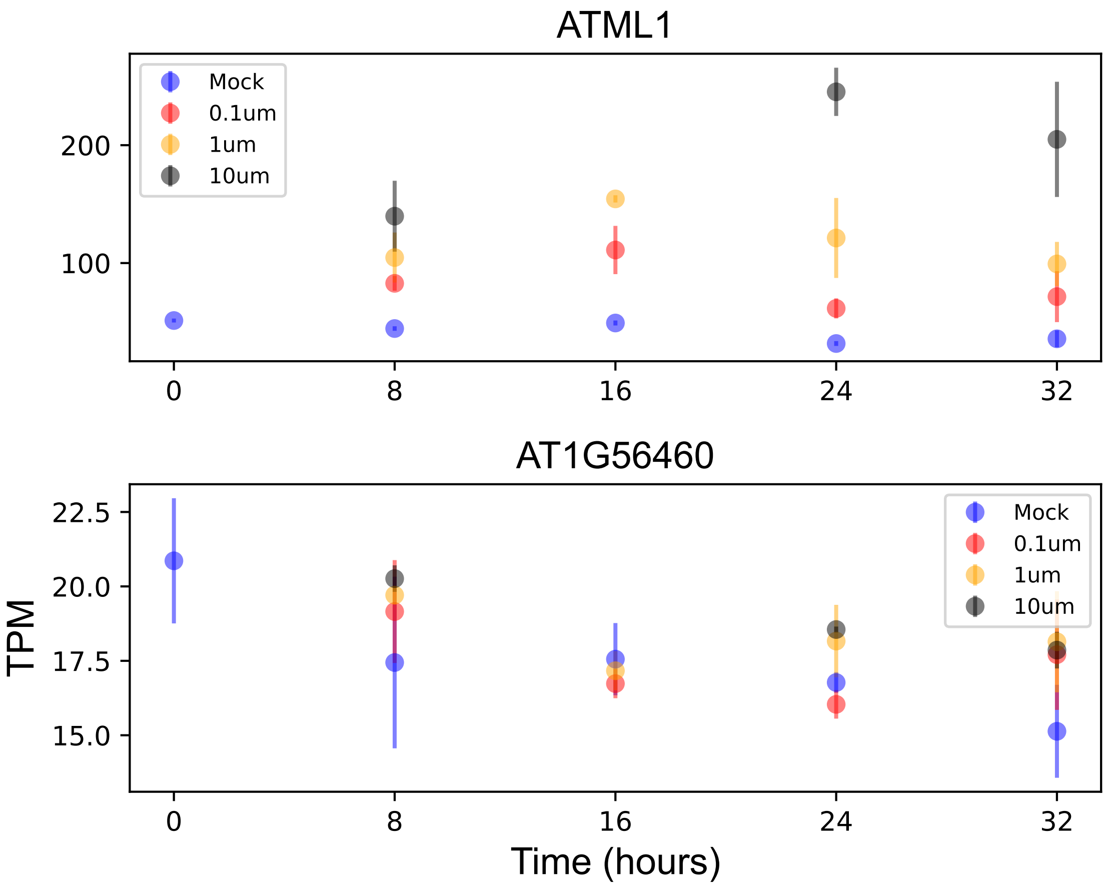

### AT1G63860.pdf

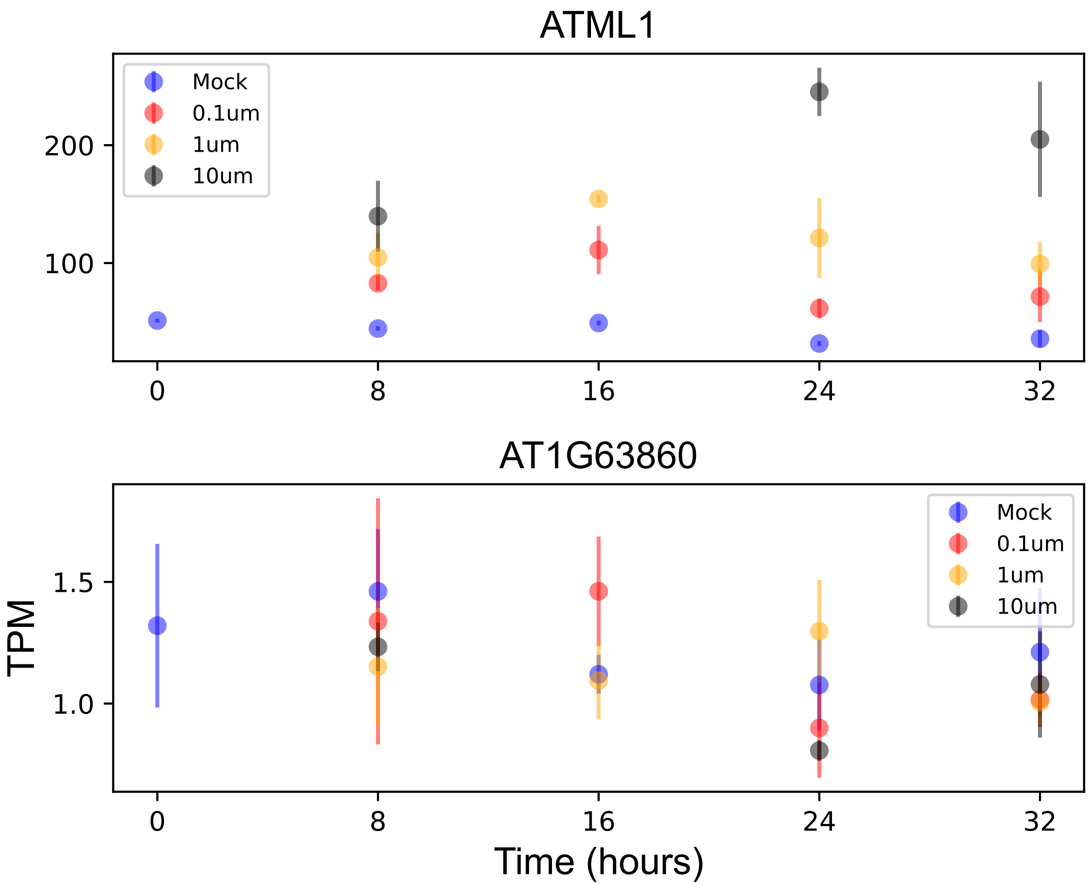

### AT1G66180.pdf

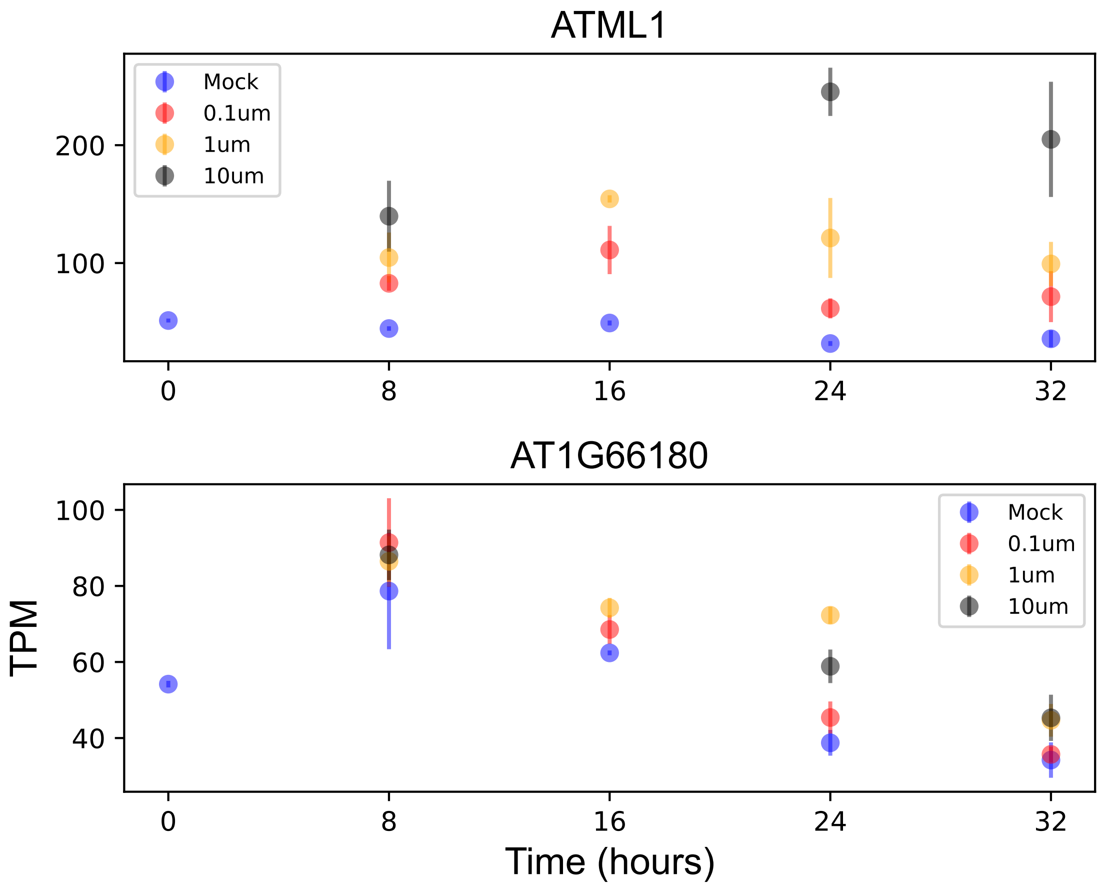

### AT1G78450.pdf

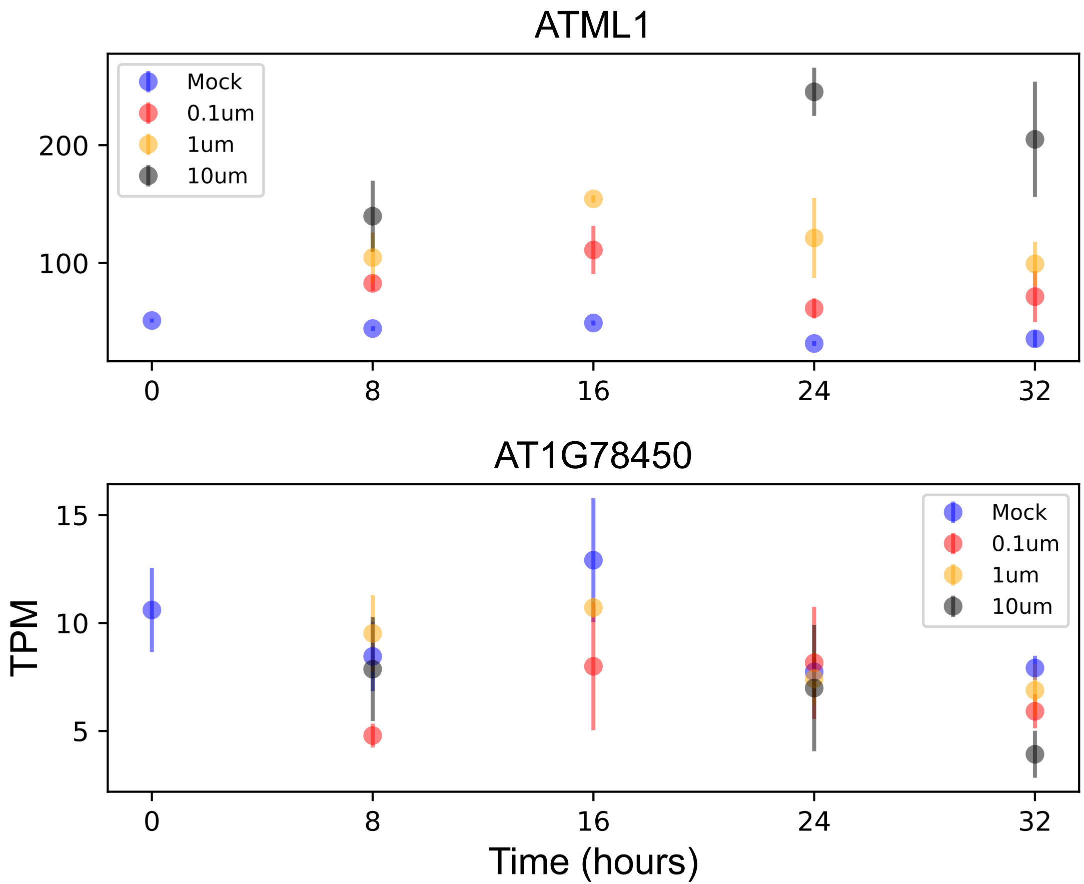

### AT2G02520.pdf

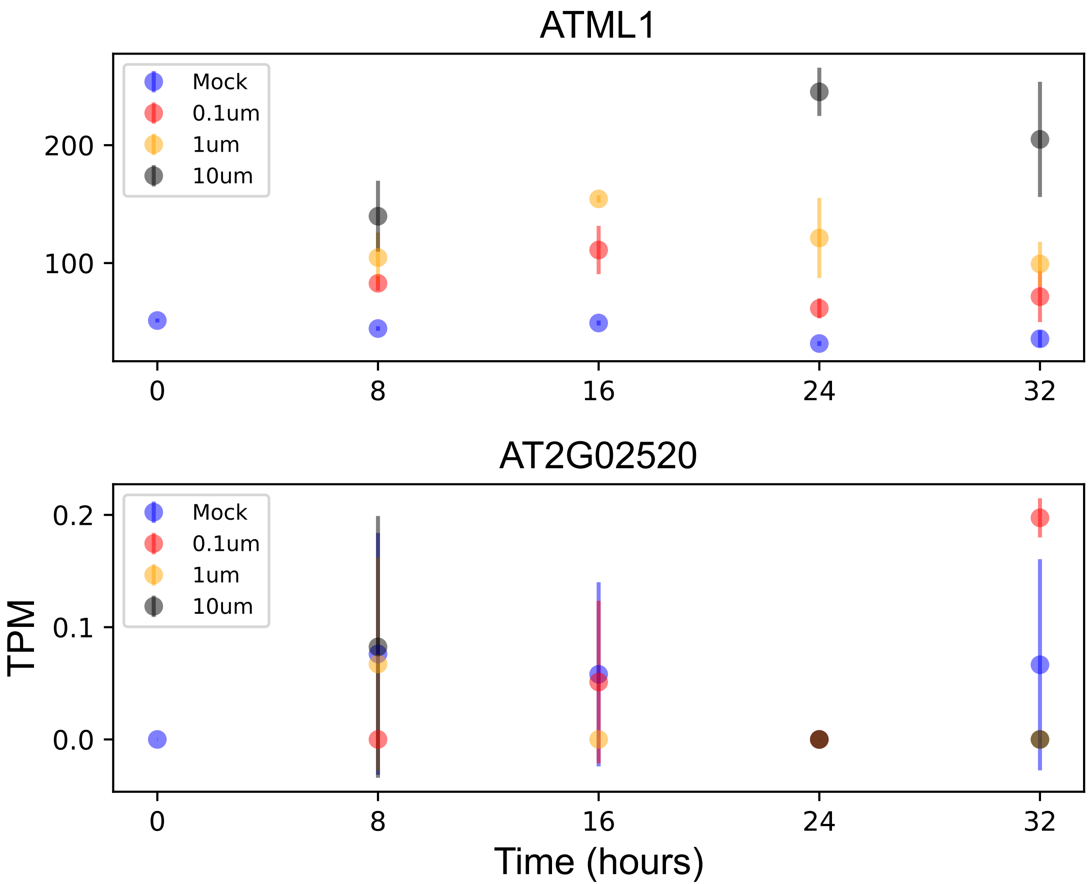

### AT2G03980.pdf

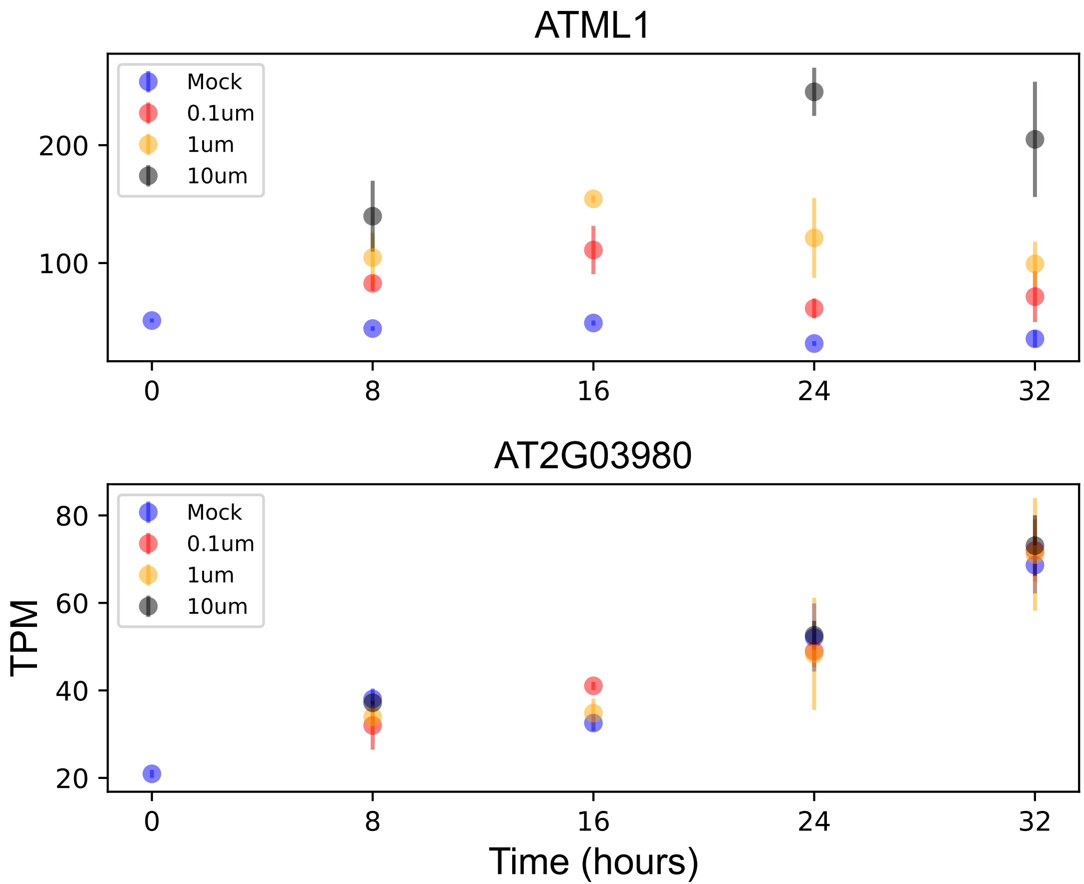

### AT2G07648.pdf

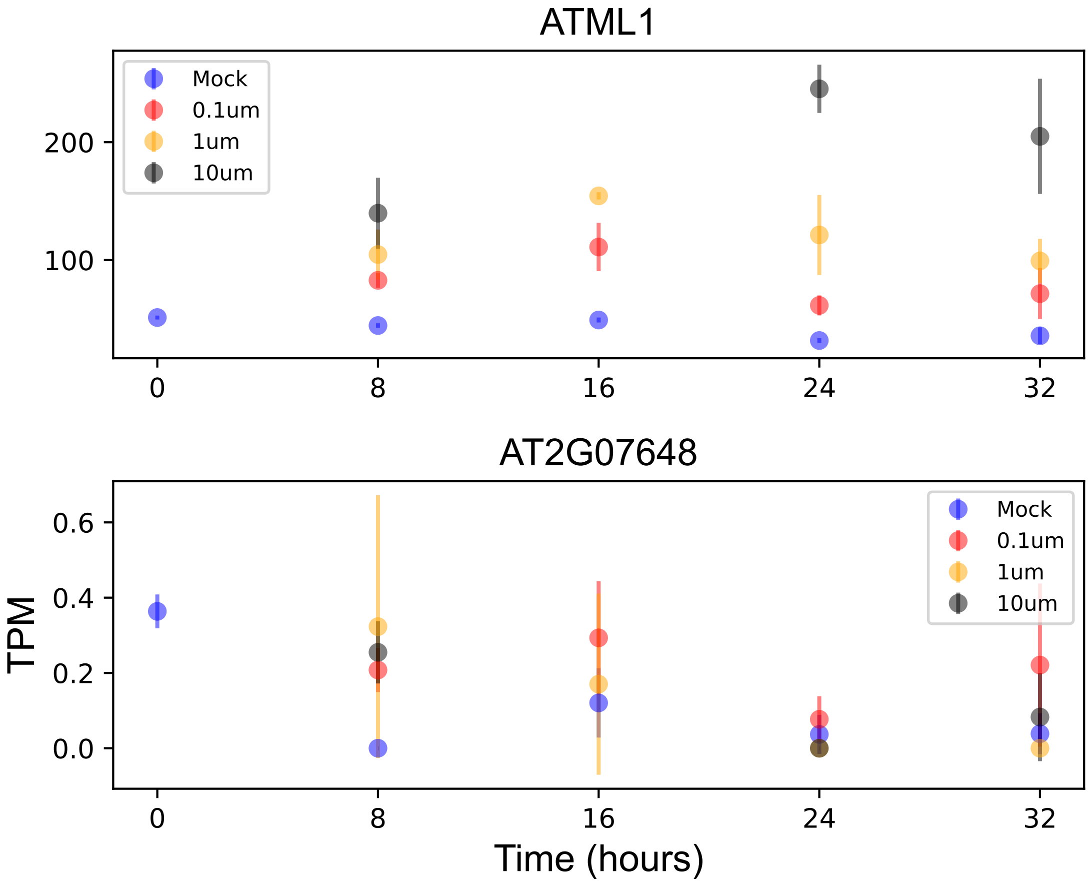

### AT2G18110.pdf

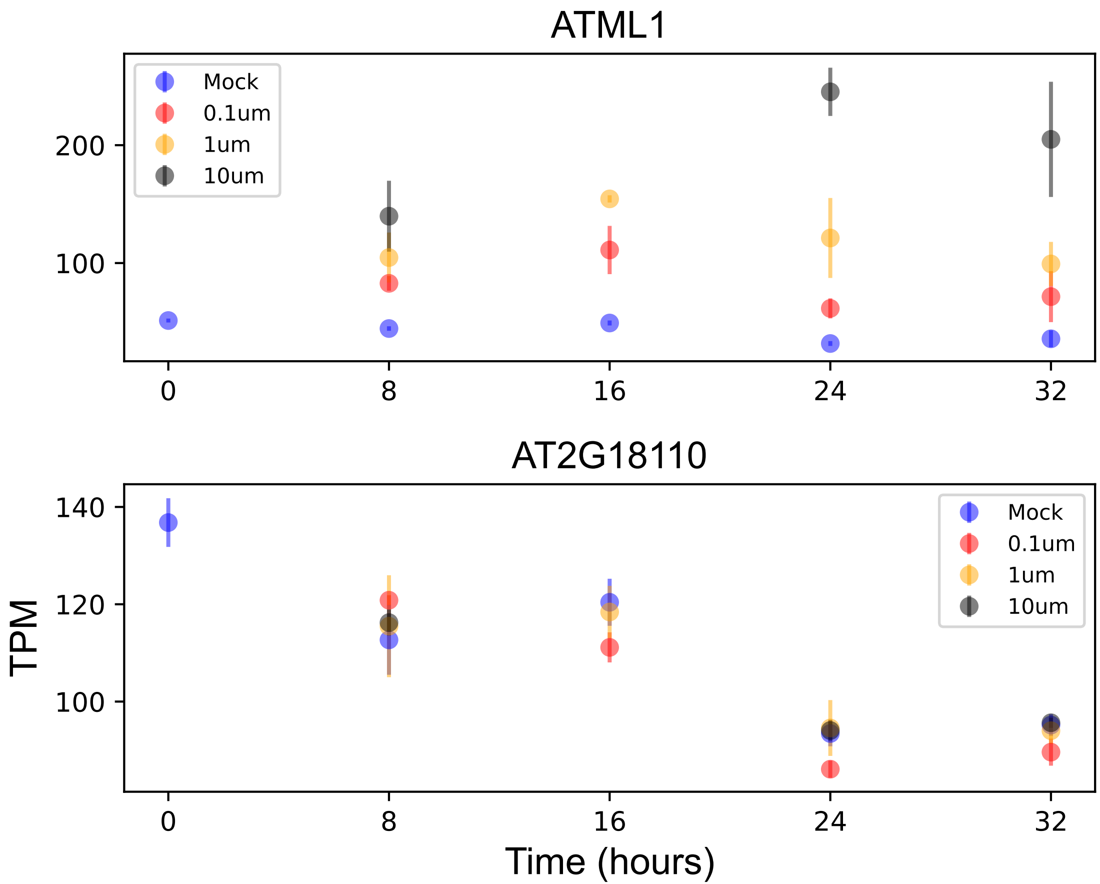

### AT2G35830.pdf

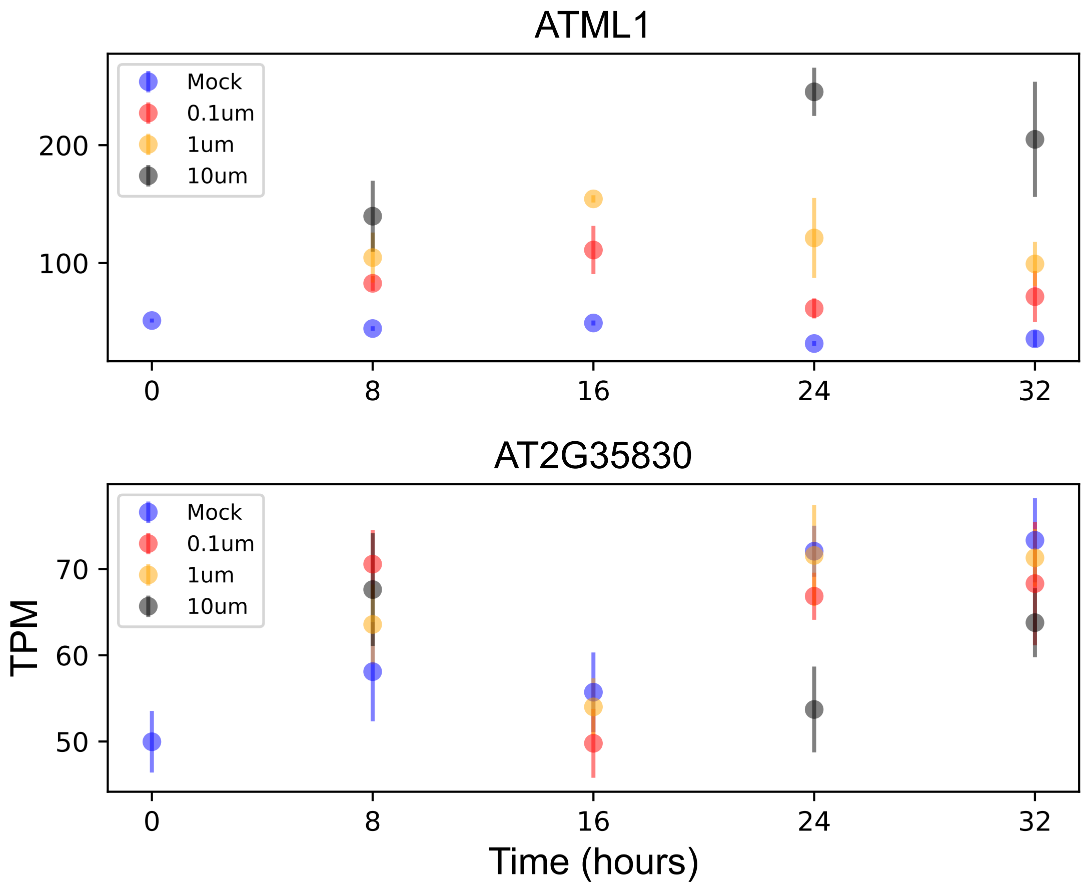

### AT2G43490.pdf

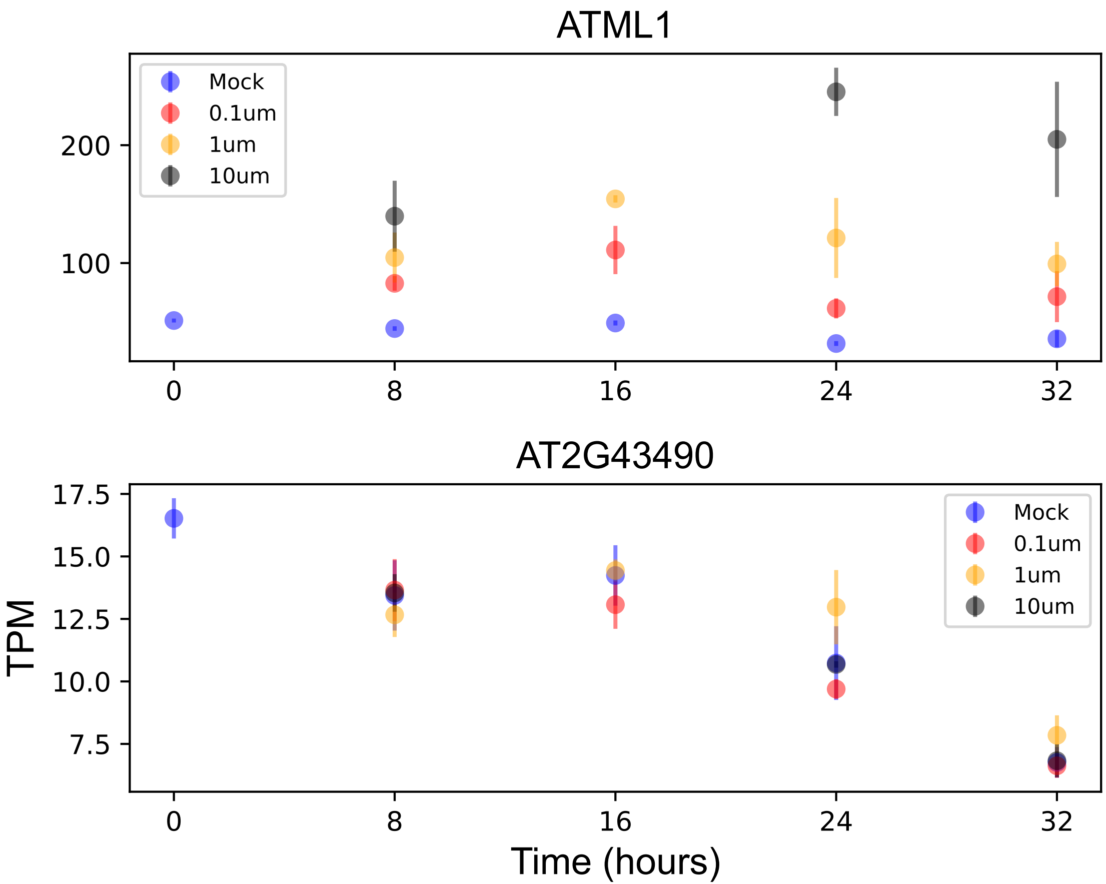

### AT2G44030.pdf

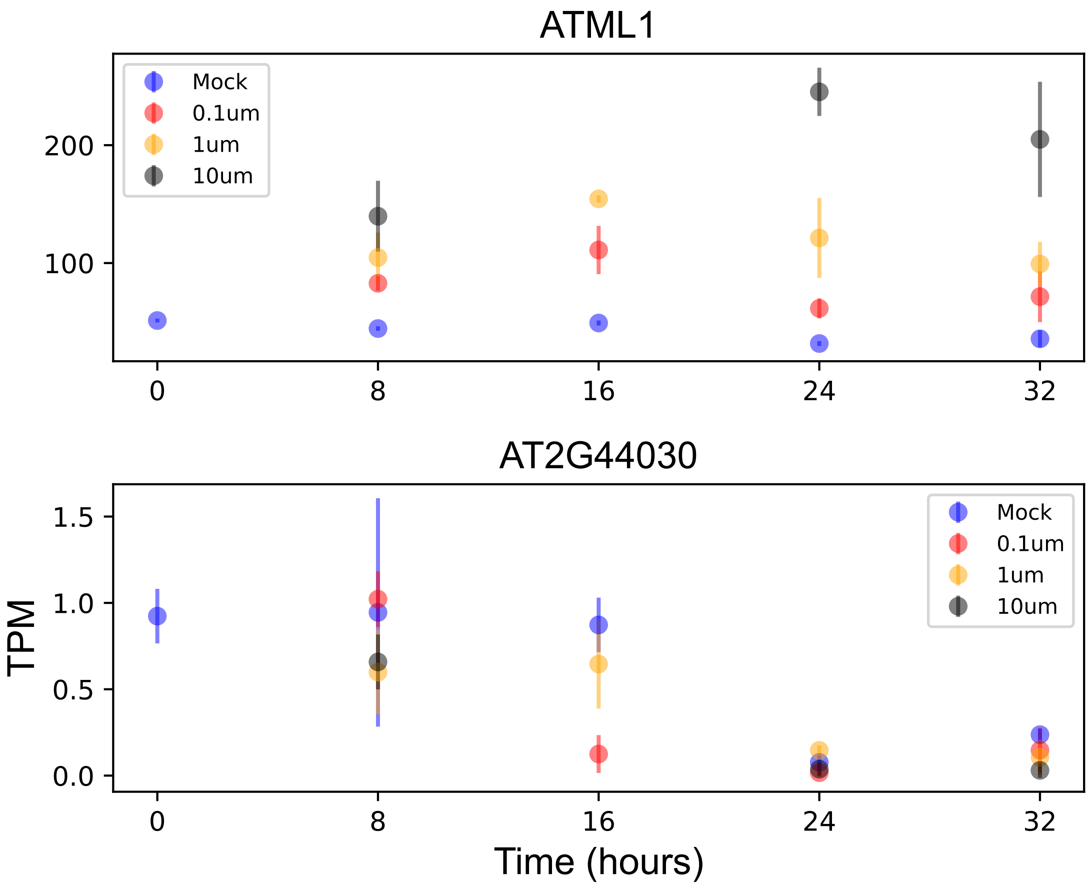

### AT2G44860.pdf

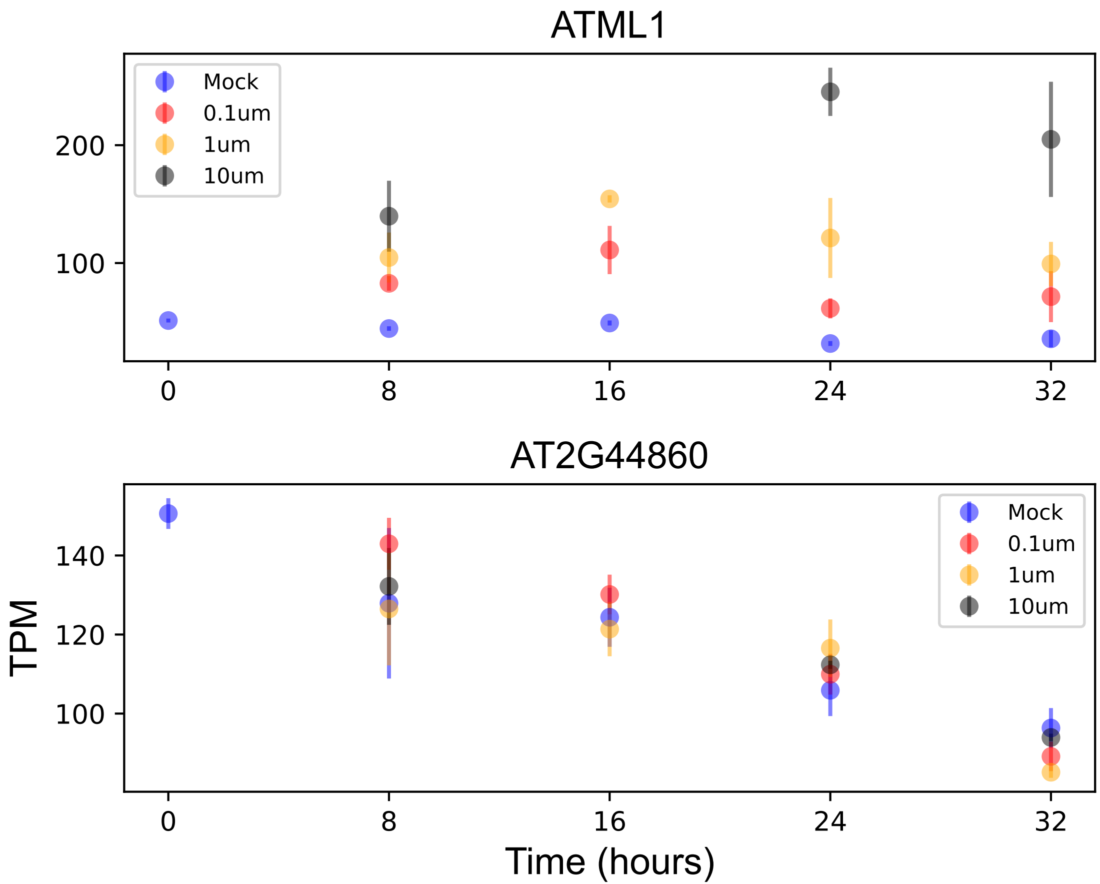

### AT3G04840.pdf

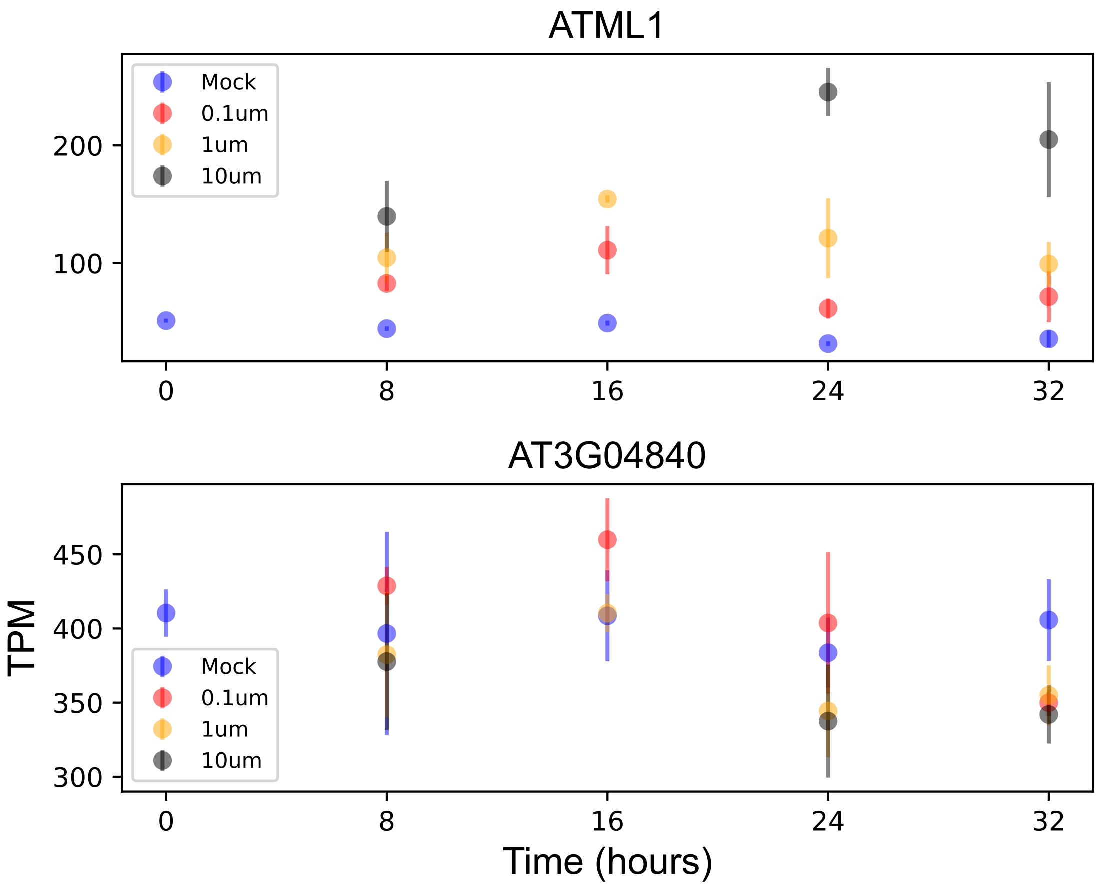

### AT3G06000.pdf

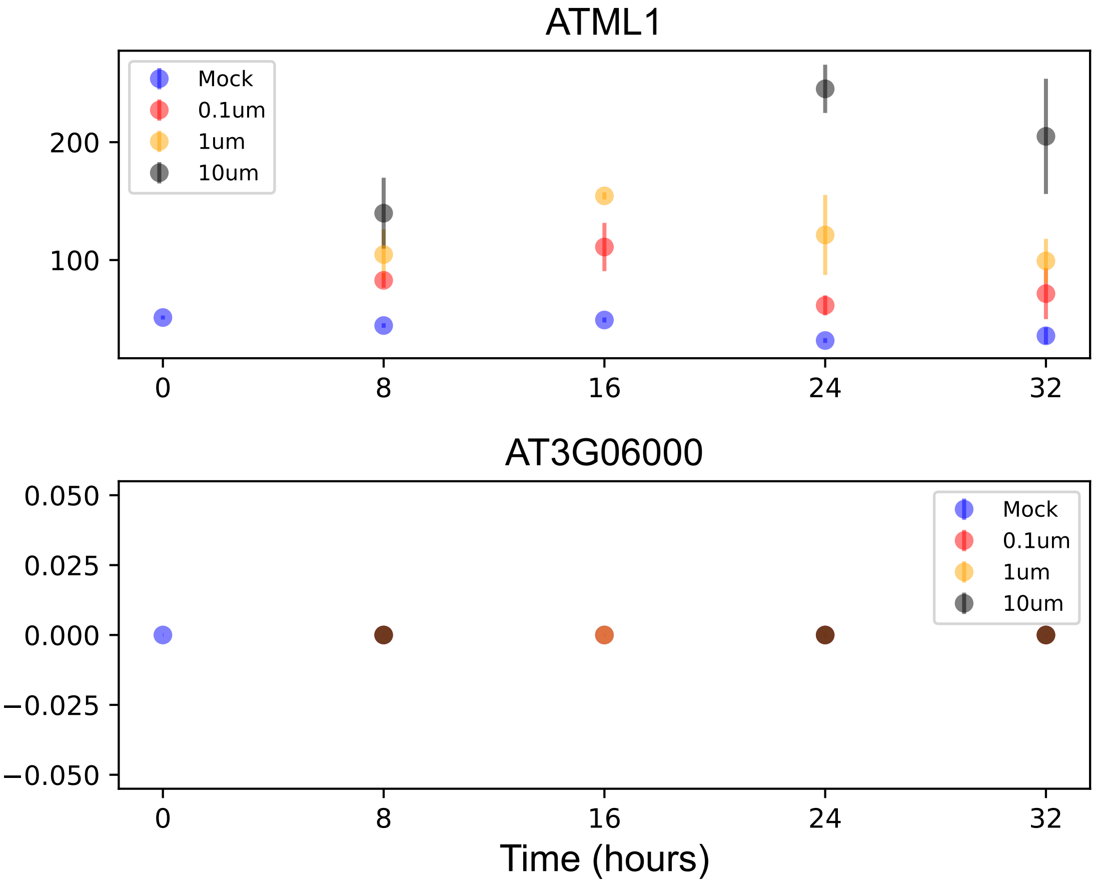

### AT3G07070.pdf

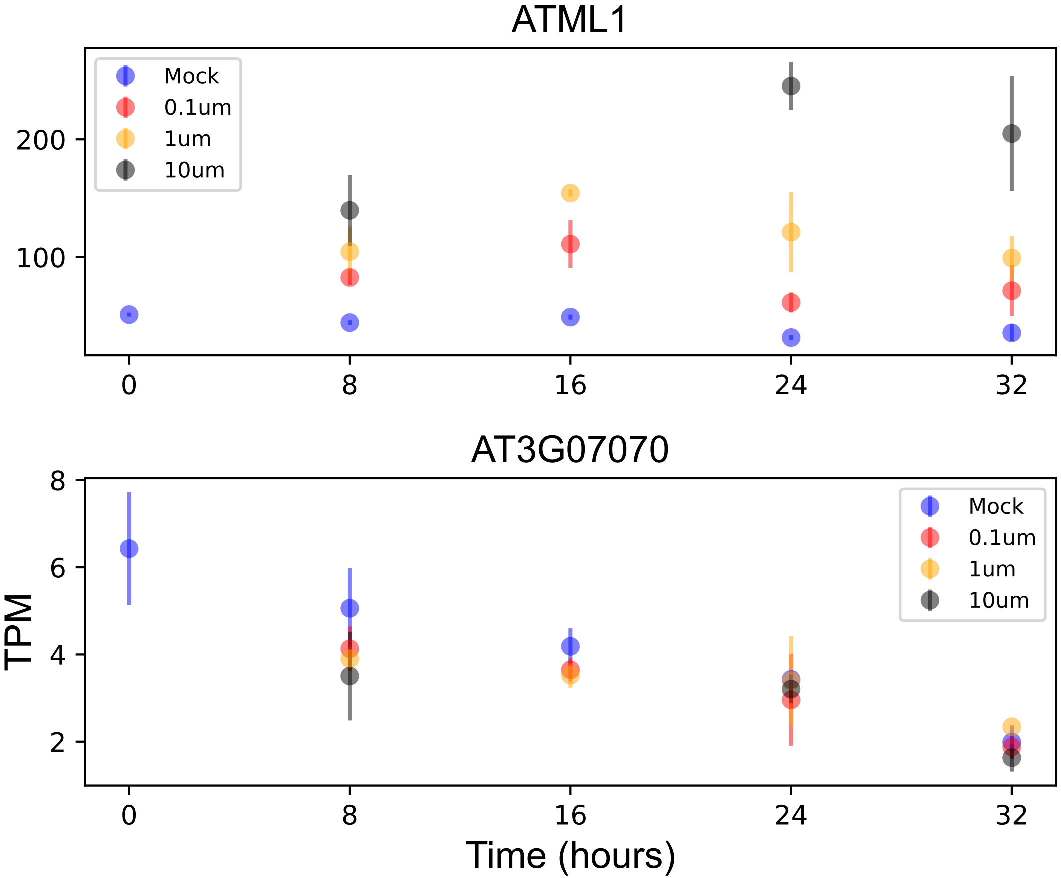

### AT3G11530.pdf

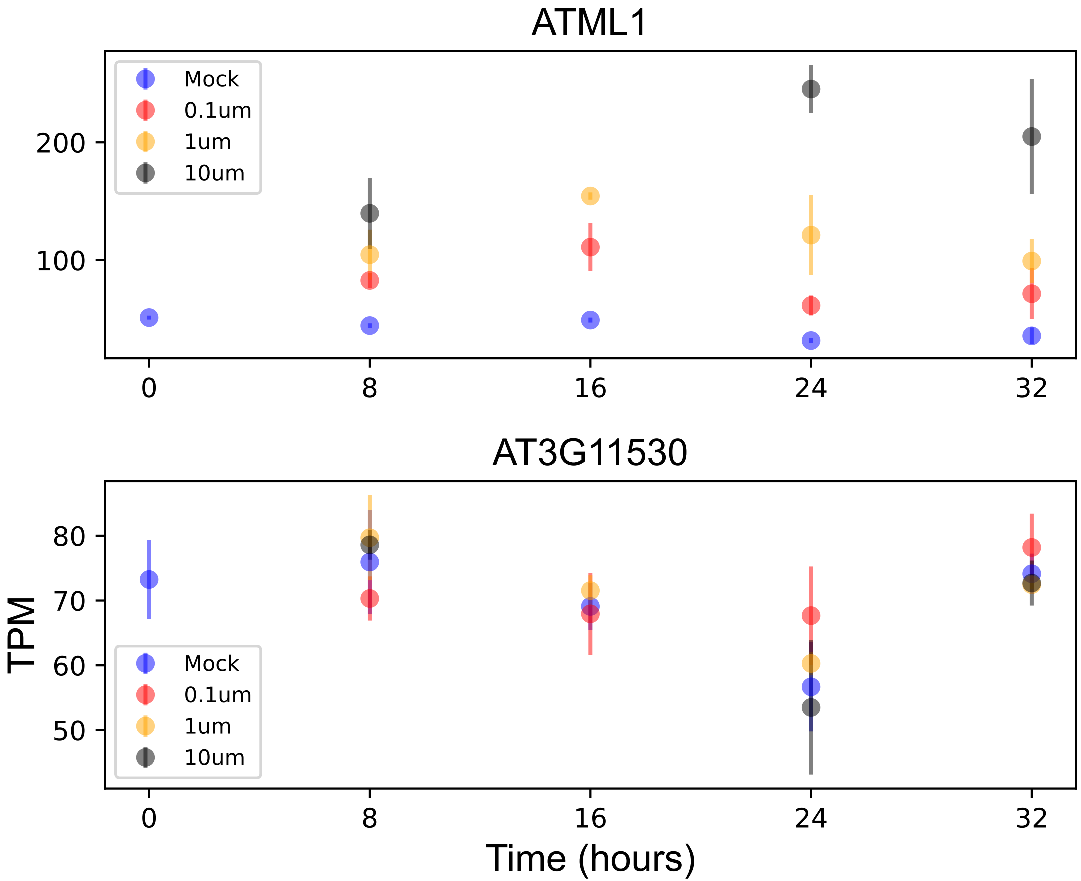

### AT3G11850.pdf

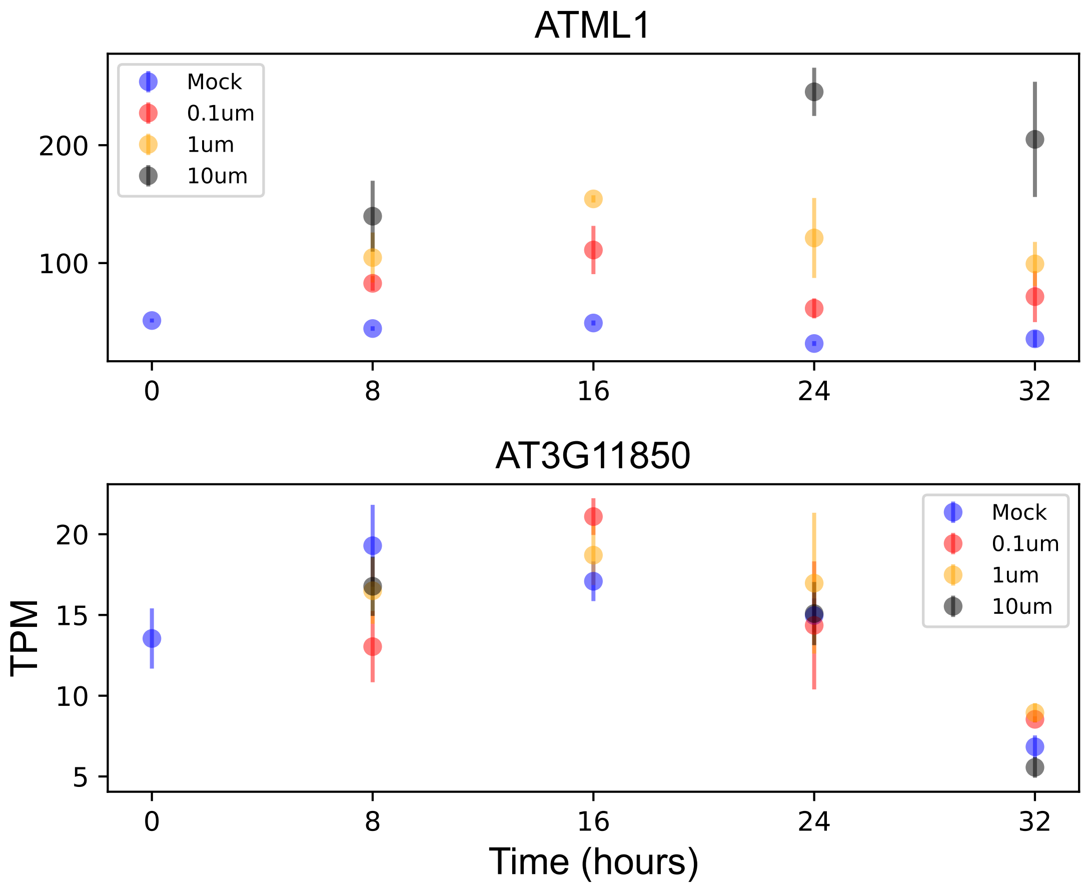

### AT3G21480.pdf

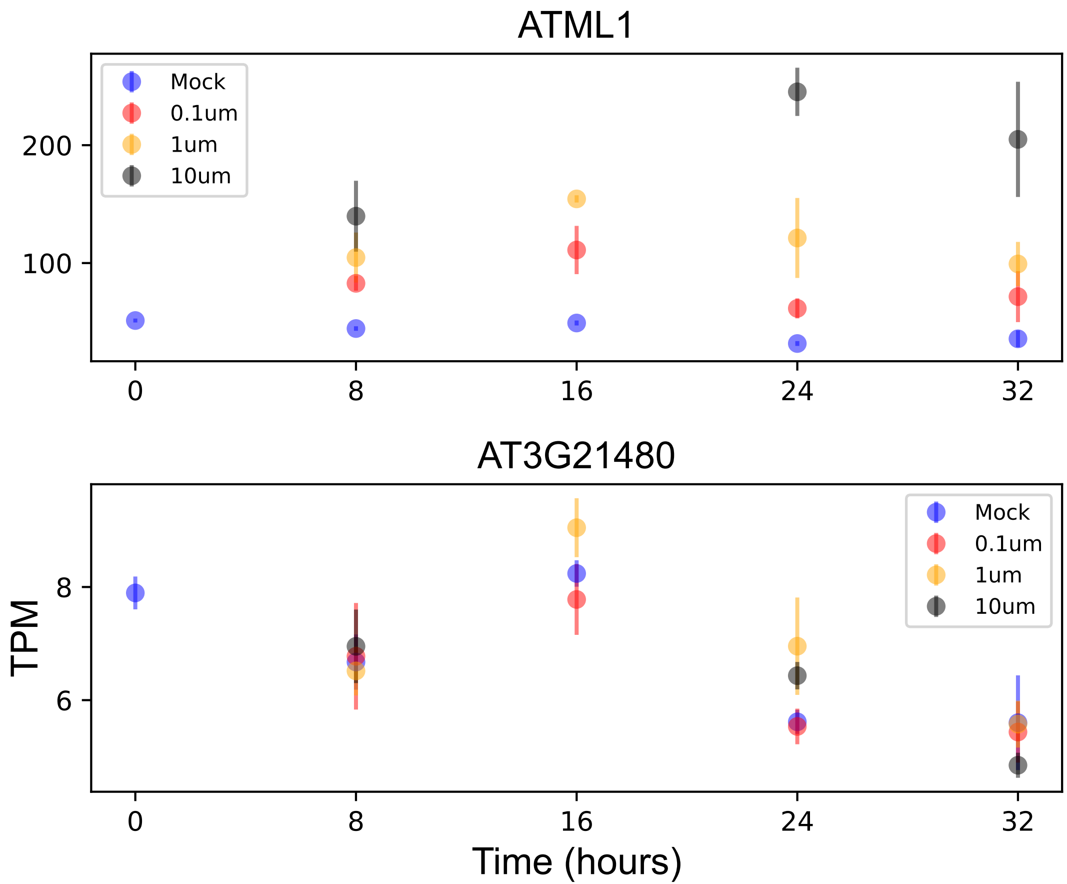

### AT3G27610.pdf

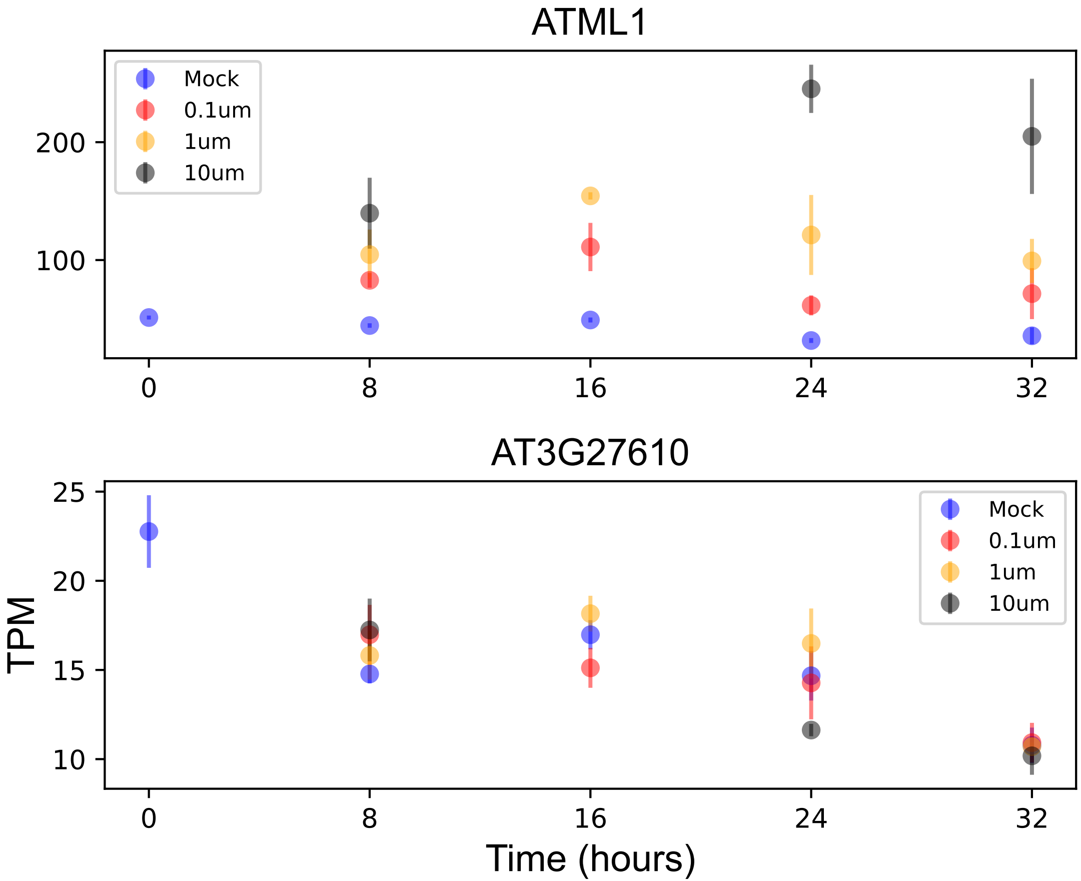

### AT3G28760.pdf

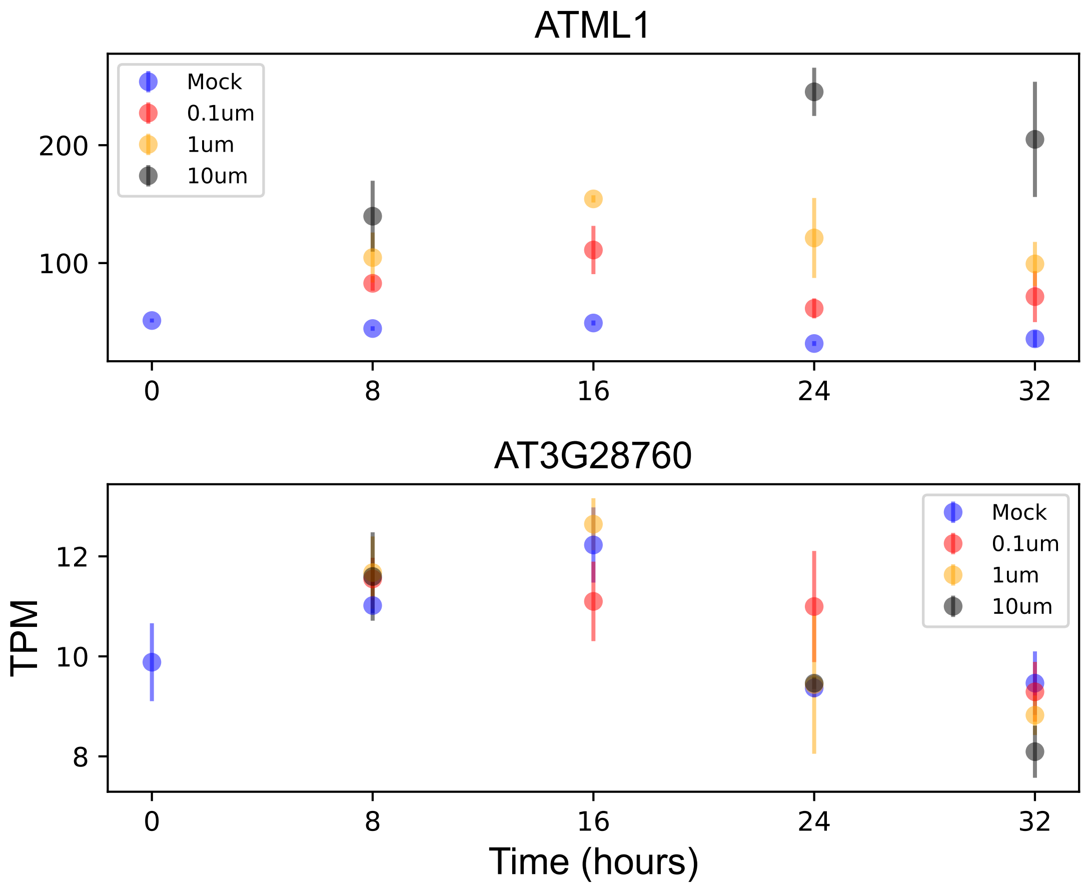

### AT3G45950.pdf

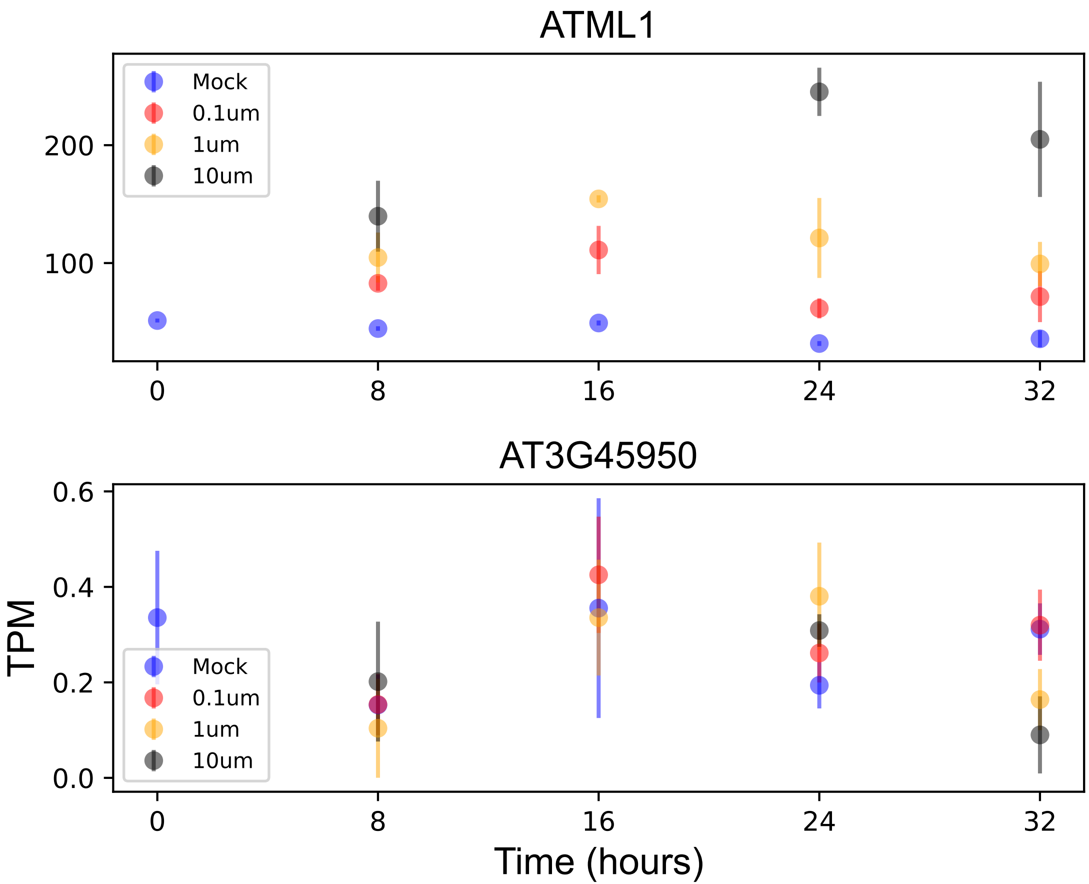

### AT3G50370.pdf

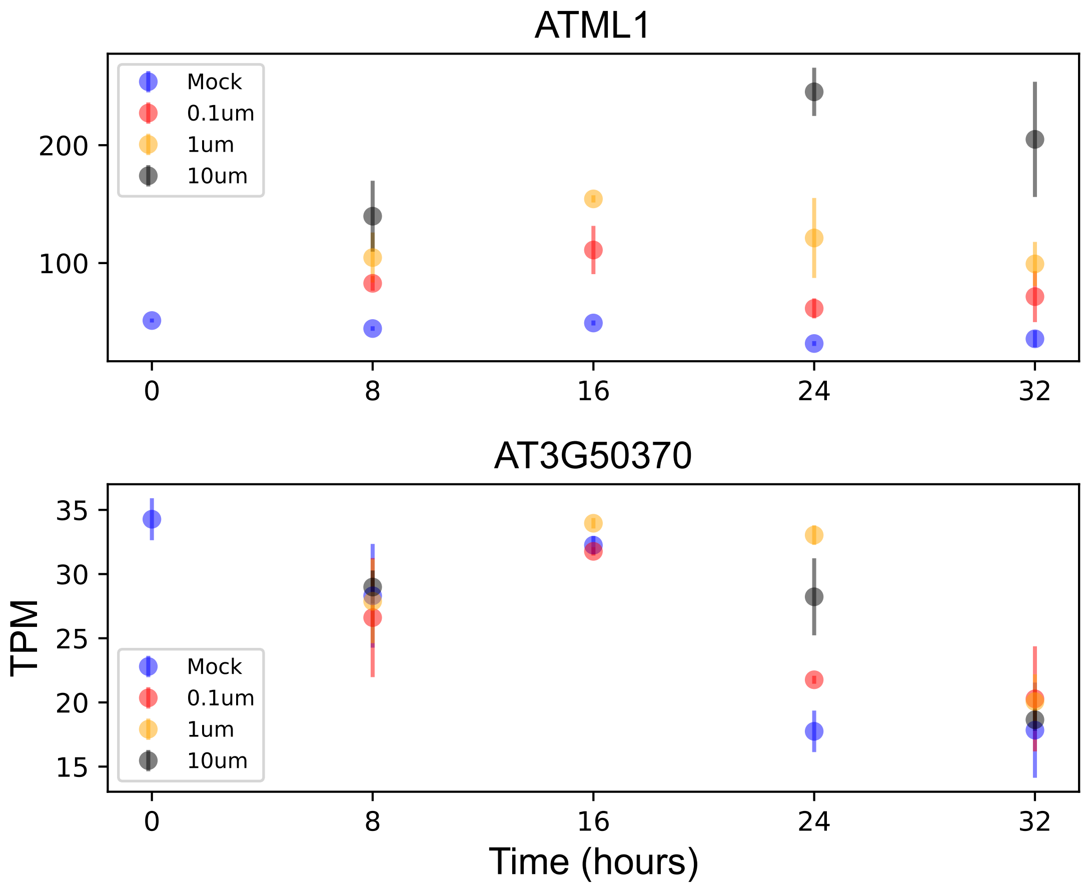

### AT3G51070.pdf

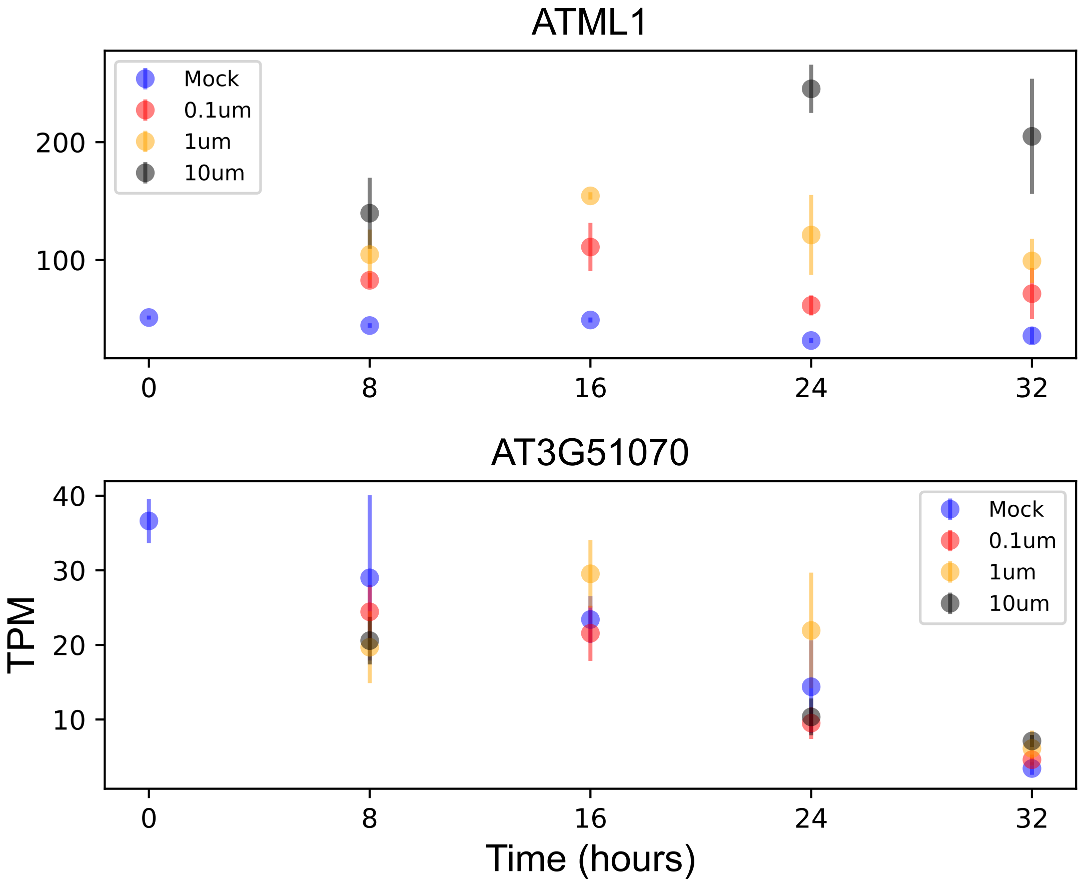

### AT3G60670.pdf

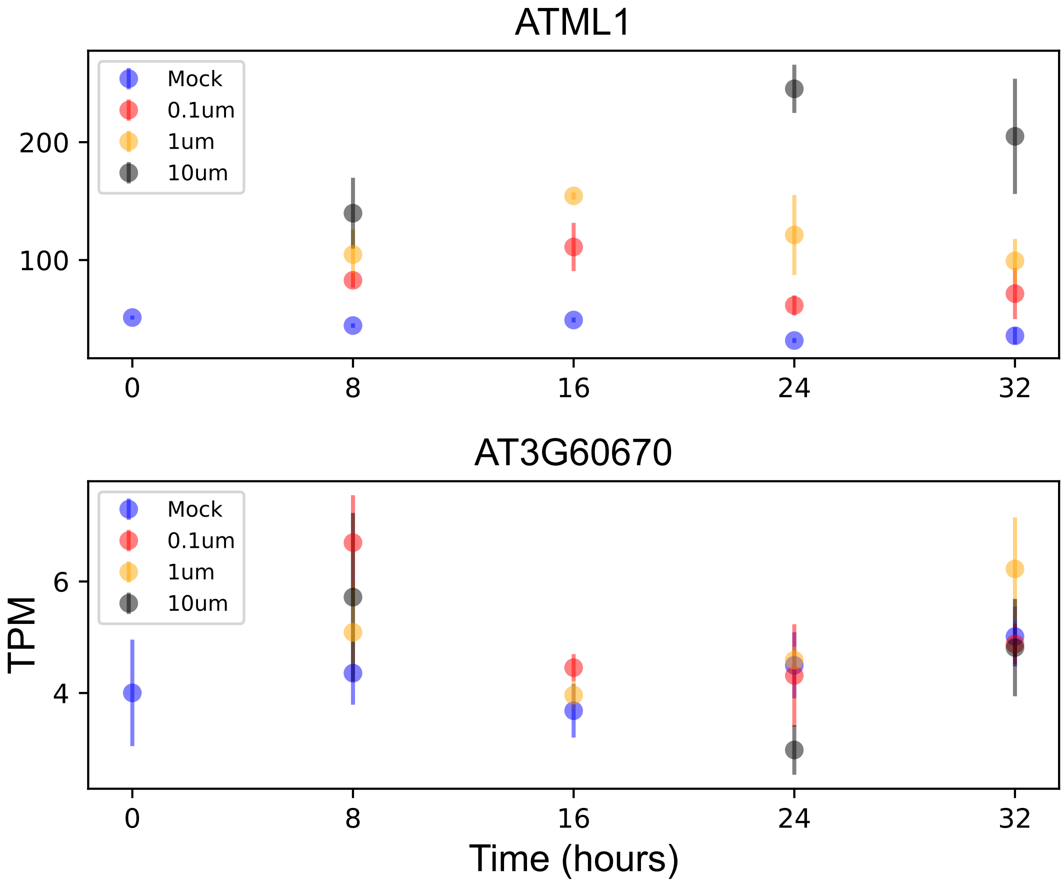

### AT3G61170.pdf

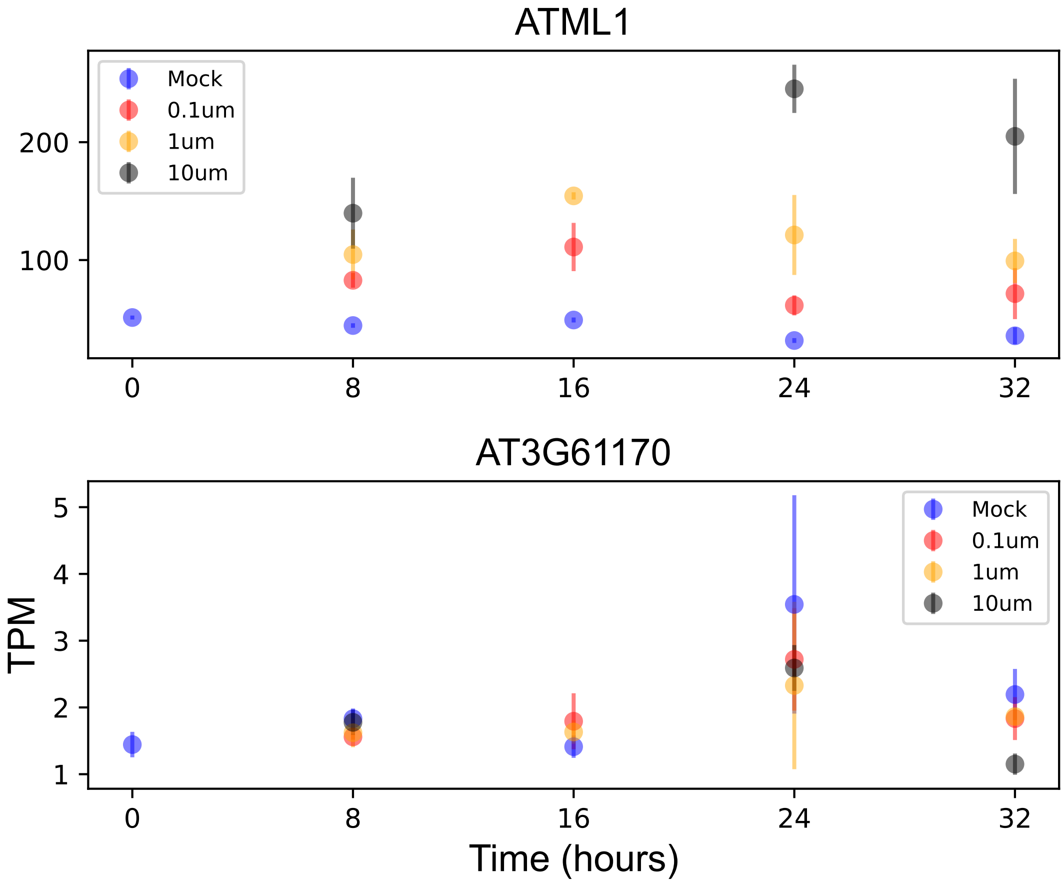

### AT4G03440.pdf

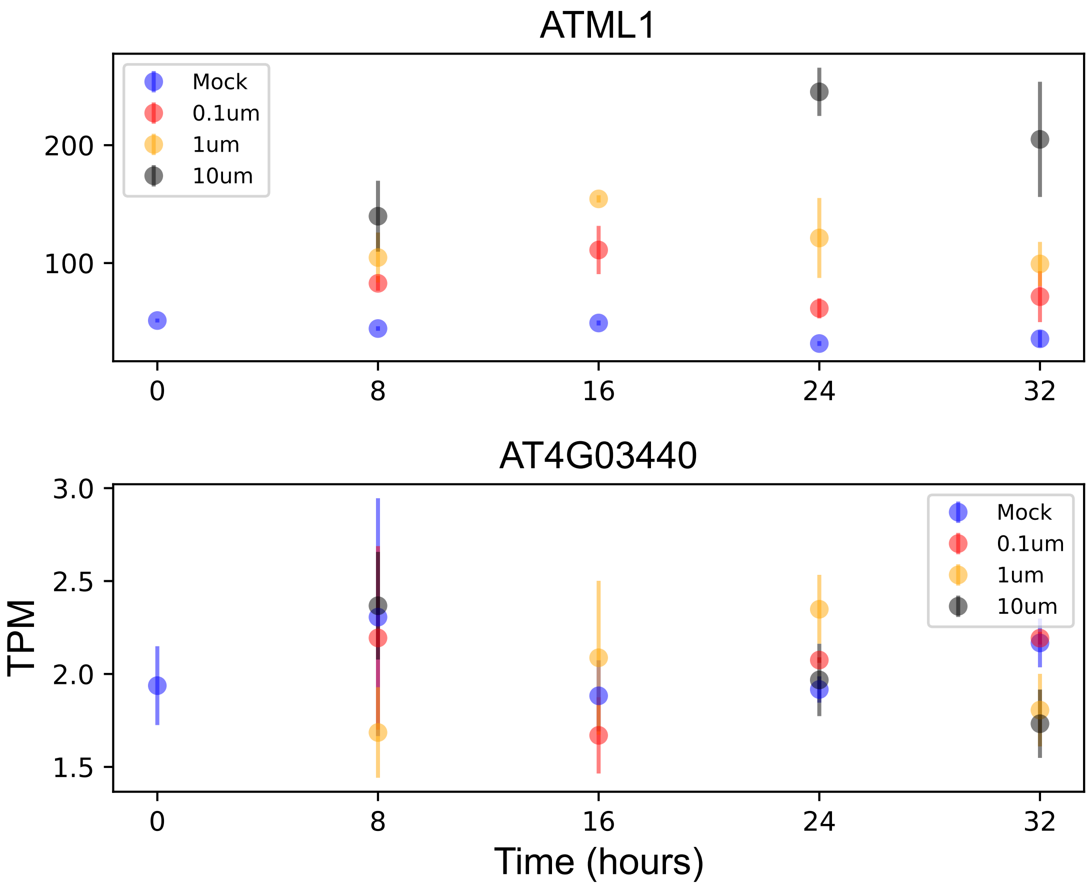

### AT4G09170.pdf

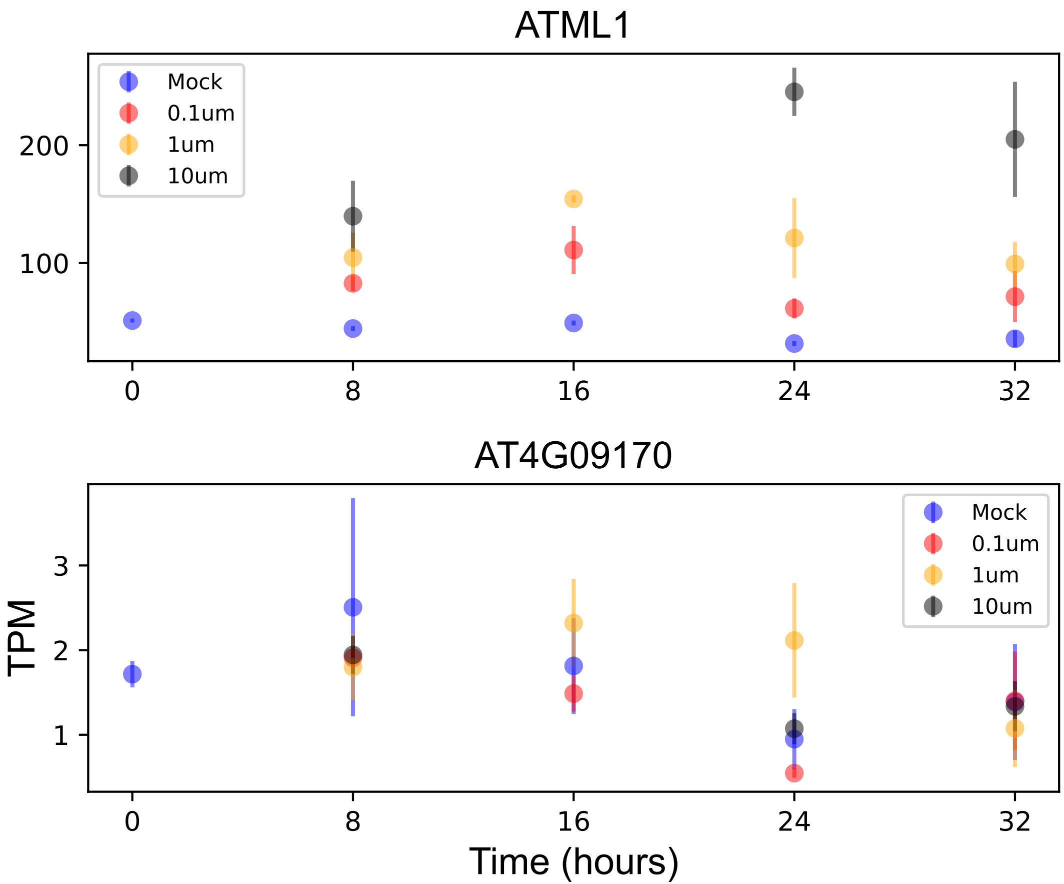

### AT4G09750.pdf

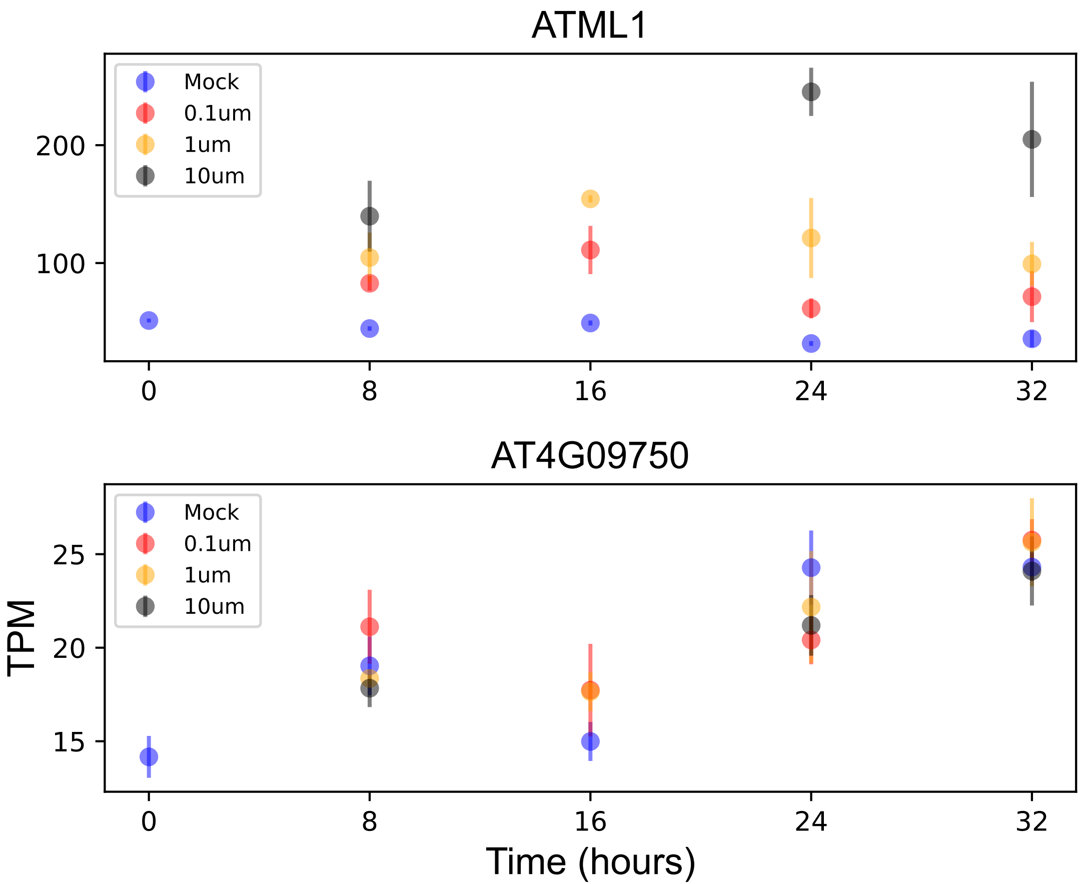

### AT4G18823.pdf

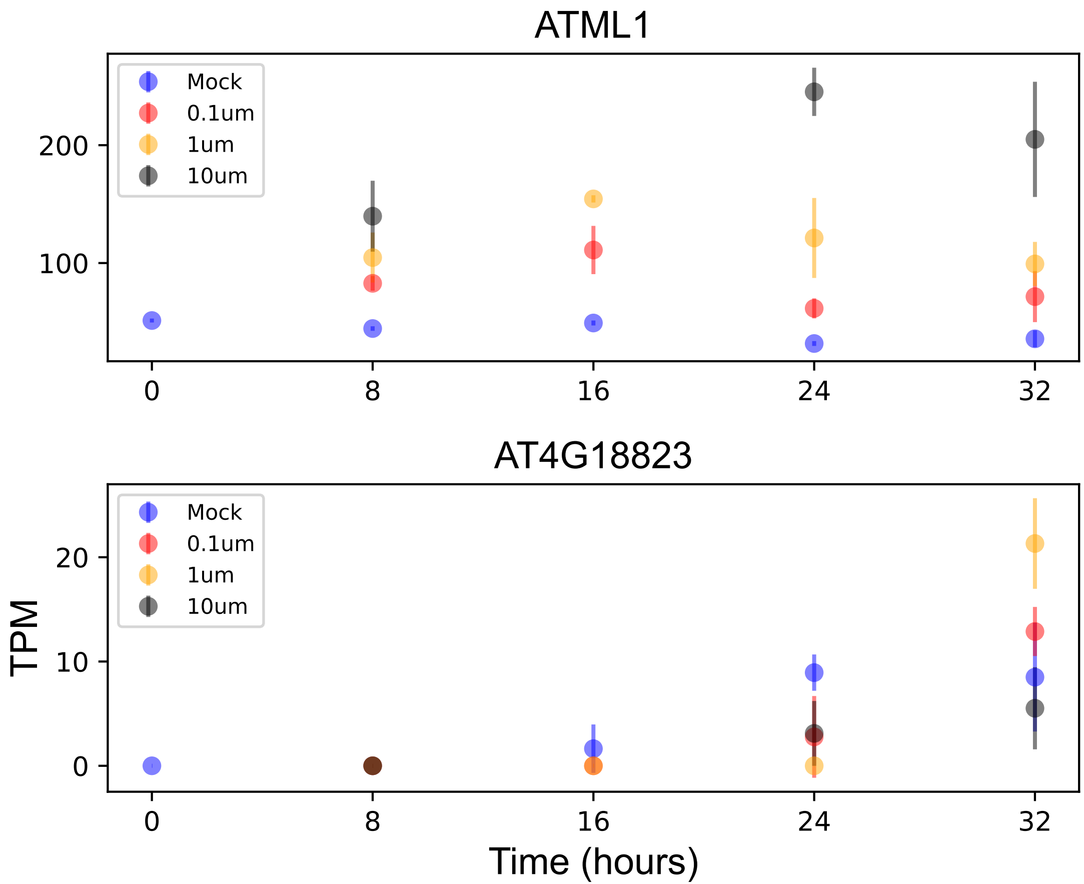

### AT4G19730.pdf

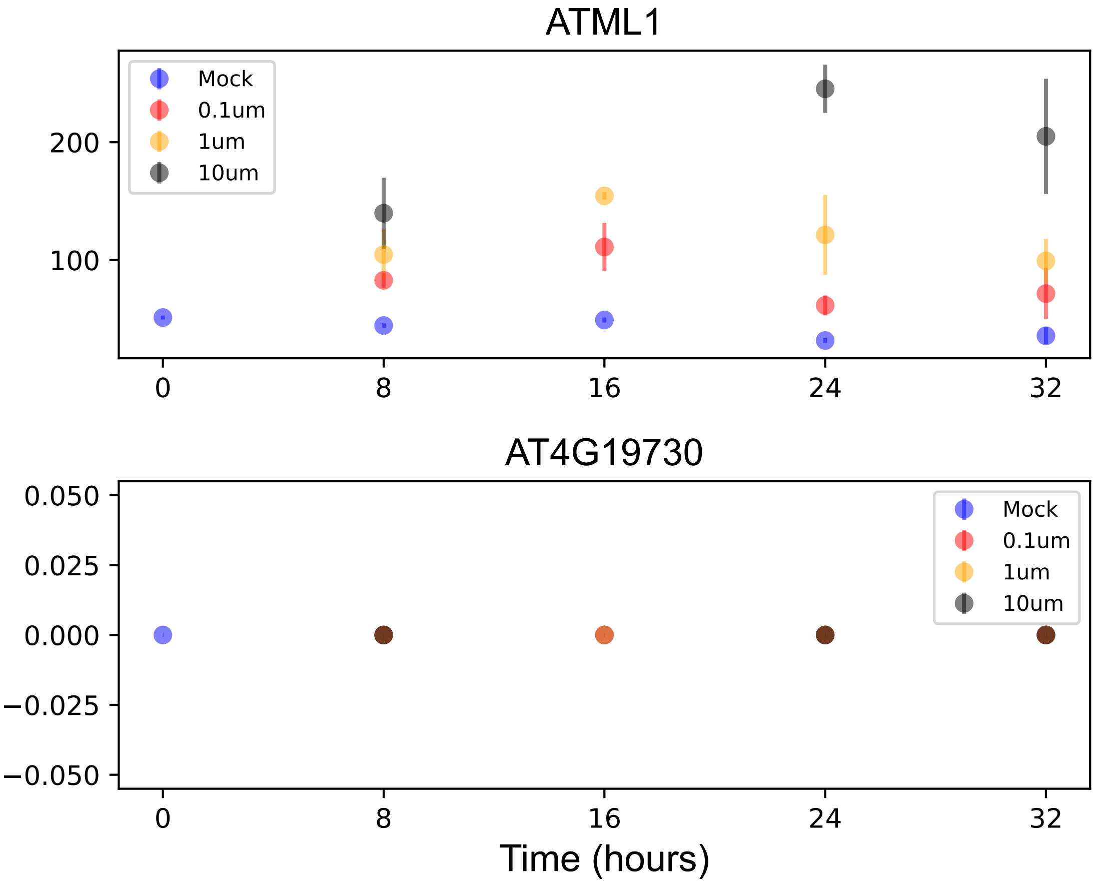

### AT4G27250.pdf

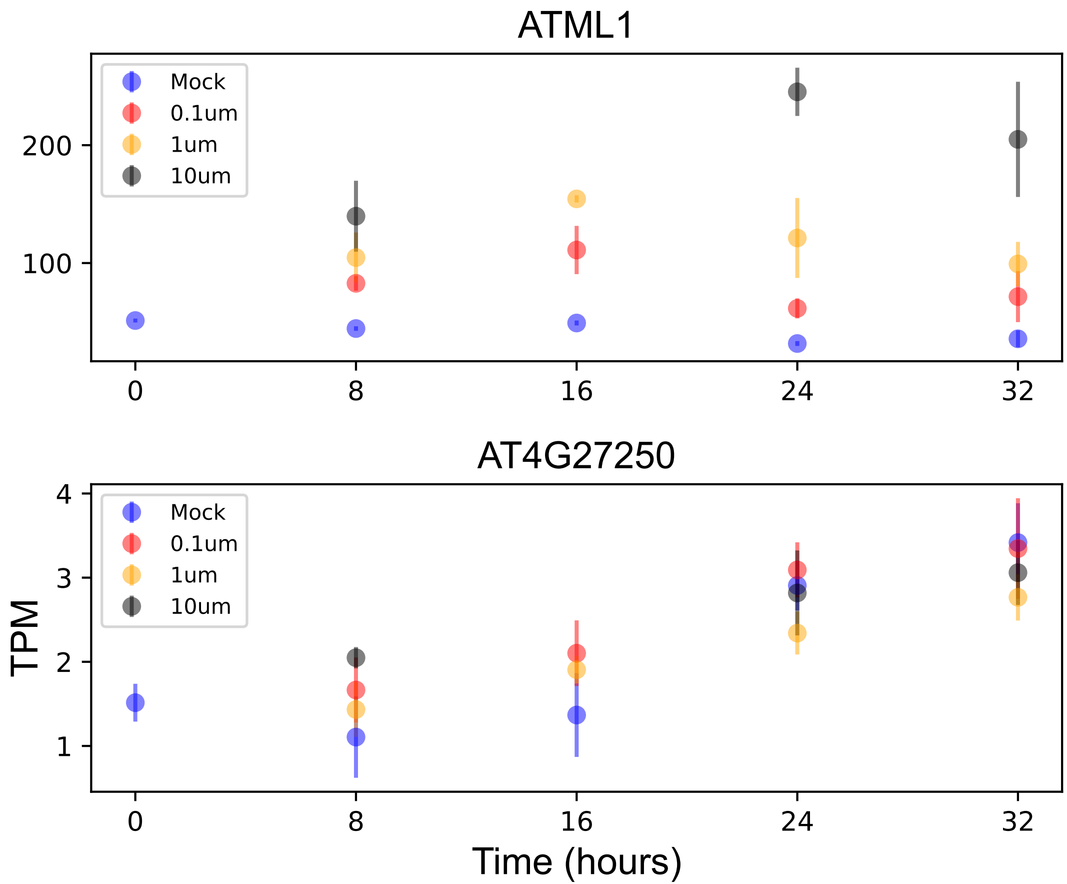

### AT4G29550.pdf

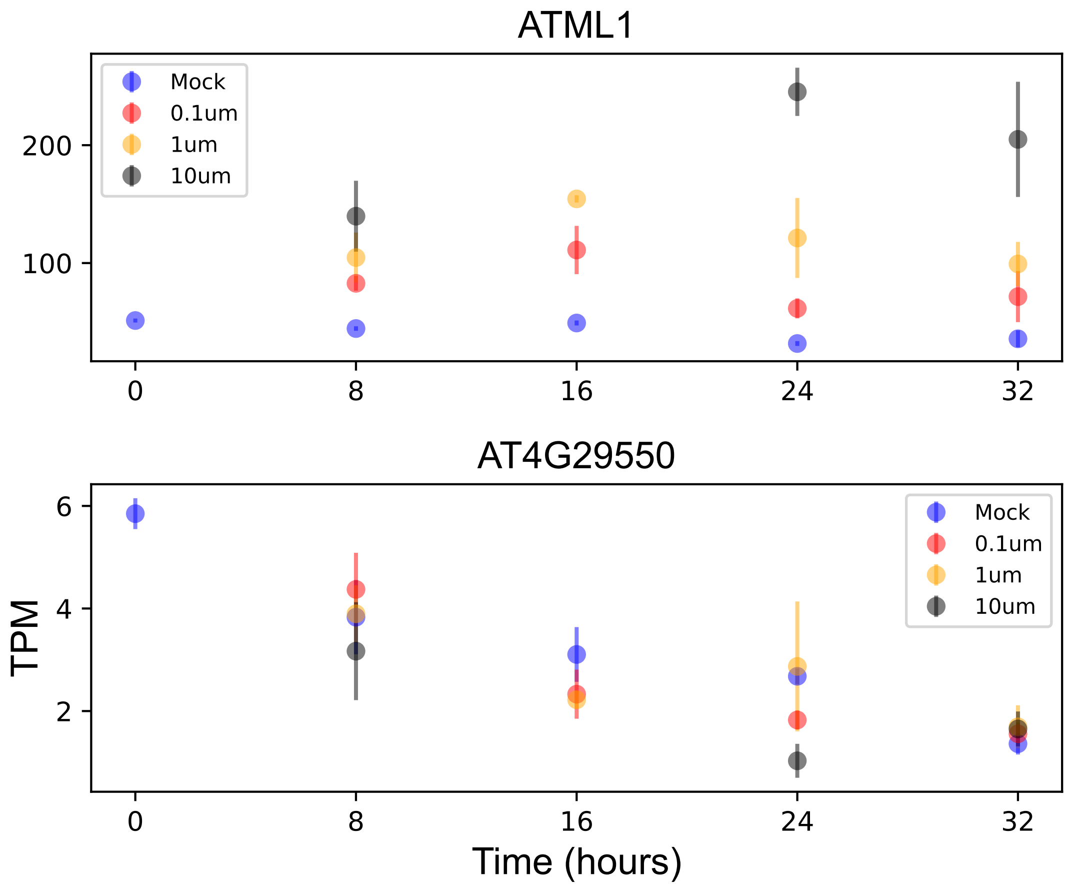

### AT4G33170.pdf

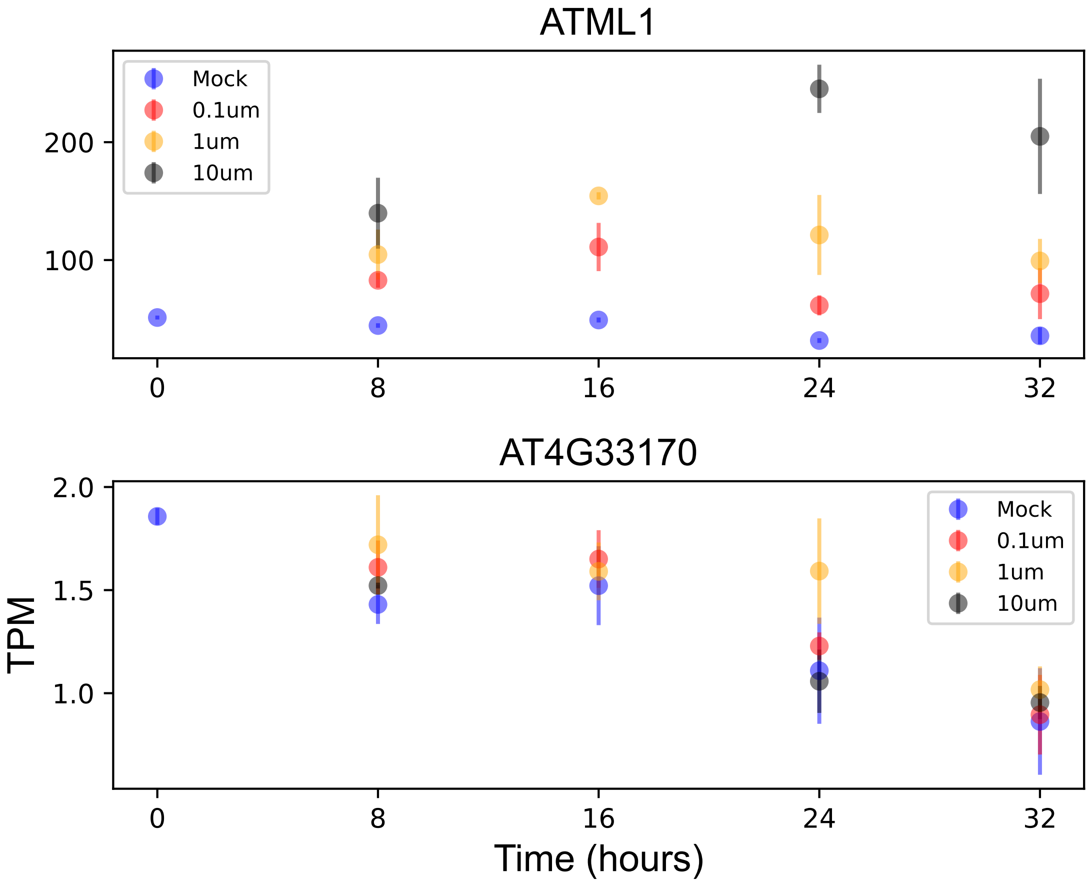

### AT4G33985.pdf

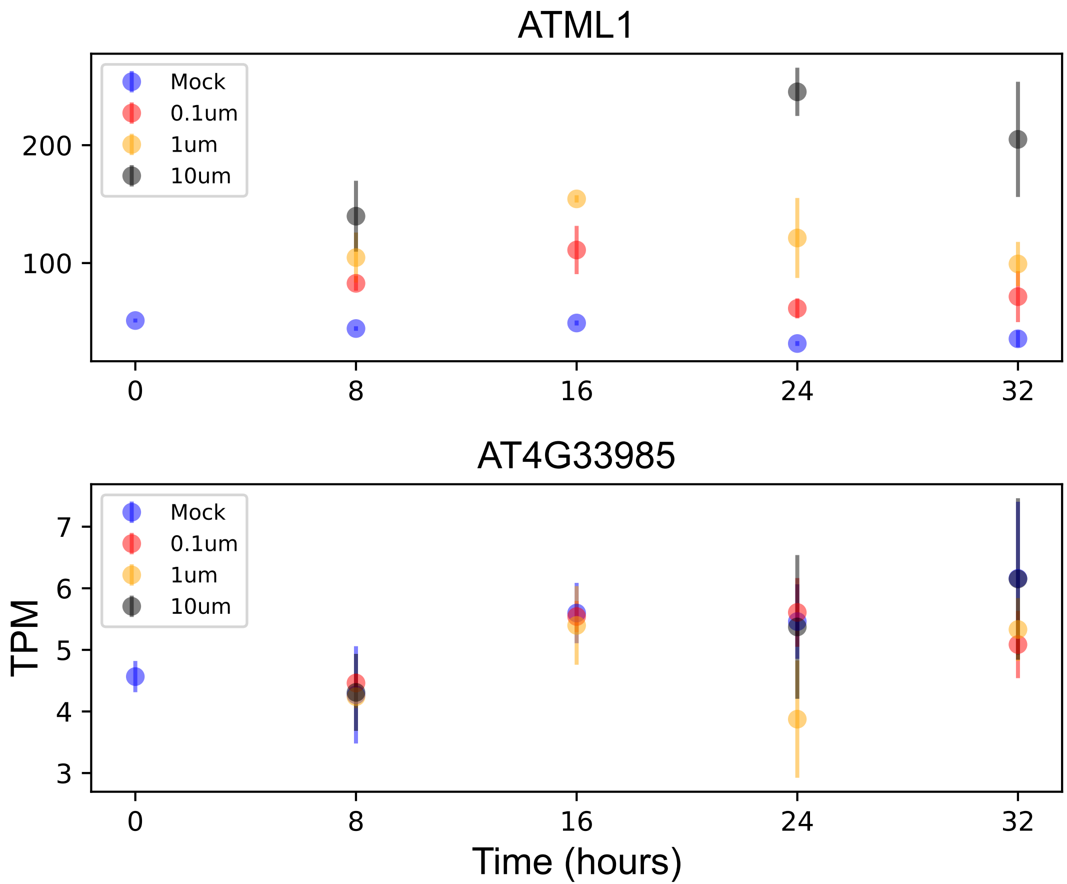

### AT5G05030.pdf

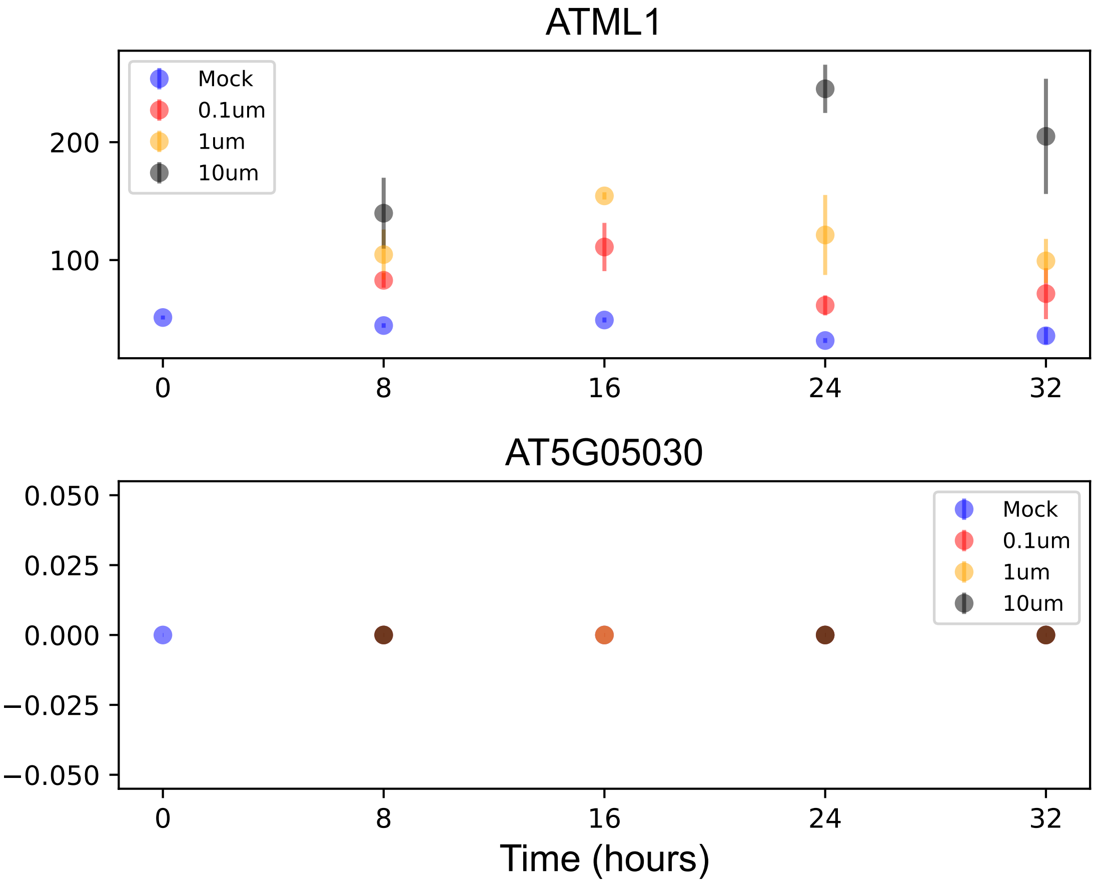

### AT5G06020.pdf

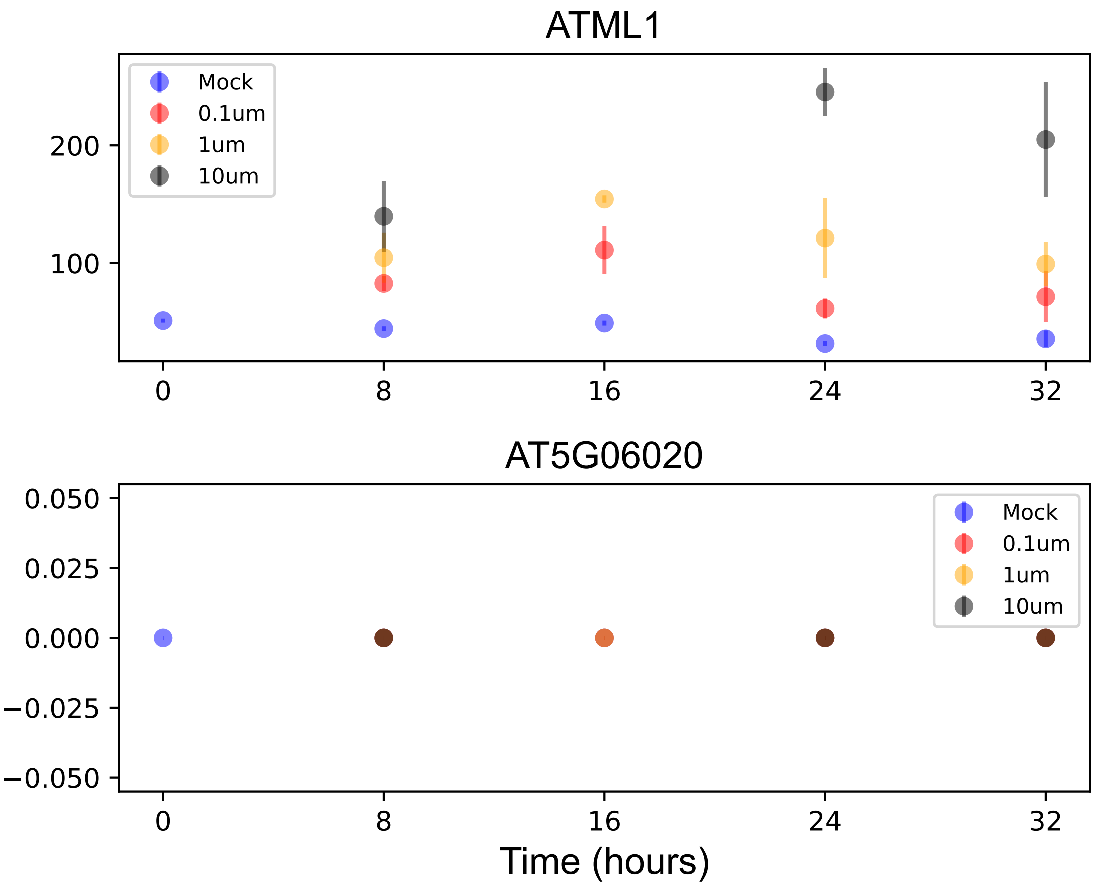

### AT5G06380.pdf

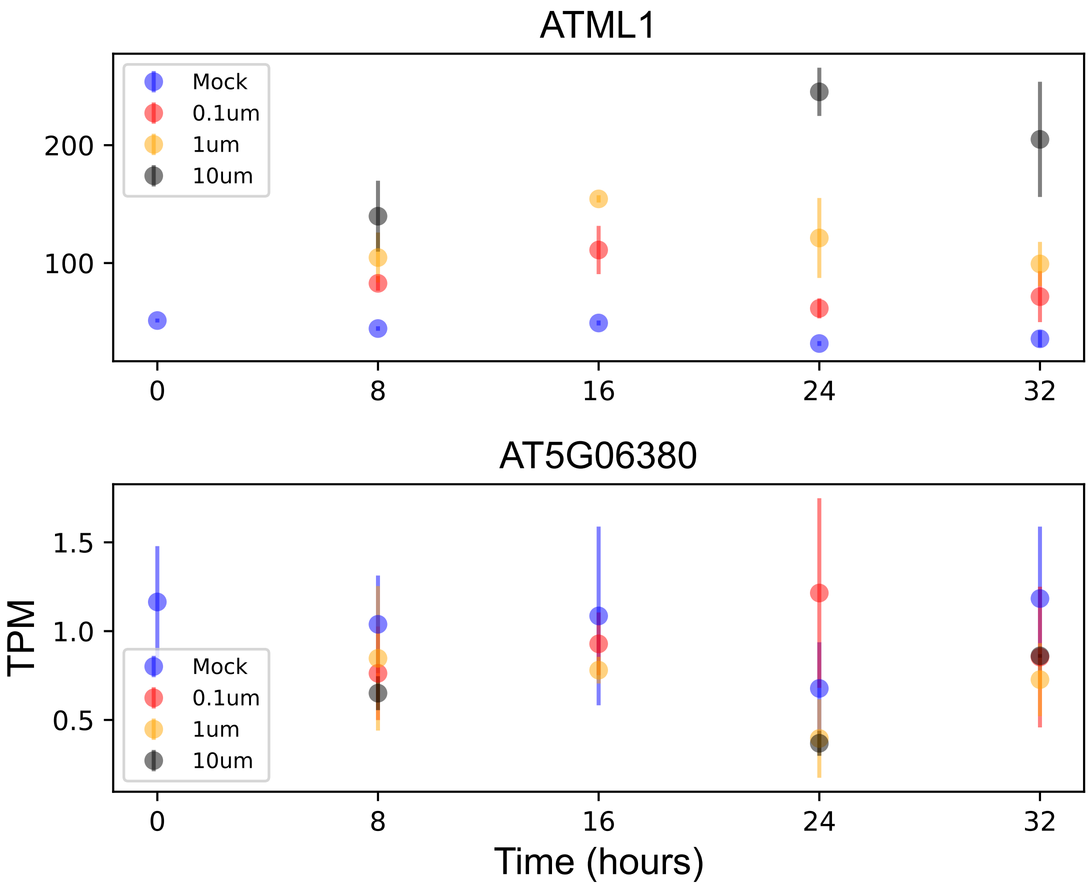

### AT5G10370.pdf

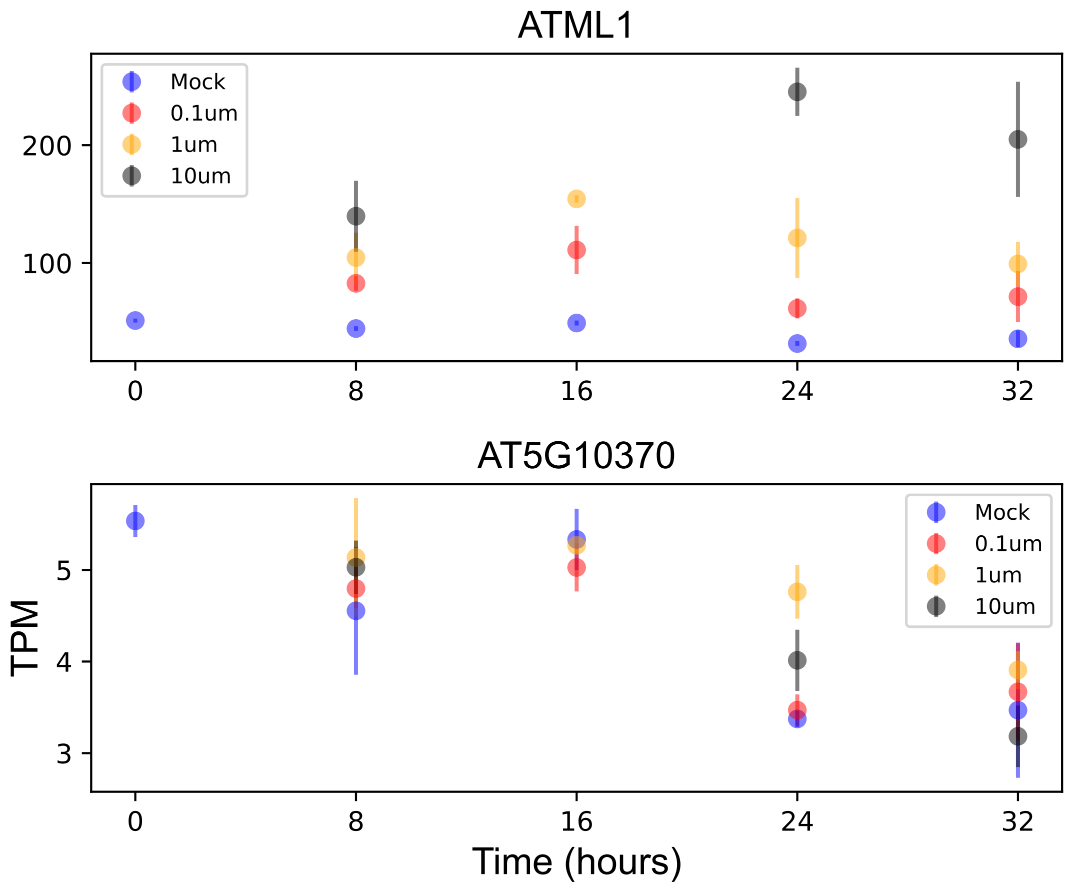

### AT5G14900.pdf

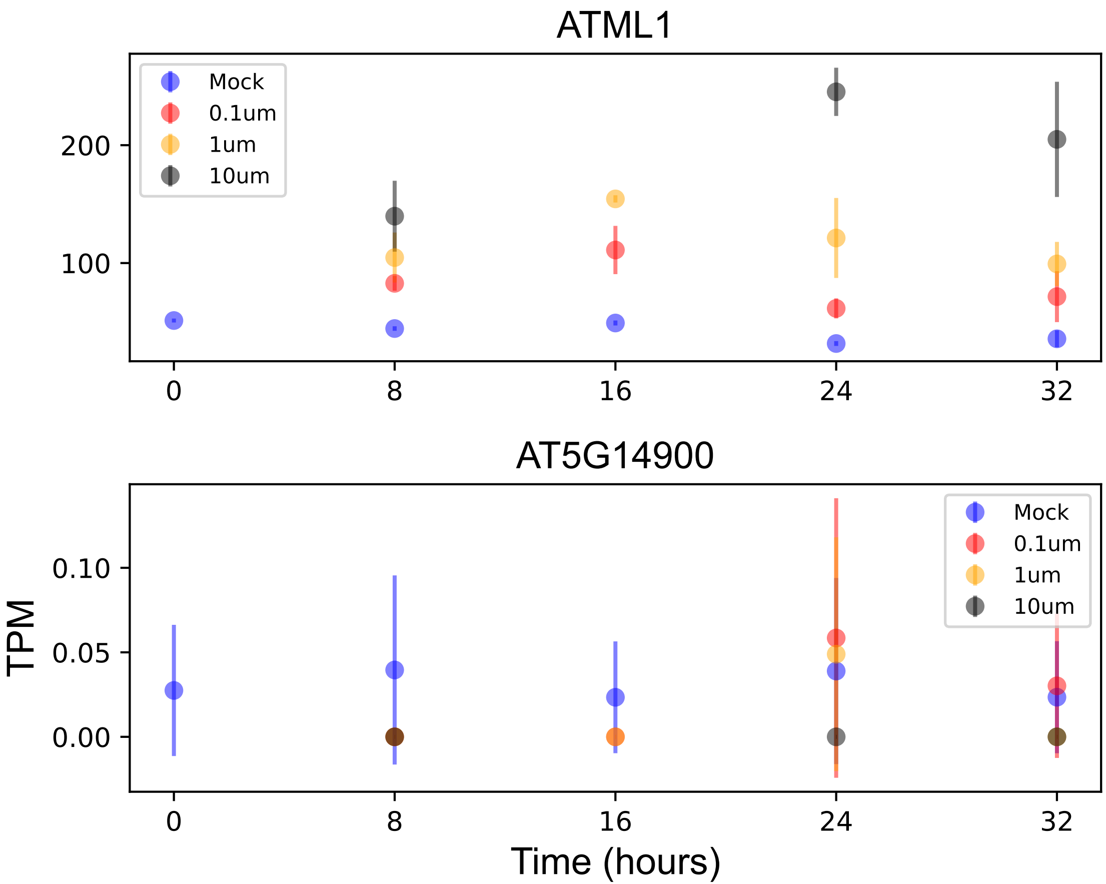

### AT5G15843.pdf

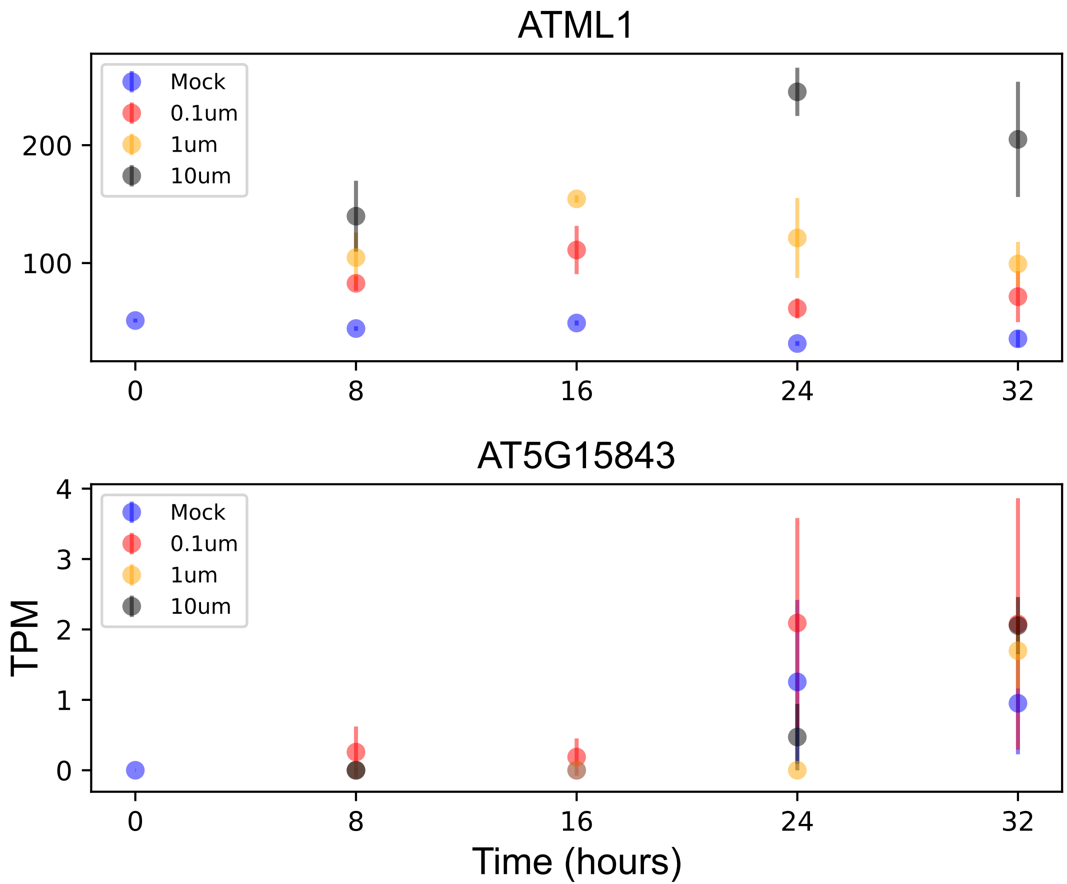

### AT5G17930.pdf

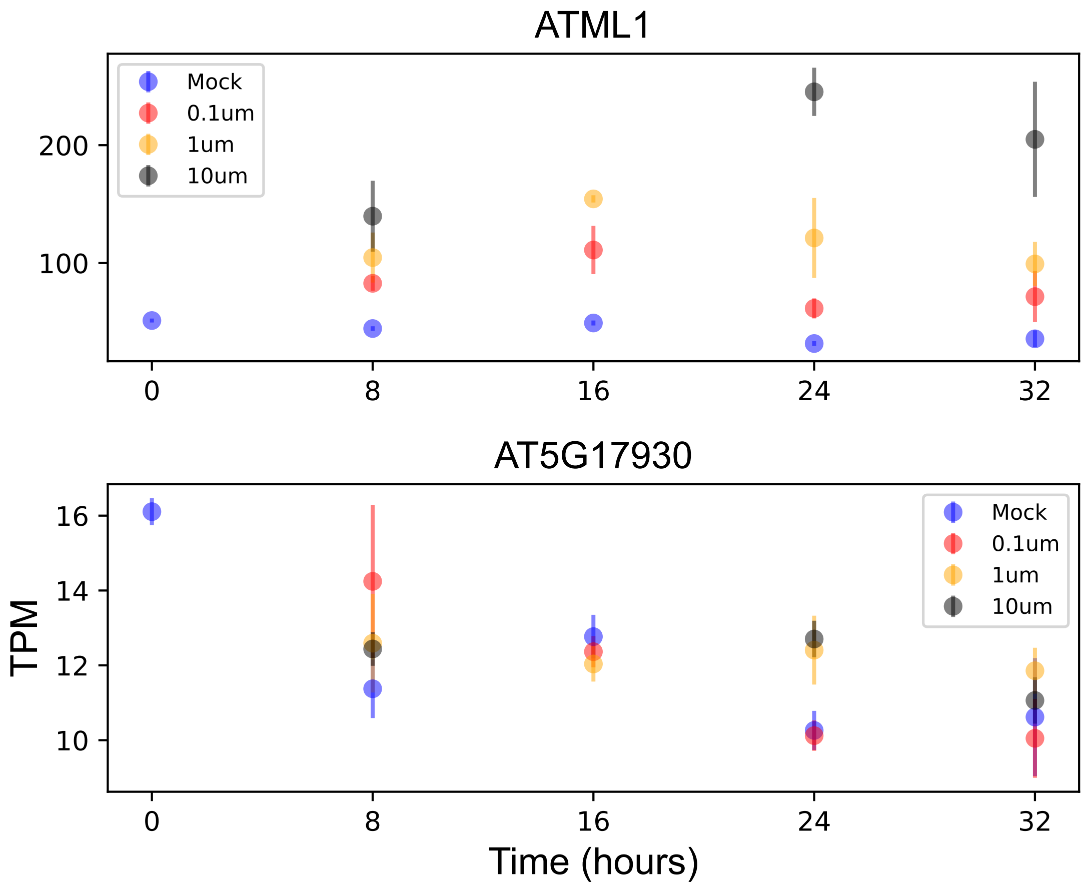

### AT5G59210.pdf

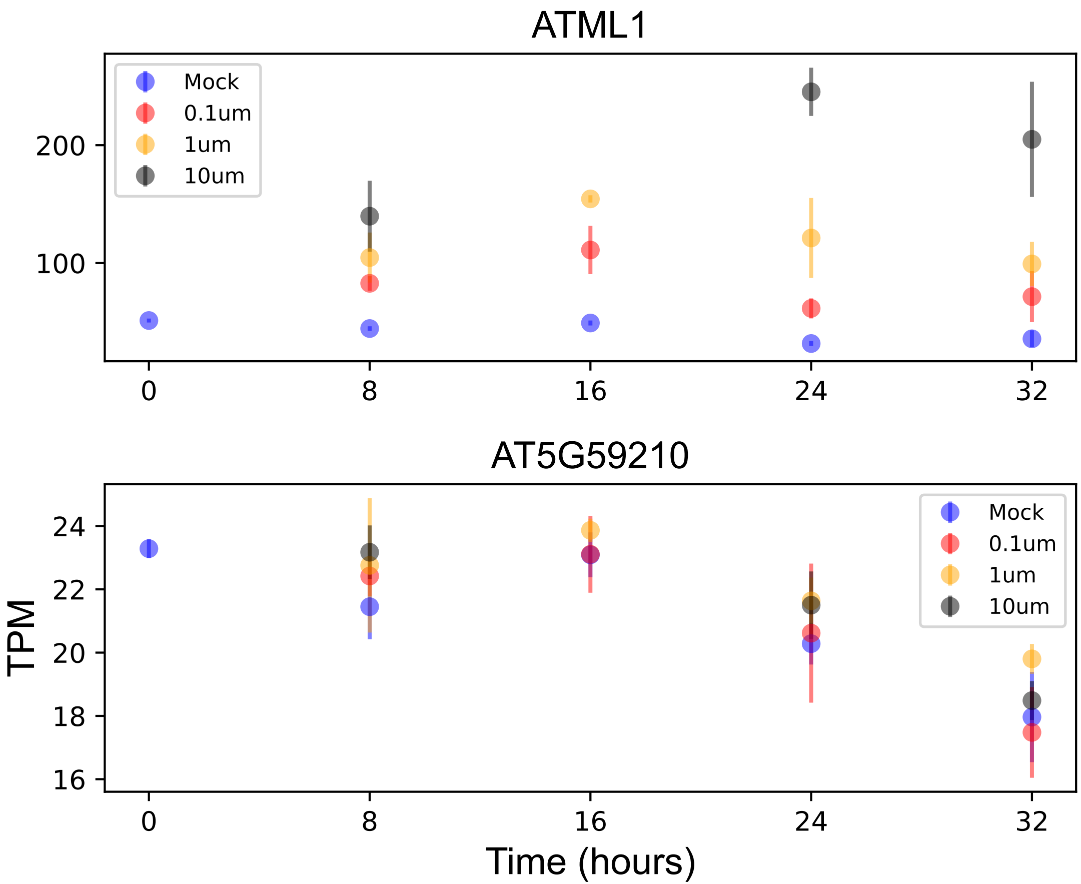

### AT5G62210.pdf

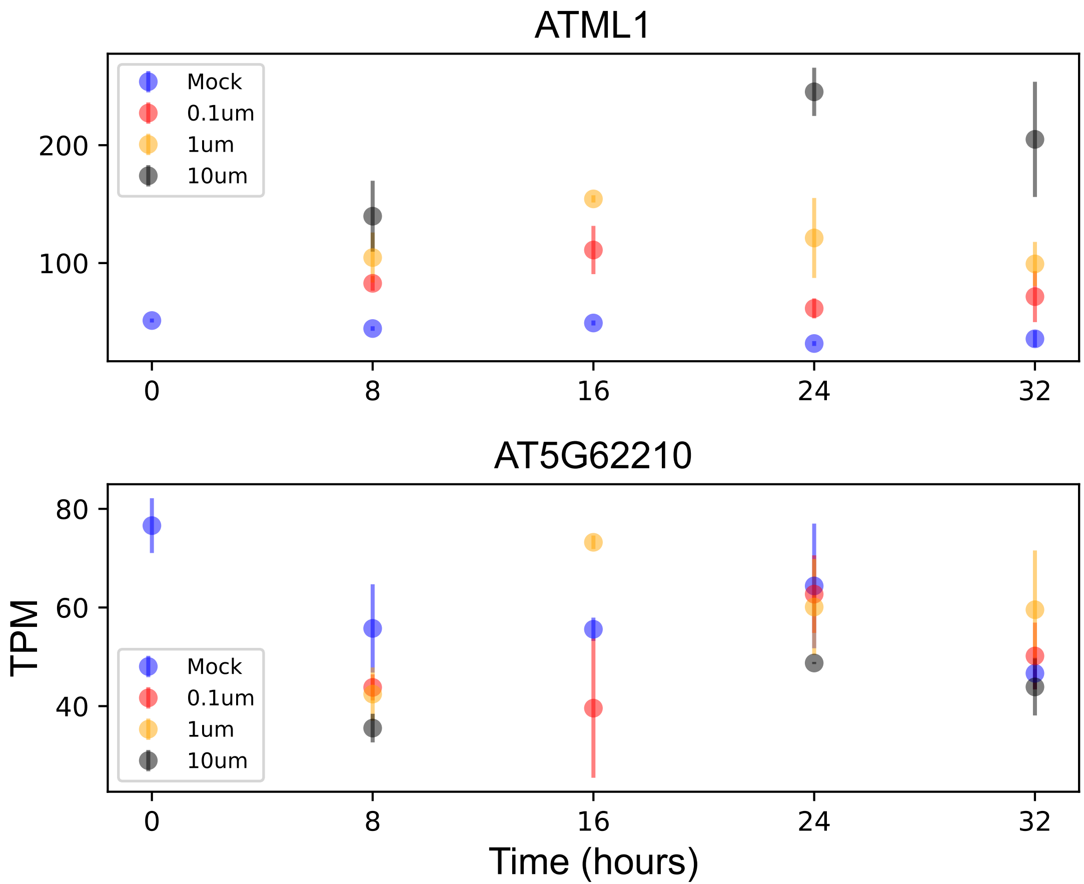

### AT-HSFA5.pdf

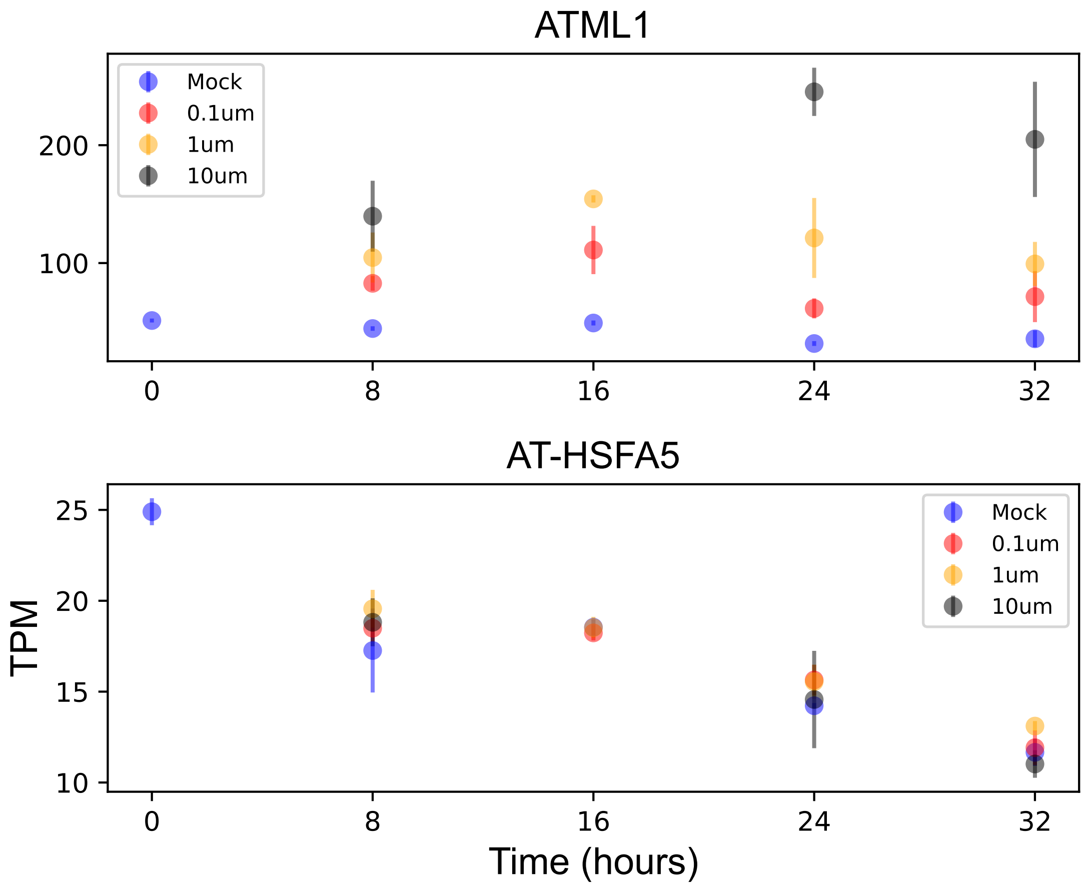

### ATARD2.pdf

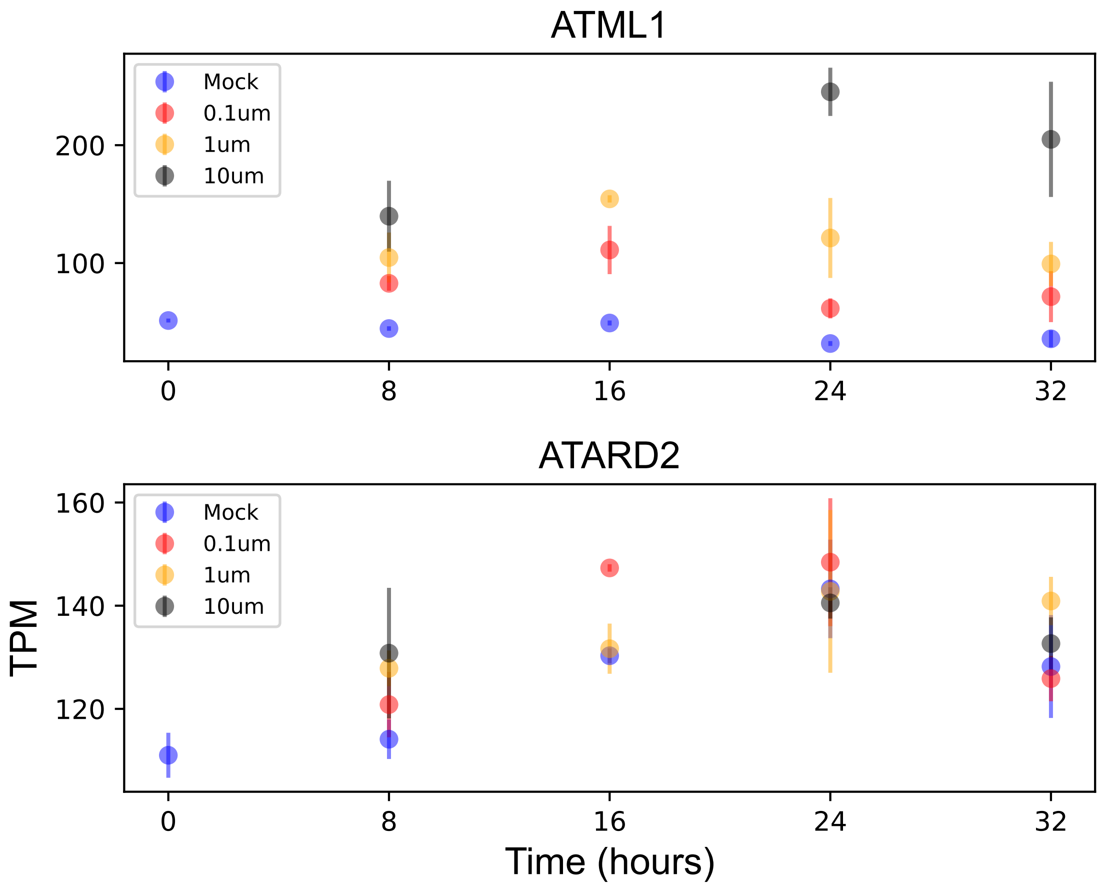

### ATO.pdf

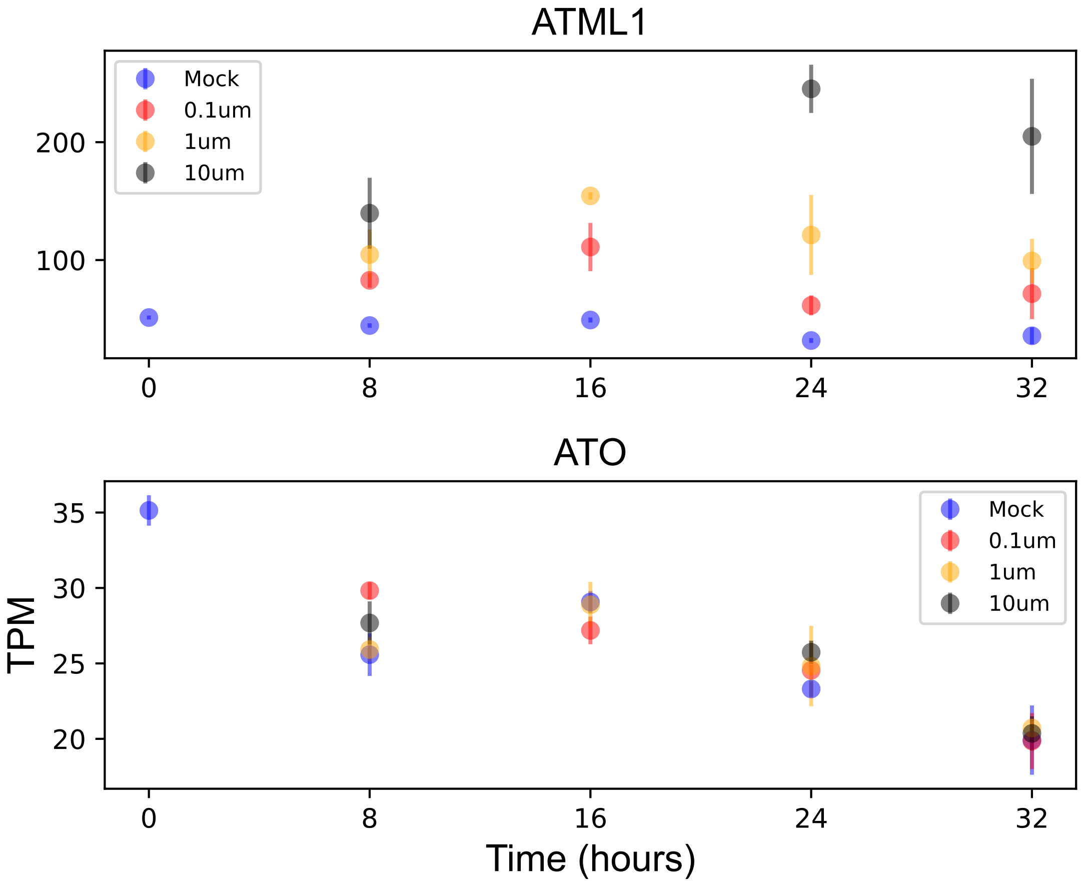

### CFM4.pdf

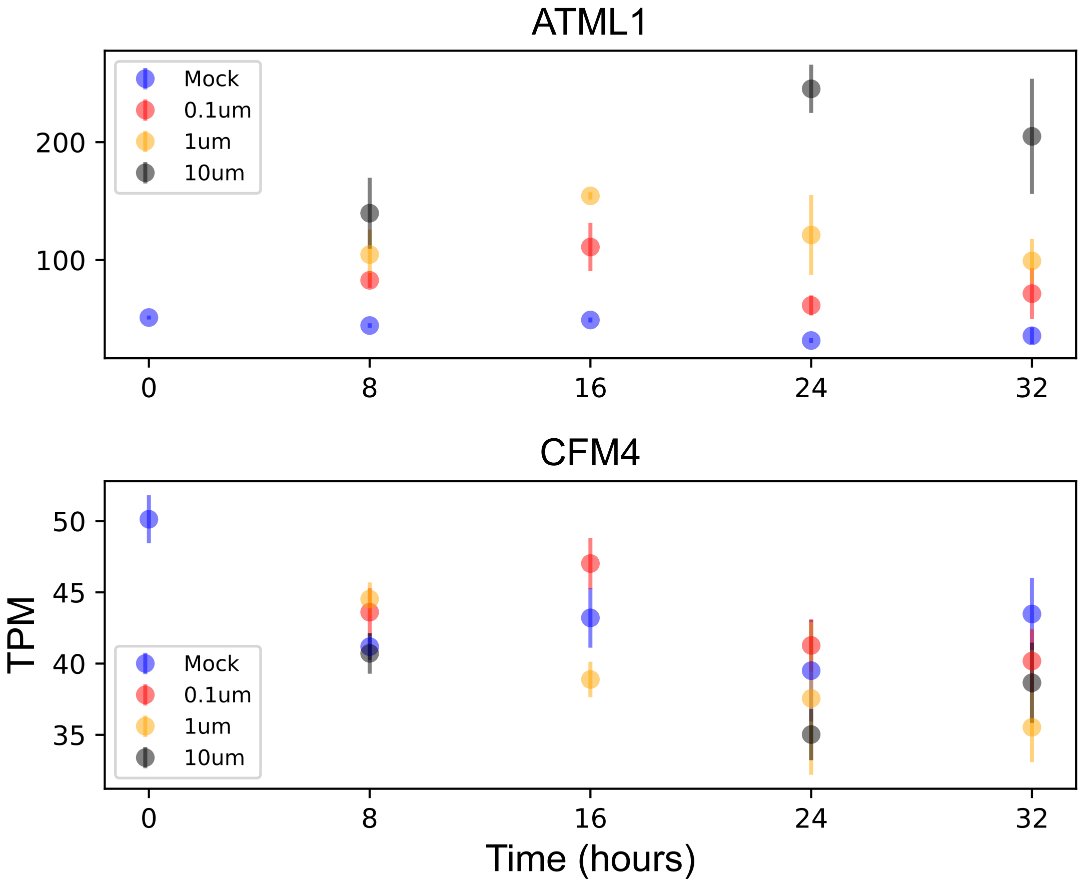

### ftsh7.pdf

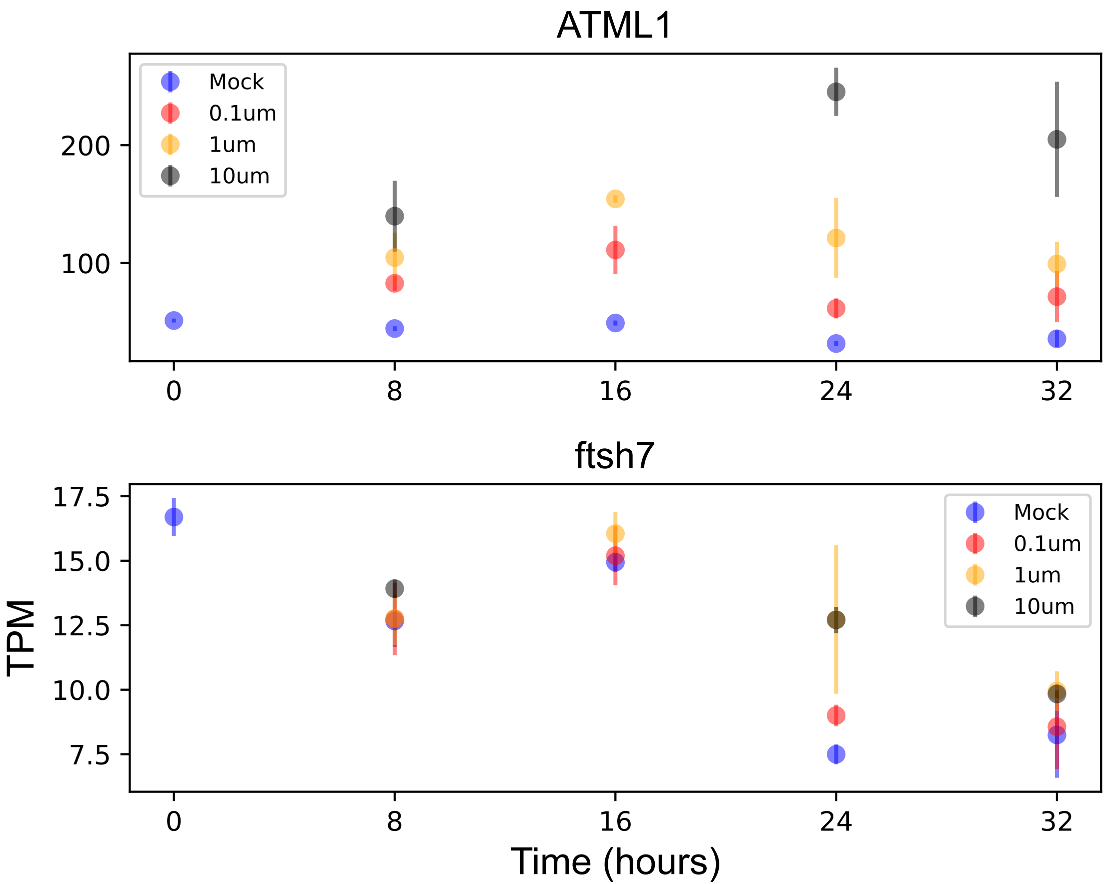

### PAP26.pdf

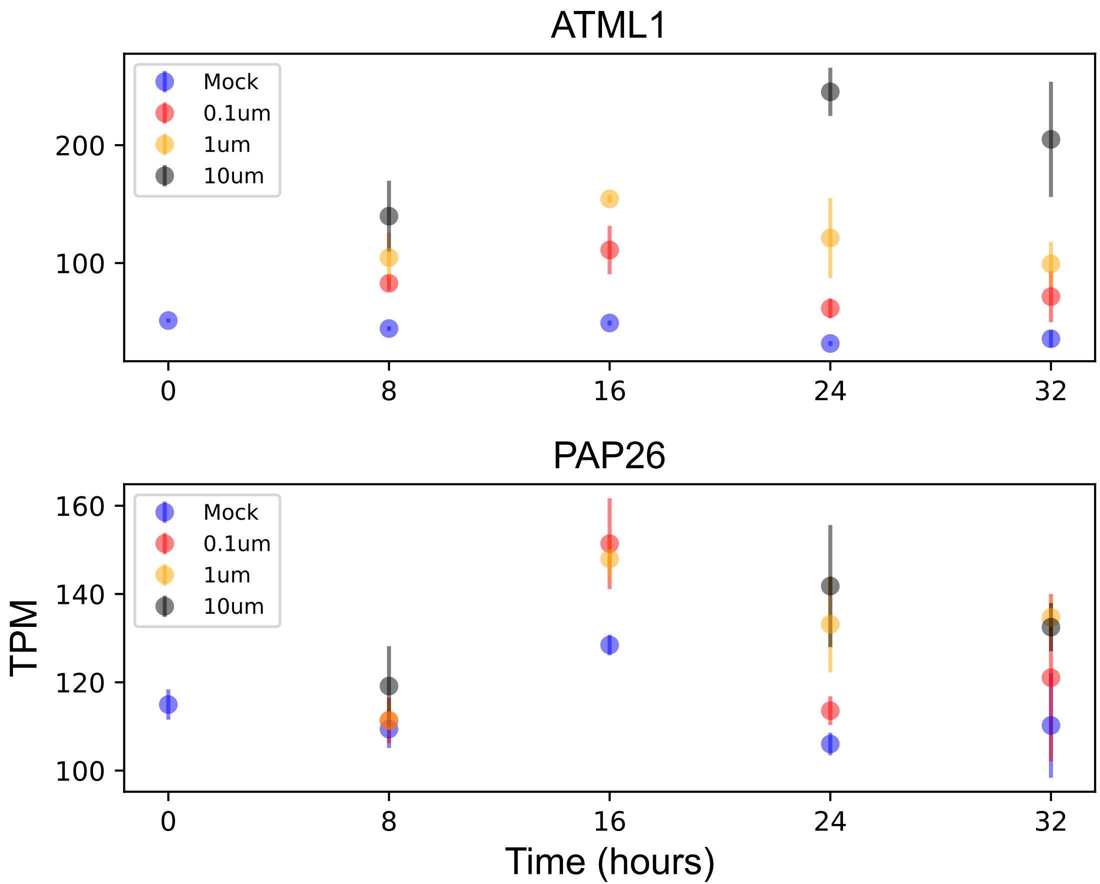

### PAS2.pdf

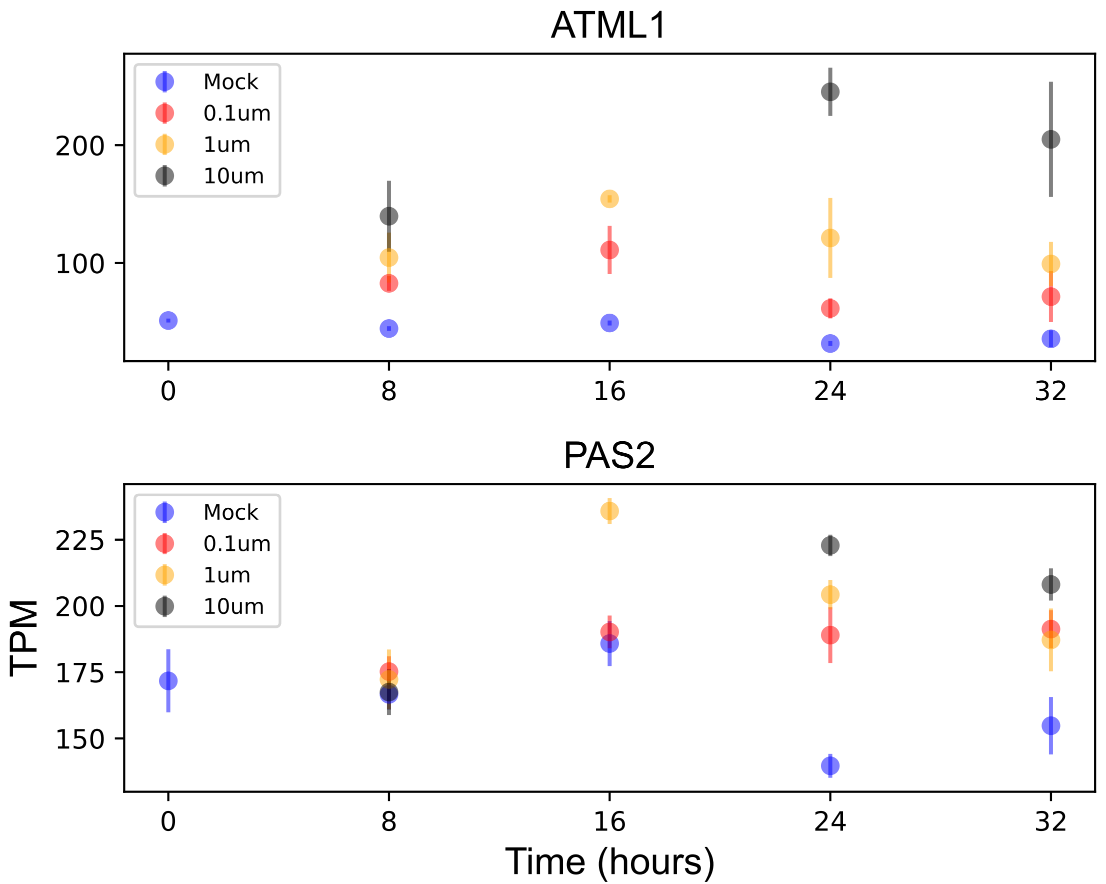

### PBP1.pdf

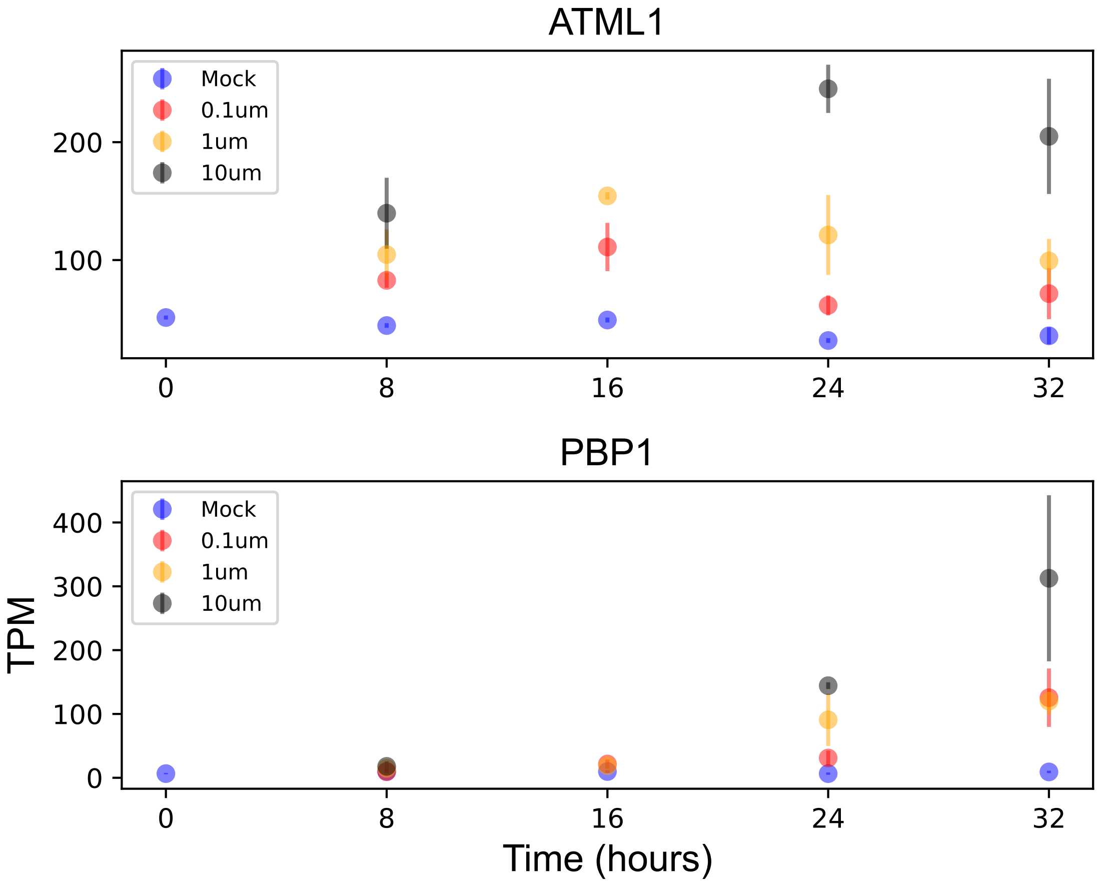

### PLP9.pdf

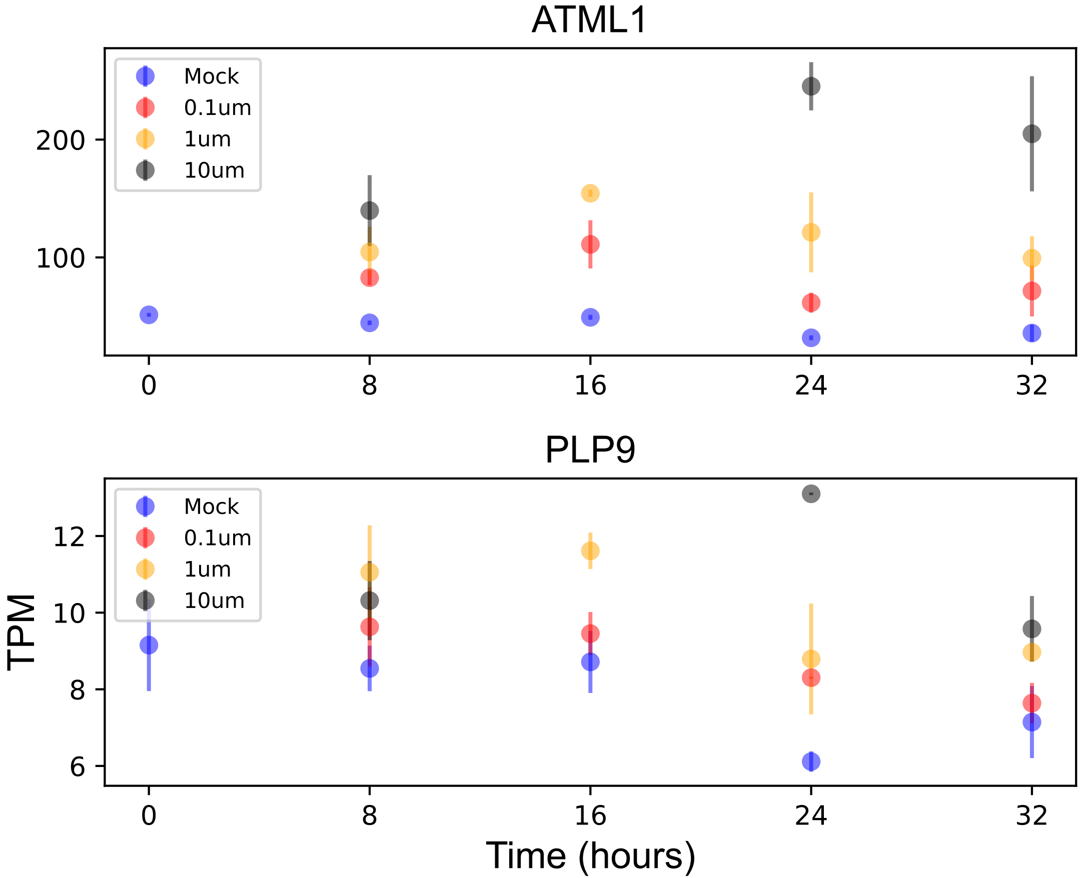

### PME44.pdf

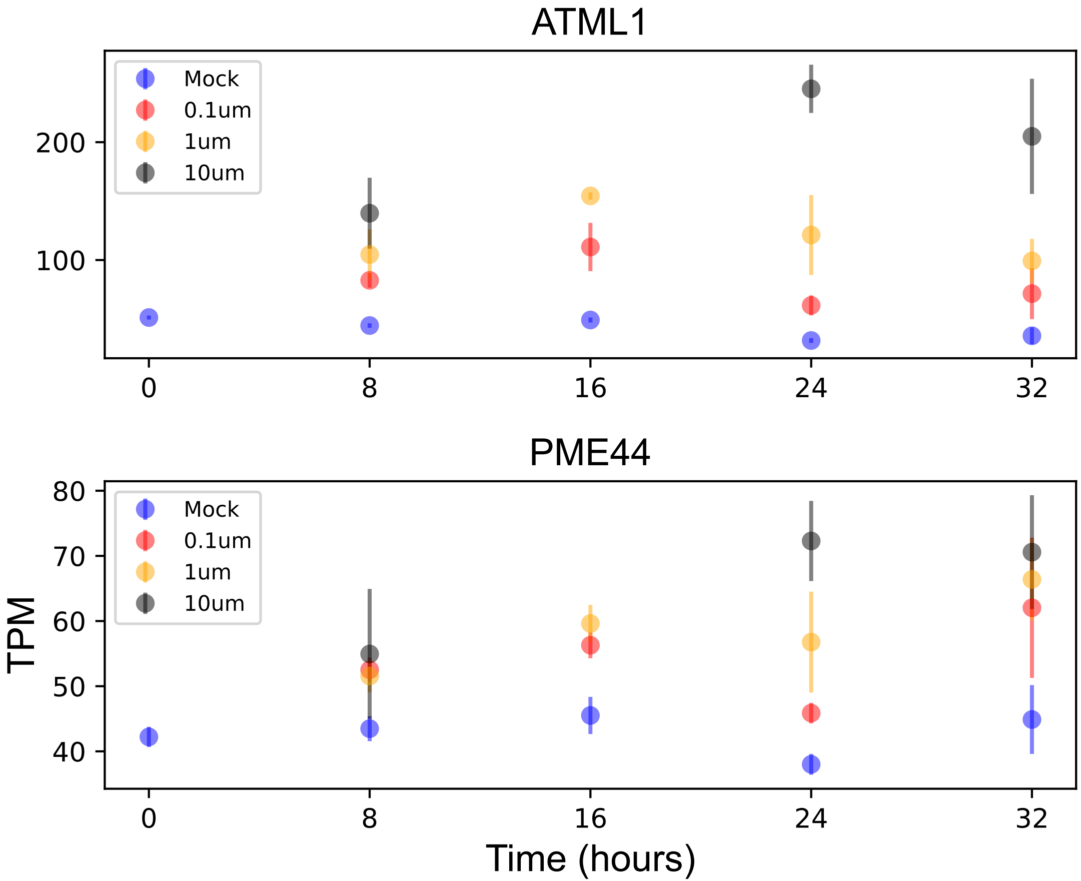

### PnsB3.pdf

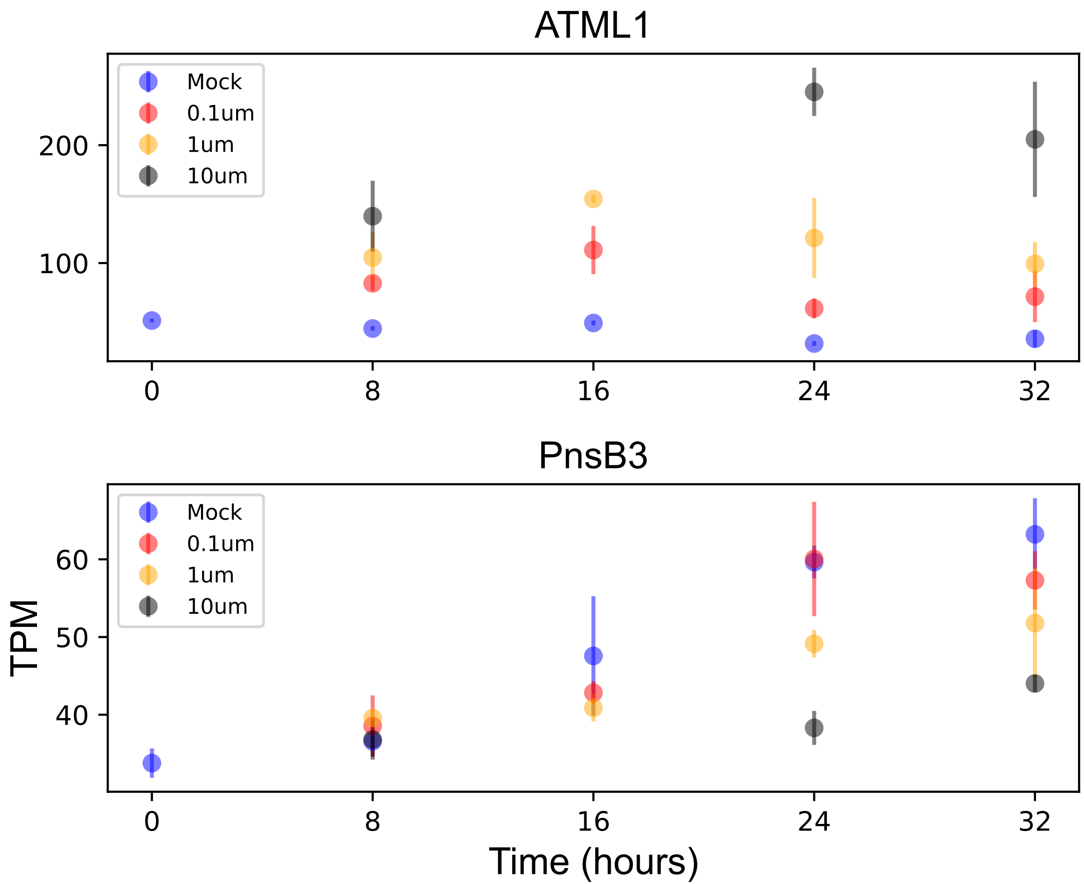

### PnsL4.pdf

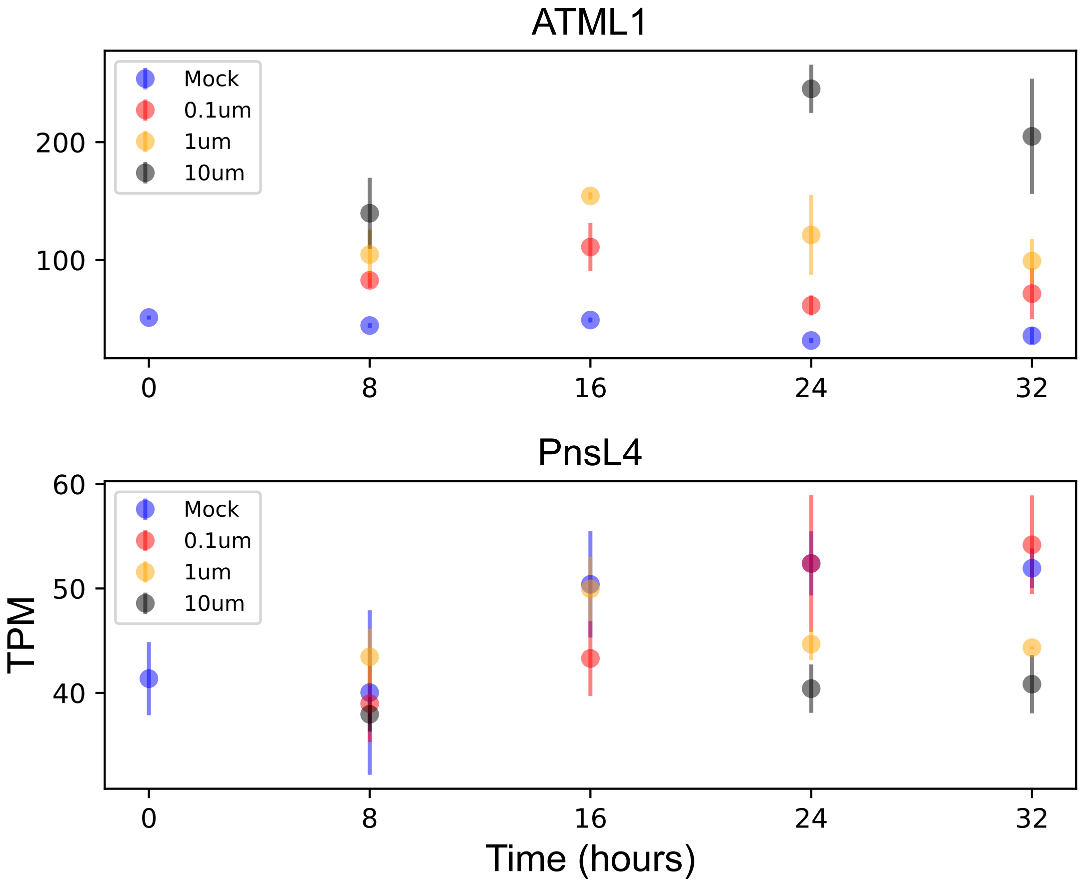

### PRIN2.pdf

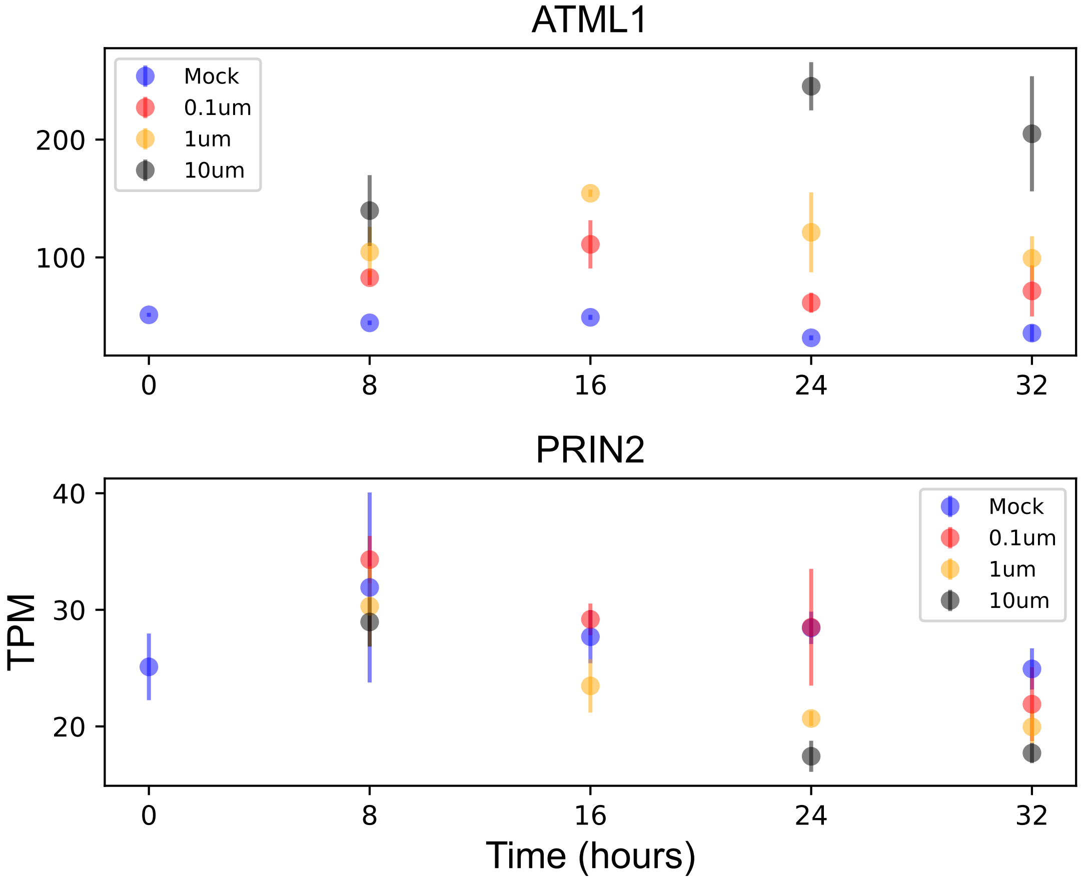

### PRXR1.pdf

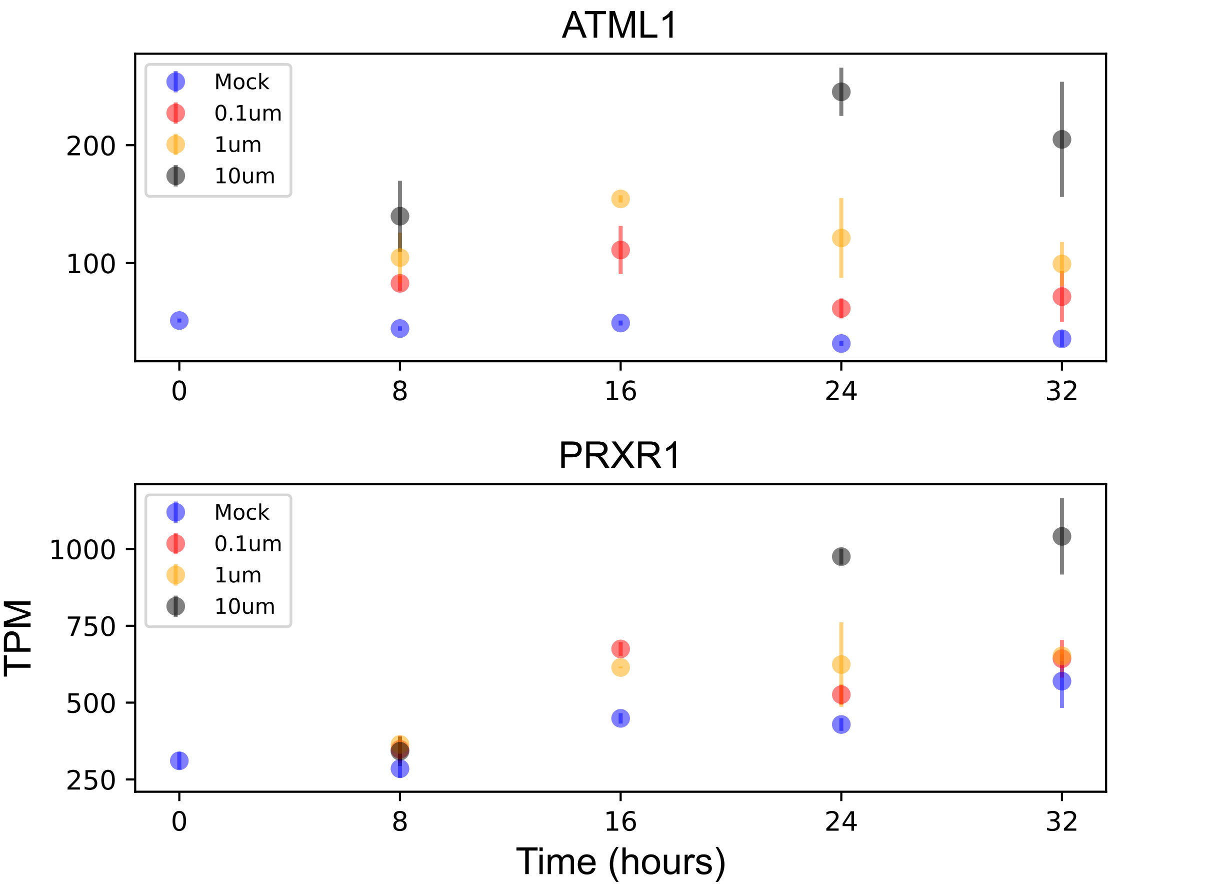

### PTAC7.pdf

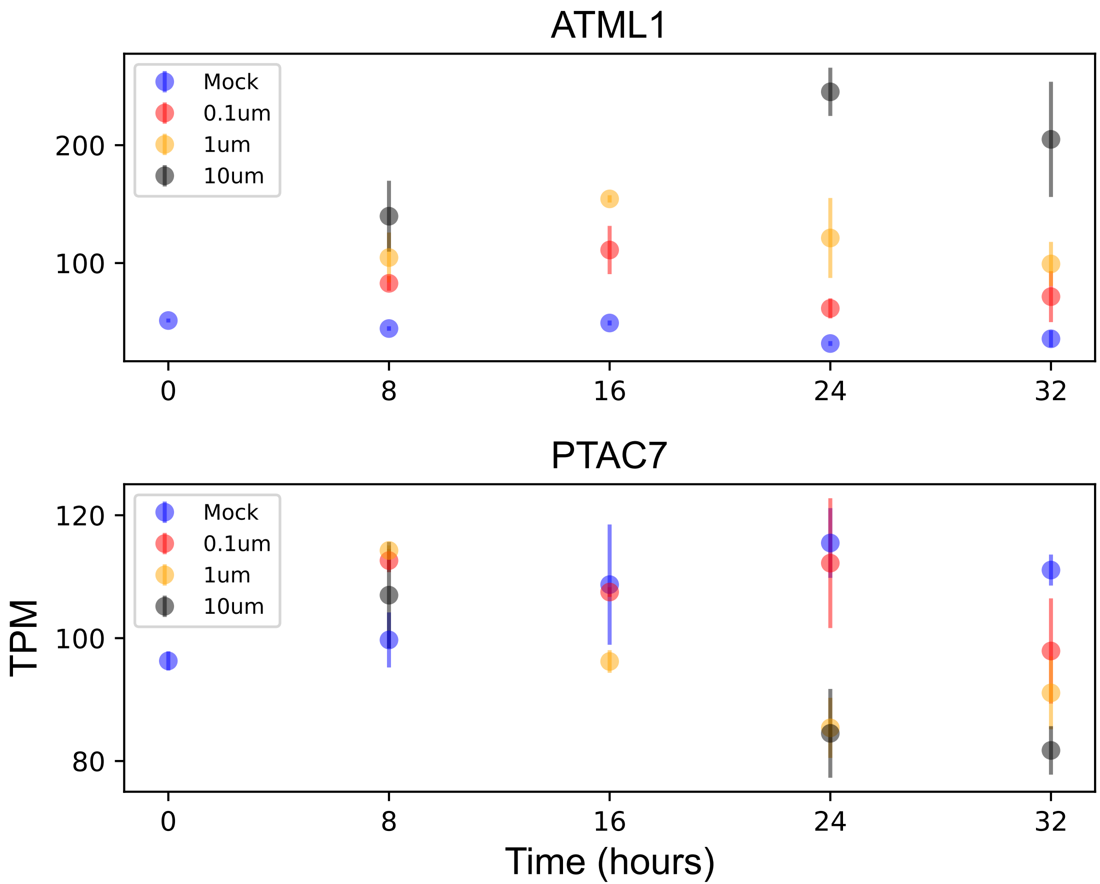

### PYD1.pdf

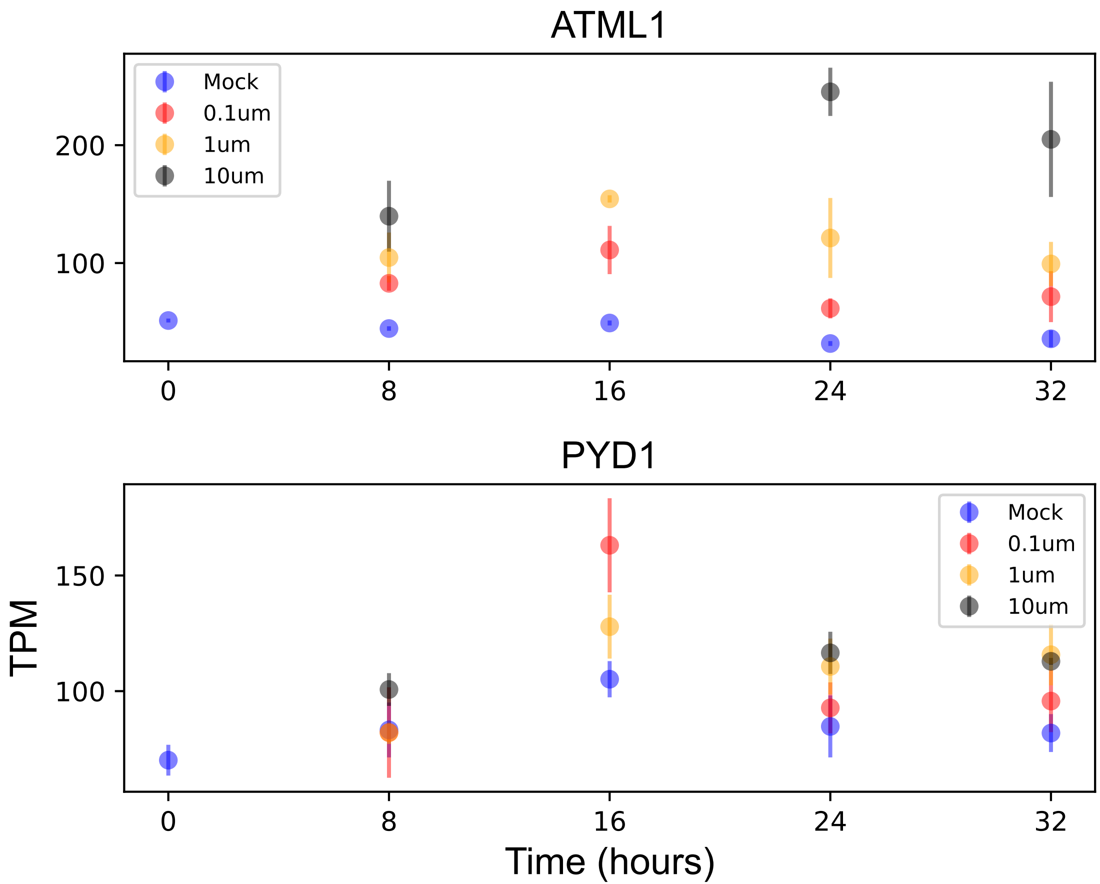

### RBCS1B.pdf

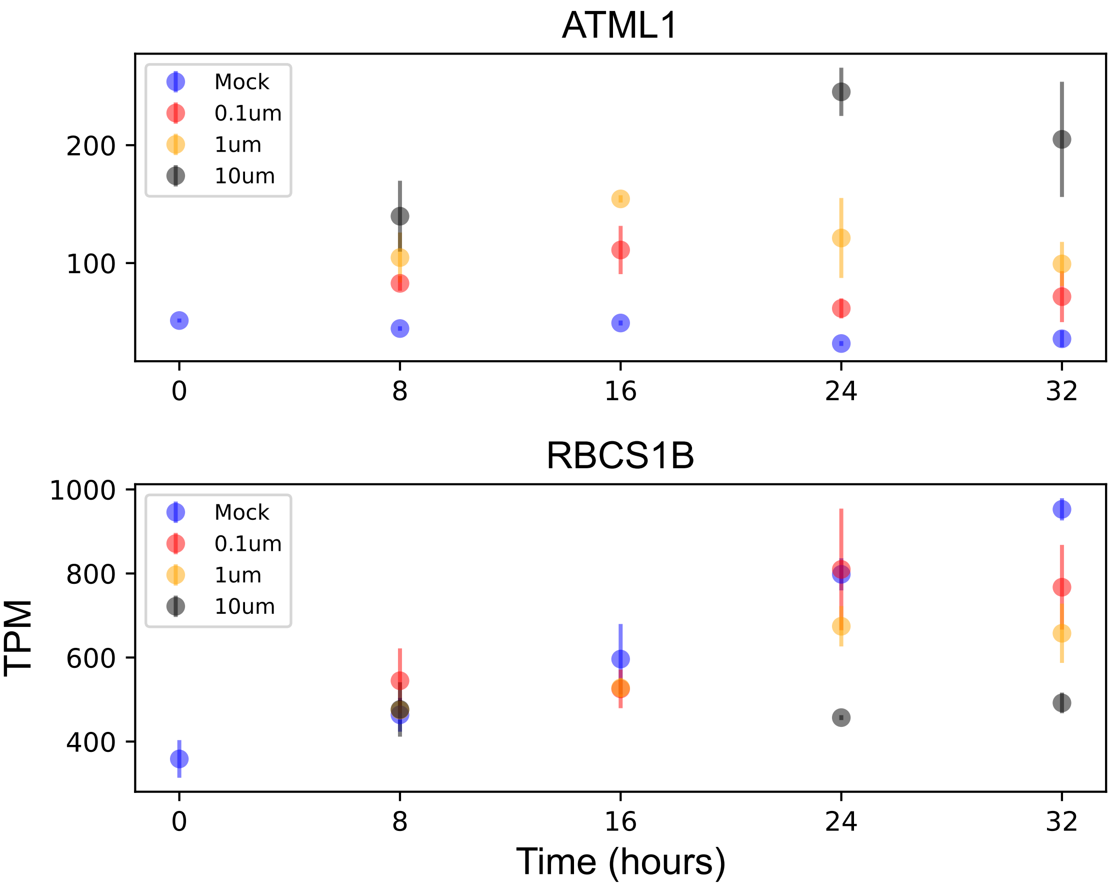

### RGXT2.pdf

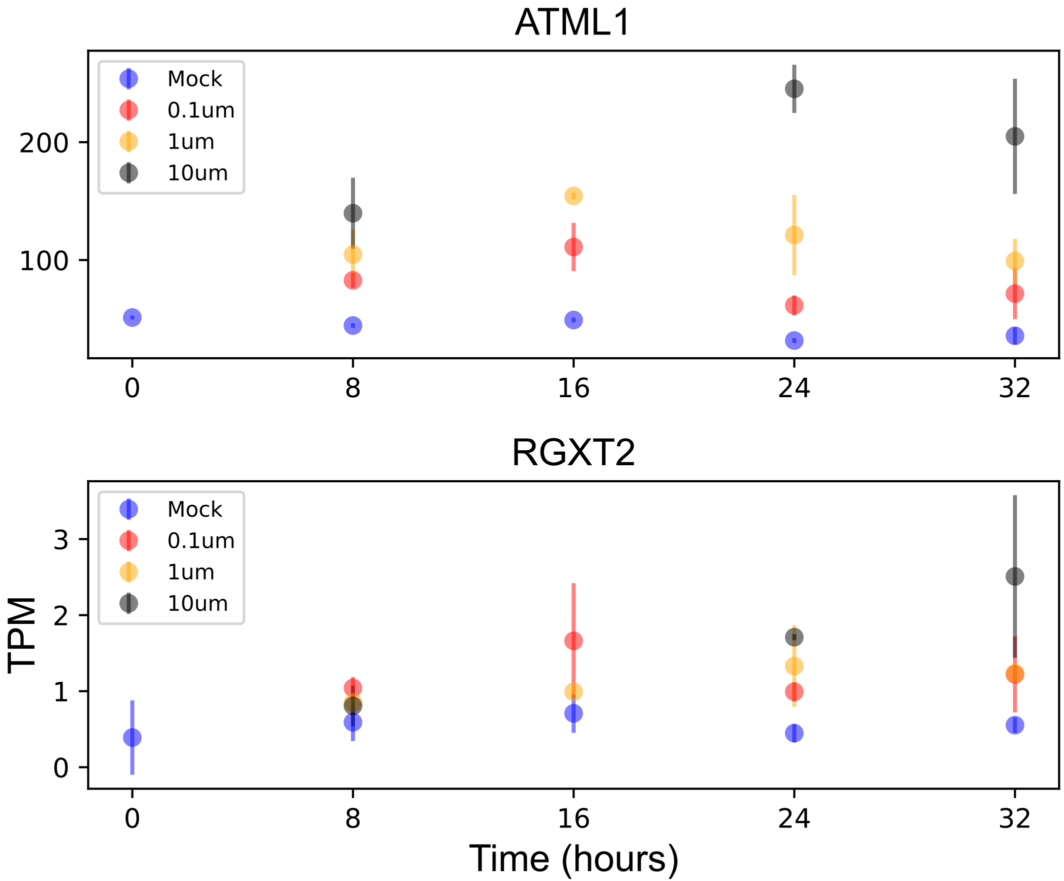

### RPL16.pdf

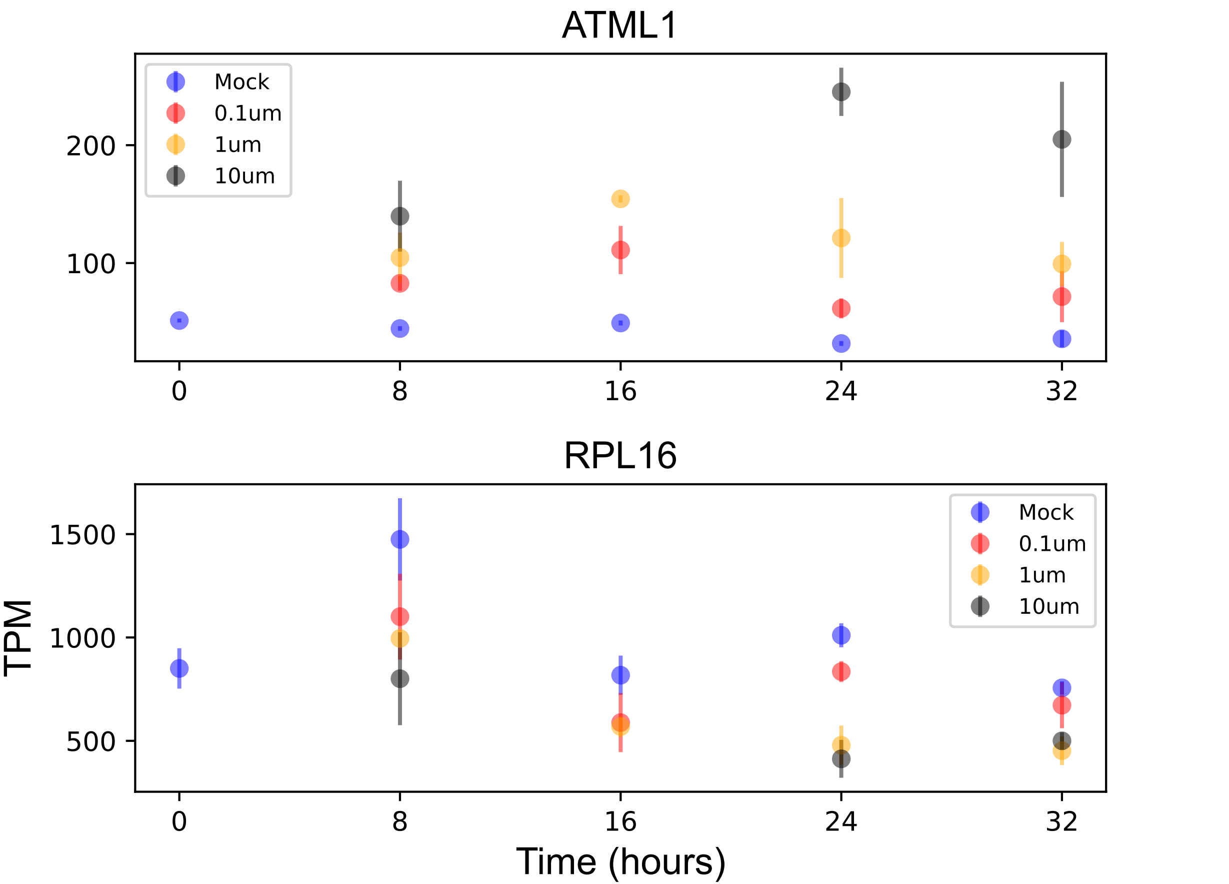

### RPL22.pdf

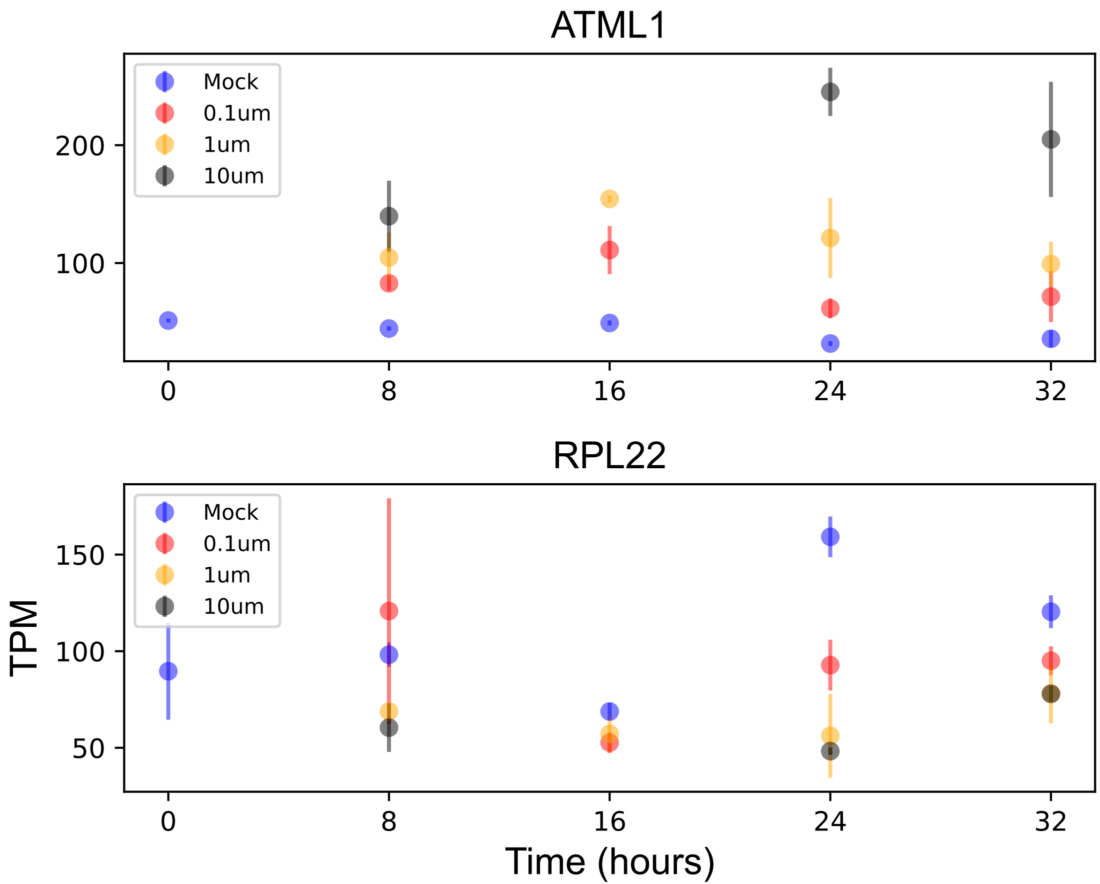

### RPS2.pdf

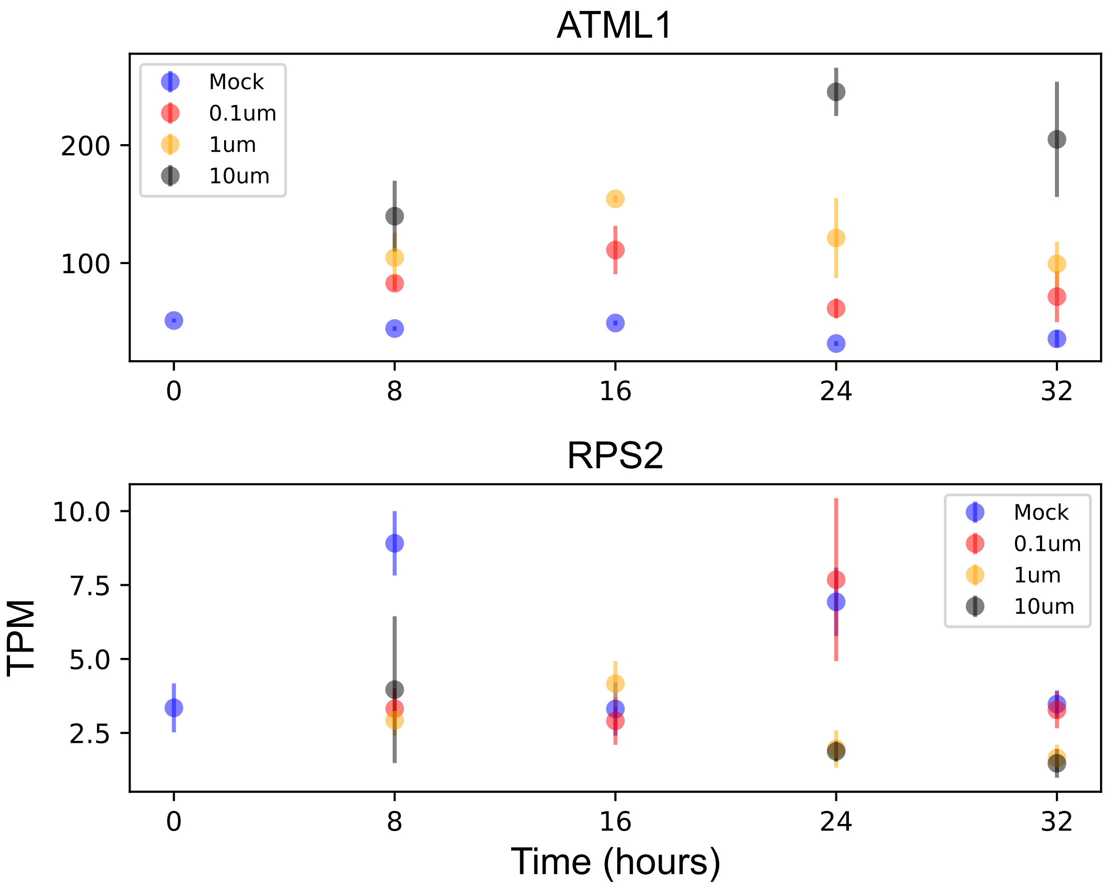

### RPS3.pdf

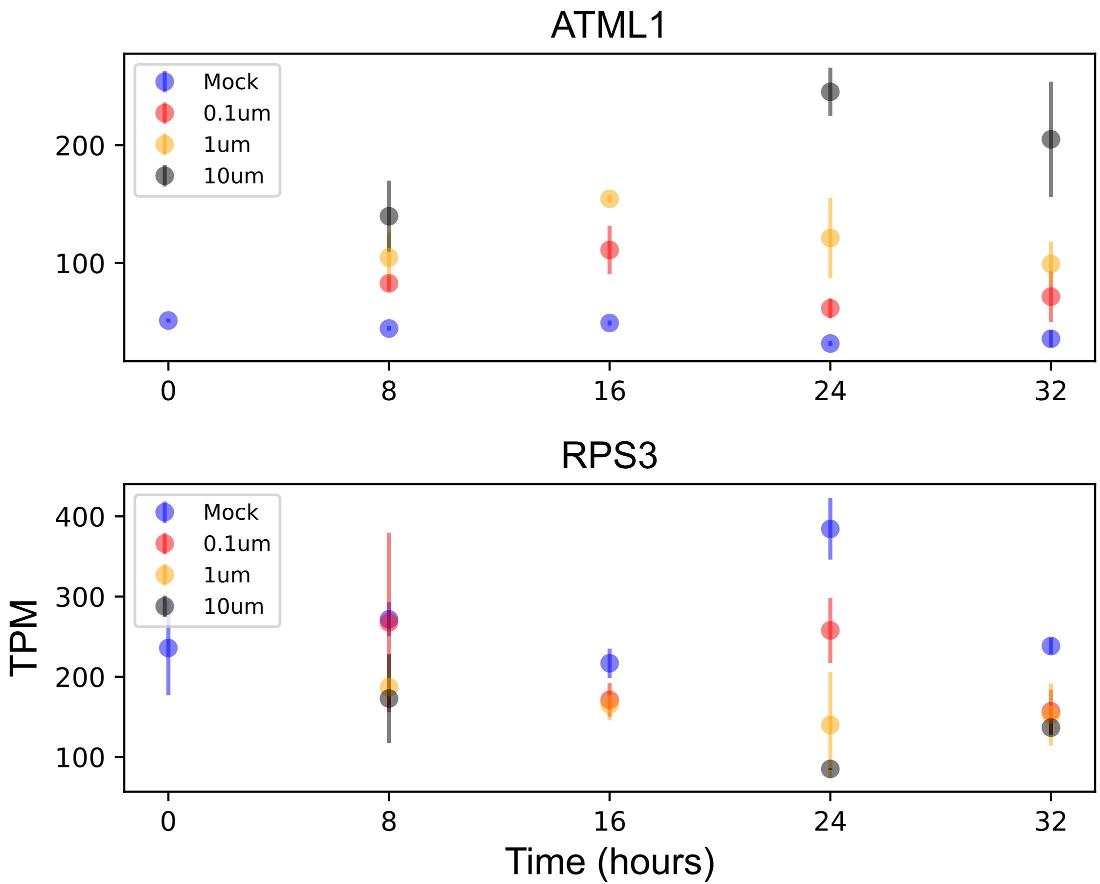

### RPS4.pdf

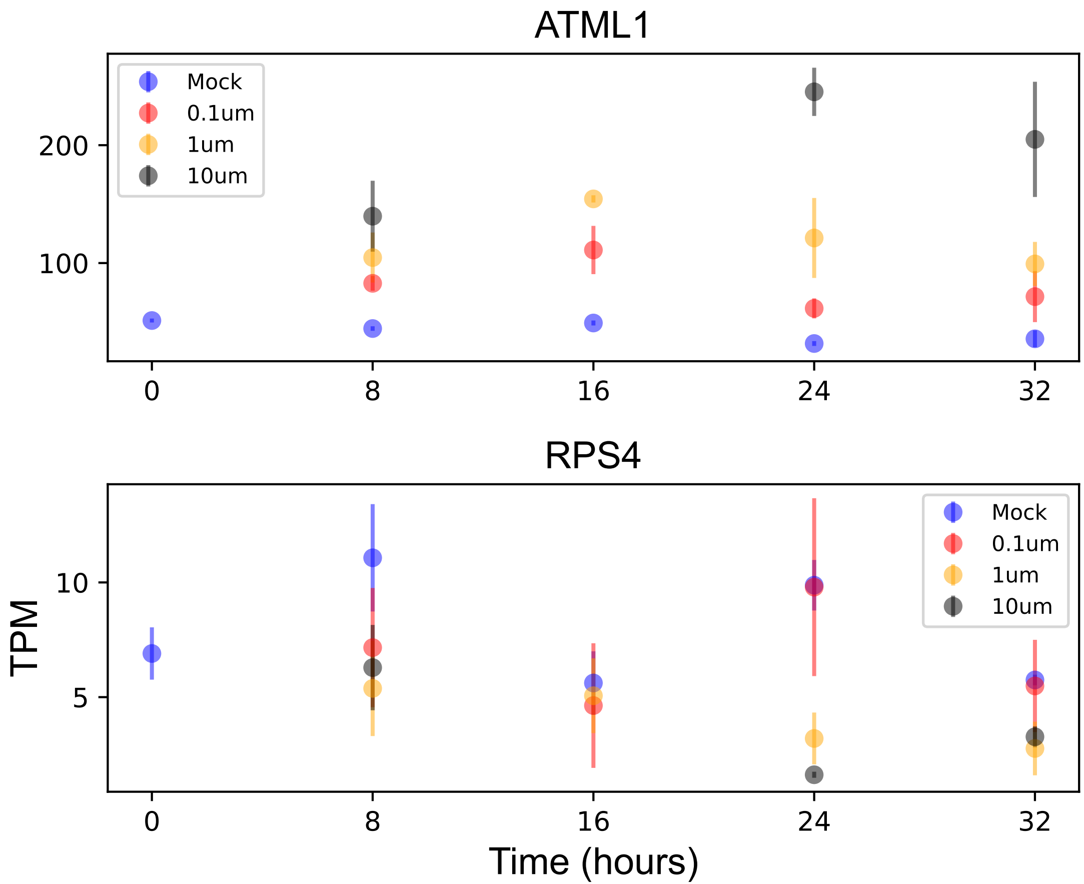

### RPS7.1.pdf

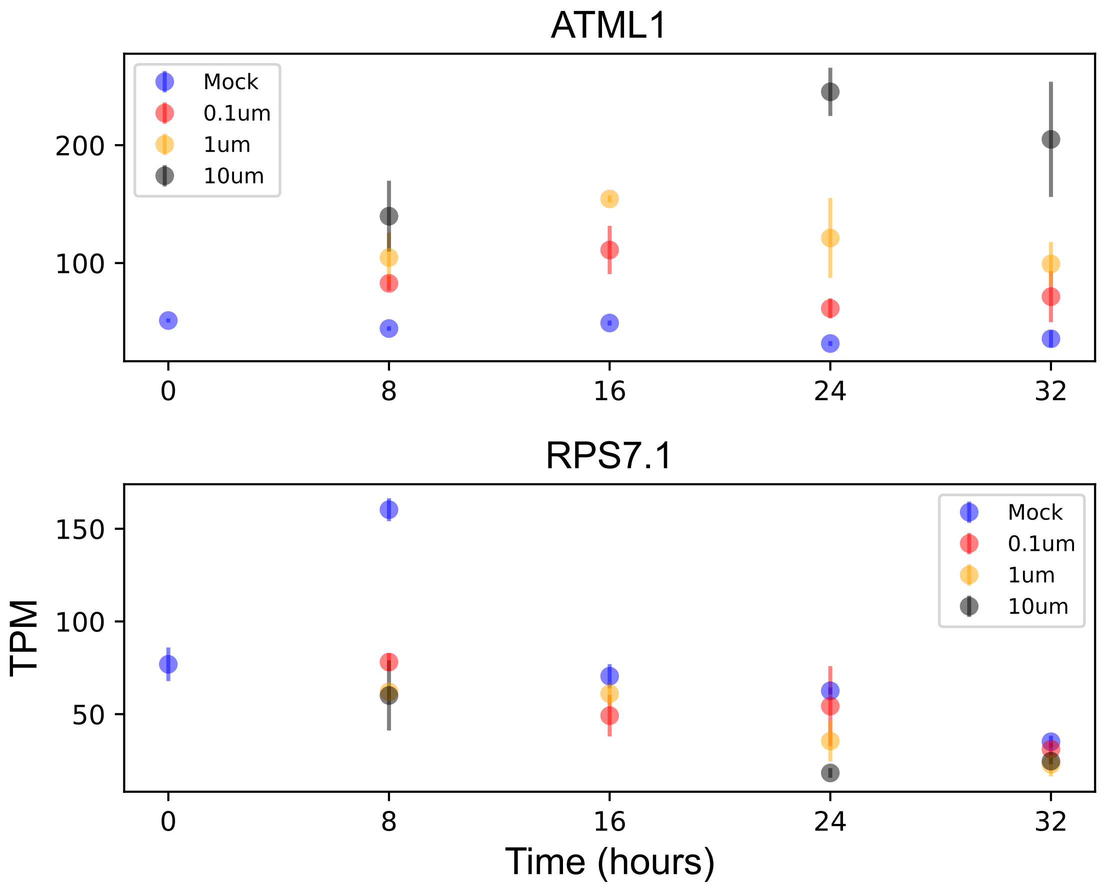

### RPS7.2.pdf

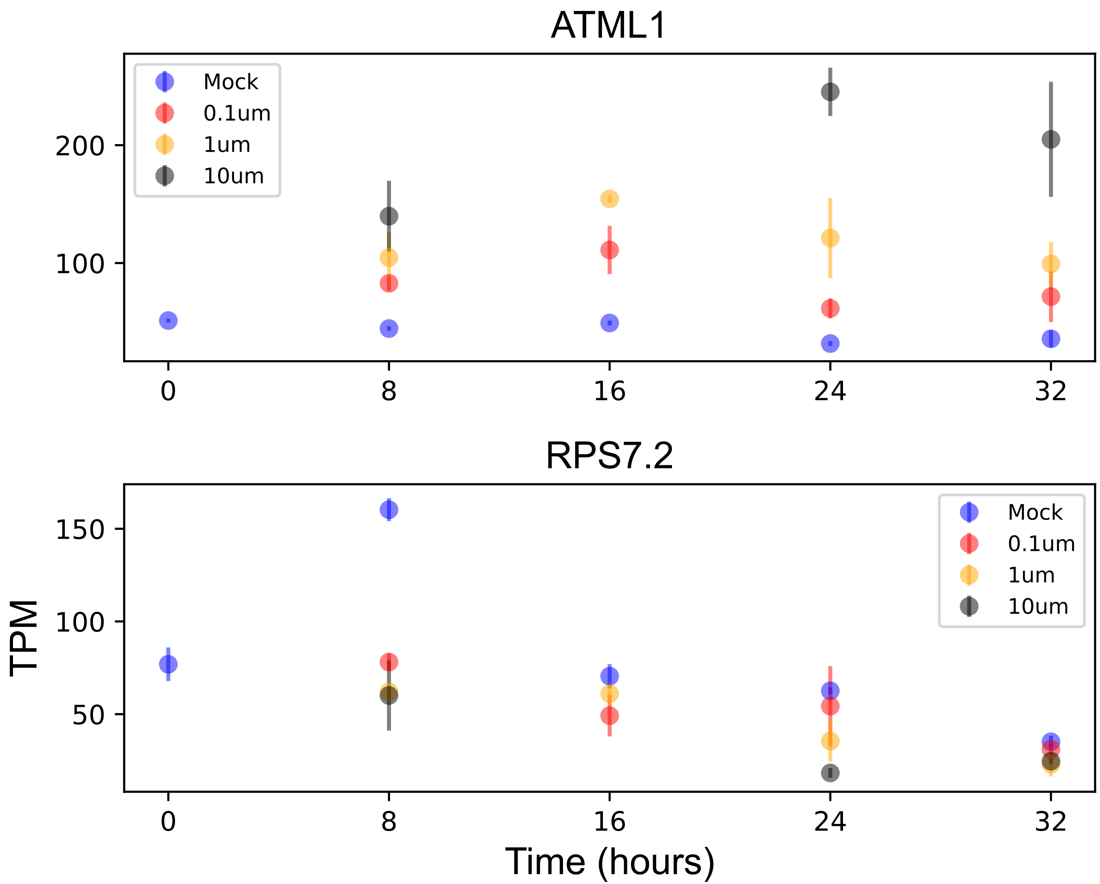

### RPS14.pdf

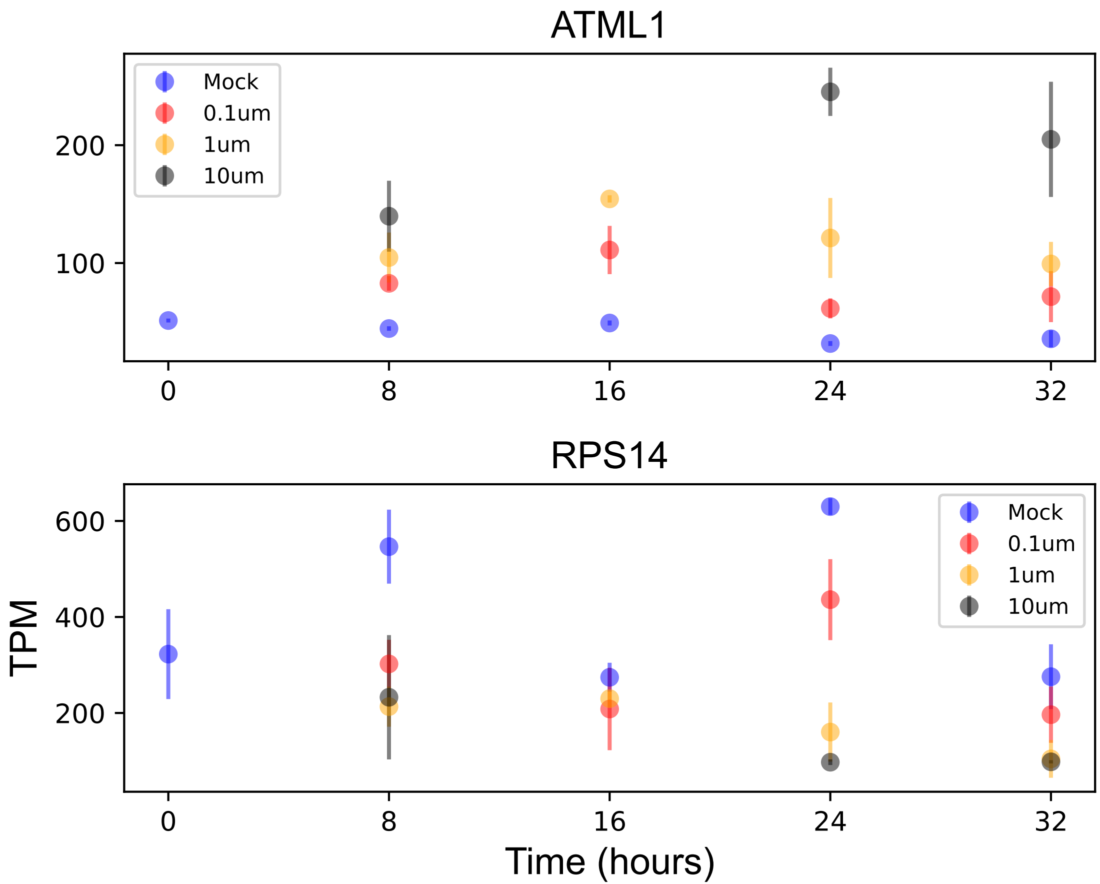

### RPT2a.pdf

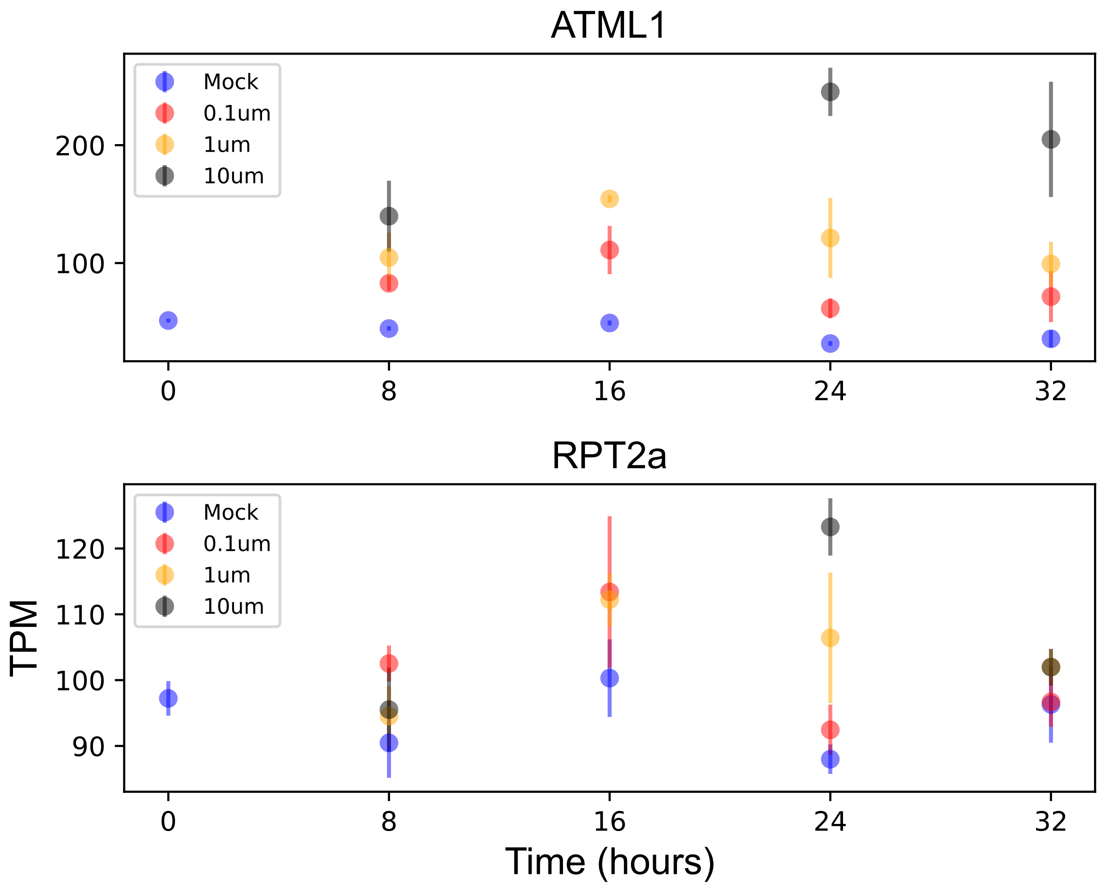

### RRT1.pdf

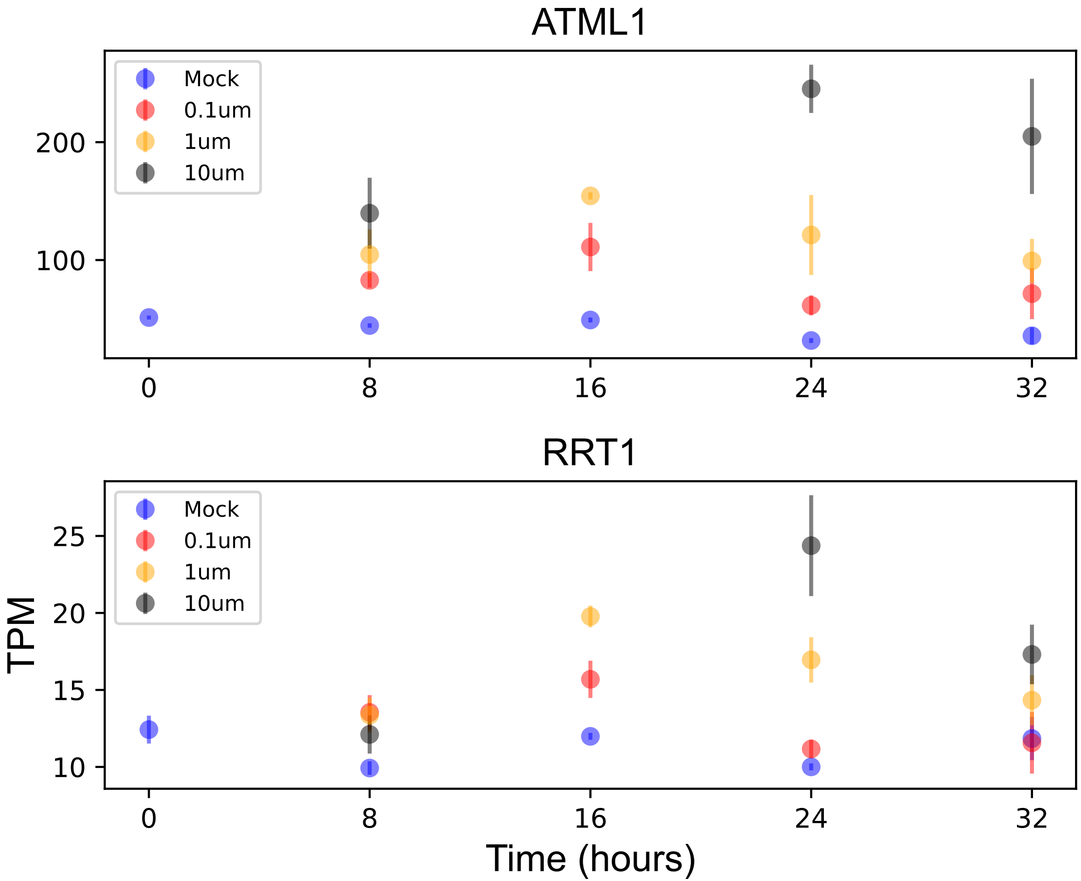

### SCPL9.pdf

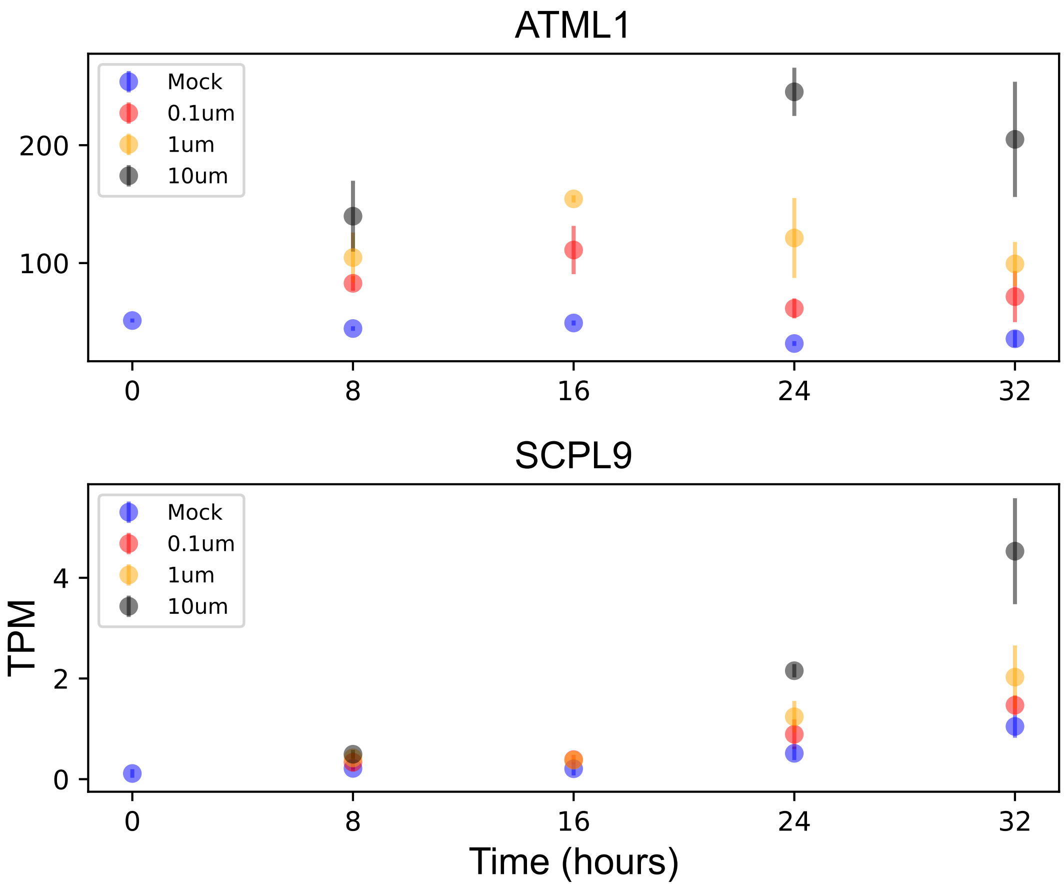

### SEP1.pdf

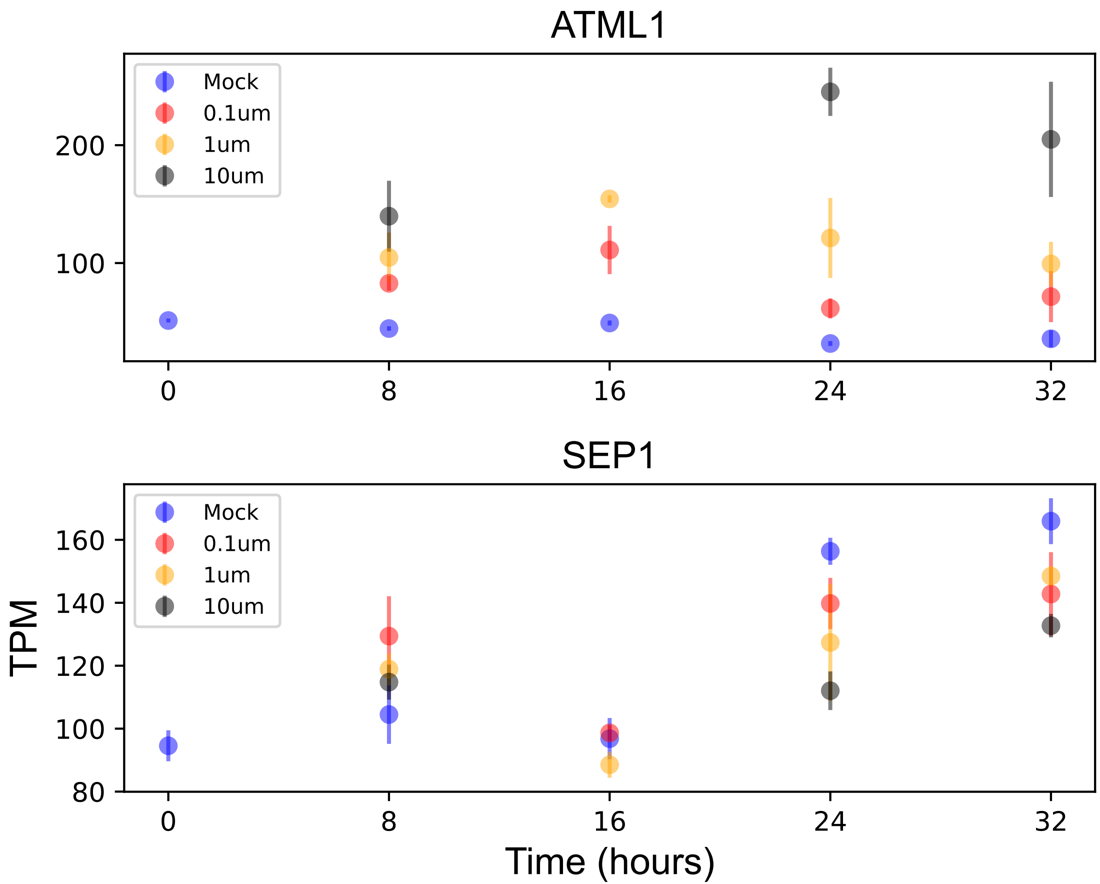

### SKS6.pdf

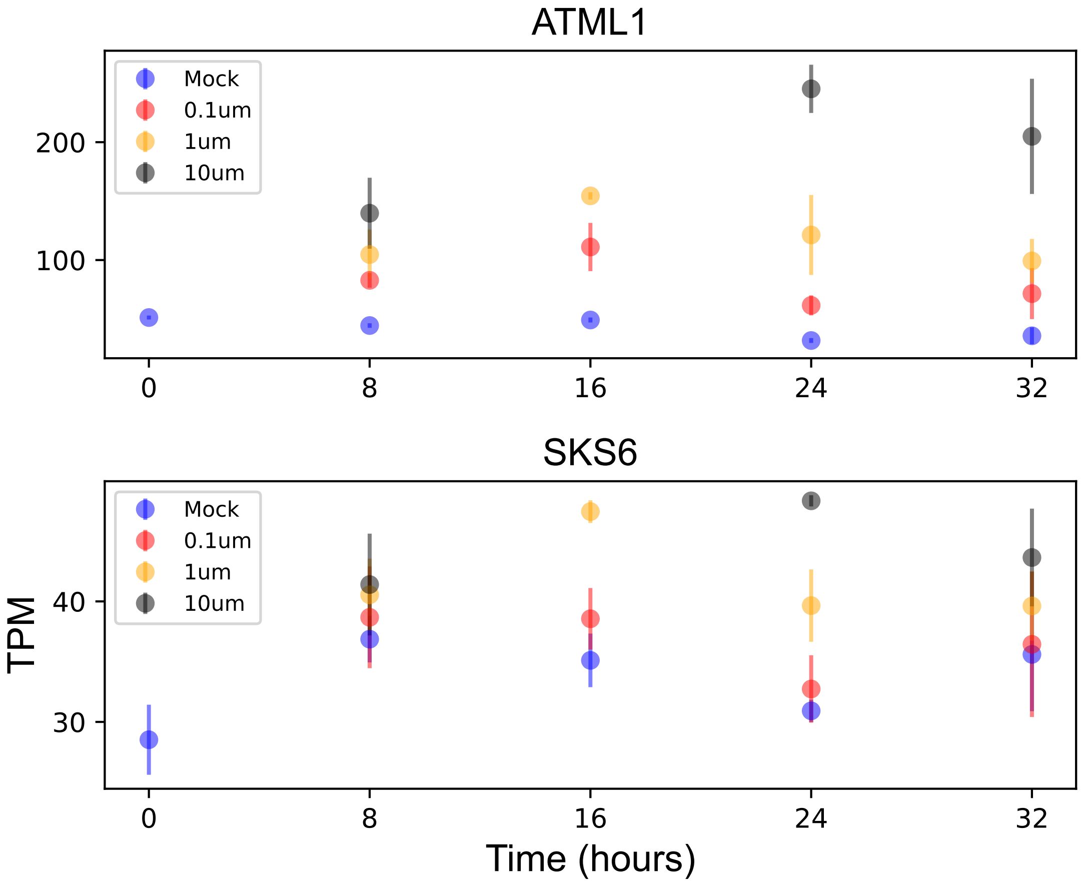

### SRC2.pdf

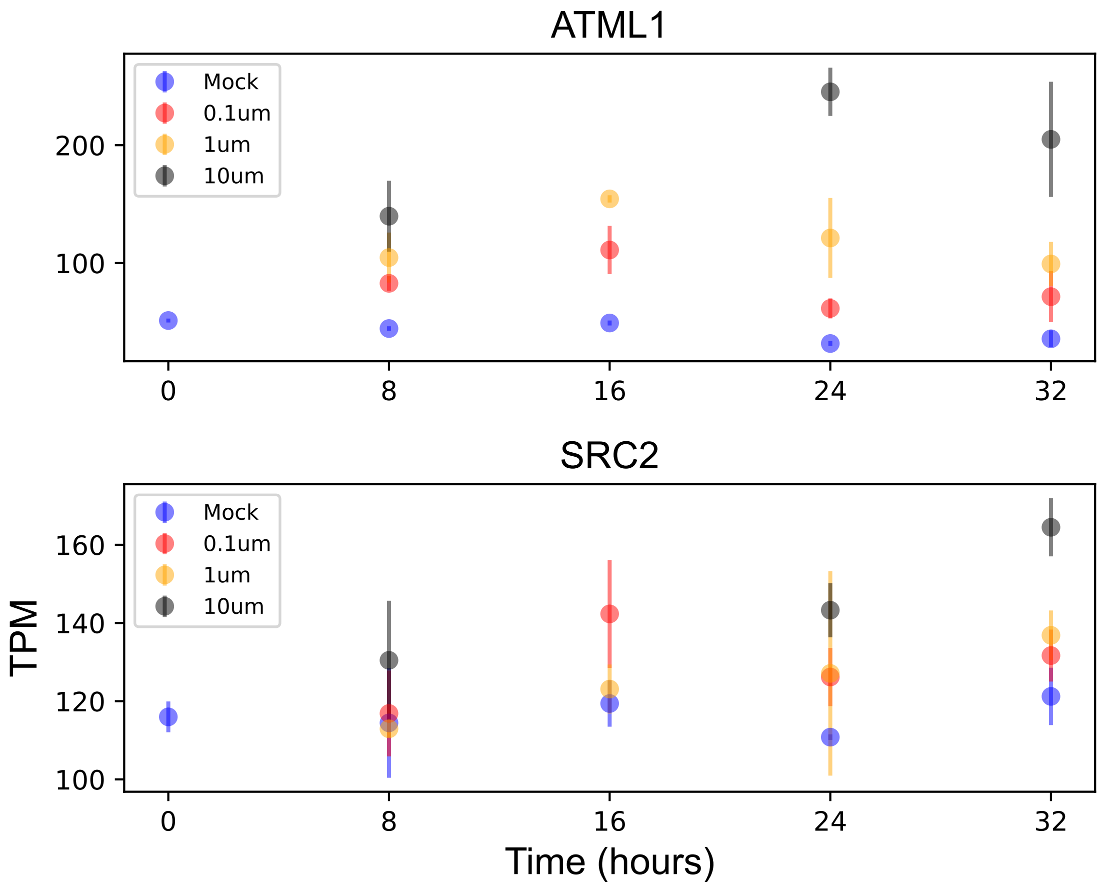

### ST2B.pdf

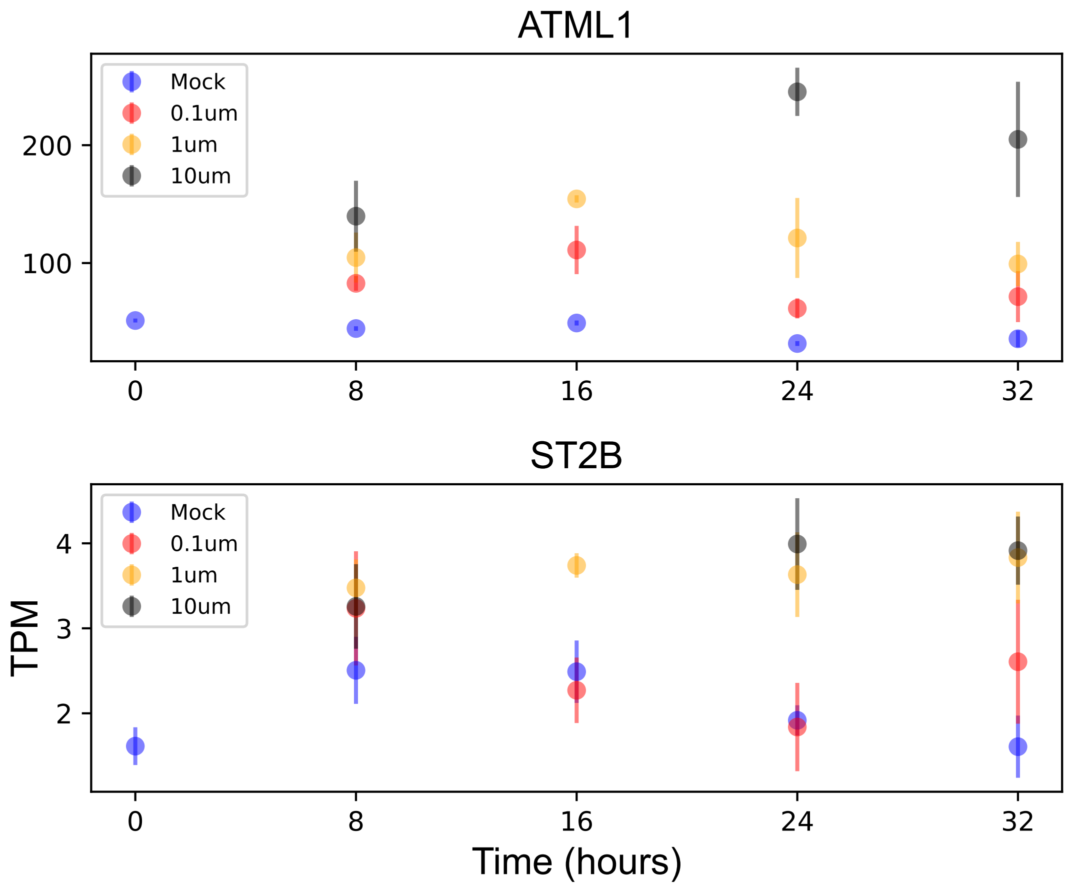

### SVB.pdf

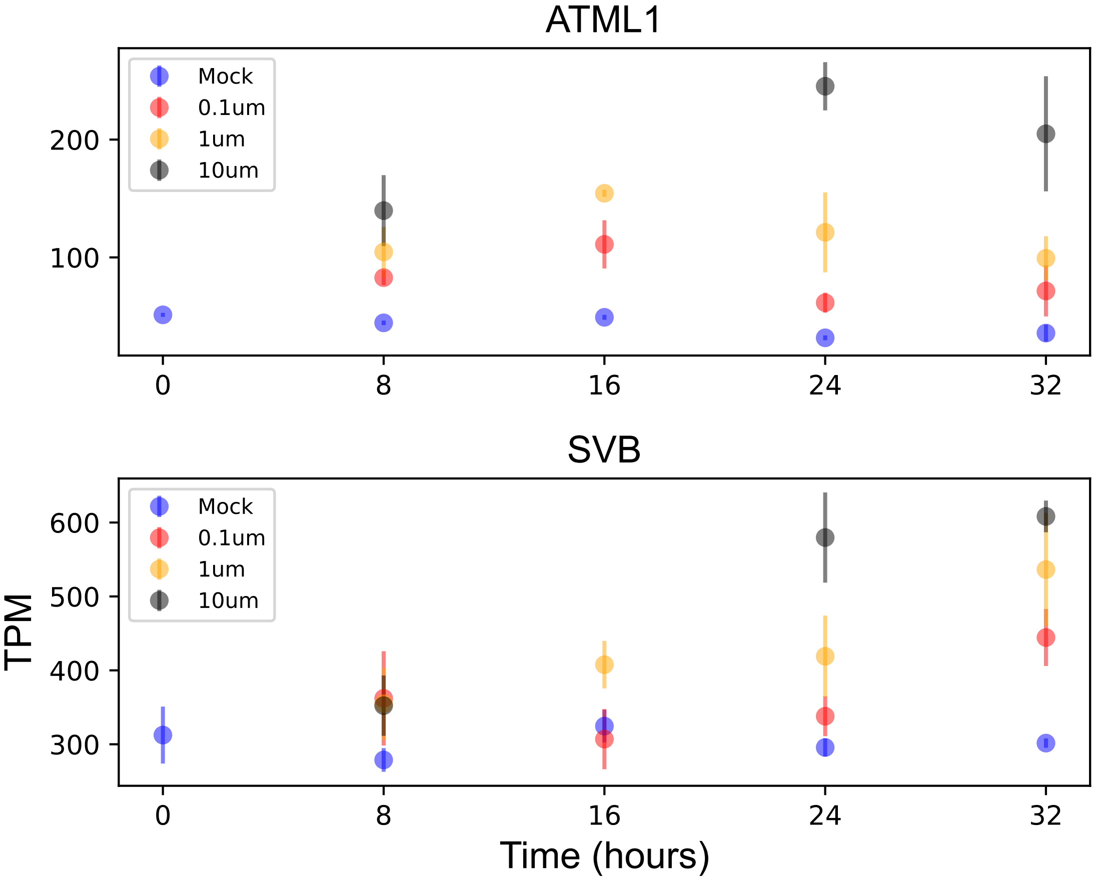

### SWIB2.pdf

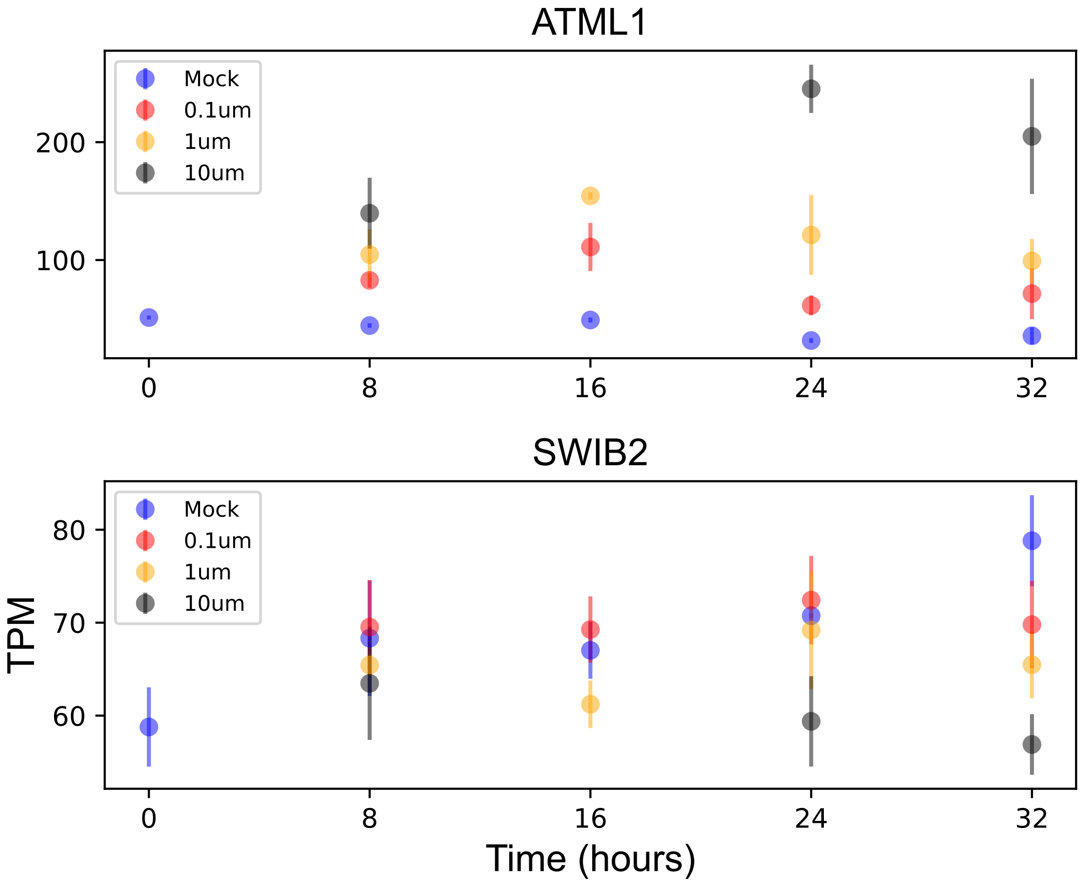

### TMM.pdf

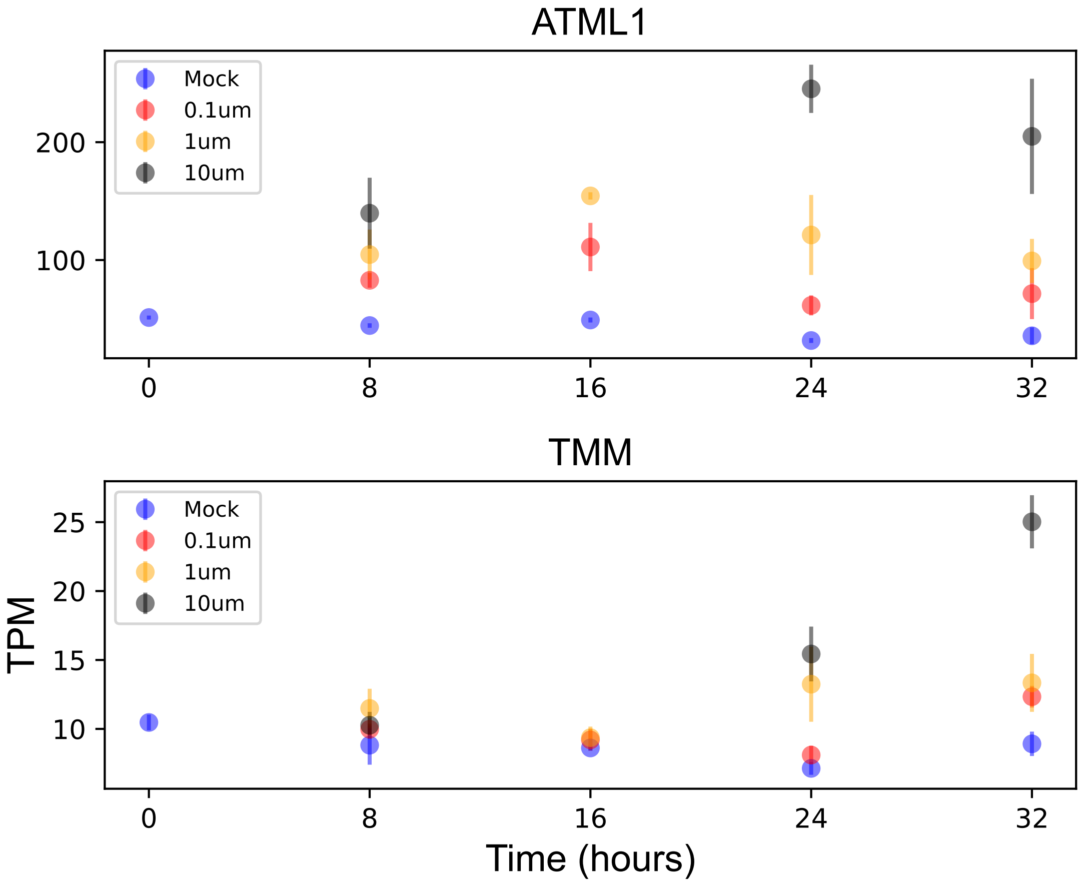

### UGD1.pdf

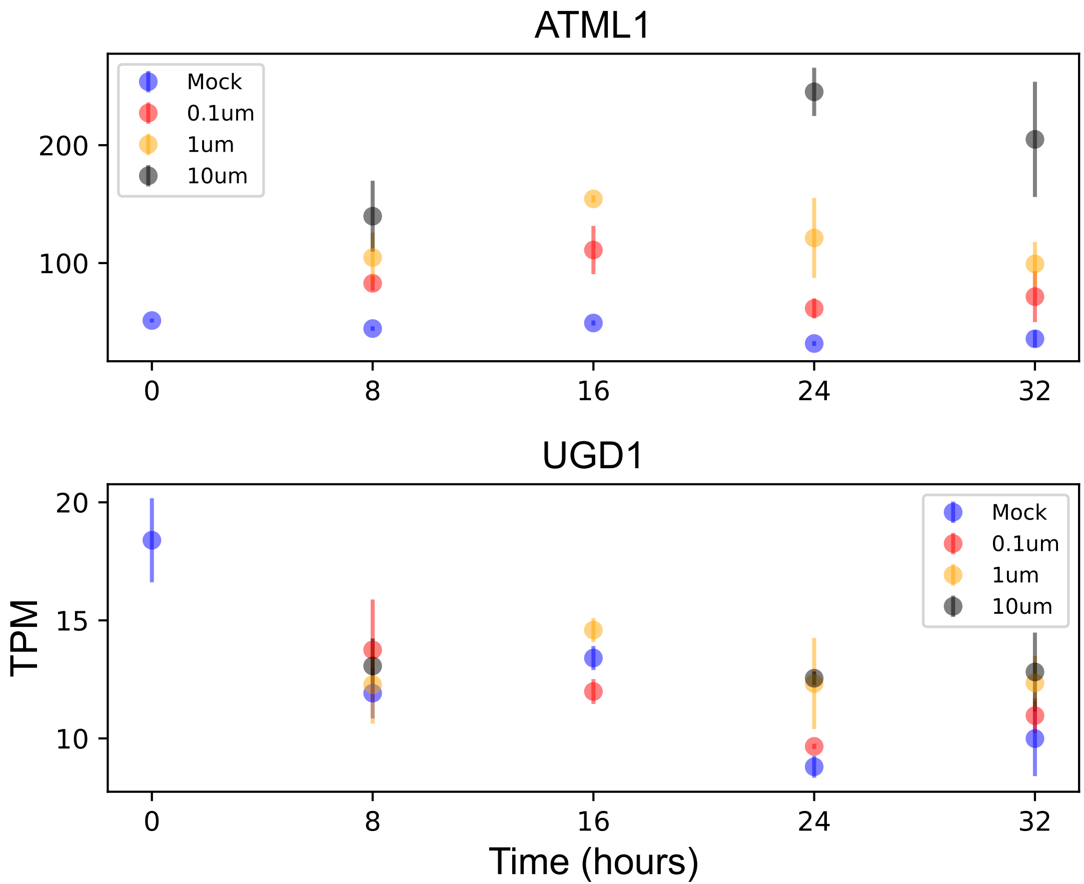

### UGT72E1.pdf

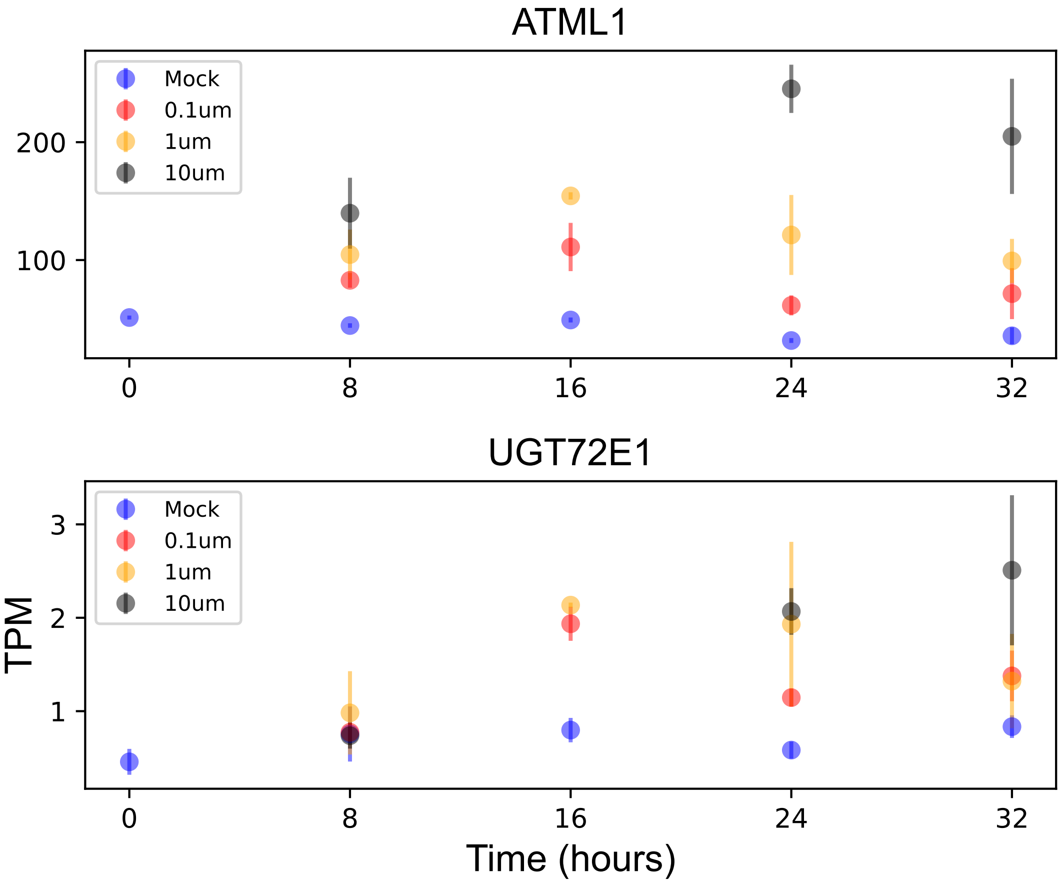

### UGT73B3.pdf

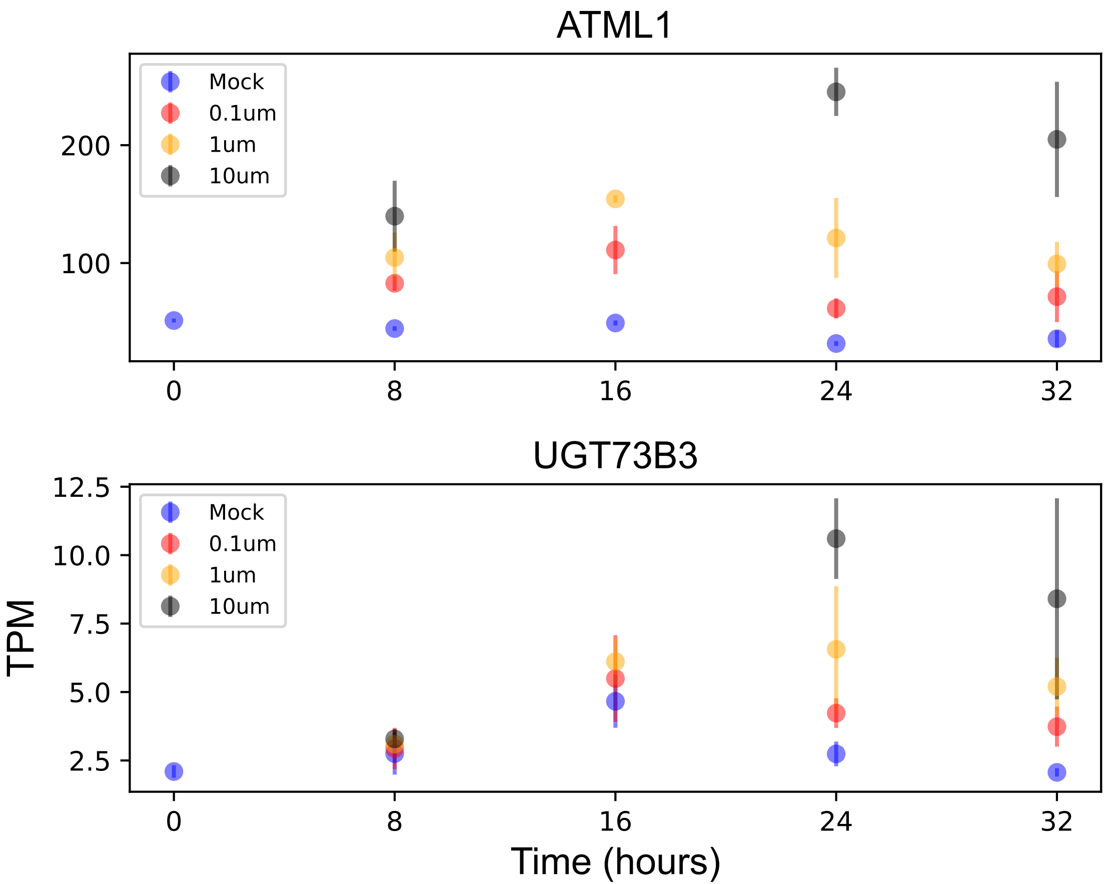

### UGT73B5.pdf

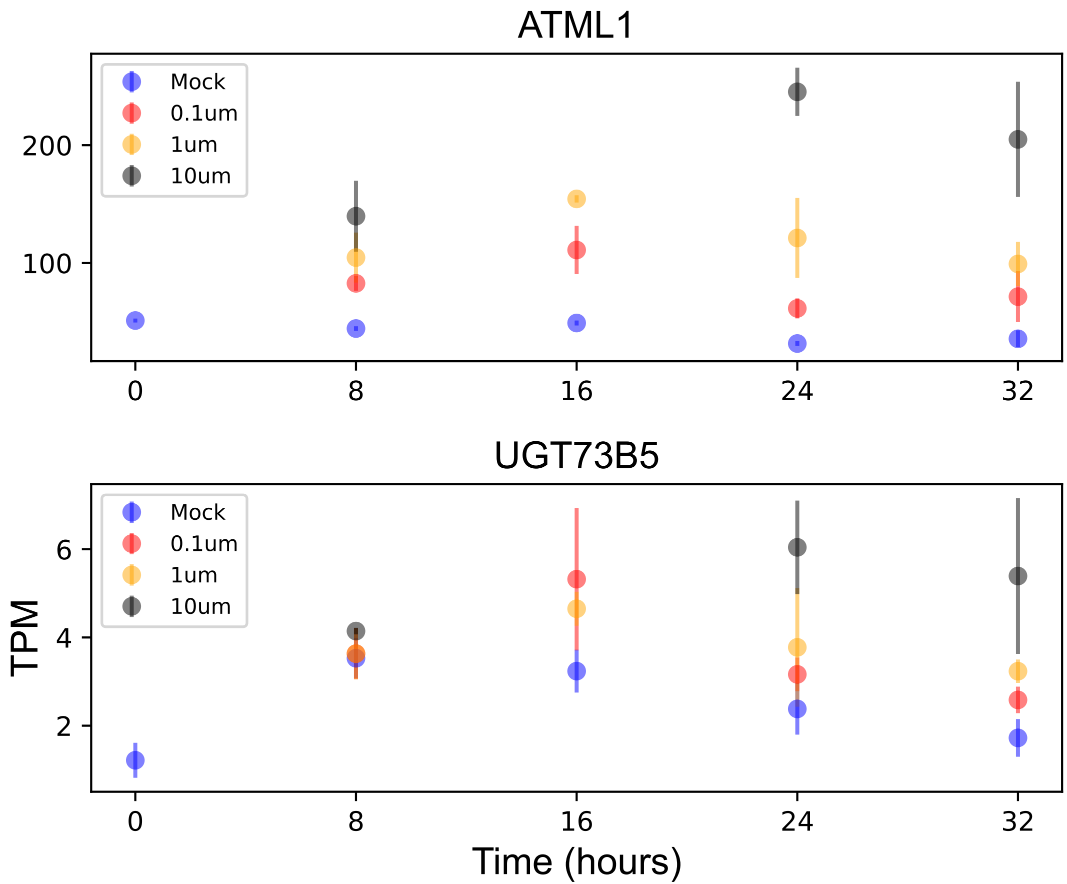

### VPT1.pdf

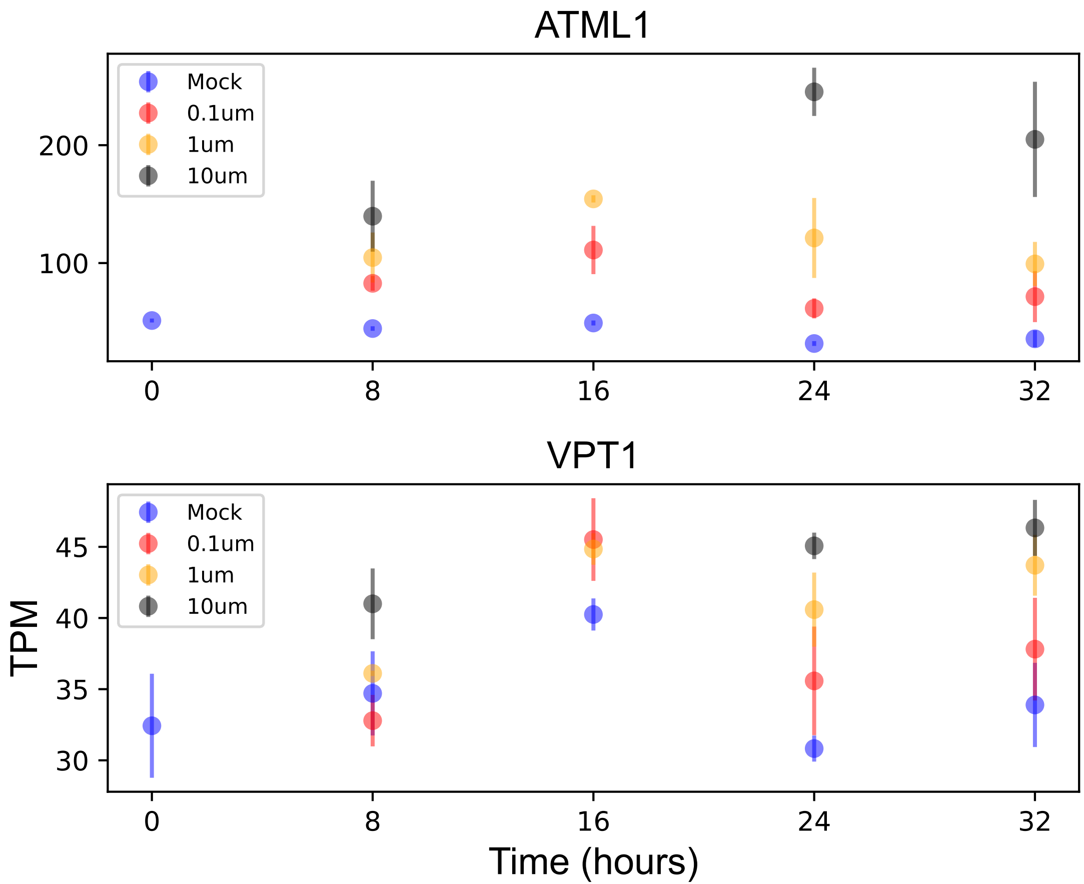

### WAVE2.pdf

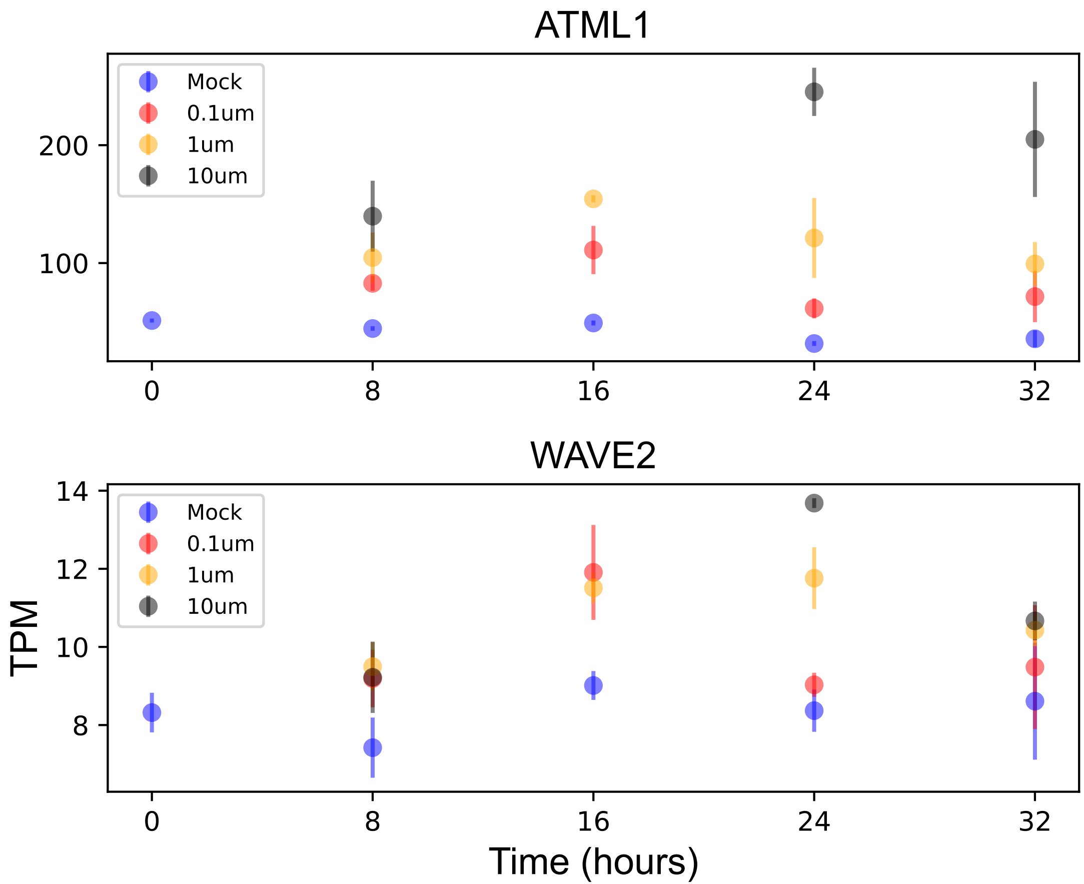

### XTH31.pdf

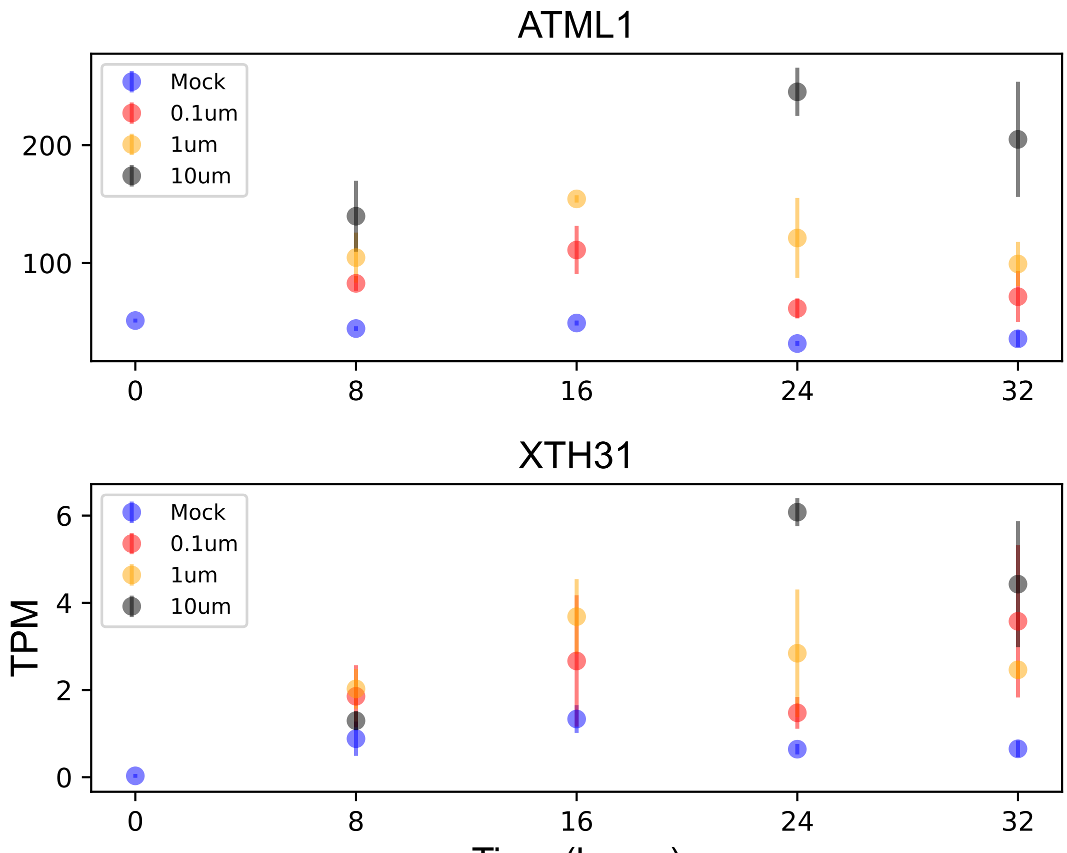

### YCF1.2.pdf

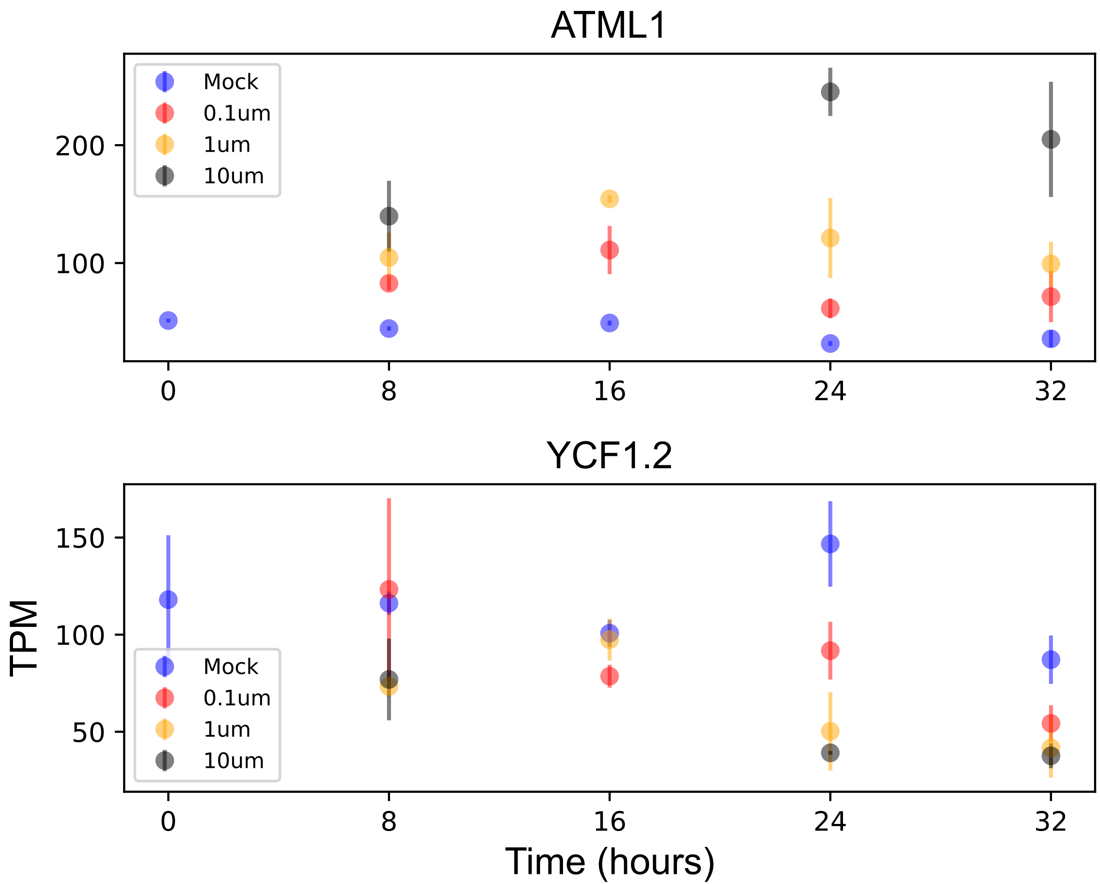

### YCF5.pdf

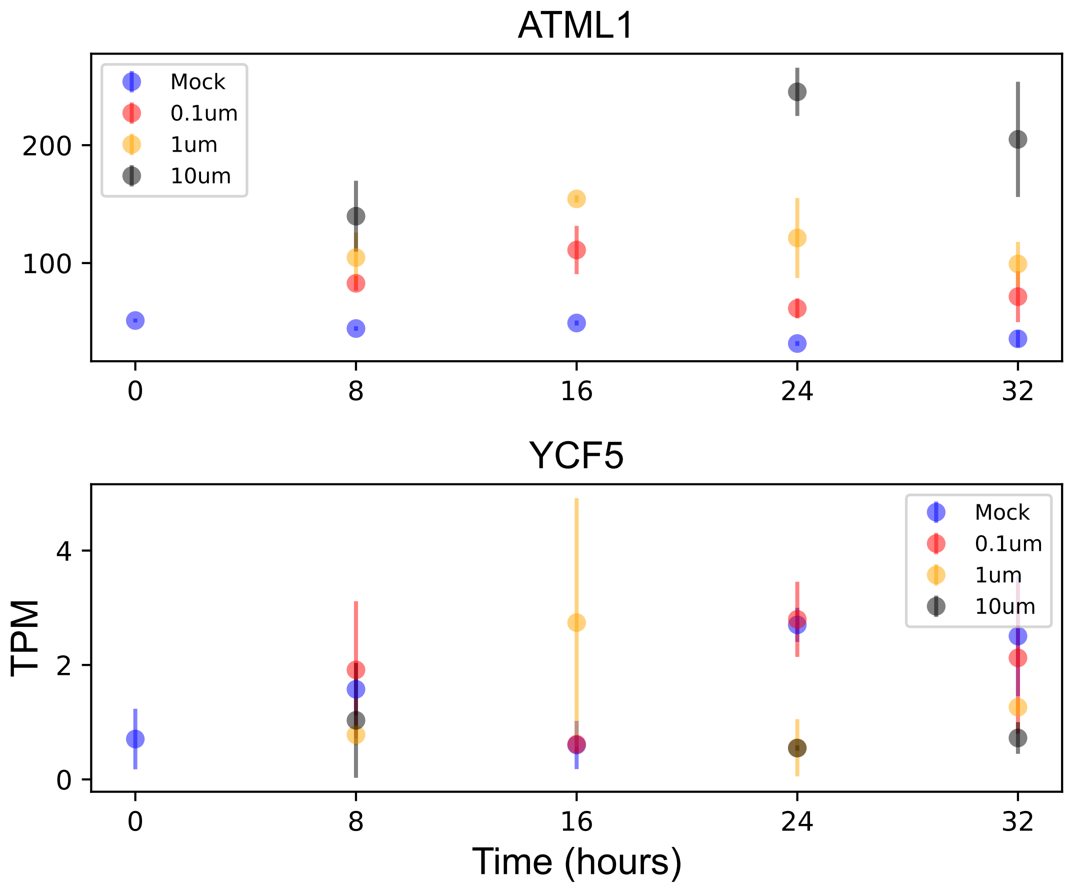
