## Supplementary figures and images for "The transcription factor ATML1 maintains giant cell identity by inducing synthesis of its own long-chain fatty acid-containing ligands"

### ftsh7.pdf

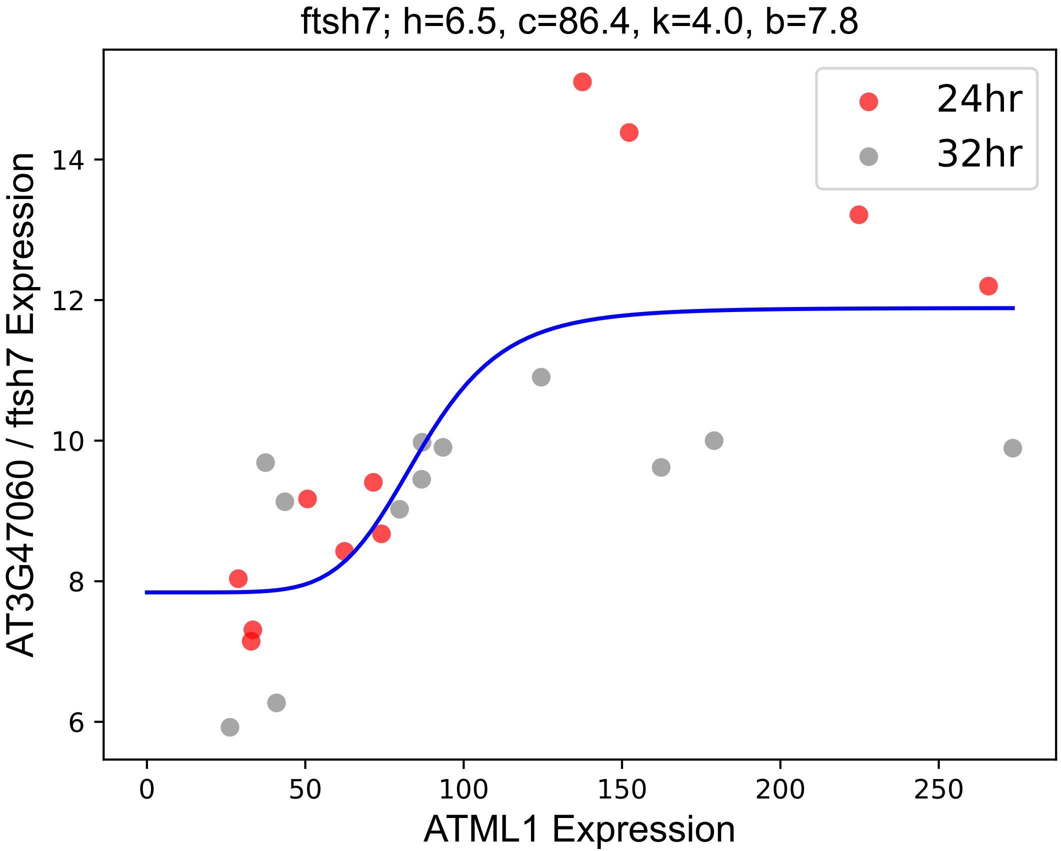

### PAS2.pdf

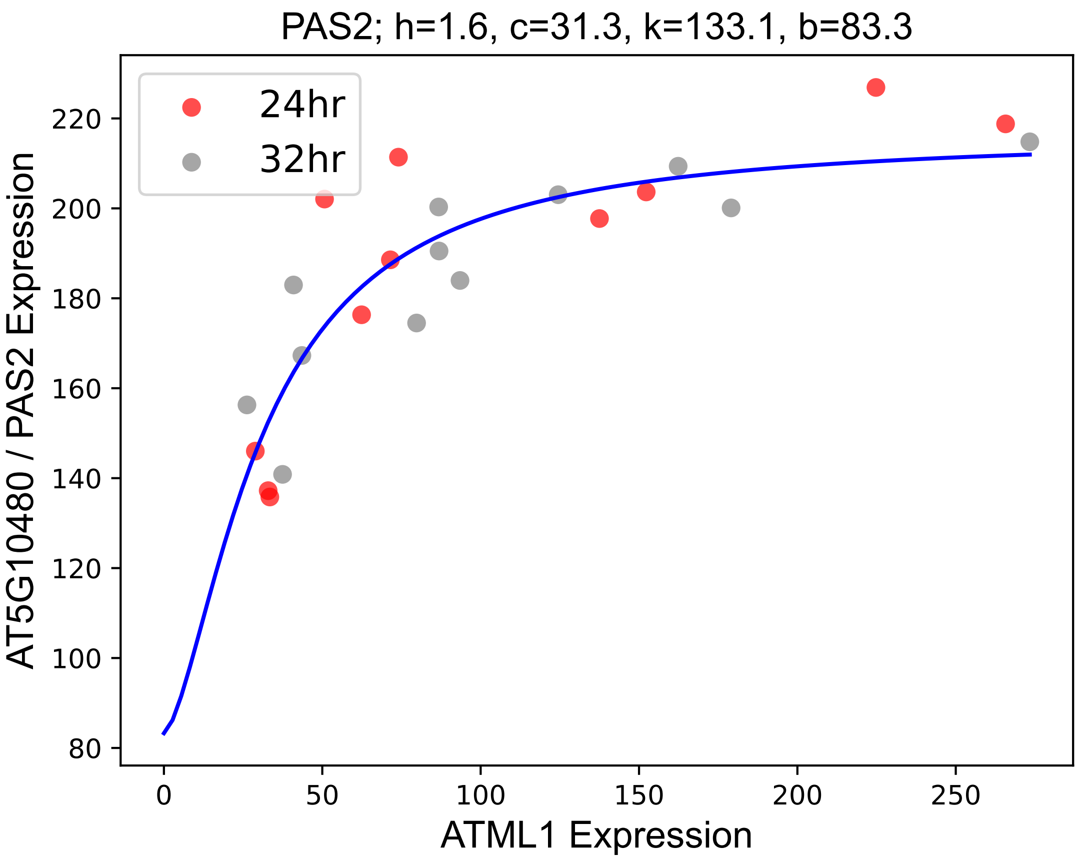

### PBP1.pdf

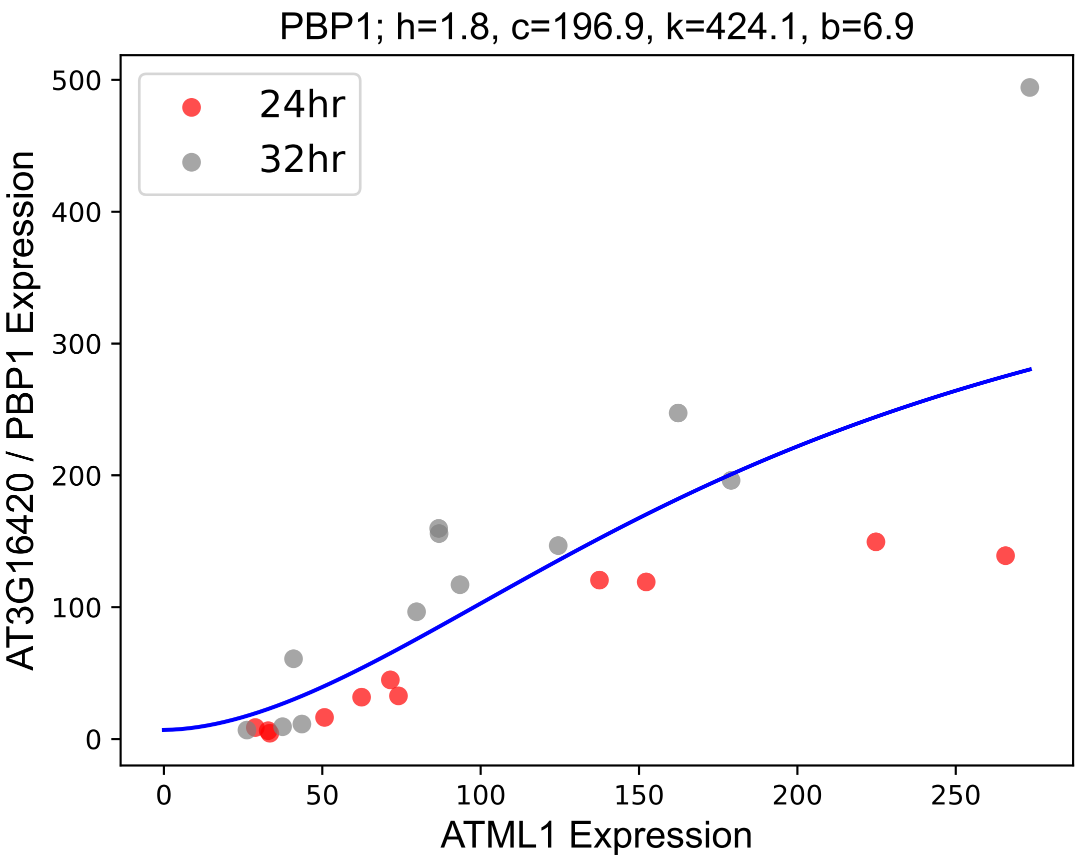

### PLP9.pdf

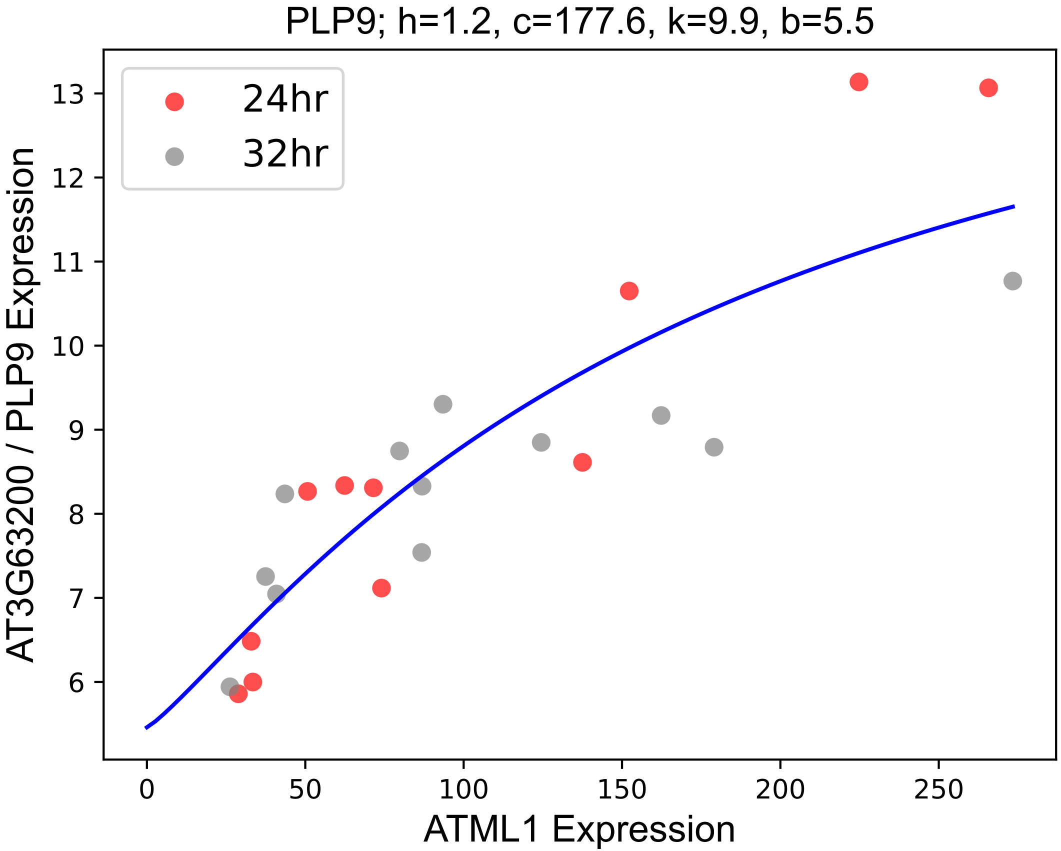

### PME44.pdf

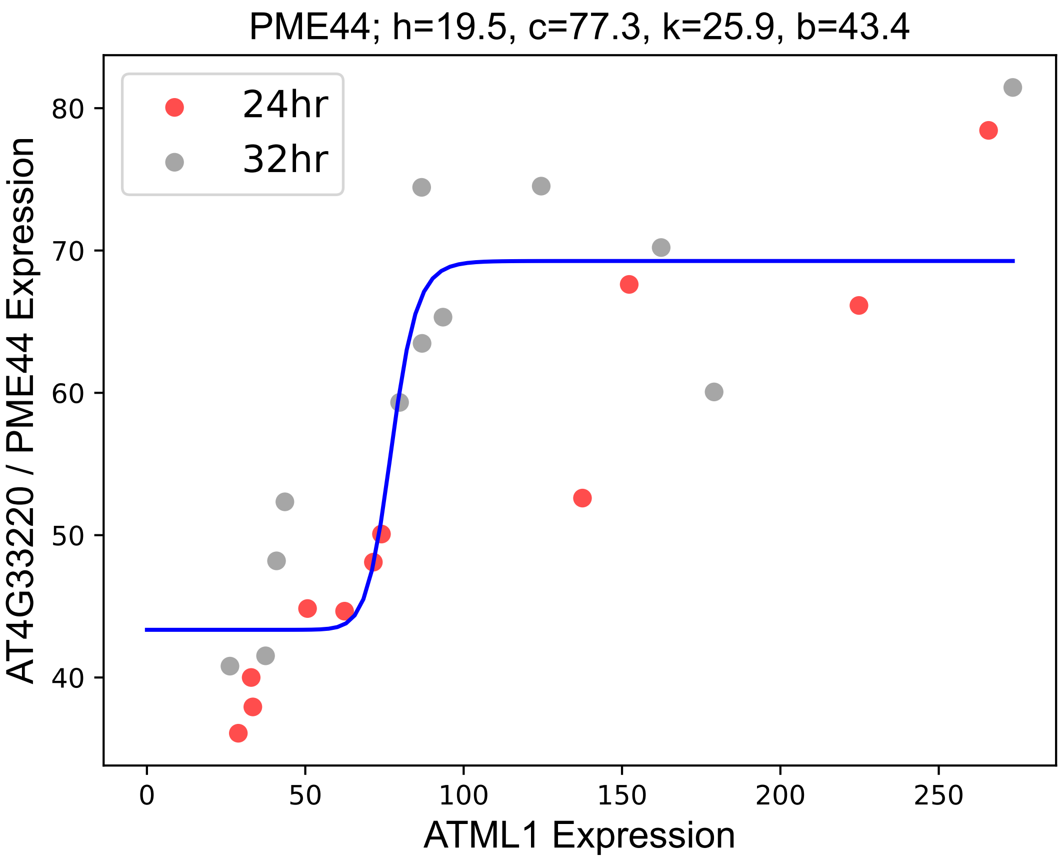

### PnsB3.pdf

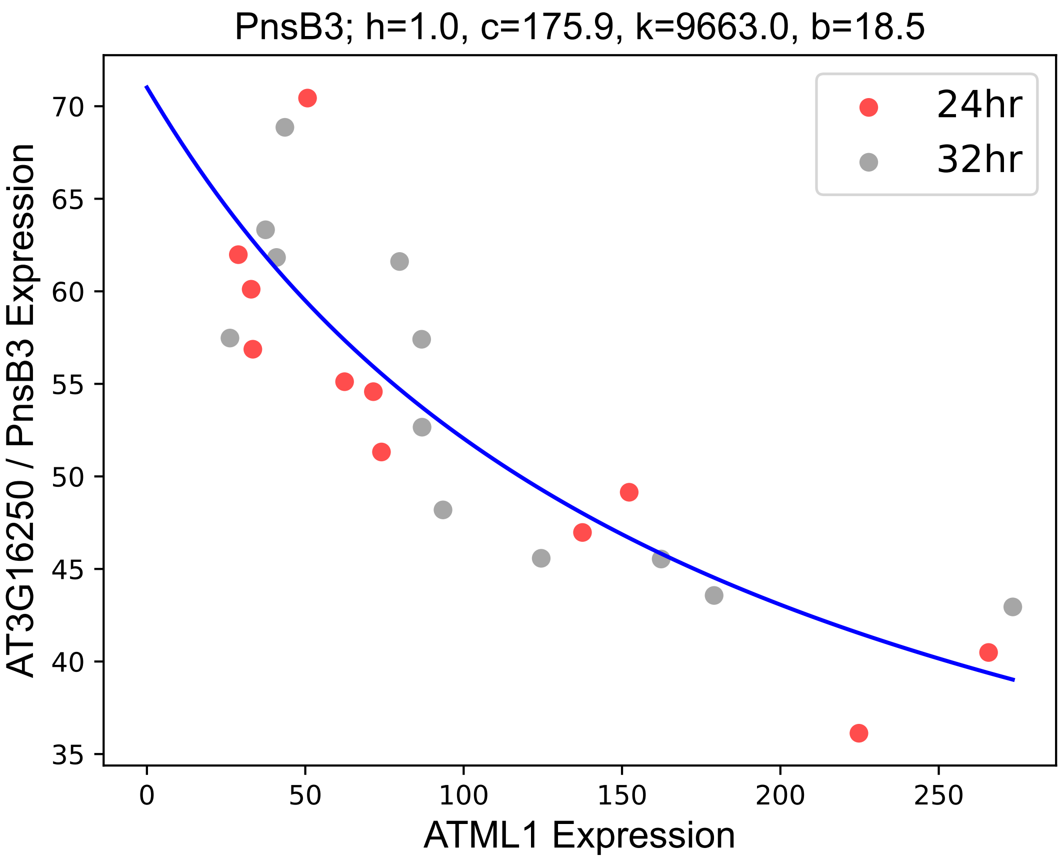

### PnsL4.pdf

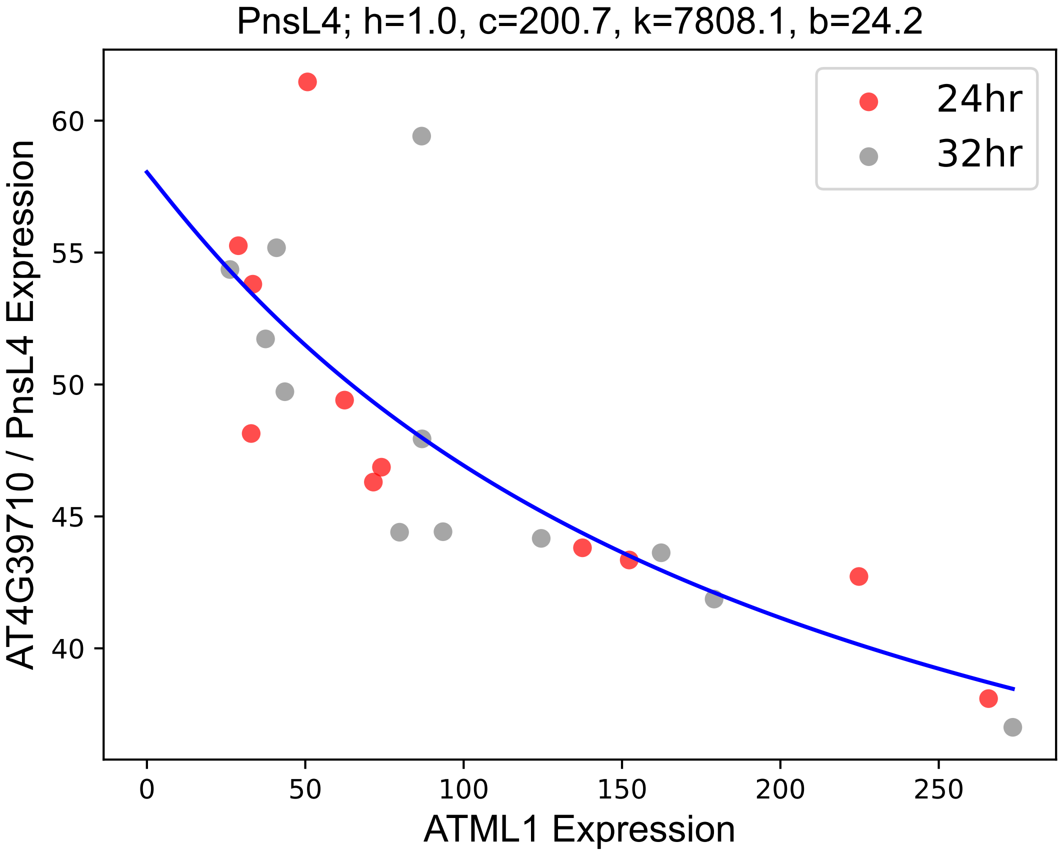

### PRIN2.pdf

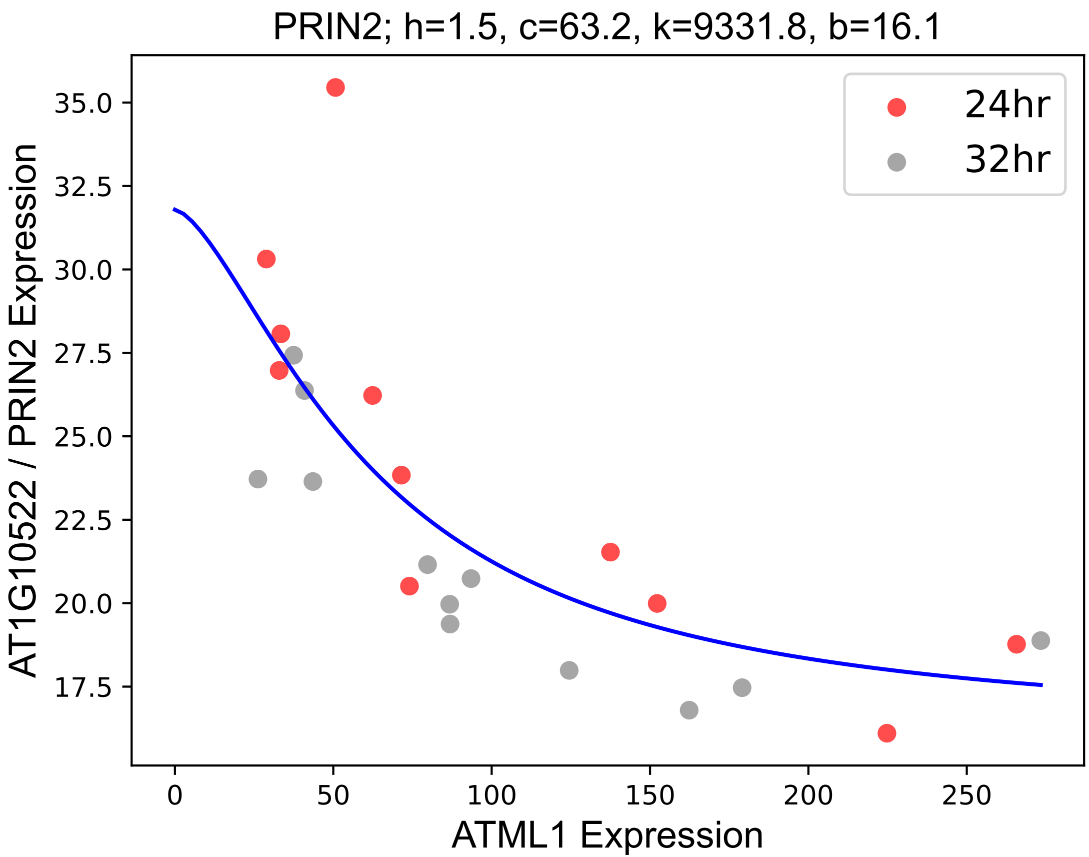

### PRXR1.pdf

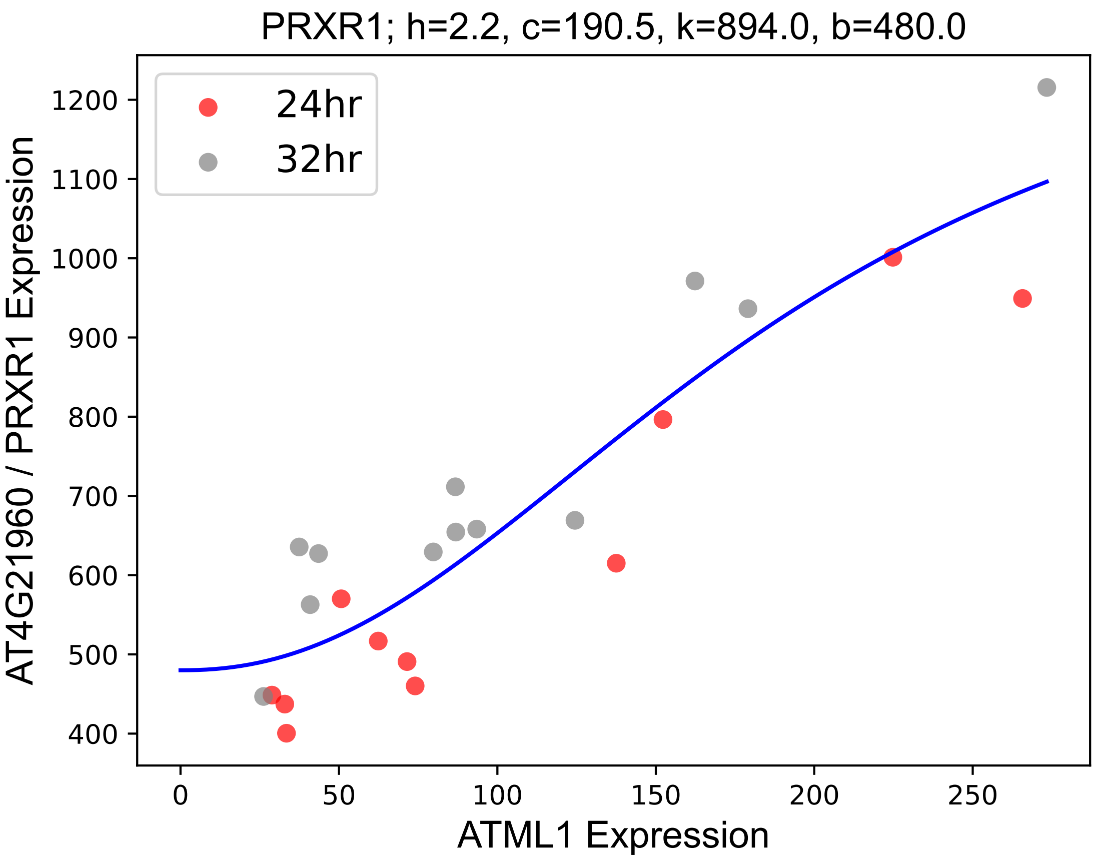

### PTAC7.pdf

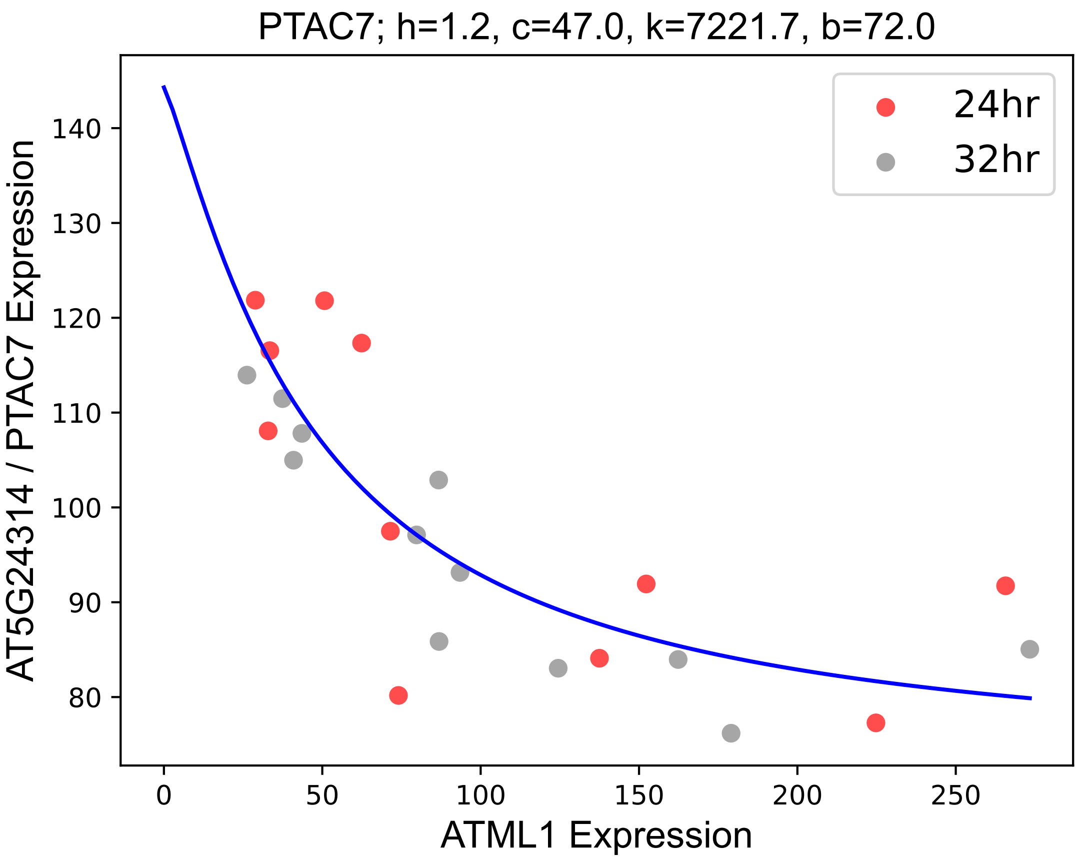

### PYD1.pdf

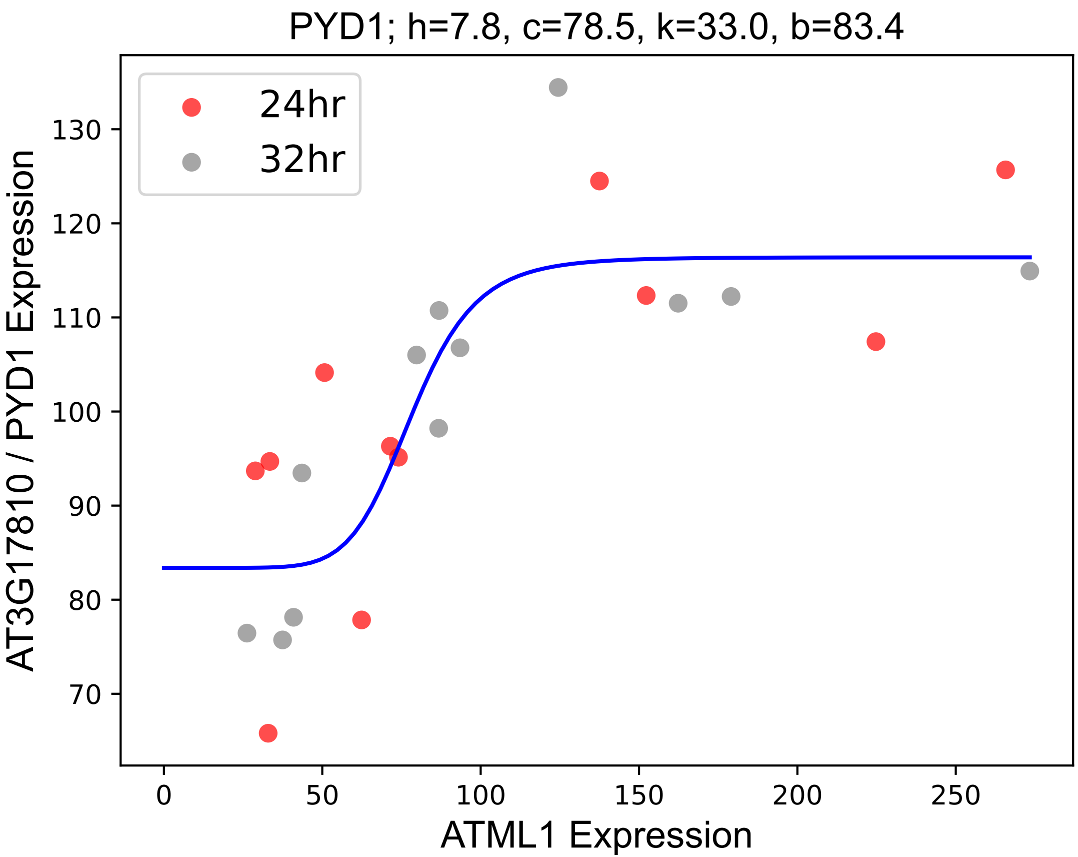

### RBCS1B.pdf

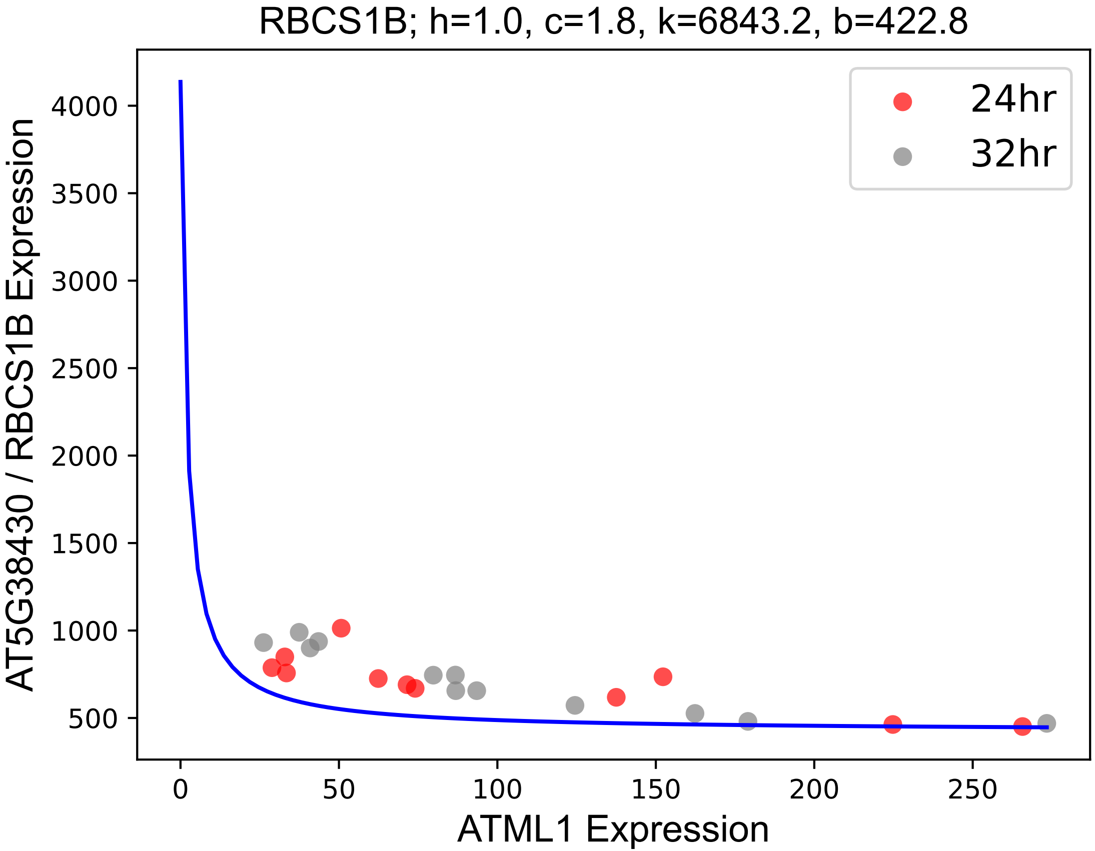

### RGXT2.pdf

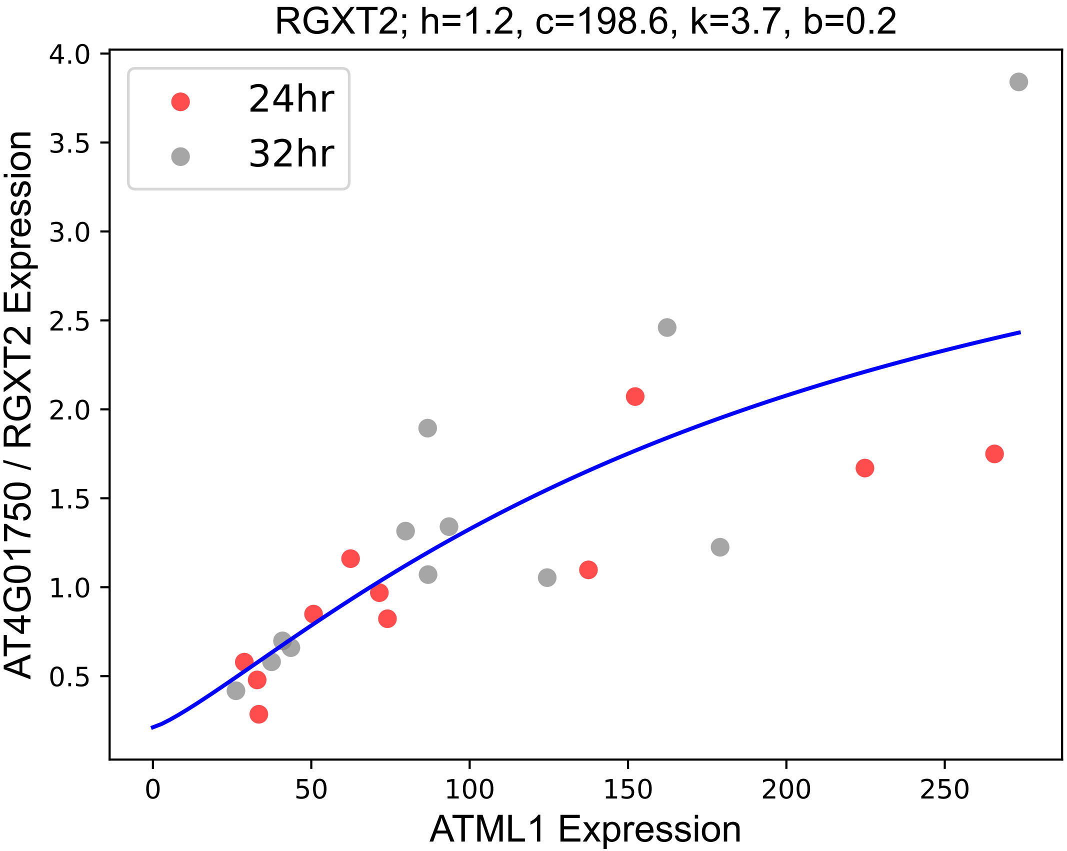

### RPL16.pdf

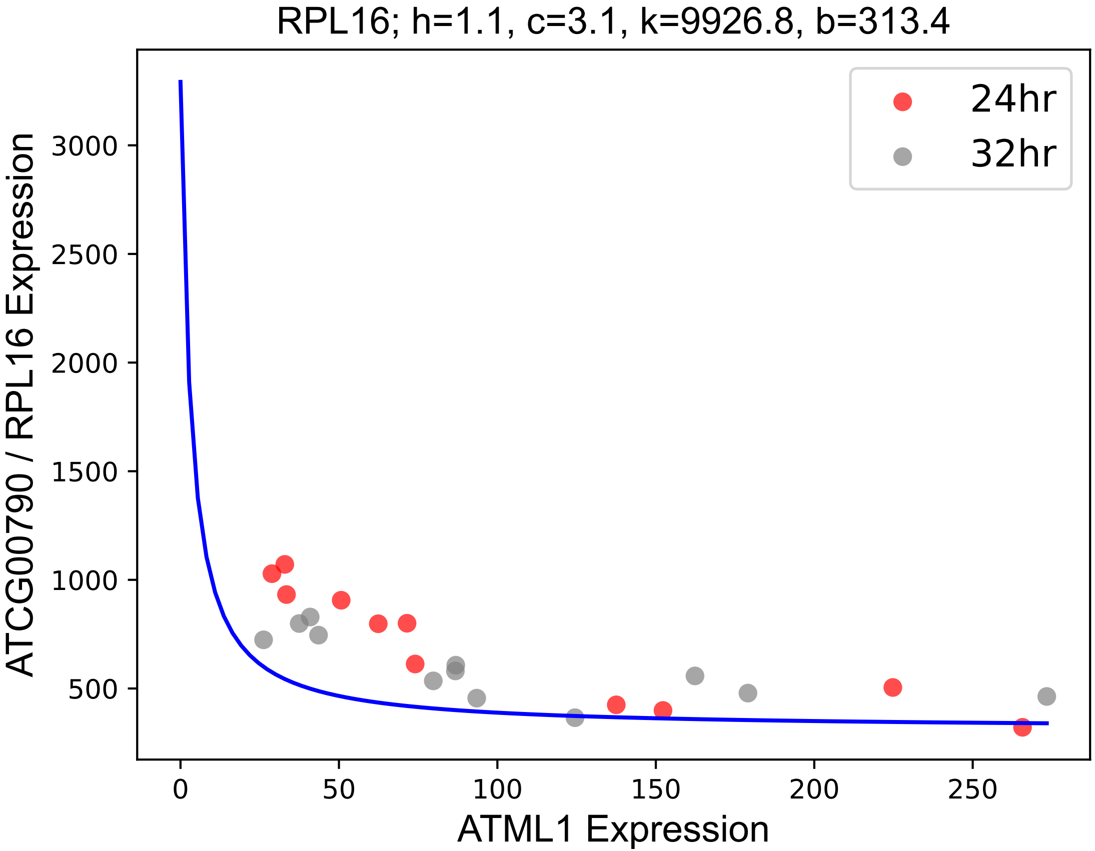

### RPL22.pdf

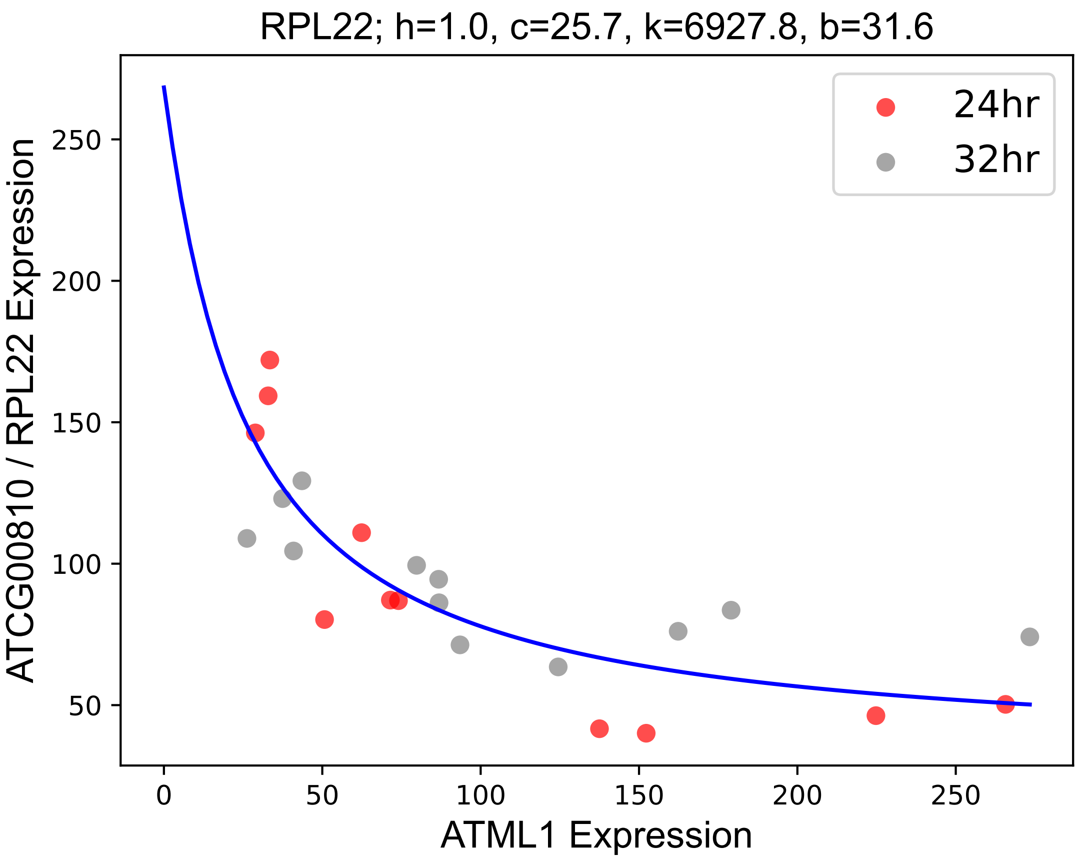

### RPS2.pdf

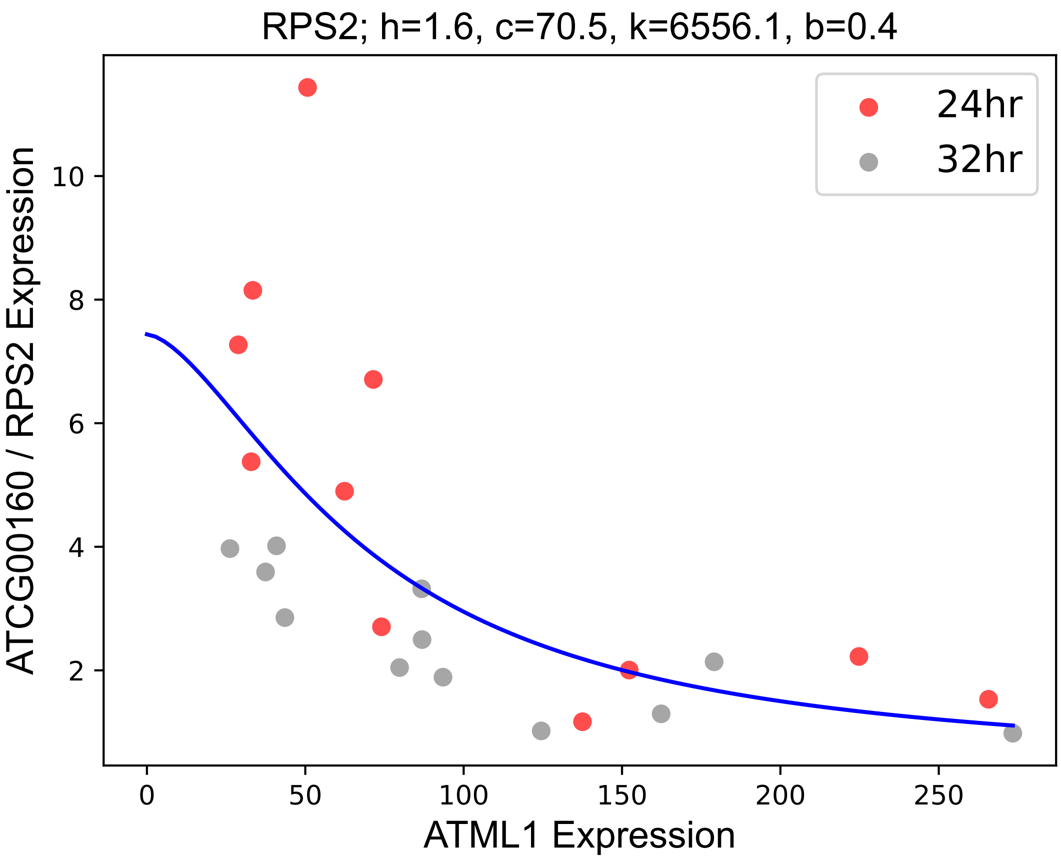

### RPS3.pdf

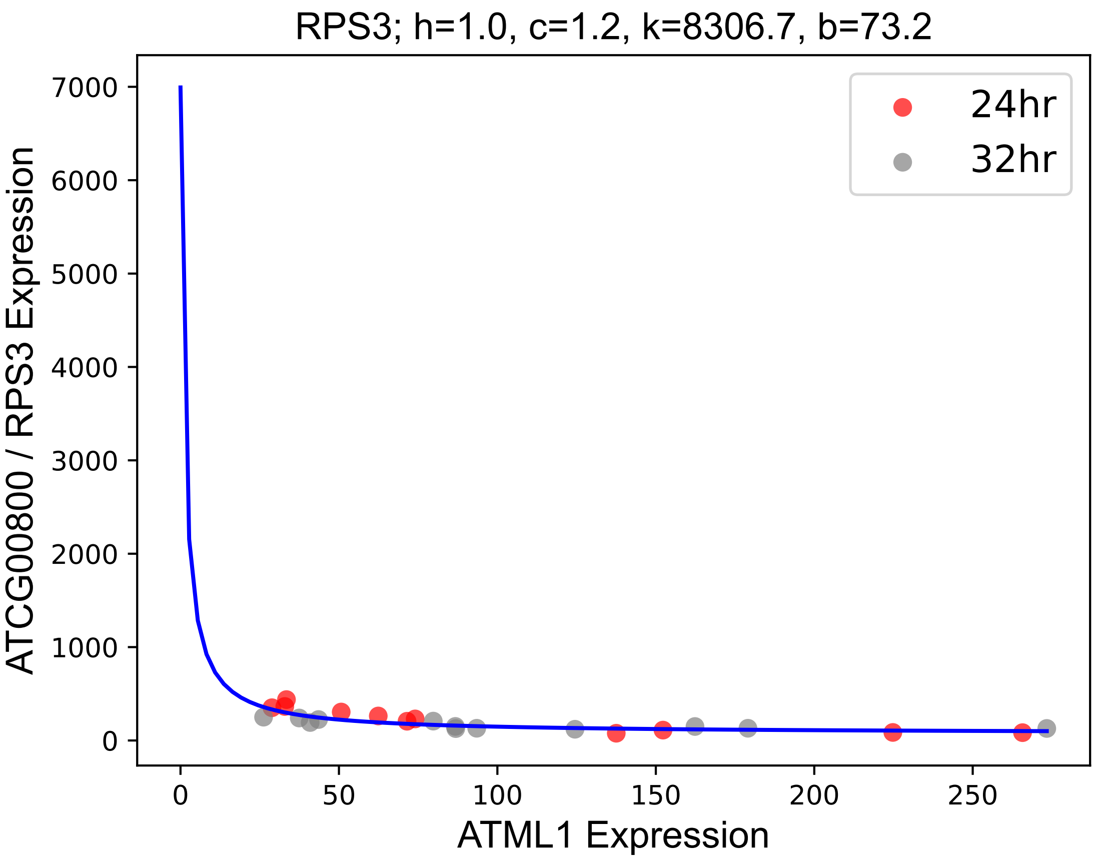

### RPS4.pdf

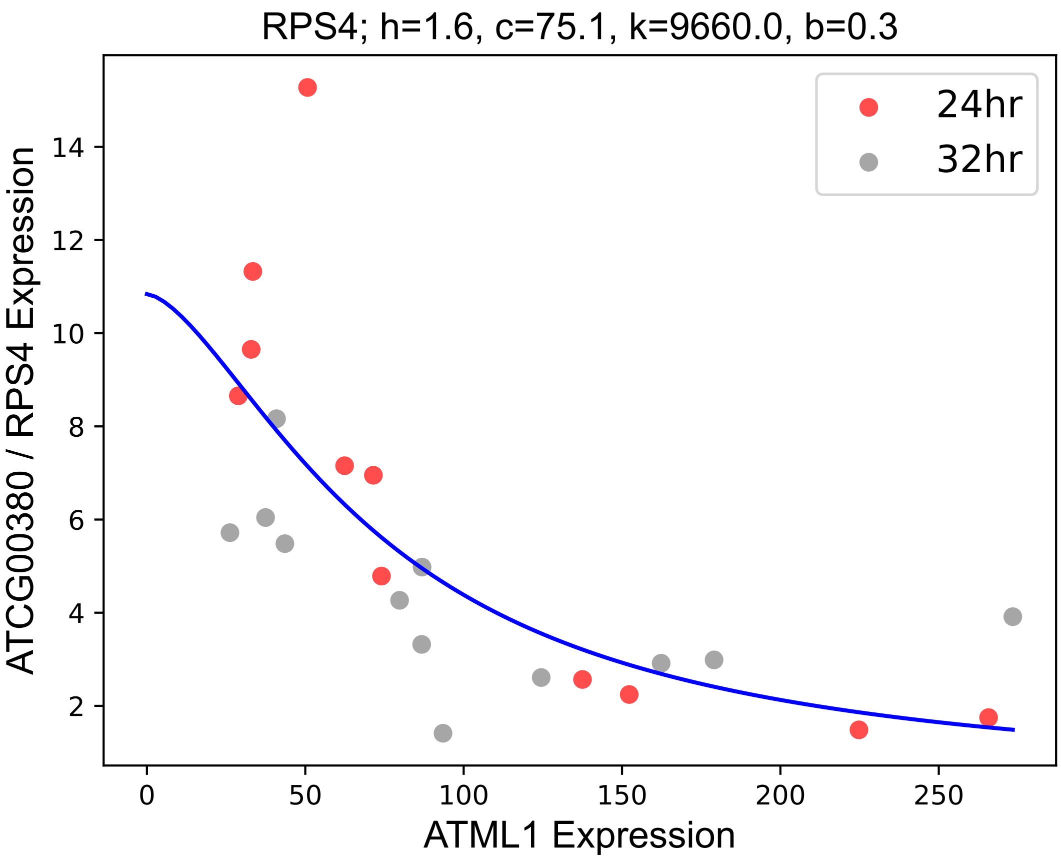

### RPS7.1.pdf

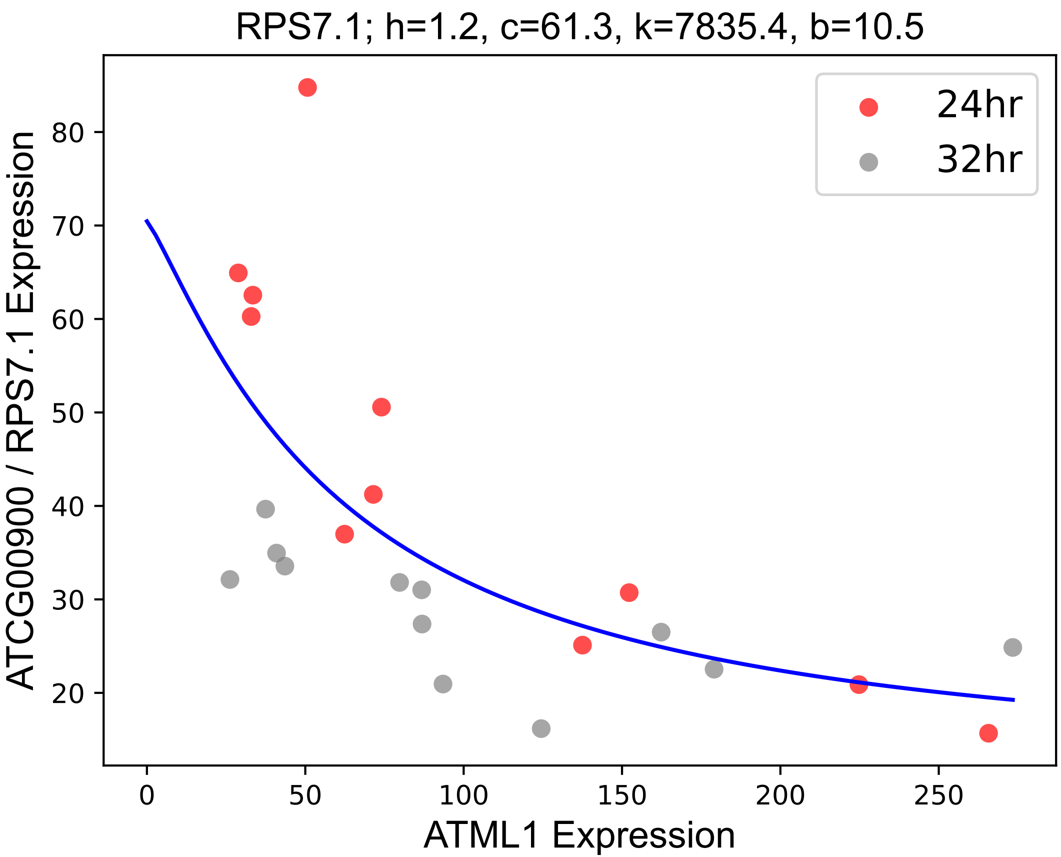

### RPS7.2.pdf

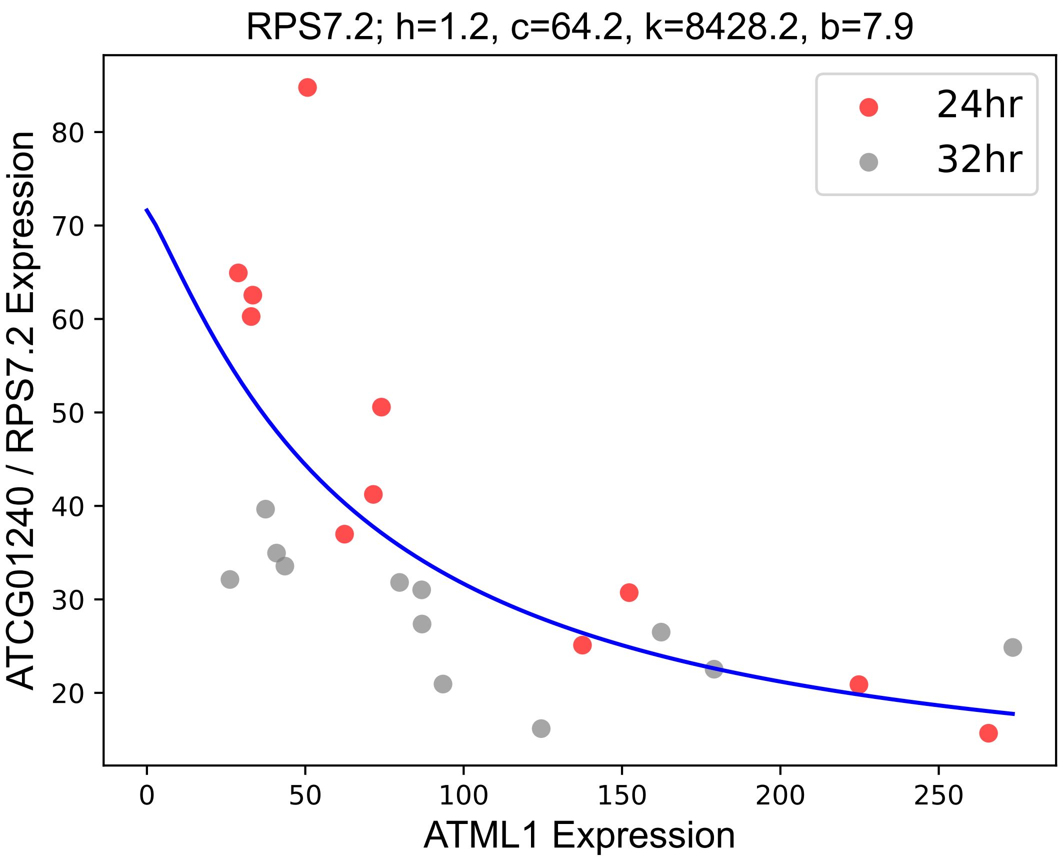

### RPS14.pdf

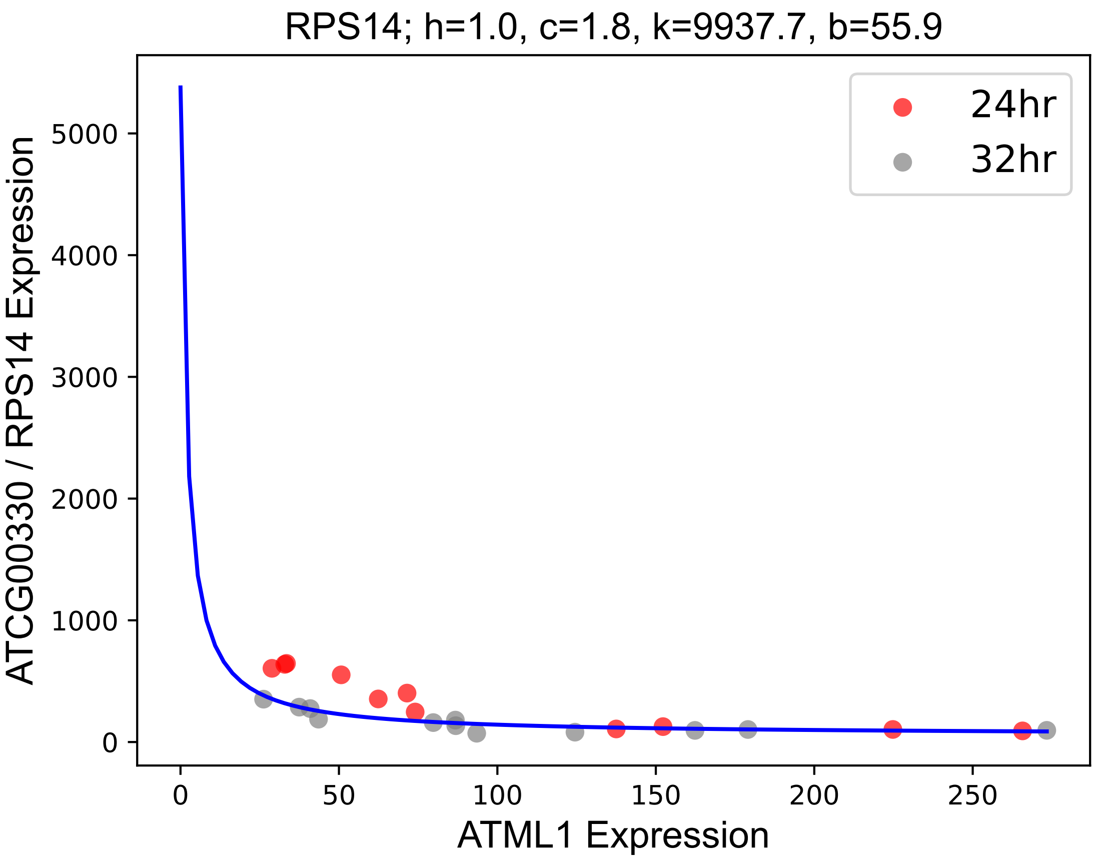

### RPT2a.pdf

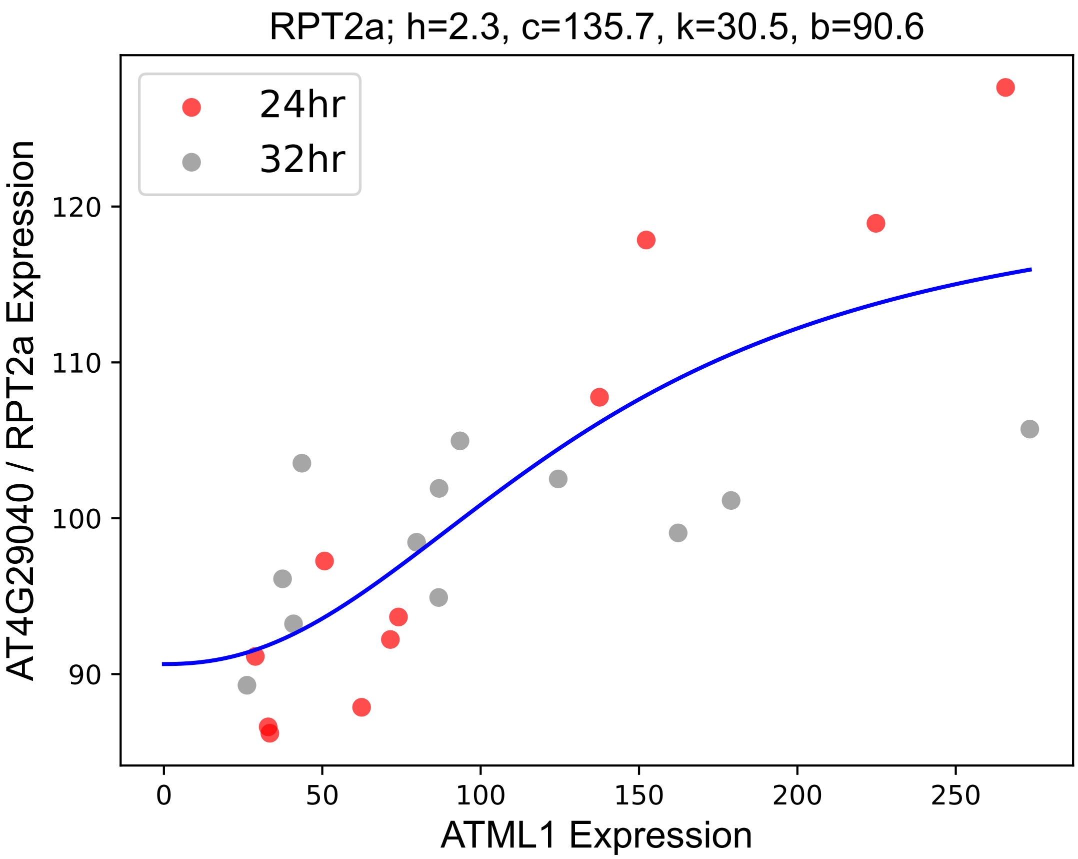

### RRT1.pdf

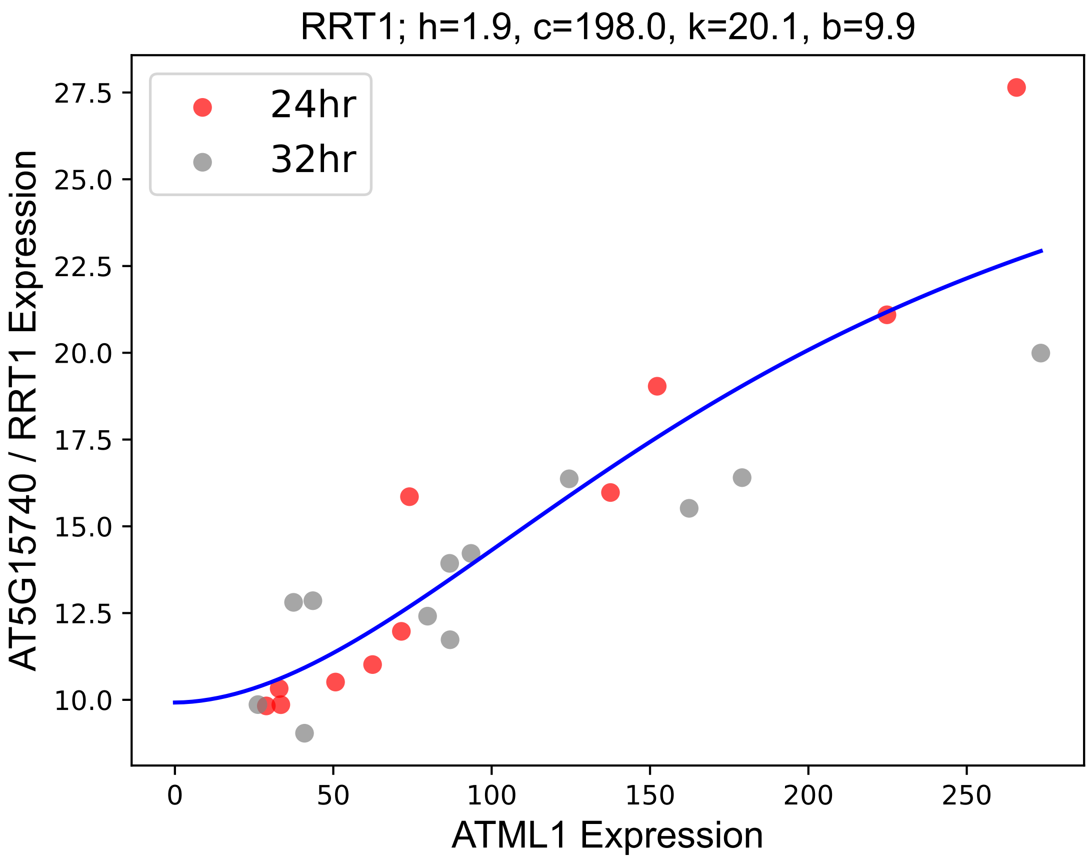

### SCPL9.pdf

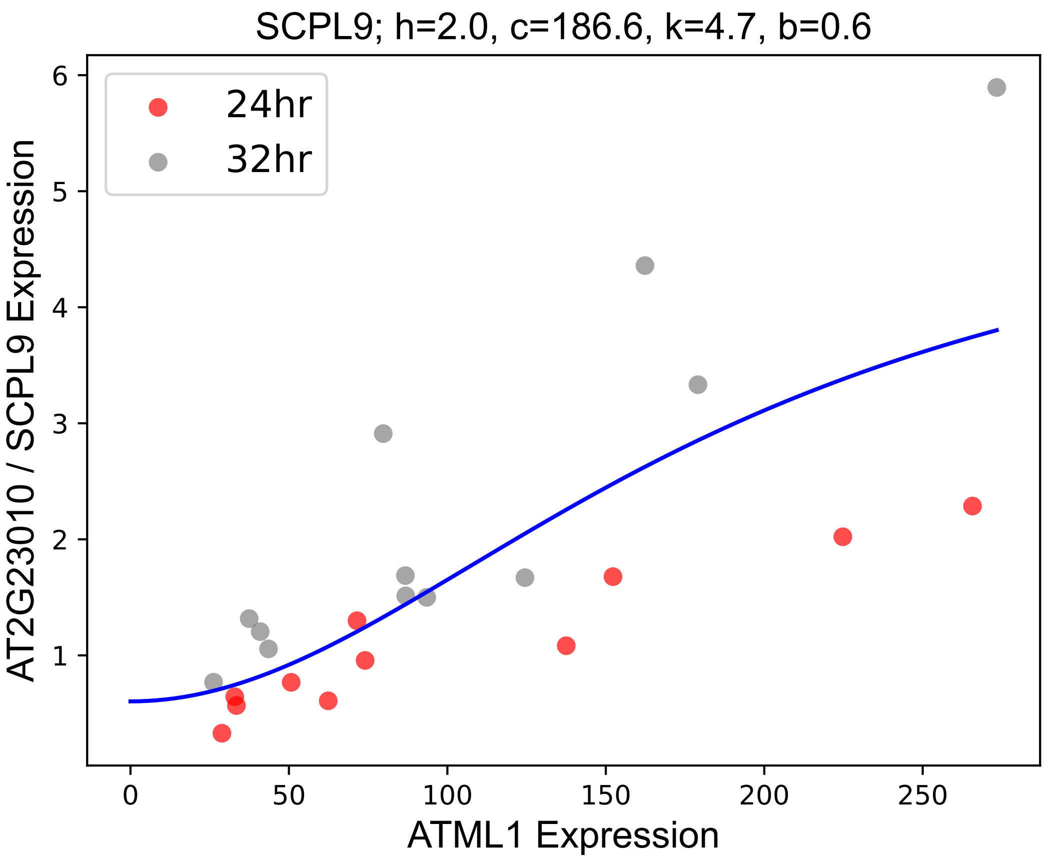

### SEP1.pdf

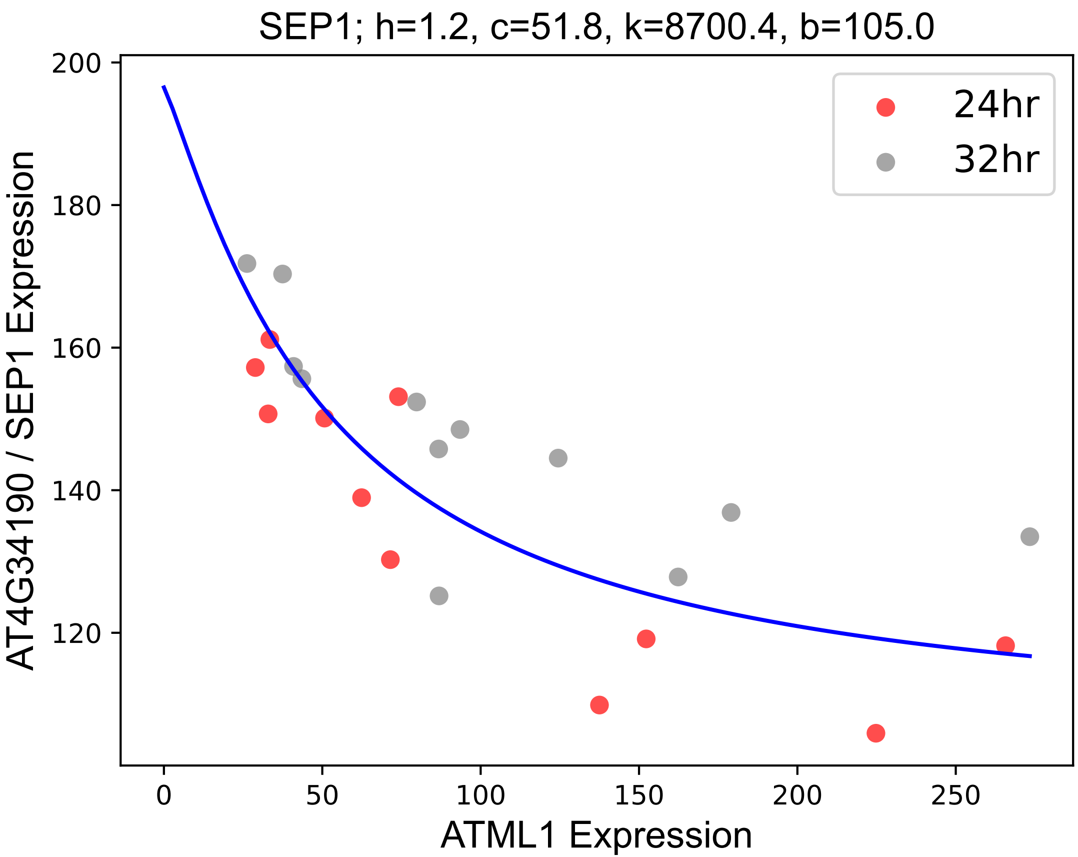

### SKS6.pdf

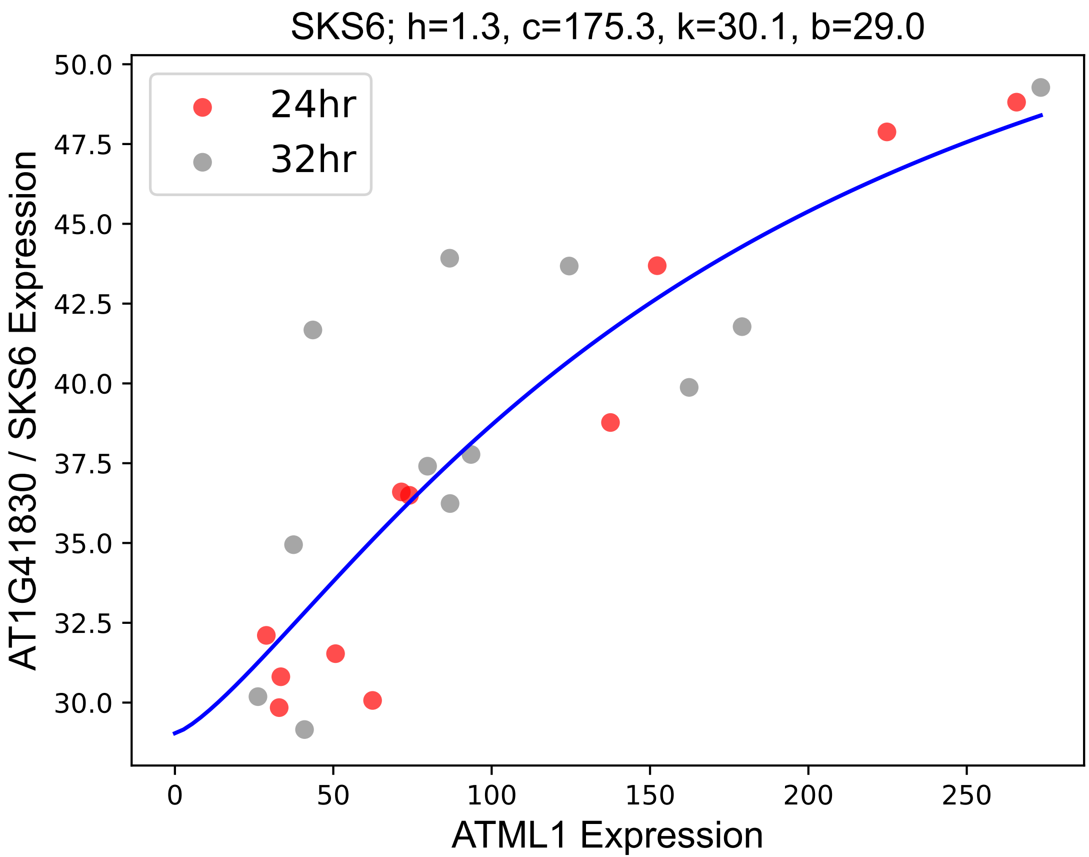

### SRC2.pdf

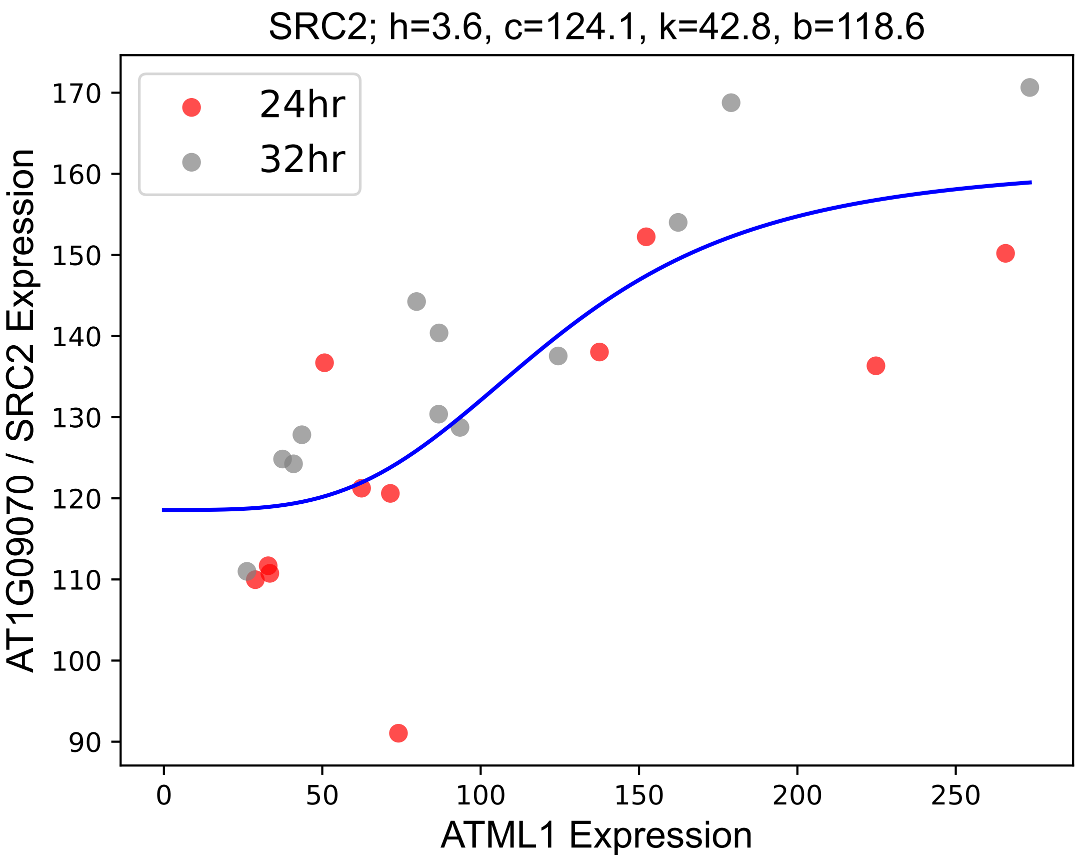

### ST2B.pdf

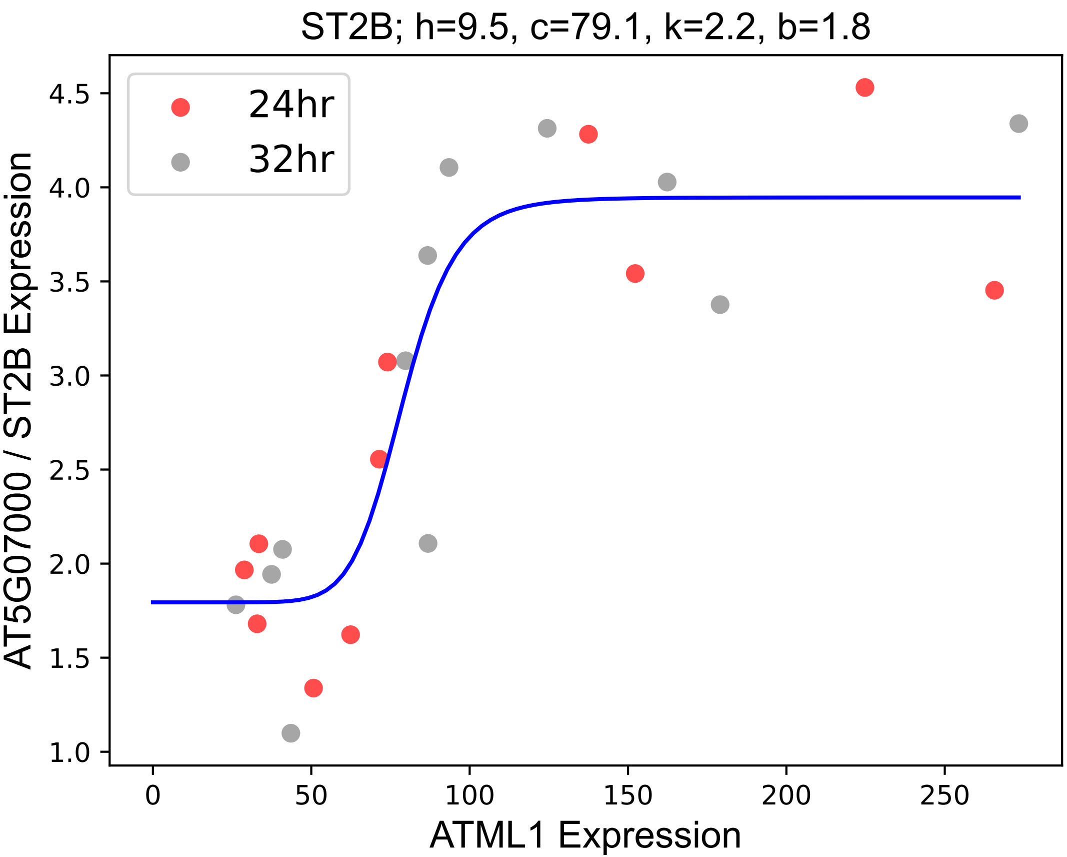

### SVB.pdf

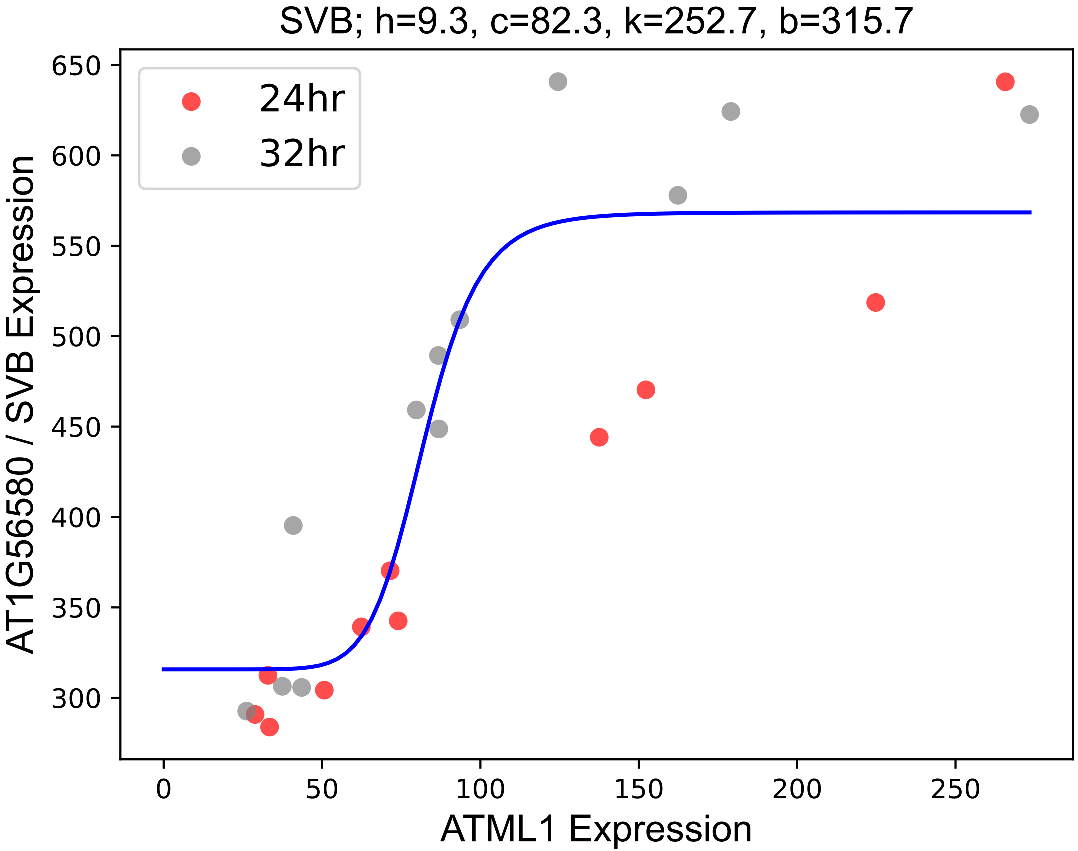

### SWIB2.pdf

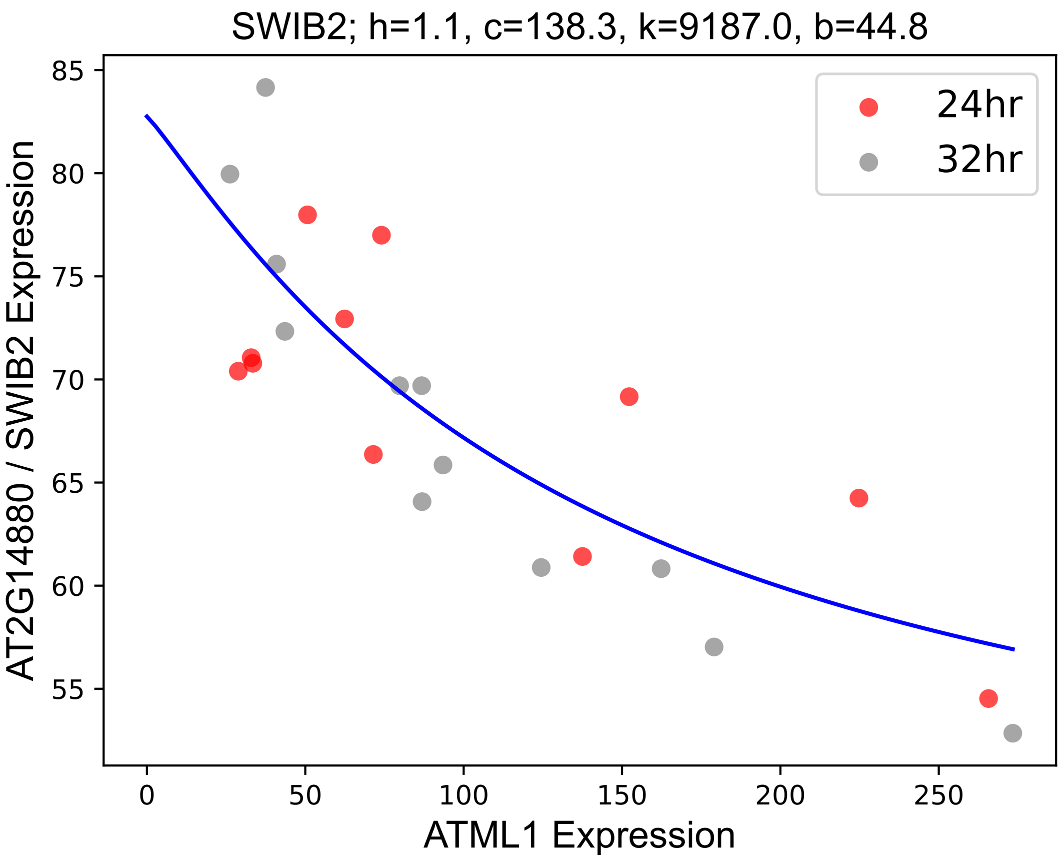

### TMM.pdf

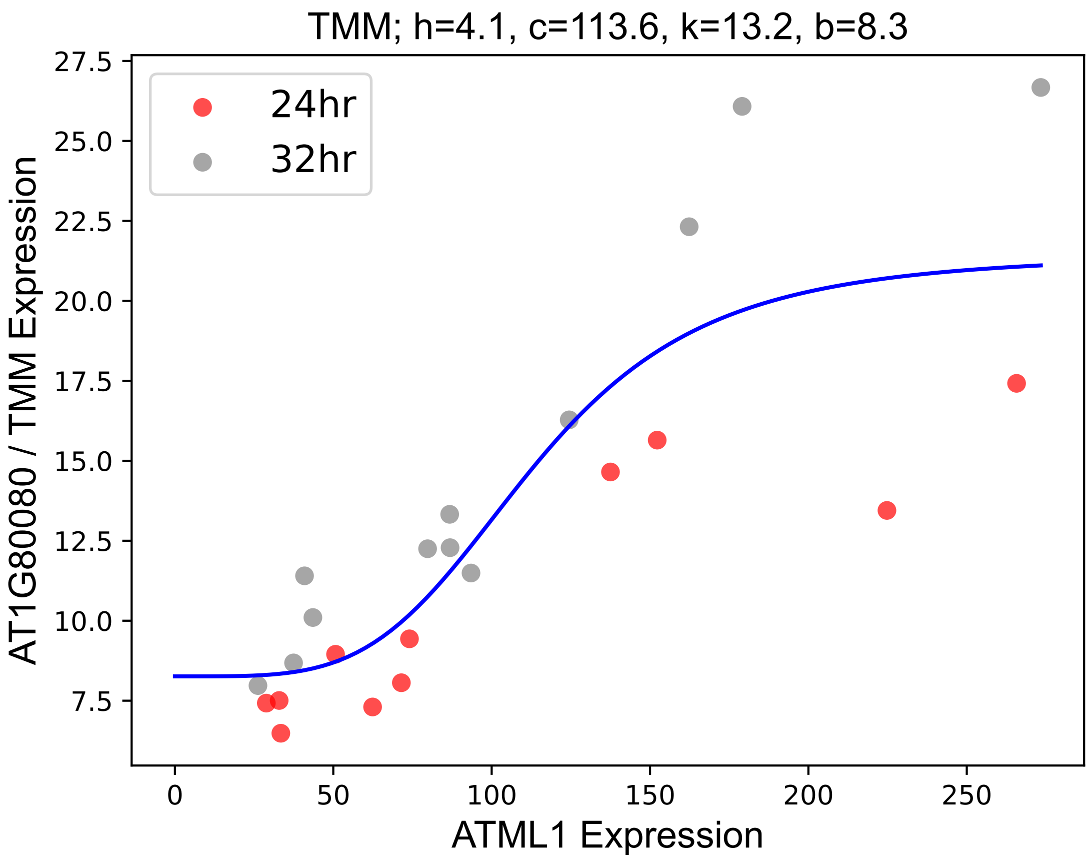

### UGD1.pdf

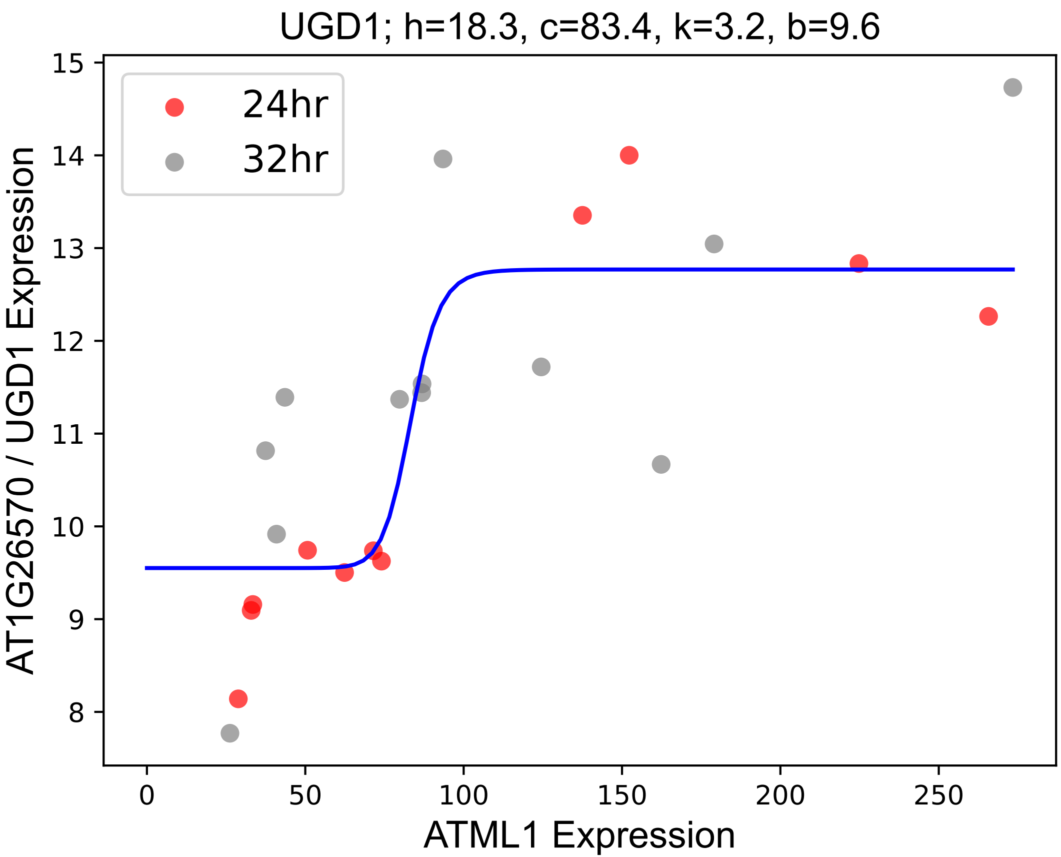

### UGT72E1.pdf

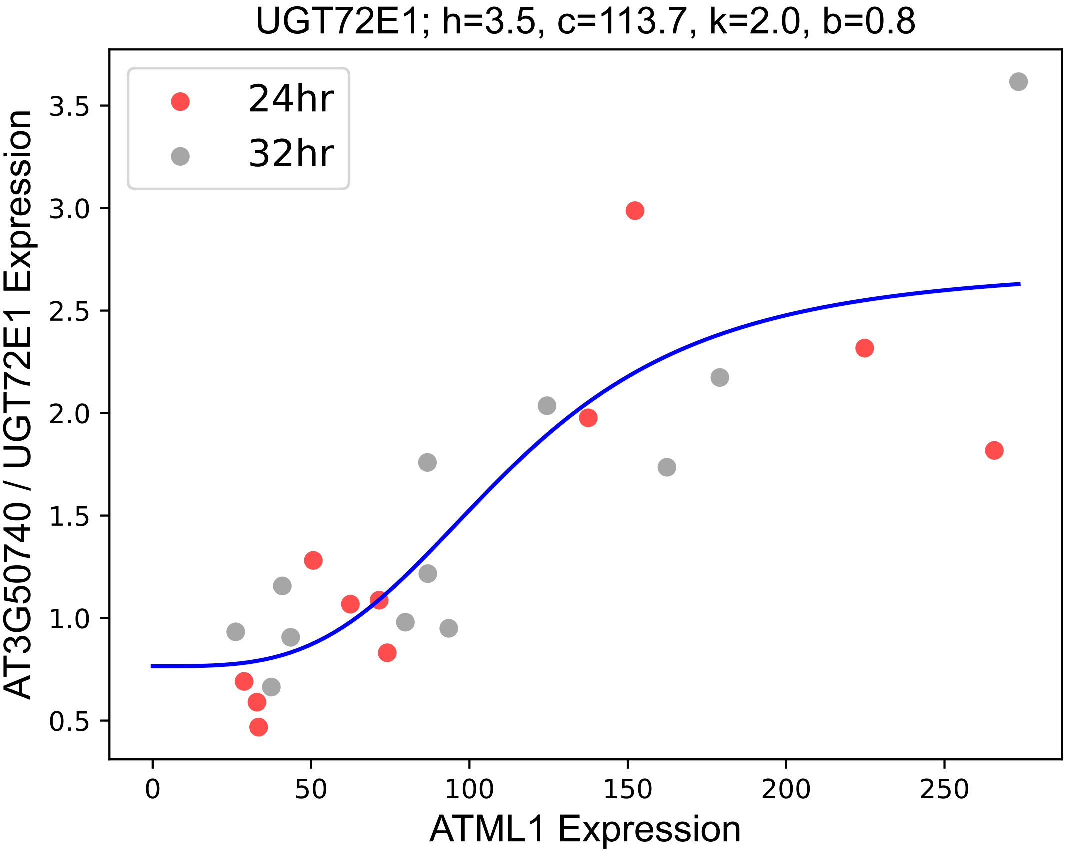

### UGT73B3.pdf

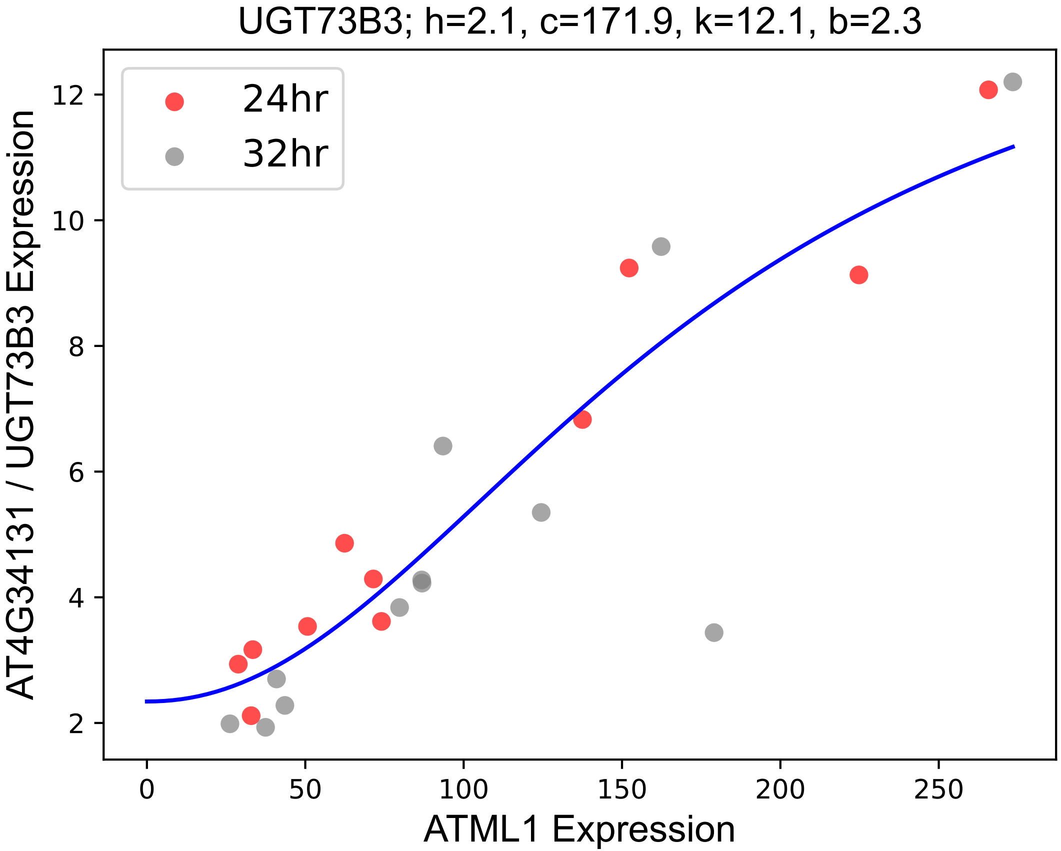

### UGT73B5.pdf

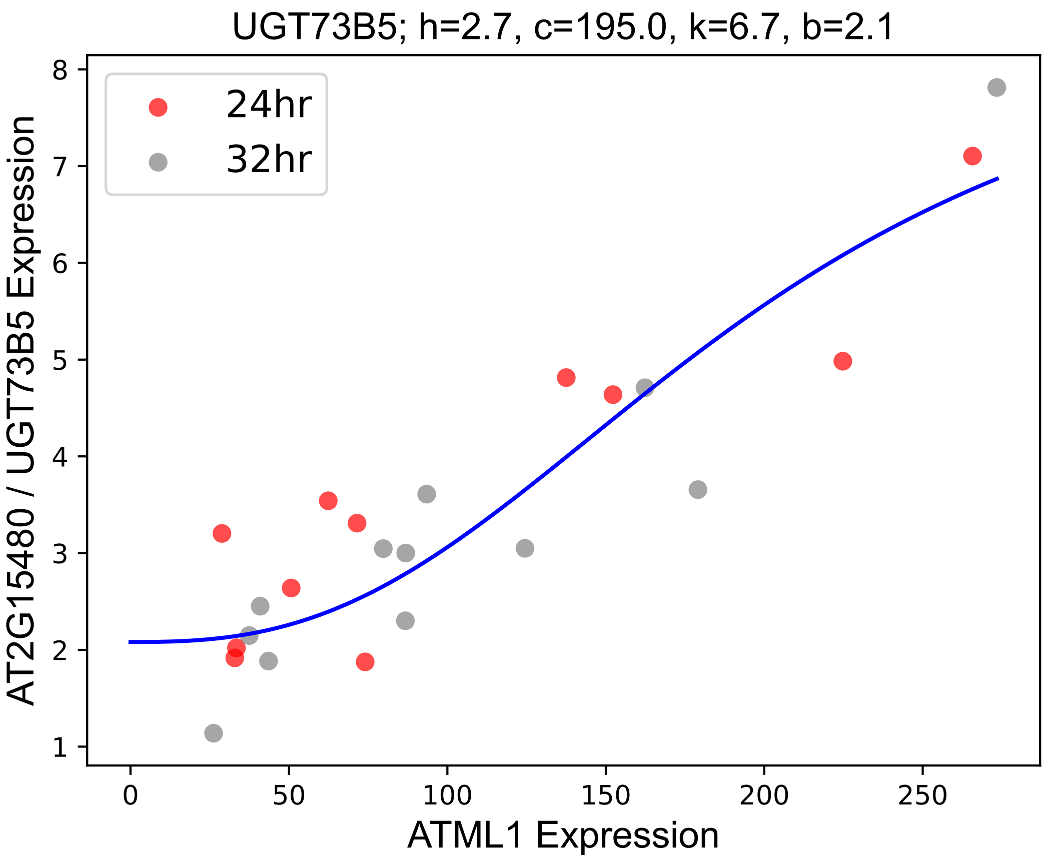

### VPT1.pdf

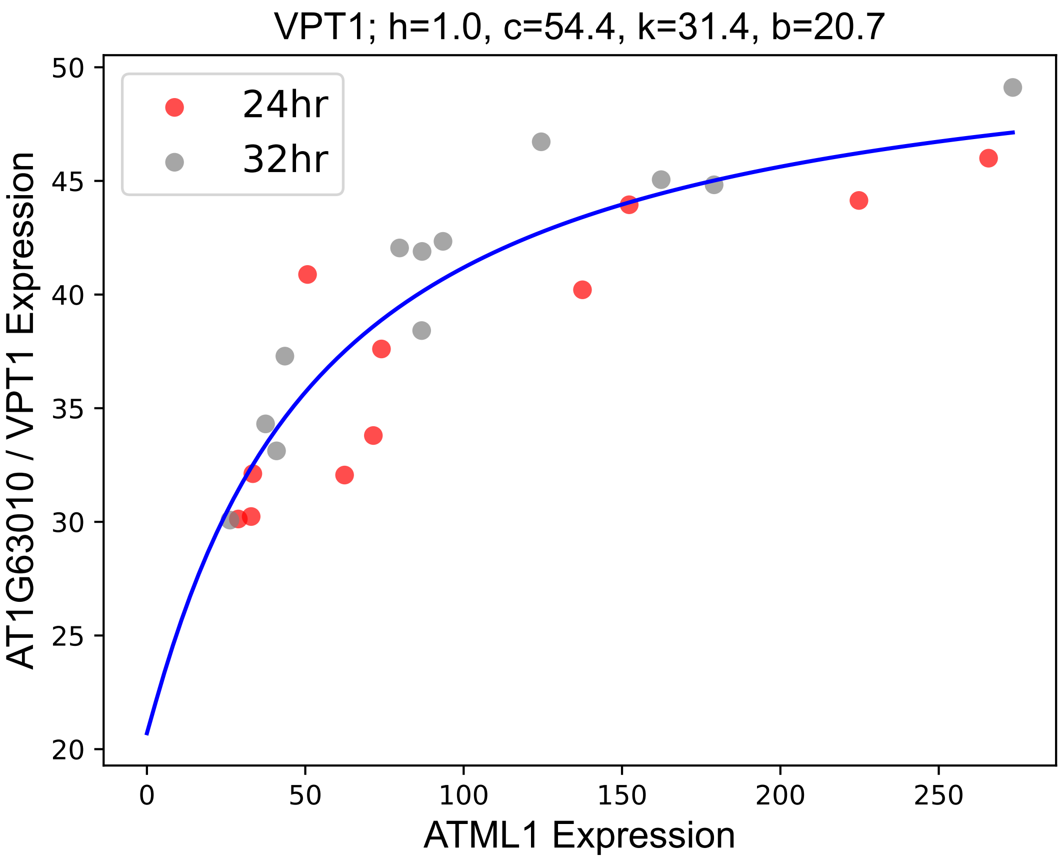

### WAVE2.pdf

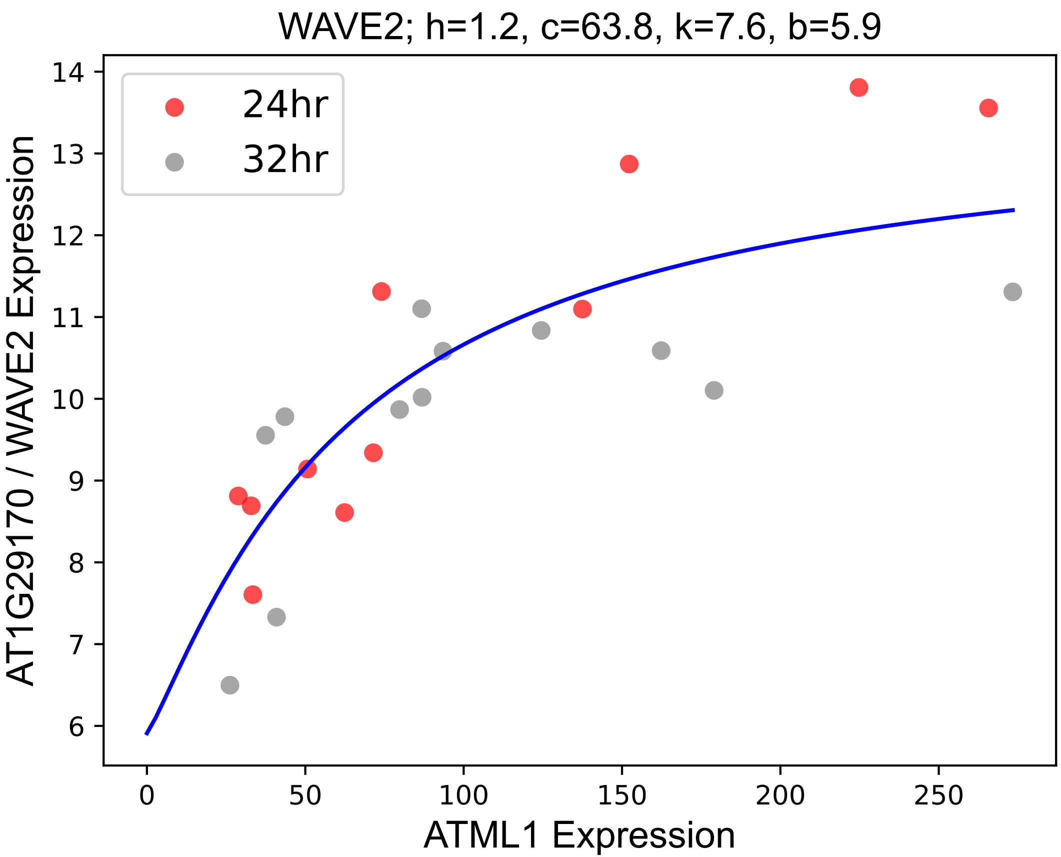

### XTH31.pdf

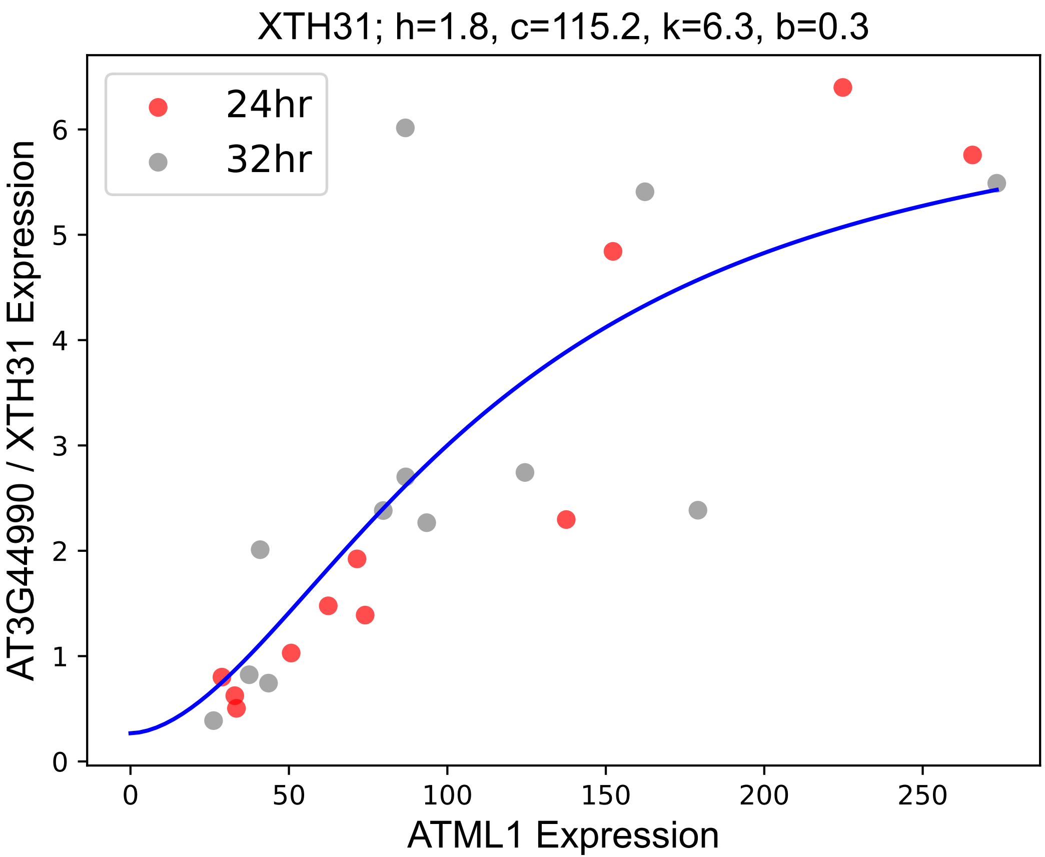

### YCF1.2.pdf

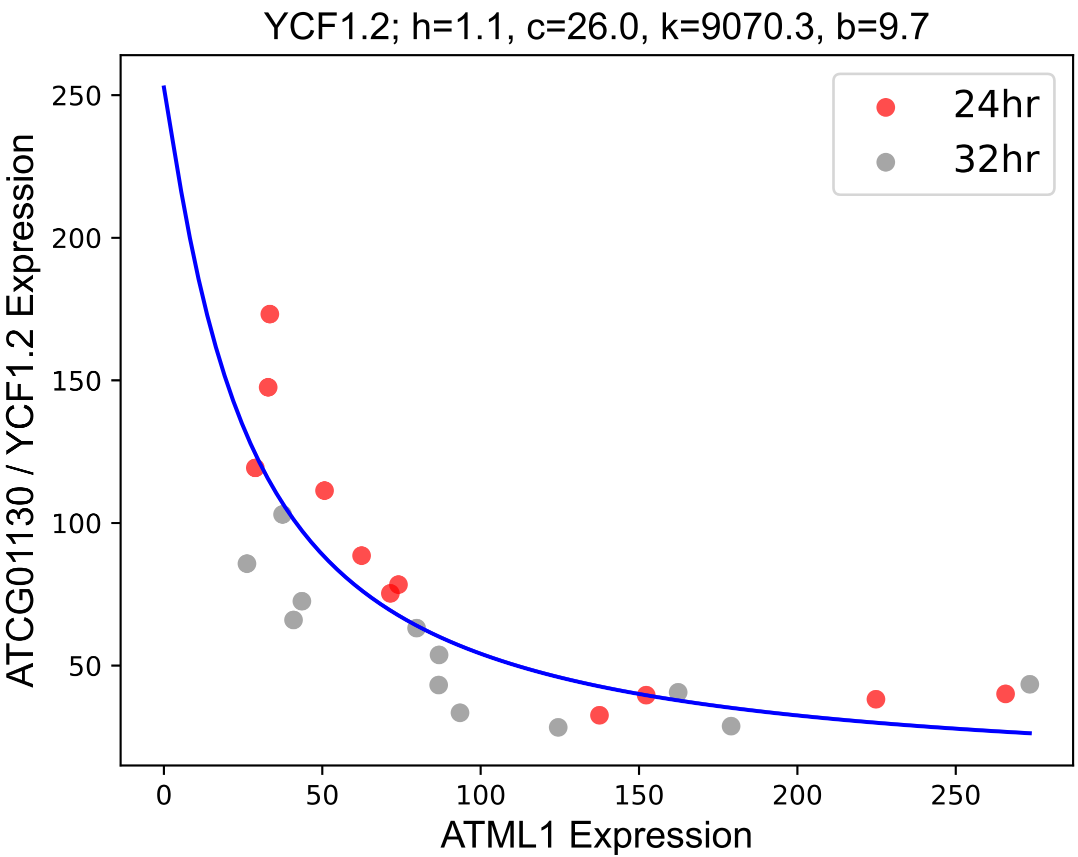

### YCF5.pdf

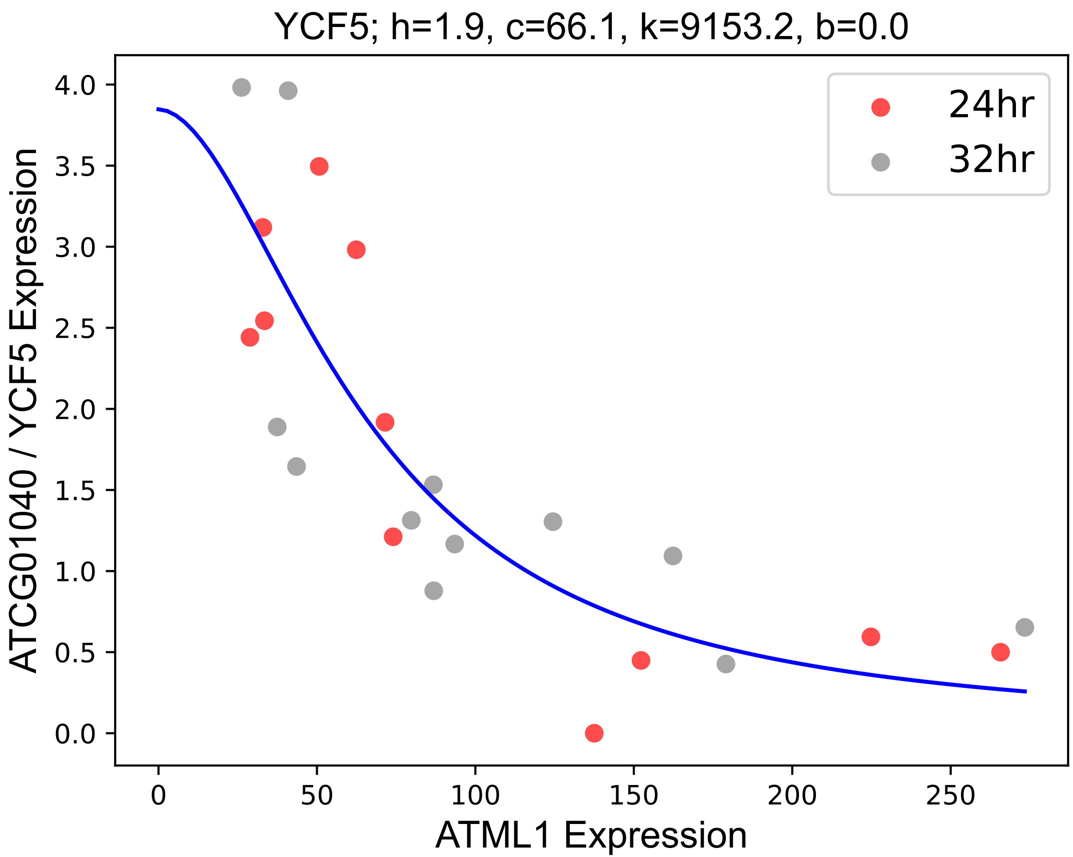
