## Supplementary figures and images for "The transcription factor ATML1 maintains giant cell identity by inducing synthesis of its own long-chain fatty acid-containing ligands"

### AAP6.pdf

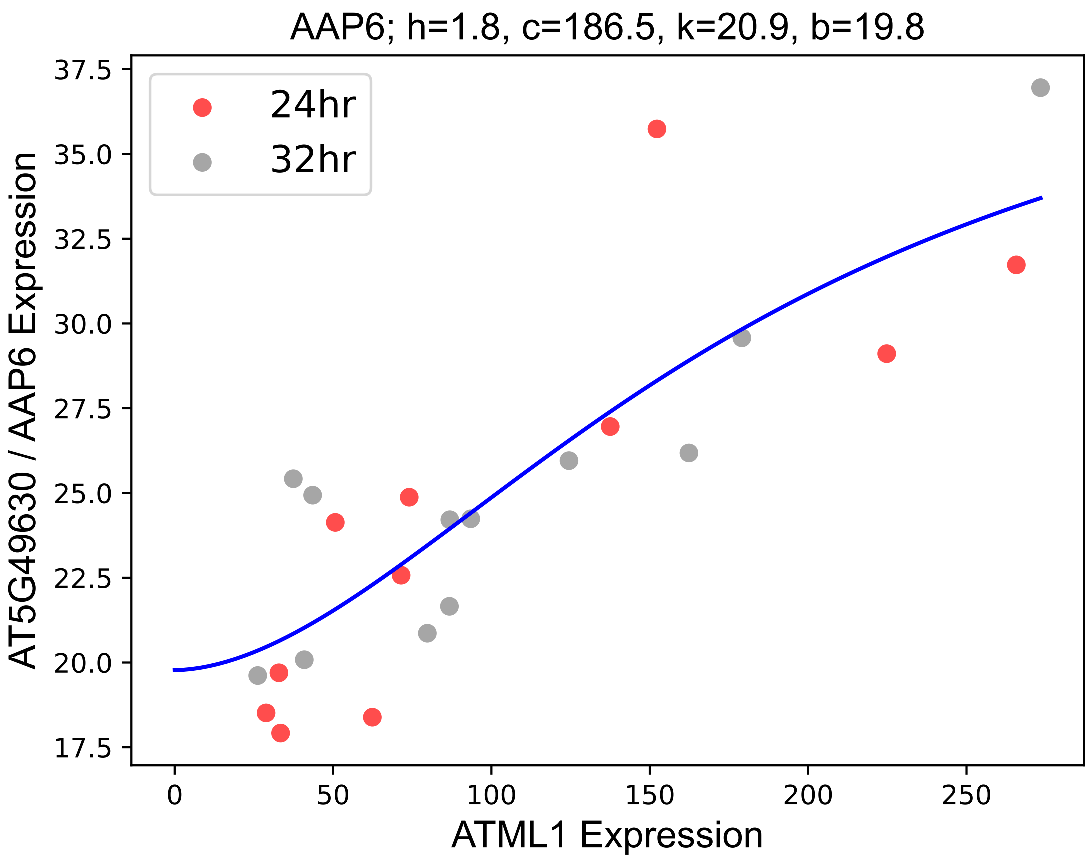

### ABCB2.pdf

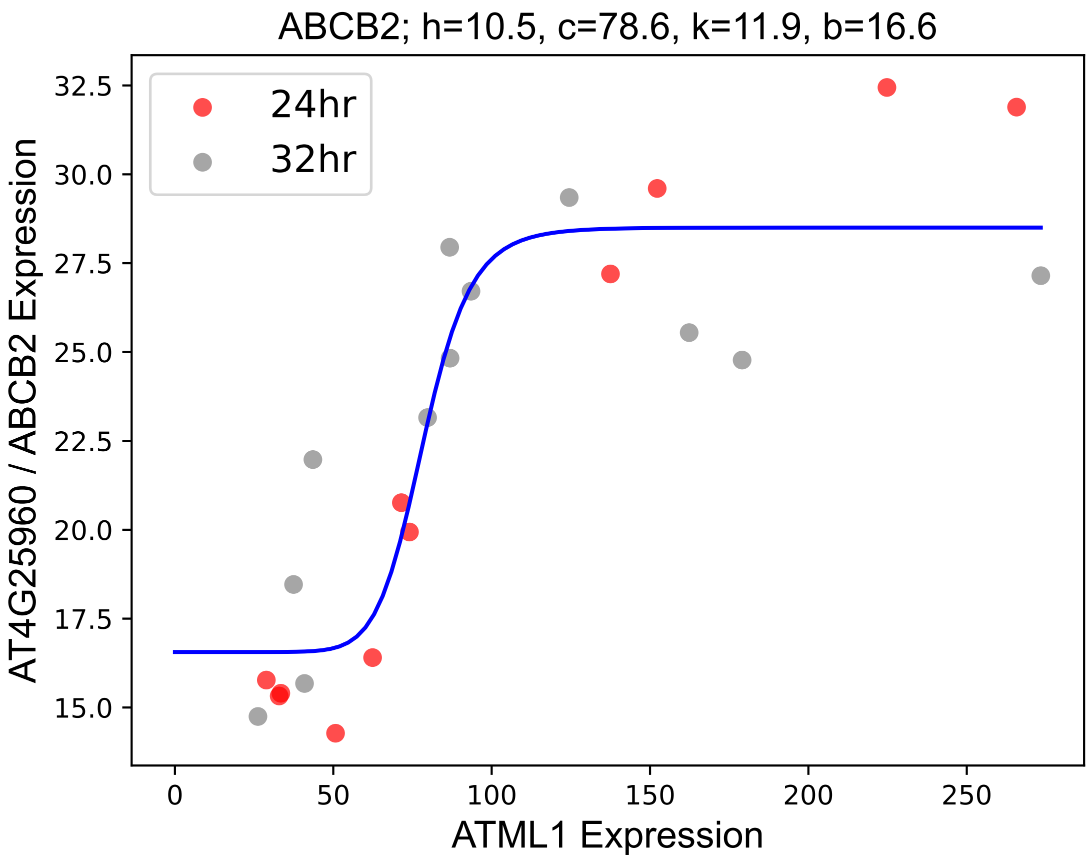

### ABCG11.pdf

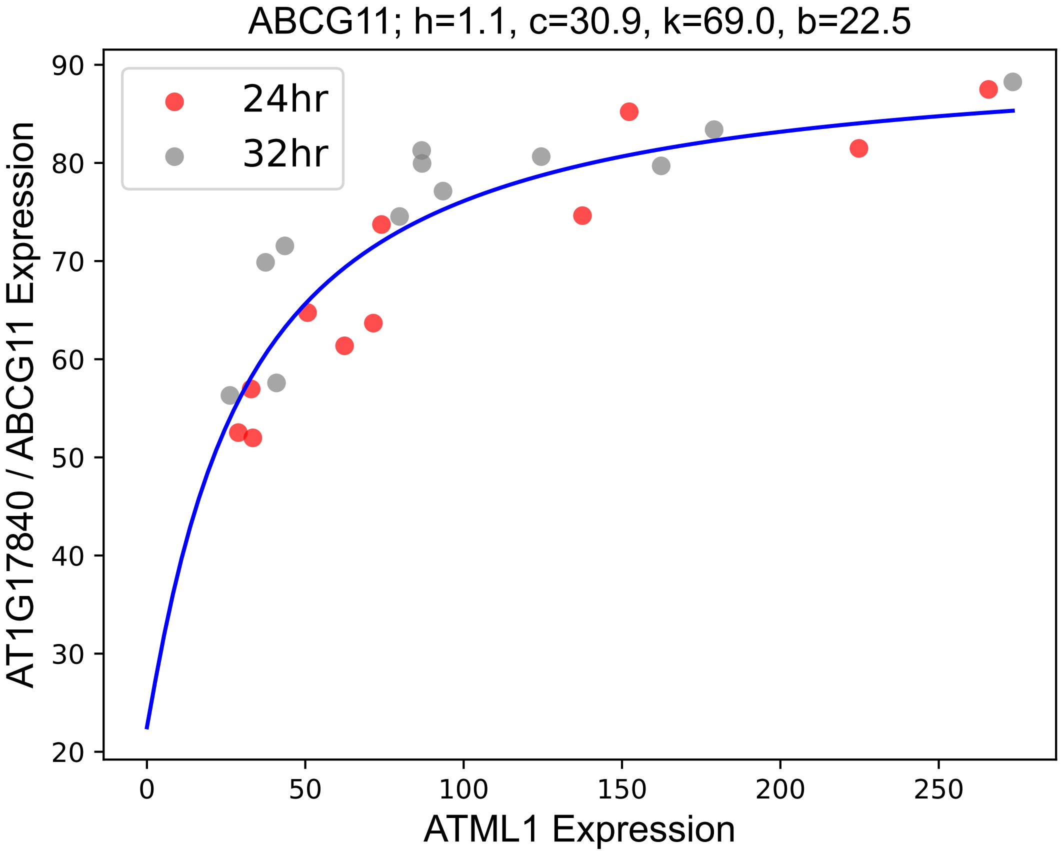

### ABCG25.pdf

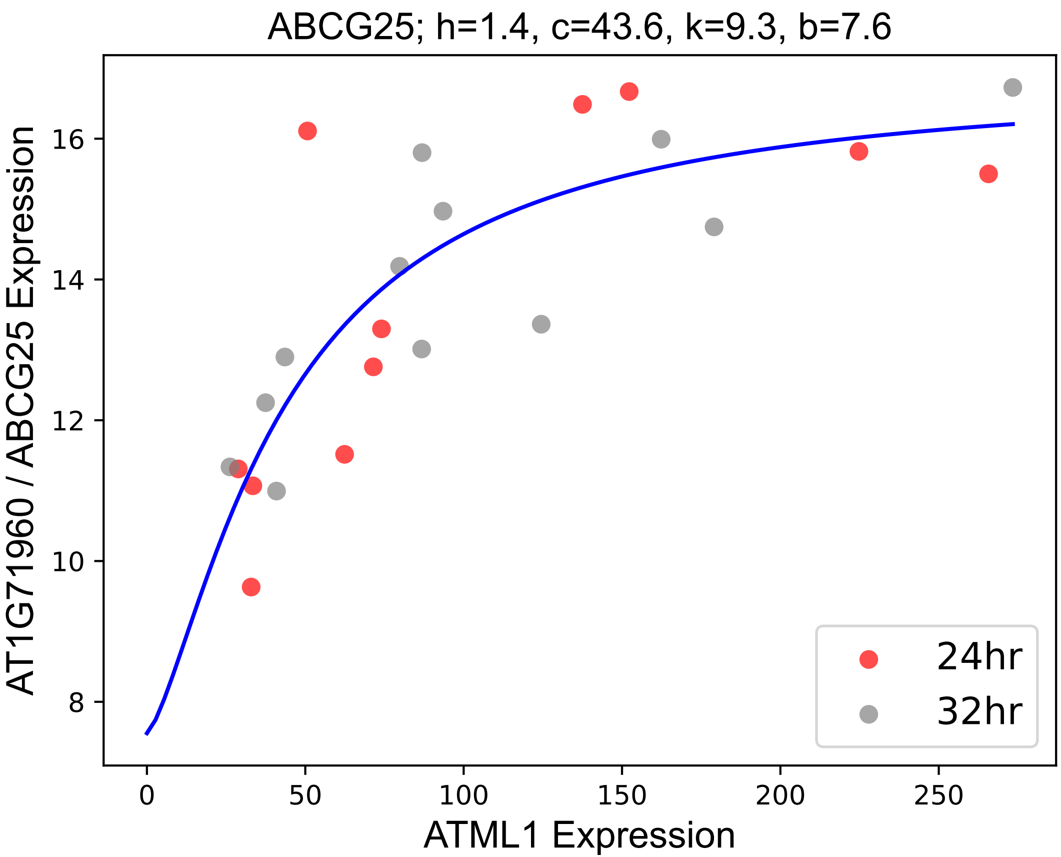

### ABCG32.pdf

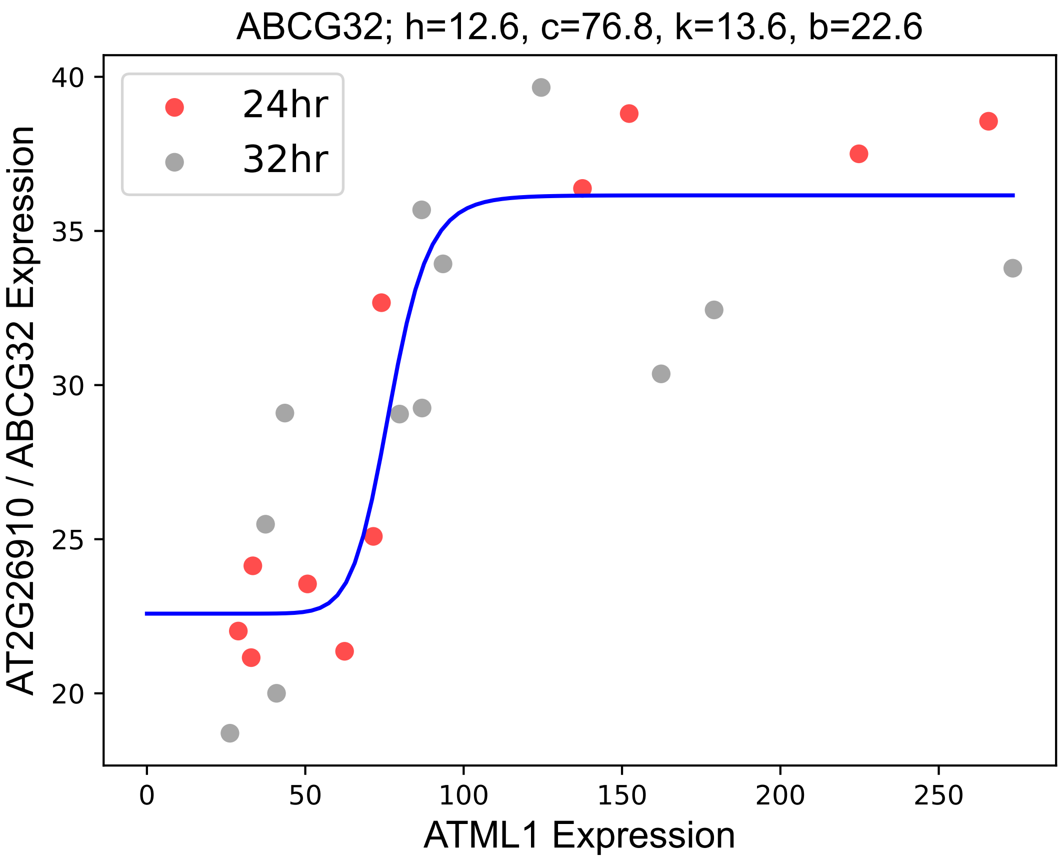

### ABCG40.pdf

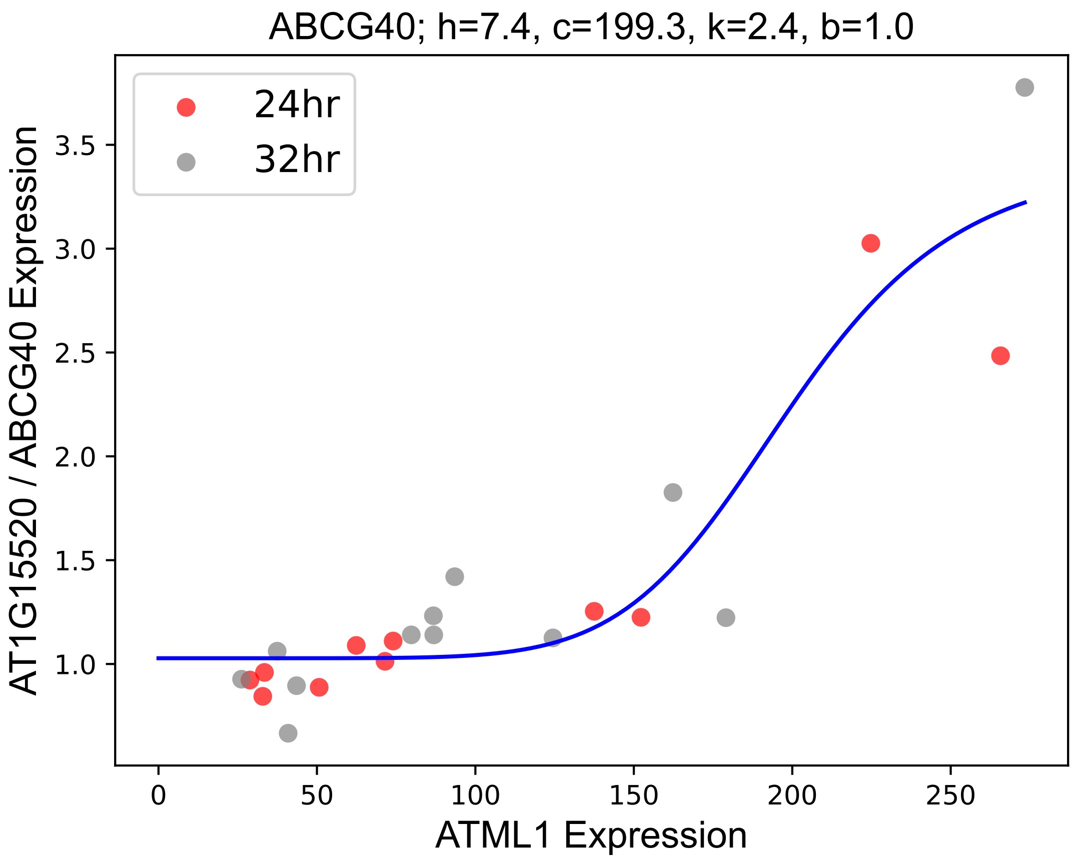

### ACX1.pdf

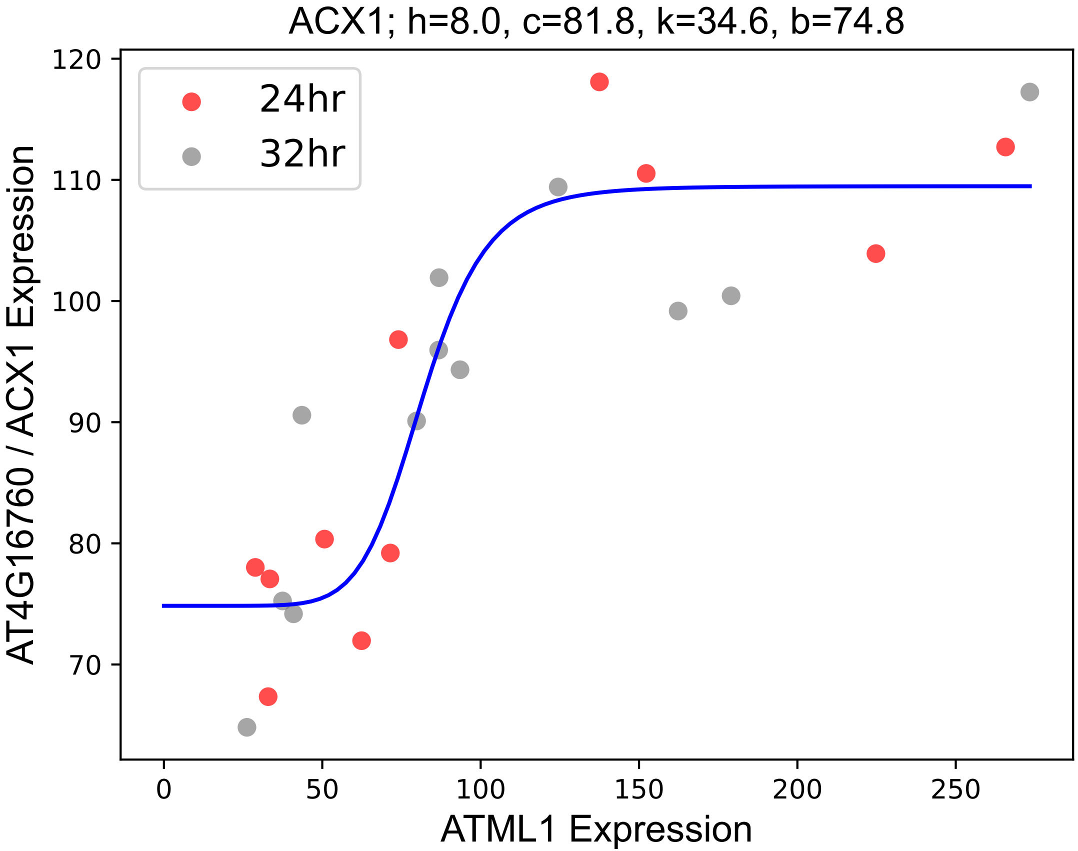

### ADC2.pdf

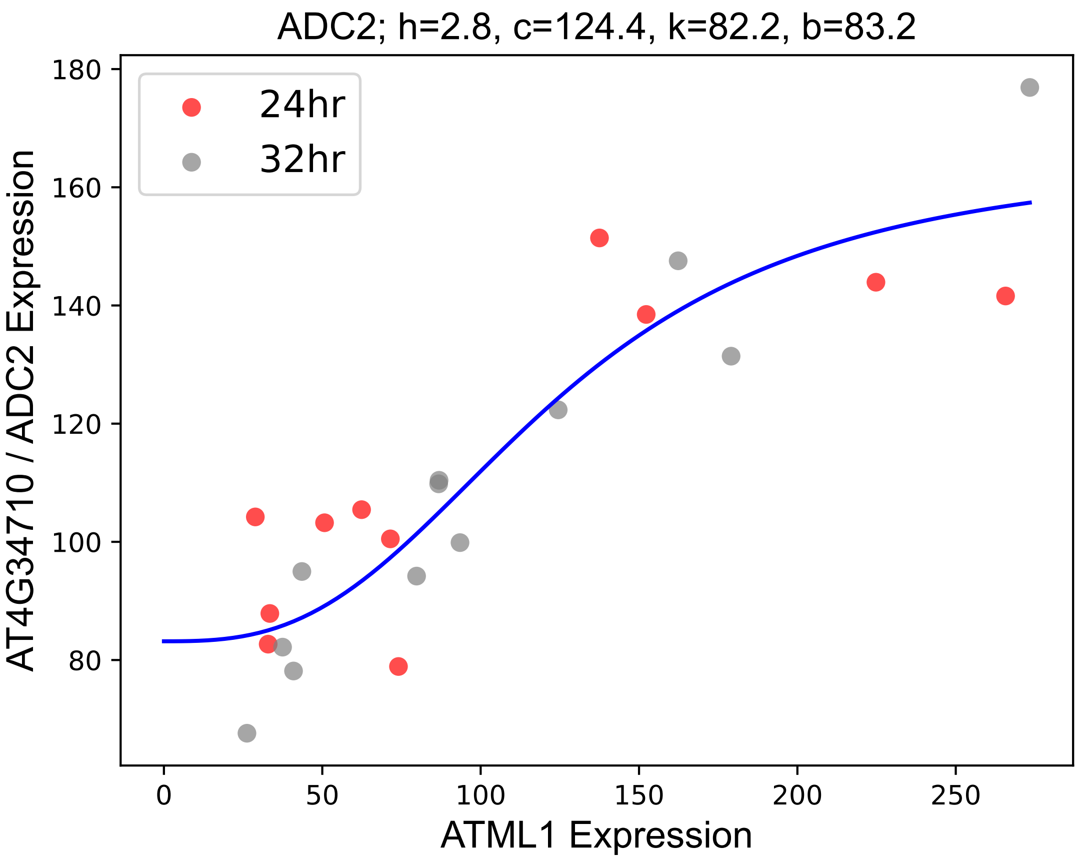

### ALF5.pdf

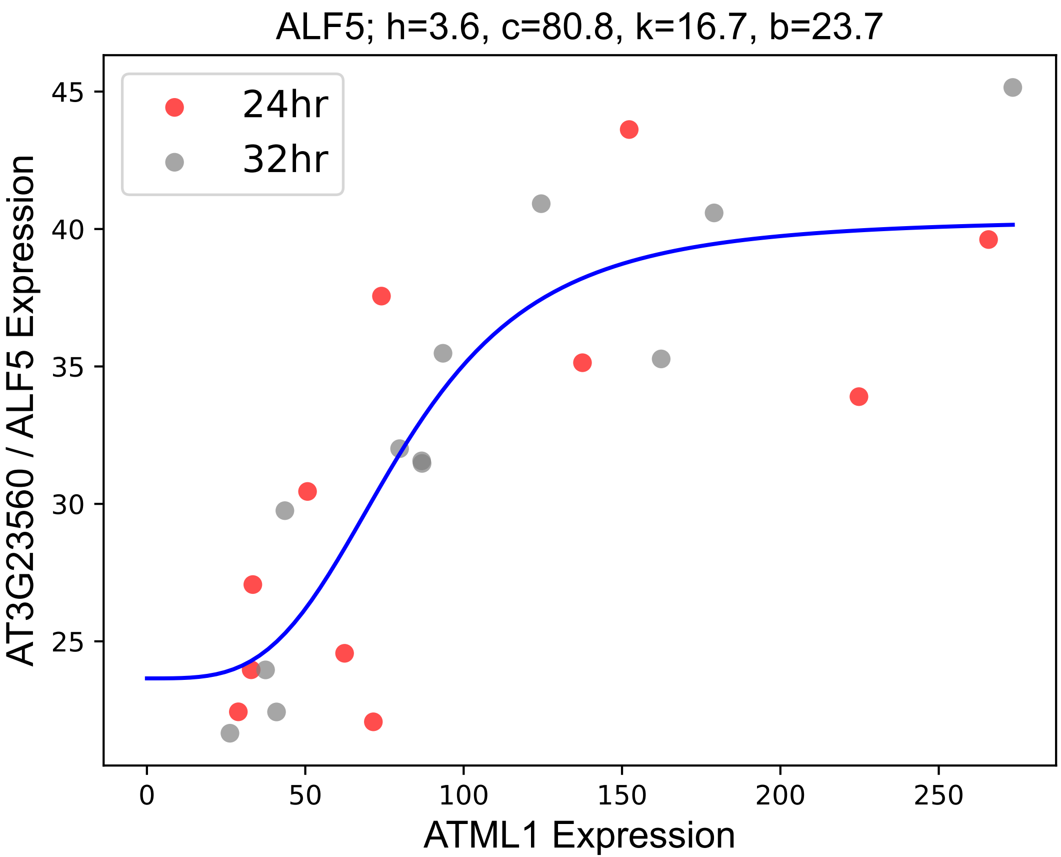

### ALKBH10B.pdf

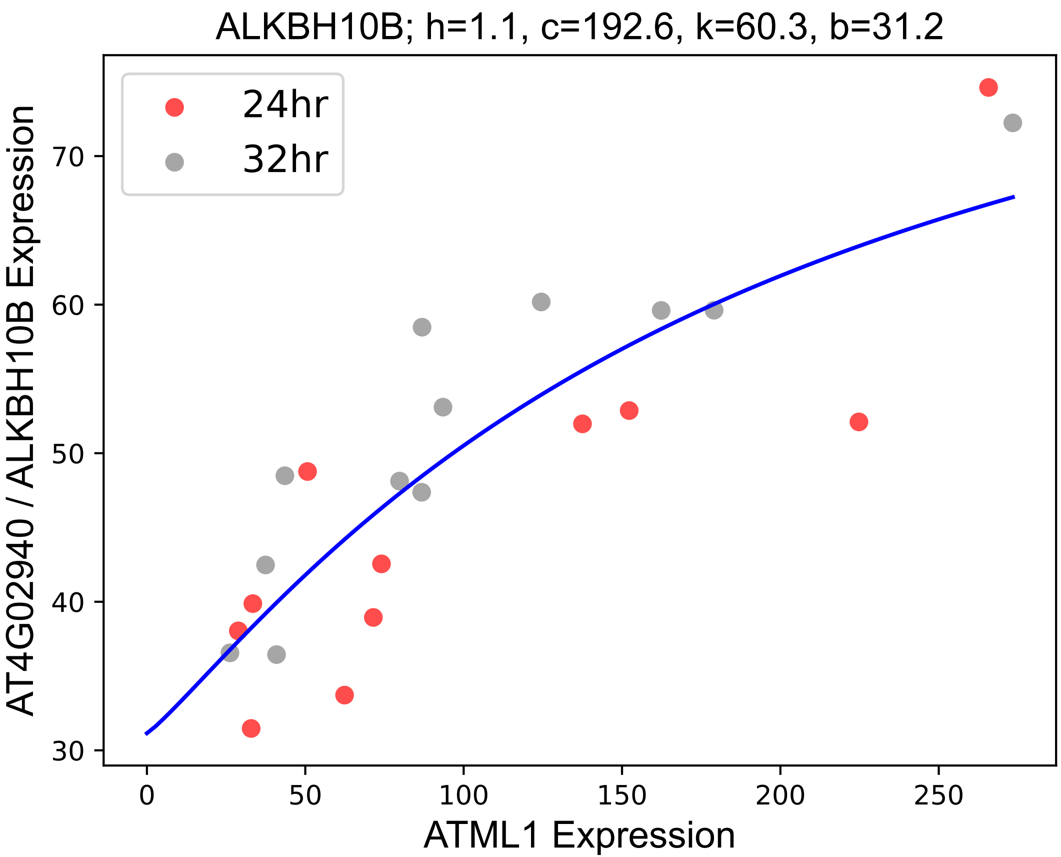

### ANNAT4.pdf

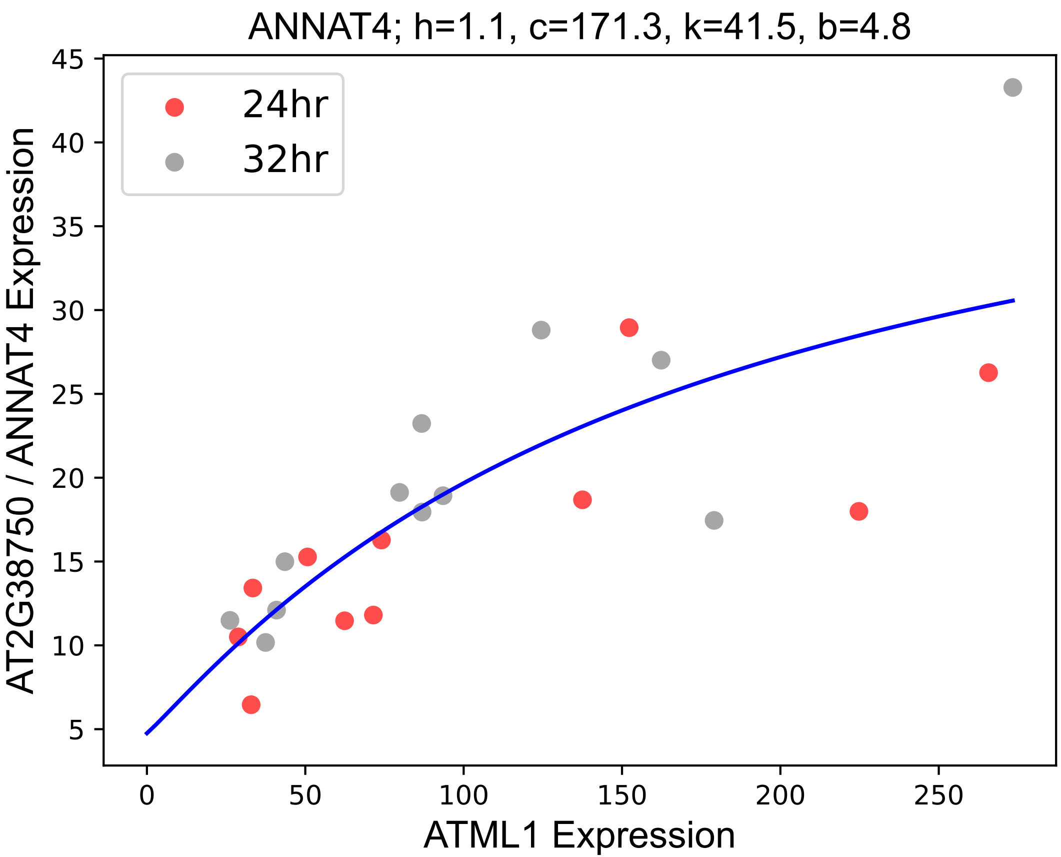

### AO1.pdf

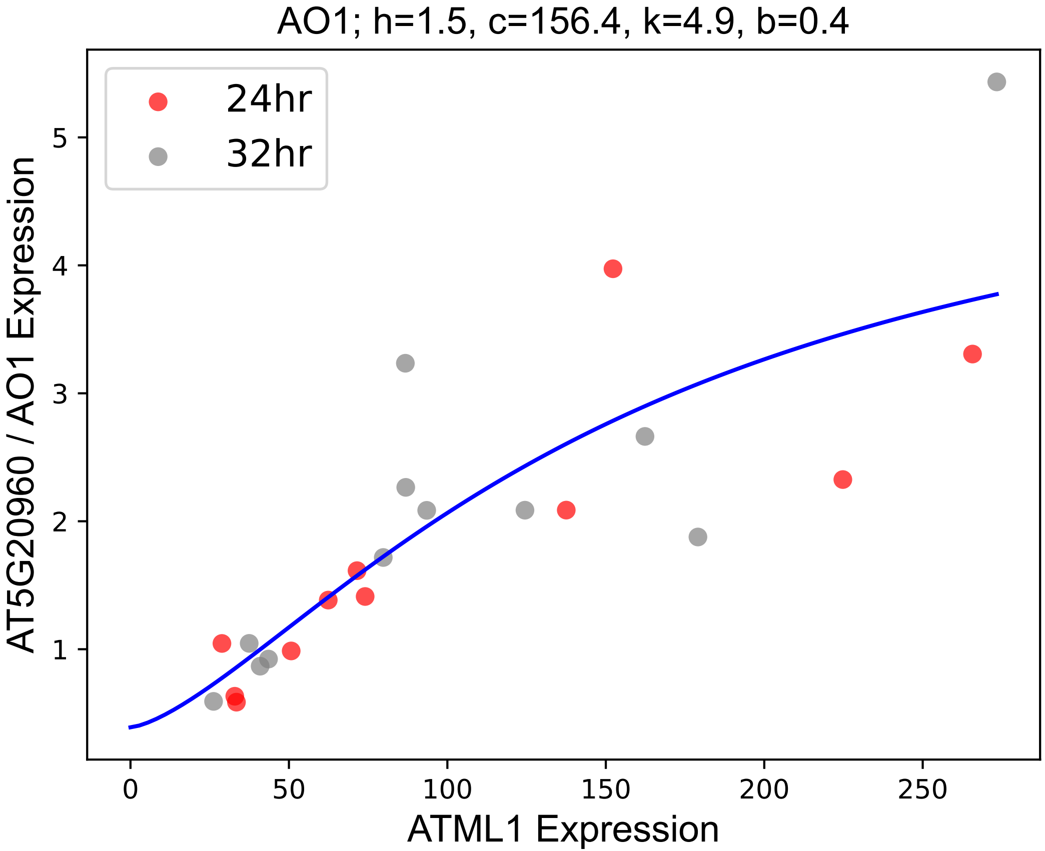

### APSE.pdf

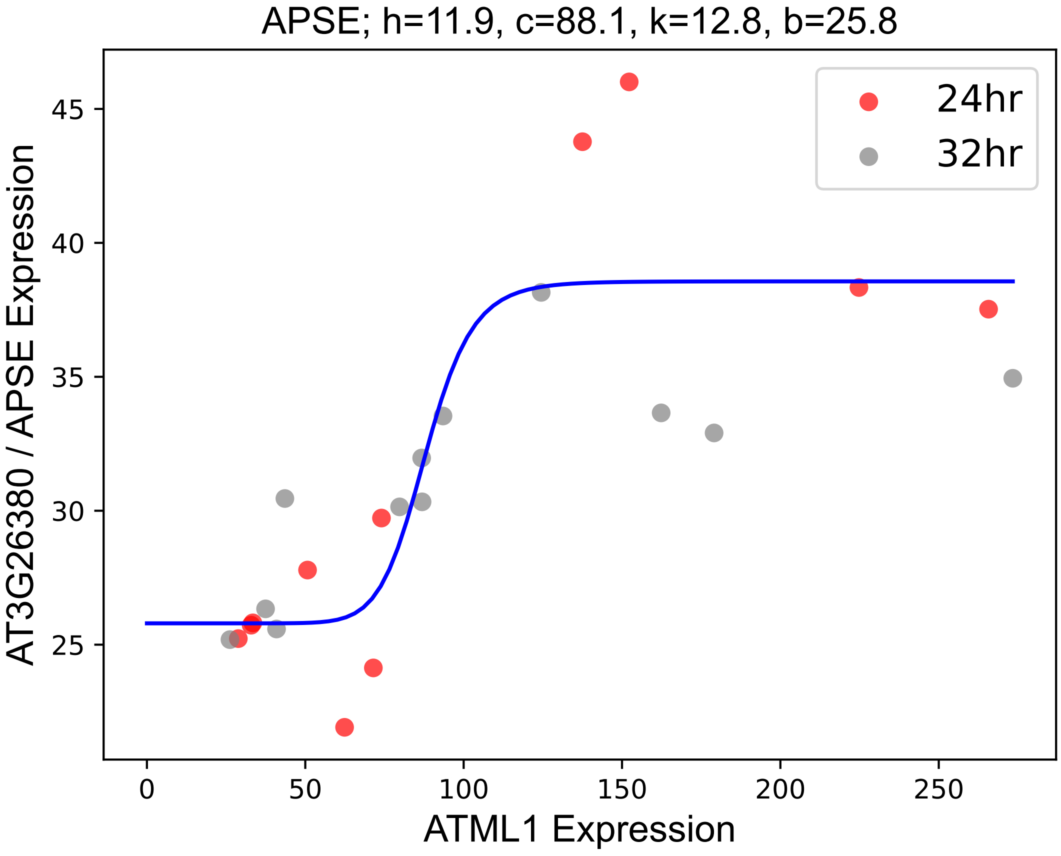

### ARA12.pdf

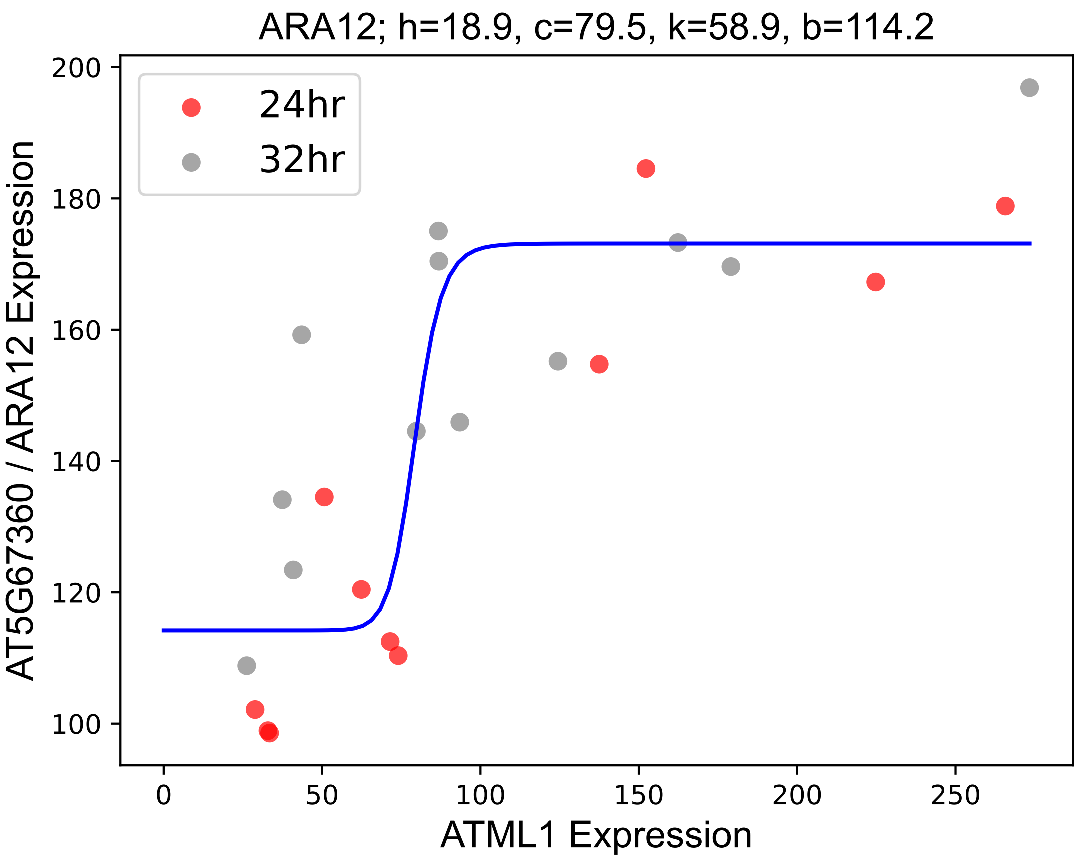

### AT1G09310.pdf

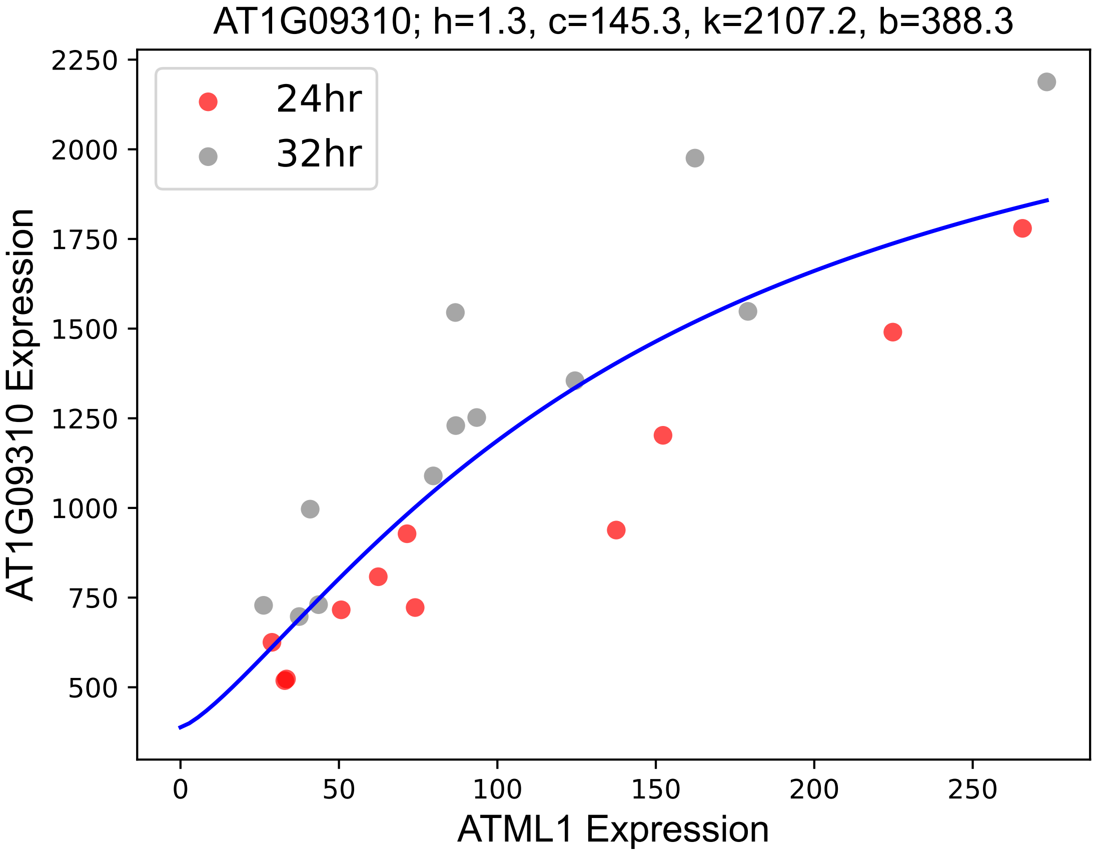

### AT1G17500.pdf

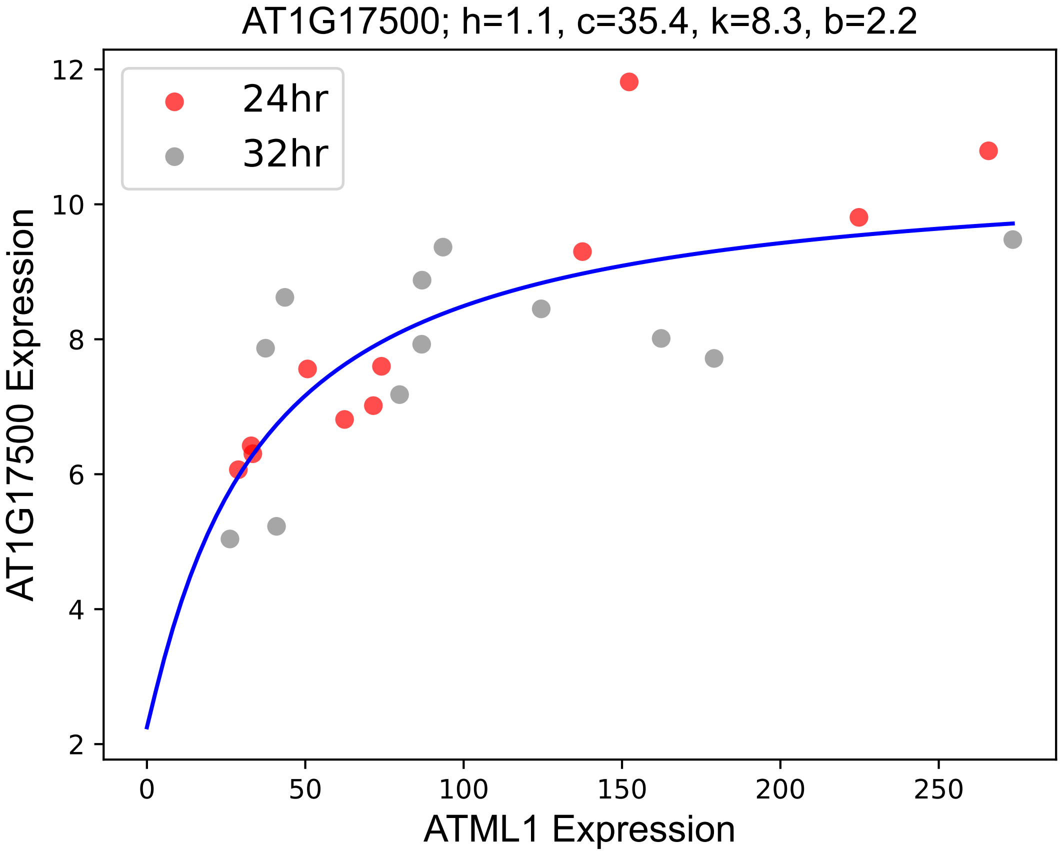

### AT1G19430.pdf

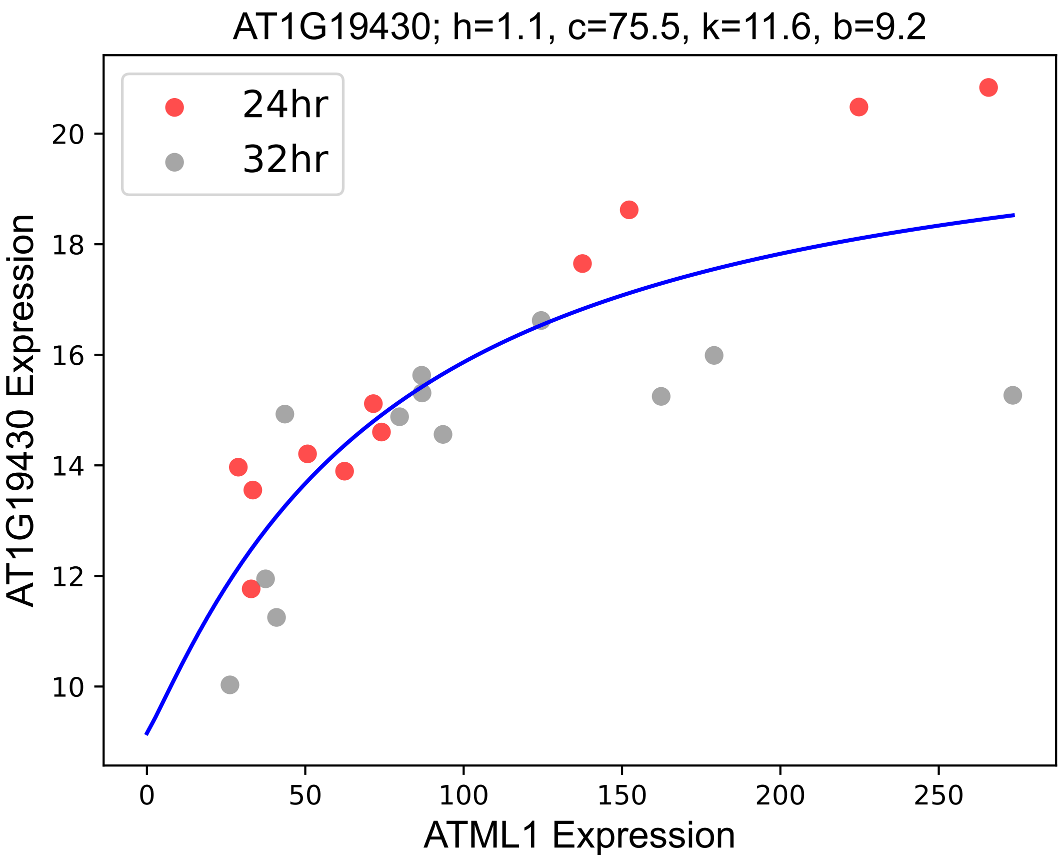

### AT1G20810.pdf

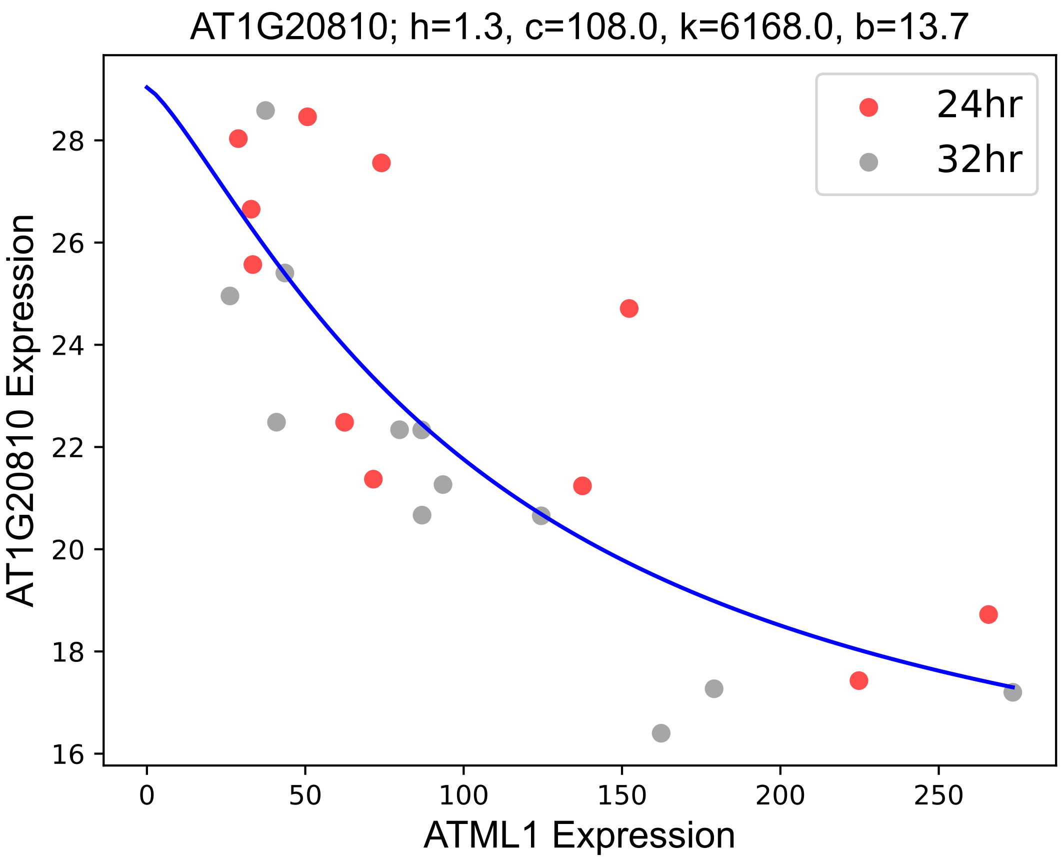

### AT1G24996.pdf

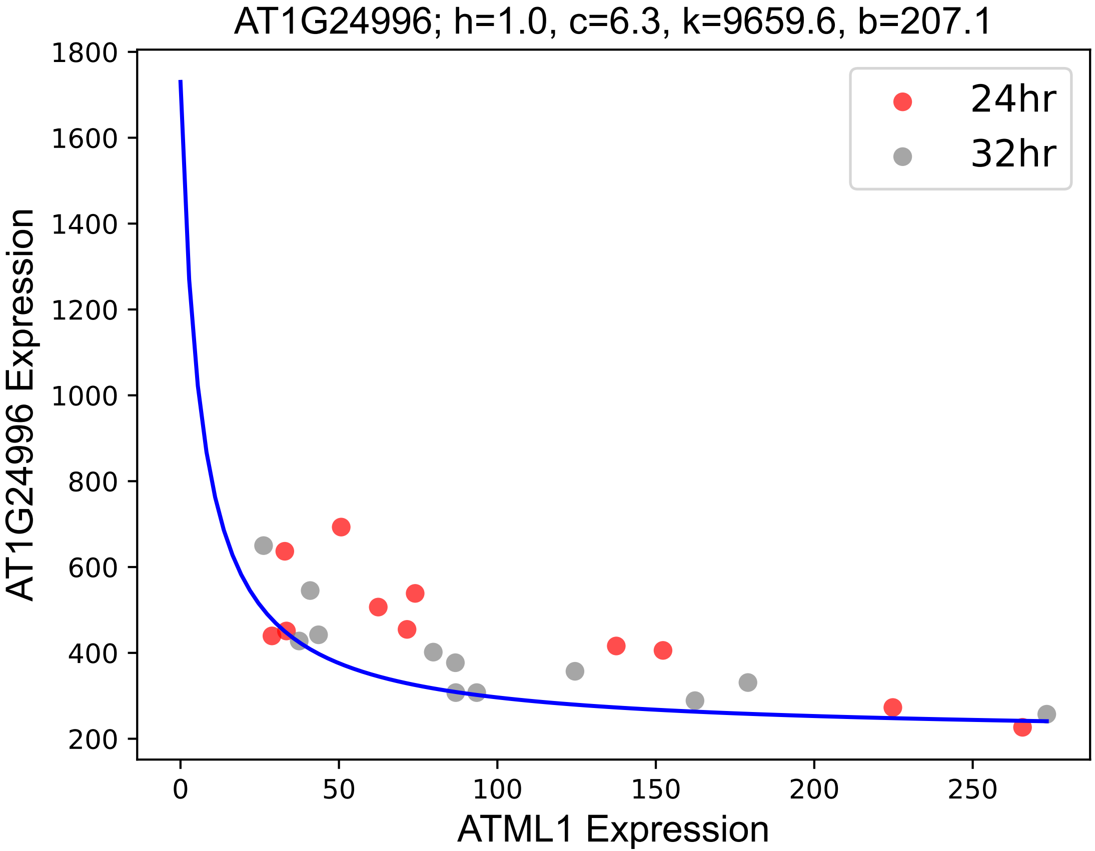

### AT1G52720.pdf

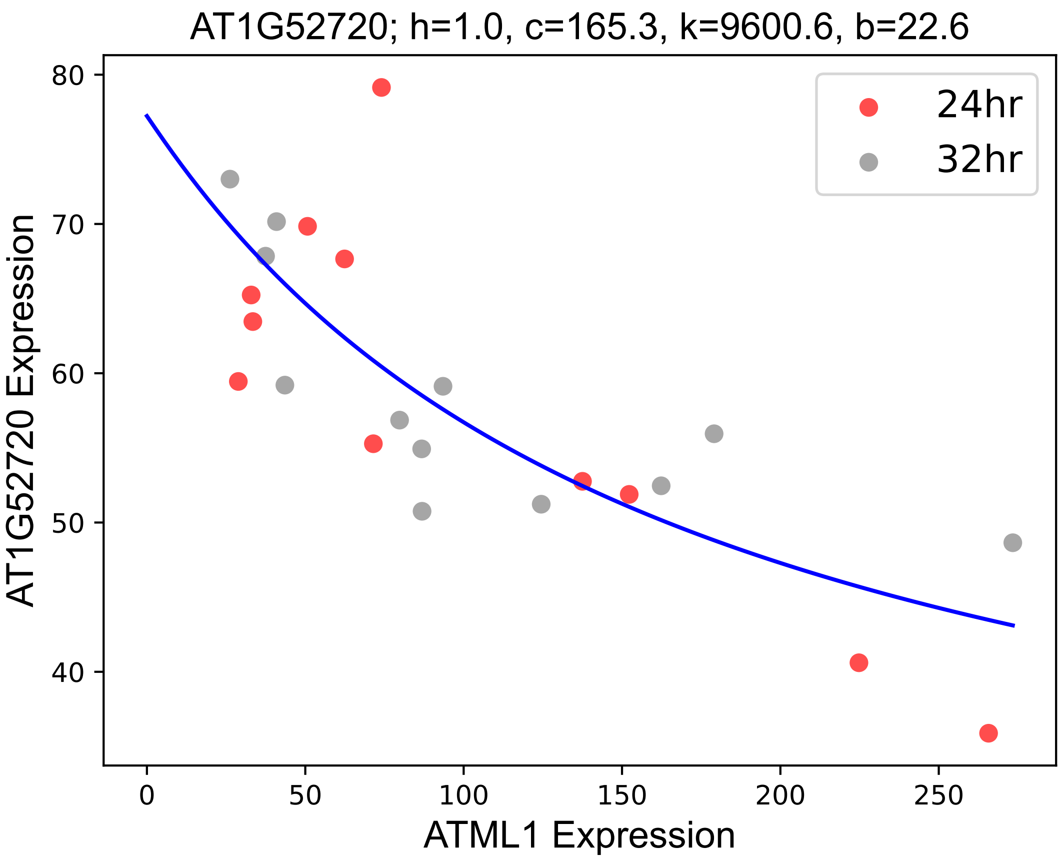

### AT1G61890.pdf

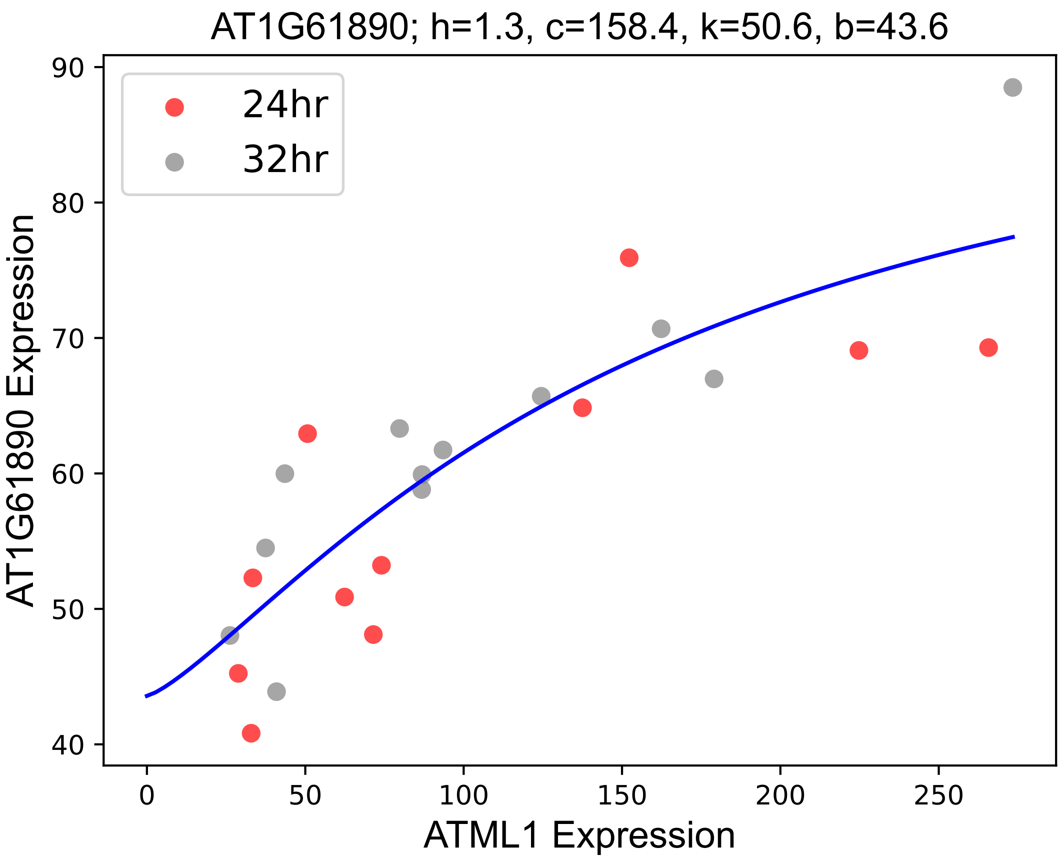

### AT1G62975.pdf

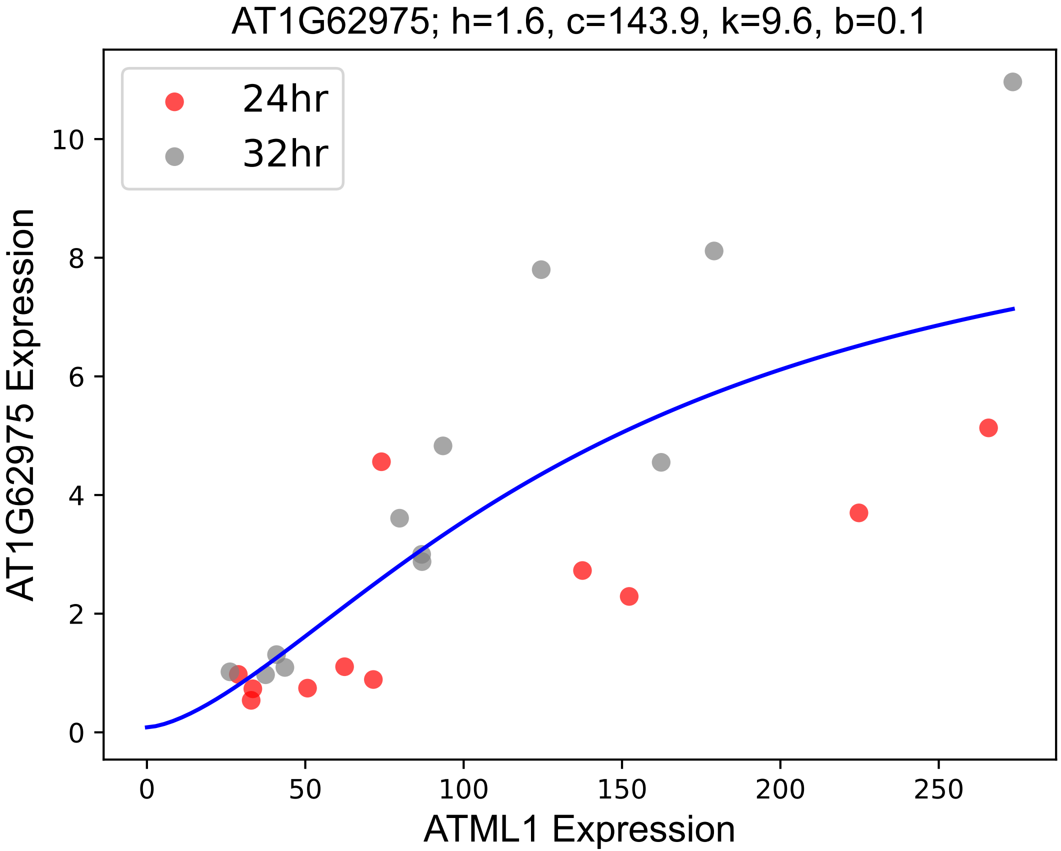

### AT1G64710.pdf

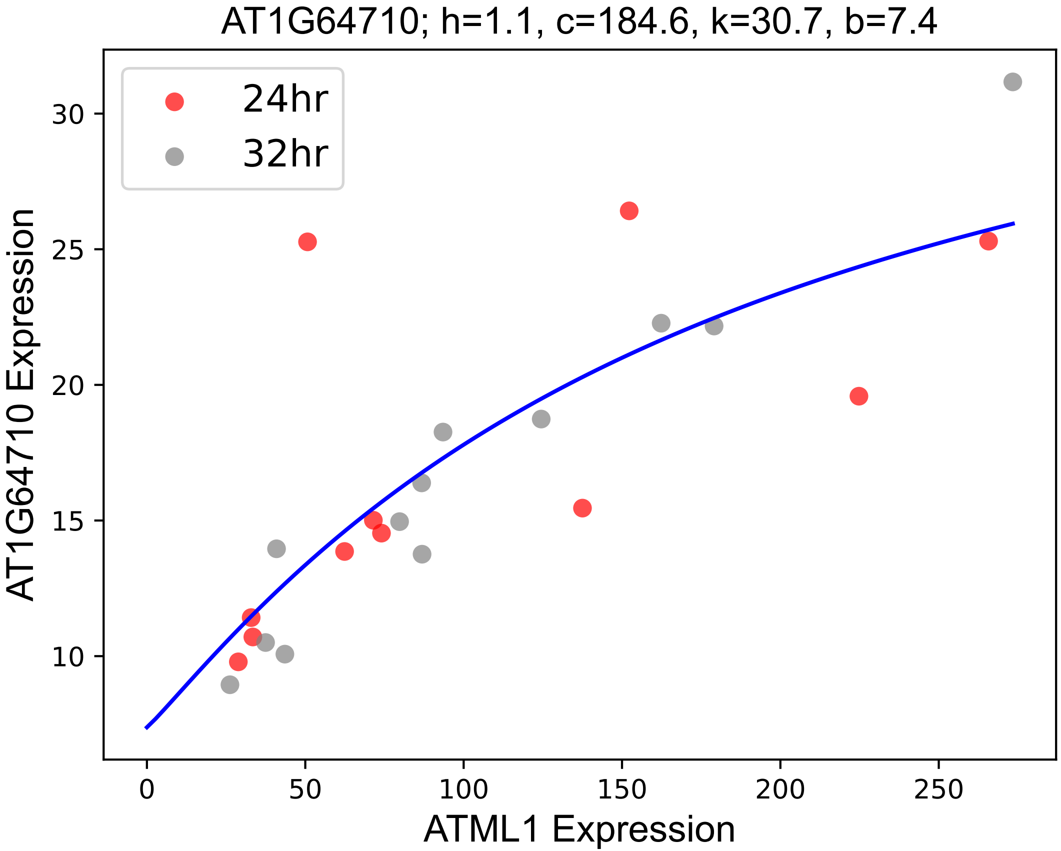

### AT1G68470.pdf

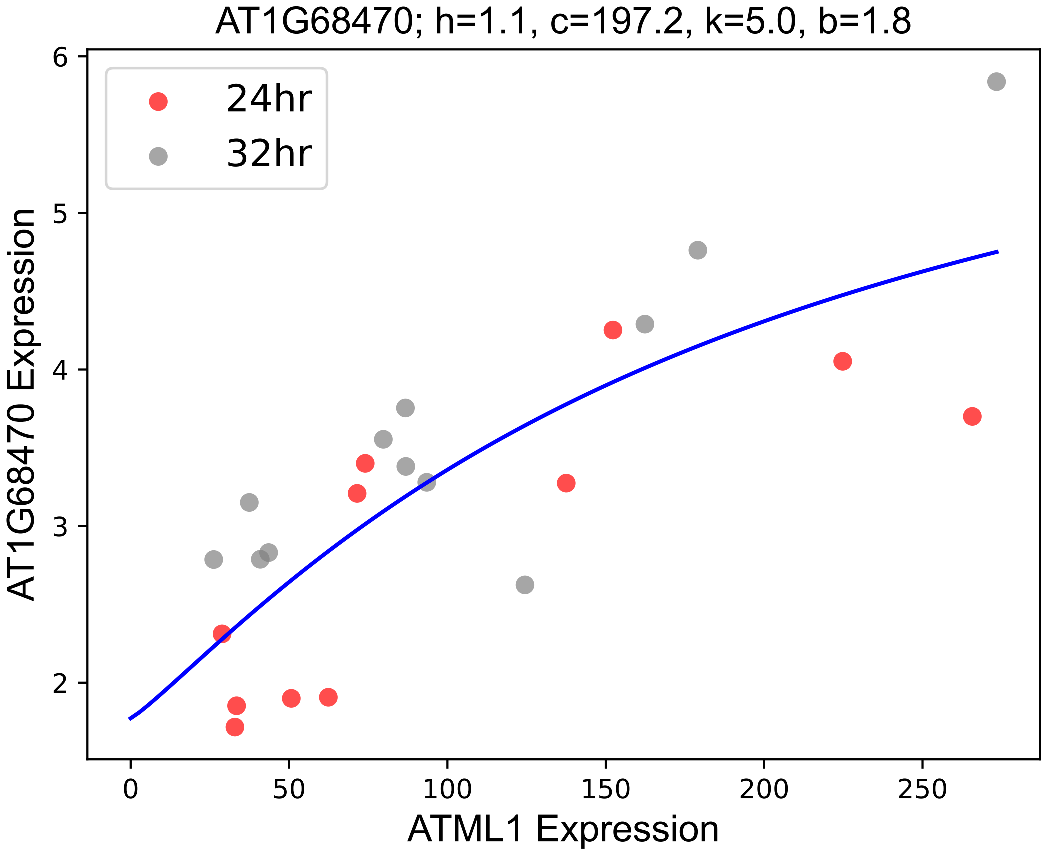

### AT1G78210.pdf

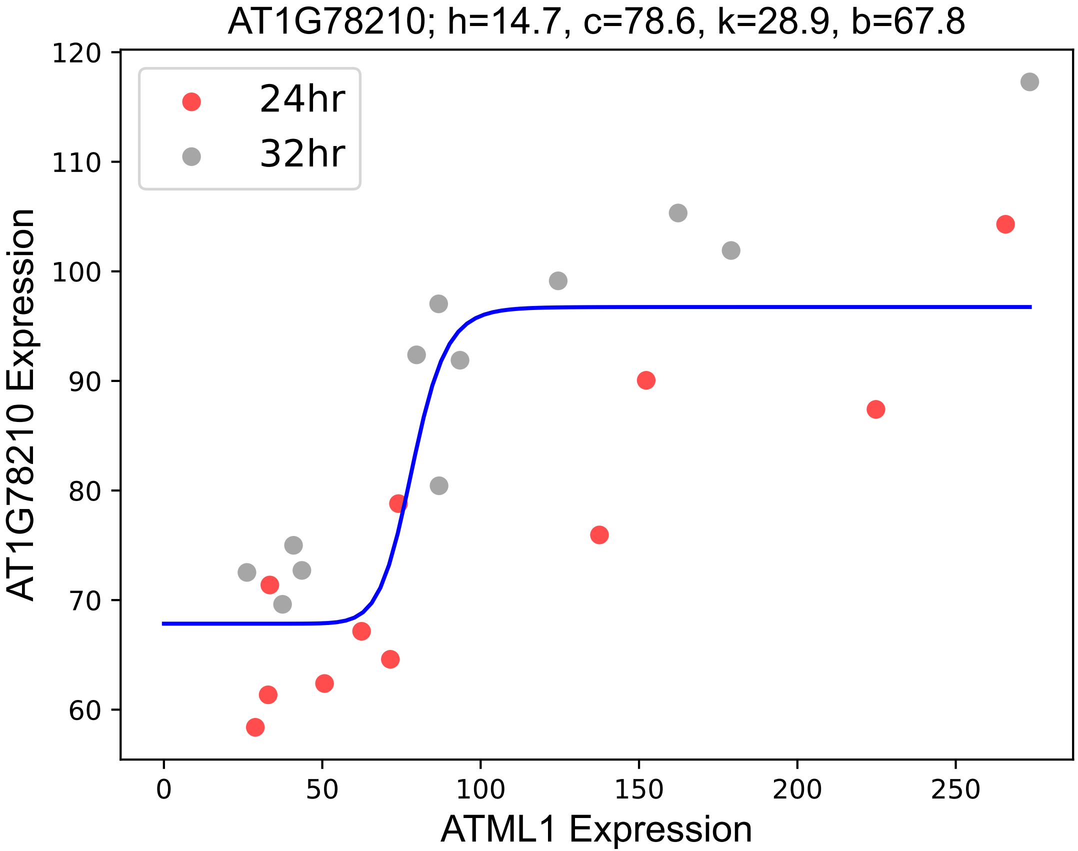

### AT2G04100.pdf

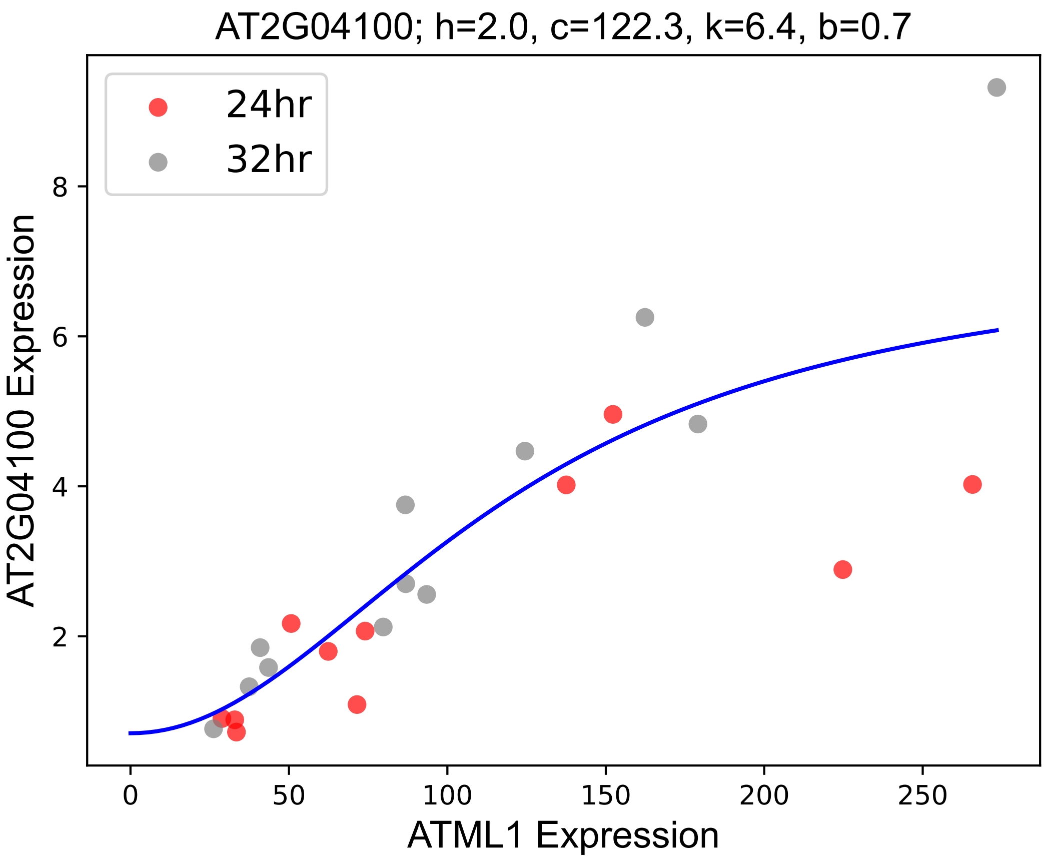

### AT2G05510.pdf

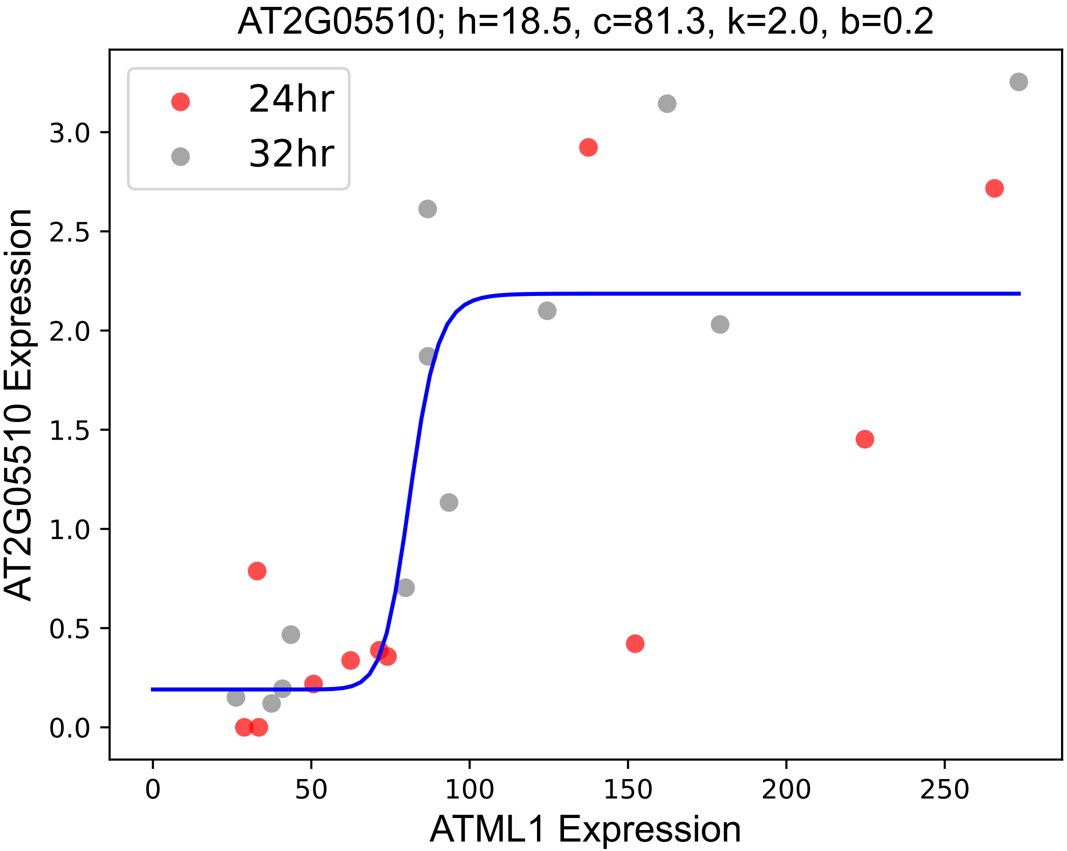

### AT2G20950.pdf

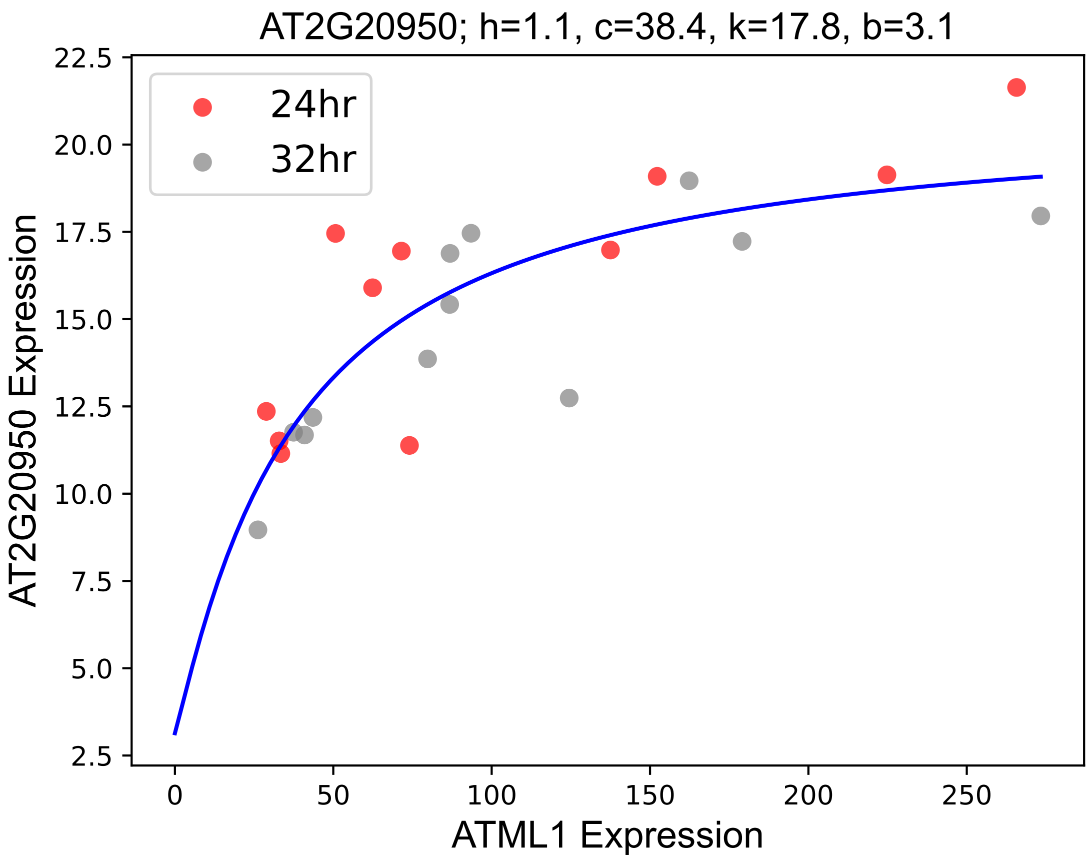

### AT2G23210.pdf

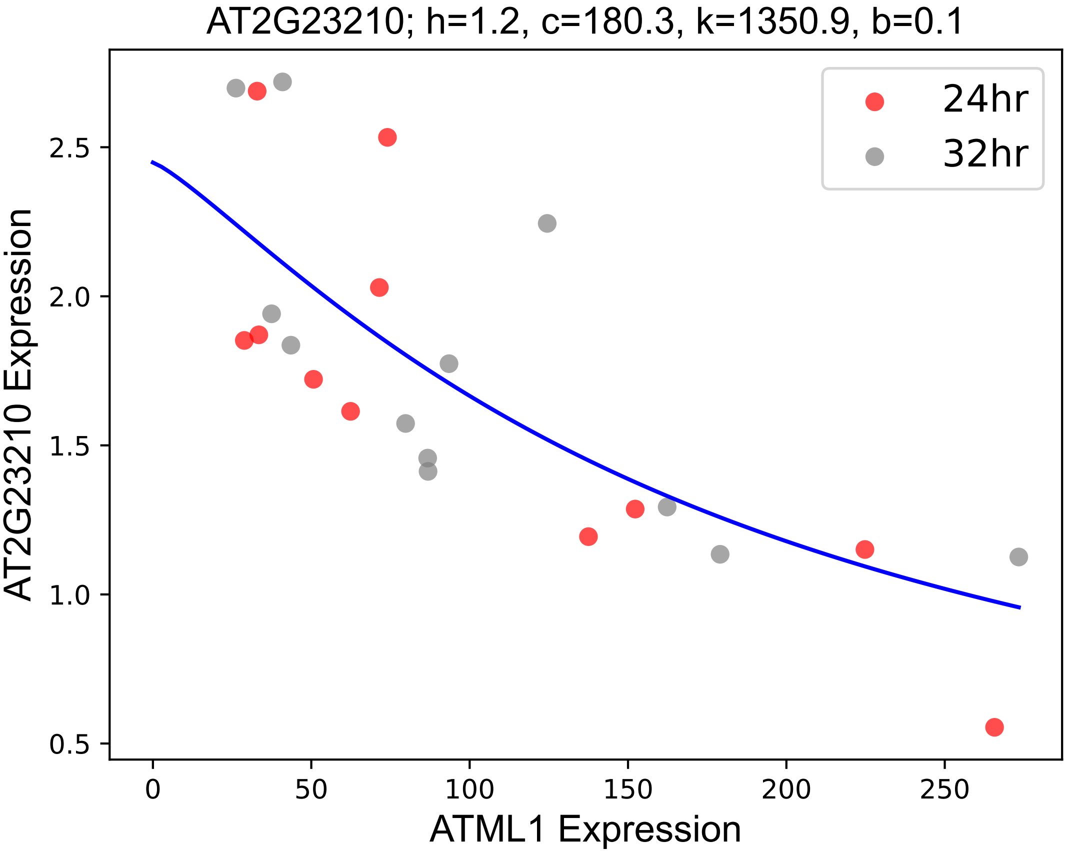

### AT2G23450.pdf

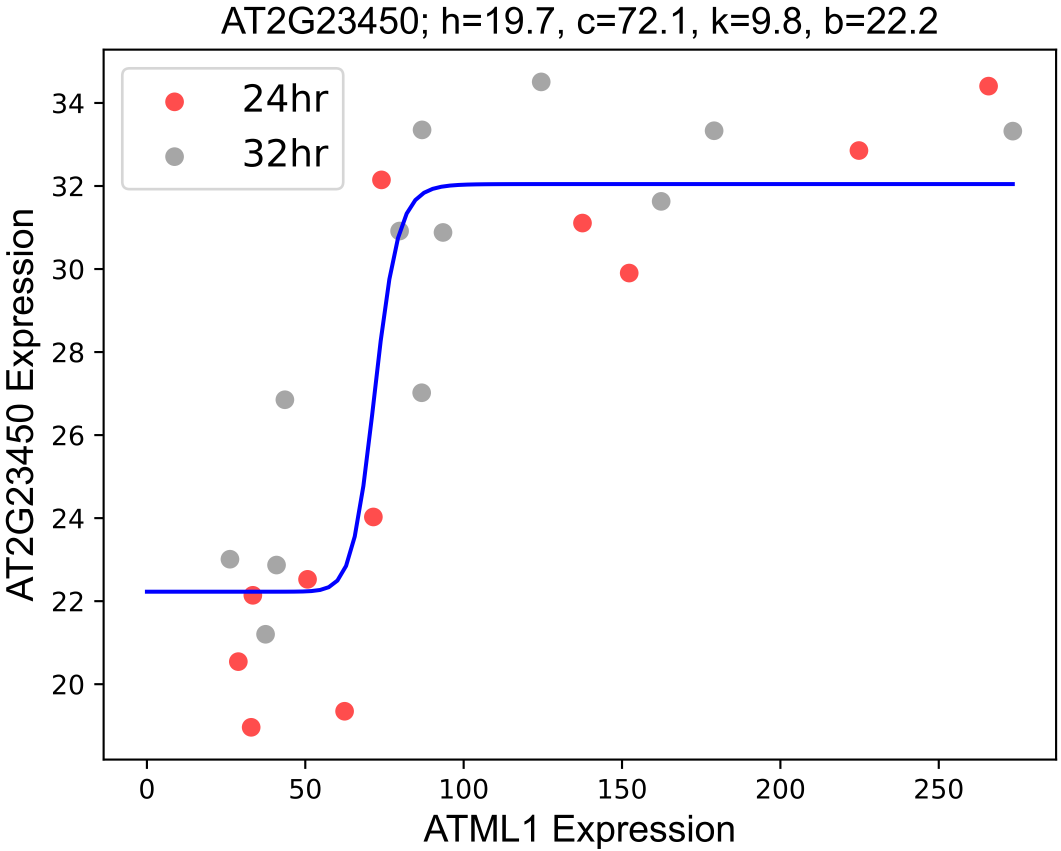

### AT2G32650.pdf

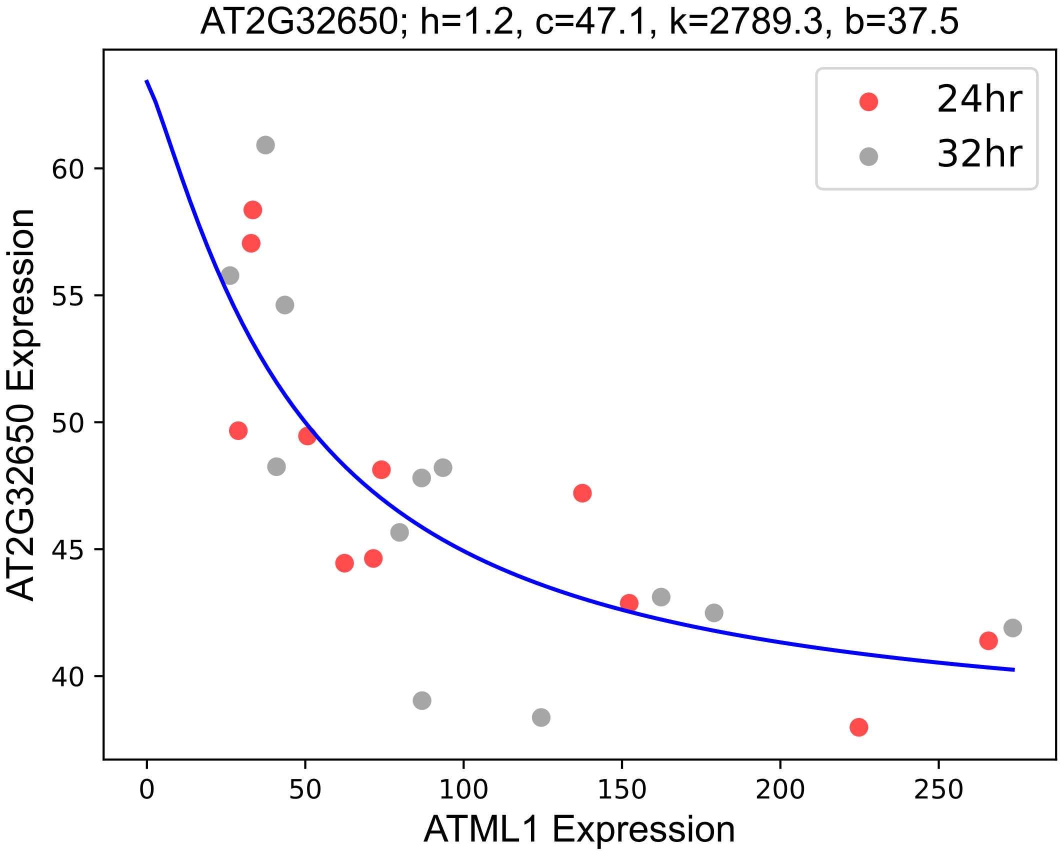

### AT2G43535.pdf

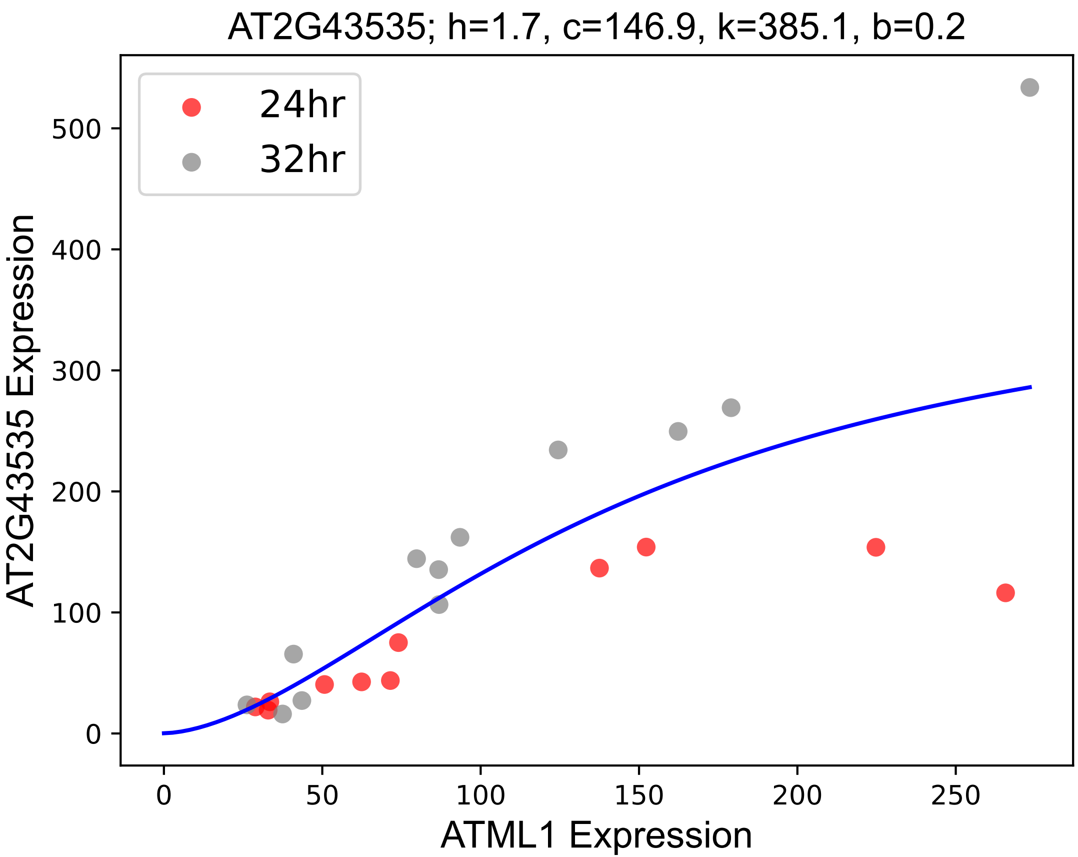

### AT3G06210.pdf

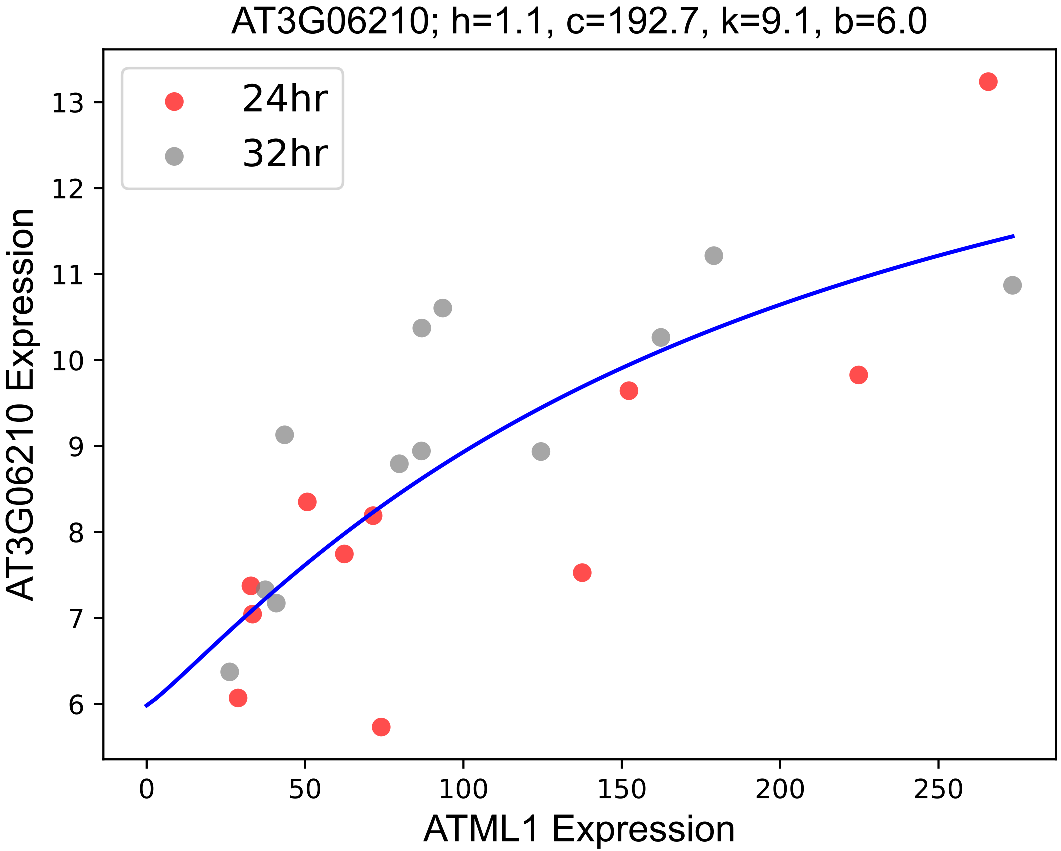

### AT3G12880.pdf

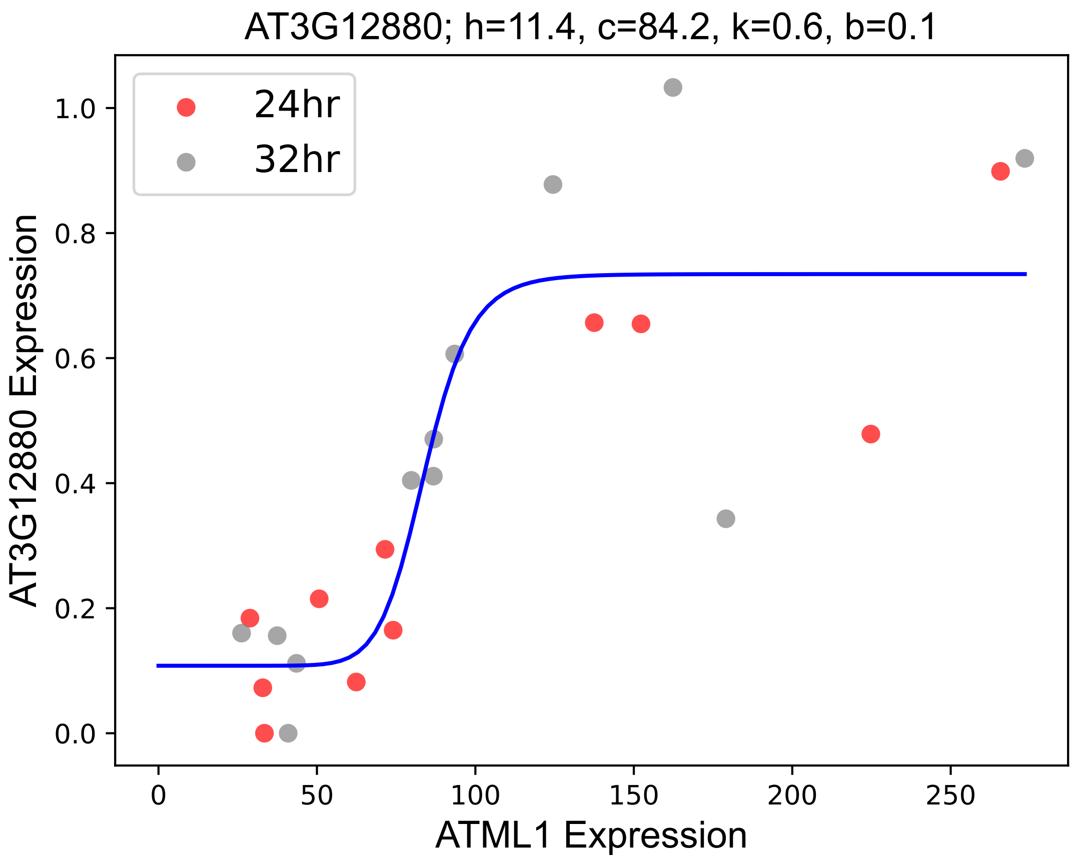

### AT3G18350.pdf

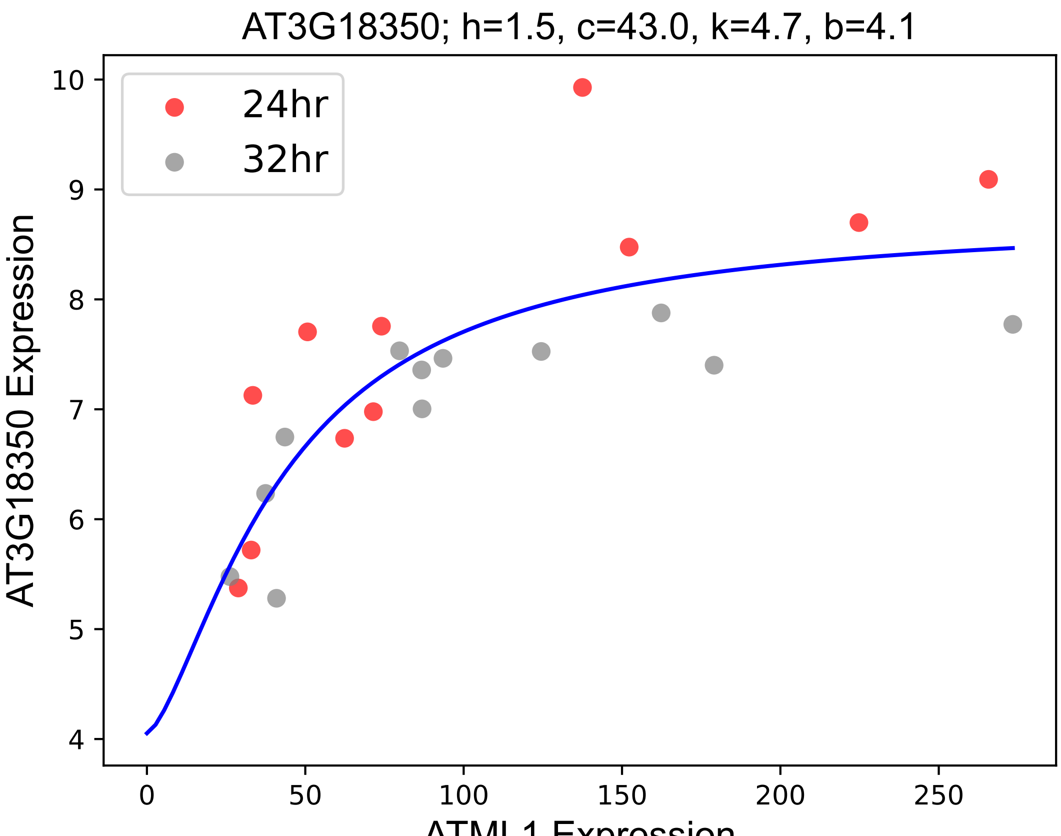

### AT3G19660.pdf

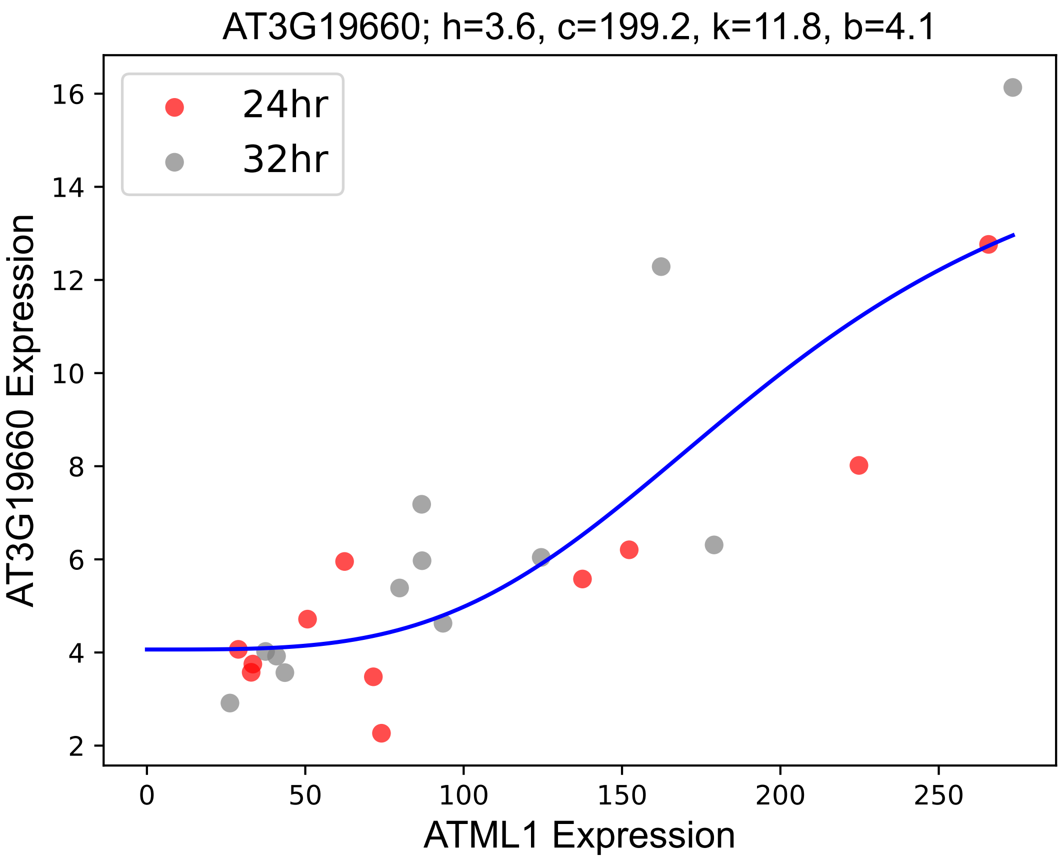

### AT3G25700.pdf

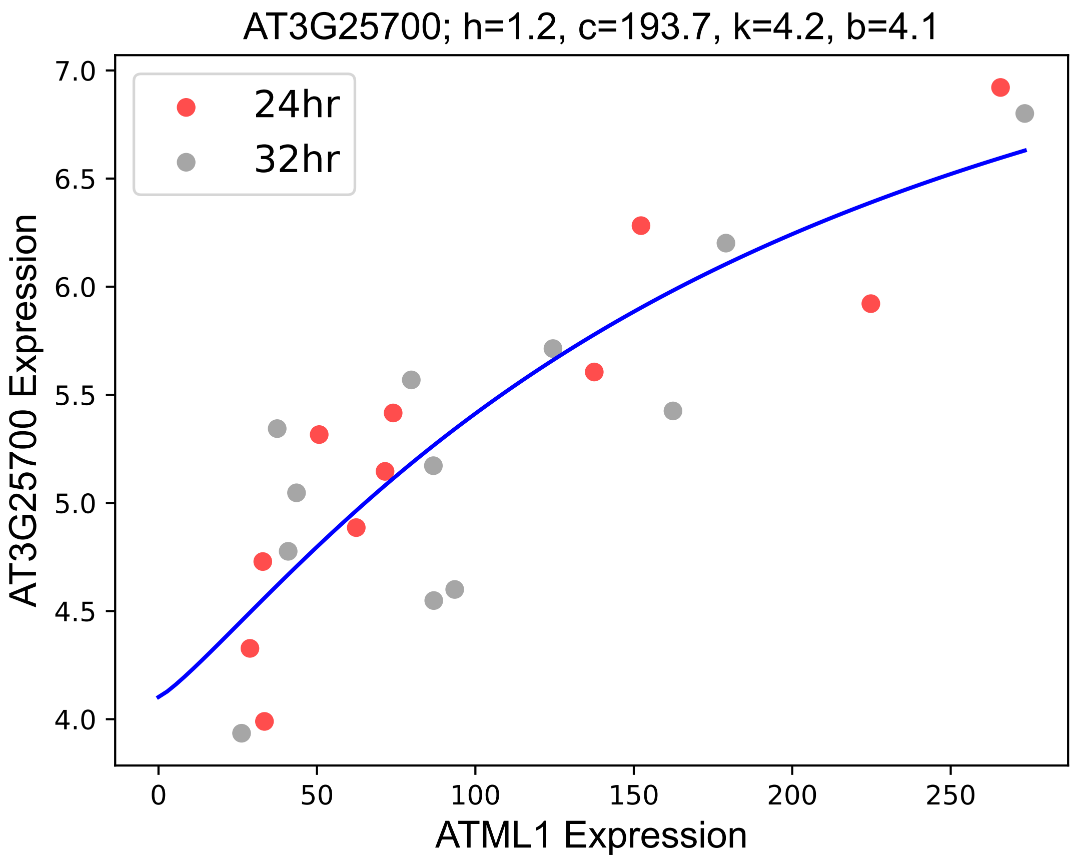

### AT3G26450.pdf

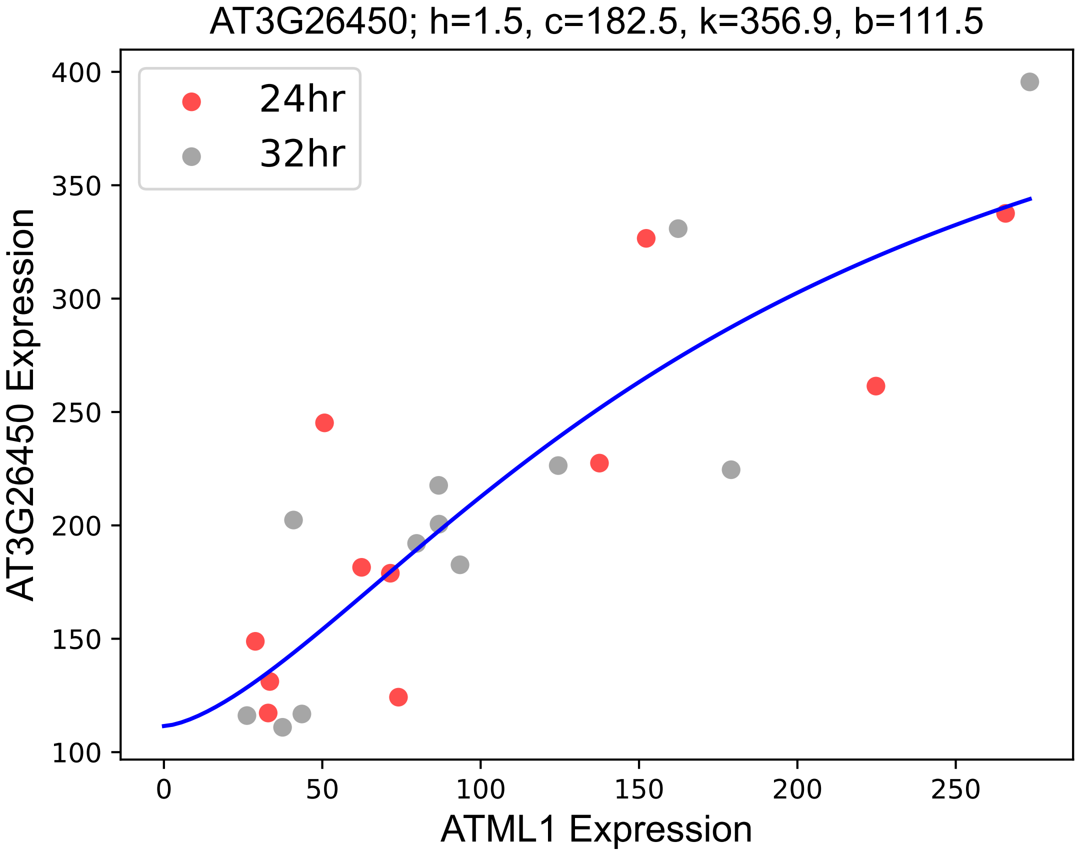

### AT3G26470.pdf

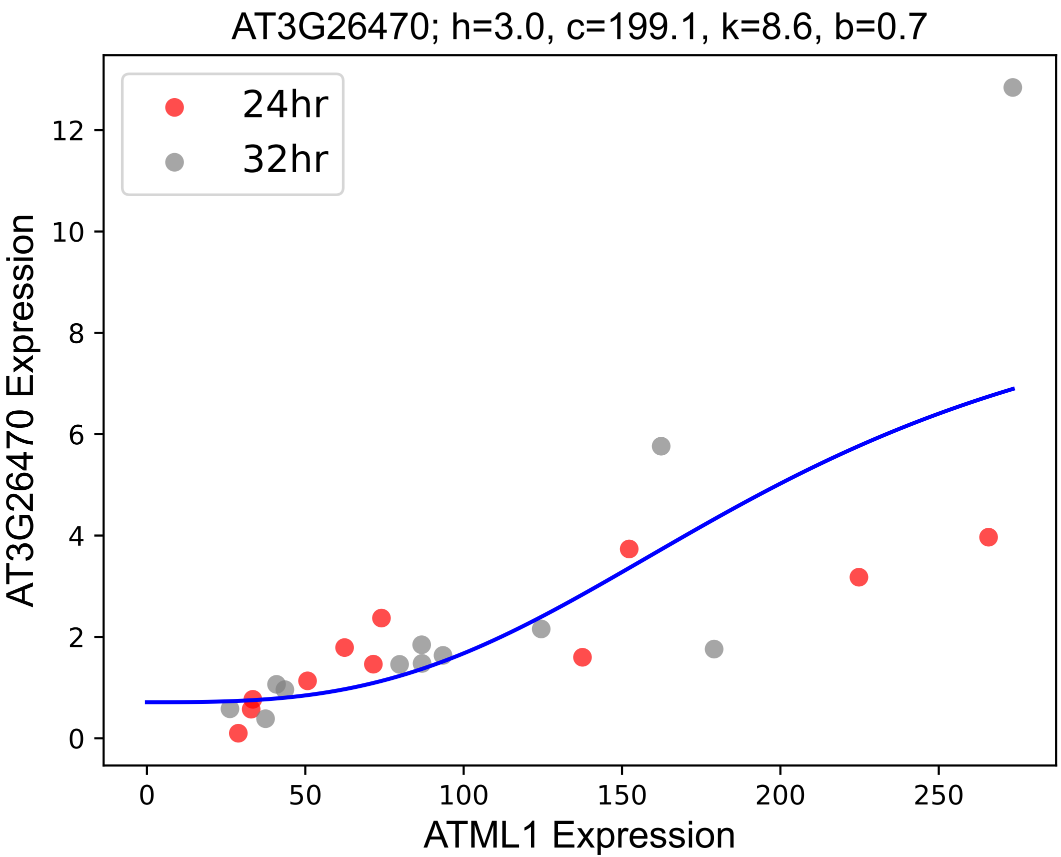

### AT3G57400.pdf

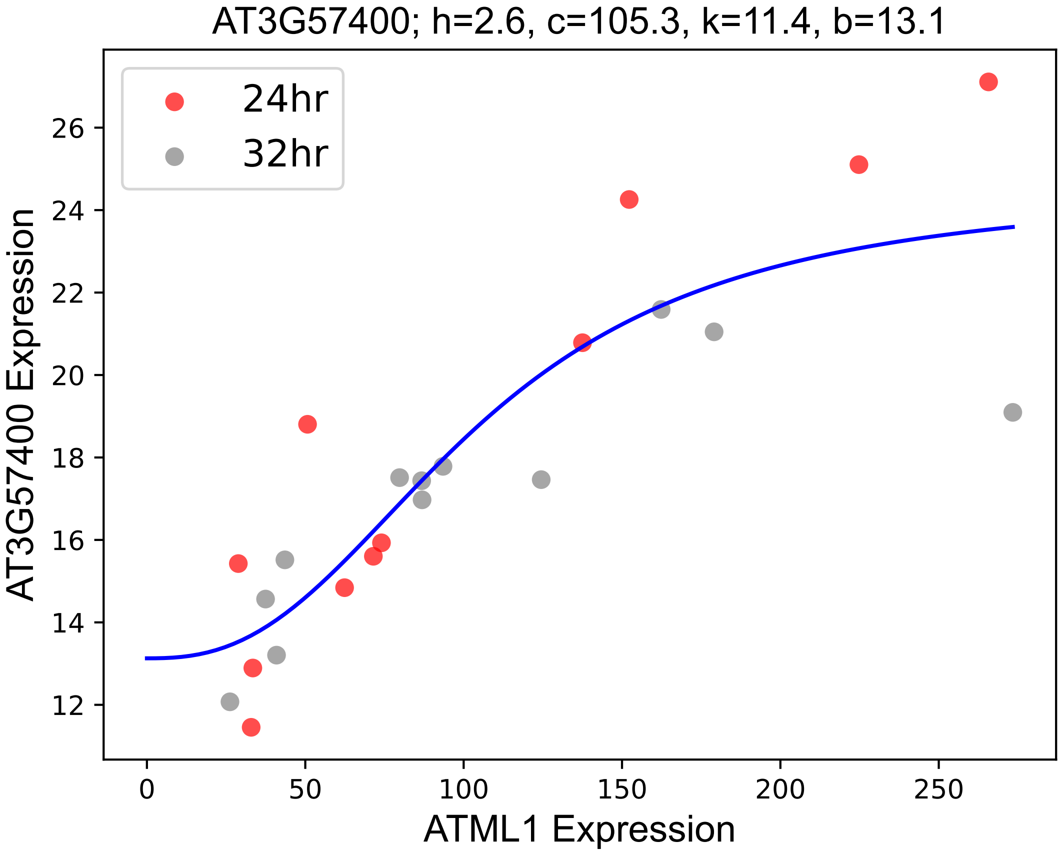

### AT3G61490.pdf

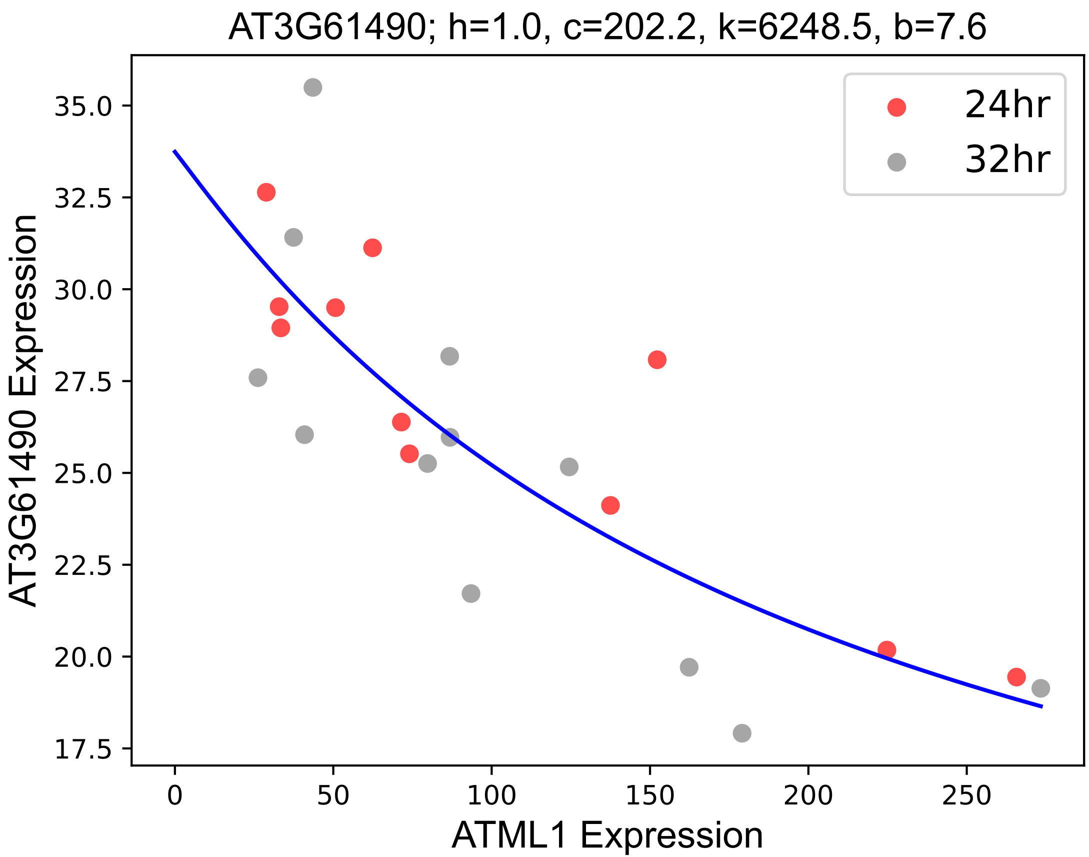

### AT4G01110.pdf

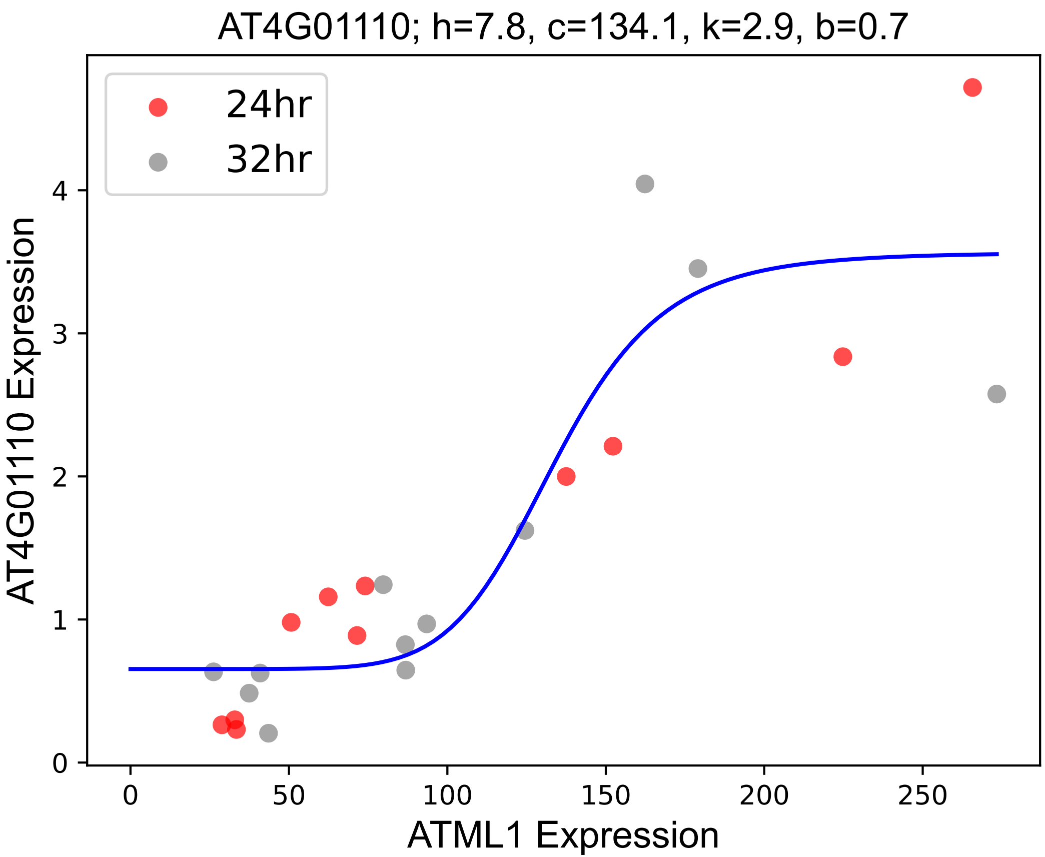

### AT4G01460.pdf

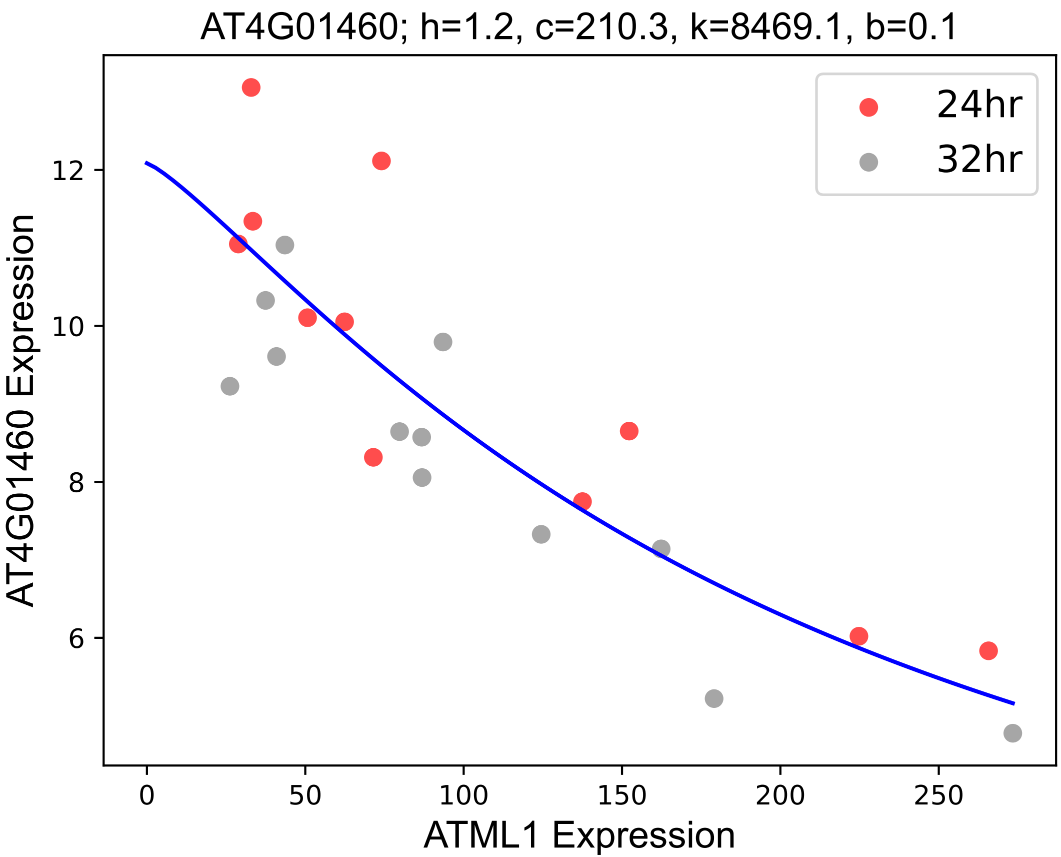

### AT4G18220.pdf

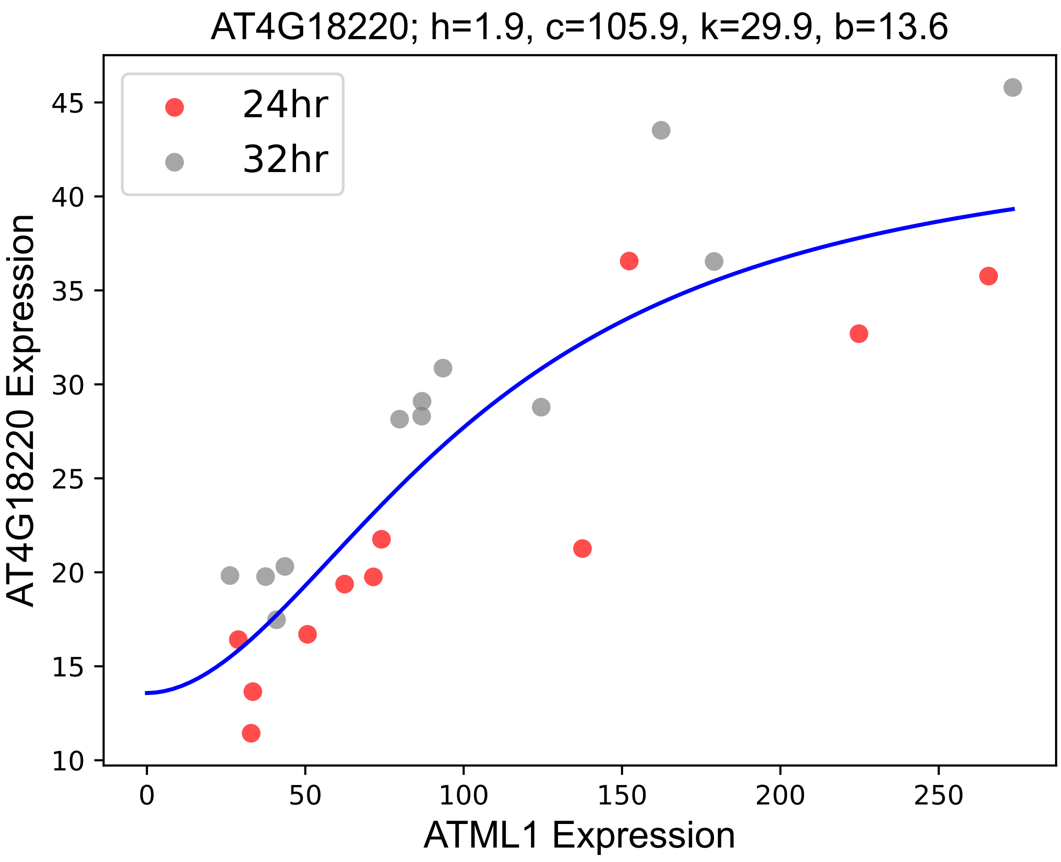

### AT4G23670.pdf

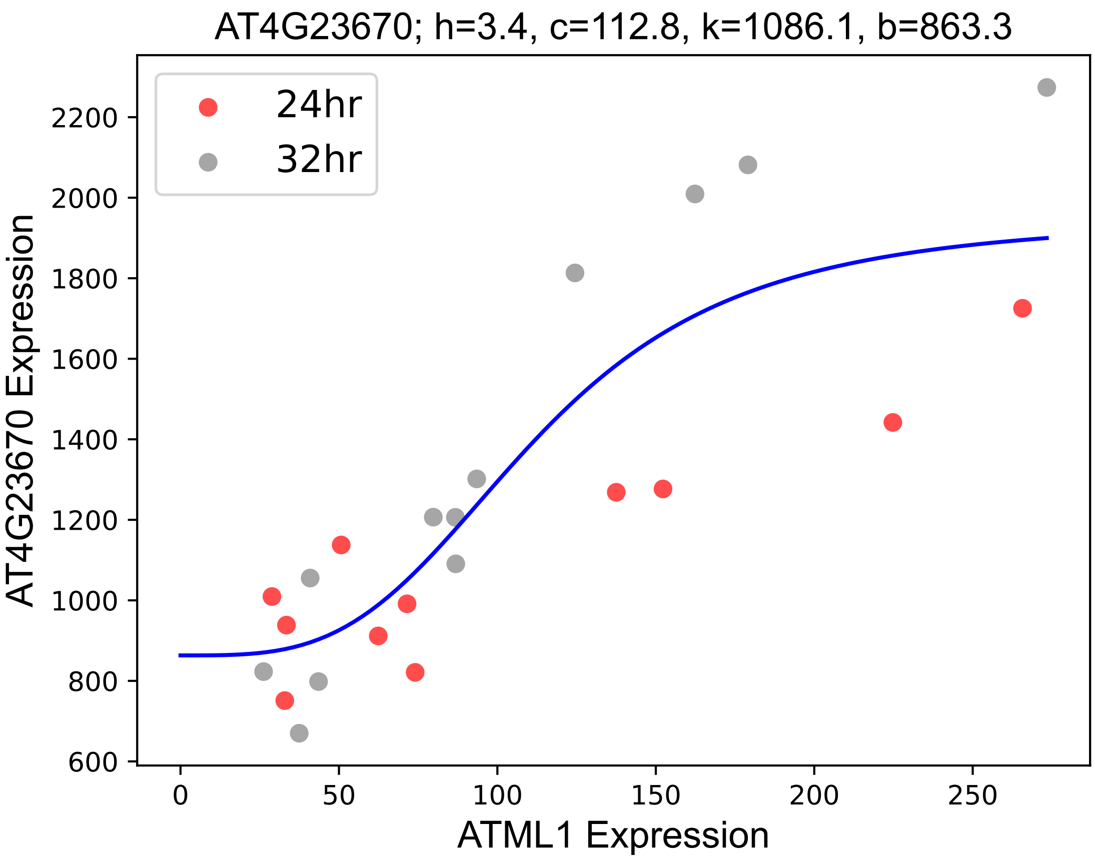

### AT4G27700.pdf

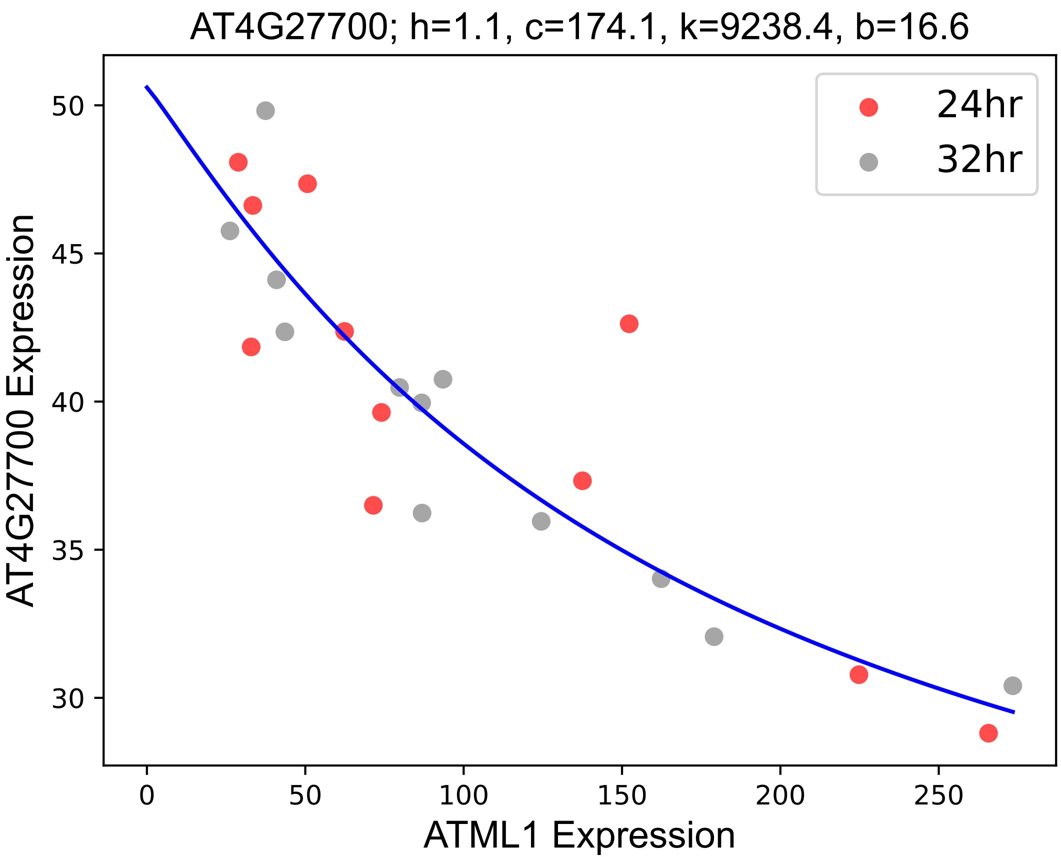

### AT4G30470.pdf

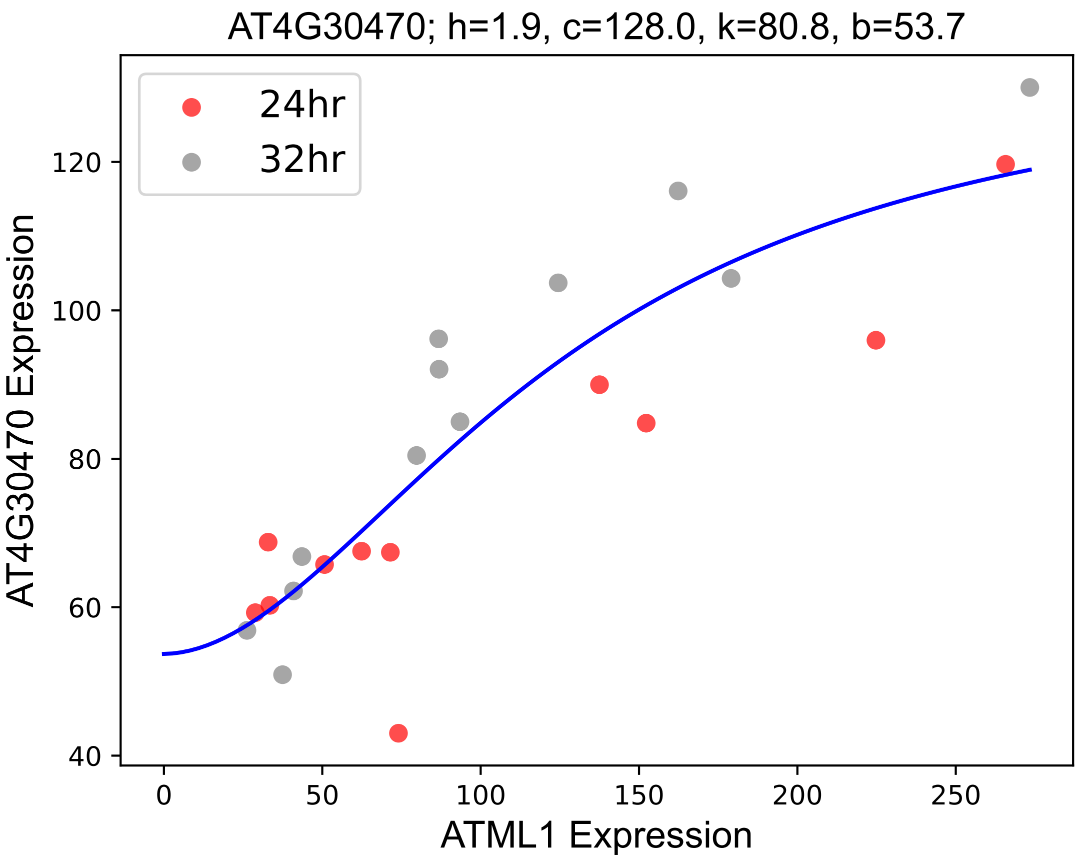

### AT4G32000.pdf

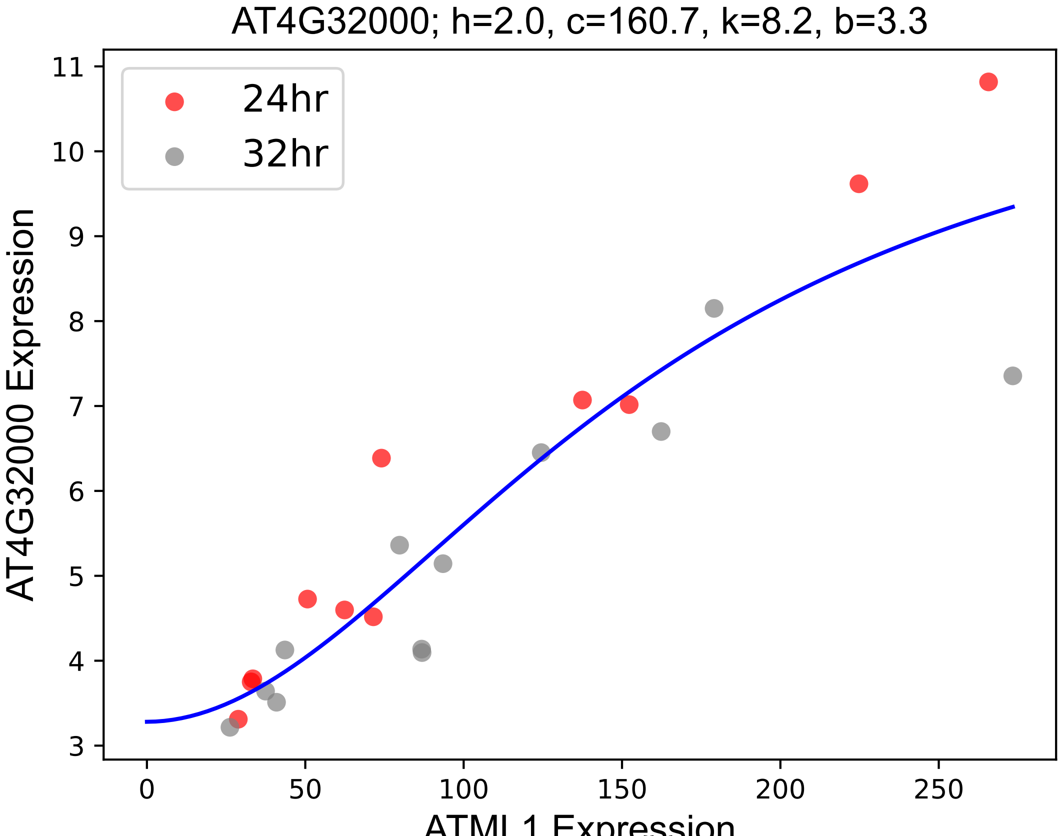

### AT4G34480.pdf

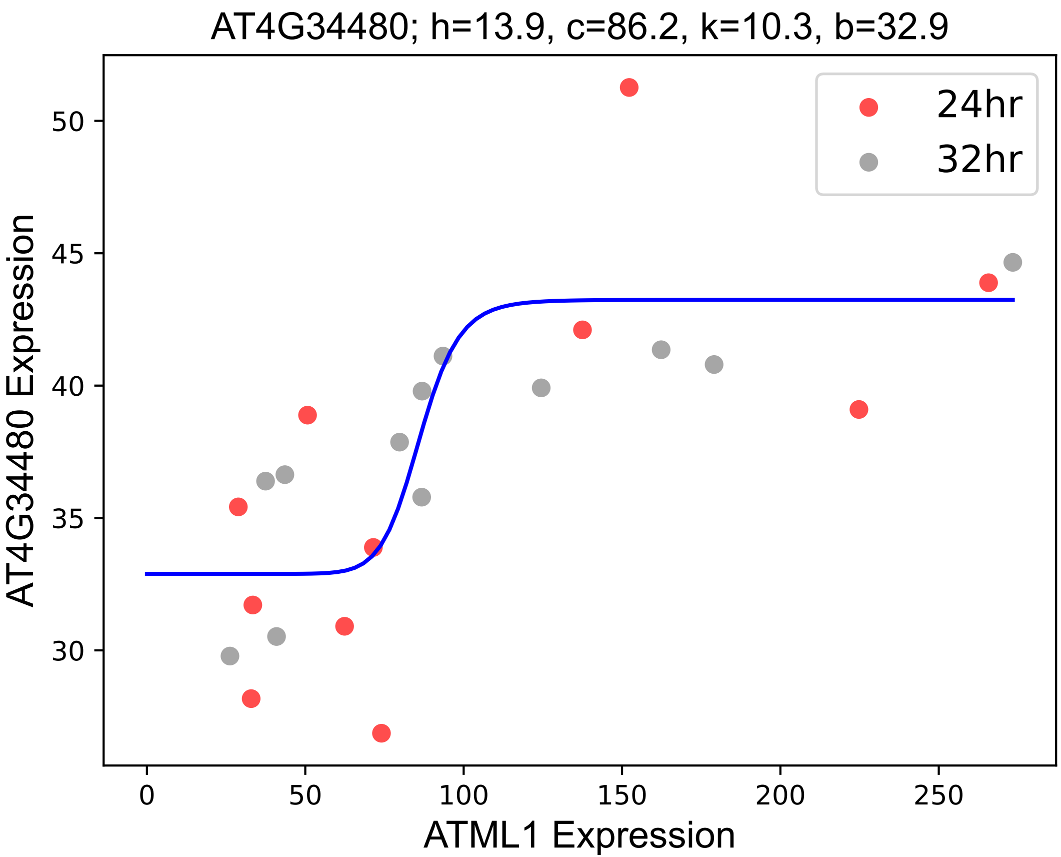

### AT5G02890.pdf

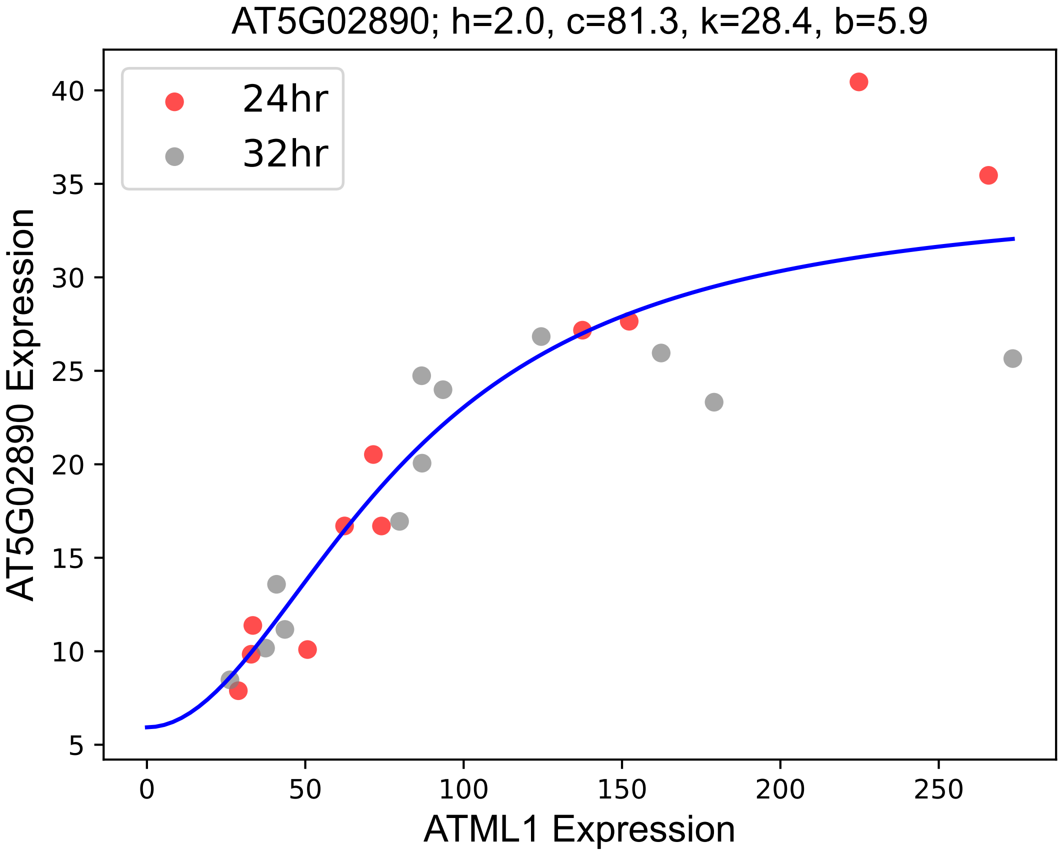

### AT5G03830.pdf

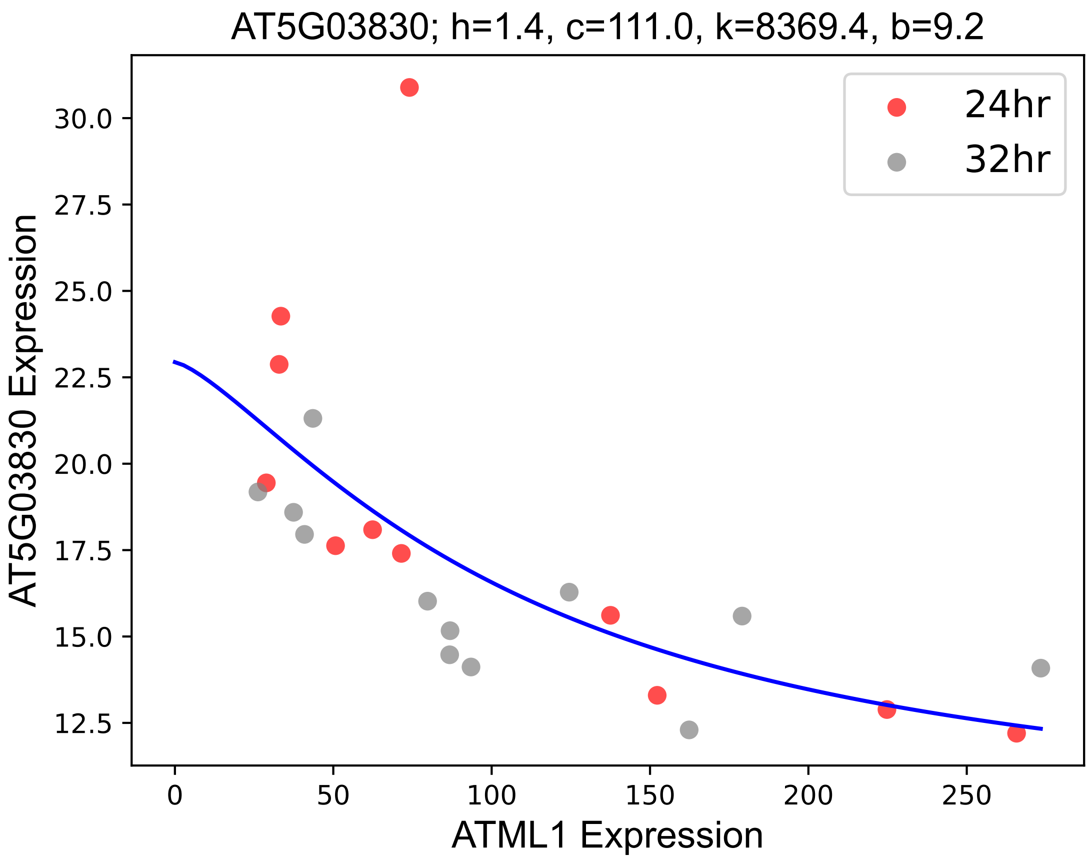

### AT5G09620.pdf

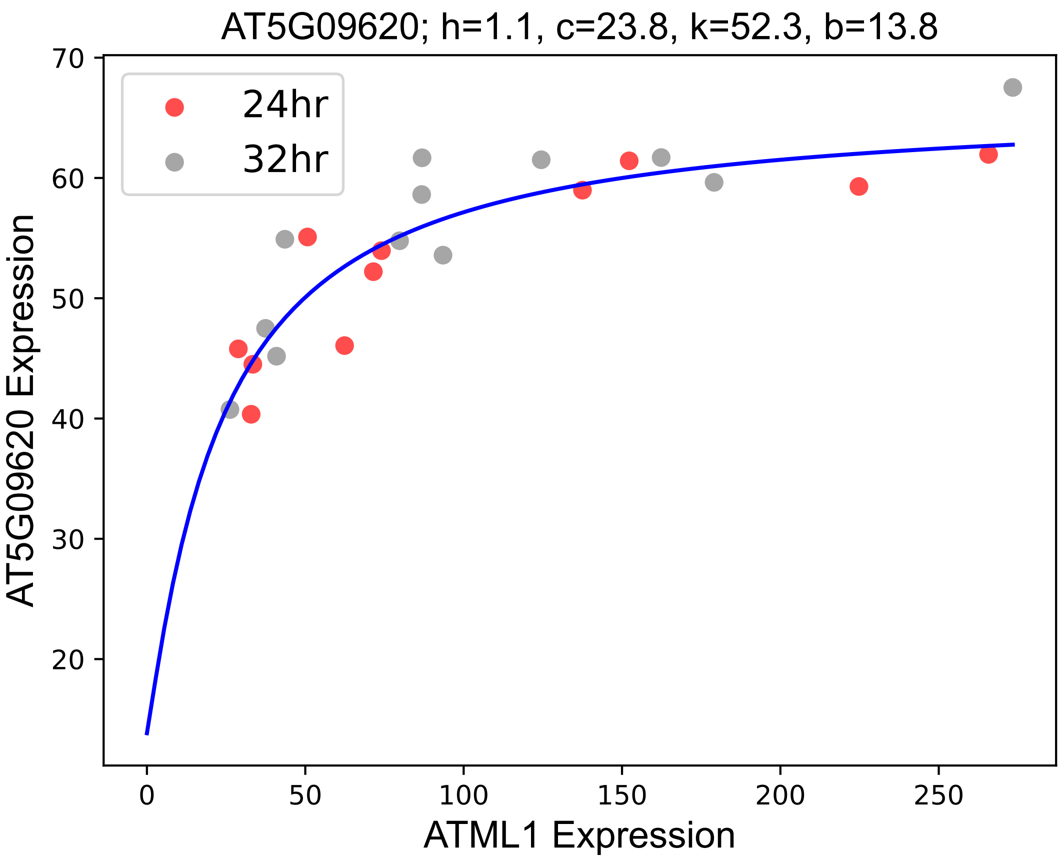

### AT5G17700.pdf

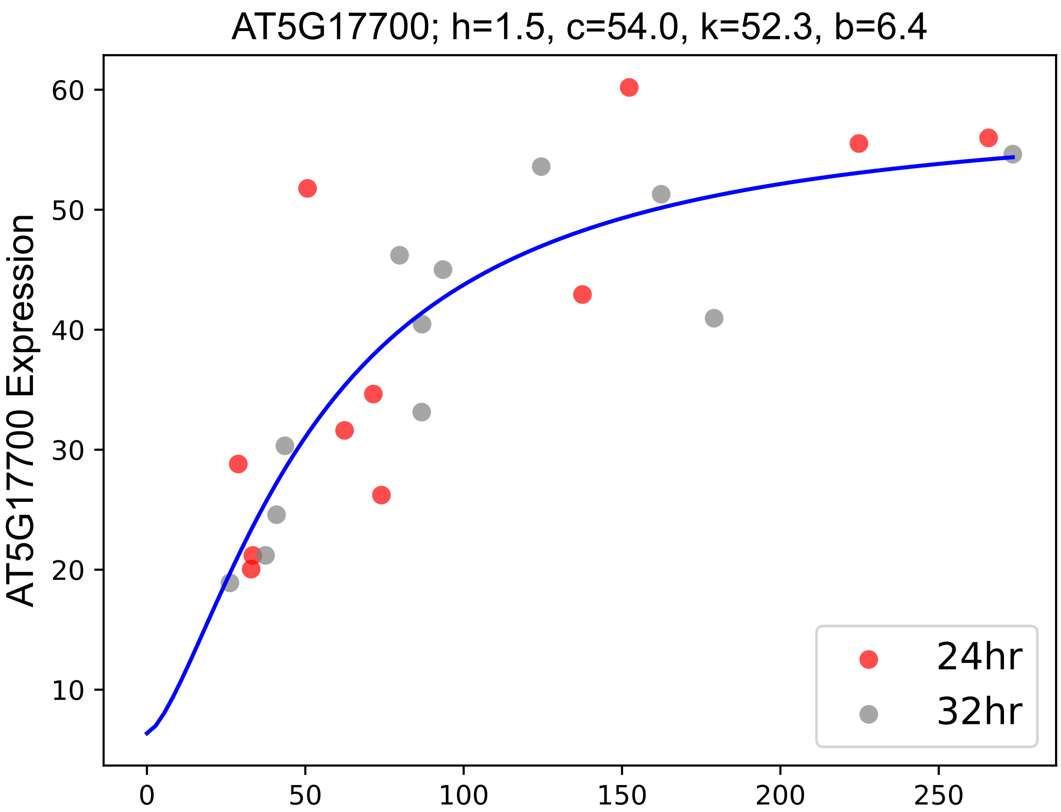

### AT5G18460.pdf

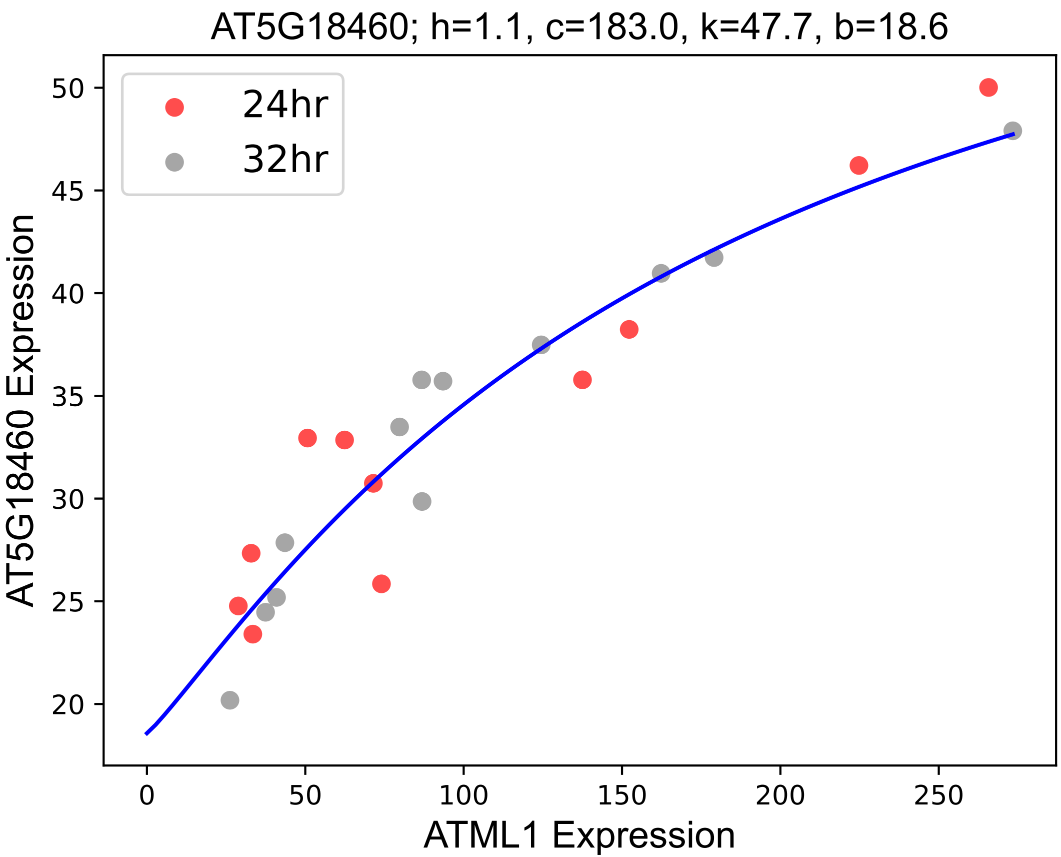

### AT5G23920.pdf

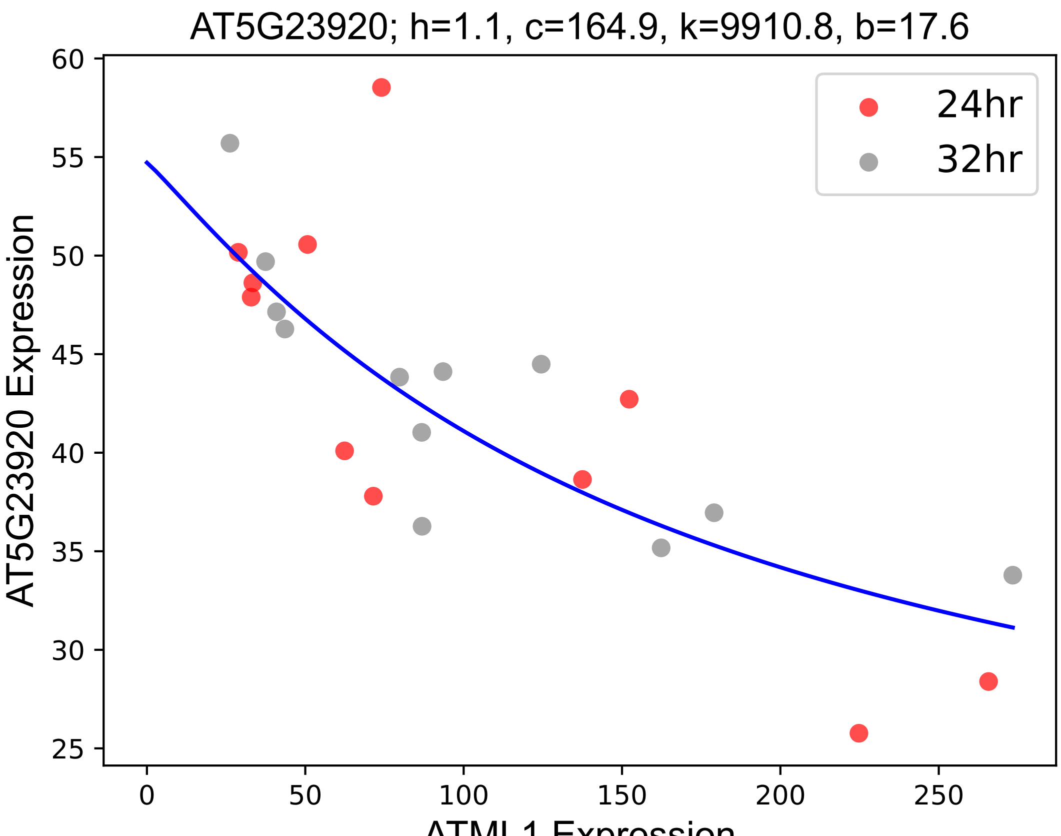

### AT5G27860.pdf

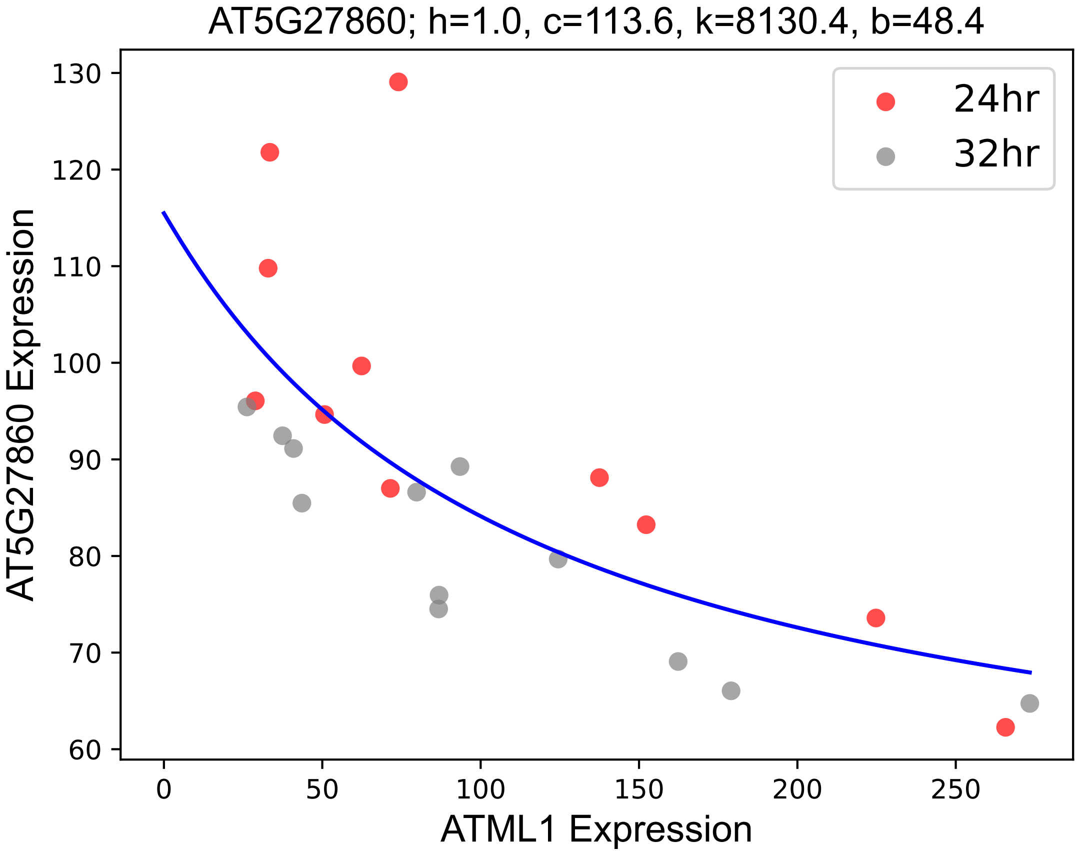

### AT5G38200.pdf

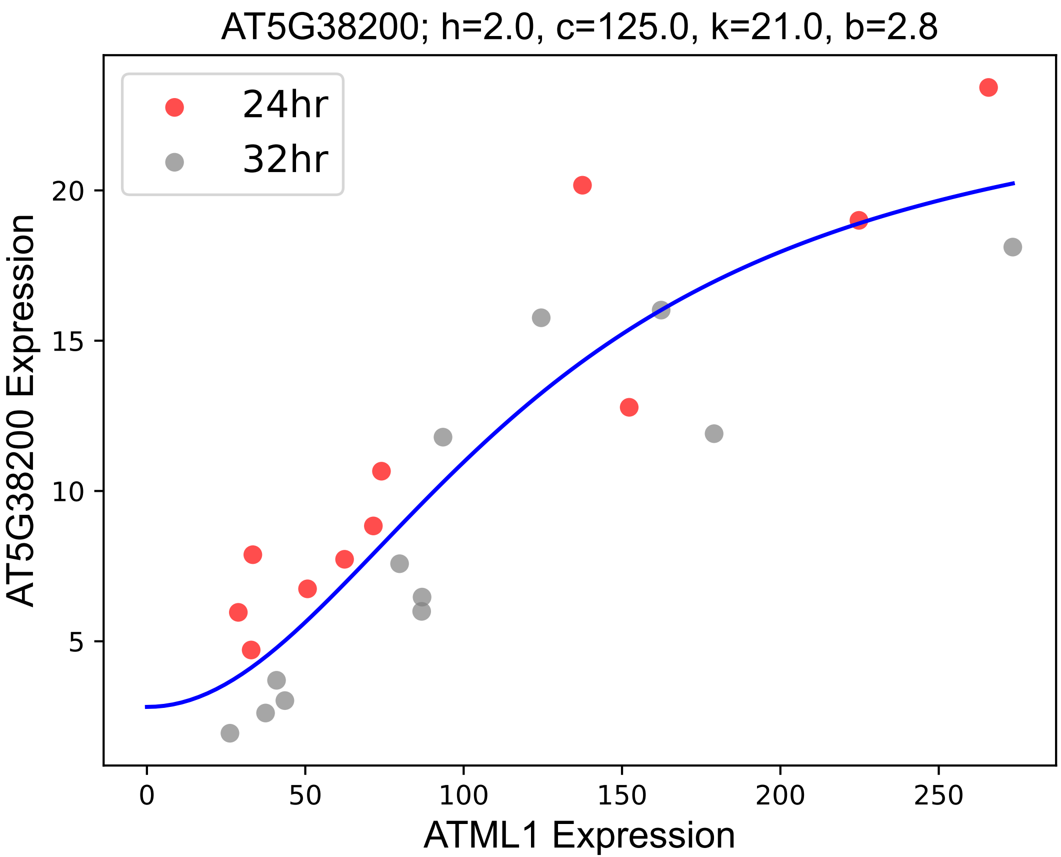

### AT5G51500.pdf

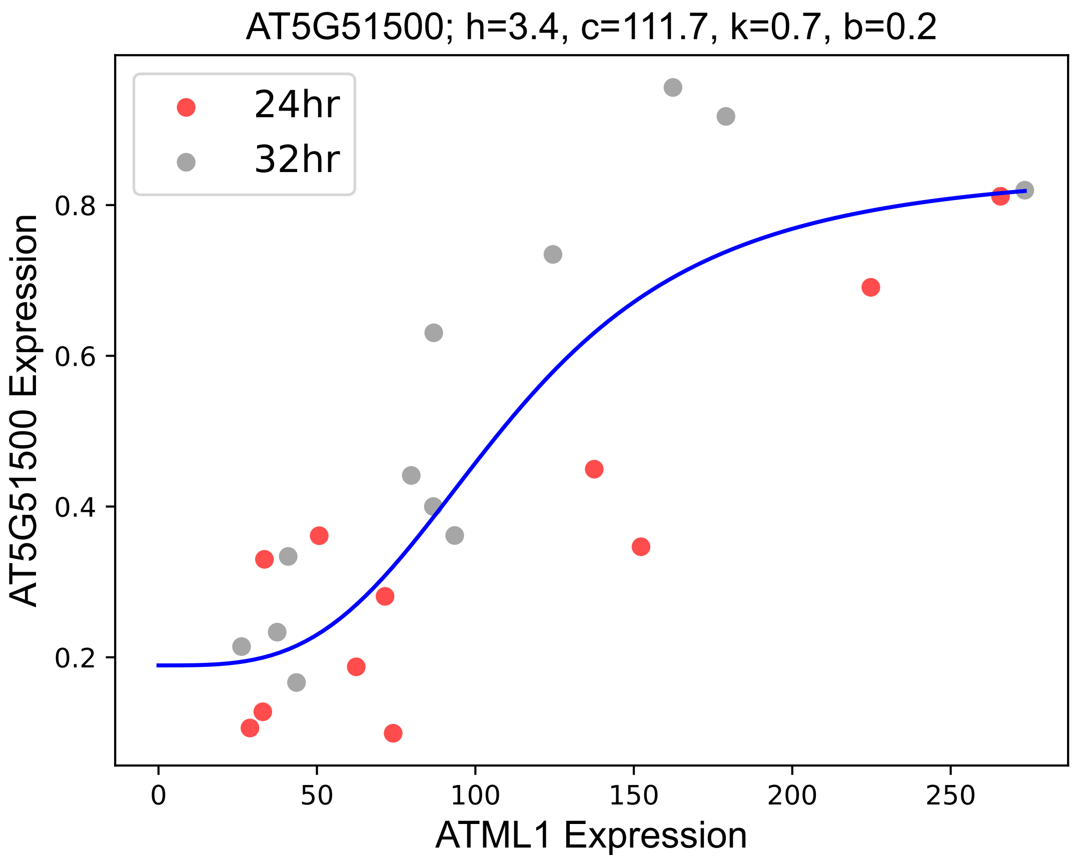

### AT5G51580.pdf

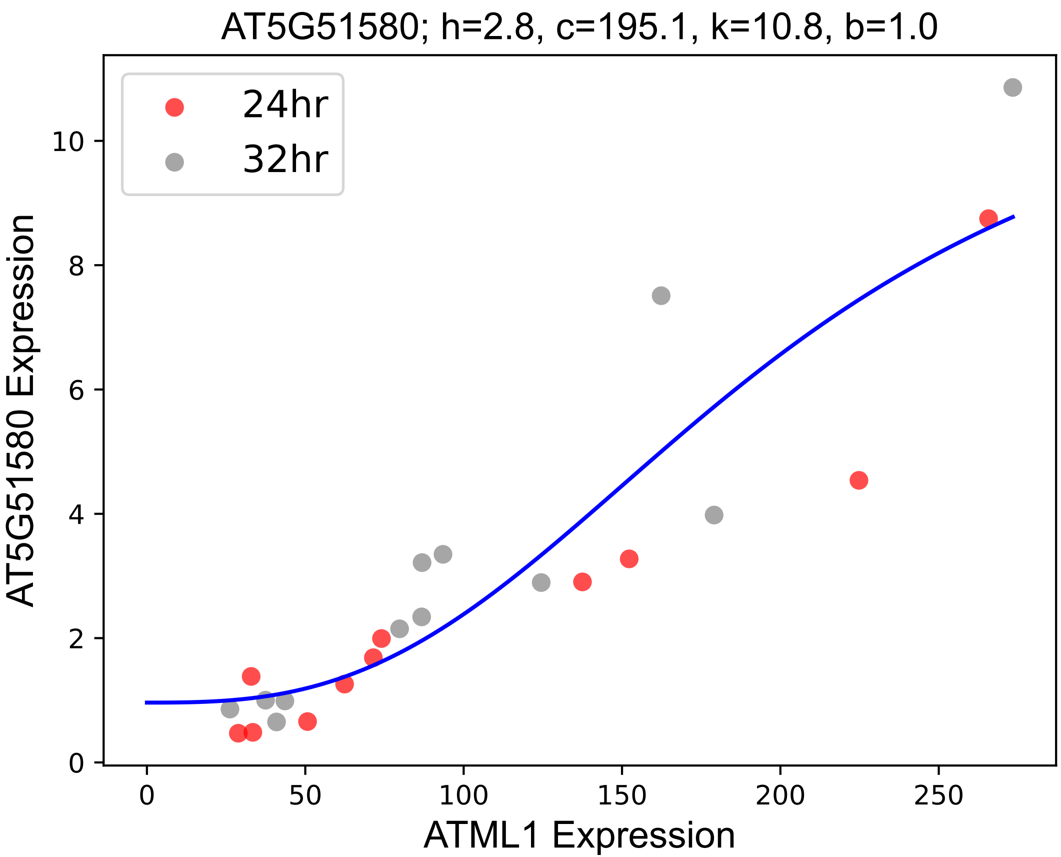

### AT5G63180.pdf

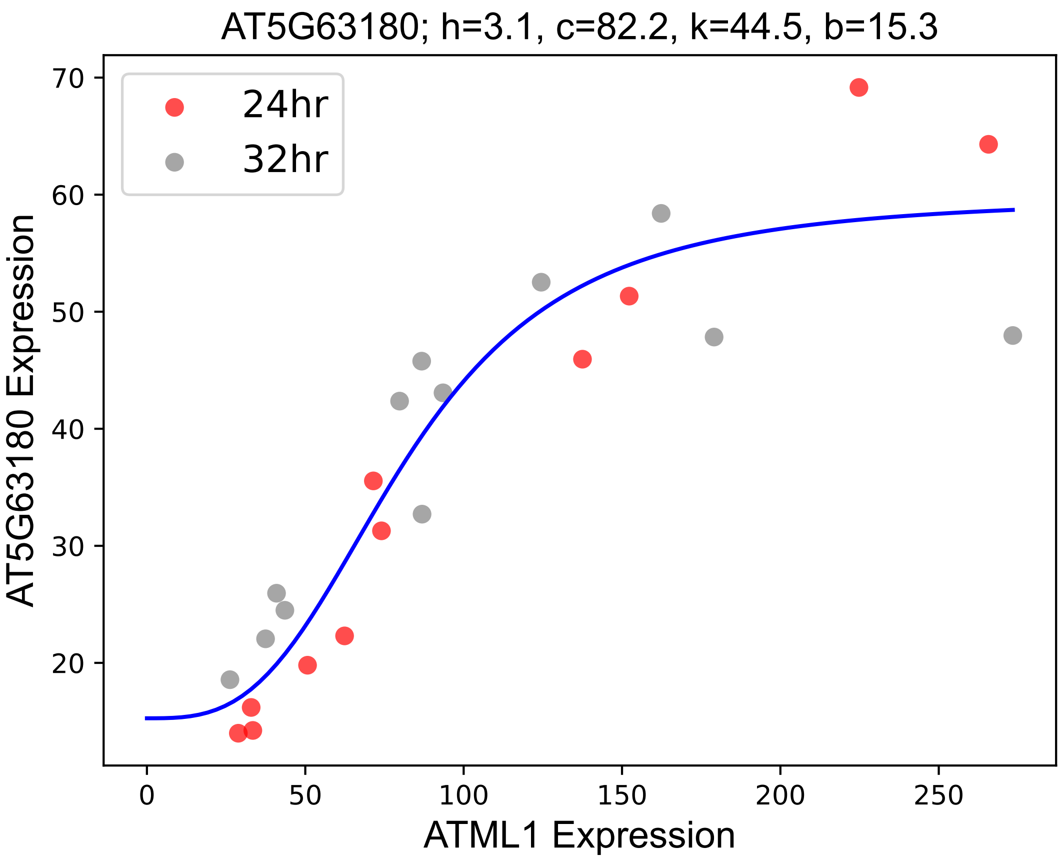

### ATPE.pdf

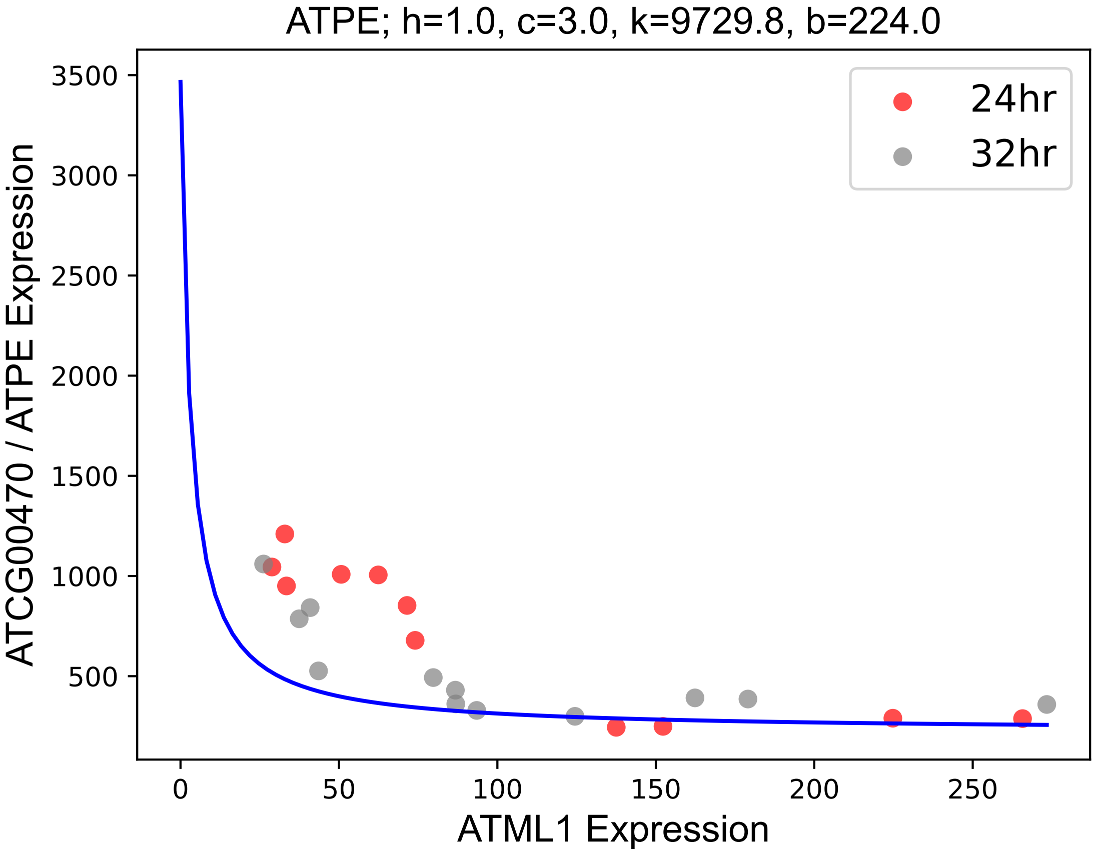

### ATPMEI10.pdf

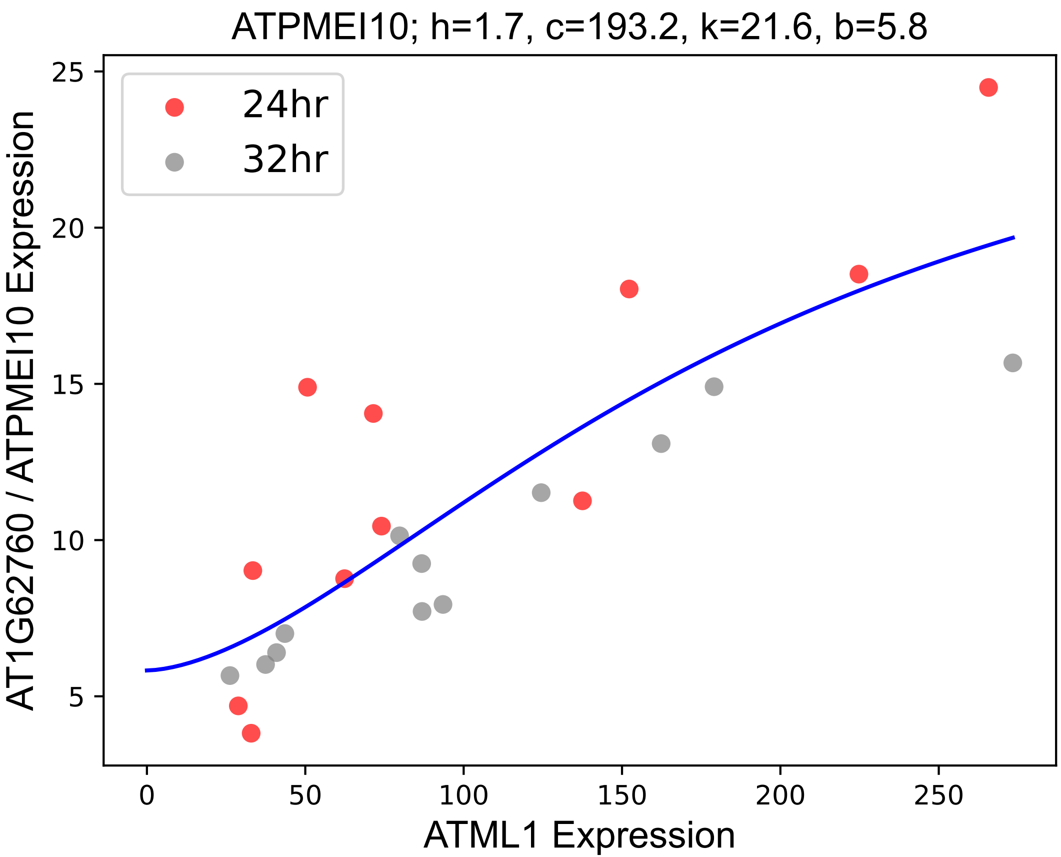

### ATSYTF.pdf

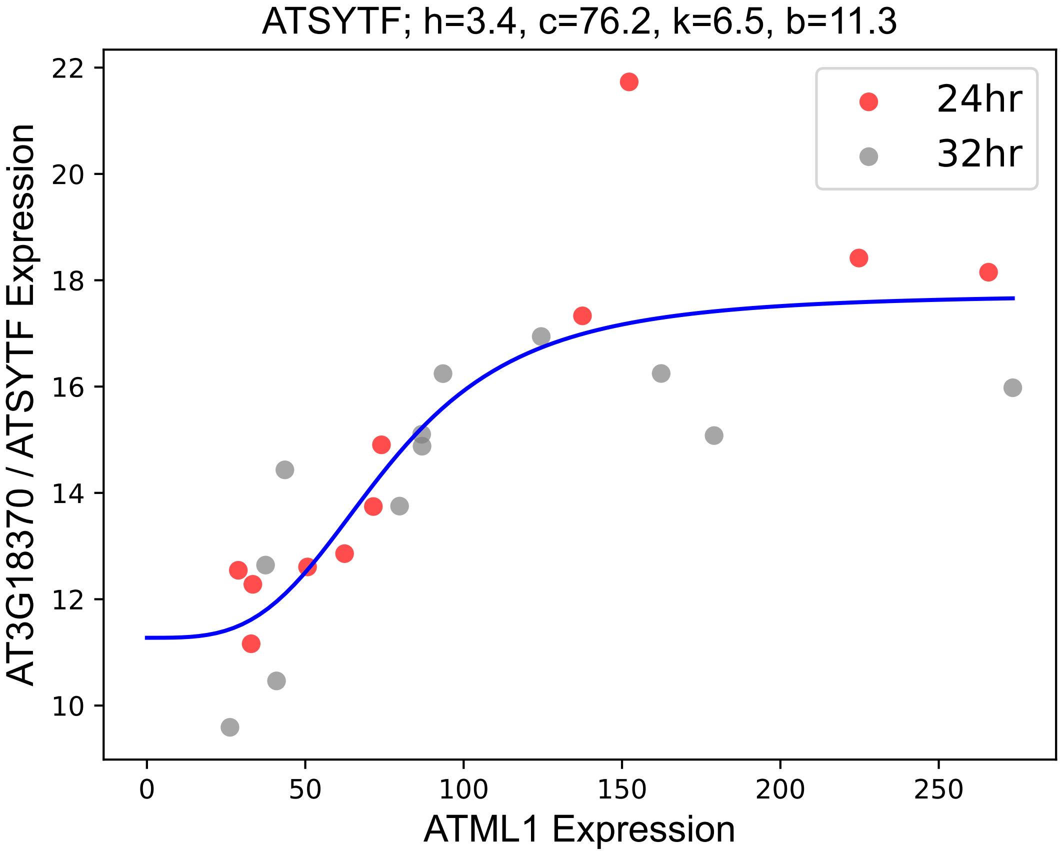

### BAM1.pdf

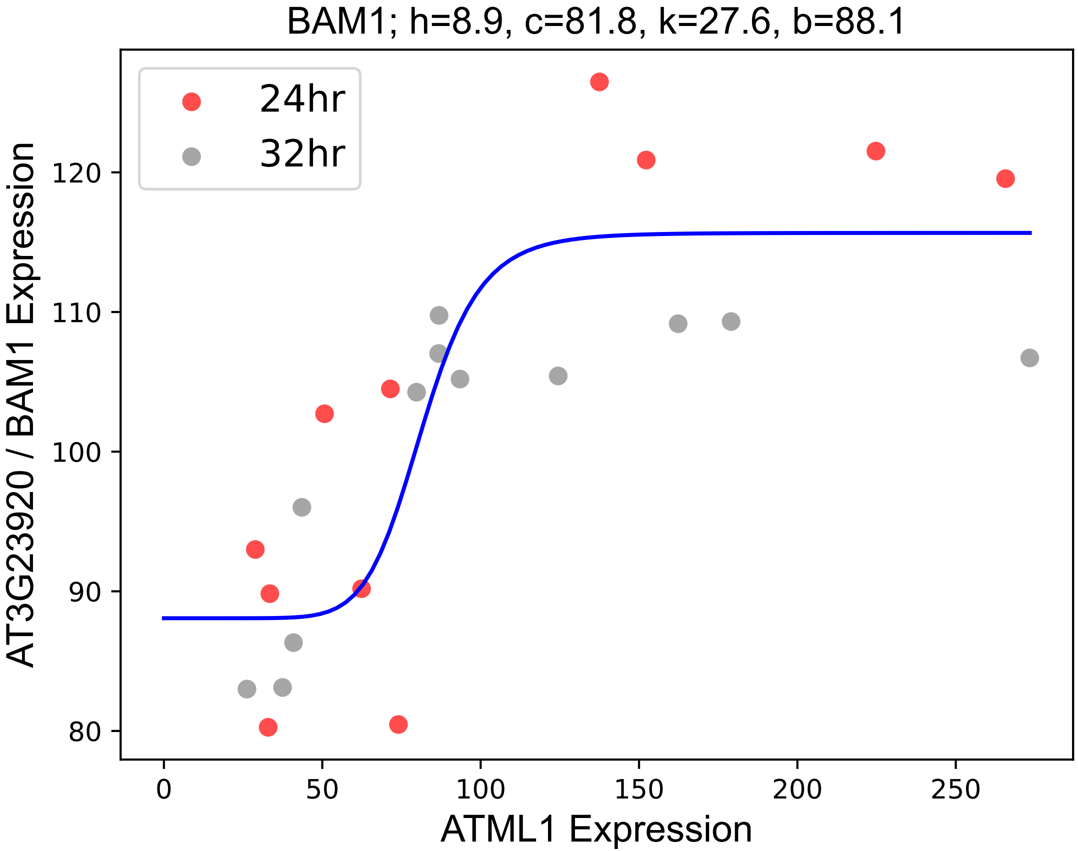

### BGLU25.pdf

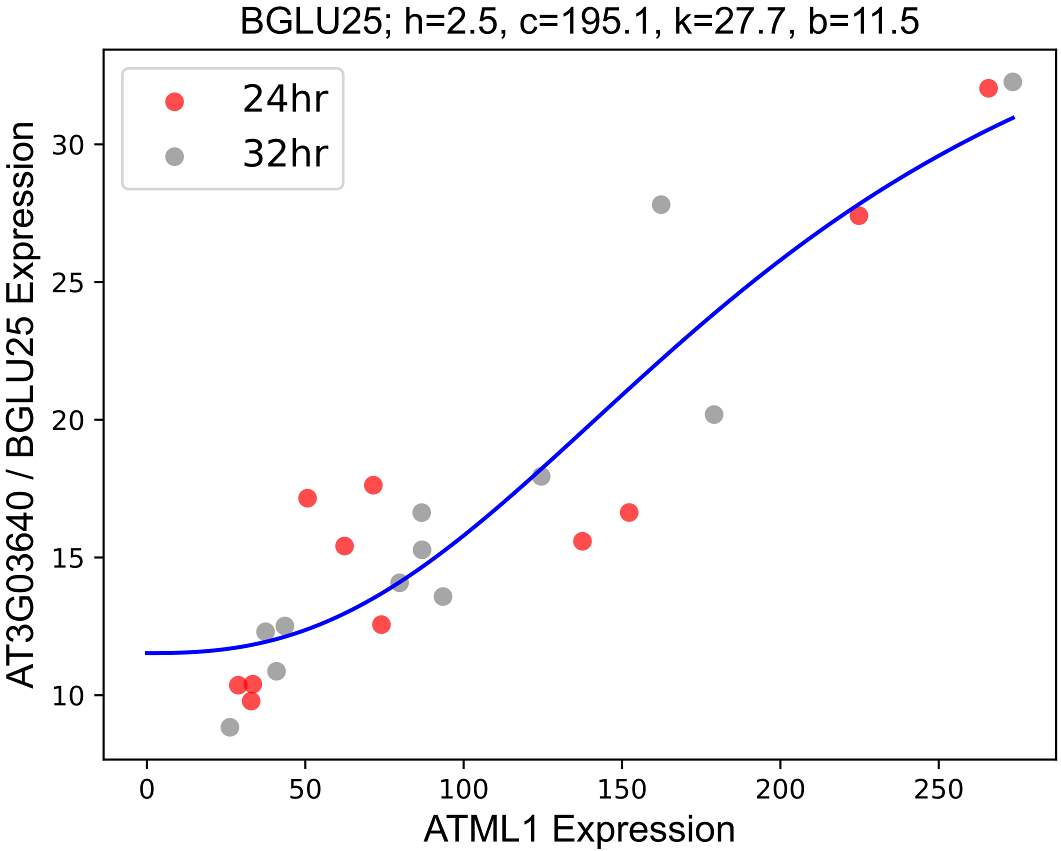

### BLH1.pdf

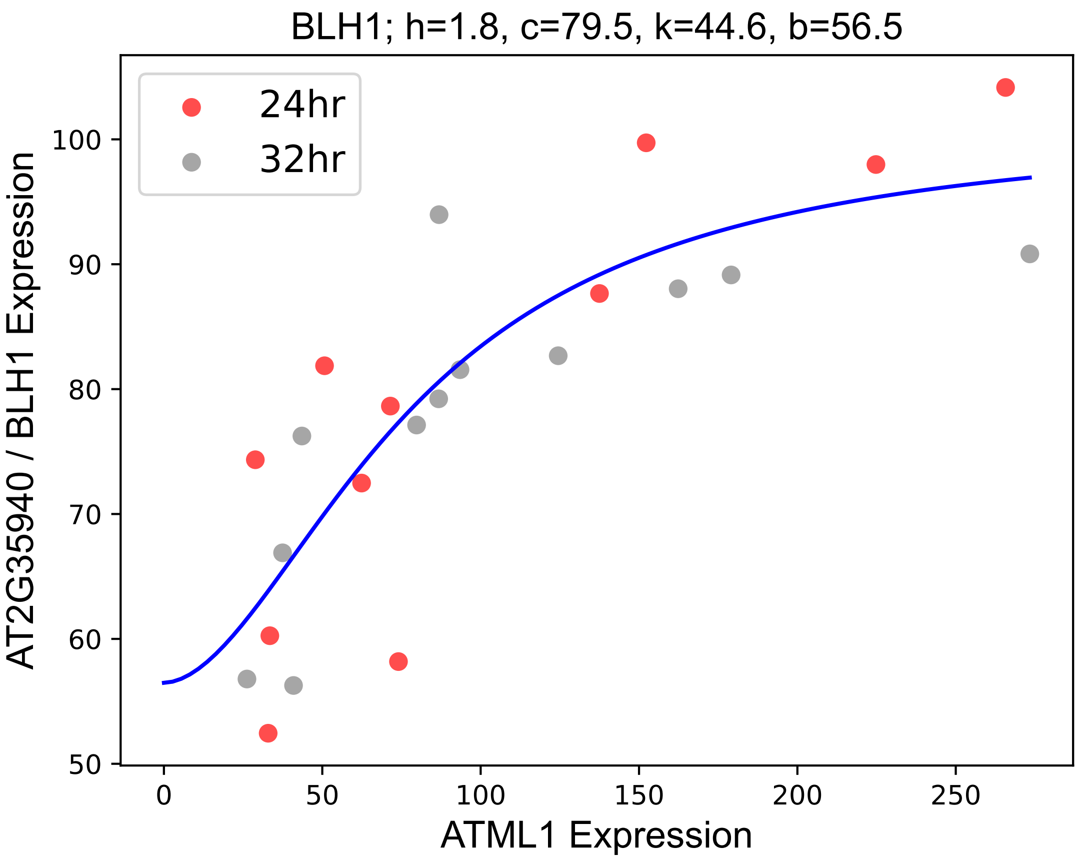

### CER1.pdf

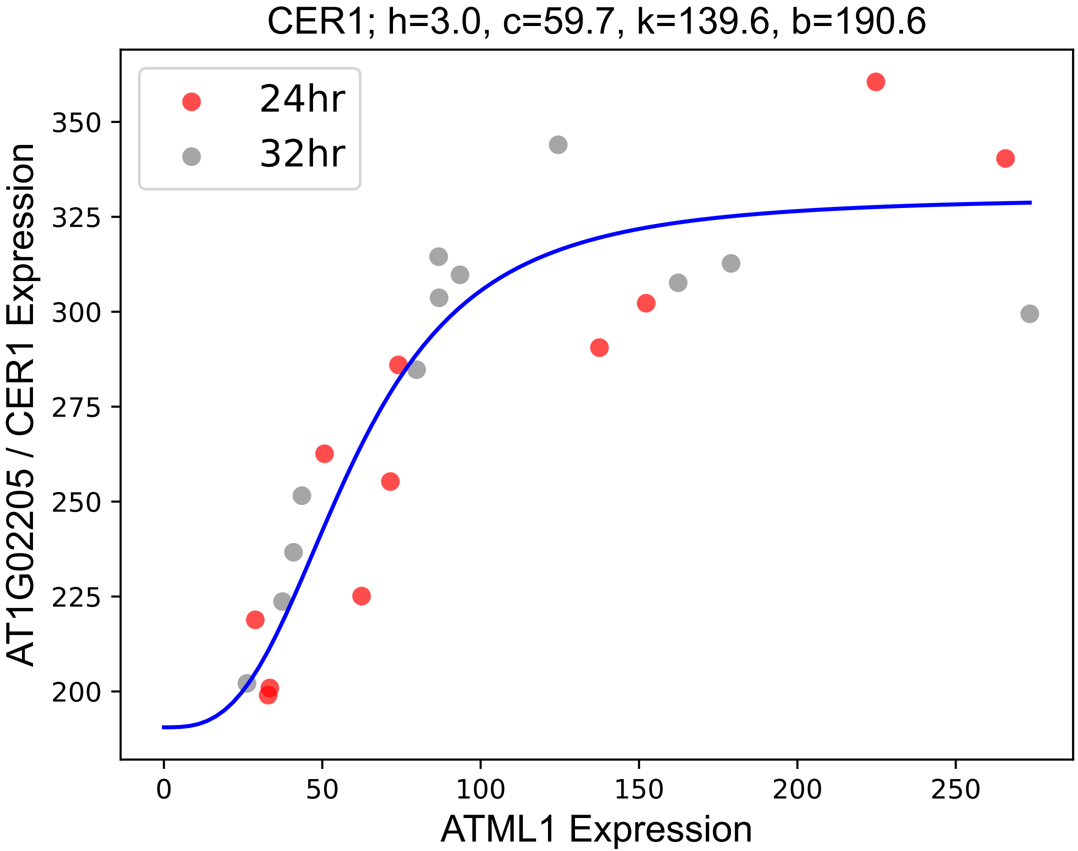

### CER3.pdf

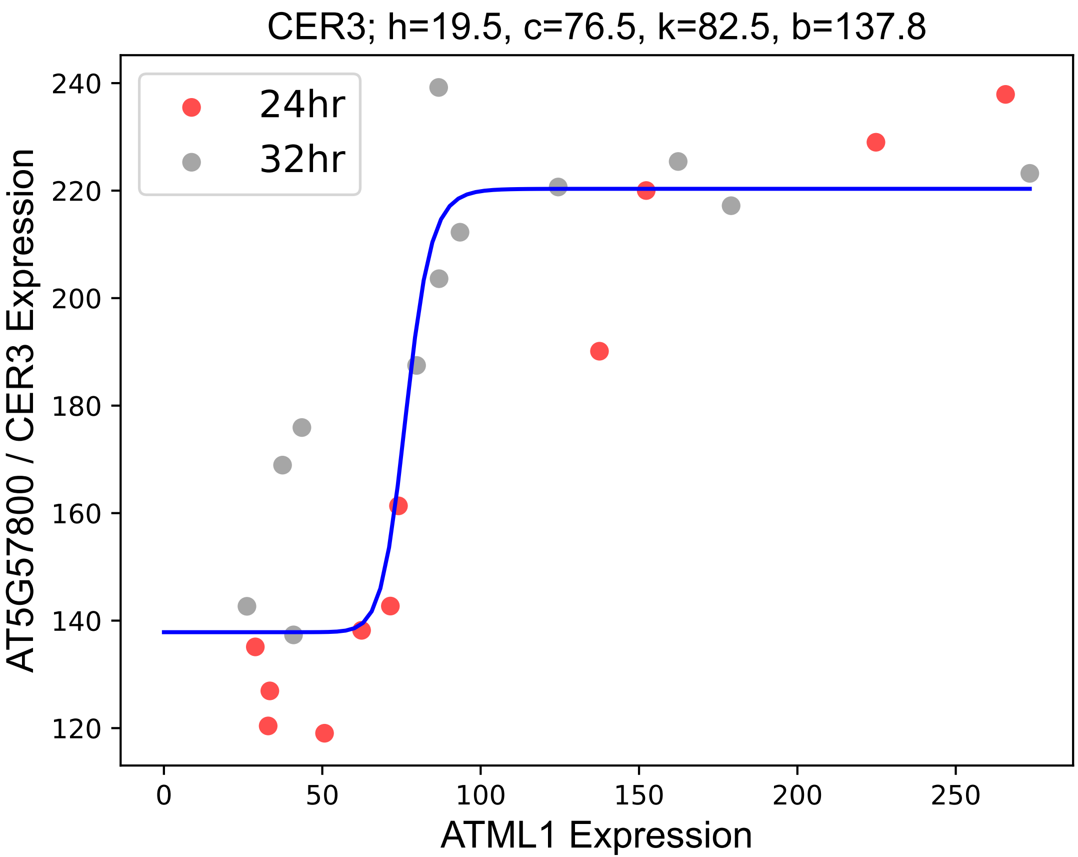

### CER5.pdf

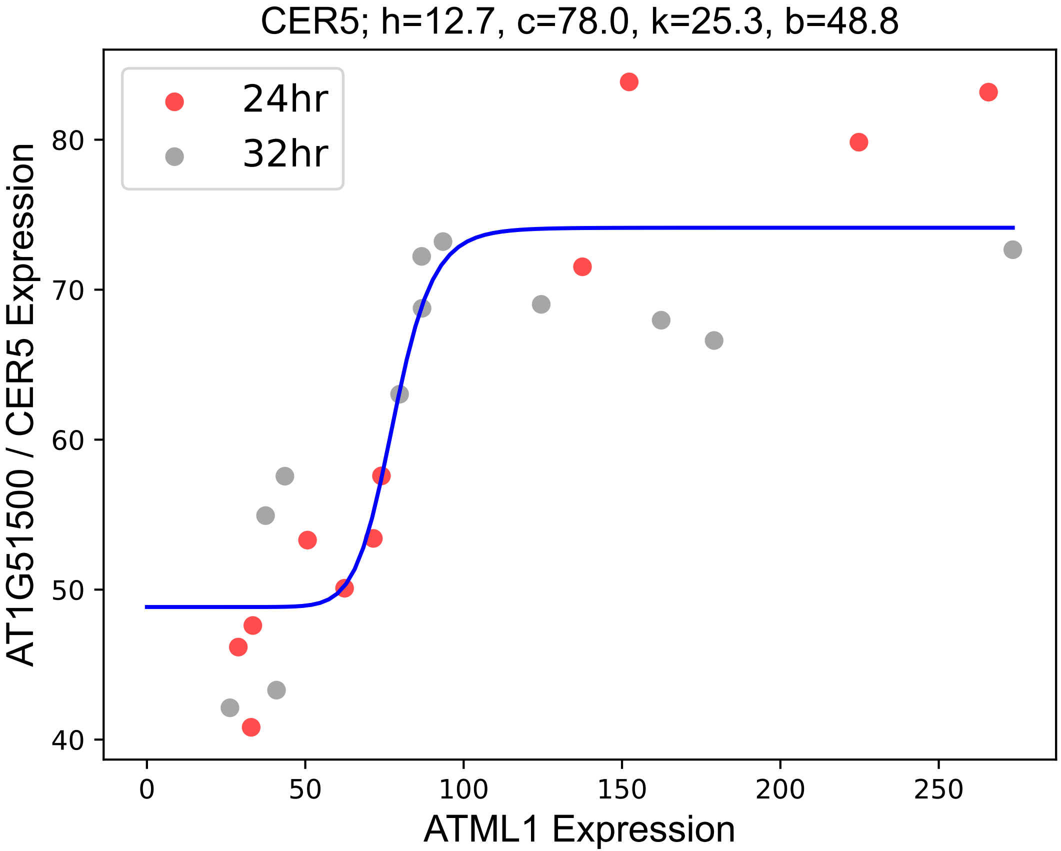

### CSLG3.pdf

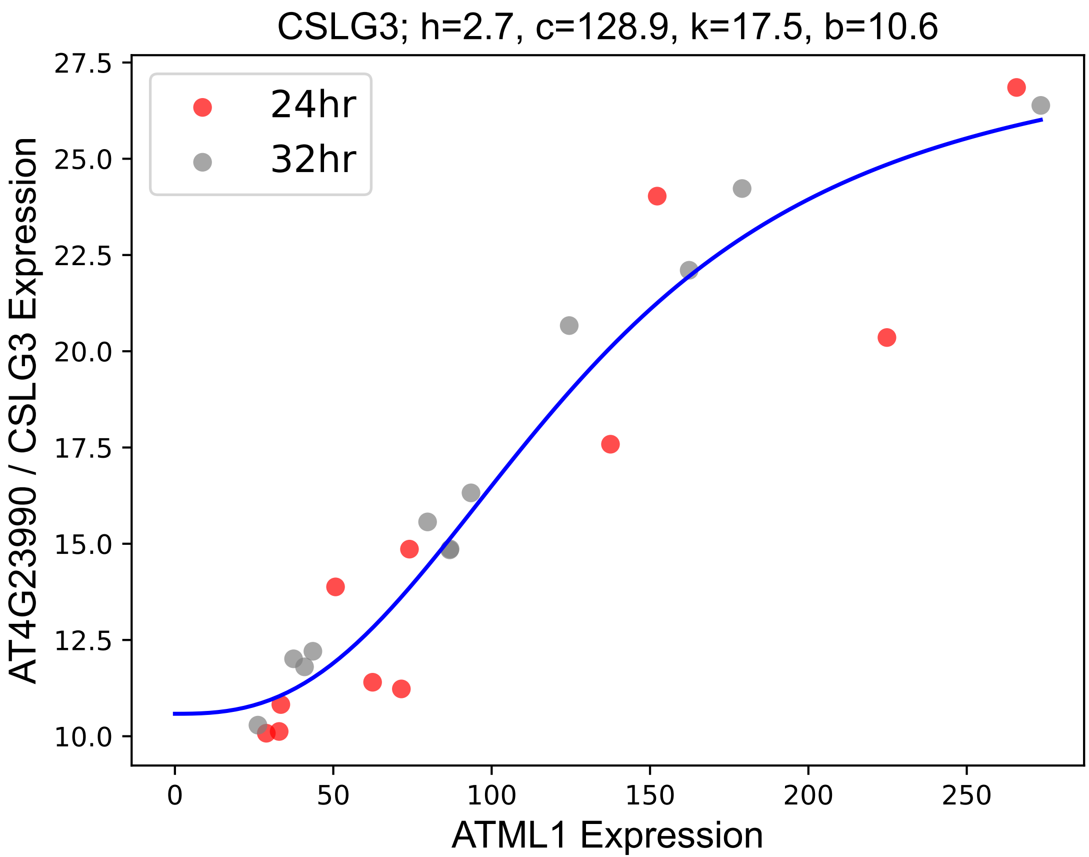

### CSY2.pdf

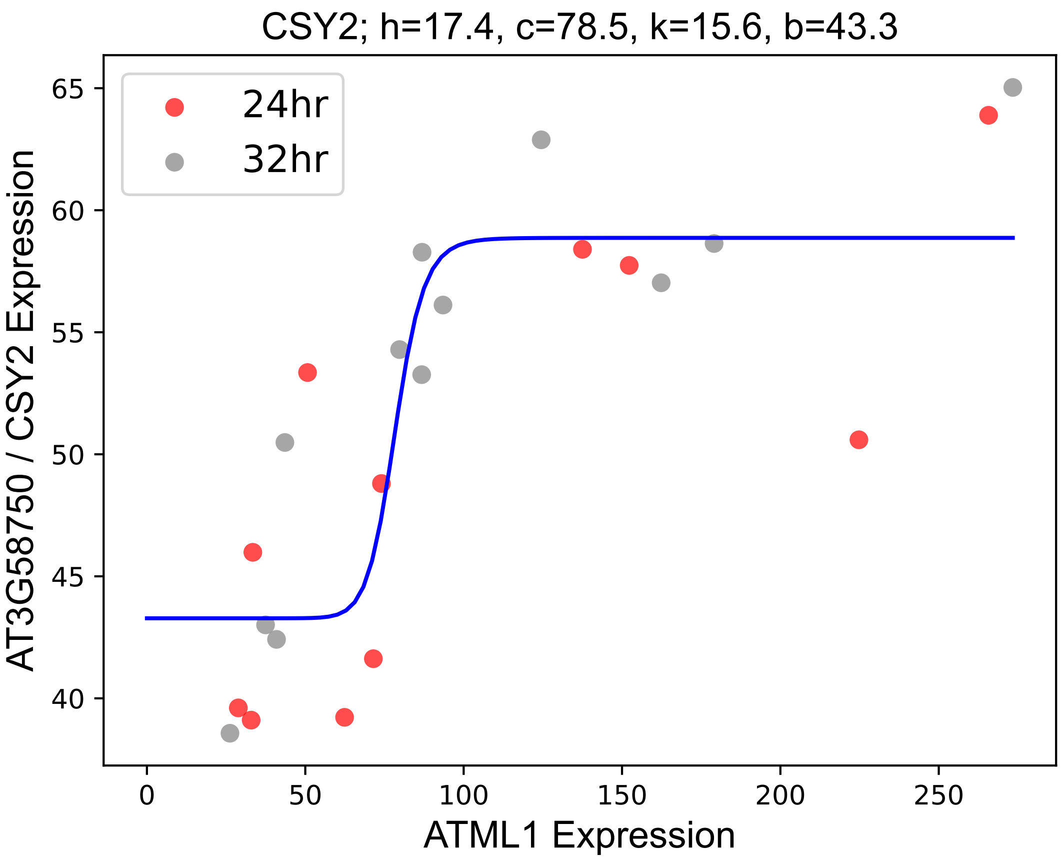

### CYP96A12.pdf

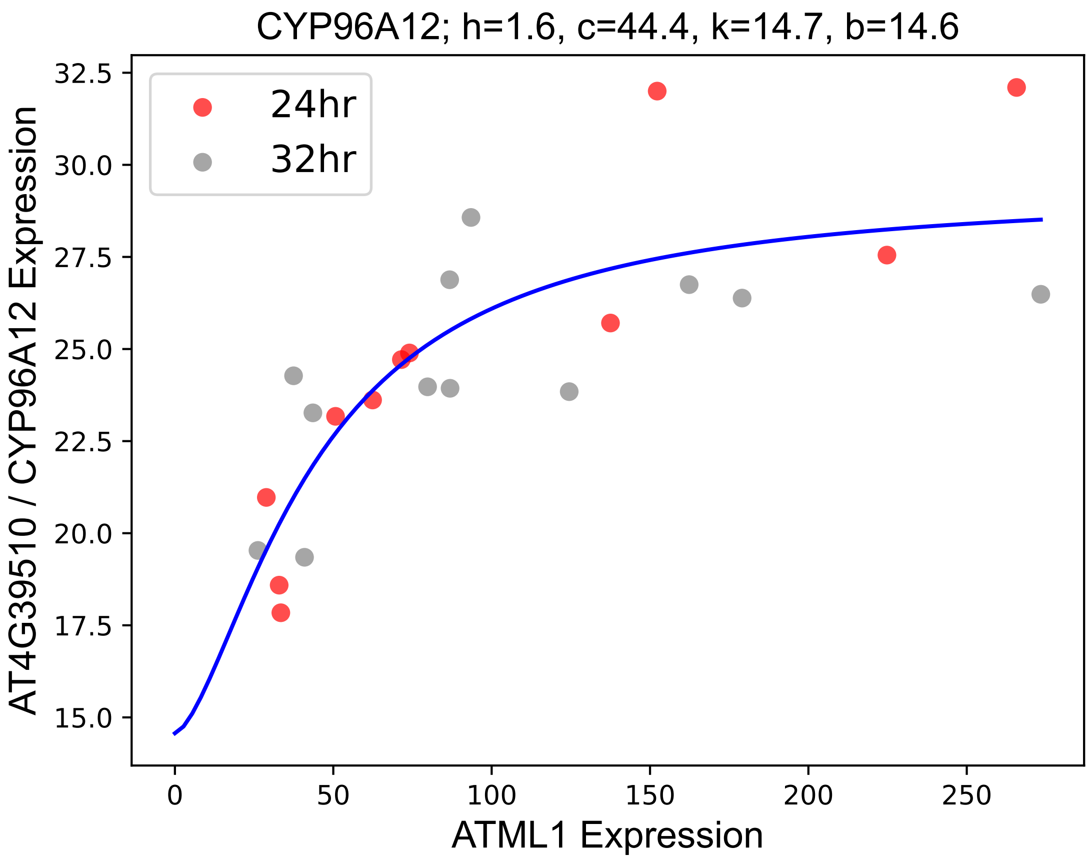

### DALL3.pdf

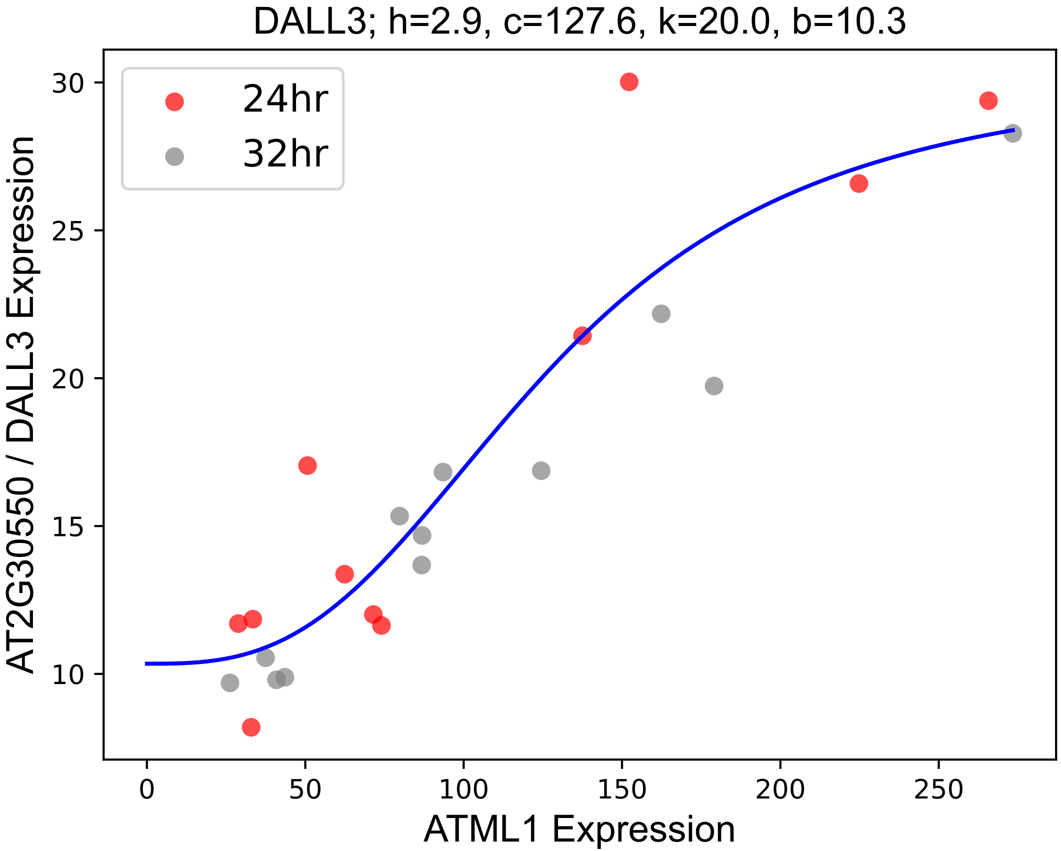

### DTX14.pdf

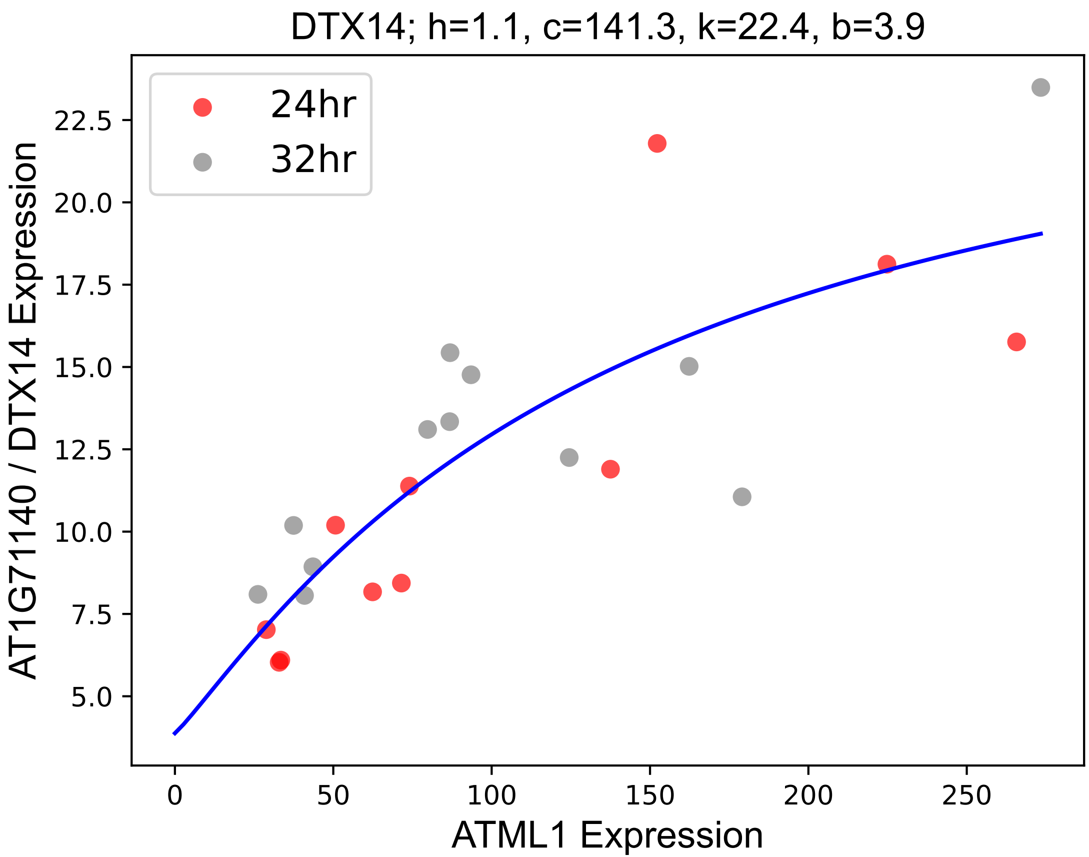

### ECA1.pdf

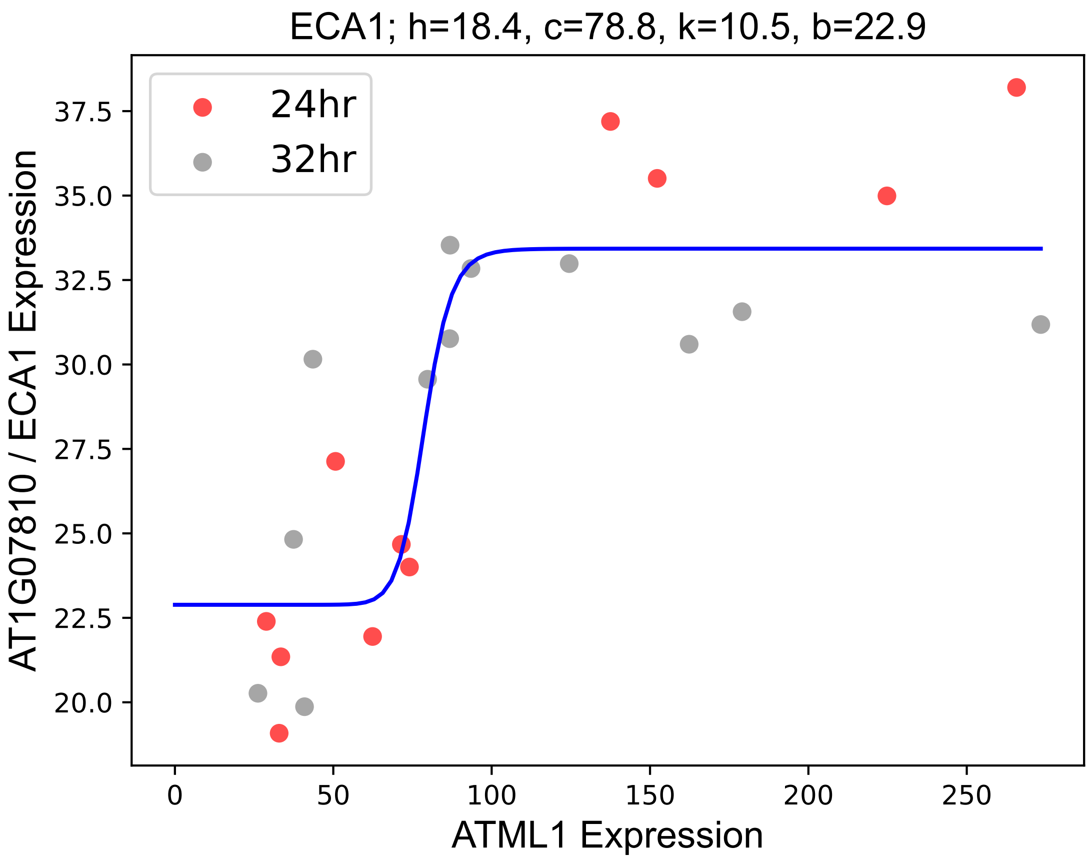

### EXPA16.pdf

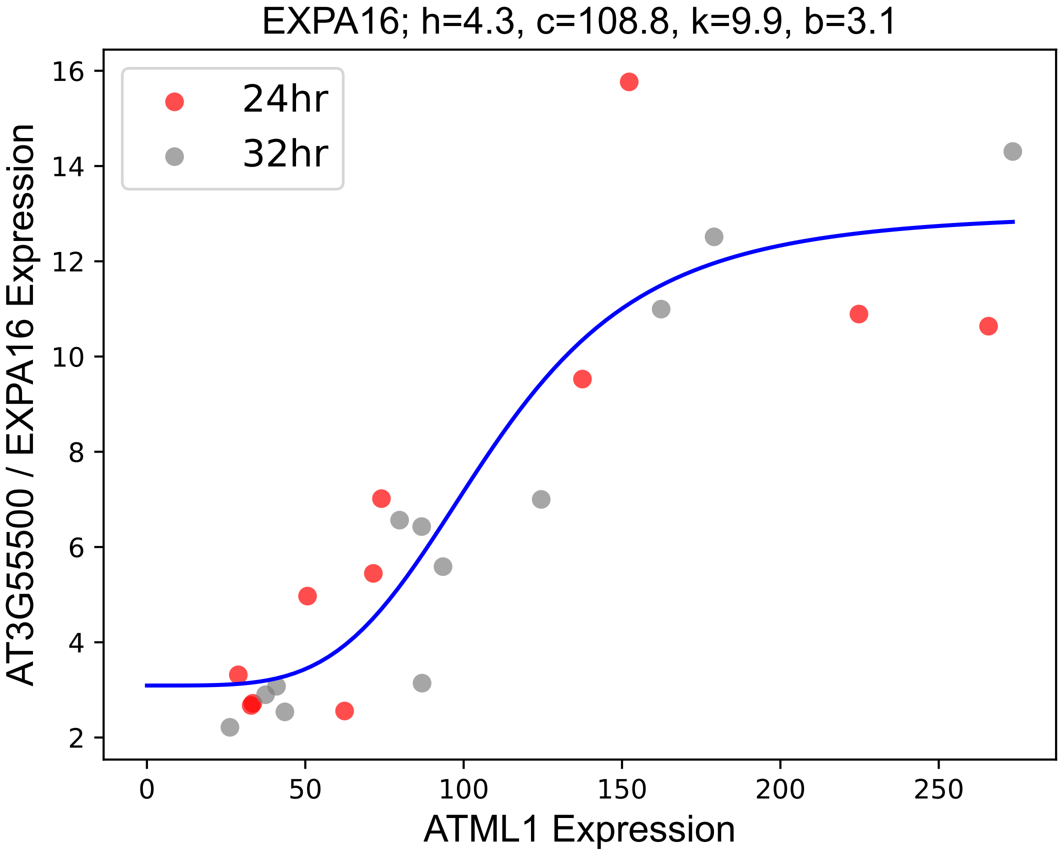

### FDH.pdf

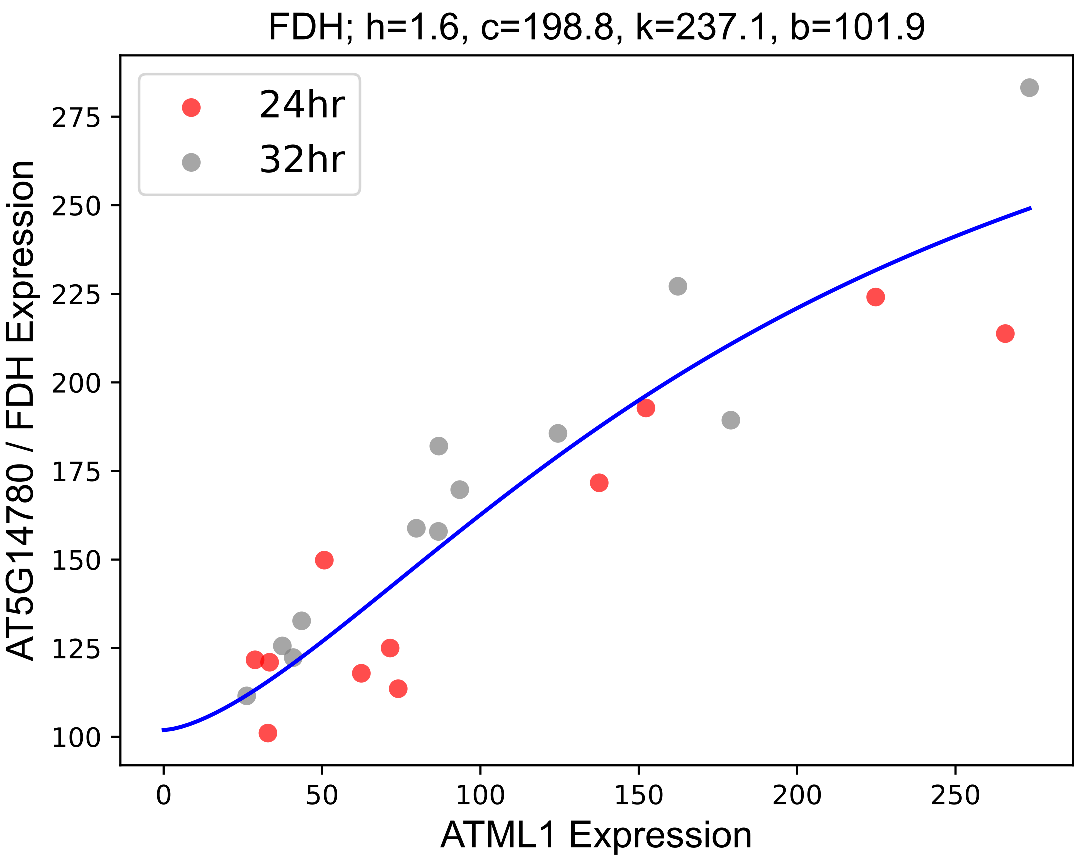

### GAD.pdf

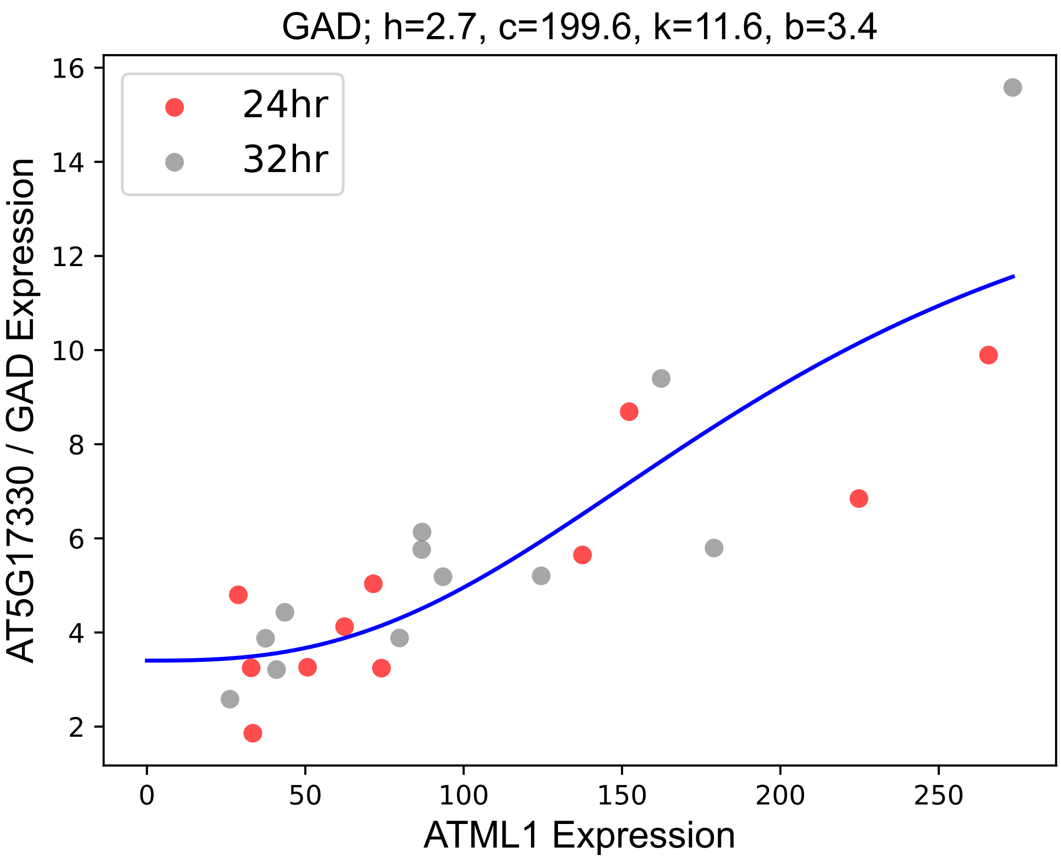

### GDH1.pdf

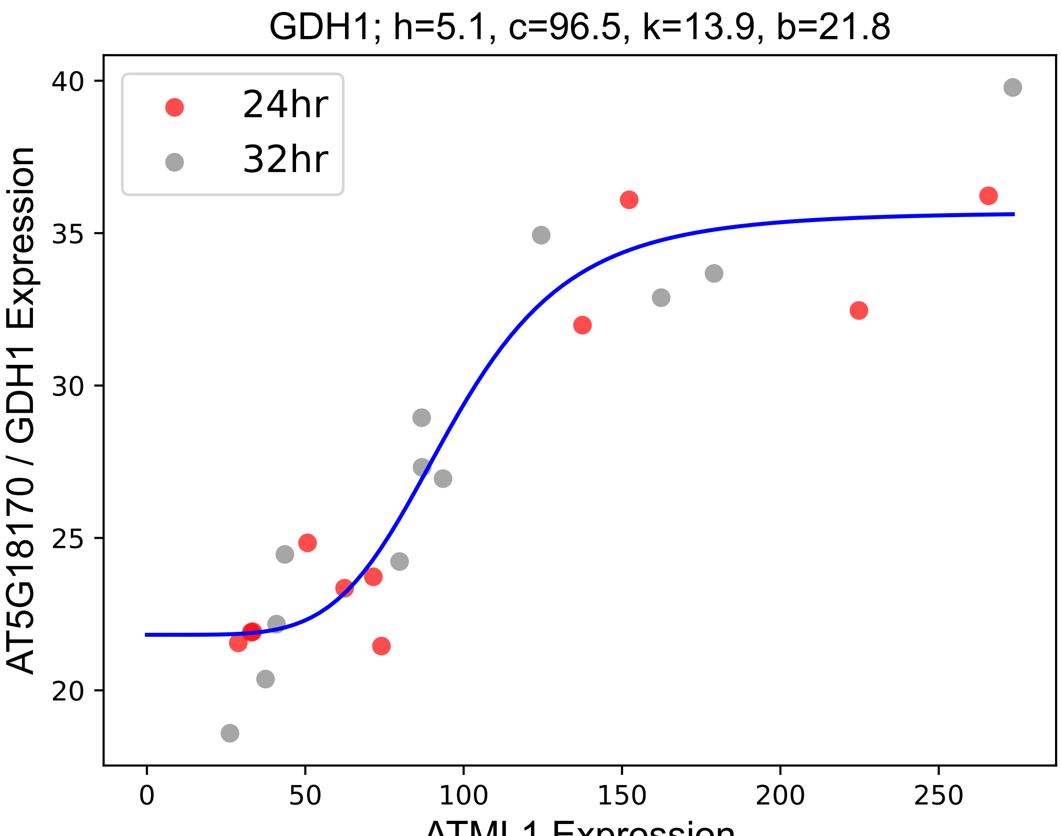

### GPAT4.pdf

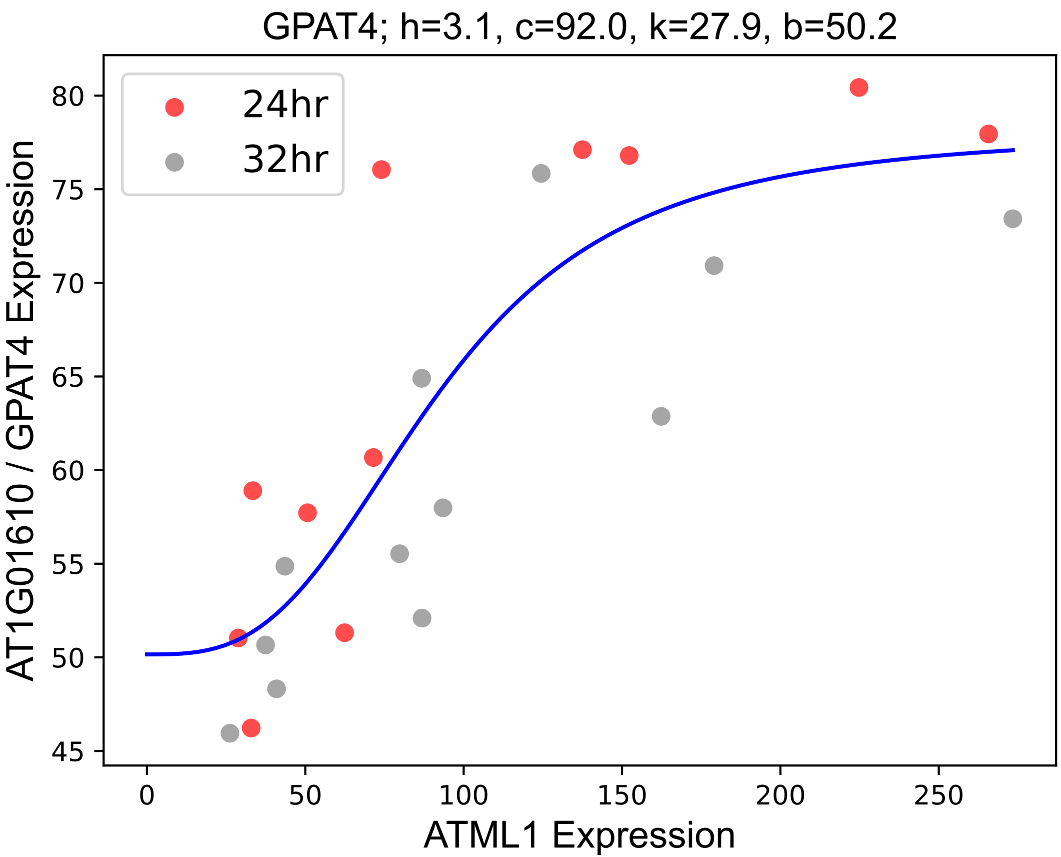

### Hop2.pdf

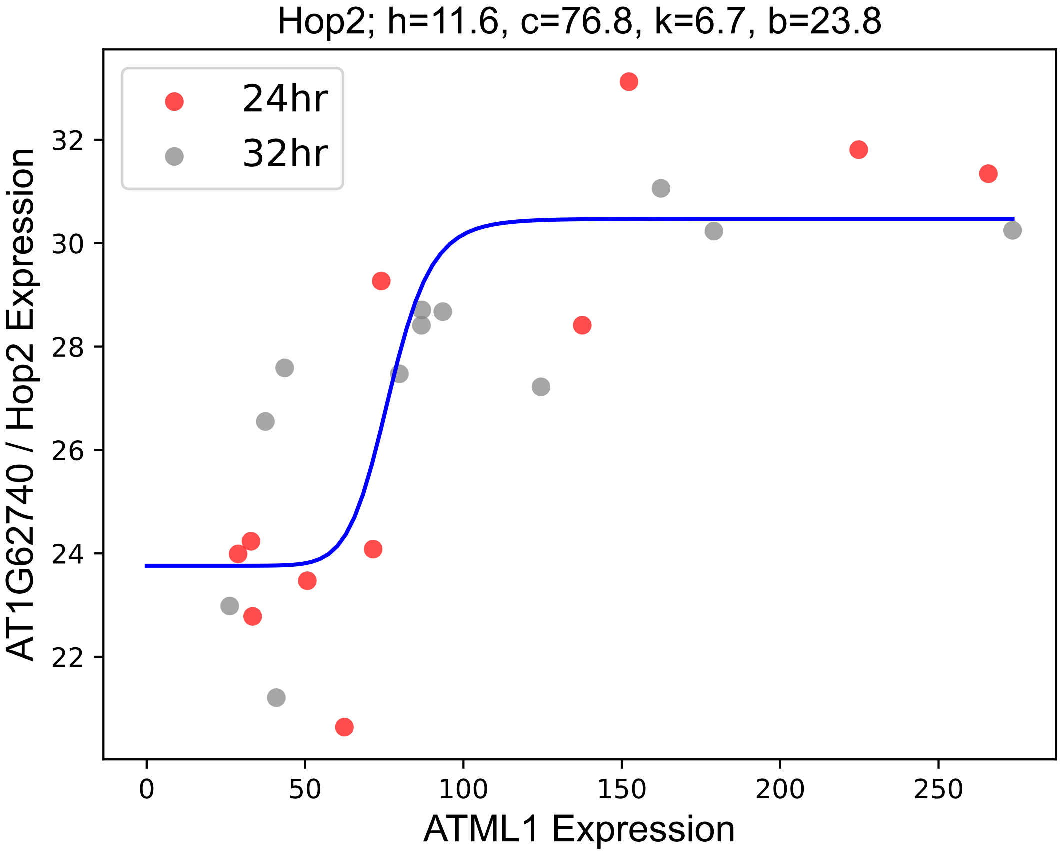

### ICE1.pdf

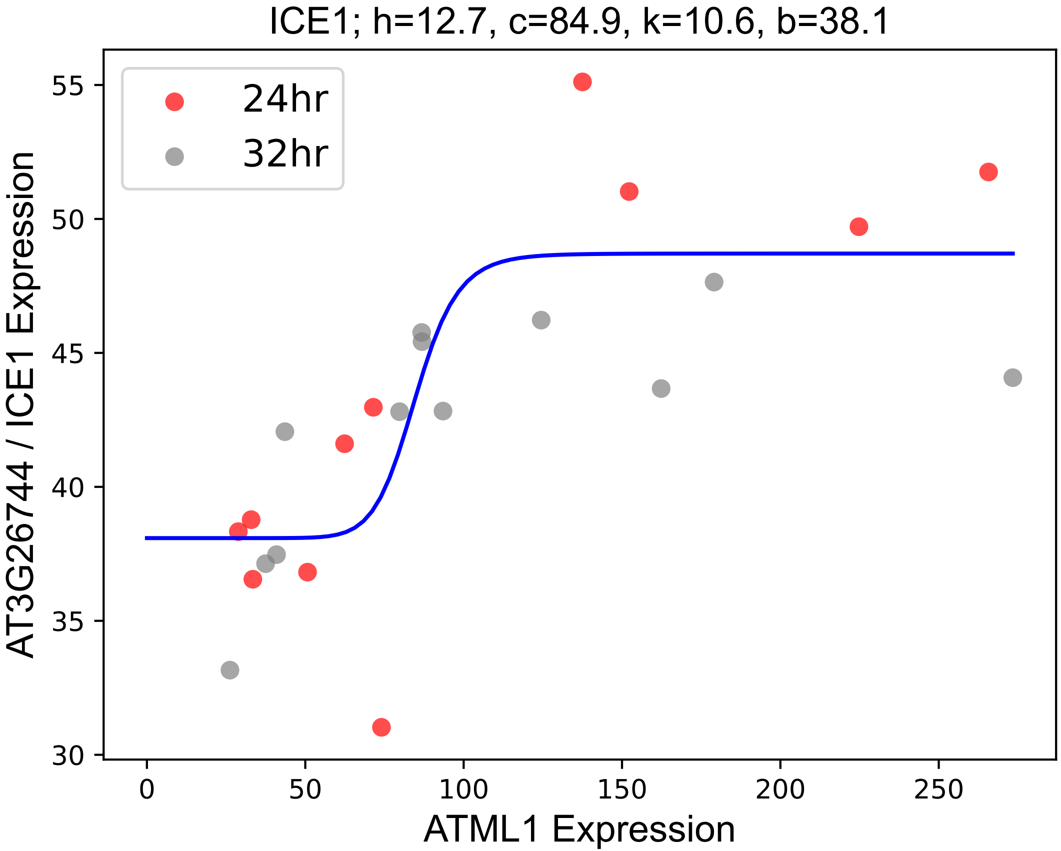

### ICK3.pdf

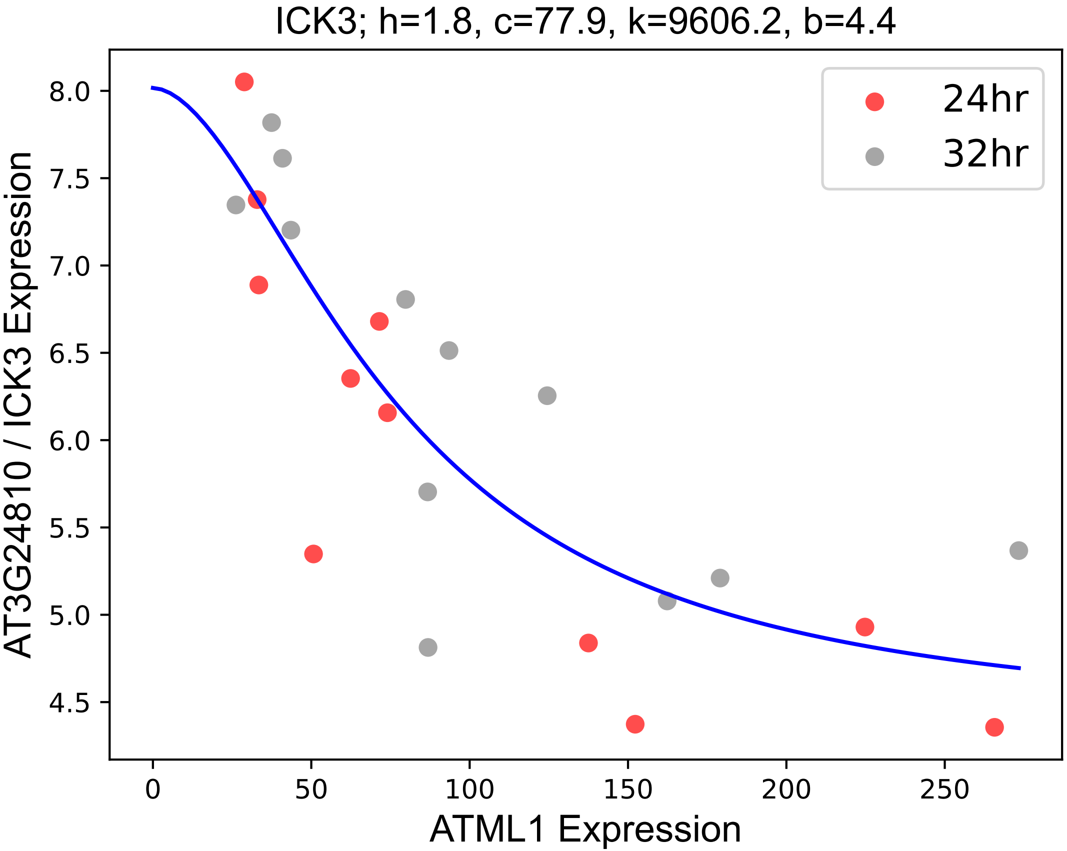

### JAL22.pdf

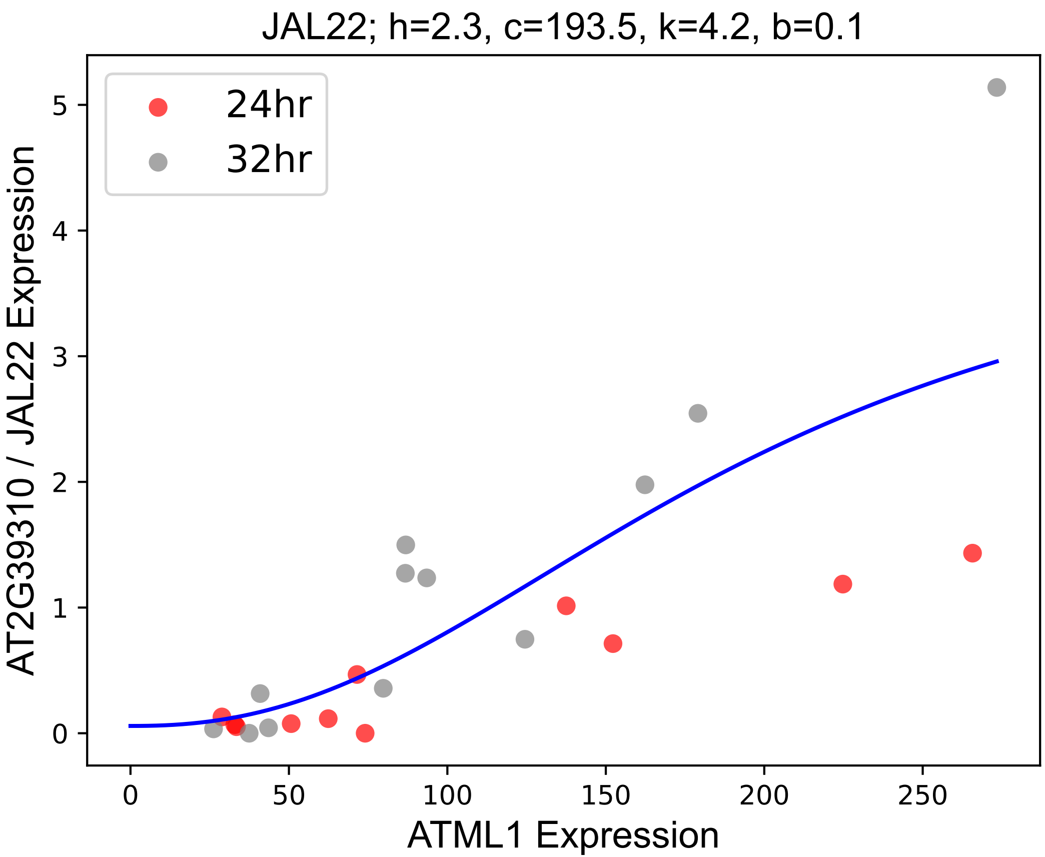

### JAL34.pdf

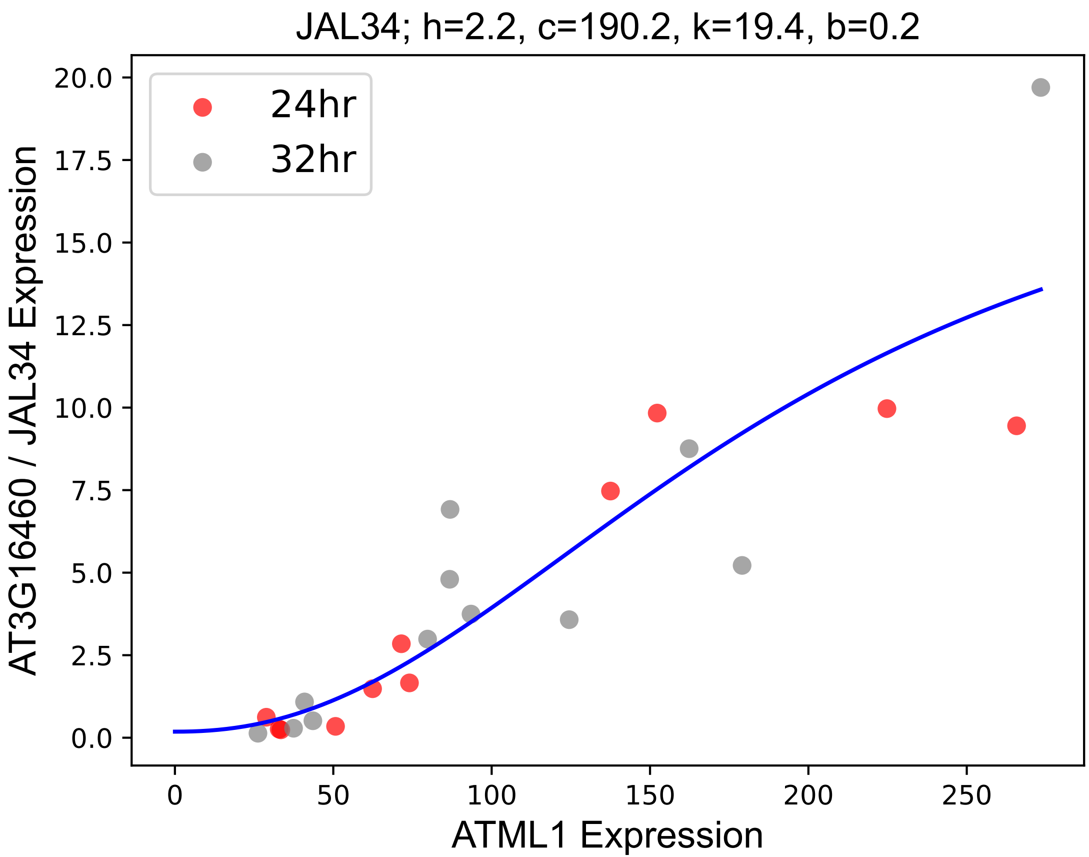

### LECRK-IV.3.pdf

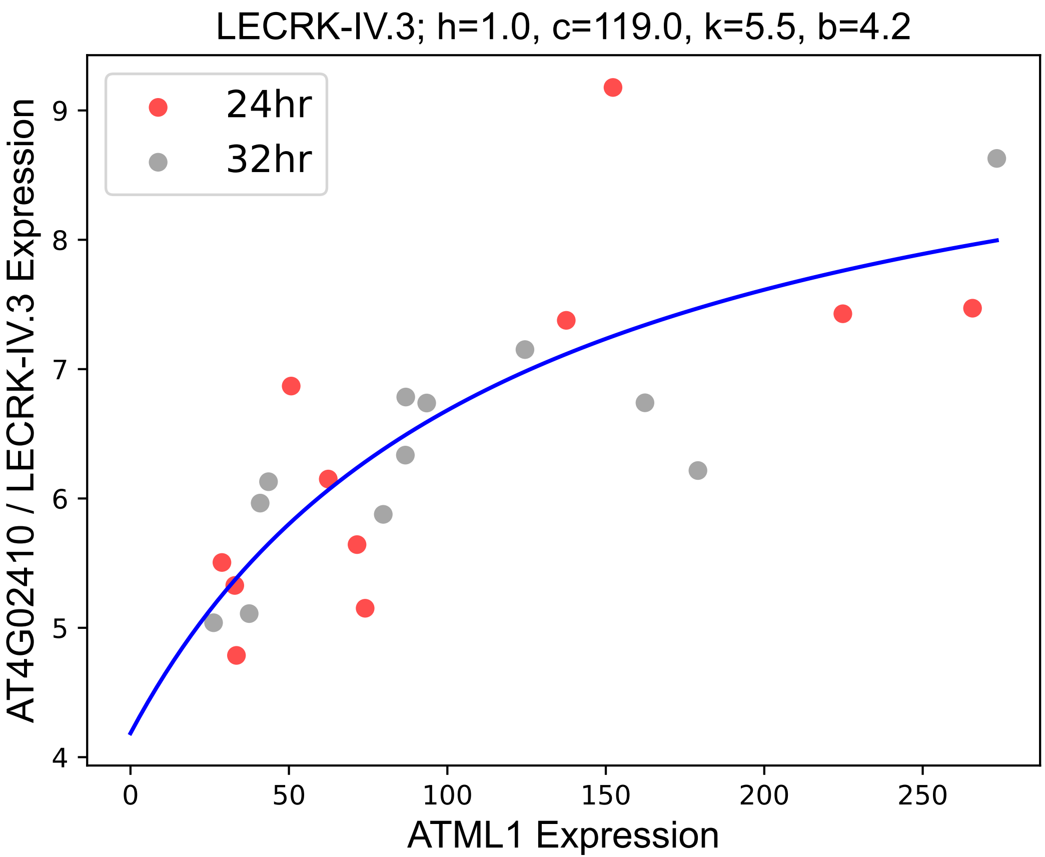

### LTI30.pdf

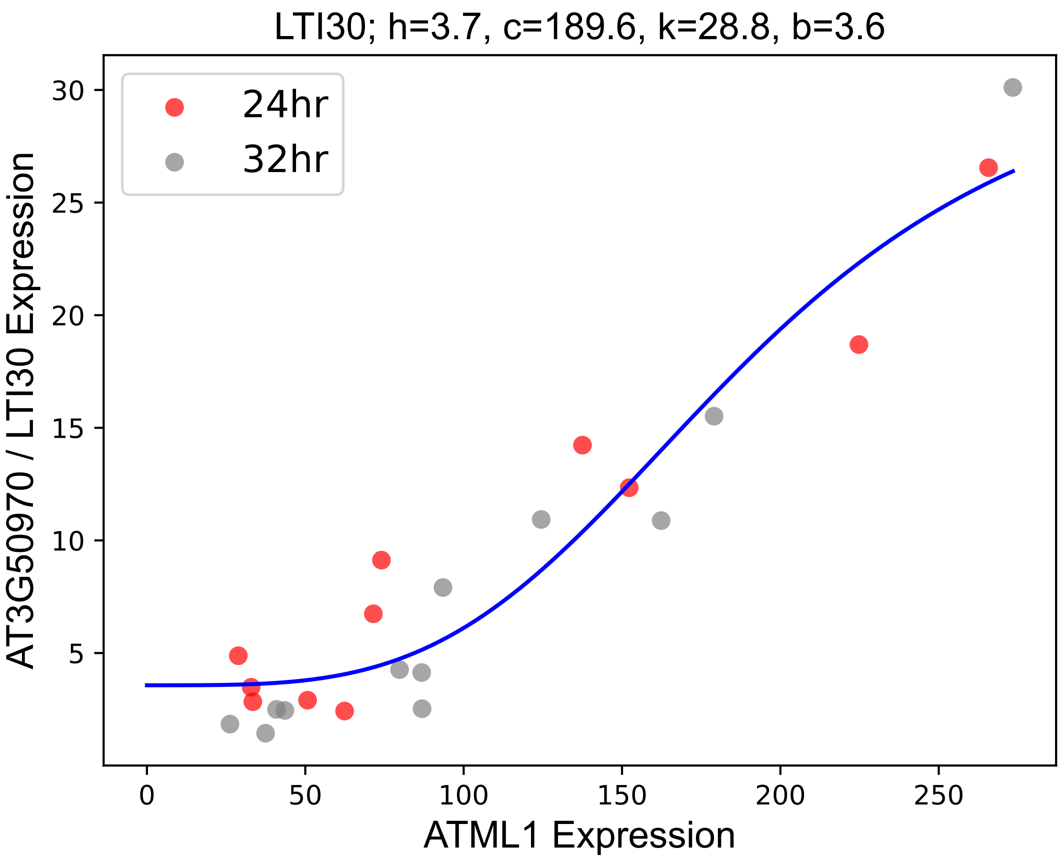

### LTP.pdf

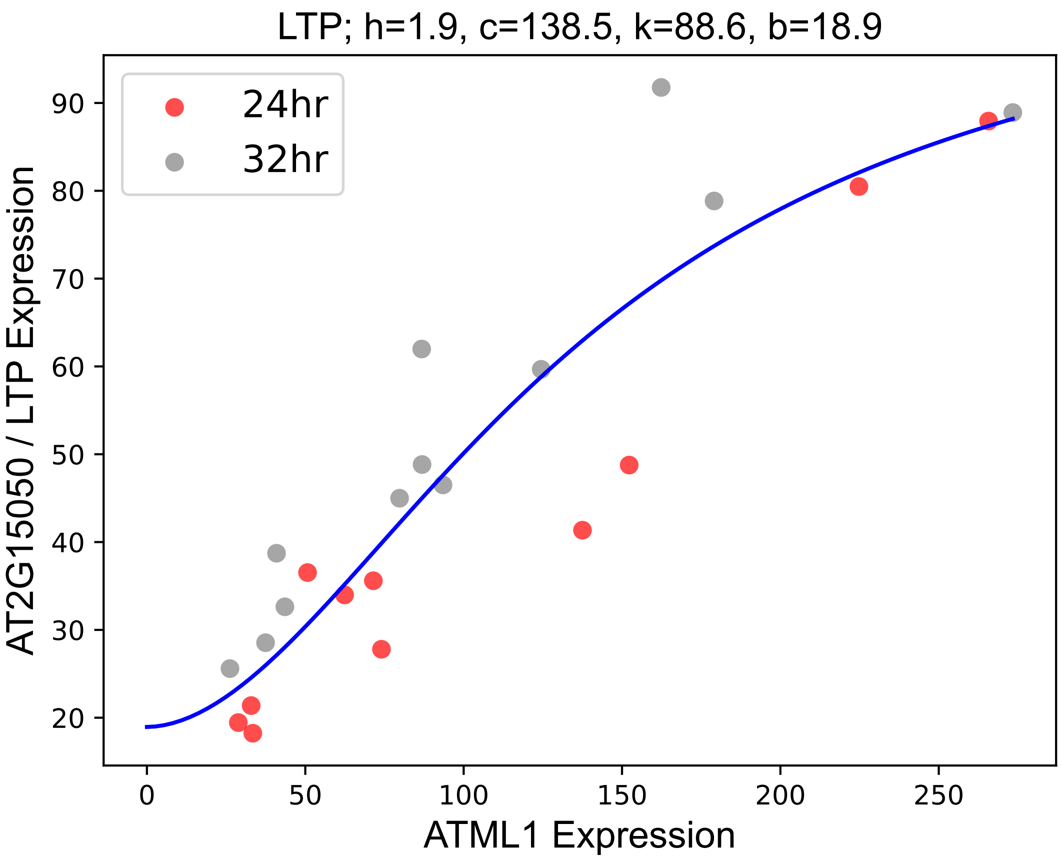

### MLO3.pdf

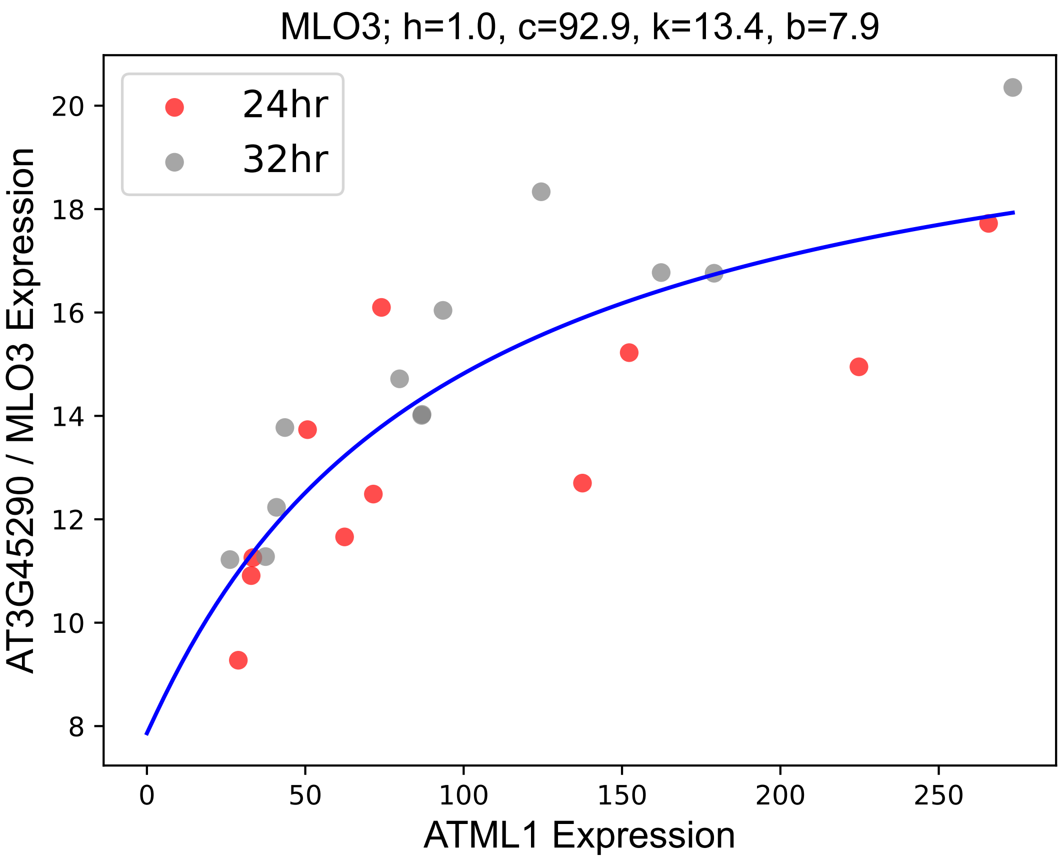

### MSS1.pdf

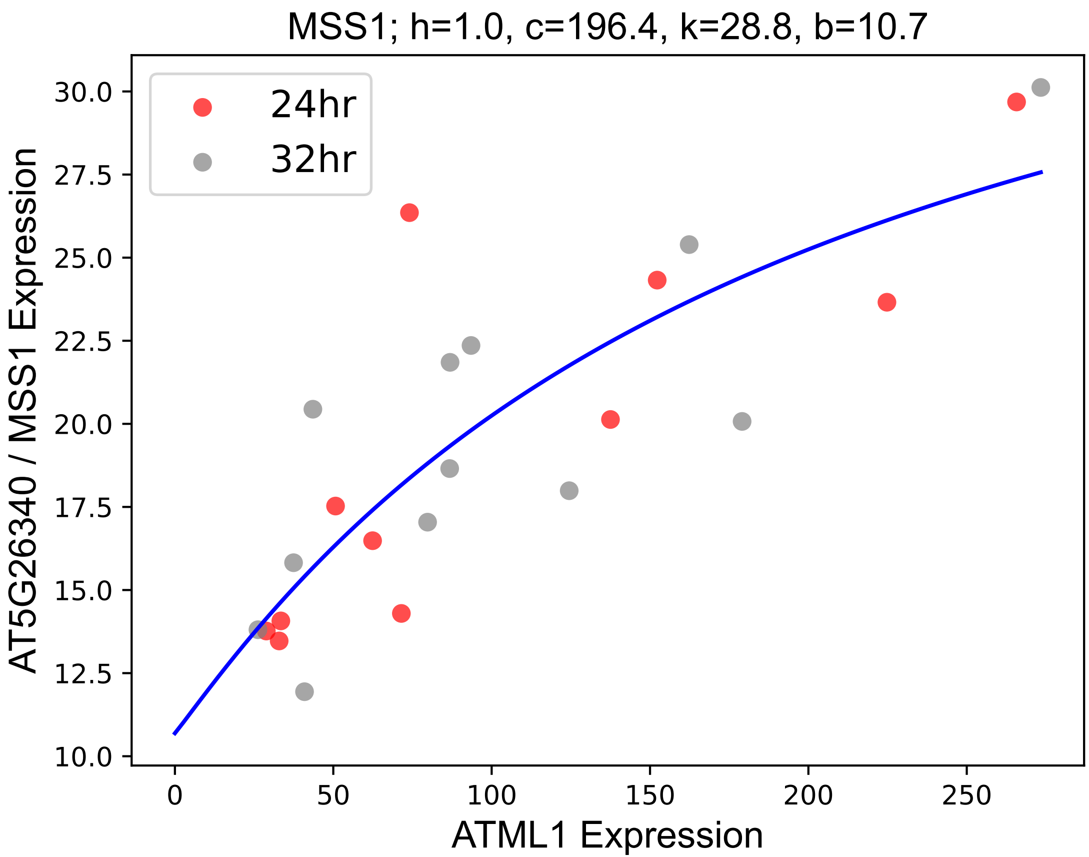

### MYB30.pdf

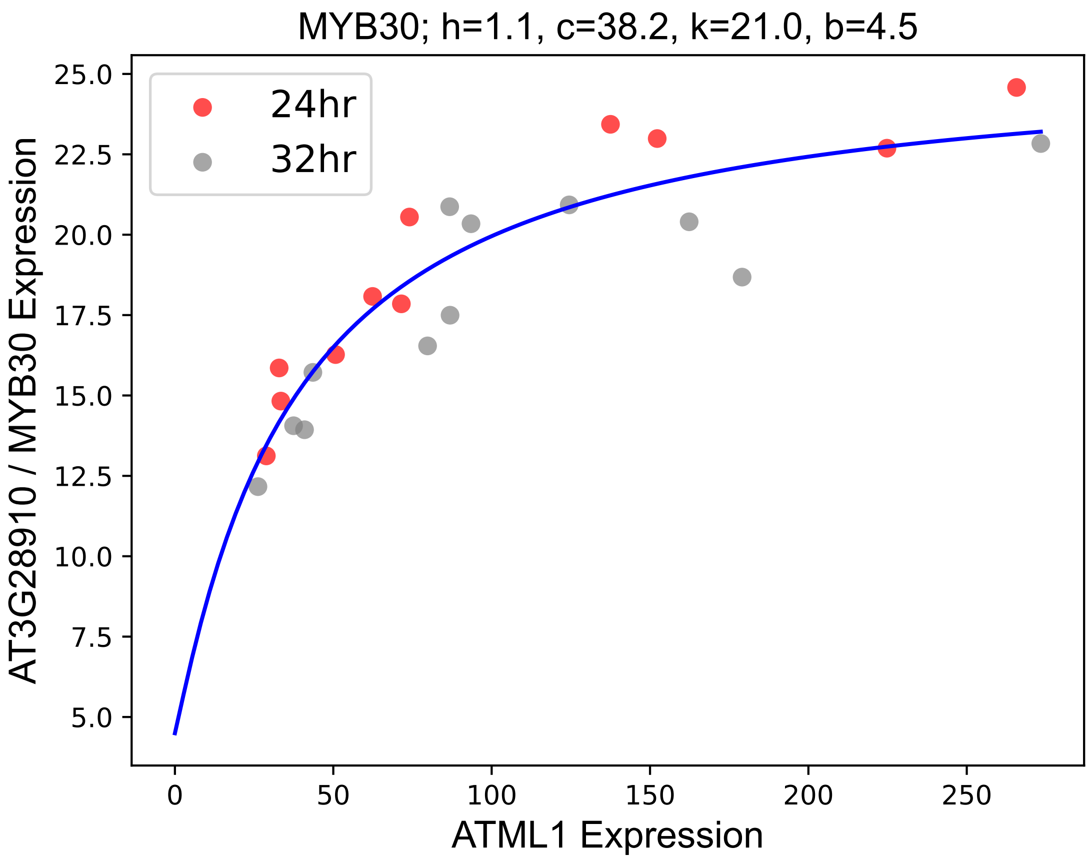

### MYB31.pdf

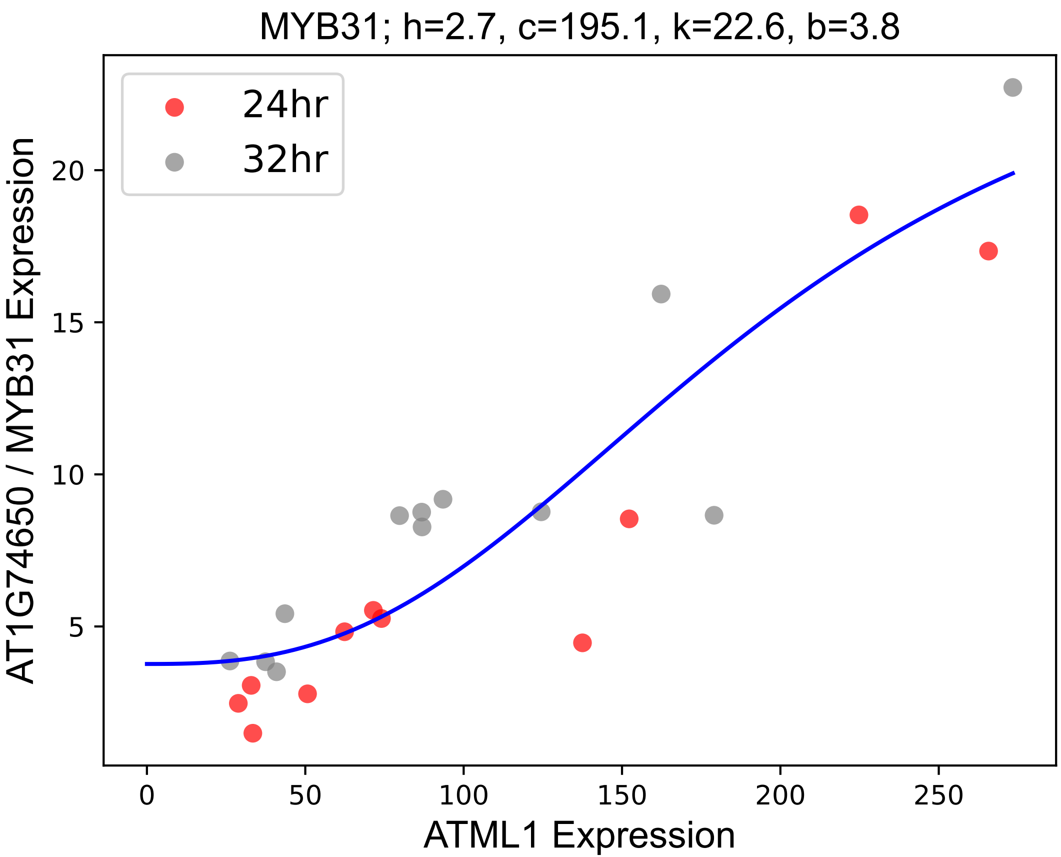

### MYB66.pdf

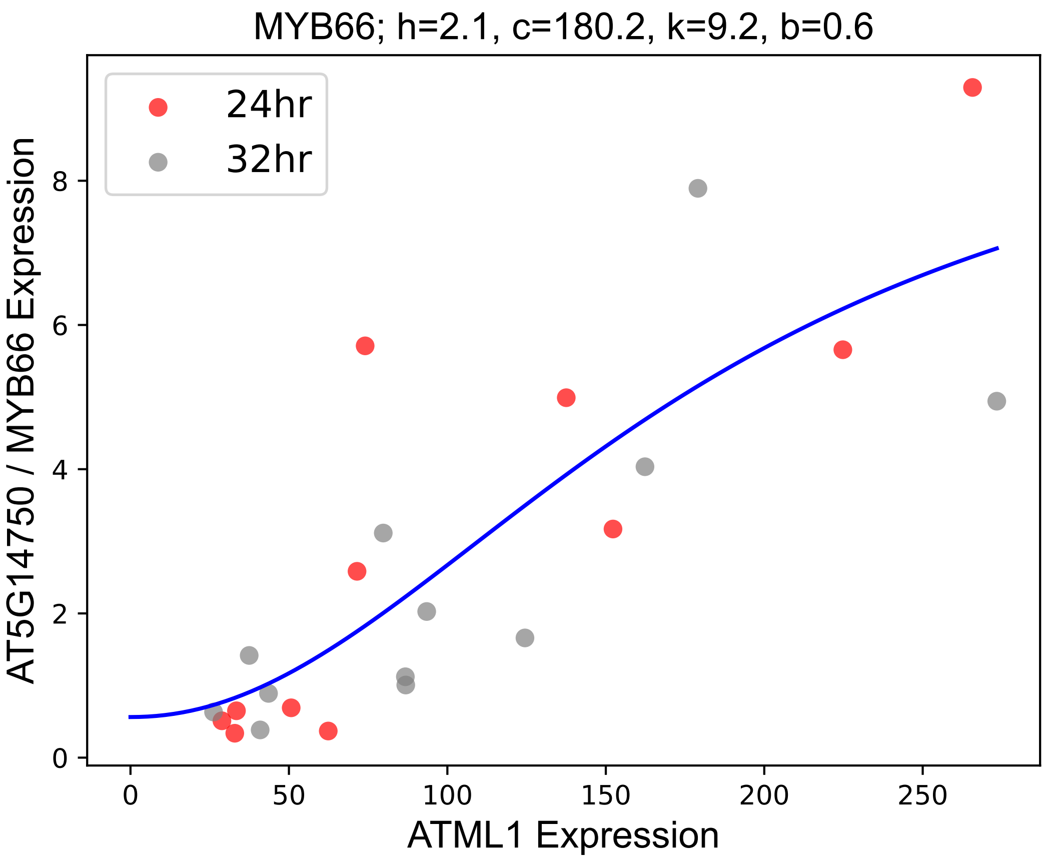

### NAI1.pdf

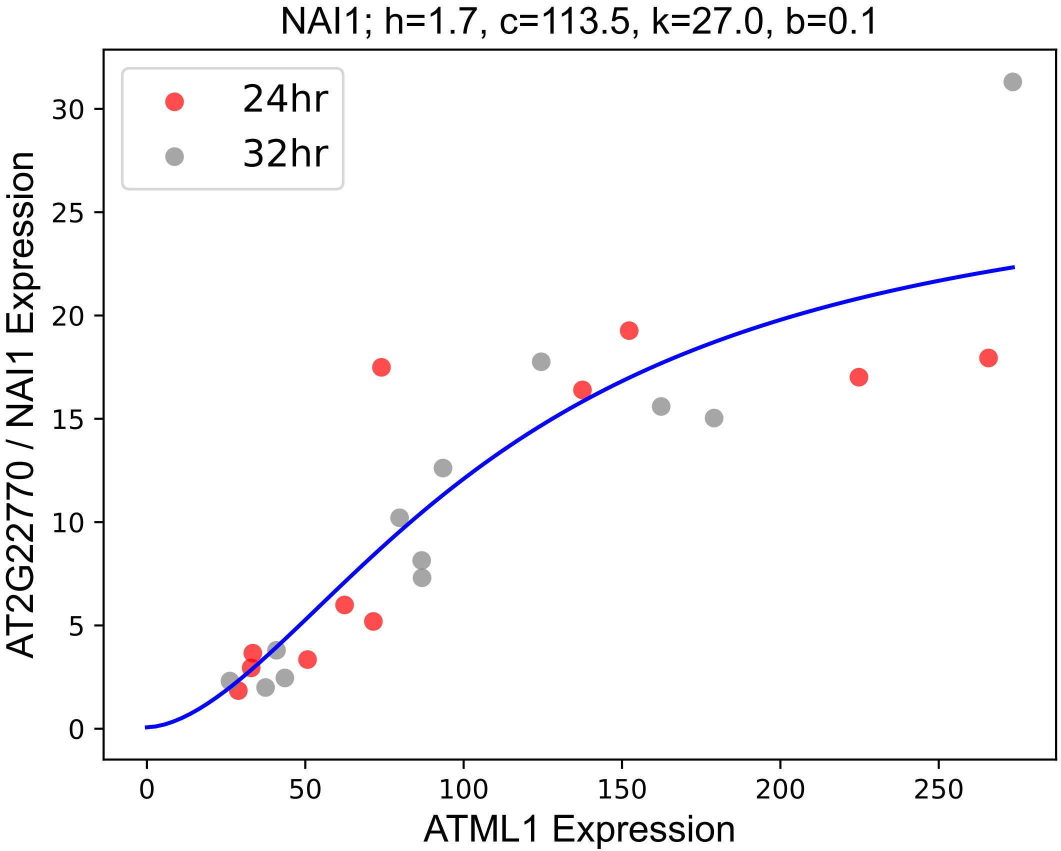

### NANA.pdf

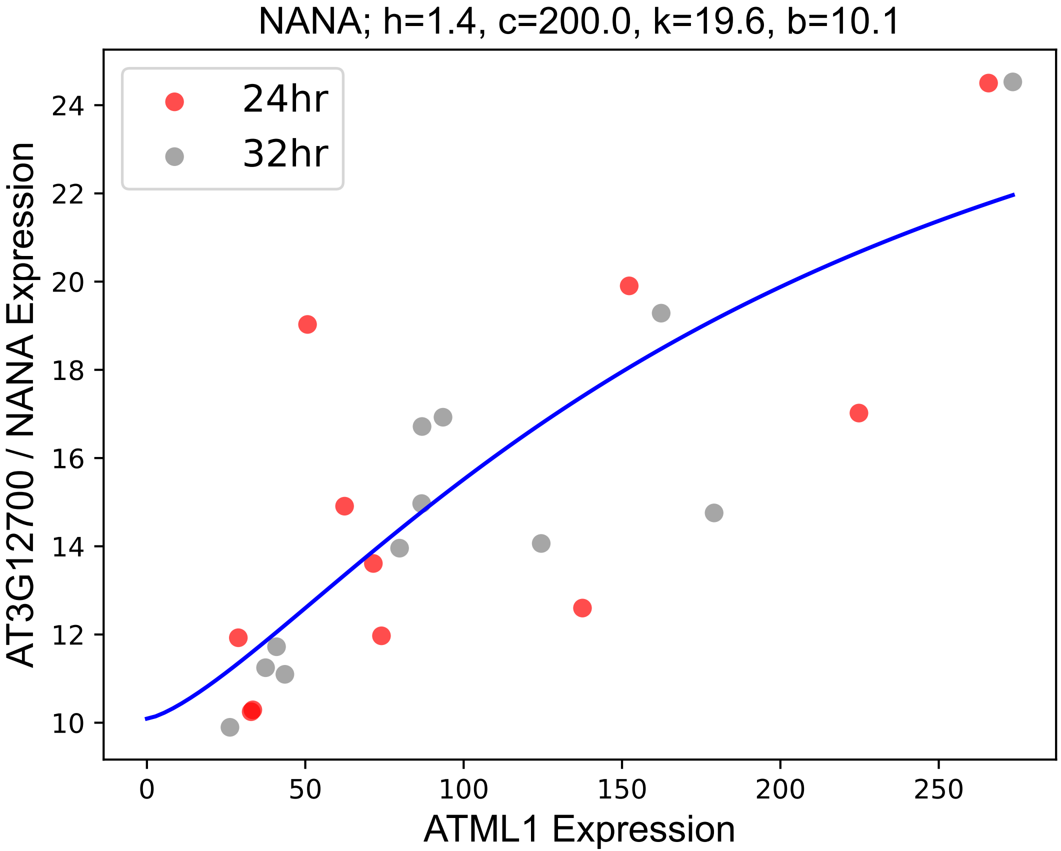

### NF-YC12.pdf

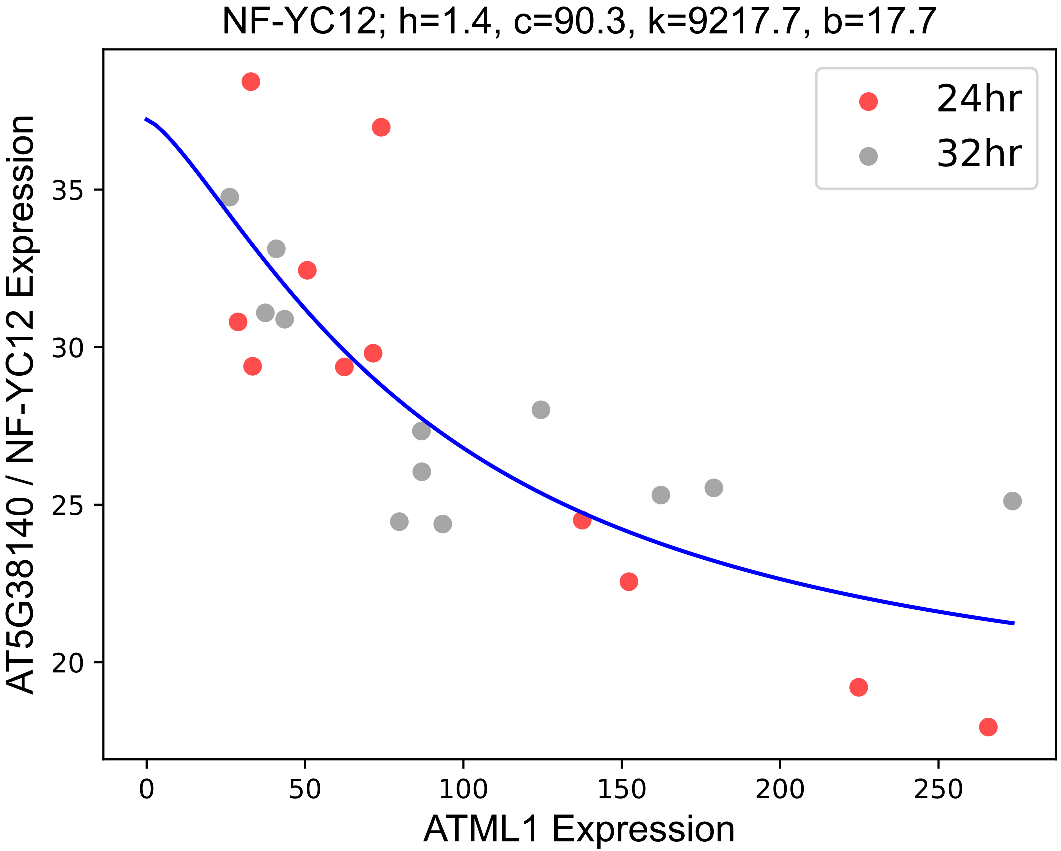

### NRAMP3.pdf

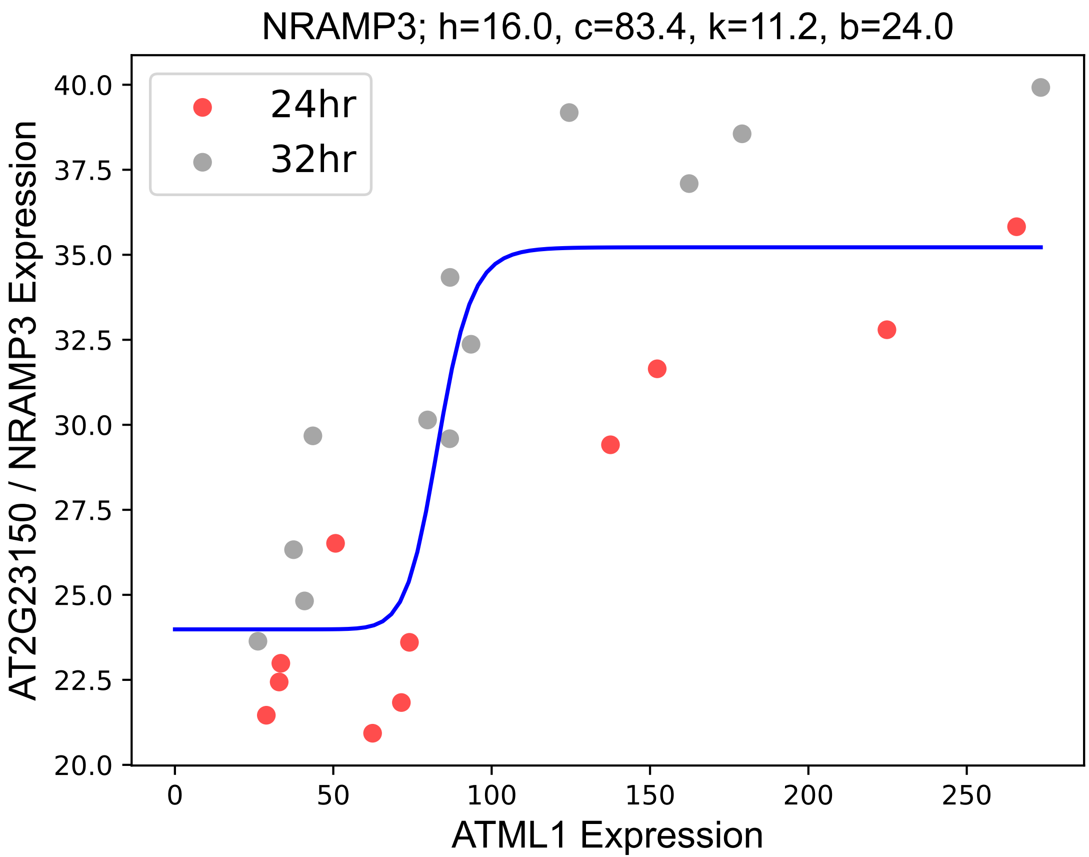

### NSP1.pdf

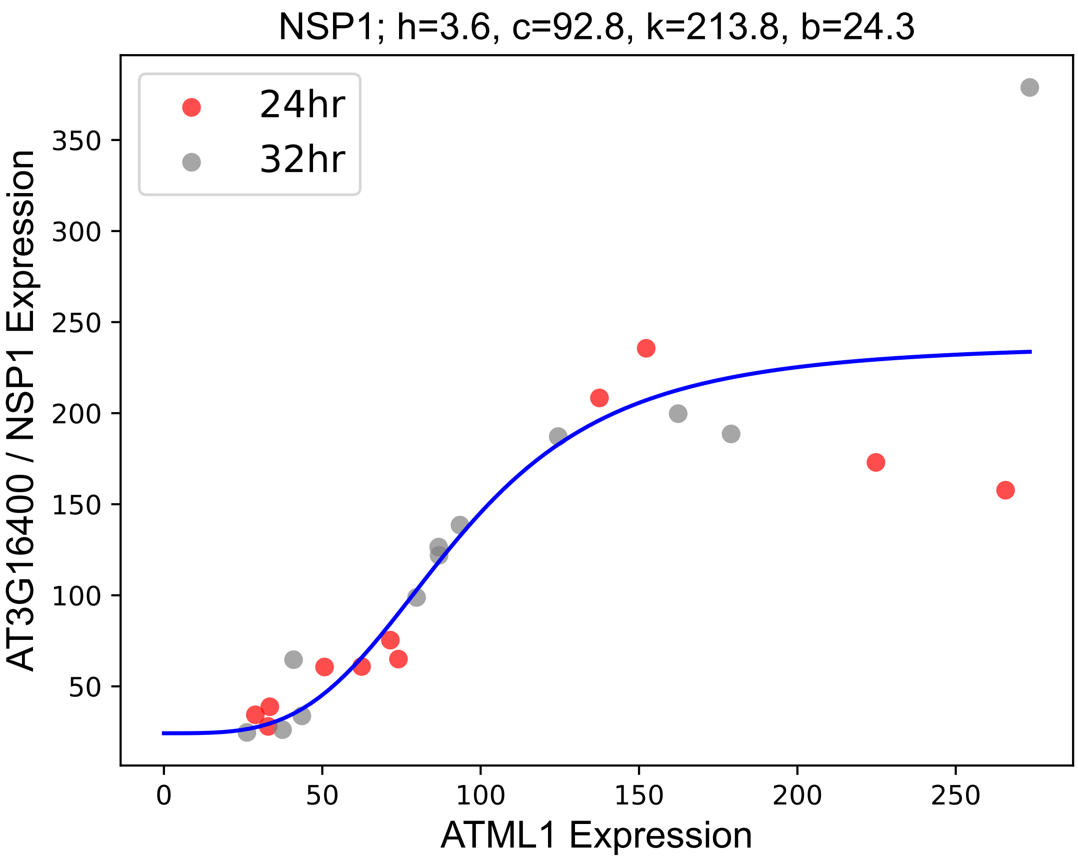

### PAE11.pdf

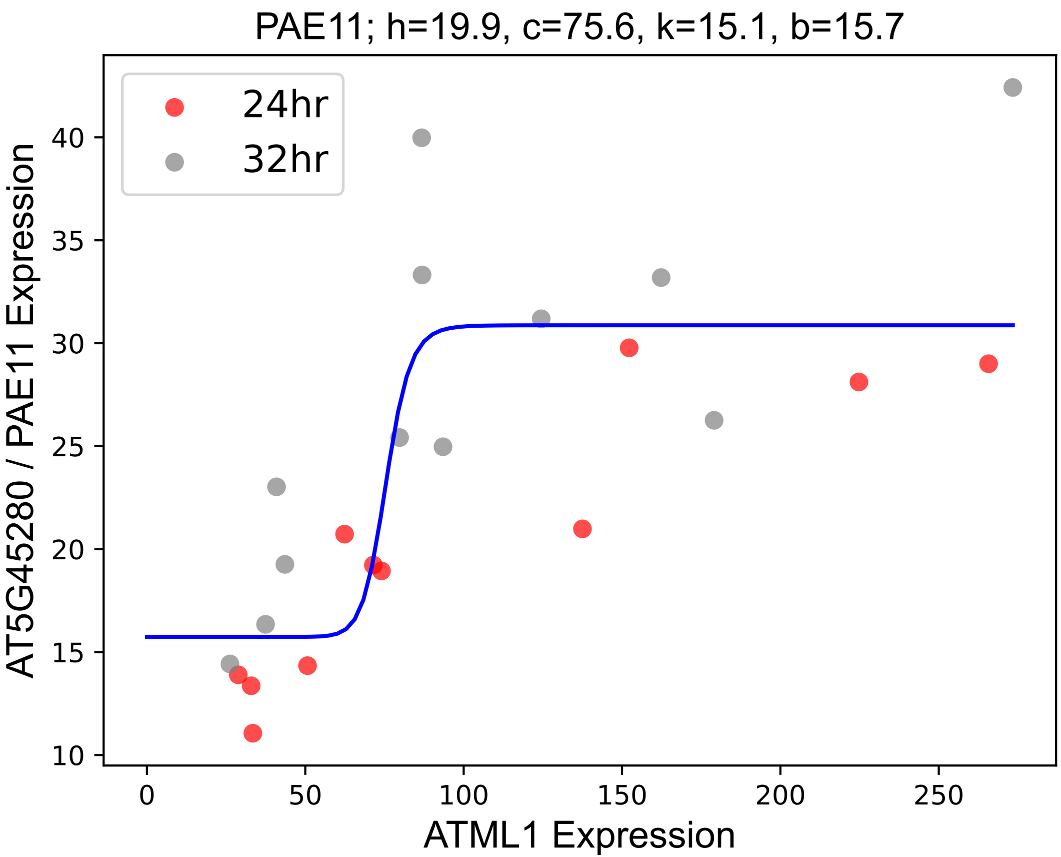

### PAP26.pdf

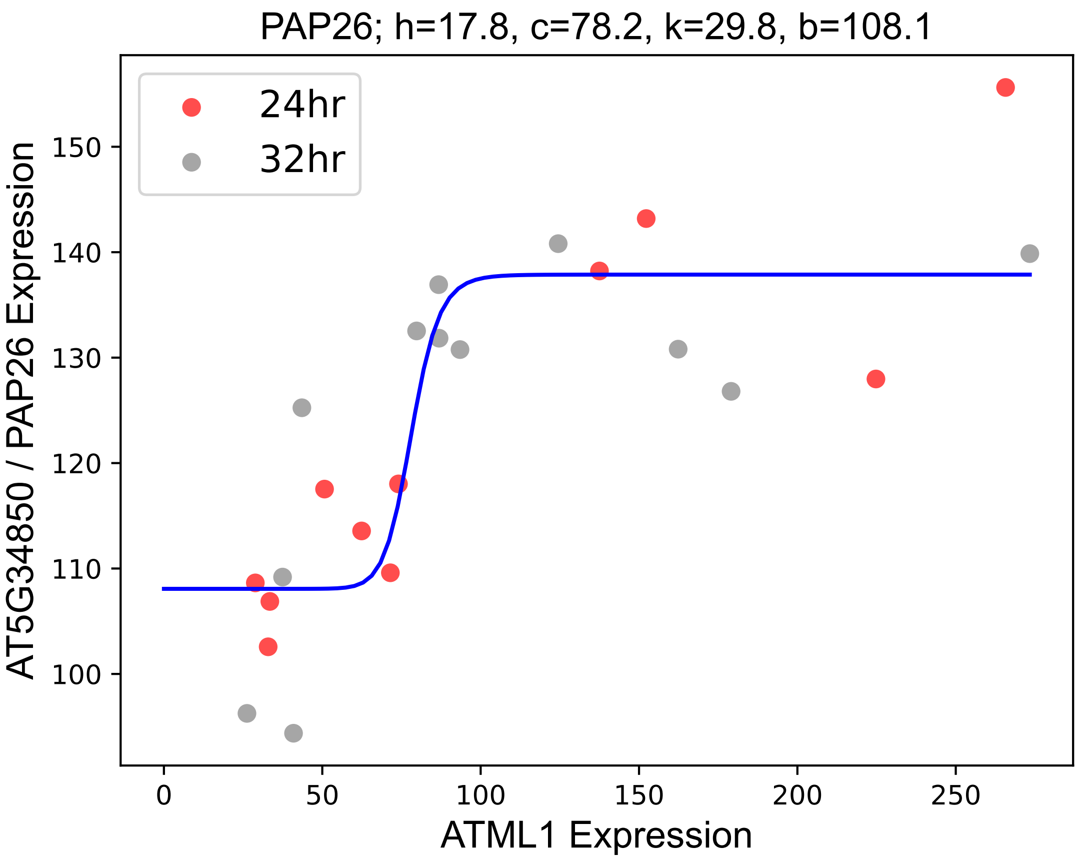
